# Diversity-oriented synthesis encoded by deoxyoligonucleotides

**DOI:** 10.1101/2022.10.16.512431

**Authors:** Liam Hudson, Jeremy W. Mason, Matthias V. Westphal, Matthieu J. R. Richter, Jonathan R. Thielman, Bruce K. Hua, Christopher J. Gerry, Guoqin Xia, Heather L. Osswald, John M. Knapp, Zher Yin Tan, Praveen Kokkonda, Ben I. C. Tresco, Shuang Liu, Andrew G. Reidenbach, Katherine S. Lim, Jennifer Poirier, John Capece, Simone Bonazzi, Christian M. Gampe, Nichola J. Smith, James E. Bradner, Connor W. Coley, Paul A. Clemons, Bruno Melillo, C. Suk-Yee Hon, Johannes Ottl, Christoph E. Dumelin, Jonas V. Schaefer, Ann Marie E. Faust, Frédéric Berst, Stuart L. Schreiber, Frédéric J. Zécri, Karin Briner

## Abstract

Diversity-oriented synthesis (DOS)is a powerful strategy to prepare molecules with underrepresented features in commercial screening collections, resulting in the elucidation of novel biological mechanisms. In parallel to the development of DOS, DNA-encoded libraries (DELs) have emerged as an effective, efficient screening strategy to identify protein binders. Despite recent advancements in this field, most DEL syntheses are limited by the presence of sensitive DNA-based constructs. Here, we describe the design, synthesis, and validation experiments performed for a 3.7 million-member DEL, generated using diverse skeleton architectures with varying exit vectors, derived from DOS, to achieve structural diversity beyond what is possible by varying appendages alone. We will make this DEL available to the academic scientific community to increase access to novel structural features and accelerate early-phase drug discovery.

## Introduction

Screening collections comprising novel chemical structures have produced valuable probes of targets of interest, including compounds that exert their biological effects through novel mechanisms of action (nMoA)^1^. One approach to generate collections with broad molecular diversity is diversity-oriented synthesis (DOS)^2–6^, which has produced numerous nMoA compounds through phenotypic screening campaigns^7–19^. Similar success with target-based screening approaches can be scarce in cases where detailed knowledge of the underlying biology is not available, where there is no knowledge of ligandable pockets or known ligands, or where desirable efficacy can only be achieved through polypharmacological modulation of the target system^20^. Despite the long-held goal of DOS — to increase the structural and especially the performance diversity of screening collections — few such collections are readily available to the scientific community from commercial vendors. Compounding the challenges to the widespread use of such collections is that these compounds generally have increased costs, with reduced or sporadic supply of key synthetic intermediates for the rapid confirmation and follow-up of identified hit compounds^21^. Consequently, access to such collections is restricted to researchers with substantial funding and/or synthetic chemistry resources, creating a barrier to new discovery in hit-generation sciences.

DNA-encoded library (DEL) technology is established as a screening tool in industry and academia to identify binders of soluble proteins via affinity-based screens. The general concept of DEL technology — to prepare large libraries of compounds through combinatorial split-and-pool chemistry with concurrent encoding of each individual molecule by a unique DNA barcode — facilitates the simultaneous parallel screening of entire libraries of compounds for binding to a target of interest^22^. The amount of material needed for these screens is greatly reduced compared to traditional high-throughput screening (HTS), as it can leverage the sensitivity and reliability of polymerase chain reaction (PCR) and next-generation sequencing (NGS) for data generation^23^. Various approaches that expand the applicability of DELs for synthetic chemistry compatibility have been developed successfully, thus expanding the chemical space coverage of libraries (typically in an aqueous environment, restricted by the sensitivities of DNA to reagents, heat, and pH)^24–28^. In addition, progress has been made in DEL screening, including alternate coding schemes and novel procedures for performing screens, that broaden the range of tractable targets and enhance the value of screening information^29–32^.

Several approaches are used for DEL generation. The “single pharmacophore library”^33^ is the most common strategy for synthesizing DELs. Here, successive steps of appendage diversification of a common skeleton (pharmacophore) are encoded by enzymatic ligation that record the synthetic history for each individual compound (**Fig. 1a**). The skeleton may be generated in situ or off-DNA, depending on the types of building blocks used in the synthesis. Another strategy (**Fig. 1b**) does not use a fixed central skeleton, and instead connects diverse building blocks in a linear fashion. An expanded strategy (**Fig. 1c**), employed in this study, makes use of multiple skeletal elements that are consistent in reactivity to allow for simultaneous appendage diversification using a common set of diverse appendages. Earlier work by Gerry et al.^33^ highlighted a library generated with this strategy; in this work, the library generation strategy is expanded with additional multifunctional skeletons, additional building blocks, and diverse chemistry types. An attractive feature of this strategy is that the common appendages vary greatly with respect to their relative exit vectors, allowing for increased diversification.

**Fig. 1.**
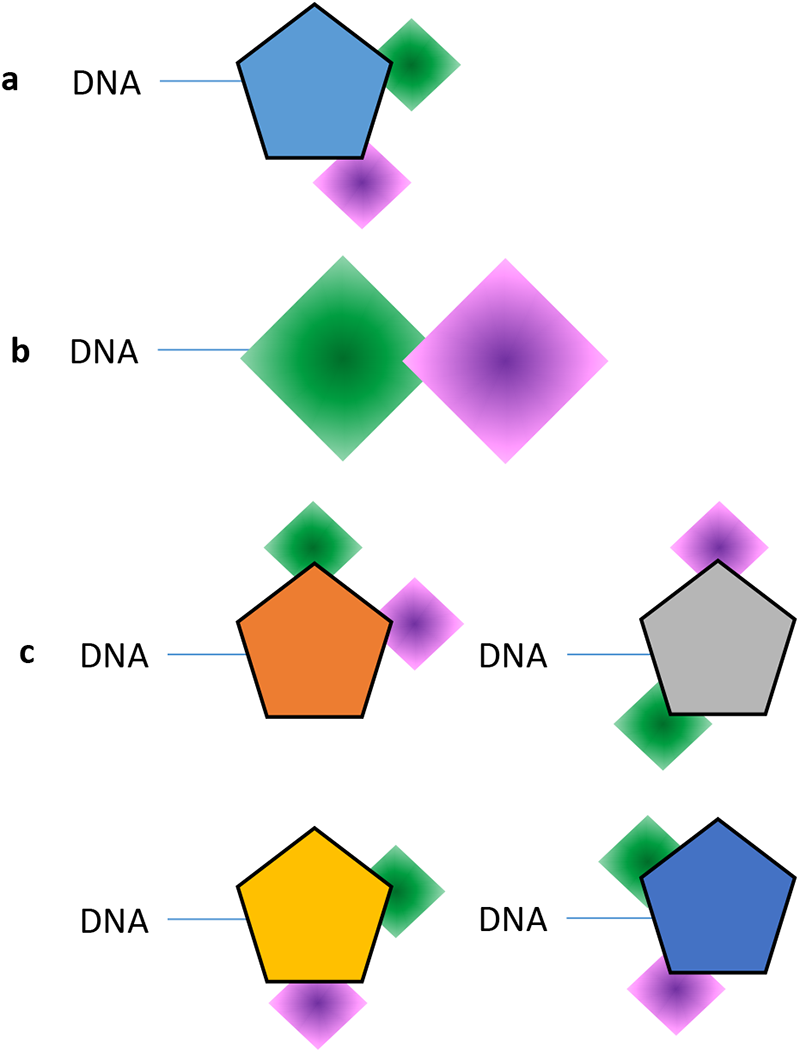
Conceptual comparison of DNA-encoded library synthesis strategies. Shapes with solid coloring represent multifunctional skeletons with defined exit vectors, and shapes with graduated coloring represent collections of building blocks with diverse pharmacophoric features. **a**, A single fixed skeleton library wherein appendage diversity results from fixed exit vectors of the central skeleton. **B**, A skeleton-free library constructed by direct linkage of diverse building blocks to each other. **C**, A library comprising multiple skeletons bearing well-defined but variable exit vectors to collections of building blocks with diverse chemical features.

We aim to facilitate DEL screening with compounds with diverse skeleton architectures that are readily derived through DOS chemistry. We call this approach diversity-oriented synthesis encoded by deoxyoligonucleotides (DOSEDO). In this work, we describe the design, synthesis, and validation experiments performed using this enhanced DEL. The DOSEDO library is used in screening campaigns to generate diverse “hits” from different protein targets, demonstrating a range of screening success. This approach is an expansion of prior work from this group, wherein a complete set of stereoisomers was encoded from a single scaffold series^33^, and this approach is complimentary to approaches such as the ‘aldehyde explosion’ approach, where alternate reaction conditions are used to modify a single on-DNA moiety to many different functional groups^34^. This library and resources for resynthesis are available to the academic scientific community through a facilitated access approach, discussed later in this report^35^.

## Results

### Selection of components

While DEL technology allows for the rapid generation of extremely large compound libraries, we prepared a moderate-sized library compared to those described in some reports^36^. The reasons for this library size were three-fold: 1) to ensure that reaction validation prior to library production could be performed with good representation of the final library and manually controlled by a single operator with high reproducibility; 2) to ensure that moderate-to-high coverage of library sequences was achievable in widely available and relatively low-cost NGS experiments; and 3) to ensure a manageable cost for acquiring the raw materials to enable library production and follow-up studies. We selected 61 multifunctional compounds to serve as skeletons^33, 37–41^, all comprising secondary amines (Fmoc-, Boc- or Ns-protected) and an aryl halide (Br or I), which we planned to use for on-DNA diversification (**Fig. 2**). Each skeleton bore either a carboxylic acid or primary hydroxyl group, which were used as the site of DNA attachment and in some cases allowed for an alternative DNA attachment point. Other DELs generated with similarly diverse skeletons have led to successful screening campaigns^43^. We included as many discrete stereo- and position isomers of these skeletons as was practical.

A branching rather than linear sequence library design was used so that the rigid central skeletons, containing well-defined exit vectors, allowed for a broad distribution of appendages in three-dimensional space^44^. We also focused on amine and aryl halide functional groups to enable the use of thoroughly validated DNA-compatible diversification steps (acylation, sulfonylation, reductive amination, and Suzuki couplings)^24^. Building blocks to be appended to the core skeletons were selected through a semi-automated process created in Knime Analytics Platform^45^ to pick a diverse set from commercially available compounds within a defined calculated property space, followed by a manual review (**Supplementary Fig. 21**). For our DNA coding scheme, we used a modified version of the double-stranded DNA scheme described previously^46^. The overall goals of these choices were to facilitate library construction and resynthesis and to maximize the structural space encompassed by the library.

**Fig. 2.**
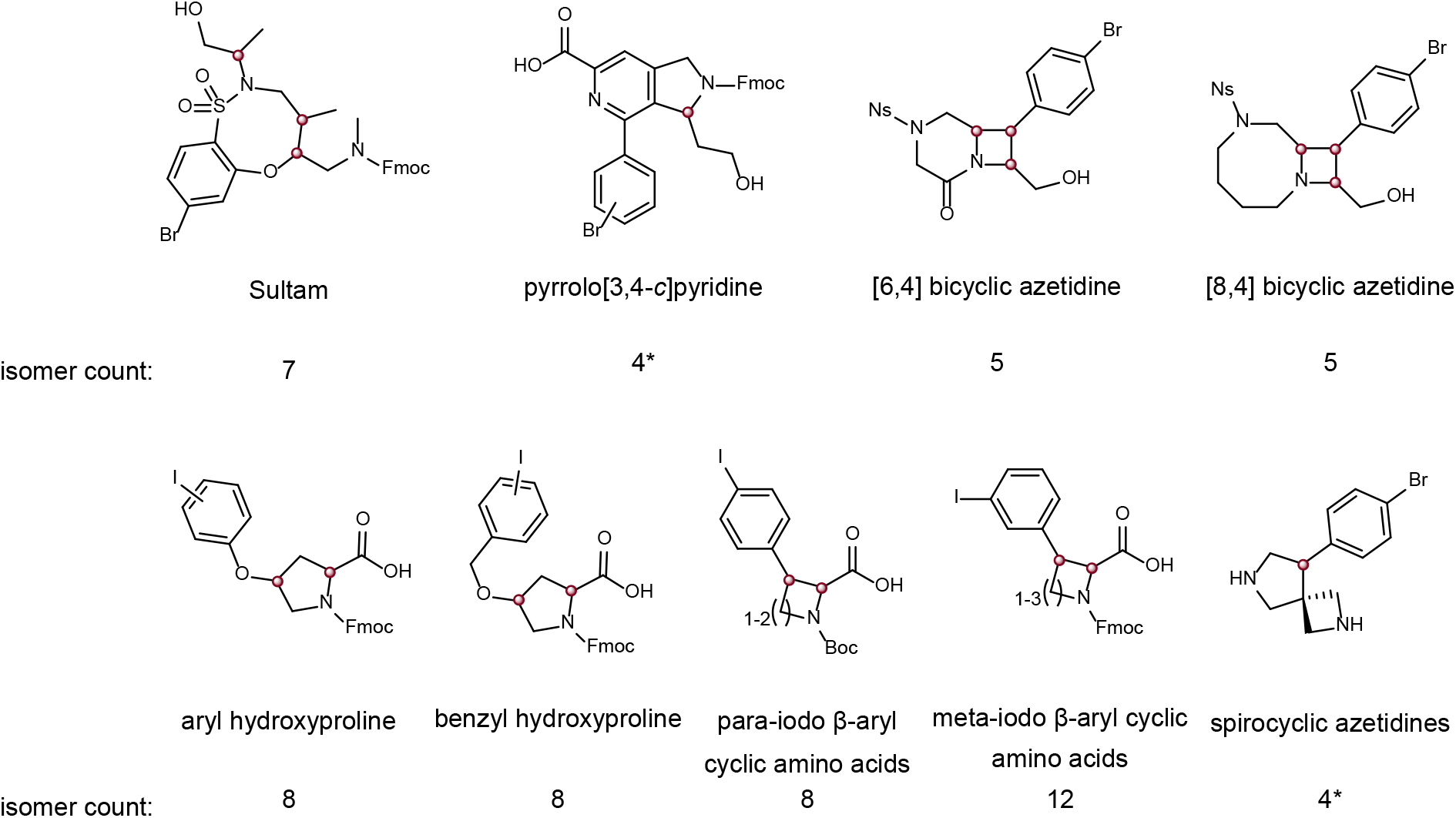
Overview of skeletons selected for inclusion in the DOSEDO library. Skeletons shown are indicative of the compounds loaded onto DNA. Isomer counts report the number of isomers (stereogenic and positional) for the indicated skeletons. Red spheres indicate two distinct isomers at each stereogenic center – in no cases were stereochemical mixtures or skeletons of unknown absolute configuration used in the library construction. Asterisks indicate skeleton series having alternate DNA attachment points.

### Development of DNA linkage chemistry

Skeletons attached to DNA through their carboxylic acids were coupled under conditions modified from earlier reports^46^ and optimized to minimize material usage (**Supplementary Tables 6–8**). Skeletons attached to DNA through their primary hydroxyl groups were coupled using an optimized carbamate-linkage protocol (**Supplementary Section 2.3.1.2**). The hydroxyl group was activated with *N,N*′-disuccinimidyl carbonate (DSC), filtered over silica to remove *N*-hydroxysuccinimide (NHS) and unreacted DSC, and incubated with amine-functionalized DNA. We also developed a DNA-compatible solution-phase Ns deprotection (**Supplementary Section 2.2.2),** similar to a recently reported method^47^, rather than using solid-supported processes^48, 49^. After linking each skeleton to DNA, conjugates were purified by preparative HPLC. The resulting HPLC eluents were concentrated and desalted by EtOH precipitation and ultrafiltration. Chromatographic data for the purified constructs following reaction validation and library production are provided in **Supplementary Table 11**.

### Reaction optimization

With purified skeleton–DNA conjugates in hand, we optimized the diversification steps of the DEL synthesis. We focused initially on reducing the residual acetate derived from preparative HPLC mobile phase, which generally presented a chemoselectivity challenge in acylation reactions. Acylation conditions were optimized using DNA-conjugated proline as a representative amine. Following preparative HPLC, multiple washes of EtOH precipitates, ultrafiltration, or desalting on G-25 Sephadex led to excellent minimization of acetylation by-products in acylation reactions. Minor adaptations of published acylation^46^, reductive amination and sulfonylation conditions were effective across many substrate–reactant combinations^33^.

We used Suzuki coupling reactions to diversify aryl bromides and iodides in the library. Using general coupling conditions that could be applied to all skeletons, we performed several optimization reactions (**Supplementary Fig. 2**). Four palladium catalysts were screened in combination with two boronic acid pinacol esters that were dissolved in either EtOH, MeCN, or DMA. A total of 288 reactions were performed, leading to the selection of PdCl2(dppf)·CH2Cl2 as the preferred catalyst. This catalyst had the highest conversion to desired product at all temperatures and in all co-solvents compared to matched reaction sets. In addition, the PdCl2(dppf)·CH2Cl2 catalyst resulted in the least variable outcomes across a range of temperature and time conditions. We also found that MeCN was the preferred solvent in combination with any of the catalysts tested, though in practice, EtOH was complementary to MeCN for building block dissolution. Therefore, a 1:1 mixture of EtOH and MeCN was used in later reactions for increased homogeneity.

### Building block validations

Am mine-capping building blocks were profiled using HPLC-purified and desalted **Pro-DNA** as a model system, following the general scheme shown in **Fig. 3a**. At the end of each reaction, one relative volume of 5% aqueous piperidine was added to quench residual electrophile, and an EtOH precipitation was performed prior to LCMS analysis. A tabulated dataset can be found in **Supplementary Table 12**, and **Fig. 3a** reports the output for each reaction class. Only building blocks with >70% area-under-the curve (AUC) conversion to the desired product and with <10% AUC unknown species were considered for inclusion in the DOSEDO library. Prior to initiating a full Suzuki-coupling validation campaign, we applied the developed conditions to a subset of DNA-conjugated skeletons with uncapped or protected amines (**Supplementary Fig. 3**) using three building blocks that generally had high conversion to desired products. The DNA-conjugated skeletons (as free amines) gave variable outcomes based on the skeleton class or stereoisomer. We also noted generally poor conversion for these couplings, mirroring the conclusions drawn for related published skeletons^33^.

We then assessed the effect of amine-capping mode on Suzuki coupling for different skeleton classes and to assess the reactivity of aryl iodides versus bromides. We performed acylations with acetic acid and benzoic acid, sulfonylations with tosyl chloride, and reductive alkylations with benzaldehyde on six DNA-conjugated aryl iodides and six aryl bromides (**Fig. 3b**). The resulting 48 products were subjected to Suzuki coupling conditions (using a 0.1 mM solution of DNA-conjugate in pH 10 bicarbonate buffer), with phenylboronic acid or its pinacol ester. During early optimization experiments, we noticed some apparent degradation of [6,4] bicyclic azetidine skeletons, possibly resulting from basic hydrolysis. Therefore, we repeated the same set of reactions with a pH 8 phosphate buffer. Analysis of the resulting 192 LCMS traces led to several conclusions. First, the use of boronic acid or pinacol ester had little differential effect on average %AUC product formation, which were 78 and 76, respectively, with average %AUC unknown DNA species of 11 in both cases. Likewise, the different amine capping modes had no substantial impact on the conversion to desired products (with average %AUC {standard deviation} for benzoic acid = 80{10}; acetic acid = 77{11}; *o*-toluenesulfonyl chloride = 77{20}; benzaldehyde = 73{11}). A key differentiating factor was the combination of buffer pH and whether the skeleton bore a bromide or iodide. **Fig. 3b** shows the change in %AUC product and unknown DNA species that occurred in response to the buffer change. In general, the iodide-containing group of reactants (comprising *meta*-iodo phenyl cyclic amino acids based on azetidine, pyrrolidine and piperidine, as either the *cis* or *trans* diastereomers) had preferred reaction profiles in pH 10 bicarbonate buffer, with considerably greater %AUC product and less %AUC unknown species present versus the otherwise identical pH 8 phosphate buffered conditions. The reverse was true for the aryl bromide group. We applied the developed conditions with our full set of boronic acids and esters using exemplar ArI and ArBr model systems (**1** and **2** respectively, **Fig. 3c**), and found that while reaction profiles were generally effective with respect to unknown byproducts, changing the buffer alone did not give a comparable validation rate between iodide and bromide model systems. A higher catalyst and boronic acid/ester loading, as well as a longer reaction time, led to an acceptable number of validated building blocks, with good concordance between the iodide and bromide validation campaigns.

**Fig. 3.**
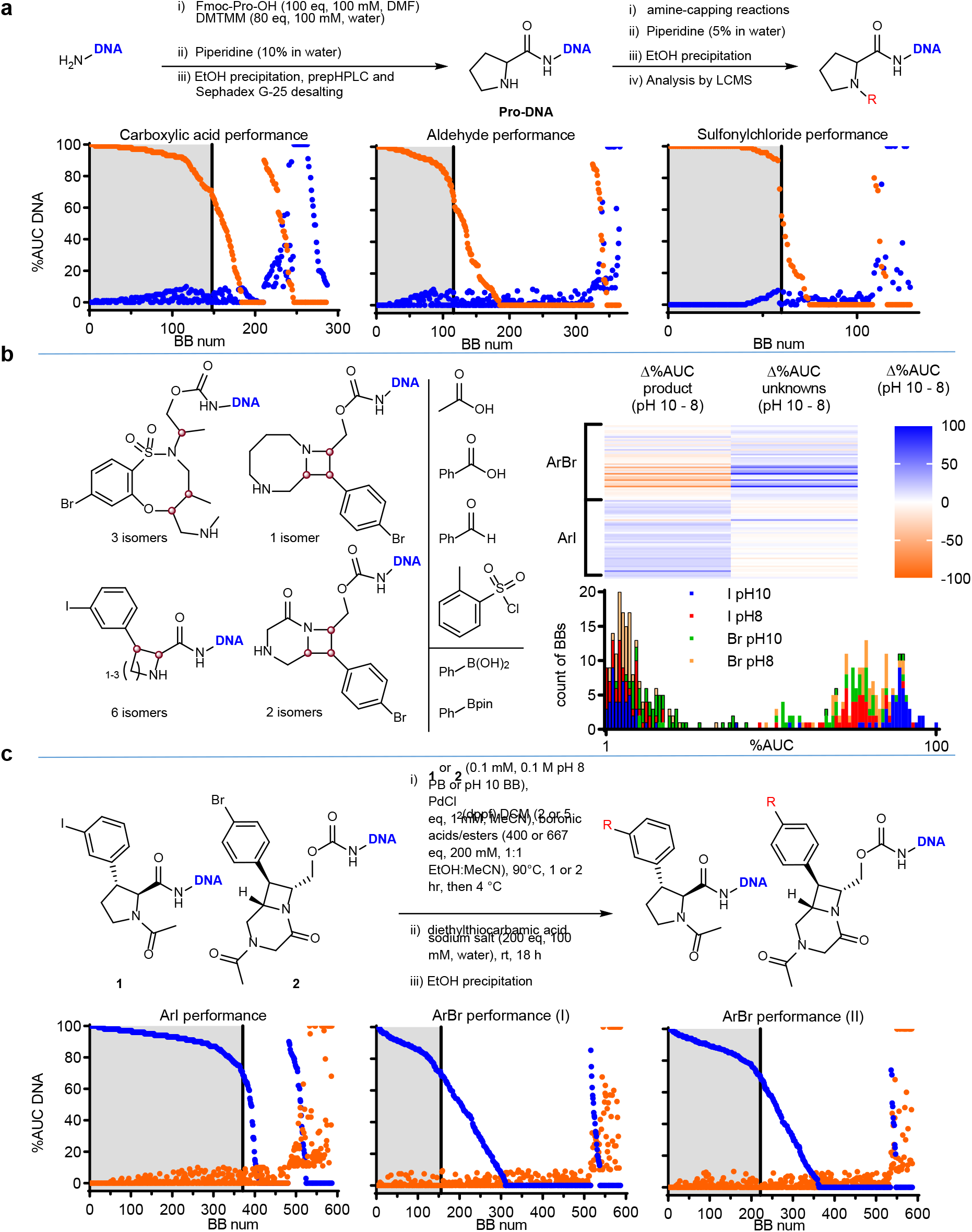
Summary of building block validations. **a**, Summary of amine-capping validation experiments, including a general synthetic scheme for the preparation and use of a proline-based model construct (Pro-DNA), and charts showing the performance of acylation, reductive alkylation, and sulfonylation. Blue points show %AUC (area under the curve) corresponding to desired product species; orange points show %AUC of unknown DNA species as assessed by MaxEnt1 deconvoluted mass spectra. “BB (building block) num” is a ranking of compounds in two phases: i) decreasing %AUC product with increasing %AUC unknowns up to 10, then ii) decreasing %AUC product with >10 %AUC unknowns. The highlighted grey region of each chart denotes building blocks that met our inclusion criteria of >70%AUC product with <10%AUC unknowns. **B**, Effect of buffer on performance of Suzuki couplings comparing a subset of skeletons [with variably capped amines] bearing ArI (aryliodide) versus ArBr (arylbromide). The skeleton series, amine capping, and model Suzuki coupling building blocks are shown, alongside a heat map representation of the difference in %AUC product and unknown species present when analogous reactions were performed at pH 10 vs. pH 8. Also displayed is a stacked bar chart comparing the performance of ArBr and ArI skeletons, on aggregate, with respect to product formation (bars without borders) and unknown species (bars with black borders) in different buffers. **C**, Summary of Suzuki coupling building-block validations including the constructs and conditions used and a comparative analysis of performance for a model ArI and ArBr (with the chart labelled “Arylbromide (II)” making use of higher catalyst and boronic acid/ester loading). Additional reaction time is also noted.

Following validation of our complete collection of amine-capping reagents and boronic acids/esters, we tested the extent of available diversity that could be incorporated into the final library design. In a basic sense, the expected reactivity was observed; for example: aryl and vinyl sulfonyl chlorides were considerably more productive than aliphatic sulfonyl chlorides (presumably due to rapid hydrolysis of the latter in aqueous mixtures), and sterically more hindered aldehydes and carboxylic acids tended to be less reactive. For each class of building block, we prepared a distance matrix generated from Morgan 2 fingerprints and applied multidimensional scaling (MDS, by Sammon mapping, **Fig. 4**). Except for alkyl sulfonyl chlorides, we generally observed that positive reaction outcomes in amine capping reactions resulted in good coverage of available chemical space across all reaction modes. However, for Suzuki couplings (**Fig. 4a–c**), the building blocks forming the core of the MDS map tend to perform better than those at the extremities, which are populated with compounds comprising the most dissimilar building blocks in the collection. We conclude that while our conditions allow for reasonable inclusion of building blocks, they are biased against less represented and more diverse chemotypes. We found that electron-rich and/or -hindered 5-membered heterocycles, as well as vinyl boronates, often constituted the bulk of poorly performing reactants. Moreover, we noticed that seeking consensus positive outcomes in the ArI and ArBr Suzuki coupling validation campaigns led to an erosion of fingerprint diversity in the highest priority set of building blocks (**Fig. 4a**), but we believe this sacrifice of appendage diversity for consistent synthetic outcomes to be a worthwhile compromise.

**Fig. 4.**
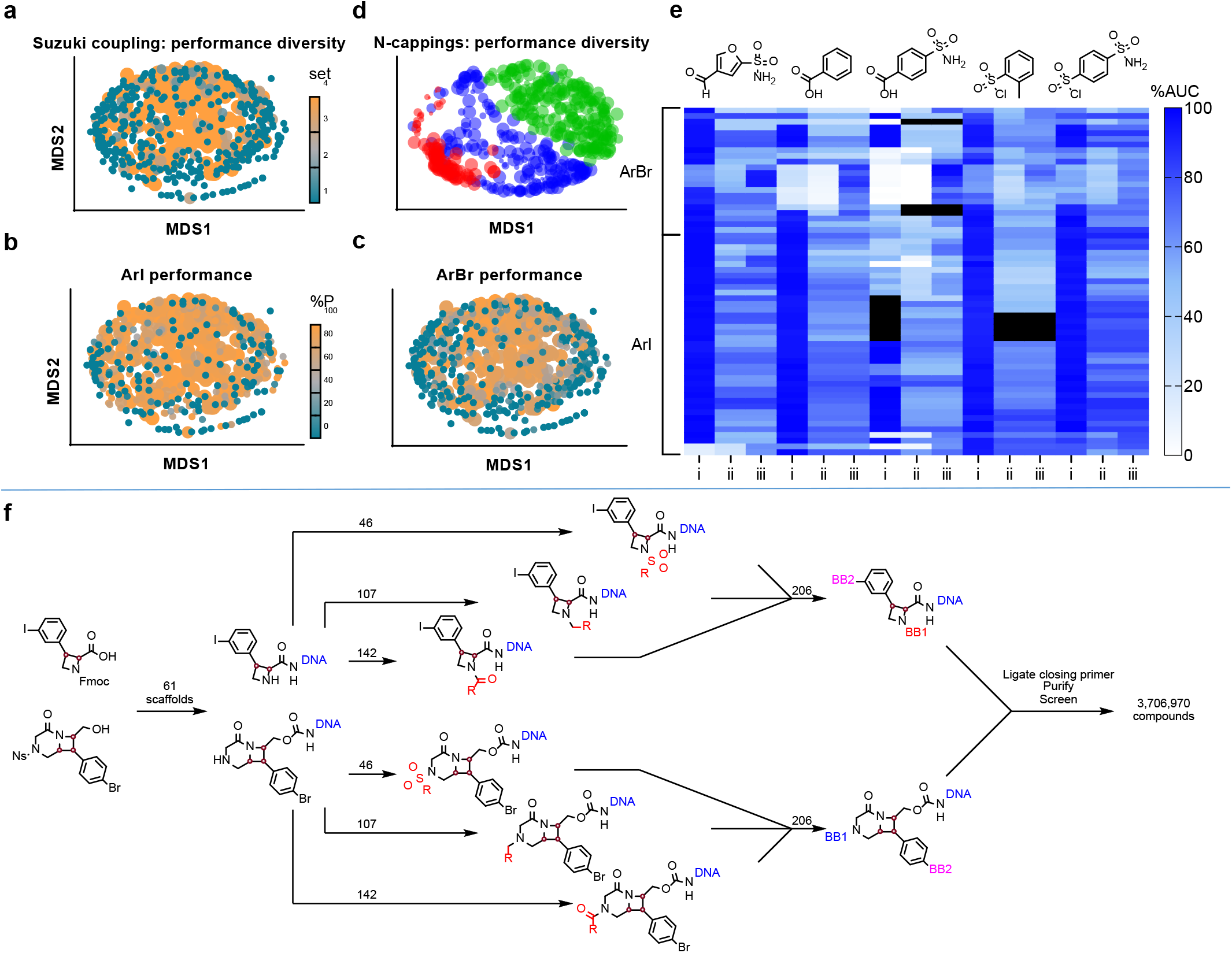
Assessment of reactivity in building block and skeleton validation experiments. **A**, Combined performance of ArI (aryliodide) and ArBr (arylbromide) Suzuki coupling reactions, where each common boronic acid/ester moiety was replaced with a methyl group in silico, then visualized by Sammon mapping of a Morgan2 fingerprint distance matrix to two dimensions, coloring and sizing by preference for inclusion in DNA-encoded libraries (DELs) (4 = highest), lowest preference building blocks (BBs) are layered above higher preference BBs for visualization. **b**, Performance of the same BBs represented in **a** specific to the ArI model system, with point size still indicating consensus preference category but colored by adjusted %AUC (area under the curve) product (%P). **c**, As for **b** showing data specific to the ArBr model system. **d**, Comparison of amine-capping performance by Sammon mapping of individual building blocks (with no in silico treatment of common reactive moieties), sized by %AUC product, blue – aldehydes, green – carboxylic acids, red – sulfonylchlorides. **e**, Heat-map representation of skeleton validation, showing data for five different amine capping reactions followed by a single Suzuki coupling with (4-sulfamoylphenyl)boronic acid. Each row represents reactions of a specific skeleton. Column labelling indicates: i – adjusted %AUC for the desired amine-capped product; ii – adjusted %AUC of the desired Suzuki coupled product; iii – adjusted combined %AUC for the desired Suzuki coupled product as well as the Suzuki product of non-*N*-capped starting material. Black cells indicate either that the chromatographic data obtained was insufficient to resolve starting material and product peaks, or that no DNA species were present based on chromatography (potentially due to loss of the DNA pellet during ethanol precipitation or instrumentation error). **f**, Synthetic plan for DOSEDO library production using a single ArI and ArBr skeleton to represent the wider collection, with numbers indicating how many building blocks were used for each step.

Based on our analysis, we reasoned that the Suzuki coupling used in the library synthesis should be performed following amine-capping reactions, and under different conditions for the bromide and iodide-containing pools of intermediates. With those defined restrictions, we designed a synthetic plan (**Fig. 4f**) to maximize the performance and consistency of individual reactions. The synthesis began with skeleton attachment to the DNA headpiece through either acylation or carbamylation. After amine deprotection, purification, and encoding individual constructs, two separate sub-library pools were generated for ArI- and ArBr-containing skeletons. These separate pools underwent amine-capping and encoding reactions, followed by Suzuki coupling and encoding reactions. The two sub-libraries were pooled, closing sequences were ligated, and the sub-libraries were purified as a single encoded collection.

### Skeleton validation

Prior to DOSEDO library synthesis, we validated the conditions for amine capping and Suzuki coupling to ensure that they were compatible with all skeletons planned for inclusion in the library. We aimed to identify any critical reactivity or stability issues, such as determining whether reactions are dependent on stereochemistry, that could impact the quality of the library or affect the interpretation of screening data or prioritization of putative hits for resynthesis. Detailed protocols for the skeleton validation campaign are described in **Supplementary Section 2.2.4**. The concentration of each purified skeleton-DNA conjugate was normalized in nuclease-free water. Conjugates were subjected to five amine-capping reactions (at least one of each reaction class). A sample was taken for LCMS analysis, and the residual crude amine capping reaction mixture was subjected to a single Suzuki coupling (under skeleton-appropriate conditions) followed by a final LCMS and analysis (**Fig. 4e**). All skeleton–DNA constructs maintained their integrity throughout these reaction sequences, and performance was acceptable for all skeletons to be included in the DOSEDO library using optimized reaction conditions. We did note variability in the performance of specific reactions. For example, β-arylated cyclic amino acids behaved consistently – with high-yielding amine cappings and concurrent high-yielding Suzuki couplings – except in the case of *trans*-configured piperidines, which performed very poorly in reductive alkylations. In addition, while sultam skeletons generally displayed slightly reduced reactivity towards amine-capping than other skeletons, they had notably reduced reactivity across all seven stereoisomers in the panel towards acylation compared to sulfonylation and reductive amination. Based on these findings, we included all 61 skeletons, 295 amine-capping building blocks (142 carboxylic acids, 107 aldehydes, and 46 sulfonyl chlorides), and 206 boronic acids/esters in the library synthesis. We also included two null-reaction conditions in the Suzuki coupling step (‘subjected to reaction conditions without boronic acid/ester present’, and ‘not subjected to reaction conditions’), resulting in a theoretical combinatoric matrix of 3,706,970 encoded compounds.

### Library synthesis

As a key goal of this work was to make this library available for screening by the research community, we sought to ensure adequate scale of production to support that effort without introducing non-validated elements into our protocols. Therefore, the synthesis scale was limited by operational considerations rather than by available materials. A 15% excess of each skeleton was used at the outset of synthesis (100 nmol) relative to the limiting forward primer-binding site duplex. This decision ensured that variability in the quantification of skeleton–headpiece conjugates was normalized during the first stage of synthesis. Skeleton identity was concurrently encoded by introducing a cycle-1 tag in two–fold excess of the forward primer binding site duplex, to ensure that all tandem ligations progressed to complete conversion (**Supplementary Fig. 6**). After pooling iodides and bromides separately, we performed cycle-2 tag ligations with a slight molar excess (10%) relative to the cycle 1 tags. Ligation efficiencies were assessed by analytical electrophoresis using pools of 12 samples per lane, and we detected no evidence of individual ligation failure by native or denaturing methods (**Supplementary Figs. 7 and 8**). Amine-capping reactions were performed, and after quenching residual electrophiles with piperidine, the products were pooled and concentrated with respect to DNA species and cleansed of small-molecule components by ultrafiltration and EtOH precipitation. For continued synthesis, we assumed that recovery was quantitative. The pools of encoded intermediates were first split, and then cycle-3 tag ligations were performed with a slight molar excess (approx. 10%) relative to the cycle-2 tag. We monitored ligation efficiencies by analytical electrophoresis with 12 pooled samples per lane (**Supplementary Fig. 9**). Suzuki couplings were then performed according to the developed conditions (**Fig. 3c**), and products were pooled as separate Br and I-derived sub-libraries.

### Input analysis

Prior to a production-scale closing ligation, we engaged in a small test library closure and sequencing experiment – DOSEDO_v1. We assumed that the efficiencies of all processes leading up to this stage were perfect and pooled a small quantity of the sub-libraries at a presumed 1:1 molar ratio at the individual encoded compound level. Closing sequences were ligated, and purification was performed by continuous flow electrophoresis (**Supplementary Section 2.4**). The ‘yield’ for DOSEDO_v1 closure over all steps of synthesis ranged from 19% (fluorometric quantification) to 37% (spectrophotometric quantification) relative to the limiting forward primer binding site duplex. DOSEDO_v1 was prepared for input sequence analysis according to the methods outlined in **Supplementary Section 3,** with the resulting analysis shown in **Fig. 5a**. We determined that pooling of sub-libraries was not equivalent; I-derived compounds were present in a 1.31–fold excess, on average, according to the two negative binomial distributions of the sub-libraries. We adjusted the pooling ratio for a production-scale closure that we termed DOSEDO_v2 (**Supplementary Section 2.5**). **Fig. 5b** shows the effect of input correction based on the v1 analysis. We also investigated the distribution of cycle-1, -2, and -3 tag ligations, on aggregate across all compounds, for both sub-libraries (**Supplementary Figs. 13 and 14**), and found that with very few exceptions, distributions were tight, with high concordance of ligation efficiency between sub-libraries. While sequencing depth for the v1 library was not ideal (7% of expected sequences were not observed), we dedicated more reads to the v2 input analysis, and as a result could assess the unobserved sequences (0.03%; 1254 sequences) for potential trends relevant to library use (**Fig. 5c**). This analysis suggested that in many cases, a specific Cy2 tag ligation failed from the I-derived pool but not the Br-derived pool. In another example, a combination of two Cy2 tags with a specific Cy3 tag was commonly not observed, with the Cy1 identity having no apparent impact on the outcome. With these two exceptions, we assume that all ligations were similarly effective in both sub-libraries, and the v2 sub-library pooling leads to an equivalent representation of individual compounds in the final library.

**Fig. 5.**
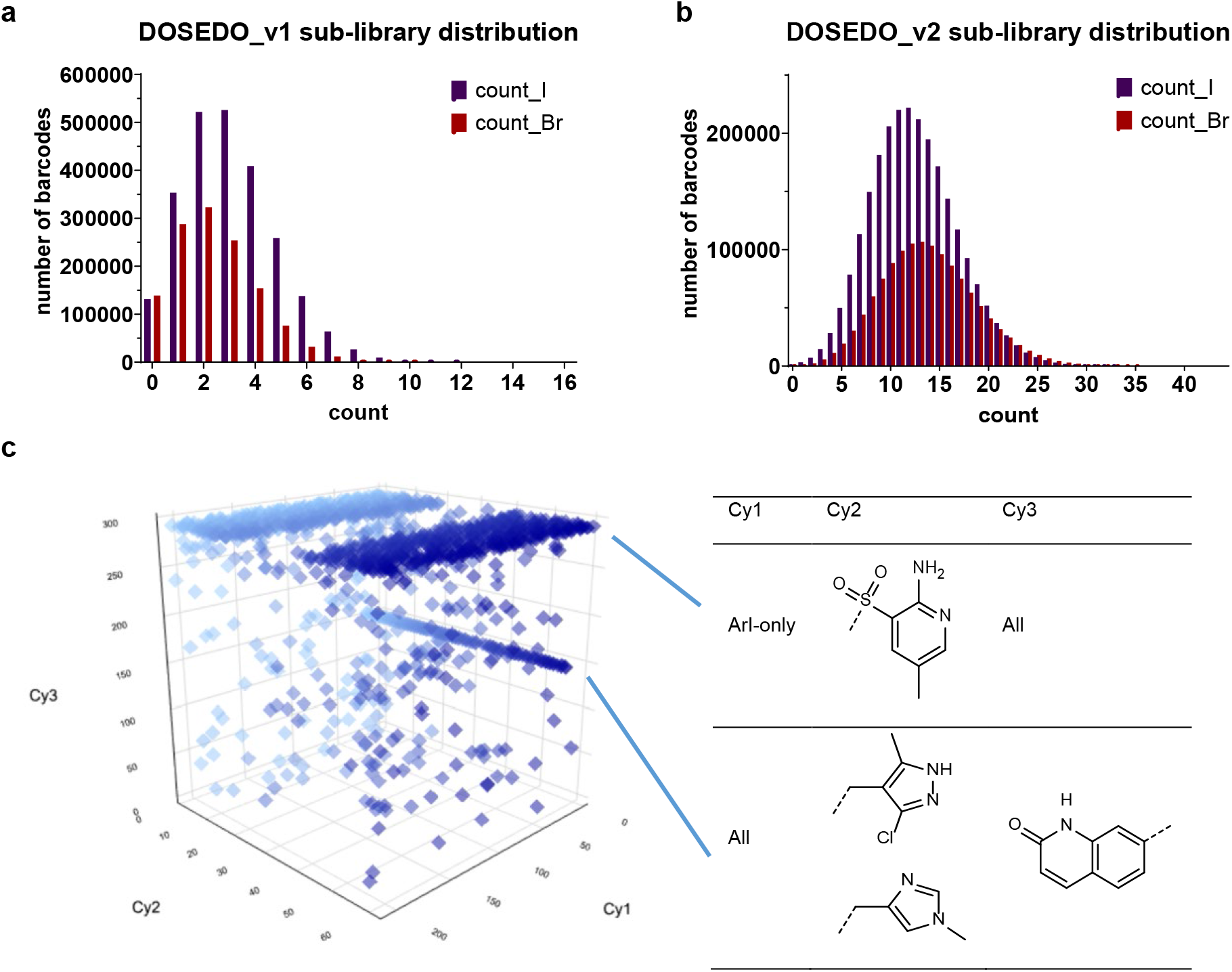
Assessment of input sequences. **a**, The number of barcodes observed at a given number of counts for the input DOSEDO_v1 library, showing the relative distribution of Br- and I-derived sub-libraries. **b**, As in **a** for DOSEDO_v2. **c.** unobserved sequences in DOSEDO_v2, with two sets of features highlighted in blue for being unobserved with unusual incidence as SAR (structure OR sequence activity relationship). Chart is graduated in blue in line with Cy2 identity to better perceive depth.

### PoC/validation screening experiments

We next wanted to ensure that the synthetic chemistry during DEL synthesis performed as expected and was encoded as intended. We selected three protein targets to screen and then synthesize putative hits off-DNA for validation purposes: carbonic anhydrase IX (CAIX), isocitrate dehydrogenase 1 (IDH1) R132H, and ubiquitin specific peptidase 7 (USP7). CA isoforms are common model systems for the DEL field because the primary sulfonamide structural motif is known to bind, often with high potency and tolerance of other functionality in the ligands^50^. Therefore, we included at least one example of a primary sulfonamide as a likely binder in each reaction set. There are also potential clinical applications for selective inhibitors of CAIX^51–53^. IDH1 mutants have been implicated in several medical conditions, including acute myeloid leukemia^54, 55^ and diffuse gliomas^56^. There are several compounds known to bind IDH1 R132H in its allosteric site^57^, some of which resemble compounds encoded within the DOSEDO library^58^. USP7 is a deubiquitinating enzyme (DUB) that can alter the degradation rate and cellular localization of specific protein substrates, some of which are of high interest in cancer progression^59^. We viewed mutant IDH1 and wild-type USP7 as more challenging proof-of-concept targets than CAIX, in that we had minimal prior knowledge of the chemical features that promote binding and did not deliberately bias the collection towards potent chemotypes, as was the case for CAIX. All screens were performed using an estimated 1 million copies of each encoded molecule per sample and using only a single cycle of panning the library over the target protein. Initial screening conditions were not optimized, to more accurately reflect the potential output of an initial experiment as it might be performed by an external research group. Thus, a diverse set of targets with potentially large differences in hit rates was chosen to test the DOSEDO library, under conditions that would be relevant to a wide range of research capabilities.

CAIX screening resulted in the expected enrichment of primary sulfonamide features above baseline noise (**Fig. 6**). With the added variable of the skeletons, intriguing SAR was noted – specifically a set of high-enrichment compounds and some low-enrichment compounds with similar structures. This finding indicated a possible difference in binding based on subtle appendage changes or skeleton stereochemistry but could also indicate a variable efficiency of on-DNA synthesis or bias induced by the screening protocol and NGS library preparation that was not fully captured by the prospective validation experiments. We probed these possibilities through off-DNA syntheses of the inferred compounds associated with high barcode enrichment.

**Fig. 6.**
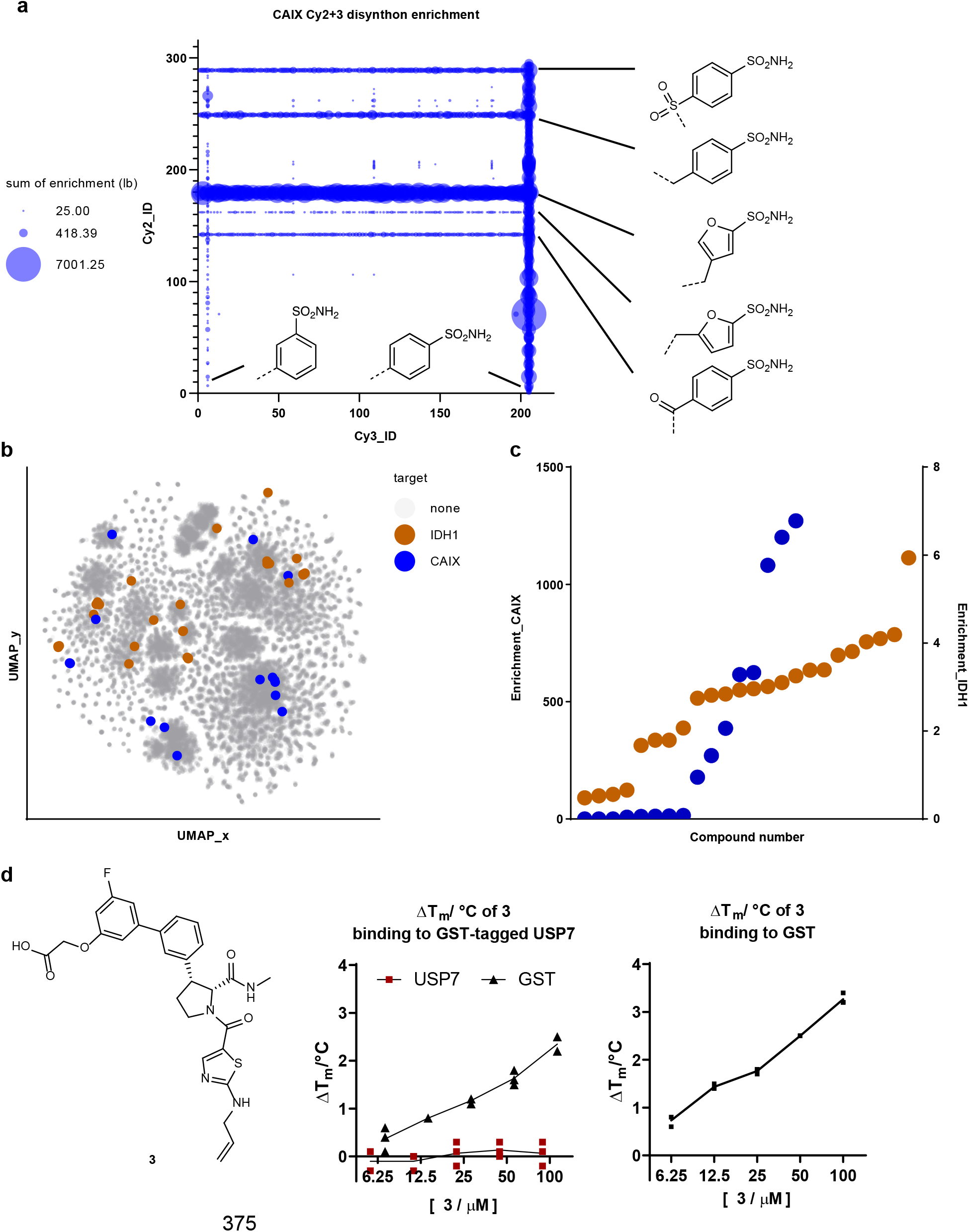
Analysis of validation screening exercises. **a**, Carbonic anhydrase IX (CAIX) screen output summarized by sum of lower bound (lb) calculated enrichments for all Cy2-Cy3 disynthons with all primary sulfonamide-containing building blocks included in the DOSEDO library shown (summed lower bound enrichments <25 was excluded from the plot for clarity). **b**, Chemical space map generated through UMAP multi-dimensional scaling of a Tanimoto distance matrix derived from Morgan fingerprints of enumerated encoded compounds. A random sample of 50,000 compounds from the enumerated DOSEDO library was used as background chemical space, along with 44 selected compounds for resynthesis. These compounds are colored by their binding target. **c**, Calculated enrichments of selected resynthesis target compounds from CAIX- and isocitrate dehydrogenase 1 (IDH1) R132H-validation screens. **d**, Structure of **3** alongside differential scanning fluorimetry (DSF) results with *N*-His-GST (glutathione S-transferase) tagged USP7 as well as His tagged GST (5 µM) in 50 mM Tris-HCl, pH 7.4. Two peaks belonging to GST (T_m_ 54.6 °C) and ubiquitin specific peptidase 7 (USP7) (T_m_ 44.9 °C) were observed in the melt curves of GST-tagged USP7. No thermal stabilization of USP7 by **3** was observed by tracking the USP7 peak; GST stabilization by **3** was observed by tracking the GST peak. **3** induces thermal stabilization of His-tagged GST in a dose-dependent manner, up to 3.3 °C at 100 µM. Data are mean ± SD, n=3 technical replicates.

An IDH1 R132H screen was performed using similar conditions; this screen yielded a far lower dynamic range of counts and calculated enrichment ratios than the CAIX screen. Nevertheless, we identified apparent SAR across multiple series encoded within the library. To validate the library synthesis, we selected compounds for resynthesis covering all reaction types, skeleton series, and levels of enrichment obtained from each screen. Wherever possible, we also captured close structural analogues with markedly different enrichment values to probe the variability of the generated sequencing data relative to off-DNA validation (**Fig. 6b and c**).

For off-DNA resynthesis, we assumed that encoded compounds were the true compounds of interest, and for the sake of simplicity, synthesized those specific compounds rather than replicating the DEL synthesis conditions to capture byproducts or truncates. Compounds were prepared following closely related published conditions^33, 38–40, 60^, often switching the order of steps relative to the DEL synthesis where it was deemed helpful. We modified all carboxylic acids that were used for DNA attachment as the methyl amides and left unmodified the corresponding hydroxyl groups that were used for carbamate DNA linkage.

Anticipated CA binders were principally assessed using a colorimetric enzyme-inhibition assay. While selected primary sulfonamides tended to validate as active, the anticipated non-binders often inhibited CA function quite potently. Given the high enrichment ratios typically observed for validated CA binders, these false negative results likely indicate poor synthetic performance for specific reaction combinations in the pool (**Supplementary Table 10**). Compounds prepared as putative IDH1 R132H binders were selected based on lower signal-to-noise sequencing data, and as such, we expected lower correlation of enrichment to off-DNA binding. This prediction held true, though 9 of the 25 compounds tested were validated as binders by SPR (**Supplementary Fig. 17**). Of the four skeleton series included in the resynthesized hits, we identified at least two binding series, with the sultam series being the more potent series. We then showed that only the tightest binding sultam (K_D_ = 0.89 µM, also the most highly enriched compound from the NGS data) effectively stabilized IDH1 R132H and displayed robust dose-dependent enzymatic inhibition (**Supplementary Figs. 19 and 20).** The specific stereochemical features of the identified hit aligned with the previously published IDH1 R132H inhibitor BRD2879^58^, bearing different substituents installed through an alternate reaction scheme.

BRD2879 was identified using a conventional high-throughput screen, providing support for the overall synthesis, encoding, screening, barcode amplification, and NGS steps of these experiments. For the USP7 screen, a single compound was nominated for off-DNA synthesis (**3**), primarily because the level of enrichment was considerably lower relative to the matrix only control screen. USP7 was screened as an N-terminally His-tagged glutathione S-transferase (GST) fusion protein, and we found that **3** selectively bound the GST portion of the construct. This finding was confirmed by DSF with isolated GST as compared to the GST-USP7 fusion (**Fig. 6d**), and further by SPR studies (**Supplementary Fig. 18**). The range of screening outcomes for this library was in line with the distinct characteristics of the protein targets screened.

## Discussion

In this work, we synthesized a DNA-encoded collection of compounds using a strategy based on skeleton and appendage diversification, extending and expanding on earlier work^33^. In particular, the expanded number and structural features of skeletons with varied exit vectors allowed us to achieve a branching library synthesis scheme to cover more three-dimensional space. The library size was sufficiently large to be inclusive of a large chemical space, but the relatively small size allowed us to perform robust analytics and have confidence in the barcode identification of hits. We screened the library against several distinct protein targets and validated that the library can be used to identify binders to targets of interest. We are confident that the on-DNA synthetic chemistry, as well as the concurrent encoding steps, was satisfactory, but our study highlights the impact of noise in DEL screening data and the critical importance of the protein targets used.

We believe that the encoded library will be of interest to the hit-discovery community because underrepresented chemical features are present in the library, including novel and rigid skeleton architectures and well-defined stereogenic elements. The use of defined novel skeleton structures whose synthesis can be scaled up in preparation for hit validation, along with optimized downstream reactions with commercially available building blocks, will facilitate the process of hit validation.

We aim to facilitate DOSEDO library screening for the academic scientific community through FAST-DEL^35^. It is our hope that this newly described DOSEDO resource will lead to the discovery of useful tool compounds for challenging targets and targets of high interest and will foster collaboration for the benefit of the broader scientific community.

## Methods

### Analytical and purification

#### NMR spectroscopy

Nuclear magnetic resonance (NMR) spectra were recorded on Bruker AV-III HD 2-channel spectrometers operating at a frequency of 400.14 MHz (^1^H) and 100.61 MHz (^13^C) and equipped with either a 5 mm BBO-F conventional (rt) probehead or a 5 mm BBO-F cryoprobe. Unless indicated otherwise, data were acquired at 25 °C using the software ICON-NMR under TopSpin program control and processed with MestReNova (Version 12.0.2-20910). Spectra were referenced to residual solvent resonance^56^. For spectra recorded at elevated temperatures, the same literature values of residual solvent resonances (at ambient temperature) were used for referencing, neglecting temperature-dependent peak shifts. Resonance signals are reported as chemical shifts (δ) in ppm with multiplicity (s = singlet, d = doublet, t = triplet, q = quartet, m = multiplet or unresolved, br = broad signal), couplings constant(s) in Hertz (Hz) and integral (not for ^13^C). ^13^C and ^19^F NMR spectra were recorded with broadband ^1^H decoupling.

#### Reaction monitoring by UPLC-MS

Samples were resolved on Waters ACQUITY ultra-performance liquid chromatography-mass spectrometry (UPLC-MS) systems equipped with C18 columns (ACQUITY UPLC BEH C18 1.7 µm, 2.1×30 mm, ACQUITY UPLC BEH C18 1.7 µm, 2.1×50 mm, or ACQUITY UPLC CSH C18 1.7 µm, 2.1×50 mm) kept at 50 °C. UV signal was recorded with ACQUITY UPLC PDA detectors (210–400 nm). Light scattering signal was recorded with ACQUITY UPLC ELSD or SofTA 1100 ELSD detectors. Low-resolution mass spectra (LRMS) of eluting species were recorded with Waters SQD, SQD-2 or QDa detectors (electron spray ionization (positive and negative modes), scan time 0.3 sec, scanning range 120-1250 Da (QDa), 120-1600 Da (SQD) or 120-2850 Da (SQD-2)). High-resolution mass spectra (HRMS) of eluting species were recorded on systems equipped with Waters Xevo G2 Qtof or Waters Xevo G2-XS Tof detectors (electron spray ionization (positive mode), scan time 0.2 sec, scanning range 100-2050 Da).<colcnt=1>

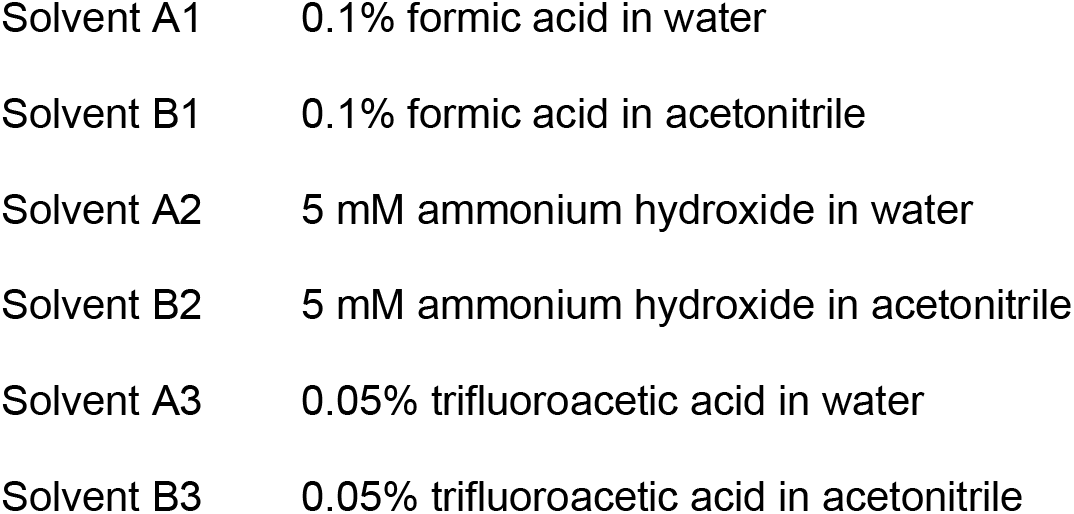

##### RXNMON-Acidic

| time | %A (solvent A1) | %B (solvent B1) |
| --- | --- | --- |
| 0.00 | 98 | 2 |
| 0.10 | 98 | 2 |
| 1.50 | 2 | 98 |
| 1.80 | 2 | 98 |
| 1.90 | 98 | 2 |
| 2.00 | 98 | 2 |

##### RXNMON-Basic

| time | %A (solvent A2) | %B (solvent B2) |
| --- | --- | --- |
| 0.00 | 98 | 2 |
| 0.10 | 98 | 2 |
| 1.50 | 2 | 98 |
| 1.80 | 2 | 98 |
| 1.90 | 98 | 2 |
| 2.00 | 98 | 2 |

##### ProductAnalysis-Acidic

| time | %A (solvent A1) | %B (solvent B1) |
| --- | --- | --- |
| 0.00 | 98 | 2 |
| 4.40 | 2 | 98 |
| 5.15 | 2 | 98 |
| 5.19 | 98 | 2 |

##### ProductAnalysis-Basic

| time | %A (solvent A2) | %B (solvent B2) |
| --- | --- | --- |
| 0.00 | 98 | 2 |
| 4.40 | 2 | 98 |
| 5.15 | 2 | 98 |
| 5.19 | 98 | 2 |

##### HRMS

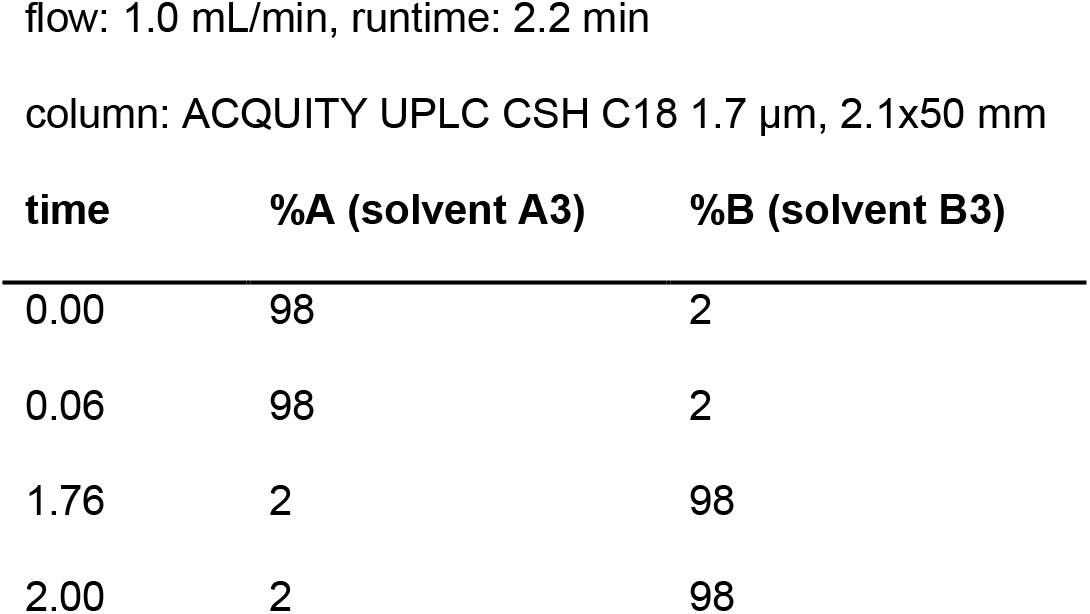
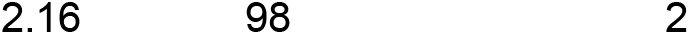

#### Ion-pair chromatographic analysis of DNA-conjugates by UPLC-MS

Samples were resolved on a Waters ACQUITY system equipped with a C18 column (ACQUITY Oligonucleotide BEH C18 1.7 µm, 2.1×50 mm, part #186003949) kept at 50 °C. UV signal was recorded with an ACQUITY TUV detector (260 nm, sampling rate 20 points/sec). Mass spectra of eluting species were recorded with a Waters SQ Detector 2 connected to a ZSpray^TM^ source (negative ion mode, scan time 0.2 sec, scanning range 500-3000 Da, MaxEnt1 deconvolution, processing 2-20 kDa).

##### Standard method (5-50% B)

| time | %A (solvent A) | %B (solvent B) |
| --- | --- | --- |
| 0.00 | 95 | 5 |
| 4.00 | 50 | 50 |
| 4.50 | 5 | 95 |
| 5.00 | 5 | 95 |
| 5.10 | 95 | 5 |
| 6.00 | 95 | 5 |

#### Preparative HPLC

Purifications were performed on a Waters high-performance liquid chromatography (HPLC) system equipped with Waters 515 pumps, a Waters 2545 binary gradient module, a Waters Acquity QDa detector, a Waters 2998 PDA detector and a Waters 2767 sample manager. The column (Waters XBridge Prep C18 OBD, 5 µm, 30 mm (inner diameter) x 50 mm) was kept at rt. Material was injected as solution in methanol/water mixtures (1.5 mL) and eluted with gradients of solvent A (water) and solvent B (acetonitrile), both modified with either 0.1% formic acid or 5 mM ammonium hydroxide (75 mL/min flowrate). Fraction collection was triggered by UV signal and mass (TIC) of eluting species. Predefined gradients were picked from a list of methods optimized for target retention time as observed during reaction monitoring by UPLC-MS.

Purification of target compounds described in Section 2.3. was performed using a Teledyne ISCO ACCQPrep HP150 equipped with a XBridge C18 column (19 x 250 mm, 5 µm), eluting with 10–100% MeCN [+1% HCO_2_H] in H_2_O [+1% HCO_2_H])

#### Preparative HPLC of oligonucleotide conjugates

Purifications were performed on an Agilent Infinity system equipped with an Infinity 1260 Bio Quat Pump (pump system, G5611A), an Infinity 1260 HiP Bio ALS (autosampler, G1330B), an Infinity 1290 TCC (thermostatted column compartment, G5667A), an Infinity 1260 DAD (diode array detector, G4212B), and an Infinity 1260 Bio FC-AS (fraction collector, G5664A) under Agilent OpenLab CDS (C.01.07 SR1) software control. The column (Waters XBridge BEH C18 OBD, 130 Å, 5 µm, 10x150 mm) was kept at 50 °C. Material was injected as aqueous solutions (up to 100 µL per injection) and eluted with gradients of solvent A (50 mM triethylammonium acetate in water) and solvent B (MeOH). Fraction collection was triggered by UV (260 nm).<colcnt=1>

##### 5-50% method

| time | %A (solvent A) | %B (solvent B) |
| --- | --- | --- |
| 0.0 | 95 | 5 |
| 15.0 | 50 | 50 |
| 16.0 | 5 | 95 |
| 18.0 | 5 | 95 |
| 18.1 | 95 | 5 |
| 20.0 | 95 | 5 |

Following separation, desired species-containing fractions were pooled and concentrated by ultrafiltration (Millipore Amicon Ultra-15/4 centrifugal filters; Ultracel 10K/3K) according to manufacturer guidelines. Three cycles of sample concentration/buffer dilution with nuclease-free water were employed.

#### Other methods and equipment

Centrifugation was performed with a Beckman Coulter Allegra X-15R or Beckman Coulter Microfuge 18. Aqueous solutions of DNA were quantified spectrophotometrically with a Thermo Scientific Nanodrop One. Electrophoresis was performed with an Invitrogen E-GEL Power Snap device using commercial gels (E-Gel EX, 4% agarose) according to manufacturer guidelines. Denaturing electrophoresis was performed using Novex™ TBE-Urea Gels (15%, Invitrogen, EC68855BOX) prepacked gels and Novex™ TBE-Urea Sample Buffer (2X, Invitrogen, LC6876) according to manufacturer guidelines. Completed gels were stained with SYBR™ Gold Nucleic Acid Gel Stain (Invitrogen, S11494) according to manufacturer directions.

#### Preparative continuous flow electrophoresis

A Bio-Rad Model 491 Prep Cell was used according to manufacturer guidelines to purify crude DNA-encoded libraries following the ligation of oligonucleotides bearing a closing primer binding site sequence. The gel assembly tube (small or large) was cast with 4% agarose in 1× TBE buffer to a height of 8.5 – 9.5 cm and left to solidify over 2 hr while circulating rt water through the cooling core at approx. 100 mL/min, carefully overlaying 2 mL of a 1:1 mixture of IPA and water on the top of the gel, then standing overnight at rt without water circulation through the cooling core. No stacking gel was cast. After standard assembly of the Prep Cell (see: http://www.bio-rad.com/webroot/web/pdf/lsr/literature/M1702925.pdf) and addition of 1× TBE buffer, the crude DEL sample was loaded in Novex™ Hi-Density TBE Sample Buffer (5X, Invitrogen LC6678), and electrophoresis was run at 12 W constant power. Fraction collection was performed using a Masterflex L/S easy-load II (Cole Parmer, 77200-50) running at approx 1 mL/min and BioFrac™ fraction collector (Bio-Rad) with a fraction time window of 2 min, i.e., approx 2 mL fractions). Total electrophoresis time was roughly 3.5 hr. Fractions were analyzed by E-GEL Power Snap device (Invitrogen, G8100) using commercial gels (Invitrogen, E-Gel EX, 4% agarose). Product containing fractions of adequate purity were pooled, then concentrated/desalted using Millipore Amicon Ultra-15/4 centrifugal filters; Ultracel 10K according to manufactures guidelines, washing at least twice with nuclease-free water.

#### Flash chromatography

Automated flash chromatography was performed on Teledyne ISCO CombiFlash^®^ systems equipped with TUV and ELSD detectors using prepacked columns (RediSep^®^ Rf, prepacked with 4 g, 12 g, 24 g, 40 g, 80 g, 120 g, or 330 g silica, 35-70 µm). Crude material was usually adsorbed on diatomaceous earth (Biotage Isolute HM-N) and subjected to chromatography (dry-loading), eluting with gradients of ethyl acetate in heptane, methanol in CH_2_Cl_2_, or as specified. Fractions containing homogeneous material according to detection method (TUV/ELSD) and/or thin layer chromatography (TLC) were pooled, and solvents were removed by rotary evaporation under reduced pressure at 30–45 °C to yield target compounds.

#### Compound synthesis and characterization

All chemical synthesis procedures and compound characterization data are provided in the **Supplementary Information in Sections 1, 2, 7,** and **8**, and all DOSEDO library member structures can be found in **Supplementary Files 2** and **3.**

### Preparation of NGS libraries

Barcodes of a DEL sample (input or eluted after screening) were PCR amplified using 3 µL i5 index primer (10 µM stock in water), 3 µL i7 index primer (10 µM stock in water), 19 µL cleaned up elution samples, and 25 µL Invitrogen Platinum™ Hot Start PCR Master Mix (2×) (Invitrogen 13000012). The PCR method is as follows: 95 °C for 2 min; 22 cycles of 95 °C (15 s), 55 °C (15 s), 72 °C (30 s); 72 °C for 7 min; hold at 4 °C. The PCR products were cleaned up using the ChargeSwitch PCR Clean-Up Kit, pooled in equimolar amounts, and the 187 bp amplicon was gel purified using a 2% E-Gel™ EX Agarose Gels (ThermoFisher Scientific G401002) and QIAquick Gel Extraction Kit (Qiagen 28704). The DNA concentration was measured using the Qubit dsDNA BR assay kit and sequenced using a HiSeq SBS v4 50 cycle kit (Illumina FC-401-4002) and HiSeq SR Cluster Kit v4 (Illumina GD-401-4001) on a HiSeq 2500 instrument (Illumina) in a single 50-base read with custom primer CTTAGCTCCCAGCGACCTGCTTCAATGTCGGATAGTG and 8-base index read 32 using custom primer CTGATGGAGGTAGAAGCCGCAGTGAGCATGGT (**Supplementary Fig. 11**).

### DEL screening

The protein targets and beads-only control (B buffer replacing the protein) were screened in duplicate using a KingFisher Duo Prime (Thermo Scientific) in a 96-well deepwell plate (Thermo Scientific 95040452) at room temperature. C-His tagged IDH1 R132H (Met1-Leu414) was purchased from G-Biosciences (BAN1708, 50 µg). N-His and GST tagged USP7 (Lys208-Glu560) were purchased from SinoBiological (11681-H20B, 100 µg) and reconstituted in 400 µL water. The buffers used are ‘B Buffer’ containing 25 mM HEPES pH 7.4, 150 mM NaCl, 0.05% Tween-20 (w/v), and ‘S Buffer’ containing 25 mM HEPES, pH 7.4, 150 mM NaCl, 0.05% Tween-20 (w/v) and 0.3 mg/mL Ultrapure Salmon Sperm DNA (ThermoFisher Scientific 15632011). His-Tag Dynabeads (Invitrogen 10103D, 10 µL per sample) was washed three times with B buffer before protein immobilization. The proteins were diluted to 1.2 µM in B buffer (100 µL per sample) and immobilized to the washed beads (1 hr, medium mix). The beads were washed once with B buffer (200 µL) and twice with S buffer (200 µL) (3 min each, medium mix). The beads were transferred to the DEL library (1 million copies per library member, 100 µL in S buffer) and incubated (1 hr, medium mix). The beads with protein (except beads-only control) and DEL bound were washed once with S buffer (200 µL) and twice with B buffer (200 µL) (3 min each, medium mix). The beads were transferred to B buffer (100 µL) and heated (90 °C, 10 min) to elute DEL compounds into the supernatant. 20 µL of the elution was restriction digested by 0.1 µL StuI (NEB R0187) in 56.5 µL 1×SmartCutter buffer (NEB B7204S) per sample (37 °C, 1 hr) and cleaned up using the ChargeSwitch PCR Clean-Up Kit (Thermo Scientific CS12000) (**Supplementary Fig. 15**).

### Assay details

#### Carbonic anhydrase

Carbonic anhydrase II (CAII) inhibition assays were performed using a commercially available kit (BioVision, #K473-100), following the manufacturer’s protocol. For each assayed DOSEDO compound, 11 concentrations (half-log dilutions from 10 µM to 100 pM) were tested, with three technical replicates performed on each of two separate 384-well µClear® medium-binding, flat-bottom polystyrene microplates (Greiner Bio-One, #82051-294) (n=6 for each compound concentration); for **53** only, six technical replicates (n=6) were performed on the same plate for each compound concentration. Each reaction well contained 3 µL of compound (prepared in 10% DMSO, 90% kit buffer), 25.5 µL of enzyme/buffer mix (1.4 µL enzyme, 24.1 µL kit buffer), and 1.5 µL of substrate, for a total volume of 30 µL. Absorbance at 405 nm was read on a Molecular Devices SpectraMax i3x. In addition to the experimental conditions, a DMSO control and a no-enzyme control were each performed with three technical replicates on each plate, except for the experiment with **53**, where six technical replicates for each control were performed on the same plate. The average change in absorbance for the no-enzyme control was subtracted from the change in absorbance over the same time interval for each experimental condition and the DMSO control on the same plate. For each concentration of each compound, relative activity was calculated separately for each replicate based on the average change in absorbance for the DMSO control on the same plate. Dose-response curves were fitted and IC50 values and 95% confidence intervals were calculated using GraphPad Prism v9.2.0. Curves were fitted using GraphPad Prism’s log(inhibitor) vs. response – variable slope nonlinear regression equation, with the top asymptote constrained to 100. Outliers were detected and removed via GraphPad Prism’s ROUT method, with a Q-value of 1%. Error bars (only shown if greater than the height of the symbol) represent 95% confidence intervals. See **Supplementary Fig. 16**.

#### Recombinant homodimeric IDH1 R132H production

The plasmid pET-22b(+) containing the recombinant IDH1 R132H gene (Met1-Leu414) fused with a His6 tag at the C terminus, via Ndel and Xhol restriction sites, was expressed in the *E. coli* BL21(DE3) strain. A starter culture was grown in 100 mL of LB media supplemented with ampicillin (50 μg/mL) at 37 °C overnight. The starter culture (1:100 v/v) was used to inoculate 3 liters of LB media supplemented with ampicillin (50 μg/mL) and grown at 37 °C to an OD600 nm= 0.6. Expression was induced with 1 mM isopropyl-β-D-1-thiogalactopyranoside (IPTG) at 20 °C for 14 hr. The cells were collected by centrifugation (Thermo Scientific™ Sorvall™ LYNX 6000 centrifuge) at 8,000 × g for 10 min. The pellets were suspended in 30 mL of lysis buffer containing 20 mM Tris-HCl, 500 mM NaCl, pH 7.4, 1% Triton X-100, lysozyme (1 mg/mL), benzonase (10 μL), 1 mM TCEP and 1 mM phenylmethylsulfonyl fluoride (0.1 M solution in ethanol, 300 uL). The cells were lysed on ice by sonication (50% amplitude, 3 s pulse on and 3 s pulse off, for total 7 min ‘on’ time). The cell debris was precipitated by centrifugation (Thermo Scientific™ Sorvall™ LYNX 6000 centrifuge) at 30,000 × g for 60 min. The cell lysates were filtered through a 0.45 µm syringe filter and purified by affinity chromatography using a 5 mL HisTrap HP column (GE Healthcare Life Sciences) with wash buffer (20 mM Tris-HCl, 500 mM NaCl, pH 7.4) and a gradient of 4-100% elution buffer (20 mM Tris-HCl, 500 mM NaCl, 250 mM imidazole, pH 7.4) over 10 CV. The fractions containing IDH1 R132H were pooled, concentrated and further purified using a Superdex S200 26/600 200 pg column in 20 mM Tris-HCl, 100 mM NaCl, pH 7.4. Pure fractions of IDH1 R132H were pooled, concentrated, aliquoted and flash frozen before storage at -80 °C.

#### IDH1 R132H and USP7 surface plasmon resonance

IDH1 R132H: SPR experiments were performed on a Biacore T200 instrument (GE Healthcare) at 25 °C using a running buffer containing 50 mM HEPES pH 7.5, 50 mM KCl, 0.005% tween-20, 1 mM DTT, 2% DMSO. The compounds were 2-fold serial diluted from 20 µM to 78 nM with a 2% DMSO only control. Biotinylated homodimeric IDH1 (RU ∼3500) was immobilized to the streptavidin sensor chip (Cytiva BR100531) preconditioned with 1M NaCl and 40 mM NaOH (60 s ×3 cycles, 100 µL/min) and running buffer (60 s, 100 µL/min). The run has a startup run of 12 cycles using the running buffer, flow rate of 50 µL/min, contact time of 60 s and dissociation time of 150—500 s depending on the compound, with two negative controls using the running buffer between each compound. The syringe was washed with 50% DMSO between injections. Data was analyzed using the affinity mode of the Biacore T200 Evaluation Software and is presented in **Supplementary Fig. 17**.

USP7 and GST: SPR experiments were performed on a Biacore T200 instrument (GE Healthcare) at 25 °C using a running buffer containing 20 mM HEPES pH 7.5, 100 mM NaCl, 0.005% tween-20, 0.2 mM TCEP, 1% DMSO. The compounds were 2-fold serial diluted from 20 µM to 78 nM with a 1% DMSO only control. His-GST-tagged USP7 (SinoBiological 11681-H20B) and His-tagged GST (Abcam ab89494) were immobilized to the streptavidin sensor chip (Cytiva BR100531) via two different channels. The streptavidin sensor chip was preconditioned with 1M NaCl and 40 mM NaOH (60 s ×3 cycles, 100 µL/min), running buffer (60 s, 100 µL/min), and functionalized to capture His-tagged proteins by 350 mM EDTA (60 s), 1 µM Tris-NTA biotin trifluoroacetate salt solution (Sigma-Aldrich 75543) (100 s), 500 µM NiCl_2_ (100 s), 3 mM EDTA (60 s) (for all functionalizing steps: flow rate 100 µL/min, reagents in running buffer). The run has a startup run of 12 cycles using the running buffer, flow rate of 50 µL/min, contact time of 60 s and dissociation time of 150 s. The syringe was washed with 50% DMSO between injections. Data were analyzed using the affinity mode of the Biacore T200 Evaluation Software, shown in **Supplementary Fig. 18**.

#### IDH1 R132H differential scanning fluorimetry

Thermal shift experiments were carried out using a QuantStudio^TM^ 7 Flex system (Applied Biosystems) in a MicroAmp Optical 384-well plate (Applied Biosystems 4309849)^57^. Each well contained 10 µL of 5 µM protein with 10× Sypro Orange in 50 mM Tris-HCl, pH 7.4. Various concentrations of compounds (6.25 – 100 µM, 1% DMSO final) were mixed with the protein for thermal stabilization studies. The samples in triplicates were subjected to temperature increases from 25 °C to 95 °C at 0.02 °C s^-1^, with optical filters of x1-m3 corresponding to excitation 470 nm and emission 586 nm respectively. Protein Thermal Shift Software v1.3 (Applied Biosystems) was used to determine the melting temperature, T_m_, from the derivative of the melt curve. Data are shown in **Supplementary Fig. 19.**

#### IDH1 R132H enzymatic inhibition

The inhibitory activity of the off-DNA compounds against IDH1 R132H were assessed by absorbance assays similarly to reported procedures^58^, using IDH1 R132H purified in-house as described above. The conversion of 2-oxoglutarate (2OG) and NADPH to 2-hydroxyglutarate (2HG) and NADP^+^ catalyzed by IDH1 R132H were monitored by measuring NADPH absorbance at 340 nm in 96-well half area clear microtiter plates (Greiner Bio-One 675001) in a final volume of 100 µL continuously over 1 hr at room temperature. The assay buffer consists of 100 mM Tris-HCl, 100 mM NaCl, 10 mM MgCl_2_, 0.005% Tween-20, pH 7.5, 0.1 mg/mL bovine serum albumin and 0.2 mM dithiothreitol (DTT).

Percentage inhibition of IDH1 R132H was measured by diluting 1 mM compound stock in DMSO to 40 µM in the assay buffer (25 µL, 10 µM compound and 1% DMSO final), incubated with IDH1 R132H (25 µL, 30 nM final) for 12 min, followed by the addition of 2OG (25 µL, 1.5 mM final) and NADPH (25 µL, 50 µM final) to initiate the reaction. The difference in absorbance, Δ*A*_340_, in the linear range of the reaction profile was converted to % residual activity with the DMSO control being the 100% residual activity reference. The % inhibition is calculated by (1 − activity with inhibitor/activity with DMSO control) × 100%. IC_50_ of the compounds were measured in a comparable manner with a 3-fold serial dilution of the compounds from 10 µM to 0.169 nM (final concentration) plus a 1% DMSO control. Percent residual activities at 11 compound concentrations were fitted using GraphPad Prism to obtain the IC_50_ value. Data are shown in **Supplementary Fig. 20**.

### In silico

#### Building block selection

Building block selection was performed through a semi-automated process. Building blocks were sourced from available internal stocks of commercial building blocks and newly ordered compounds (primarily from Enamine). Knime was used to filter to a diverse set of building blocks in each desired reactivity class for manual review and selection for purchasing. The detailed workflow and explanation are shown in **Supplementary Fig. 21**.

A vendor catalogue was used as input to the described workflow, with structures in SMILES format. In the workflow illustration, the input structures were all ‘amines’ in the Enamine catalogue at the time (28,976 compounds). Input structures were desalted, converted to canonical SMILES, and replicate molecules were removed (27,296 compounds). A manual definition of a target functional group was defined as substructure filter, in this example, a primary amine moiety without other restrictions using the SMARTS query: [#7;h2] (15,700 compounds). A set of hierarchical substructure filters were applied to remove functionality that would be undesirable in the specified building block set. Filters included various alkyl halides, carboxylic acids, thiols, thioethers, hydrazines, amino alcohols, and in the case of primary amines, for example, any additional secondary amine was excluded (11,289 compounds). A ‘dummy’ library was enumerated using this filtered set of building blocks by fixing the skeleton and ‘other’ building block of a three-component combinatoric library as representative members. As an example, the reaction SMARTS for the set of primary amines was: [#7;h2:1] >>[#6]-[#7]-[#6](=O)-[#6]-1-[#6](-[#6]-[#6]-[#7]-1-[#6]-[#6]-1=[#6]-[#6](C#N)=[#6](-[#6]-2-[#6]-[#6]-2)-[#6](F)=[#6]-1)-[#6]-1=[#6]-[#6](-[#7;h1:1])=[#6]-[#6]=[#6]-1. Molecular descriptors were calculated for this dummy library, including exactMW, SLogP, HBD, HBA, fsp3, and TPSA. Compounds were further filtered by hard cut-offs of SLogP <6.0 and exactMW <600, which in this ‘primary amine’ example reduced the available pool of building blocks to 7,699. RDKit Diversity Picker was employed under default settings (based on Morgan 2 fingerprint diversity) to select 500 compounds from the pool. These 500 were naturally biased towards larger building blocks, so to enrich the building block selection towards those with more desirable physicochemical properties, a second round of selection was performed. The second round focused on building blocks with SLogP <5 and exactMW <450 (when incorporated as dummy enumerated library members). Typically, as many as available of the second-round building blocks were selected for inclusion in the final list of selected building blocks; however, if several hundred met the selection criteria, a set of 200 was chosen, using the initial 500 as template structures to bias away from. Final lists were written to a file, and their structures were reviewed manually before ordering. In general, labile moieties were omitted, and racemic building blocks were only included if they were considered interesting by the reviewing chemist(s).

Once delivered, selected building blocks were split into multiple 2D-barcoded matrix tubes to facilitate development of optimal conditions for a given building block’s use in different contexts. This organization was achieved through a semi-automated process of Tecan-guided dissolution in a highly volatile solvent (or solvent mixture), followed by transfer and gentle solvent removal using a Genevac HT-12 centrifugal evaporator. Building blocks were stored in a Hamilton robotic tube handler as dry powder/oil in roughly 1 mg portions.

#### Library enumeration

Enumeration of the entire DEL (final targets associated to the merged Cy1-4 sequences) was performed using Knime Analyics Platform. To represent the assumed pharmacophore of any future putative hit, the skeletons were normalized in several ways, detailed below:

1. Skeletons containing alkyl linkers to their DNA attachment site (e.g., O=C(O)CCCC(=O)N1C[C@H](c2ccc(I)cc2)C2(CNC2)C1) were truncated down to the corresponding methyl amides, marking the DNA attachment site with a tritium atom (e.g., [3H]CC(=O)N1C[C@H](c2ccc(I)cc2)C2(CNC2)C1).
2. Skeletons linked to DNA via a carbamate linkage (e.g., OC[C@@H]1[C@H](c2ccc(Br)cc2)[C@H]2CNCCCCN12) were represented in the enumerated library as methyl ethers, again marking the DNA attachment point with a tritium atom (e.g., [3H]COC[C@@H]1[C@H](c2ccc(Br)cc2)[C@H]2CNCCCCN21).
3. Skeletons linked to DNA via an amide linkage (e.g., O=C(O)[C@@H]1C[C@H](Oc2ccc(I)cc2)CN1) were represented in the enumerated library as methyl amides, again marking the DNA attachment point with a tritium atom (e.g., [3H]CNC(=O)[C@@H]1C[C@H](Oc2ccc(I)cc2)CN1).
4. Skeletons containing a functionality known to completely and cleanly hydrolyze during synthesis were modified accordingly (e.g., the ethyl ester of CCOC(=O)c1cc2c(c(-c3cccc(Br)c3)n1)[C@H](CCO)NC2 becomes [H]OC(=O)c1cc2c(c(-c3cccc(Br)c3)n1)[C@H](CCOC[3H])NC2).

Amine-capping reactions were performed *in silico* on the pool of modified skeletons according to the below reaction definitions, followed by filtering to products derived from reactivity at the most nucleophilic N-atom in each skeleton.

> N-capping reaction definitions (as SMARTS):
>
> Sulfonylation: [#7:1].[Cl:2][S:3](=[O:4])=[O:5]>>[#7:1][S:3](=[O:4])=[O:5]
>
> Reductive amination: [#7:1].[*:4]-[#6:2]=[O:3]>>[#7:1]-[#6;h2:2]-[*:4]
>
> Acylation: [#7:1].[#8;h1:2]-[#6:3]=[O:4]>>[#7:1]-[#6:3]=[O:4]

Encoding information was joined to structure based on canonical SMILES representations of the skeletons and building blocks used for each ‘reaction’. Products of N-capping reactions were labelled as ‘intermediate’.

Suzuki couplings were performed *in silico* on the pool of intermediates according to the reaction definitions shown below, again joining encoding information to enumerated structures based on canonical SMILES representations of molecules used in each ‘reaction’. Null reactions (i.e., no boronic acid/ester present and either with Pd-catalyst and heated, or without Pd-catalyst and not heated) were also encoded accordingly.

Suzuki coupling reaction definitions (as SMARTS):

> I[*:1].[#8]-[#5](-[#8])-[*:2]>>[#6:2]-[*:1]
>
> Br[*:1].[#8]-[#5](-[#8])-[*:2]>>[#6:2]-[*:1]

No effort was made to directly enumerate all expected side products deriving from each building block (e.g., carboxylic ester and nitrile hydrolysis and TFA-protecting group removal).

Complete enumeration afforded 3,724,965 unique barcodes, representing 3,688,975 unique compounds. Duplicate structures arose from: i) ‘null Suzuki couplings’, and ii) two analogous Suzuki couplings using a boronic acid and its pinacol ester (CC1(C)OB(c2ccc(S(C)(=O)=O)cc2)OC1(C)C and CS(=O)(=O)c1ccc(B(O)O)cc1).

A selection of properties was calculated for the enumerated library and plotted for a quick overview of the whole collection (**Supplementary Fig. 22**).

## Supporting information

SMILES strings for DOSEDO-Br library members

SMILES strings for DOSEDO-I library

## Data Availability

The datasets generated and/or analyzed during the current study are available from the corresponding author upon reasonable request.

## Acknowledgements

We thank all members of the Novartis DNA-encoded library platform group for many fruitful discussions and their valuable input; Ritesh Tichkule, Thomas Dice, Aaron Hohos, Greg Wendel and Scott Bowes for their guidance with compound management; Phil Michaels, Max Cushner and Carmelina Rakiec for analytical support; Rishi Arora for his SPR advice, as well as Patricia Horton and Michael Romanowski for providing biotinylated IDH1 R132H; and Ellen Crawford and Ayako Honda for support with outsourcing of chemical synthesis. The research was supported in part by the National Institute of General Medical Sciences (R35GM127045 awarded to S.L.S.) and by the NIBR Scholar’s Program.

## Author Contributions

L.H., B.K.H., C.J.G., H.L.O., S.L., A.G.R., S.B., J.E.B., P.A.C., B.M., C.S-Y.H, F.B., S.L.S., F.J.Z, and K.B. contributed to the study design. L.H., J.W.M., C.J.G., and G.X. performed skeleton/scaffold synthesis. L.H., J.W.M., B.K.H., and C.J.G. performed on-DNA synthesis. M.J.R.R., J.R.T., J.M.K., Z.Y.T., P.K., and B.I.C.T. performed hit synthesis. S.L., A.G.R., and K.S.L. performed screens. M.V.W., H.L.O., J.P., J.C., and C.M.G. contributed to data generation. M.V.W. and K.S.L. performed data analysis. L.H., J.R.T., B.K.H., S.L., A.G.R., S.B., N.J.S., and C.W.C. performed data analysis and interpretation. J.O., C.E.D., J.V.S., and A.M.E.F. are responsible for technical enablement of the work described. L.H., S.L., A.G.R., and K.S.L. drafted the manuscript and Supplemental Information. L.H., S.B., J.V.S., F.B., and A.M.E.F. revised the manuscript and Supplementary Information.

