## Supplemental Methods, Tables, and Figures for "Diversity-oriented synthesis encoded by deoxyoligonucleotides"

#### 1. Synthesis: off-DNA

##### 1.1. β-CH arylated cyclic amino acids

Synthesis followed published protocols[2] with minor adaptations as detailed below with reference to the general scheme outlined in **Supplementary Scheme 1**.

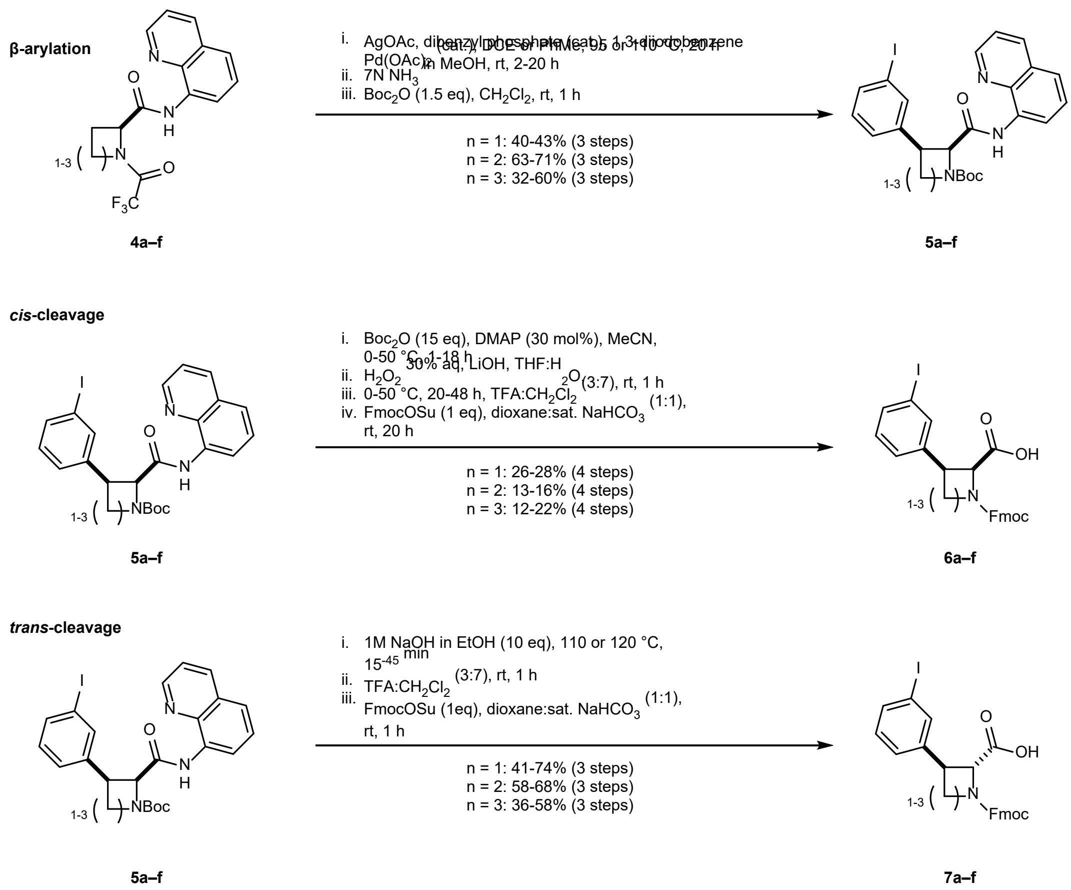

**Supplementary Scheme 1 | Overview of synthetic route to meta-I-β-aryl cyclic Fmoc amino acids**

General β-arylation sequence protocol:

Arylation precursor (**4a–f**, 1 eq), dibenzyl hydrogen phosphate (0.2 eq), AgOAc (2 eq), $Pd(OAc)_2$ (0.1 eq) and 1,3-diiodobenzene (2 eq) were added to a reaction vessel (rbf or vial). Reaction solvent (DCE or toluene, 1 M) was added and the reaction vessel was evacuated of air by five cycles of vacuum and $N_2$ supply. The reaction vessel was placed in a preheated reaction block at 95 or 110 °C, and the mixture was stirred vigorously for 20 hr. After cooling to rt 7N $NH_3$ in MeOH (3 mL of solution per 0.2 mmol reactant) was added to the residue and stirred at rt until complete TFA-cleavage was observed by LCMS analysis, then removed volatiles under reduced pressure. The resulting residue was dissolved as far as possible in $CH_2Cl_2$ (0.1 M) before adding $Boc_2O$ (1.5 eq) and stirred at rt until complete Boc-protection could be observed. Mono-Boc intermediate, **5a–f,** was partially purified by normal phase flash chromatography in a solvent system of 0-X% EtOAC in heptane (azetidines: X = 40; pyrrolidines: X = 30; piperidines: X = 20).

*cis*-cleavage:

**5a–f** (1 eq) was added to a reaction vial before dissolving in MeCN (0.2 M), adding $Boc_2O$ (15 eq) and DMAP (0.1–0.3 eq). The mixture was then stirred between rt and 50 °C until complete conversion of mono- to di-Boc-protected intermediate was observed. Volatiles were removed under reduced pressure, and the deep red residue was dissolved in a 2:1 mixture of THF and water (0.2 M) before cooling to 0 °C. 30% $H_2O_2$ (10 eq, water) was added, followed by LiOH (6 eq), and stirring was continued at 0 °C for 1 hr, then warmed to 50 °C for 20 h. The crude reaction mixture was neutralized with 4 N HCl in 1,4-dioxane and volatiles were removed under reduced pressure. The residue was treated with TFA (as a 3:7 mixture with $CH_2Cl_2$; 10 mL per mmol of reactant). Upon completion of Boc-cleavage volatiles were removed under reduced pressure. To the residue was added 1,4-dioxane (5 mL per 1 mmol of reactant), saturated $NaHCO_3$ (5 mL per 1 mmol of reactant), and FmocOSu (1 eq) before stirring at rt overnight. The crude reaction mixture was diluted with 1 N aq. HCl and extracted with $CH_2Cl_2$ (three times). Combined organic portions were dried over $Na_2SO_4$, filtered, and solvent removed under reduced pressure. Crude Fmoc-protected amino acid, **6a–f**, was purified by preparative HPLC.

*trans*-cleavage:

**5a–f** (1 eq) was added to a Biotage μW vial and dissolved in a solution of NaOH (10 eq, 1 M in EtOH). The resulting solution was heated to either 110 or 120 °C for between 15 and 45 min under μW irradiation. The crude reaction mixture was neutralized with 4N HCl in 1,4-dioxane and volatiles were removed under reduced pressure. The bright orange residue was treated with TFA (as a 3:7 mixture with $CH_2Cl_2$; 10 mL per mmol of reactant). Upon completion of Boc-cleavage volatiles were removed under reduced pressure. To the residue was added 1,4-dioxane (5 mL per 1 mmol of reactant), saturated $NaHCO_3$ (5 mL per 1 mmol of reactant), and FmocOSu (1 eq) before stirring at rt overnight. The crude reaction mixture was diluted with 1 N aq. HCl and extracted with $CH_2Cl_2$ (three times). Combined organic portions were dried over $Na_2SO_4$, filtered, and solvent removed under reduced pressure. Crude Fmoc-protected amino acid, **7a–f**, was purified by preparative HPLC.

List of general changes to published route[1]:

1) Normal phase flash column chromatography was only performed for β-arylated N-Boc intermediates (**5a–f**), following β-arylation, TFA-protecting group removal, and mono-Boc installation.
2) All steps commencing with β-arylated N-Boc intermediates (**5a–f**) were telescoped without intermediate purification through to final Fmoc amino acids (**6a–f** and **7a–f**).
3) Target Fmoc-protected amino acids (**6a–f** and **7a–f**) were isolated by preparative HPLC.

Ring-size specific notes:

1) Directed β-arylation of piperidines performed well at slightly reduced temperatures (i.e., 90–100 °C), and it could be noted through chiral SFC analysis that high temperatures of above 110 °C (e.g., due to hotplate instability, or from 'hot spots' if heated under μW irradiation) can lead to epimerization at the α-position.
2) Installation of the second Boc group (*cis*-cleavage pathway) was facile for azetidines under published conditions, required approx 15 eq of $Boc_2O$ and 10 mol% DMAP for pyrrolidines, and 15 eq of $Boc_2O$ with 30 mol% DMAP and heating at 50 °C for >24 hr for piperidines.
3) Directing group cleavage (*trans*-configuration) was performed under μW irradiation. Azetidines required 15 min at 110 °C, pyrrolidines required 25 min at 110 °C and piperidines required 45 min at 120 °C. For all ring sizes some Boc-cleavage at the cyclic amine was observed, and the crude mixture could be progressed directly through complete Boc-deprotection and Fmoc installation prior to preparative HPLC purification.

Proton NMR data are reported at temperatures giving the greatest clarity for assignment of structure. In some cases, this may mean a temperature at which rotamers collapse to well-defined multiples; in other cases, rt proton NMR yielded clearly distinct rotamers, whereas higher temperatures lead to rotamer collapse with poor resolution.

(2*R*,3*S*)-1-(((9H-fluoren-9-yl)methoxy)carbonyl)-3-(3-iodophenyl)azetidine-2-carboxylic acid (**6a**)

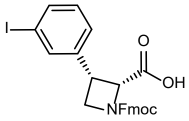

$^{1}$H NMR (400 MHz, DMSO+$D_2O$, 80 °C) δ 7.78 (d, *J* = 7.4 Hz, 2H), 7.63 – 7.51 (m, 4H), 7.33 (t, *J* = 7.6 Hz, 2H), 7.26 (q, *J* = 7.5 Hz, 3H), 7.06 (t, *J* = 7.8 Hz, 1H), 4.88 (d, *J* = 8.7 Hz, 1H), 4.24 – 4.09 (m, 6H). $^{13}$C NMR (101 MHz, $CD_3OD$) δ 170.38, 156.69, 143.74, 141.20, 139.39, 137.05, 136.46, 129.82, 127.47, 127.29, 126.82, 124.90, 124.83, 119.54, 93.41, 67.40, 66.74, 52.24, 46.25, 36.71. HRMS (ESI$^+$) m/z calculated for $C_{25}H_{20}INO_4$ $[M+H]^+$ 526.0510; found 526.0543.

(2*S*,3*R*)-1-(((9H-fluoren-9-yl)methoxy)carbonyl)-3-(3-iodophenyl)azetidine-2-carboxylic acid (**6b**)

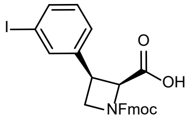

$^{1}$H NMR (400 MHz, DMSO+$D_2O$, 80 °C) δ 7.78 (d, *J* = 7.4 Hz, 2H), 7.63 – 7.51 (m, 4H), 7.33 (t, *J* = 7.6 Hz, 2H), 7.26 (q, *J* = 7.5 Hz, 3H), 7.06 (t, *J* = 7.8 Hz, 1H), 4.88 (d, *J* = 8.7 Hz, 1H), 4.24 – 4.09 (m, 6H). $^{13}$C NMR (101 MHz, $CD_3OD$) δ 143.76, 141.20, 139.46, 137.07, 136.43, 129.81, 127.46, 127.30, 126.81, 124.91, 124.85, 119.53, 93.40, 67.42, 66.87, 52.19, 46.32, 36.72. HRMS (ESI$^+$) m/z calculated for $C_{25}H_{20}INO_4$ $[M+H]^+$ 526.0510; found 526.0532.

(2*R*,3*R*)-1-(((9H-fluoren-9-yl)methoxy)carbonyl)-3-(3-iodophenyl)pyrrolidine-2-carboxylic acid (**6c**)

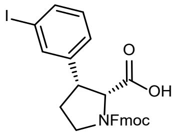

$^{1}$H NMR (400 MHz, DMSO+$D_2O$) (55:45 mixture of rotamers) δ 7.90 (app. dd, *J* = 12.3, 7.5 Hz, 2H), 7.74 – 7.62 (m, 4H), 7.48 – 7.30 (m, 5H), 7.16 (app. dt, *J* = 11.2, 7.7 Hz, 1H), 4.62 (d, *J* = 8.6 Hz, 0.45H), 4.46 (d, *J* = 8.7 Hz, 0.55H), 4.37 – 4.16 (m, 3H), 3.85 – 3.70 (m, 2H), 3.54 – 3.44 (m, 1H), 2.50 – 2.40 (m, 1H), 2.19 – 2.11 (m, 1H). $^{13}$C NMR (101 MHz, $CD_3OD$) δ 172.95, 172.79, 155.08, 154.90, 143.91, 143.89, 143.78, 143.68, 141.27, 141.23, 141.17, 141.08, 139.53, 137.00, 136.95, 136.27, 136.25, 129.79, 129.77,

127.42, 126.85, 126.80, 124.87, 124.76, 124.72, 119.56, 119.52, 119.49, 93.47, 93.44, 67.90, 67.38, 64.08, 63.79, 46.08, 45.69, 28.05, 27.11. HRMS (ESI$^+$) m/z calculated for $C_{26}H_{22}INO_4$ $[M+H]^+$ 540.0666; found 540.0693.

(2*S*,3*S*)-1-(((9H-fluoren-9-yl)methoxy)carbonyl)-3-(3-iodophenyl)pyrrolidine-2-carboxylic acid (**6d**)

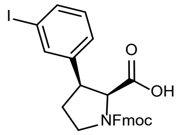

$^{1}$H NMR (400 MHz, DMSO+$D_2O$) (55:45 mixture of rotamers) δ 7.89 (app. dd, *J* = 12.1, 7.5 Hz, 2H), 7.72 – 7.60 (m, 4H), 7.46 – 7.27 (m, 5H), 7.14 (app. dt, *J* = 11.9, 7.7 Hz, 1H), 4.60 (d, *J* = 8.6 Hz, 0.45H), 4.43 (d, *J* = 8.6 Hz, 0.55H), 4.34 – 4.14 (m, 3H), 3.85 – 3.67 (m, 2H), 3.51 – 3.41 (m, 1H), 2.49 – 2.36 (m, 1H), 2.17 – 2.07 (m, 1H). $^{13}$C NMR (101 MHz, $CD_3OD$) δ 172.89, 172.74, 155.08, 154.89, 143.91, 143.89, 143.77, 143.67, 141.27, 141.23, 141.18, 141.08, 139.51, 139.48, 136.99, 136.95, 136.29, 136.26, 129.80, 129.77, 127.41, 126.85, 126.81, 124.87, 124.76, 124.72, 119.56, 119.53, 119.50, 93.48, 93.45, 67.89, 67.39, 64.04, 63.72, 46.07, 45.69, 28.03, 27.07 HRMS (ESI$^+$) m/z calculated for $C_{26}H_{22}INO_4$ $[M+H]^+$ 540.0666; found 540.0697.

(2*R*,3*R*)-1-(((9H-fluoren-9-yl)methoxy)carbonyl)-3-(3-iodophenyl)piperidine-2-carboxylic acid (**6e**)

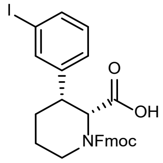

$^{1}$H NMR (400 MHz, DMSO+$D_2O$, 80 °C) δ 7.84 (d, *J* = 7.5 Hz, 2H), 7.64 (d, *J* = 7.5 Hz, 2H), 7.60 – 7.55 (m, 2H), 7.41 (t, *J* = 7.5 Hz, 2H), 7.33 (tt, *J* = 7.4, 1.6 Hz, 2H), 7.26 – 7.17 (m, 1H), 7.10 (t, *J* = 8.0 Hz, 1H), 4.83 – 4.75 (m, 1H), 4.51 – 4.40 (m, 2H), 4.28 (t, *J* = 6.4 Hz, 1H), 3.96 – 3.85 (m, 1H), 3.31 – 3.21 (m, 1H), 2.94 – 2.84 (m, 1H), 2.09 (app. qd, *J* = 13.0, 3.5 Hz, 1H), 1.85 – 1.74 (m, 2H), 1.54 – 1.41 (m, 1H). $^{13}$C NMR (101 MHz, $CD_3OD$) (not all quaternary carbon peaks resolved) δ 143.70, 143.07, 136.41, 135.58, 129.63, 127.43, 127.04, 126.76, 124.72, 124.48, 119.70, 119.57, 93.36, 67.39, 59.03, 47.17,

43.10, 40.31, 24.66, 23.69. HRMS (ESI$^+$) m/z calculated for $C_{27}H_{24}INO_4$ $[M+H]^+$ 554.0823; found 554.0853.

(2*S*,3*S*)-1-(((9H-fluoren-9-yl)methoxy)carbonyl)-3-(3-iodophenyl)piperidine-2-carboxylic acid (**6f**)

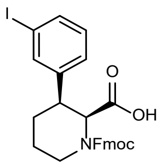

$^1$H NMR (400 MHz, DMSO+$D_2O$, 60 °C) δ 7.87 (d, *J* = 7.5 Hz, 2H), 7.65 (d, *J* = 7.4 Hz, 2H), 7.57 (d, *J* = 7.6 Hz, 2H), 7.42 (t, *J* = 7.4 Hz, 2H), 7.37 – 7.31 (m, 2H), 7.27 – 7.16 (m, 1H), 7.11 (t, *J* = 7.8 Hz, 1H), 4.85 – 4.68 (m, 1H), 4.50 – 4.35 (m, 2H), 4.29 (t, *J* = 6.5 Hz, 1H), 3.98 – 3.82 (m, 1H), 3.35 – 3.19 (m, 1H), 3.01 – 2.80 (m, 1H), 2.14 – 1.99 (m, 1H), 1.78 (app. ddd, *J* = 16.2, 12.9, 6.7 Hz, 2H), 1.46 (app. dtd, *J* = 18.2, 9.1, 4.8 Hz, 1H). $^{13}$C NMR (101 MHz, $CD_3OD$) (not all quaternary carbon peaks resolved) δ 143.78, 136.46, 135.58, 129.63, 127.43, 127.06, 126.77, 124.71, 119.57, 93.36, 67.13, 59.42, 59.05, 43.11, 40.30, 24.70, 23.70. HRMS (ESI$^+$) m/z calculated for $C_{27}H_{24}INO_4$ $[M+H]^+$ 554.0823; found 554.0860.

(2*S*,3*S*)-1-(((9H-fluoren-9-yl)methoxy)carbonyl)-3-(3-iodophenyl)azetidine-2-carboxylic acid (**7a**)

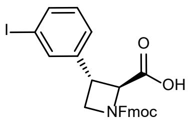

$^1$H NMR (400 MHz, DMSO, 80 °C) δ 7.86 (dt, *J* = 7.6, 1.0 Hz, 2H), 7.76 (t, *J* = 1.8 Hz, 1H), 7.71 – 7.64 (m, 3H), 7.45 – 7.39 (m, 3H), 7.34 (td, *J* = 7.4, 1.2 Hz, 2H), 7.22 (t, *J* = 7.8 Hz, 1H), 4.53 (d, *J* = 5.9 Hz, 1H), 4.38 – 4.33 (m, 2H), 4.31 – 4.23 (m, 2H), 3.88 – 3.83 (m, 1H), 3.83 – 3.75 (m, 1H). $^{13}$C NMR (101 MHz, DMSO) (C=O signals assigned from 2D spectra) δ 144.07, 143.10, 141.18, 136.59, 136.16, 131.33, 128.18, 127.59, 127.00, 125.72, 120.57, 95.67, 67.35, 66.95, 53.17, 47.12, 38.07. HRMS (ESI$^+$) m/z calculated for $C_{25}H_{20}INO_4$ $[M+H]^+$ 526.0510; found 526.0540.

(2*R*,3*R*)-1-(((9H-fluoren-9-yl)methoxy)carbonyl)-3-(3-iodophenyl)azetidine-2-carboxylic acid (**7b**)

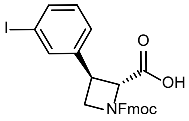

$^1$H NMR (400 MHz, DMSO, 80 °C) δ 7.86 (dd, *J* = 7.5, 1.1 Hz, 2H), 7.76 (t, *J* = 1.8 Hz, 1H), 7.71 – 7.64 (m, 3H), 7.44 – 7.38 (m, 3H), 7.33 (td, *J* = 7.5, 1.2 Hz, 2H), 7.22 (t, *J* = 7.8 Hz, 1H), 4.53 (d, *J* = 5.8 Hz, 1H), 4.38 – 4.33 (m, 2H), 4.30 – 4.22 (m, 2H), 3.88 – 3.82 (m, 1H), 3.82 – 3.75 (m, 1H). $^{13}$C NMR (101 MHz, DMSO) (<u>C</u>=O signals assigned from 2D spectra) δ 144.07, 142.53, 141.18, 136.17, 135.24, 131.33, 128.18, 127.60, 127.01, 125.72, 120.58, 95.68, 66.23, 66.11, 53.21, 47.12, 37.21. HRMS (ESI$^+$) m/z calculated for $C_{25}H_{20}INO_4$ [M+H]$^+$ 526.0510; found 526.0543.

(2*S*,3*R*)-1-(((9H-fluoren-9-yl)methoxy)carbonyl)-3-(3-iodophenyl)pyrrolidine-2-carboxylic acid (**7c**)

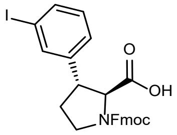

$^1$H NMR (400 MHz, DMSO) (approx. 55:45 mixture of rotamers [all peaks overlap]) δ 12.86 (s, 1H), 7.90 (app. dd, *J* = 14.4, 7.5 Hz, 2H), 7.76 – 7.60 (m, 4H), 7.47 – 7.28 (m, 5H), 7.18 (app. dt, *J* = 12.2, 7.8 Hz, 1H), 4.42 – 4.08 (m, 4H), 3.70 – 3.36 (m, 3H), 2.24 (app. ddd, *J* = 12.5, 8.5, 5.3 Hz, 1H), 2.00 (app. dt, *J* = 12.7, 7.8 Hz, 1H). $^{13}$C NMR (101 MHz, DMSO) (<u>C</u>=O signals assigned from 2D spectra) δ 144.79, 144.18, 141.24, 136.38, 136.26, 131.30, 128.17, 127.62, 125.85, 125.62, 120.60, 95.63, 67.53, 66.70, 48.49, 47.14, 46.32, 33.60. HRMS (ESI$^+$) m/z calculated for $C_{26}H_{22}INO_4$ [M+H]$^+$ 540.0666; found 540.0701.

(2*R*,3*S*)-1-(((9H-fluoren-9-yl)methoxy)carbonyl)-3-(3-iodophenyl)pyrrolidine-2-carboxylic acid (**7d**)

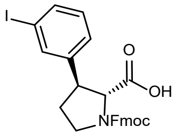

$^1$H NMR (400 MHz, DMSO) (approx. 55:45 mixture of rotamers [all peaks overlap]) δ 7.90 (app. dd, *J* = 14.4, 7.5 Hz, 2H), 7.76 – 7.60 (m, 4H), 7.47 – 7.28 (m, 5H), 7.18 (app. dt, *J* = 12.2, 7.8 Hz, 1H), 4.41 –

4.08 (m, 4H), 3.70 – 3.38 (m, 4H), 2.24 (app. dq, $J$ = 12.4, 5.9, 4.2 Hz, 1H), 2.06 – 1.94 (m, 1H). $^{13}$C NMR (101 MHz, DMSO) δ 173.06, 154.20, 154.11, 144.27, 144.18, 141.23, 141.11, 136.38, 136.26, 131.34, 131.29, 128.17, 127.65, 127.62, 127.01, 126.92, 125.72, 125.67, 125.63, 125.56, 120.64, 120.60, 120.54, 95.62, 67.54, 67.12, 65.89, 65.46, 49.16, 48.15, 47.14, 46.33, 40.61, 33.22, 32.21. HRMS (ESI$^+$) m/z calculated for $C_{26}H_{22}INO_4$ [M+H]$^+$ 540.0666; found 540.0701.

(2*S*,3*R*)-1-(((9H-fluoren-9-yl)methoxy)carbonyl)-3-(3-iodophenyl)piperidine-2-carboxylic acid (**7e**)

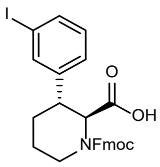

$^1$H NMR (400 MHz, DMSO, 80 °C) δ 7.86 (dt, $J$ = 7.4, 1.2 Hz, 2H), 7.69 (t, $J$ = 1.8 Hz, 1H), 7.65 – 7.57 (m, 3H), 7.41 (td, $J$ = 7.5, 3.5 Hz, 2H), 7.34 – 7.26 (m, 3H), 7.12 (t, $J$ = 7.8 Hz, 1H), 4.86 (d, $J$ = 2.5 Hz, 1H), 4.50 – 4.38 (m, 2H), 4.31 (t, $J$ = 6.6 Hz, 1H), 3.85 (d, $J$ = 13.1 Hz, 1H), 3.53 (q, $J$ = 3.9 Hz, 1H), 3.11 (td, $J$ = 12.3, 4.2 Hz, 2H), 1.87 – 1.71 (m, 2H), 1.50 – 1.30 (m, 2H). $^{13}$C NMR (101 MHz, DMSO) δ 172.53, 145.92, 144.21, 141.22, 136.65, 135.48, 130.98, 128.14, 127.61, 127.11, 125.43, 120.62, 95.53, 67.53, 57.17, 49.07, 47.11, 41.66, 38.29, 27.65, 19.29. HRMS (ESI$^+$) m/z calculated for $C_{27}H_{24}INO_4$ [M+H]$^+$ 554.0823; found 554.0859.

(2*R*,3*S*)-1-(((9H-fluoren-9-yl)methoxy)carbonyl)-3-(3-iodophenyl)piperidine-2-carboxylic acid (**7f**)

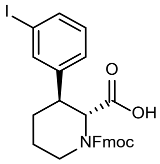

$^1$H NMR (400 MHz, DMSO, 80 °C) δ 7.86 (d, $J$ = 7.6 Hz, 2H), 7.69 (t, $J$ = 1.9 Hz, 1H), 7.62 (dd, $J$ = 11.7, 7.6 Hz, 3H), 7.41 (td, $J$ = 7.4, 3.5 Hz, 2H), 7.33 – 7.26 (m, 3H), 7.12 (t, $J$ = 7.7 Hz, 1H), 4.86 (d, $J$ = 2.5 Hz, 1H), 4.50 – 4.37 (m, 2H), 4.30 (t, $J$ = 6.8 Hz, 1H), 3.85 (d, $J$ = 13.1 Hz, 1H), 3.57 – 3.50 (m, 1H), 3.10 (dt, $J$ = 12.5, 7.2 Hz, 2H), 1.86 – 1.71 (m, 2H), 1.49 – 1.33 (m, 2H). $^{13}$C NMR (101 MHz, DMSO) δ 172.53, 145.92, 144.21, 141.22, 136.65, 135.47, 130.97, 128.14, 127.61, 127.11, 125.43, 120.61, 95.53, 67.52,

57.16, 49.07, 47.11, 41.65, 38.32, 27.65, 19.29. HRMS ($ESI^+$) m/z calculated for $C_{27}H_{24}INO_4$ $[M+H]^+$ 554.0823; found 554.0865.

Azetidine chiral SFC analysis [Column: Chiralpak IC 4.6 x 100 mm 5 μm; Mobile Phase: 5-55% MeOH w/ 10 mM $NH_4OH/CO_2$ 5 mL/min 120 bar]:

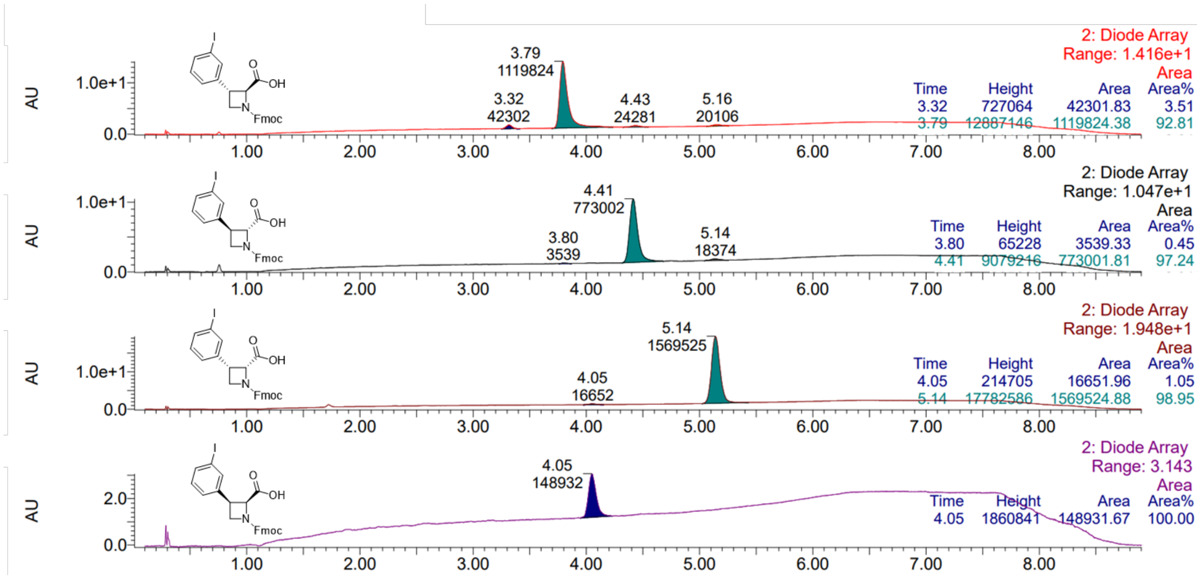

Pyrrolidine chiral SFC analysis [Column: Chiralcel OJ-H 4.6 x 100 mm 5 μm; Mobile Phase: 10% MeOH w/ 10 mM $NH_4OH/CO_2$ 5 mL/min 120 bar]:

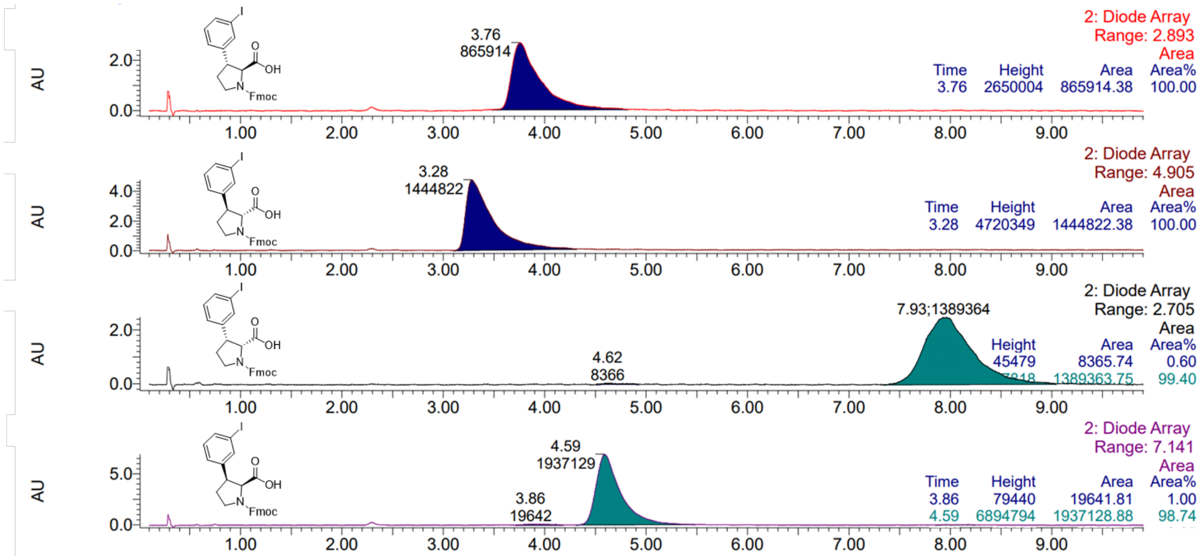

Piperidine chiral SFC analysis [Column: Chiralpak AD-3 3 x 100 mm; Mobile Phase: 5-55% 1:1 MeOH:IPA w/ 10 mM $NH_4OH/CO_2$ 2.5 mL/min 120 bar]:

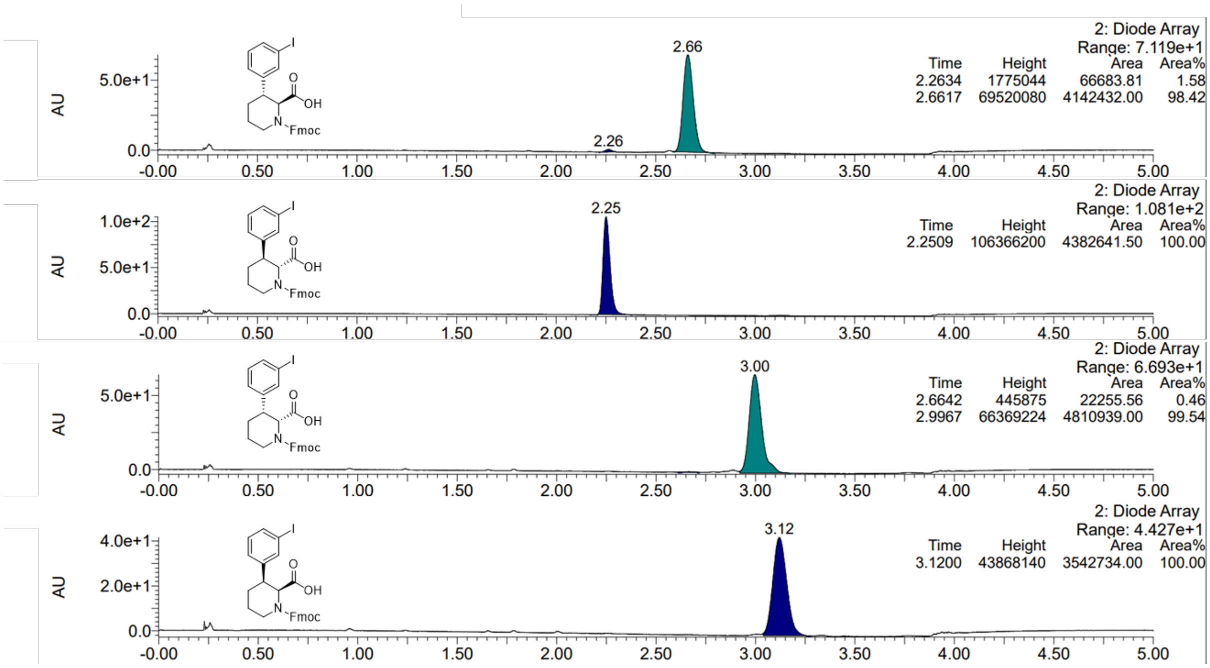

#### 1.2. Arylated and benzylated hydroxyproline stereoisomers

General preparation (A) of O-aryl hydroxyproline derivatives:

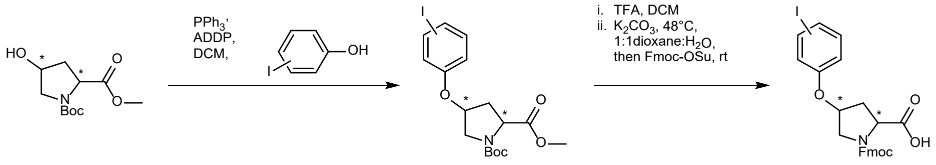

**8a–d** **9a–di–ii** **10a–di–ii**

**Supplementary Scheme 2 | General preparation of O-aryl hydroxyproline derivatives**

Part 1: A commercially sourced chirally pure isomer of N-Boc hydroxyproline methyl ester (**8a–d**, 1 eq) was added to a reaction vessel, followed by $PPh_3$ (1.4 eq), 3- or 4-iodophenol (1.2 eq) and $CH_2Cl_2$ (0.1 M). The solution was cooled to 0 °C before the addition of ADDP (1.2 eq), and the reaction was stirred overnight while allowing to warm to rt. The reaction mixture was diluted with $H_2O$ and extracted three times with $CH_2Cl_2$. Combined organic portions were evaporated onto Isolute HM-N and purified by flash chromatography to afford **9a–di–ii** as white foaming solids.

Part 2: A purified sample of **9a–di–ii** was dissolved in $CH_2Cl_2$ (0.16 M), then added TFA (40 eq) while stirring at rt. After complete Boc-deprotection could be observed by LCMS analysis the mixture was diluted with $CH_2Cl_2$ and volatile components were removed under reduced pressure. To the residue was added 1:1 1,4-dioxane:water (0.13 M) followed by $K_2CO_3$ (3–6 eq), and the resulting mixture was heated with stirring to 48 °C until complete ester hydrolysis could be observed by LCMS analysis. The mixture was cooled to rt before Fmoc-OSu (0.9 eq) was added and the mixture was stirred until complete Fmoc protection could be observed by LCMS. The crude reaction mixture was separated between ice-water and EtOAc, then pH was adjusted to approx 2 with 1 M HCl. The aqueous layer was extracted three times with EtOAc, and pooled organic portions were dried over $MgSO_4$, filtered and solvent removed under reduced pressure. **10a–di–ii** was isolated by flash chromatography with a gradient of EtOAc+0.1% AcOH in heptane.

General preparation (B) of O-benzyl hydroxyproline derivatives:

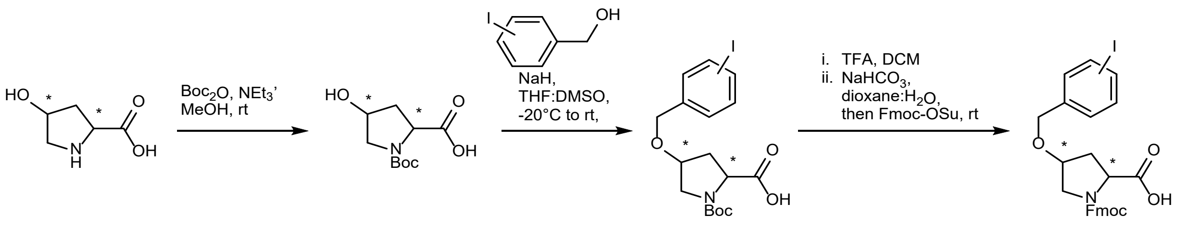

**11a–d** **12a–d** **13a–di–ii** **14a–di–ii**

**Supplementary Scheme 3 | General preparation of O-benzyl hydroxyproline derivatives**

Part 1: A commercially sourced chirally pure isomer of hydroxyproline (**11a–d**, 1 eq) was dissolved in MeOH (0.19 M) before adding $NEt_3$ (1.9 eq) and $Boc_2O$ (2 eq). The resulting mixture was heated at reflux for 2 h, then cooled to rt and stirred overnight. Volatiles were removed under reduced pressure before adding water (25 rel. vol.) and EtOAc (25 rel. vol.). The mixture was cooled to approx 0 °C before addition of $NaH_2PO_4$ (approx 10% wt/wt), adjusting pH to approx 2 with 0.2 M HCl (aq) and collecting the organic portion. The aqueous portion was extracted twice with EtOAc, and pooled organic portions were dried over $MgSO_4$, filtered and solvent removed under reduced pressure to afford **12a–d** as white foaming solids.

Part 2: NaH [60% in mineral oil] (2.5 eq) was added to a sample with septum before removing the mineral oil by rinsing with heptanes and drying under a stream of $N_2$. The resulting white solid was cooled to -20 °C with a bath of 7:3 $H_2O$:MeOH and dry ice, then a solution of **12a–d** (1 eq, 0.74 M) in a 10:1 mixture of THF:DMSO was added. After 20 min the reaction was warmed to 0 °C for 10 min, then cooled again to -20 °C before adding a solution of 3- or 4-iodobenzylbromide (1.3 eq, 0.84 M, THF). After approx 20 min the reaction mixture was warmed to rt and stirred until LCMS indicated complete consumption of starting material. The crude reaction mixture was diluted with water and EtOAc, then pH was adjusted to approx. 2 with 1 M HCl. The organic phase was isolated, and the aqueous portion was extracted twice with EtOAc. Combined organic portions were dried over $MgSO_4$, filtered and solvent was removed under reduced pressure. **13a–di–ii** were purified by flash chromatography (EtOAc/heptane) to afford a white foaming solid.

Part 3: A purified sample of **13a–di–ii** was dissolved in $CH_2Cl_2$ (0.16 M), then added TFA (20–40 eq) while stirring at rt. After complete Boc-deprotection could be observed by LCMS analysis the mixture was diluted with $CH_2Cl_2$ and volatiles were removed under reduced pressure. To the residue was added 5:1 1,4-dioxane:water (0.10 M) followed by $NaHCO_3$ (3–6 eq) and Fmoc-OSu (1.2 eq), and the mixture stirred until complete Fmoc protection could be observed by LCMS. The crude reaction mixture was diluted with 0.1 M HCl and extracted three times with EtOAc. Pooled organic portions were dried over $MgSO_4$, filtered and solvent removed under reduced pressure. **14a–di–ii** were isolated by preparative HPLC as a white solid.

(2*S*,4*S*)-1-*tert*-butyl 2-methyl 4-(4-iodophenoxy)pyrrolidine-1,2-dicarboxylate (**9ai**): Synthesized following general procedure A, Part 1. Yield: 81%.

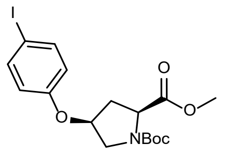

$^1$H NMR (400 MHz, $CDCl_3$) (57:43 mixture of rotamers) δ 7.58 – 7.51 (m, 2H), 6.58 (app. dd, *J* = 8.8, 3.4 Hz, 2H), 4.89 – 4.82 (m, 1H), 4.54 (dd, *J* = 8.3, 3.2 Hz, 0.43H), 4.42 (dd, *J* = 8.3, 3.5 Hz, 0.57H), 3.82 –

3.74 (m, 1H), 3.72 (app. d, *J* = 6.5 Hz, 3H), 3.71 – 3.60 (m, 1H), 2.50 – 2.40 (m, 2H), 1.46 (app. d, *J* = 18.1 Hz, 9H).

(2*S*,4*R*)-1-*tert*-butyl 2-methyl 4-(4-iodophenoxy)pyrrolidine-1,2-dicarboxylate (**9bi**): Synthesized following general procedure A, Part 1. Yield: 72%.

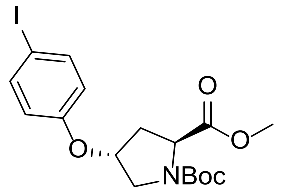

$^{1}$H NMR (400 MHz, $CDCl_3$) (3:2 mixture of rotamers) δ 7.55 – 7.49 (m, 2H), 6.64 – 6.58 (m, 2H), 4.88 – 4.80 (m, 1H), 4.45 (t, *J* = 7.7 Hz, 0.4H), 4.38 (t, *J* = 8.0 Hz, 0.6H), 3.77 – 3.73 (m, 1.6H), 3.72 (s, 3H), 3.62 (d, *J* = 11.9 Hz, 0.4H), 2.54 – 2.42 (m, 1H), 2.23 – 2.14 (m, 1H), 1.41 (app. d, *J* = 11.8 Hz, 9H). $^{13}$C NMR (101 MHz, $CDCl_3$) δ 173.24, 173.03, 156.74, 154.25, 153.57, 138.42, 117.90, 83.70, 80.45, 75.52, 74.80, 57.91, 57.54, 52.34, 52.15, 52.00, 51.76, 36.43, 35.48, 28.36, 28.24. HRMS (ESI$^{+}$) m/z calculated for $C_{17}H_{22}INO_5$ [M-Boc+H]$^{+}$ 348.0097; found 348.0141.

(2*R*,4*R*)-1-*tert*-butyl 2-methyl 4-(4-iodophenoxy)pyrrolidine-1,2-dicarboxylate (**9ci**): Synthesized following general procedure A, Part 1. Yield: 60%.

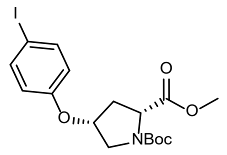

$^{1}$H NMR (400 MHz, $CDCl_3$) δ 7.53 – 7.47 (m, 2H), 6.55 (dd, *J* = 8.9, 3.4 Hz, 2H), 4.82 (dtd, *J* = 9.4, 4.7, 2.3 Hz, 1H), 4.50 (dd, *J* = 7.9, 3.4 Hz, 0.43H), 4.39 (dd, *J* = 8.5, 3.2 Hz, 0.57H), 3.74 (app. dd, *J* = 12.2, 5.0 Hz, 1H), 3.69 (s, 3H), 3.66 – 3.58 (m, 1H), 2.49 – 2.33 (m, 2H), 1.42 (app. d, *J* = 17.8 Hz, 9H). $^{13}$C NMR (101 MHz, $CDCl_3$) δ 172.35, 171.97, 156.47, 154.10, 153.71, 138.38, 117.92, 83.62, 80.29, 80.21, 75.63, 74.59, 57.77, 57.42, 52.23, 52.10, 51.87, 51.48, 36.14, 35.26, 28.39, 28.28. HRMS (ESI$^{+}$) m/z calculated for $C_{17}H_{22}INO_5$ [M-Boc+H]$^{+}$ 348.0097; found 348.0125.

(2*R*,4*S*)-1-*tert*-butyl 2-methyl 4-(4-iodophenoxy)pyrrolidine-1,2-dicarboxylate (**9di**): Synthesized following general procedure A, Part 1. Yield: 80%.

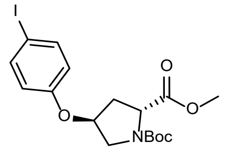

$^{1}$H NMR (400 MHz, $CDCl_3$) (3:2 mixture of rotamers) δ 7.51 (dd, *J* = 8.6, 6.0 Hz, 2H), 6.63 – 6.55 (m, 2H), 4.87 – 4.79 (m, 1H), 4.44 (t, *J* = 7.8 Hz, 0.4H), 4.37 (t, *J* = 8.0 Hz, 0.6H), 3.76 – 3.72 (m, 1.6H), 3.71 (s, 3H), 3.61 (d, *J* = 11.9 Hz, 0.4H), 2.54 – 2.41 (m, 1H), 2.23 – 2.13 (m, 1H), 1.40 (app. d, *J* = 11.9 Hz, 9H). $^{13}$C NMR (101 MHz, $CDCl_3$) δ 173.22, 173.01, 156.73, 154.24, 153.57, 138.41, 117.89, 83.68, 80.44, 75.51, 74.79, 57.91, 57.54, 52.32, 52.14, 51.99, 51.75, 36.41, 35.47, 28.41, 28.22. HRMS (ESI$^+$) m/z calculated for $C_{17}H_{22}INO_5$ [M-Boc+H]$^+$ 348.0097; found 348.0134.

(2*S*,4*S*)-1-*tert*-butyl 2-methyl 4-(3-iodophenoxy)pyrrolidine-1,2-dicarboxylate (**9aii**): Synthesized following general procedure A, Part 1. Yield: 76%.

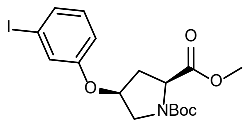

$^{1}$H NMR (400 MHz, $CDCl_3$) (3:2 mixture of rotamers) δ 7.31 – 7.24 (m, 1H), 7.16 – 7.11 (m, 1H), 7.00 – 6.93 (m, 1H), 6.78 – 6.72 (m, 1H), 4.89 – 4.81 (m, 1H), 4.53 (dd, *J* = 8.0, 3.3 Hz, 0.4H), 4.41 (dd, *J* = 8.6, 3.2 Hz, 0.6H), 3.80 – 3.73 (m, 1H), 3.71 (d, *J* = 8.6 Hz, 3H), 3.69 – 3.61 (m, 1H), 2.53 – 2.38 (m, 2H), 1.45 (app. d, *J* = 19.4 Hz, 9H). $^{13}$C NMR (101 MHz, $CDCl_3$) δ 172.35, 171.96, 157.18, 154.14, 153.74, 130.96, 130.56, 124.78, 124.71, 115.11, 94.44, 94.41, 80.37, 80.28, 75.78, 74.69, 57.79, 57.43, 52.28, 52.14, 51.95, 51.53, 36.21, 35.29, 28.42, 28.29. HRMS (ESI$^+$) m/z calculated for $C_{17}H_{22}INO_5$ [M-Boc+H]$^+$ 348.0097; found 348.0111.

(2*S*,4*R*)-1-*tert*-butyl 2-methyl 4-(3-iodophenoxy)pyrrolidine-1,2-dicarboxylate (**9bii**): Synthesized following general procedure A, Part 1. Yield: 52%.

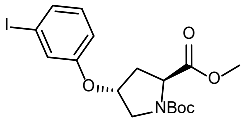

$^1$H NMR (400 MHz, $CDCl_3$) (3:2 mixture of rotamers) δ 7.34 – 7.27 (m, 1H), 7.23 – 7.19 (m, 1H), 7.03 – 6.96 (m, 1H), 6.85 – 6.79 (m, 1H), 4.91 – 4.84 (m, 1H), 4.48 (t, *J* = 7.8 Hz, 0.4H), 4.41 (t, *J* = 8.0 Hz, 0.6H), 3.80 – 3.73 (m, 4.6H), 3.66 (d, *J* = 11.9 Hz, 0.4H), 2.58 – 2.45 (m, 1H), 2.26 – 2.17 (m, 1H), 1.44 (app. d, *J* = 13.5 Hz, 9H). $^{13}$C NMR (101 MHz, $CDCl_3$) δ 173.28, 173.10, 157.42, 154.30, 153.62, 130.97, 130.70, 130.63, 124.84, 124.76, 115.11, 114.96, 94.46, 80.55, 75.70, 74.87, 57.93, 57.55, 52.37, 52.17, 52.05, 51.81, 36.49, 35.54, 28.37, 28.24. HRMS (ESI$^+$) m/z calculated for $C_{17}H_{22}INO_5$ [M-Boc+H]$^+$ 348.0097; found 348.0135.

(2*R*,4*R*)-1-*tert*-butyl 2-methyl 4-(3-iodophenoxy)pyrrolidine-1,2-dicarboxylate (**9cii**): Synthesized following general procedure A, Part 1. Yield: 86%.

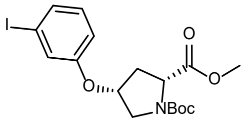

$^1$H NMR (400 MHz, $CDCl_3$) (3:2 mixture of rotamers) δ 7.27 (dd, *J* = 8.1, 4.9 Hz, 1H), 7.15 – 7.10 (m, 1H), 7.00 – 6.92 (m, 1H), 6.77 – 6.71 (m, 1H), 4.88 – 4.81 (m, 1H), 4.52 (dd, *J* = 8.0, 3.5 Hz, 0.4H), 4.40 (dd, *J* = 8.6, 3.1 Hz, 0.6H), 3.79 – 3.72 (m, 1H), 3.70 (app. d, *J* = 8.2 Hz, 3H), 3.64 (t, *J* = 13.6 Hz, 1H), 2.51 – 2.37 (m, 2H), 1.44 (app. d, *J* = 19.4 Hz, 9H). $^{13}$C NMR (101 MHz, $CDCl_3$) δ 172.33, 171.95, 157.18, 154.10, 153.71, 130.96, 130.54, 124.78, 124.70, 115.11, 94.41, 80.31, 80.23, 75.78, 74.68, 57.78, 57.42, 52.26, 52.13, 51.94, 51.52, 36.20, 35.28, 28.41, 28.29. HRMS (ESI$^+$) m/z calculated for $C_{17}H_{22}INO_5$ [M-Boc+H]$^+$ 348.0097; found 348.0124.

(2*R*,4*S*)-1-*tert*-butyl 2-methyl 4-(3-iodophenoxy)pyrrolidine-1,2-dicarboxylate (**9dii**): Synthesized following general procedure A, Part 1. Yield: 71%.

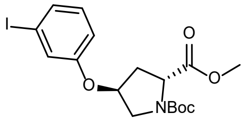

$^{1}H$ NMR (400 MHz, $CDCl_3$) (3:2 mixture of rotamers) δ 7.28 (dd, *J* = 8.2, 6.3 Hz, 1H), 7.18 (dd, *J* = 2.4, 2.0 Hz, 1H), 6.97 (q, *J* = 7.5 Hz, 1H), 6.82 – 6.76 (m, 1H), 4.88 – 4.81 (m, 1H), 4.46 (t, *J* = 7.8 Hz, 0.4H), 4.38 (t, *J* = 8.0 Hz, 0.6H), 3.77 – 3.71 (m, 4.6H), 3.64 (d, *J* = 11.9 Hz, 0.4H), 2.56 – 2.42 (m, 1H), 2.24 – 2.14 (m, 1H), 1.42 (app. d, *J* = 13.1 Hz, 9H). $^{13}C$ NMR (101 MHz, $CDCl_3$) δ 173.23, 173.04, 157.44, 157.39, 154.27, 153.60, 130.97, 130.66, 130.58, 124.83, 124.74, 115.08, 114.94, 94.46, 80.51, 75.69, 74.85, 57.91, 57.54, 52.34, 52.16, 52.05, 51.80, 36.46, 35.51, 28.35, 28.23. HRMS (ESI$^+$) m/z calculated for $C_{17}H_{22}INO_5$ $[M\text{-}Boc+H]^+$ 348.0097; found 348.0121.

(2*S*,4*S*)-1-(*tert*-butoxycarbonyl)-4-((4-iodobenzyl)oxy)pyrrolidine-2-carboxylic acid (**13ai**): Synthesized following general procedure B, Parts 1 and 2. Yield: 37% (2 steps).

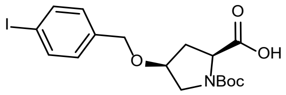

$^{1}H$ NMR (400 MHz, $CDCl_3$) (1:1 mixture of rotamers) δ 10.97 (s, 1H), 7.62 (app. t, *J* = 6.7 Hz, 2H), 7.08 – 6.92 (m, 2H), 4.53 – 4.29 (m, 3H), 4.13 – 4.09 (m, 1H), 3.69 – 3.50 (m, 2H), 2.65 (d, *J* = 13.9 Hz, 0.5H), 2.46 – 2.31 (m, 1H), 2.24 – 2.12 (m, 0.5H), 1.46 (app. d, *J* = 18.7 Hz, 9H). $^{13}C$ NMR (101 MHz, $CDCl_3$) δ 177.06, 177.04, 177.02, 174.94, 174.89, 174.85, 156.02, 154.06, 137.43, 137.31, 129.62, 129.39, 93.14,

81.48, 80.72, 76.71, 76.04, 70.06, 69.89, 57.84, 57.73, 52.87, 51.39, 35.96, 33.32, 28.38, 28.28. HRMS (ESI$^+$) m/z calculated for $C_{17}H_{22}INO_5$ [M-Boc+H]$^+$ 348.0097; found 348.0104.

(2*S*,4*R*)-1-(*tert*-butoxycarbonyl)-4-((4-iodobenzyl)oxy)pyrrolidine-2-carboxylic acid (**13bi**): Synthesized following general procedure B, Parts 1 and 2. Yield: 54% (2 steps).

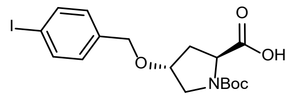

$^1$H NMR (400 MHz, $CDCl_3$) (55:45 mixture of rotamers) δ 9.62 (s, 1H), 7.72 – 7.61 (m, 2H), 7.11 – 7.00 (m, 2H), 4.51 – 4.34 (m, 3H), 4.21 – 4.14 (m, 1H), 3.73 (d, *J* = 11.8 Hz, 0.55H), 3.63 – 3.51 (m, 1.45H), 2.50 – 2.31 (m, 1H), 2.31 – 2.21 (m, 0.45H), 2.18 – 2.08 (m, 0.55H), 1.44 (app. d, *J* = 18.8 Hz, 9H). $^{13}$C NMR (101 MHz, $CDCl_3$) δ 178.11, 176.03, 155.54, 153.99, 137.56, 137.41, 137.37, 129.51, 129.43, 93.35, 81.29, 80.90, 76.68, 76.21, 70.56, 70.40, 57.91, 57.76, 51.93, 51.31, 36.63, 34.96, 28.39, 28.24.

(2*R*,4*R*)-1-(*tert*-butoxycarbonyl)-4-((4-iodobenzyl)oxy)pyrrolidine-2-carboxylic acid (**13ci**): Synthesized following general procedure B, Parts 1 and 2. Yield: 47% (2 steps).

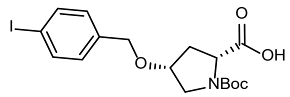

$^1$H NMR (400 MHz, $CDCl_3$) (1:1 mixture of rotamers) δ 11.41 (s, 1H), 7.38 (t, *J* = 7.2 Hz, 2H), 6.77 (t, *J* = 7.9 Hz, 2H), 4.30 – 4.08 (m, 3H), 3.90 – 3.84 (m, 1H), 3.45 – 3.31 (m, 2H), 2.31 (d, *J* = 13.7 Hz, 0.5H), 2.15 (t, *J* = 5.2 Hz, 1H), 2.06 – 1.93 (m, 0.5H), 1.24 (app. d, *J* = 18.9 Hz, 9H). $^{13}$C NMR (101 MHz, $CDCl_3$) δ 176.89, 175.59, 155.42, 154.14, 137.44, 137.33, 129.49, 129.37, 93.04, 81.01, 80.64, 76.85, 76.03, 69.98, 69.89, 57.63, 52.58, 51.38, 35.95, 33.97, 28.42, 28.28. HRMS (ESI$^+$) m/z calculated for $C_{17}H_{22}INO_5$ [M-Boc+H]$^+$ 348.0097; found 348.0102.

(2*R*,4*S*)-1-(*tert*-butoxycarbonyl)-4-((4-iodobenzyl)oxy)pyrrolidine-2-carboxylic acid (**13di**): Synthesized following general procedure B, Parts 1 and 2. Yield: 42% (2 steps).

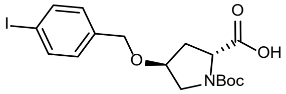

$^{1}$H NMR (400 MHz, $CDCl_3$) (3:2 mixture of rotamers) δ 10.67 (s, 1H), 7.66 (app. dd, *J* = 8.2, 3.0 Hz, 2H), 7.05 (d, *J* = 8.1 Hz, 2H), 4.50 – 4.34 (m, 3H), 4.21 – 4.13 (m, 1H), 3.73 (d, *J* = 11.7 Hz, 0.6H), 3.63 – 3.52 (m, 1.4H), 2.48 – 2.32 (m, 1H), 2.28 – 2.20 (m, 0.4H), 2.17 – 2.07 (m, 0.6H), 1.44 (app. d, *J* = 17.6 Hz, 9H). $^{13}$C NMR (101 MHz, $CDCl_3$) δ 178.04, 176.23, 155.42, 154.01, 137.43, 129.52, 129.43, 93.35, 81.19, 80.91, 76.71, 76.21, 70.54, 70.38, 57.92, 57.73, 51.93, 51.32, 36.63, 35.03, 28.40, 28.25. HRMS (ESI$^{+}$) m/z calculated for $C_{17}H_{22}INO_5$ [M-Boc+H]$^{+}$ 348.0097; found 348.0103.

(2*S*,4*S*)-1-(*tert*-butoxycarbonyl)-4-((3-iodobenzyl)oxy)pyrrolidine-2-carboxylic acid (**14aii**): Synthesized following general procedure B, Parts 1 and 2. Yield: 49% (2 steps).

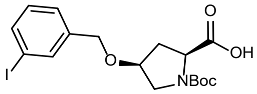

$^{1}$H NMR (400 MHz, $CDCl_3$) (1:1 mixture of rotamers) δ 9.94 (s, 1H), 7.68 – 7.55 (m, 2H), 7.31 – 7.22 (m, 1H), 7.06 (t, *J* = 7.8 Hz, 1H), 4.55 – 4.30 (m, 3H), 4.12 – 4.09 (m, 1H), 3.60 (d, *J* = 30.9 Hz, 2H), 2.66 (d, *J* = 13.8 Hz, 0.5H), 2.36 (s, 1H), 2.23 – 2.12 (m, 0.5H), 1.46 (app. d, *J* = 21.4 Hz, 9H). $^{13}$C NMR (101 MHz, $CDCl_3$) δ 176.75, 174.50, 156.18, 154.04, 140.07, 136.75, 136.53, 136.33, 130.26, 126.98, 126.72, 94.31, 81.61, 80.76, 76.62, 76.00, 69.86, 69.62, 57.87, 57.71, 52.93, 51.41, 35.79, 33.07, 28.37, 28.29. HRMS (ESI$^{+}$) m/z calculated for $C_{17}H_{22}INO_5$ [M-Boc+H]$^{+}$ 348.0097; found 348.0125.

(2*S*,4*R*)-1-(*tert*-butoxycarbonyl)-4-((3-iodobenzyl)oxy)pyrrolidine-2-carboxylic acid (**14bii**): Synthesized following general procedure B, Parts 1 and 2. Yield: 56% (2 steps).

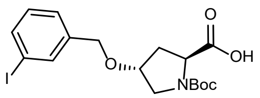

$^{1}$H NMR (400 MHz, $CDCl_3$) δ 10.12 (s, 1H), 7.66 (s, 1H), 7.63 – 7.57 (m, 1H), 7.26 (d, *J* = 7.7 Hz, 1H), 7.06 (td, *J* = 7.8, 3.3 Hz, 1H), 4.50 – 4.34 (m, 3H), 4.21 – 4.14 (m, 1H), 3.73 (d, *J* = 11.8 Hz, 1H), 3.63 – 3.53 (m, 1H), 2.50 – 2.32 (m, 1H), 2.29 – 2.20 (m, 0H), 2.18 – 2.09 (m, 1H), 1.44 (d, *J* = 17.3 Hz, 9H). $^{13}$C NMR (101 MHz, $CDCl_3$) δ 177.89, 176.16, 155.39, 154.05, 140.11, 136.86, 136.82, 136.40, 136.37, 130.25, 126.73, 126.68, 94.46, 81.19, 80.94, 76.79, 76.34, 70.25, 70.07, 57.93, 57.72, 51.92, 51.31, 36.61, 35.02, 28.42, 28.26.

(2*R*,4*R*)-1-(*tert*-butoxycarbonyl)-4-((3-iodobenzyl)oxy)pyrrolidine-2-carboxylic acid (**14cii**): Synthesized following general procedure B, Parts 1 and 2. Yield: 59% (2 steps).

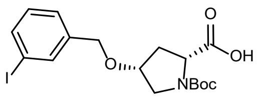

$^{1}$H NMR (400 MHz, $CDCl_3$) (1:1 mixture of rotamers) δ 11.65 (s, 1H), 7.55 (s, 1H), 7.51 (d, *J* = 8.1 Hz, 1H), 7.22 – 7.16 (m, 1H), 7.02 – 6.96 (m, 1H), 4.44 – 4.24 (m, 3H), 4.07 – 3.99 (m, 1H), 3.61 – 3.45 (m, 2H), 2.44 (d, *J* = 13.7 Hz, 0.5H), 2.32 – 2.27 (m, 1H), 2.23 – 2.08 (m, 0.5H), 1.40 (app. d, *J* = 20.6 Hz, 9H). $^{13}$C NMR (101 MHz, $CDCl_3$) δ 176.62, 175.46, 155.36, 154.13, 140.19, 136.58, 136.28, 136.16, 130.25, 126.81, 126.67, 94.30, 80.96, 80.66, 76.81, 76.01, 69.76, 69.62, 57.60, 57.54, 52.57, 51.38, 35.78, 33.89, 28.42, 28.27. HRMS (ESI$^{+}$) m/z calculated for $C_{17}H_{22}INO_5$ $[M\text{-}Boc+H]^+$ 348.0097; found 348.0121.

(2*R*,4*S*)-1-(*tert*-butoxycarbonyl)-4-((3-iodobenzyl)oxy)pyrrolidine-2-carboxylic acid (**14dii**): Synthesized following general procedure B, Parts 1 and 2. Yield: 35% (2 steps).

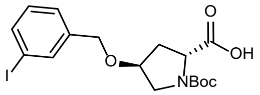

$^{1}$H NMR (400 MHz, $CDCl_3$) (3:2 mixture of rotamers) δ 10.80 (s, 1H), 7.65 (app. t, *J* = 1.7 Hz, 1H), 7.60 (app. dd, *J* = 7.9, 3.4 Hz, 1H), 7.26 (d, *J* = 7.6 Hz, 1H), 7.06 (app. td, *J* = 7.6, 3.7 Hz, 1H), 4.49 – 4.34 (m, 3H), 4.21 – 4.13 (m, 1H), 3.73 (d, *J* = 11.8 Hz, 0.6H), 3.63 – 3.54 (m, 1.4H), 2.49 – 2.32 (m, 1H), 2.28 – 2.20 (m, 0.4H), 2.13 (ddd, *J* = 13.0, 7.8, 5.0 Hz, 0.6H), 1.44 (app. d, *J* = 17.1 Hz, 9H). $^{13}$C NMR (101 MHz, $CDCl_3$) δ 177.87, 176.22, 155.35, 154.04, 140.12, 136.84, 136.81, 136.39, 136.36, 130.24, 126.73, 126.68, 94.45, 81.15, 80.93, 76.79, 76.33, 70.24, 70.06, 57.92, 57.71, 51.91, 51.31, 36.61, 35.03, 28.41, 28.25. HRMS (ESI$^+$) m/z calculated for $C_{17}H_{22}INO_5$ [M-Boc+H]$^+$ 348.0097; found 348.0097.

(2*S*,4*S*)-1-(((9H-fluoren-9-yl)methoxy)carbonyl)-4-(4-iodophenoxy)pyrrolidine-2-carboxylic acid (**10ai**): Synthesized following general procedure A, Part 2. Yield: 52%.

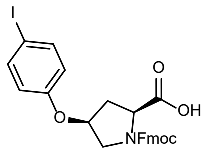

$^{1}$H NMR (400 MHz, Methanol-$d_4$) (1:1 mixture of rotamers) δ 7.80 – 7.74 (m, 2H), 7.70 – 7.59 (m, 2H), 7.59 – 7.52 (m, 2H), 7.40 – 7.34 (m, 2H), 7.34 – 7.24 (m, 2H), 6.76 – 6.68 (m, 2H), 4.93 (ddp, *J* = 12.8, 5.0, 2.3 Hz, 1H), 4.49 – 4.18 (m, 4H), 3.87 (dd, *J* = 11.9, 5.5 Hz, 0.5H), 3.76 (dd, *J* = 11.9, 5.3 Hz, 0.5H), 3.71 – 3.60 (m, 1H), 2.59 (ddd, *J* = 14.3, 9.3, 5.2 Hz, 0.5H), 2.49 (ddd, *J* = 14.4, 9.5, 5.2 Hz, 0.5H), 2.45 – 2.32 (m, 1H). $^{13}$C NMR (101 MHz, $CD_3OD$) δ 176.23, 175.85, 157.12, 157.02, 155.44, 155.26, 144.23, 144.15, 143.80, 143.75, 141.28, 141.15, 141.04, 138.10, 138.08, 127.38, 126.85, 126.77, 125.14, 125.04, 124.84, 124.80, 119.51, 119.46, 119.44, 118.08, 118.00, 82.57, 82.48, 75.59, 74.80, 67.74, 67.26, 59.27, 59.18, 52.06, 51.60, 47.03, 36.11, 35.13. HRMS (ESI$^+$) m/z calculated for $C_{26}H_{22}INO_5$ [M+H]$^+$ 556.0621; found 556.0627.

(2*S*,4*R*)-1-(((9H-fluoren-9-yl)methoxy)carbonyl)-4-(4-iodophenoxy)pyrrolidine-2-carboxylic acid (**10bi**): Synthesized following general procedure A, Part 2. Yield: 64%.

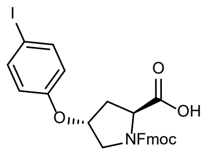

$^{1}$H NMR (400 MHz, $CDCl_3$) (3:2 mixture of rotamers) δ 11.37 (s, 1H), 7.75 (app. dd, *J* = 7.6, 5.5 Hz, 2H), 7.65 – 7.48 (m, 4H), 7.45 – 7.37 (m, 2H), 7.35 – 7.21 (m, 2H), 6.69 – 6.59 (m, 2H), 4.93 – 4.86 (m, 0.6H), 4.86 – 4.80 (m, 0.4H), 4.64 (t, *J* = 8.0 Hz, 0.6H), 4.57 – 4.38 (m, 2.4H), 4.27 (t, *J* = 7.0 Hz, 0.6H), 4.22 – 4.18 (m, 0.4H), 3.95 – 3.75 (m, 2H), 2.68 – 2.55 (m, 1H), 2.40 – 2.27 (m, 1H). $^{13}$C NMR (101 MHz, $CDCl_3$) δ 177.63, 177.48, 156.62, 156.53, 155.41, 154.69, 143.92, 143.68, 143.63, 143.61, 141.32, 141.28, 138.63, 138.54, 127.84, 127.78, 127.15, 125.02, 124.96, 120.06, 117.91, 84.08, 84.01, 75.27, 74.61, 68.04, 58.04, 57.49, 52.49, 51.88, 47.17, 47.07, 36.60, 35.30. HRMS (ESI$^+$) m/z calculated for $C_{26}H_{22}INO_5$ [M+H]$^+$ 556.0621; found 556.0630.

(2*R*,4*R*)-1-(((9H-fluoren-9-yl)methoxy)carbonyl)-4-(4-iodophenoxy)pyrrolidine-2-carboxylic acid (**10ci**): Synthesized following general procedure A, Part 2. Yield: 72%.

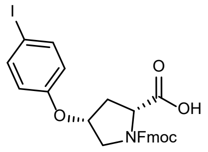

$^{1}$H NMR (400 MHz, $CDCl_3$) (55:45 mixture of rotamers) δ 11.57 (s, 1H), 7.75 (app. dd, *J* = 7.5, 4.7 Hz, 2H), 7.60 – 7.47 (m, 4H), 7.44 – 7.34 (m, 2H), 7.34 – 7.24 (m, 2H), 6.58 (app. dd, *J* = 22.0, 8.7 Hz, 2H), 4.94 – 4.87 (m, 0.55H), 4.87 – 4.81 (m, 0.45H), 4.69 – 4.32 (m, 3H), 4.28 (t, *J* = 6.9 Hz, 0.55H), 4.21 (t, *J* = 6.5 Hz, 0.45H), 3.82 – 3.69 (m, 2H), 2.64 – 2.51 (m, 1H), 2.45 (app. ddt, *J* = 13.9, 9.2, 4.3 Hz, 1H). $^{13}$C NMR (101 MHz, $CDCl_3$) δ 176.90, 176.32, 156.20, 156.17, 155.11, 154.64, 143.91, 143.71, 143.61, 141.36, 141.27, 138.52, 138.46, 127.79, 127.75, 127.14, 127.11, 125.15, 125.01, 124.91, 120.01, 118.19, 118.04, 84.16, 84.06, 75.55, 74.58, 67.85, 67.78, 57.88, 57.48, 52.10, 51.74, 47.16, 36.36, 35.15. HRMS (ESI$^+$) m/z calculated for $C_{26}H_{22}INO_5$ [M+H]$^+$ 556.0621; found 556.0648.

(2*R*,4*S*)-1-(((9H-fluoren-9-yl)methoxy)carbonyl)-4-(4-iodophenoxy)pyrrolidine-2-carboxylic acid (**10di**): Synthesized following general procedure A, Part 2. Yield: 36%.

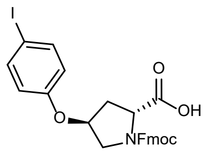

$^{1}$H NMR (400 MHz, DMSO) δ 7.90 – 7.83 (m, 2H), 7.78 (dd, *J* = 20.8, 7.5 Hz, 1H), 7.65 – 7.53 (m, 3H), 7.44 – 7.36 (m, 2H), 7.36 – 7.29 (m, 1H), 7.23 (dt, *J* = 12.7, 7.5 Hz, 1H), 6.82 – 6.76 (m, 2H), 4.99 – 4.93 (m, 1H), 4.28 – 4.09 (m, 3H), 4.06 – 3.97 (m, 1H), 3.73 (app. dt, *J* = 11.7, 5.0 Hz, 1H), 3.57 (app. ddd, *J* = 16.5, 11.7, 2.0 Hz, 1H), 2.33 (dd, *J* = 7.2, 4.6 Hz, 1H), 2.22 (dd, *J* = 7.4, 4.4 Hz, 1H). $^{13}$C NMR (101 MHz, $CD_3OD$) δ 177.10, 176.99, 157.91, 157.83, 155.47, 155.31, 144.29, 144.17, 143.86, 143.76, 141.27, 141.16, 141.13, 141.03, 130.67, 130.65, 130.04, 129.98, 127.43, 127.40, 126.89, 126.87, 126.82, 126.79, 125.28, 125.12, 124.91, 124.88, 119.58, 119.50, 114.92, 114.84, 93.65, 75.63, 74.94, 67.76, 67.23, 59.80, 52.08, 51.62, 36.33, 35.30. HRMS (ESI$^+$) m/z calculated for $C_{26}H_{22}INO_5$ $[M+H]^+$ 556.0621; found 556.0637.

(2*S*,4*S*)-1-(((9H-fluoren-9-yl)methoxy)carbonyl)-4-(3-iodophenoxy)pyrrolidine-2-carboxylic acid (**10aii**): Synthesized following general procedure A, Part 2. Yield: 29%.

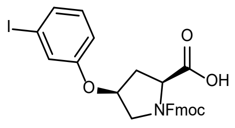

$^{1}$H NMR (400 MHz, Methanol-*d*$_4$) (1:1 mixture of rotamers) δ 7.68 – 7.60 (m, 2H), 7.60 – 7.46 (m, 2H), 7.28 – 7.12 (m, 6H), 6.94 – 6.84 (m, 1H), 6.79 (app. td, *J* = 9.0, 2.4 Hz, 1H), 4.84 – 4.70 (m, 2H), 4.36 (dd, *J* = 9.4, 3.0 Hz, 0.5H), 4.23 (d, *J* = 6.5 Hz, 2H), 4.17 (dd, *J* = 9.4, 3.3 Hz, 0.5H), 4.13 – 4.04 (m, 1H), 3.77 (dd, *J* = 11.9, 5.6 Hz, 0.5H), 3.64 (dd, *J* = 11.8, 5.4 Hz, 0.5H), 3.58 (dd, *J* = 11.9, 2.5 Hz, 0.5H), 3.51 (dd, *J* = 11.7, 2.5 Hz, 0.5H), 2.53 – 2.43 (m, 0.5H), 2.41 – 2.27 (m, 1H), 2.23 (dt, *J* = 13.8, 3.1 Hz, 0.5H). $^{13}$C NMR (101 MHz, $CD_3OD$) δ 177.10, 176.99, 157.91, 157.83, 155.47, 155.31, 144.29, 144.17, 143.86, 143.76, 141.27, 141.16, 141.13, 141.03, 130.67, 130.65, 130.04, 129.98, 127.43, 127.40, 126.89, 126.87, 126.82, 126.79, 125.28, 125.12, 124.91, 124.88, 119.58, 119.50, 114.92, 114.84, 93.65, 75.63, 74.94,

67.76, 67.23, 59.80, 52.08, 51.62, 36.33, 35.30. HRMS (ESI$^+$) m/z calculated for $C_{26}H_{22}INO_5$ $[M+H]^+$ 556.0621; found 556.0634.

(2*S*,4*R*)-1-(((9H-fluoren-9-yl)methoxy)carbonyl)-4-(3-iodophenoxy)pyrrolidine-2-carboxylic acid (**10bii**): Synthesized following general procedure A, Part 2. Yield: 38%.

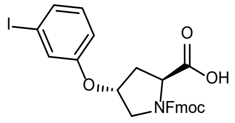

$^1$H NMR (400 MHz, Methanol-*d*$_4$) (43:57 mixture of rotamers) δ 7.67 – 7.53 (m, 2.86H), 7.42 (dd, *J* = 7.5, 1.0 Hz, 0.57H), 7.37 (dd, *J* = 7.5, 1.0 Hz, 0.57H), 7.29 – 7.13 (m, 4.86H), 7.11 (td, *J* = 7.5, 1.1 Hz, 0.57H), 7.05 (td, *J* = 7.5, 1.1 Hz, 0.57H), 6.97 – 6.89 (m, 1H), 6.85 – 6.80 (m, 0.43H), 6.78 (ddd, *J* = 8.4, 2.5, 0.9 Hz, 0.57H), 4.92 – 4.86 (m, 0.43H), 4.84 – 4.79 (m, 0.57H), 4.42 (t, *J* = 8.0 Hz, 0.43H), 4.34 (dd, *J* = 10.4, 6.6 Hz, 0.57H), 4.23 – 4.03 (m, 3H), 3.75 (dd, *J* = 12.0, 4.0 Hz, 0.43H), 3.70 (dt, *J* = 12.0, 1.9 Hz, 0.43H), 3.61 (dd, *J* = 12.0, 3.9 Hz, 0.57H), 3.52 (dt, *J* = 12.0, 1.6 Hz, 0.57H), 2.56 – 2.48 (m, 0.43H), 2.38 (ddt, *J* = 13.8, 7.8, 2.0 Hz, 0.57H), 2.22 (ddd, *J* = 13.3, 8.0, 4.8 Hz, 0.43H), 2.07 (ddd, *J* = 13.6, 8.8, 4.7 Hz, 0.57H). $^{13}$C NMR (101 MHz, $CD_3OD$) δ 178.08, 178.07, 157.82, 157.73, 155.73, 155.26, 144.26, 144.01, 143.75, 143.69, 141.18, 141.17, 141.10, 141.00, 130.88, 130.80, 130.17, 130.15, 127.39, 127.36, 126.92, 126.89, 126.74, 126.69, 125.35, 125.17, 124.70, 124.67, 124.61, 124.50, 119.57, 119.50, 119.46, 114.76, 114.72, 93.87, 93.73, 75.76, 75.23, 68.01, 67.03, 60.41, 60.16, 52.25, 51.65, 46.97, 36.82, 35.87. HRMS (ESI$^+$) m/z calculated for $C_{26}H_{22}INO_5$ $[M+H]^+$ 556.0621; found 556.0643.

(2*R*,4*R*)-1-(((9H-fluoren-9-yl)methoxy)carbonyl)-4-(3-iodophenoxy)pyrrolidine-2-carboxylic acid (**10cii**): Synthesized following general procedure A, Part 2. Yield: 29%.

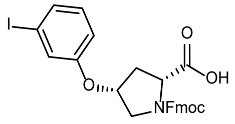

$^1$H NMR (400 MHz, $CDCl_3$) (1:1 mixture of rotamers) δ 7.80 – 7.72 (m, 2H), 7.63 – 7.55 (m, 2H), 7.47 – 7.25 (m, 5H), 7.16 (app. dt, *J* = 19.1, 2.0 Hz, 1H), 6.99 (app. dt, *J* = 21.4, 8.2 Hz, 1H), 6.77 (app. ddd, *J* = 20.6, 8.3, 2.5 Hz, 1H), 4.98 – 4.87 (m, 1H), 4.69 – 4.51 (m, 2H), 4.51 – 4.34 (m, 2H), 4.27 (app. dt, *J* =

25.2, 6.5 Hz, 1H), 3.77 (app. dd, *J* = 22.4, 3.0 Hz, 2H), 2.63 (app. dd, *J* = 25.4, 14.0 Hz, 1H), 2.54 – 2.41 (m, 1H). $^{13}$C NMR (101 MHz, $CDCl_3$) δ 177.37, 176.45, 156.86, 156.81, 155.18, 154.54, 143.93, 143.86, 143.76, 143.66, 141.40, 141.29, 131.02, 130.96, 130.88, 130.80, 127.79, 127.71, 127.12, 125.11, 124.96, 124.87, 119.99, 115.26, 115.05, 94.48, 77.26, 75.52, 74.60, 67.85, 67.67, 57.88, 57.41, 52.11, 51.82, 47.19, 36.42, 35.10. HRMS (ESI$^+$) m/z calculated for $C_{26}H_{22}INO_5$ $[M+H]^+$ 556.0621; found 556.0649.

(2*R*,4*S*)-1-(((9H-fluoren-9-yl)methoxy)carbonyl)-4-(3-iodophenoxy)pyrrolidine-2-carboxylic acid (**10dii**): Synthesized following general procedure A, Part 2. Yield: 60%.

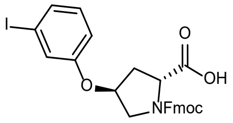

$^1$H NMR (400 MHz, Methanol-*d*$_4$) (55:45 mixture of rotamers) δ 7.68 – 7.54 (m, 3H), 7.44 (d, *J* = 7.5 Hz, 1H), 7.38 (d, *J* = 7.5 Hz, 1H), 7.30 – 7.09 (m, 6H), 7.06 (t, *J* = 7.5 Hz, 1H), 6.99 – 6.90 (m, 1H), 6.87 – 6.82 (m, 0H), 6.79 (dd, *J* = 8.5, 2.5 Hz, 1H), 4.94 – 4.88 (m, 0H), 4.85 – 4.81 (m, 1H), 4.45 – 4.33 (m, 1H), 4.24 – 4.05 (m, 3H), 3.75 (dd, *J* = 11.9, 4.0 Hz, 0H), 3.70 (dt, *J* = 12.2, 1.9 Hz, 0H), 3.61 (dd, *J* = 12.1, 3.9 Hz, 1H), 3.52 (dt, *J* = 12.0, 1.4 Hz, 1H), 2.56 – 2.49 (m, 0H), 2.43 – 2.35 (m, 1H), 2.22 (ddd, *J* = 13.3, 7.9, 4.8 Hz, 0H), 2.08 (ddd, *J* = 13.6, 8.7, 4.7 Hz, 1H). $^{13}$C NMR (101 MHz, $CD_3OD$) δ 178.01, 177.92, 157.83, 157.75, 155.73, 155.25, 144.27, 144.03, 143.75, 143.69, 141.19, 141.17, 141.10, 141.00, 130.87, 130.79, 130.17, 130.14, 127.38, 127.34, 126.91, 126.88, 126.73, 126.68, 125.34, 125.16, 124.71, 124.65, 124.60, 124.49, 119.54, 119.47, 119.43, 114.77, 114.73, 93.81, 93.68, 75.75, 75.23, 68.01, 67.02, 60.31, 60.12, 52.24, 51.62, 46.96, 36.80, 35.88. HRMS (ESI$^+$) m/z calculated for $C_{26}H_{22}INO_5$ $[M+H]^+$ 556.0621; found 556.0643.

(2*S*,4*S*)-1-(((9H-fluoren-9-yl)methoxy)carbonyl)-4-((4-iodobenzyl)oxy)pyrrolidine-2-carboxylic acid (**14ai**): Synthesized following general procedure B, Part 3. Yield: 62%.

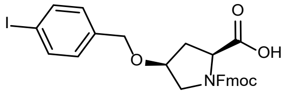

$^1$H NMR (400 MHz, $CDCl_3$) (57:43 mixture of rotamers) δ 7.73 – 7.57 (m, 2H), 7.56 – 7.37 (m, 4H), 7.35 – 7.17 (m, 4H), 6.85 (dd, *J* = 31.3, 7.9 Hz, 2H), 4.50 – 3.99 (m, 7H), 3.67 – 3.42 (m, 2H), 2.54 (d, *J* = 13.7

Hz, 0.57H), 2.38 – 2.05 (m, 1.43H). $^{13}$C NMR (101 MHz, $CDCl_3$) δ 176.44, 174.85, 156.03, 154.66, 143.75, 143.61, 141.34, 141.24, 137.46, 137.16, 129.49, 129.29, 127.80, 127.14, 125.12, 124.98, 124.87, 120.01, 93.21, 93.09, 77.26, 75.99, 68.01, 67.68, 58.18, 57.46, 52.48, 52.10, 47.18, 36.24, 33.81. HRMS (ESI$^+$) m/z calculated for $C_{27}H_{24}INO_5$ $[M+H]^+$ 570.0777; found 570.0792.

(2*S*,4*R*)-1-(((9H-fluoren-9-yl)methoxy)carbonyl)-4-((4-iodobenzyl)oxy)pyrrolidine-2-carboxylic acid (**14bi**): Synthesized following general procedure B, Part 3. Yield: 79%.

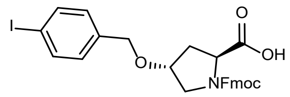

$^1$H NMR (400 MHz, Methanol-$d_4$) (55:45 mixture of rotamers) δ 7.65 – 7.57 (m, 2H), 7.52 – 7.37 (m, 4H), 7.25 – 7.17 (m, 2H), 7.12 (qd, *J* = 7.9, 3.0 Hz, 2H), 6.90 (dd, *J* = 8.3, 4.4 Hz, 2H), 4.34 (t, *J* = 7.8 Hz, 0.45H), 4.29 – 4.09 (m, 4.55H), 4.07 – 3.97 (m, 2H), 3.61 (d, *J* = 11.7 Hz, 0.45H), 3.47 – 3.40 (m, 1H), 3.37 (dd, *J* = 11.7, 4.3 Hz, 0.55H), 2.41 – 2.24 (m, 1H), 2.01 (ddd, *J* = 13.1, 7.6, 5.0 Hz, 0.45H), 1.92 (ddd, *J* = 13.9, 8.3, 5.3 Hz, 0.55H). $^{13}$C NMR (101 MHz, $CD_3OD$) δ 174.63, 174.39, 155.30, 155.22, 143.88, 143.85, 143.68, 143.63, 141.20, 141.14, 141.07, 137.99, 137.90, 137.26, 129.32, 129.22, 127.49, 126.86, 124.89, 124.85, 124.75, 119.65, 119.62, 92.45, 77.10, 76.28, 69.89, 69.79, 67.91, 67.30, 57.97, 57.73, 51.94, 51.51, 46.97, 46.91, 36.41, 35.38. HRMS (ESI$^+$) m/z calculated for $C_{27}H_{24}INO_5$ $[M+H]^+$ 570.0777; found 570.0794.

(2*R*,4*R*)-1-(((9H-fluoren-9-yl)methoxy)carbonyl)-4-((4-iodobenzyl)oxy)pyrrolidine-2-carboxylic acid (**14ci**): Synthesized following general procedure B, Part 3. Yield: 75%.

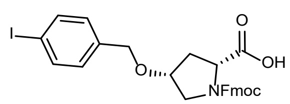

$^1$H NMR (400 MHz, Methanol-$d_4$) δ 7.58 (d, *J* = 7.5 Hz, 2H), 7.50 – 7.37 (m, 4H), 7.20 (t, *J* = 7.5 Hz, 2H), 7.17 – 7.07 (m, 2H), 6.87 (t, *J* = 7.3 Hz, 2H), 4.29 – 4.06 (m, 5H), 4.01 (dt, *J* = 13.9, 6.8 Hz, 1H), 3.94 – 3.87 (m, 1H), 3.47 – 3.31 (m, 2H), 2.28 – 2.18 (m, 1H), 2.15 – 2.02 (m, 1H). $^{13}$C NMR (101 MHz, $CD_3OD$) δ 175.80, 175.55, 157.12, 157.05, 145.85, 145.81, 145.61, 145.53, 143.07, 143.02, 142.95, 139.88, 138.98, 138.95, 131.04, 131.02, 129.37, 128.75, 126.78, 126.72, 126.68, 121.52, 94.10, 94.05, 79.11,

78.23, 71.38, 69.50, 69.21, 59.65, 59.54, 53.99, 53.71, 48.90, 48.83, 37.38, 36.39. HRMS (ESI$^+$) m/z calculated for $C_{27}H_{24}INO_5$ [M+H]$^+$ 570.0777; found 570.0791.

(2*R*,4*S*)-1-(((9H-fluoren-9-yl)methoxy)carbonyl)-4-((4-iodobenzyl)oxy)pyrrolidine-2-carboxylic acid (**14di**): Synthesized following general procedure B, Part 3. Yield: 72%.

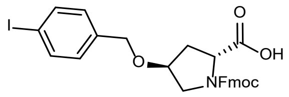

$^1$H NMR (400 MHz, Methanol-*d*$_4$) (55:45 mixture of rotamers) δ 7.59 (dd, *J* = 8.1, 3.4 Hz, 2H), 7.52 – 7.35 (m, 4H), 7.25 – 7.16 (m, 2H), 7.16 – 7.05 (m, 2H), 6.88 (dd, *J* = 8.2, 3.7 Hz, 2H), 4.34 (t, *J* = 7.8 Hz, 0.45H), 4.28 – 4.07 (m, 4.55H), 4.05 – 3.95 (m, 2H), 3.60 (d, *J* = 11.7 Hz, 0.45H), 3.48 – 3.40 (m, 1H), 3.36 (dd, *J* = 11.7, 4.3 Hz, 0.55H), 2.40 – 2.22 (m, 1H), 2.00 (ddd, *J* = 13.0, 7.6, 4.9 Hz, 0.45H), 1.91 (ddd, *J* = 13.5, 8.1, 5.0 Hz, 0.55H). $^{13}$C NMR (101 MHz, $CD_3OD$) δ 174.63, 174.39, 155.29, 155.21, 143.88, 143.84, 143.67, 143.63, 141.19, 141.18, 141.13, 141.07, 137.96, 137.87, 137.26, 129.32, 129.22, 127.51, 126.88, 124.91, 124.87, 124.77, 119.67, 119.64, 92.50, 77.10, 76.27, 69.89, 69.80, 67.91, 67.31, 57.98, 57.75, 51.95, 51.53, 46.97, 46.91, 36.41, 35.37. HRMS (ESI$^+$) m/z calculated for $C_{27}H_{24}INO_5$ [M+H]$^+$ 570.0777; found 570.0788.

(2*S*,4*S*)-1-(((9H-fluoren-9-yl)methoxy)carbonyl)-4-((3-iodobenzyl)oxy)pyrrolidine-2-carboxylic acid (**14aii**): Synthesized following general procedure B, Part 3. Yield: 49%.

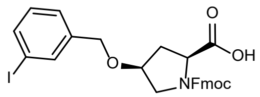

$^1$H NMR (400 MHz, $CDCl_3$) (57:43 mixture of rotamers) δ 10.11 (br. s, 1H), 7.62 (dd, *J* = 25.6, 7.6 Hz, 2H), 7.54 – 7.36 (m, 4H), 7.33 – 7.03 (m, 5H), 6.91 (app. dt, *J* = 13.9, 7.7 Hz, 1H), 4.47 – 4.12 (m, 5.57H), 4.08 – 3.92 (m, 1.43H), 3.62 – 3.45 (m, 2H), 2.44 (d, *J* = 13.7 Hz, 0.57H), 2.30 (d, *J* = 13.7 Hz, 0.43H), 2.24 – 2.05 (m, 1H). $^{13}$C NMR (101 MHz, $CDCl_3$) δ 176.65, 175.40, 155.75, 154.73, 143.75, 143.66, 141.33, 141.22, 139.97, 136.77, 136.37, 136.23, 130.28, 127.81, 127.72, 127.18, 126.79, 126.66, 125.18,

125.04, 124.96, 120.03, 94.34, 76.74, 75.88, 69.68, 67.96, 67.79, 58.01, 57.44, 52.37, 52.19, 47.19, 36.04, 34.01. HRMS (ESI$^+$) m/z calculated for $C_{27}H_{24}INO_5$ $[M+H]^+$ 570.0777; found 570.0790.

(2*S*,4*R*)-1-(((9H-fluoren-9-yl)methoxy)carbonyl)-4-((3-iodobenzyl)oxy)pyrrolidine-2-carboxylic acid (**14bii**): Synthesized following general procedure B, Part 3. Yield: 80%.

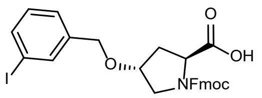

$^1$H NMR (400 MHz, Methanol-$d_4$) (54:46 mixture of rotamers) δ 7.63 – 7.55 (m, 2H), 7.51 (d, *J* = 5.5 Hz, 1H), 7.48 – 7.35 (m, 3H), 7.24 – 7.16 (m, 2H), 7.16 – 7.06 (m, 3H), 6.90 (q, *J* = 7.6 Hz, 1H), 4.34 (t, *J* = 7.9 Hz, 0.46H), 4.30 – 4.11 (m, 4.54H), 4.06 – 3.95 (m, 2H), 3.62 (dt, *J* = 12.0, 2.1 Hz, 0.46H), 3.43 (dd, *J* = 12.1, 3.4 Hz, 1H), 3.35 (dd, *J* = 11.7, 4.3 Hz, 0.54H), 2.40 – 2.23 (m, 1H), 2.00 (ddd, *J* = 13.0, 7.5, 4.9 Hz, 0.46H), 1.91 (ddd, *J* = 13.2, 7.9, 4.8 Hz, 0.54H). $^{13}$C NMR (101 MHz, $CD_3OD$) δ 174.64, 174.39, 155.29, 155.23, 143.87, 143.84, 143.69, 143.65, 141.19, 141.13, 141.07, 140.72, 140.66, 136.47, 136.19, 136.12, 129.98, 129.96, 127.50, 126.89, 126.53, 126.47, 124.91, 124.89, 124.76, 124.73, 119.65, 119.62, 93.75, 93.71, 77.17, 76.40, 69.61, 69.51, 67.93, 67.30, 57.96, 57.74, 51.92, 51.49, 46.98, 46.93, 36.41, 35.36. HRMS (ESI$^+$) m/z calculated for $C_{27}H_{24}INO_5$ $[M+H]^+$ 570.0777; found 570.0810.

(2*R*,4*R*)-1-(((9H-fluoren-9-yl)methoxy)carbonyl)-4-((3-iodobenzyl)oxy)pyrrolidine-2-carboxylic acid (**14cii**): Synthesized following general procedure B, Part 3. Yield: 86%.

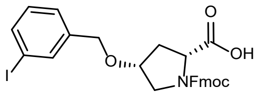

$^1$H NMR (400 MHz, Methanol-$d_4$) (1:1 mixture of rotamers) δ 7.82 – 7.75 (m, 2H), 7.74 – 7.57 (m, 4H), 7.38 (td, *J* = 7.3, 2.4 Hz, 2H), 7.35 – 7.27 (m, 3H), 7.09 (td, *J* = 7.7, 5.9 Hz, 1H), 4.50 – 4.31 (m, 5H), 4.27 (t, *J* = 6.6 Hz, 0.5H), 4.23 – 4.15 (m, 1.5H), 3.64 (dd, *J* = 11.8, 4.9 Hz, 0.5H), 3.61 – 3.54 (m, 1.5H), 2.47 – 2.28 (m, 2H). $^{13}$C NMR (101 MHz, $CD_3OD$) δ 174.15, 173.87, 155.39, 155.31, 144.05, 144.00, 143.75, 143.72, 141.27, 141.21, 141.19, 141.10, 140.81, 136.31, 136.27, 136.05, 136.03, 129.84, 129.82, 127.43, 126.83, 126.46, 124.85, 124.79, 124.77, 119.52, 119.48, 93.37, 93.32, 77.31, 76.47, 69.38, 69.33, 67.63,

67.38, 57.82, 57.74, 52.08, 51.78, 47.08, 47.02, 35.47, 34.52. HRMS (ESI$^+$) m/z calculated for $C_{27}H_{24}INO_5$ [M+H]$^+$ 570.0777; found 570.0793.

(2*R*,4*S*)-1-(((9H-fluoren-9-yl)methoxy)carbonyl)-4-((3-iodobenzyl)oxy)pyrrolidine-2-carboxylic acid (**14dii**): Synthesized following general procedure B, Part 3. Yield: 90%.

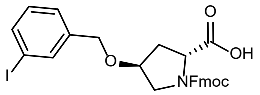

$^1$H NMR (400 MHz, Methanol-$d_4$) (55:45 mixture of rotamers) δ 7.62 – 7.55 (m, 2H), 7.51 (d, *J* = 5.3 Hz, 1H), 7.47 – 7.36 (m, 3H), 7.24 – 7.15 (m, 2H), 7.15 – 7.06 (m, 3H), 6.90 (q, *J* = 7.7 Hz, 1H), 4.34 (t, *J* = 7.9 Hz, 0.45H), 4.29 – 4.19 (m, 2.55H), 4.19 – 4.09 (m, 2H), 4.05 – 3.95 (m, 2H), 3.65 – 3.57 (m, 0.45H), 3.47 – 3.39 (m, 1H), 3.34 (dd, *J* = 11.7, 4.3 Hz, 0.55H), 2.39 – 2.23 (m, 1H), 2.00 (ddd, *J* = 13.1, 7.6, 4.9 Hz, 0.45H), 1.91 (ddd, *J* = 13.2, 7.9, 4.8 Hz, 0.55H). $^{13}$C NMR (101 MHz, $CD_3OD$) δ 174.64, 174.39, 155.29, 155.23, 143.86, 143.84, 143.69, 143.65, 141.18, 141.12, 141.07, 140.71, 140.65, 136.47, 136.19, 136.11, 129.99, 129.97, 127.50, 126.89, 126.54, 126.47, 124.91, 124.90, 124.76, 124.74, 119.66, 119.63, 93.77, 93.73, 77.17, 76.40, 69.61, 69.52, 67.93, 67.31, 57.96, 57.74, 51.92, 51.49, 46.98, 46.93, 36.41, 35.35. HRMS (ESI$^+$) m/z calculated for $C_{27}H_{24}INO_5$ [M+H]$^+$ 570.0777; found 570.0800.

#### 1.3. Spirocyclic azetidines

(*R*)-4-benzyl-3-((3*S*,4*R*)-1-benzyl-4-(4-iodophenyl)pyrrolidine-3-carbonyl)oxazolidin-2-one **(15)**

(*R*)-4-benzyl-3-((3*R*,4*S*)-1-benzyl-4-(4-iodophenyl)pyrrolidine-3-carbonyl)oxazolidin-2-one **(16)**

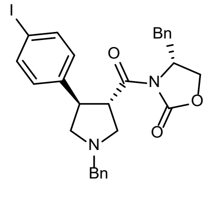

**15**

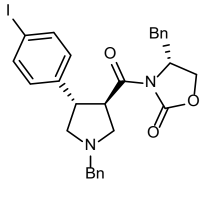

**16**

To a 250 mL round-bottom flask was added 4-iodobenzaldehyde (11.6 g, 50 mmol, 1 eq) and malonic acid (6.25 g, 60 mmol, 1.2 eq), the mixture was dissolved with pyridine (50 mL), then 0.1 mL of piperidine was added. The resulting mixture was stirred under reflux for 6 h, cooled to rt and poured into 500 mL ice-water, then adjusted the pH to ~3 by adding concentrated HCl. The precipitated white solid was collected by filtration and washed with water for 3 times. (*E*)-3-(4-iodophenyl)acrylic acid (13.8 g, quantitative yield), a white solid, was dried under air and used for next step directly.

To a 250 mL round-bottom flask was added *E*)-3-(4-iodophenyl)acrylic acid (13.7 g, 50 mmol) and dissolved with $CH_2Cl_2$ (120 mL), 5 drops of DMF was added. To this solution was added oxalyl chloride (6.0 mL, 70 mmol, 1.4 eq) slowly while stirring, after which the reaction mixture was stirred for 2 hr at rt. The volatiles were evaporated under reduced pressure and dried under vacuum to afford (*E*)-3-(4-iodophenyl)acryloyl chloride (14.7 g, quantitative yield, stored under $N_2$ before use) as a white solid.

To a 250 mL round-bottom flask was added NaH (2.4 g, 60 mmol, 60% wt in mineral oil) and MeCN (120 mL). The mixture was stirred under an ice-bath and (*R*)-4-benzyloxazolidin-2-one (8.9 g, 50 mmol) was added in portions. The reaction mixture was stirred at rt for 2 h, then cooled with an ice-bath again. (*E*)-3-(4-iodophenyl)acryloyl chloride (14.6 g, 50 mmol) was added in portions, then the mixture was stirred at rt overnight. The reaction was quenched with 5 mL of water and the organic solvent was evaporated under reduced pressure. To the resulting solid was added 200 mL of water and stirred for 30 min, the solid was collected by filtration and washed with water twice. (*R*,*E*)-4-benzyl-3-(3-(4-iodophenyl)acryloyl)oxazolidin-2-one (19.0 g, 88% yield) was isolated as an off-white solid and used directly in the next step.

To a 250 mL round-bottom flask was added (*R*,*E*)-4-benzyl-3-(3-(4-iodophenyl)acryloyl)oxazolidin-2-one (17.4 g, 40 mmol, 1 eq) and *N*-benzyl-1-methoxy-*N*-((trimethylsilyl)methyl)methanamine (10.5 g, 44 mmol, 1.1 eq), the mixture was dissolved with anhydrous $CH_2Cl_2$ (150 mL) and cooled to 0 °C. To the cooled solution was added 5 drops of trifluoroacetic acid and the reaction mixture was stirred for 30 min 0 °C. The reaction was warmed to rt and stirred for another 4 h. The reaction mixture was washed with 1 M aq. $NaHCO_3$ solution, the organic phase was collected and dried with anhydrous $Na_2SO_4$. The solvent evaporated and the residue was purified by flash column chromatography (hexanes/ethyl acetate: 3:1 to 2:1). The two diastereomers were obtained in 42% yield **15** (Rf: ~0.4, hexanes/ethyl acetate = 2:1) and 43% yield for **16** (Rf: ~0.25, hexanes/ethyl acetate = 2:1) respectively.

(*R*)-4-benzyl-3-((*3S,4R*)-1-benzyl-4-(4-iodophenyl)pyrrolidine-3-carbonyl)oxazolidin-2-one (**15**)

Pale yellow solid, upper spot on TLC, 42% yield. $[\alpha]^{20}_D$ = -68.00 (c=1, $CHCl_3$); $^1$H NMR (600 MHz, $CDCl_3$) δ 7.63 (d, *J* = 8.4 Hz, 2H), 7.41 (d, *J* = 7.4 Hz, 2H), 7.36 (t, *J* = 7.5 Hz, 2H), 7.32 – 7.28 (m, 4H), 7.22 – 7.18 (m, 2H), 7.11 (d, *J* = 8.4 Hz, 2H), 4.71 – 4.63 (m, 1H), 4.17 (q, *J* = 5.2, 4.6 Hz, 2H), 4.09 (h, *J* = 7.4 Hz, 2H), 3.81 (d, *J* = 13.0 Hz, 1H), 3.70 (d, *J* = 13.5 Hz, 1H), 3.33 (t, *J* = 9.0 Hz, 1H), 3.28 – 3.24 (m, 1H), 3.23 (m, 1H), 2.92-2.86 (m, 1H), 2.81 (dd, *J* = 13.4, 9.3 Hz, 1H), 2.70 (t, *J* = 8.0 Hz, 1H). $^{13}$C NMR (151 MHz, $CDCl_3$) δ 173.10, 153.22, 137.80, 135.14, 130.09, 129.62, 129.14, 128.94, 128.59, 127.61, 127.44, 92.13, 66.47, 61.69, 59.90, 58.24, 55.41, 51.47, 44.91, 37.99; HRMS (ESI-TOF) m/z calculated for $C_{28}H_{28}IN_2O_3$ $[M+H]^+$ 567.1139; found: 567.1142.

(*R*)-4-benzyl-3-((*3R,4S*)-1-benzyl-4-(4-iodophenyl)pyrrolidine-3carbonyl)oxazolidin-2-one (**16**)

Pale yellow solid, lower spot on TLC, 43% yield. $[\alpha]^{20}_D$ = -4.10 (c=1, $CHCl_3$); $^1$H NMR (600 MHz, $CDCl_3$) δ 7.67 (d, *J* = 8.4 Hz, 2H), 7.39 (d, *J* = 7.2 Hz, 2H), 7.34 (t, *J* = 7.5 Hz, 2H), 7.30 – 7.23 (m, 4H), 7.15 (d, *J* = 8.4 Hz, 2H), 7.08 (dd, *J* = 7.6, 1.7 Hz, 2H), 4.73 – 4.66 (m, 1H), 4.26 (ddd, *J* = 9.4, 7.5, 6.1 Hz, 1H), 4.22 – 4.18 (m, 1H), 4.15 (dd, *J* = 9.0, 2.9 Hz, 1H), 4.05 (q, *J* = 7.7 Hz, 1H), 3.81 – 3.66 (m, 2H), 3.27 (t, *J* = 9.6 Hz, 1H), 3.22 – 3.16 (m, 2H), 2.84 (dd, *J* = 9.7, 6.1 Hz, 1H), 2.78 (dd, *J* = 13.5, 9.1 Hz, 1H), 2.74 – 2.69 (m, 1H); $^{13}$C NMR (151 MHz, $CDCl_3$) δ 173.49, 153.16, 137.79, 135.03, 130.04, 129.54, 129.09, 128.90, 128.52, 127.56, 127.34, 92.13, 66.24, 61.57, 59.95, 58.06, 55.55, 51.08, 45.70, 37.75; HRMS (ESI-TOF) m/z calculated for $C_{28}H_{28}IN_2O_3$ $[M+H]^+$ 567.1139; found 567.1144.

1-(*tert*-butyl) 3-methyl (*3R,4S*)-4-(4-iodophenyl)pyrrolidine-1,3-dicarboxylate (**17**)

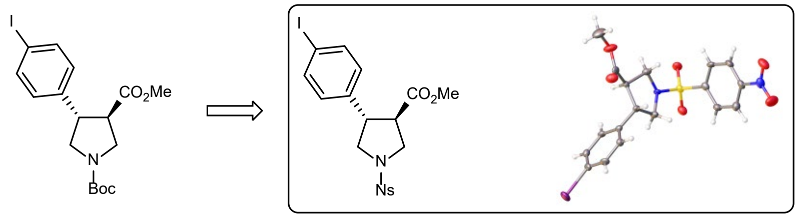

To a 250-mL round-bottom flask was added **15** (8.5 g, 15 mmol, 1 eq), dissolved with toluene (120 mL), then 1-chloroethyl chloroformate (4.3 g, 30 mmol) was added. The resulting solution was stirred under reflux, a lot of solid was precipitated during the reaction, the reflux was continued until the solid disappeared again (2-4 hr). The resulting solution was cooled to rt, and the solvent was evaporated under reduced pressure, the residue was dissolved with 50 mL of MeOH and heated at reflux for 1 h. The solvent was evaporated, and the residue was dissolved with saturated aq. $NaHCO_3$ solution (100 mL) and THF (50 mL), then $Boc_2O$ (4.4 g, 20 mmol, 1.3 eq) was added, the resulting solution was stirred at rt overnight. The reaction solution was extracted with ethyl acetate twice and the combined organic portion was dried with anhydrous $Na_2SO_4$. After filtration the solution was concentrated, and the residue was purified by flash column chromatography (hexanes/ethyl acetate: 3:1 to 2:1) to obtain *tert*-butyl (3*R*,4*S*)-3-((*R*)-4-benzyl-2-oxooxazolidine-3-carbonyl)-4-(4-iodophenyl)pyrrolidine-1-carboxylate (7.9 g, 13.7 mmol, 91% yield).

To a 250-mL round-bottom flask was added *tert*-butyl (3*R*,4*S*)-3-((*R*)-4-benzyl-2-oxooxazolidine-3-carbonyl)-4-(4-iodophenyl)pyrrolidine-1-carboxylate (7.5 g, 13 mmol, 1 eq), dissolved in THF (50 mL), then aq. LiOH (30 mL, 2.0 M) was added, followed by the addition of aq. $H_2O_2$ (4 mL, 30%v/v). The resulting solution was stirred at rt overnight. The mixture was acidified with aq. HCl (50 mL, 2.0 M) and extracted with EtOAc (5 times). The organic portions were combined and evaporated under reduced pressure, the residue was dissolved with DMF (50 mL), $K_2CO_3$ (3.6 g, 26 mmol, 2 eq) and MeI (1.7 mL, 26 mmol, 2 eq) were added sequentially, the resulting mixture was stirred at rt for 8 h. The reaction solution was diluted with 100 mL of water and extracted with EtOAc 3 times. The organic portions were combined and washed with water twice and then dried with anhydrous $Na_2SO_4$. The solvent was concentrated, and the residue was purified by flash column chromatography (hexanes/ethyl acetate: 3:1) to provide **17** (3.75 g, 8.7 mmol, 67% yield) as a white solid.

1-(*tert*-butyl) 3-methyl (*3R,4S*)-4-(4-iodophenyl)pyrrolidine-1,3-dicarboxylate (**17**)

99% ee, absolute configuration confirmed by x-ray of its Ns analogue, $[\alpha]^{20}D$ = 42.10 (c=1, $CHCl_3$). $^1H$ NMR (600 MHz, $CDCl_3$) δ 7.66 (d, *J* = 8.3 Hz, 2H), 7.02 (d, *J* = 8.4 Hz, 2H), 3.95 – 3.78 (m, 2H), 3.65 (s, 3H), 3.63 – 3.54 (m, 2H), 3.38 (q, *J* = 8.6 Hz, 1H), 3.15 (q, *J* = 9.1 Hz, 1H), 1.48 (s, 9H). $^{13}C$ NMR (151 MHz, $CDCl_3$) δ 172.93, 154.59, 139.52, 138.47, 129.94, 93.32, 80.49, 52.80, 50.90, 49.98, 49.33, 47.37, 29.09. HRMS (ESI-TOF) m/z calculated for $C_{17}H_{23}INO_4$ $[M+H]^+$ 432.0666; found 432.0669.

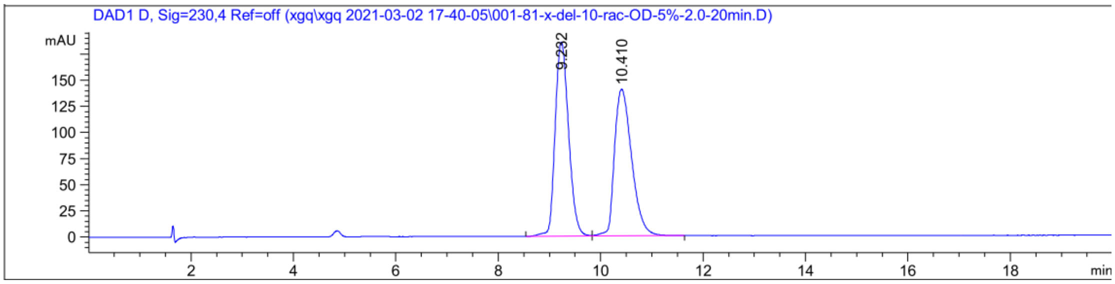

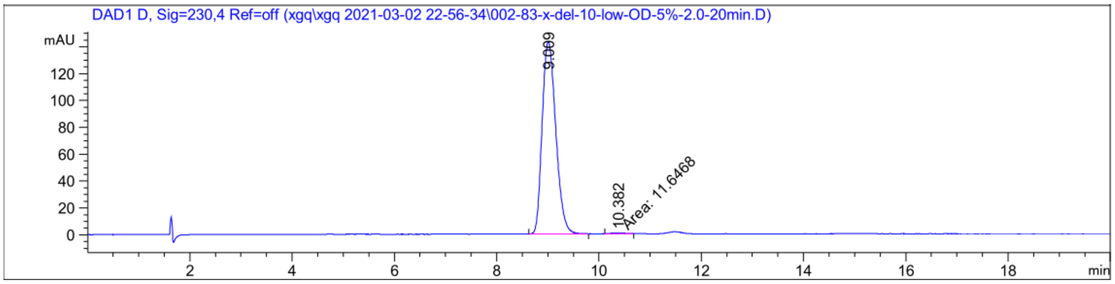

Signal 3: DAD1 D, Sig=230,4 Ref=off

| Peak # | RetTime [min] | Type | Width [min] | Area [mAU*s] | Height [mAU] | Area % |
|---|---|---|---|---|---|---|
| 1 | 9.009 | BB | 0.2908 | 2634.95508 | 143.08714 | 99.5599 |
| 2 | 10.382 | MM | 0.3021 | 11.64684 | 6.42448e-1 | 0.4401 |
| Totals : | | | | 2646.60192 | 143.72959 | |

1-(*tert*-butyl) 3-methyl (3*S*,4*R*)-4-(4-iodophenyl)pyrrolidine-1,3-dicarboxylate (**18**) [prepared as for **17**]

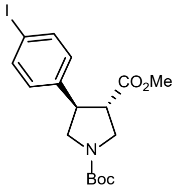

White solid, 96.5% ee. $[\alpha]^{20}_{D}$ = -41.10 (c=1, $CHCl_3$). $^1$H NMR (600 MHz, $CDCl_3$) δ 7.66 (d, *J* = 8.3 Hz, 2H), 7.02 (d, *J* = 8.4 Hz, 2H), 3.94 – 3.81 (m, 2H), 3.65 (s, 3H), 3.63 – 3.56 (m, 2H), 3.38 (q, *J* = 8.6 Hz, 1H), 3.15 (q, *J* = 9.1 Hz, 1H), 1.48 (s, 9H). $^{13}$C NMR (151 MHz, $CDCl_3$) δ 172.49, 154.15, 139.08, 138.03,

129.50, 92.88, 80.05, 52.36, 50.46, 49.54, 48.89, 46.93, 28.65. HRMS (ESI-TOF) m/z calculated for $C_{17}H_{23}INO_4$ $[M+H]^+$ 432.0666; found 432.0667.

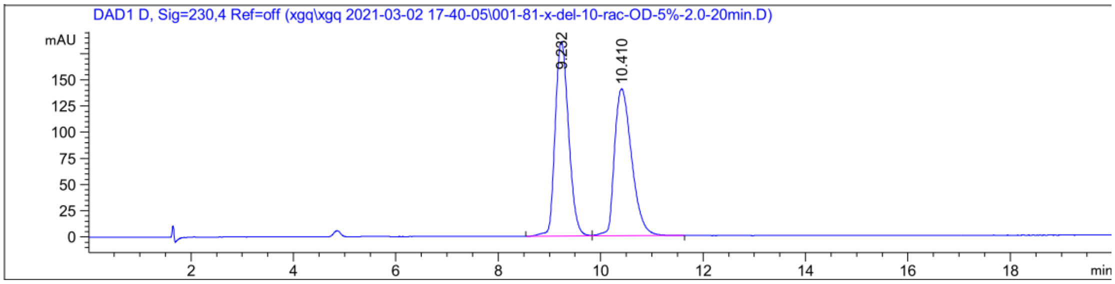

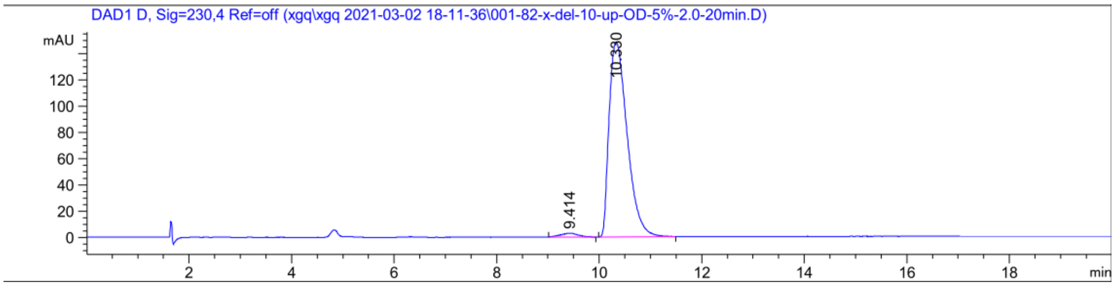

Signal 3: DAD1 D, Sig=230,4 Ref=off

| Peak # | RetTime [min] | Type | Width [min] | Area [mAU*s] | Height [mAU] | Area % |
|---|---|---|---|---|---|---|
| 1 | 9.414 | BB | 0.3026 | 65.23700 | 2.77138 | 1.7703 |
| 2 | 10.330 | BB | 0.3866 | 3619.87378 | 148.36899 | 98.2297 |
| Totals : | | | | 3685.11078 | 151.14037 | |

*tert*-butyl (*R*)-8-(4-iodophenyl)-2-((4-nitrophenyl)sulfonyl)-2,6-diazaspiro[3.4] octane-6-carboxylate (**19**)

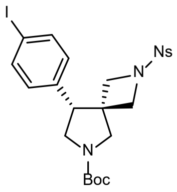

A flame-dried 100 mL round-bottom flask equipped with a magnetic stir bar was evacuated and backfilled with $N_2$ 3 times. To this flask was added diisopropylamine (0.45 mL, 3.2 mmol, 1.6 eq) and anhydrous THF (20 mL), cooled to -78 °C. $^n$BuLi solution (1.2 mL, 2.5 M in hexane, 3.0 mmol, 1.5 eq) was added to the reaction slowly via a syringe, after addition the reaction was stirred at -78 °C for 15 min. **17** (0.87 g, 2.0 mmol, 1 eq) was dissolved in anhydrous THF (5 mL) and added to the reaction slowly via a syringe, after addition the reaction was stirred -78 °C for 2 h. 2-(Bromomethyl)isoindoline-1,3-dione (0.96 g, 4 mmol, 2 eq) was dissolved in anhydrous THF (5 mL) and added to the reaction slowly via a syringe, after addition the reaction was warmed up to rt slowly and stirred overnight. Saturated aq. $NH_4Cl$ (50 mL) was added to the reaction and the reaction mixture was extracted with EtOAc (30 mL) 3 times. The combined organic portions were dried with anhydrous $Na_2SO_4$ and then filtrated, the solvent was removed under reduced pressure and the residue was purified by flash column chromatography (hexanes/ethyl acetate: 3:1) to give the alkylated product 1-(*tert*-butyl) 3-methyl (3*R*,4*R*)-3-((1,3-dioxoisoindolin-2-yl)methyl)-4-(4-iodophenyl)pyrrolidine-1,3-dicarboxylate (602 mg, 51% yield), which was used directly in subsequent steps.

To a 12 mL reaction vial was added 1-(*tert*-butyl) 3-methyl (3*R*,4*R*)-3-((1,3-dioxoisoindolin-2-yl)methyl)-4-(4-iodophenyl)pyrrolidine-1,3-dicarboxylate (590 mg, 1.0 mmol, 1 eq), dissolved with methanol (5 mL), then hydrazine monohydrate (0.1 mL, 2.0 mmol, 2 eq) was added, the vial was capped and stirred at 50 °C for 12 h. The reaction was cooled to rt and diluted with $CH_2Cl_2$ (15 mL), the white solid precipitate was filtered off and washed with $CH_2Cl_2$ (10 mL). The combined organic solution was concentrated and re-dissolved in anhydrous THF (5 mL), then lithium borohydride (1.5 mL, 2.0 M in THF, 3 eq) was added, and the resulting solution was stirred at 50 °C for 12 h. After the reaction was cooled to rt the solution was poured into 30 mL of ice-water, the aqueous solution was then extracted with EtOAc (20 mL) 5 times. The organic portions were combined and concentrated to provide *tert*-butyl (3*R*,4*R*)-3-(aminomethyl)-3-(hydroxymethyl)-4-(4-iodophenyl)pyrrolidine-1-carboxylate as a white foam, which was used directly in the next step.

To a 12 mL vial was added *tert*-butyl (3*R*,4*R*)-3-(aminomethyl)-3-(hydroxymethyl)-4-(4-iodophenyl)pyrrolidine-1-carboxylate (~1.0 mmol, 1 eq) which was prepared in the last step and 2-nitrobenzenesufonyl chloride (270 mg, 1.2 mmol, 1.2 eq), dissolved in $CH_2Cl_2$ (6 mL), then 2,6-lutidine (0.5 mL, 4.3 mmol, 4.3 eq) was added. The resulting solution was stirred at rt for 12 h, after which the reaction mixture was washed with 0.5 M aq. HCl (20 mL), and the aqueous phase was extracted with

$CH_2Cl_2$ (10 mL) 3 times. The combined organic portions were dried with anhydrous $Na_2SO_4$ and concentrated. The residue was re-dissolved with $CH_2Cl_2$ (10 mL) and $NEt_3$ (0.3 mL, 2.2 mmol, 2.2 eq) was added. The resulting solution was cooled to 0 °C and methanesulfonyl chloride (0.15 mL, 2.0 mmol, 2 eq) was added dropwise, then after addition the reaction was stirred at 0 °C for 30 min, then quenched with saturated aq. $NH_4Cl$ (10 mL). The aqueous phase was extracted with $CH_2Cl_2$ (10 mL) twice and dried with anhydrous $Na_2SO_4$. The solvent was concentrated, and the residue was re-dissolved MeCN (6 mL), then $K_2CO_3$ (280 mg, 2.0 mmol, 2 eq) was added. The resulting mixture was stirred at 80 °C for 12 h, cooled to rt and poured into water (20 mL), the aqueous solution was then extracted with EtOAc (20 mL) 3 times. The combined organic portions were dried with anhydrous $Na_2SO_4$, filtered, and concentrated under reduced pressure. The residue was purified by flash column chromatography (hexanes/ethyl acetate: 3:1 to 2:1) to give the spiro azetidine **19** (366 mg, 51% over 5 steps) as a pale brown foam.

Mixture of 2 rotamers, 97.5% ee. $[\alpha]^{20}_D$ = 1.30 (c=1, $CHCl_3$). $^1H$ NMR (600 MHz, $CDCl_3$) δ 7.86-7.77 (m, 1.6H), 7.70-7.69 (m, 0.23H), 7.66-7.64 (m, 1.77H), 7.57 (d, *J* = 8.4 Hz, 1.4H), 7.30 (t, *J* = 7.2 Hz, 0.72H), 7.24 (t, *J* = 7.2 Hz, 0.28H), 7.12 (d, *J* = 7.2 Hz, 0.58H), 6.86 (d, *J* = 7.2 Hz, 1.42H), 4.04-3.97 (m, 2H), 3.89 (d, *J* = 8.4 Hz, 0.29H), 3.81 (d, *J* = 8.4 Hz, 0.71H), 3.74-3.54 (m, 5H), 3.34-3.30 (m, 0.28H), 3.28-3.21 (m, 0.72H), 1.49 (s, 9H). $^{13}C$ NMR (151 MHz, $CDCl_3$) δ 154.32, 148.49, 138.03, 134.04, 131.95, 130.78, 129.99, 124.34, 93.54, 80.46, 61.15, 60.22, 56.16, 55.96, 55.14, 54.35, 50.15, 49.93, 49.22, 49.05, 42.95, 42.25, 28.61. HRMS (ESI-TOF) m/z calculated for $C_{23}H_{27}IN_3O_6S$ $[M+H]^+$ 600.0660; found 600.0664.

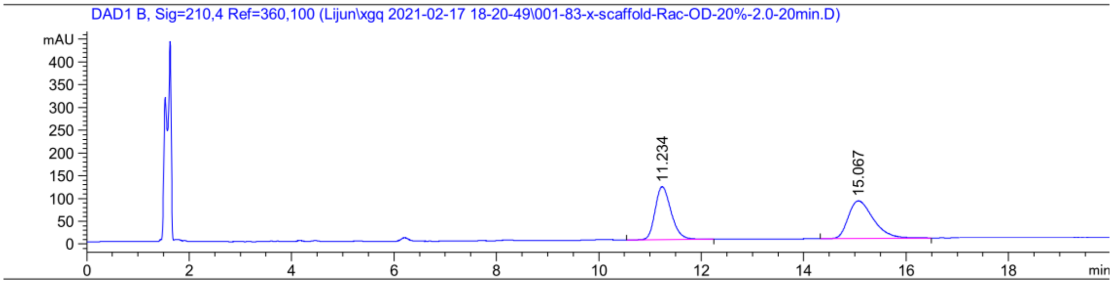

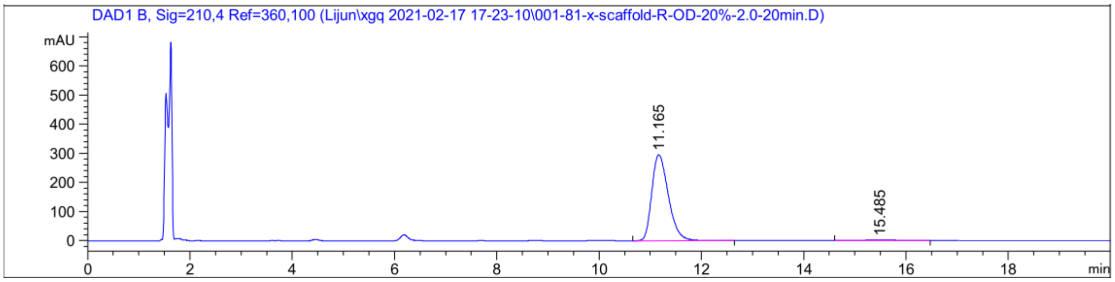

| Peak # | RetTime [min] | Type | Width [min] | Area [mAU*s] | Height [mAU] | Area % |
|---|---|---|---|---|---|---|
| 1 | 11.165 | BB | 0.3482 | 1069.92212 | 47.59830 | 98.7404 |
| 2 | 15.447 | MM | 0.5913 | 13.64903 | 3.84688e-1 | 1.2596 |
| Totals : | | | | 1083.57115 | 47.98299 | |

*tert*-butyl (*S*)-8-(4-iodophenyl)-2-((4-nitrophenyl)sulfonyl)-2,6-diazaspiro[3.4] octane-6-carboxylate (**20**) [prepared as for **19**]

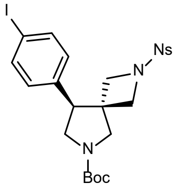

Pale yellow solid, 98.3% ee. $[\alpha]^{20}_{D}$ = -1.50 (c=1, $CHCl_3$). $^1H$ NMR (600 MHz, $CDCl_3$) δ 7.90 – 7.75 (m, 2H), 7.67 (d, *J* = 7.8 Hz, 2H), 7.64 – 7.54 (m, 2H), 6.92-6.84 (m, 2H), 4.07 – 3.95 (m, 2H), 3.87-3.80 (m, 1H), 3.76 – 3.55 (m, 5H), 3.33 – 3.20 (m, 1H), 1.52 (s, 9H). $^{13}C$ NMR (151 MHz, $CDCl_3$) δ 154.39, 154.31, 148.50, 148.47, 138.00, 134.12, 134.02, 132.03, 131.99, 130.78, 130.76, 130.04, 129.03, 128.00, 127.85, 124.36, 124.32, 93.57, 80.41, 80.26, 60.89, 60.06, 56.16, 56.00, 55.16, 54.53, 49.90, 49.08, 42.95, 42.40, 28.64, 28.61. HRMS (ESI-TOF) m/z calculated for $C_{23}H_{27}IN_3O_6S$ $[M+H]^+$ 600.0660; found: 600.0661.

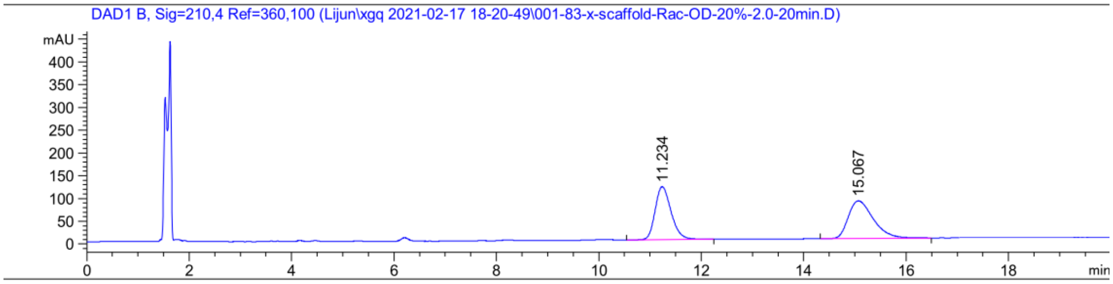

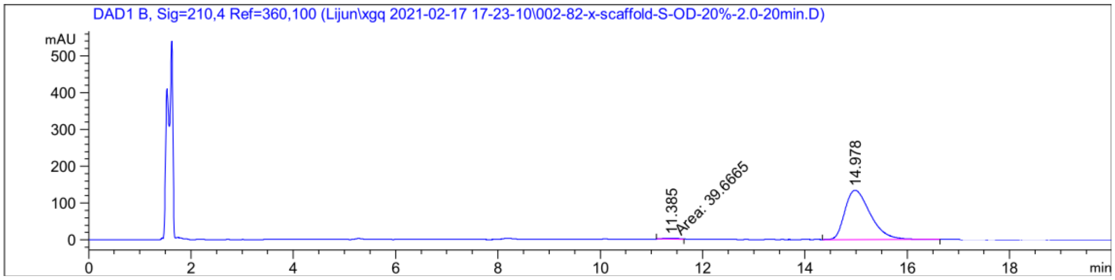

| Peak # | RetTime [min] | Type | Width [min] | Area [mAU*s] | Height [mAU] | Area % |
|---|---|---|---|---|---|---|
| 1 | 11.385 | MM | 0.3543 | 39.66647 | 1.86587 | 0.8539 |
| 2 | 14.978 | BB | 0.5354 | 4605.60449 | 133.87163 | 99.1461 |
| Totals : | | | | 4645.27096 | 135.73749 | |

(*R*)-5-(8-(4-iodophenyl)-2-((4-nitrophenyl)sulfonyl)-2,6-diazaspiro[3.4]octan-6-yl)-5-oxopentanoic acid (**21**)

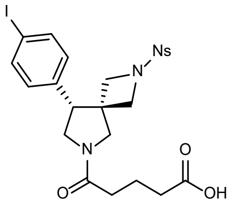

To a 12-mL vial was **19** (60 mg, 100 μmol, 1 eq), dissolved with $CH_2Cl_2$ (2 mL), then trifluoroacetic acid (0.5 mL) was added. The resulting solution was stirred at rt for 2 hr and then concentrated, the residue was dissolved with $CH_2Cl_2$ (2 mL), then $NEt_3$ (100 mg, 1.0 mmol, 10 eq) and glutaric anhydride (35 mg, 300 μmol, 3 eq) were added sequentially, and the resulting solution was stirred at rt overnight. The solution was washed with 2.0 M aq. HCl (10 mL) and extracted with $CH_2Cl_2$ (10 mL) 3 times. The combined organic portions were dried over anhydrous $Na_2SO_4$. The solution was concentrated and purified by preparative TLC chromatography to **21** (58 mg, 95 μmol, 95% yield) as a colorless foam and mixture of rotamers.

$[\alpha]^{20}_D$ = -3.20 (c=1, $CHCl_3$); $^1$H NMR (600 MHz, DMSO) δ 7.98 – 7.89 (m, 2H), 7.79 – 7.69 (m, 2H), 7.54 (dd, *J* = 13.5, 7.9 Hz, 2H), 6.94 (dd, *J* = 27.3, 7.6 Hz, 2H), 3.97 – 3.85 (m, 2.65H), 3.79 – 3.58 (m, 5.35H), 3.51 (dd, *J* = 19.8, 12.5 Hz, 1H), 2.34 – 2.20 (m, 4H), 1.78 – 1.67 (m, 2H). $^{13}$C NMR (151 MHz, DMSO) δ 174.38, 174.36, 170.44, 170.41, 148.18, 137.23, 136.57, 136.28, 134.90, 132.38, 132.36, 130.59, 130.56,

130.54, 127.02, 124.18, 94.04, 93.97, 58.01, 56.64, 56.29, 54.99, 54.79, 48.81, 48.29, 48.23, 46.69, 42.75, 41.15, 33.07, 33.03, 32.61, 32.48, 30.47, 19.82, 19.78. HRMS (ESI-TOF) m/z calculated for $C_{23}H_{25}IN_3O_7S$ $[M+H]^+$ 614.0452; found 614.0457.

(*S*)-5-(8-(4-iodophenyl)-2-((4-nitrophenyl)sulfonyl)-2,6-diazaspiro[3.4]octan-6-yl)-5-oxopentanoic acid (**22**) [prepared as for **21**]

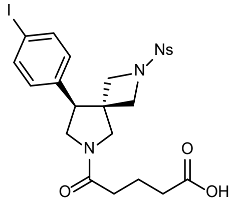

Colorless foam, mixture of rotamers. $[\alpha]^{20}_D$ = 3.00 (c=1, $CHCl_3$). $^1$H NMR (600 MHz, DMSO) δ 7.97 – 7.89 (m, 2H), 7.78 – 7.70 (m, 2H), 7.54 (dd, *J* = 14.0, 7.9 Hz, 2H), 6.94 (dd, *J* = 27.3, 7.9 Hz, 2H), 3.97 – 3.85 (m, 2.65H), 3.82 – 3.56 (m, 5.35H), 3.51 (dd, *J* = 18.6, 11.4 Hz, 1H), 2.34 – 2.20 (m, 4H), 1.73 (dt, *J* = 11.3, 7.1 Hz, 2H). $^{13}$C NMR (151 MHz, DMSO) δ 174.38, 174.36, 170.43, 170.41, 148.17, 139.23, 137.23, 136.56, 136.28, 134.90, 134.88, 132.38, 132.36, 130.59, 130.56, 130.54, 127.02, 124.97, 124.18, 94.04, 93.97, 58.01, 56.64, 56.29, 54.99, 54.79, 48.81, 48.29, 48.23, 46.69, 42.75, 41.15, 34.42, 33.07, 33.03, 32.61, 32.48, 30.47, 21.09, 19.81, 19.78. HRMS (ESI-TOF) m/z calculated for $C_{23}H_{25}IN_3O_7S$ $[M+H]^+$ 614.0452; found 614.0454.

(*R*)-5-(6-(((9H-fluoren-9-yl)methoxy)carbonyl)-8-(4-iodophenyl)-2,6-diazaspiro[3.4]octan-2-yl)-5-oxopentanoic acid (**23**)

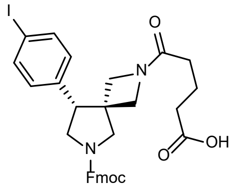

To a 12 mL vial was added **19** (60 mg, 0.1 mmol, 1 eq) and $K_2CO_3$ (28 mg, 0.2 mmol, 2 eq), dissolved with DMF (1 mL), then thiophenol (13 mg, 0.12 mmol, 1.2 eq) was added. The resulting solution was stirred at rt for 30 min and then concentrated, the residue was then dissolved with $CH_2Cl_2$ (2 mL), and $NEt_3$ (50 mg, 0.5 mmol, 5 eq) and glutaric anhydride (35 mg, 0.3 mmol, 3 eq) were added sequentially, the resulting solution was stirred at rt overnight. The solution was washed with 2.0 M aq. HCl (10 mL) and extracted with $CH_2Cl_2$ (10 mL) 3 times, then the organic portions were combined and dried over anhydrous $Na_2SO_4$. The solution was concentrated and purified by preparative TLC chromatography to give (*R*)-5-(6-(*tert*-butoxycarbonyl)-8-(4-iodophenyl)-2,6-diazaspiro[3.4]octan-2-yl)-5-oxopentanoic acid (47 mg, 89 μmol, 88% yield). The above isolated material was dissolved in a 12 mL vial with $CH_2Cl_2$ (2 mL), then trifluoroacetic acid (0.5 mL) was added. The resulting solution was stirred at rt for 2 hr and then concentrated, the residue was dissolved with $CH_2Cl_2$ (2 mL), $NEt_3$ (100 mg, 1.0 mmol, 12 eq) and fluorenylmethyloxycarbonyl chloride (52 mg, 0.2 mmol, 2.2 eq) were added sequentially, the resulting solution was stirred at rt for 4 h. The solution was washed with 1.0 M aq. HCl (10 mL) and extracted with $CH_2Cl_2$ (10 mL) 3 times. The organic portions were combined and dried over anhydrous $Na_2SO_4$. The solution was concentrated and purified by preparative TLC chromatography to give **23** (48 mg, 74 μmol, 85% yield).

Colorless foam, mixture of rotamers. $[\alpha]^{20}_{D}$ = -4.20 (c=1, $CHCl_3$). $^1$H NMR (600 MHz, $CDCl_3$) δ 7.77 (dd, *J* = 16.9, 7.5 Hz, 2H), 7.67 (q, *J* = 8.3 Hz, 2H), 7.59 (dd, *J* = 14.3, 7.4 Hz, 2H), 7.41 (dt, *J* = 13.2, 7.4 Hz, 2H), 7.36 – 7.29 (m, 2H), 6.92 – 6.83 (m, 2H), 4.58-4.46 (m, 2H), 4.25 (dt, *J* = 12.6, 6.6 Hz, 1H), 4.05 – 3.96 (m, 1H), 3.91 (d, *J* = 10.6 Hz, 1H), 3.87 – 3.50 (m, 6H), 3.33 – 3.24 (m, 1H), 2.45 – 2.32 (m, 2H), 2.15 (q, *J* = 8.2 Hz, 1H), 2.10 – 2.02 (m, 1H), 1.88 (dp, *J* = 39.2, 7.7, 7.2 Hz, 2H); $^{13}$C NMR (151 MHz, $CDCl_3$) δ 177.10, 172.98, 154.77, 143.91, 141.49, 138.24, 129.79, 129.75, 127.98, 127.94, 127.28, 127.24, 125.07, 125.04, 120.21, 120.15, 93.48, 93.45, 67.45, 60.97, 58.56, 58.08, 54.89, 54.76, 54.68, 54.57, 54.39, 51.88, 50.50, 50.20, 49.99, 49.87, 49.50, 49.39, 47.50, 43.42, 42.65, 33.16, 30.33, 30.30, 19.92, 19.89, 19.87; HRMS (ESI-TOF) m/z calculated for $C_{32}H_{32}IN_2O_5$ $[M+H]^+$ 651.1350; found 651.1351.

(*S*)-5-(6-(((9H-fluoren-9-yl)methoxy)carbonyl)-8-(4-iodophenyl)-2,6-diazaspiro[3.4]octan-2-yl)-5-oxopentanoic acid (**24**)

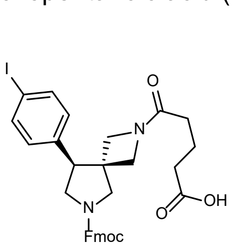

Colorless foam, mixture of rotamers. $[\alpha]^{20}_{D}$ = 4.10 (c=1, $CHCl_3$); $^1H$ NMR (600 MHz, $CDCl_3$) δ 7.77 (dd, *J* = 16.8, 7.5 Hz, 2H), 7.68 (q, *J* = 8.1 Hz, 2H), 7.59 (dd, *J* = 14.0, 7.3 Hz, 2H), 7.41 (dt, *J* = 13.4, 7.3 Hz, 2H), 7.37 – 7.29 (m, 2H), 6.94 – 6.81 (m, 2H), 4.60 – 4.45 (m, 2H), 4.26 (dt, *J* = 12.5, 6.4 Hz, 1H), 4.05 – 3.96 (m, 1H), 3.91 (d, *J* = 10.6 Hz, 1H), 3.87 – 3.49 (m, 6H), 3.29 (p, *J* = 11.0, 10.3 Hz, 1H), 2.44 – 2.33 (m, 2H), 2.15 (q, *J* = 8.7, 8.3 Hz, 1H), 2.10 – 2.03 (m, 1H), 1.95 – 1.82 (m, 2H); $^{13}C$ NMR (151 MHz, $CDCl_3$) δ 177.20, 172.95, 154.78, 144.01, 143.96, 143.90, 141.51, 138.31, 138.26, 129.80, 127.96, 127.94, 127.25, 125.05, 120.17, 93.46, 67.46, 60.99, 58.56, 54.40, 51.88, 50.54, 50.01, 49.89, 49.53, 49.42, 47.52, 43.41, 42.68, 33.20, 33.18, 33.12, 30.34, 30.31, 19.93, 19.90, 19.88; HRMS (ESI-TOF) m/z calculated for $C_{32}H_{32}IN_2O_5$ $[M+H]^+$ 651.1350; found 651.1354.

X-ray data for methyl (3*R*,4*S*)-4-(4-iodophenyl)-1-((4-nitrophenyl)sulfonyl)pyrrolidine-3-carboxylate (**25**)

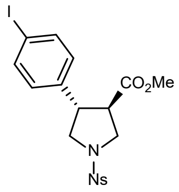

The single crystal X-ray diffraction studies were carried out on a Bruker ApexII-Ultra CCD diffractometer equipped with Mo $K_{\alpha}$ radiation ($\lambda$ =0.7107 Å).

Crystals of the subject compound were used as received. A 0.22 x 0.02 x 0.01 mm piece of a colorless crystal was mounted on a Cryoloop with Paratone oil. Data were collected in a nitrogen gas stream at 100(2) K using $\varpi$ scans. Crystal-to-detector distance was 40 mm and exposure time was 30 seconds depending on the $2\theta$ range per frame using a scan width of 0.70°. Data collection was 99.9 % complete to 25.242° in $\theta$. A total of 10995 reflections were collected covering the indices, -6<=h<=6, -14<=k<=14, -17<=l<=20. 4137 reflections were found to be symmetry independent, with a $R_{int}$ of 0.0469. Indexing and unit cell refinement indicated a **P**rimitive, **Monoclinic** lattice. The space group was found to be $\boldsymbol{P2_12_12}$. The data were integrated using the Bruker SAINT Software program and scaled using the SADABS software program. Solution by direct methods (SHELXT) produced a complete phasing model consistent with the proposed structure.

All nonhydrogen atoms were refined anisotropically by full-matrix least-squares (SHELXL-2014). All carbon bonded hydrogen atoms were placed using a riding model. Their positions were constrained relative to their parent atom using the appropriate HFIX command in SHELXL-2014. Crystallographic data are summarized in **Supplementary Table 1**.

Absolute stereochemistry was conclusively assigned (Flack = 0.14(12)). There is one copy of the molecule in the asymmetric unit. The chemical formula of the compound is: $C_{18}H_{17}IN_2O_6S$.

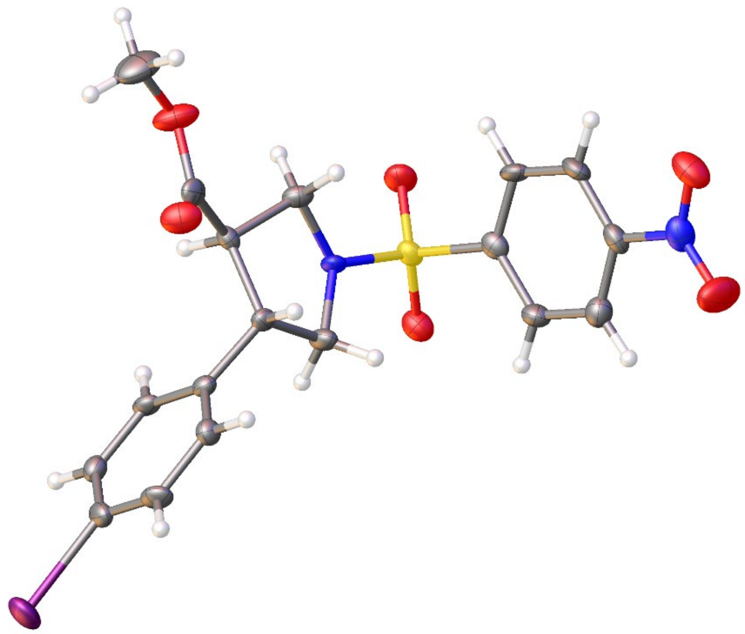

**Supplementary Table 1 | Crystal data and structure refinement.**

| | |
|---|---|
| Empirical formula | C18 H17 I N2 O6 S |
| Formula weight | 516.29 |
| Temperature | 100.0 K |
| Wavelength | 0.71073 Å |

| | | |
|---|---|---|
| Crystal system | Monoclinic | |
| Space group | P 1 21 1 | |
| Unit cell dimensions | a = 5.2498(12) Å | a = 90°. |
| | b = 11.580(2) Å | b = 96.166(7)°. |
| | c = 16.099(3) Å | g = 90°. |
| Volume | 973.0(4) $Å^3$ | |
| Z | 2 | |
| Density (calculated) | 1.762 $Mg/m^3$ | |
| Absorption coefficient | 1.791 mm-1 | |
| F(000) | 512 | |
| Crystal size | 0.22 x 0.02 x 0.01 $mm^3$ | |
| Theta range for data collection | 2.171 to 26.732°. | |
| Index ranges | -6<=h<=6, -14<=k<=14, -17<=l<=20 | |
| Reflections collected | 10995 | |
| Independent reflections | 4137 [R(int) = 0.0469] | |
| Completeness to theta = 25.242° | 99.90% | |
| Absorption correction | Semi-empirical from eq | |
| Max. and min. transmission | 0.4912 and 0.4352 | |
| Refinement method | Full-matrix least-squares on $F^2$ | |
| Data / restraints / parameters | 4137 / 1 / 254 | |
| Goodness-of-fit on $F^2$ | 1.032 | |
| Final R indices [I>2sigma(I)] | R1 = 0.0278, wR2 = 0.0587 | |
| R indices (all data) | R1 = 0.0300, wR2 = 0.0594 | |
| Absolute structure parameter | 0.014(12) | |
| Largest diff. peak and hole | 0.724 and -0.368 e.$Å^{-3}$ | |

**Supplementary Table 2 | Atomic coordinates (x $10^4$) and equivalent isotropic displacement parameters ($Å^2$ x $10^3$). U(eq) is defined as one third of the trace of the orthogonalized $U^{ij}$ tensor.**

| | x | y | z | U(eq) |
|---|---|---|---|---|
| I(1) | 5063(1) | 1190(1) | 9851(1) | 24(1) |
| S(1) | -90(2) | 6773(1) | 5621(1) | 13(1) |
| O(1) | 8377(7) | 7080(3) | 2865(2) | 24(1) |
| O(2) | 6931(9) | 5360(3) | 2582(3) | 33(1) |
| O(3) | -1969(6) | 5874(3) | 5517(2) | 17(1) |
| O(4) | -877(6) | 7955(3) | 5679(2) | 18(1) |
| O(5) | 9161(6) | 6565(3) | 8201(2) | 21(1) |
| O(6) | 7049(7) | 8244(3) | 8180(3) | 24(1) |
| N(1) | 6919(7) | 6277(5) | 2962(2) | 18(1) |
| N(2) | 1753(7) | 6485(3) | 6459(2) | 12(1) |
| C(1) | 3287(9) | 7598(4) | 4582(3) | 15(1) |
| C(2) | 4954(9) | 7464(4) | 3980(3) | 16(1) |
| C(3) | 5074(8) | 6409(4) | 3589(2) | 15(1) |
| C(4) | 3569(9) | 5474(4) | 3755(3) | 16(1) |
| C(5) | 1912(9) | 5607(4) | 4371(3) | 16(1) |
| C(6) | 1816(9) | 6667(4) | 4777(3) | 13(1) |
| C(7) | 3609(9) | 7330(4) | 6855(3) | 13(1) |
| C(8) | 4775(8) | 6643(4) | 7617(3) | 10(1) |
| C(9) | 7245(9) | 7113(4) | 8024(3) | 14(1) |
| C(10) | 9311(10) | 8793(5) | 8610(4) | 31(1) |
| C(11) | 4995(9) | 5403(4) | 7280(3) | 12(1) |
| C(12) | 2671(10) | 5291(4) | 6608(3) | 15(1) |
| C(13) | 5104(8) | 4443(4) | 7917(3) | 11(1) |
| C(14) | 3265(9) | 4360(4) | 8479(3) | 12(1) |
| C(15) | 3284(9) | 3448(4) | 9035(3) | 13(1) |
| C(16) | 5140(9) | 2597(4) | 9030(3) | 15(1) |
| C(17) | 6986(9) | 2656(4) | 8484(3) | 15(1) |

| | | | | |
|---|---|---|---|---|
| C(18) | 6965(9) | 3591(4) | 7931(3) | 13(1) |

**Supplementary Table 3 | Bond lengths [Å] and angles [°].**

| | | | | | |
|---|---|---|---|---|---|
| I(1)-C(16) | 2.100(4) | C(16)-C(17) | 1.378(6) | C(9)-C(8)-C(7) | 114.6(4) |
| S(1)-O(3) | 1.432(3) | C(17)-H(17) | 0.95 | C(9)-C(8)-H(8) | 108.2 |
| S(1)-O(4) | 1.435(3) | C(17)-C(18) | 1.401(7) | C(9)-C(8)-C(11) | 113.5(4) |
| S(1)-N(2) | 1.609(4) | C(18)-H(18) | 0.95 | C(11)-C(8)-H(8) | 108.2 |
| S(1)-C(6) | 1.776(5) | O(3)-S(1)-O(4) | 120.1(2) | O(5)-C(9)-O(6) | 123.6(4) |
| O(1)-N(1) | 1.225(6) | O(3)-S(1)-N(2) | 106.98(19) | O(5)-C(9)-C(8) | 125.4(4) |
| O(2)-N(1) | 1.227(6) | O(3)-S(1)-C(6) | 107.4(2) | O(6)-C(9)-C(8) | 111.0(4) |
| O(5)-C(9) | 1.198(5) | O(4)-S(1)-N(2) | 107.14(19) | O(6)-C(10)-H(10A) | 109.5 |
| O(6)-C(9) | 1.339(6) | O(4)-S(1)-C(6) | 107.8(2) | O(6)-C(10)-H(10B) | 109.5 |
| O(6)-C(10) | 1.455(6) | N(2)-S(1)-C(6) | 106.8(2) | O(6)-C(10)-H(10C) | 109.5 |
| N(1)-C(3) | 1.479(5) | C(9)-O(6)-C(10) | 116.2(4) | H(10A)-C(10)-H(10B) | 109.5 |
| N(2)-C(7) | 1.476(6) | O(1)-N(1)-O(2) | 123.8(4) | H(10A)-C(10)-H(10C) | 109.5 |
| N(2)-C(12) | 1.476(6) | O(1)-N(1)-C(3) | 118.2(4) | H(10B)-C(10)-H(10C) | 109.5 |
| C(1)-H(1) | 0.95 | O(2)-N(1)-C(3) | 117.9(4) | C(8)-C(11)-H(11) | 107.9 |
| C(1)-C(2) | 1.383(6) | C(7)-N(2)-S(1) | 122.3(3) | C(8)-C(11)-C(12) | 103.9(4) |
| C(1)-C(6) | 1.381(6) | C(12)-N(2)-S(1) | 119.2(3) | C(12)-C(11)-H(11) | 107.9 |
| C(2)-H(2) | 0.95 | C(12)-N(2)-C(7) | 111.2(4) | C(13)-C(11)-C(8) | 116.5(3) |
| C(2)-C(3) | 1.380(6) | C(2)-C(1)-H(1) | 120.6 | C(13)-C(11)-H(11) | 107.9 |
| C(3)-C(4) | 1.383(6) | C(6)-C(1)-H(1) | 120.6 | C(13)-C(11)-C(12) | 112.5(4) |
| C(4)-H(4) | 0.95 | C(6)-C(1)-C(2) | 118.8(4) | N(2)-C(12)-C(11) | 104.7(4) |
| C(4)-C(5) | 1.397(7) | C(1)-C(2)-H(2) | 120.6 | N(2)-C(12)-H(12A) | 110.8 |
| C(5)-H(5) | 0.95 | C(3)-C(2)-C(1) | 118.9(4) | N(2)-C(12)-H(12B) | 110.8 |
| C(5)-C(6) | 1.394(6) | C(3)-C(2)-H(2) | 120.6 | C(11)-C(12)-H(12A) | 110.8 |
| C(7)-H(7A) | 0.99 | C(2)-C(3)-N(1) | 117.9(4) | C(11)-C(12)-H(12B) | 110.8 |

| | | | | | |
|---|---|---|---|---|---|
| C(7)-H(7B) | 0.99 | C(2)-C(3)-C(4) | 123.4(4) | H(12A)-C(12)-H(12B) | 108.9 |
| C(7)-C(8) | 1.532(6) | C(4)-C(3)-N(1) | 118.7(4) | C(14)-C(13)-C(11) | 120.9(4) |
| C(8)-H(8) | 1 | C(3)-C(4)-H(4) | 121.2 | C(18)-C(13)-C(11) | 120.7(4) |
| C(8)-C(9) | 1.492(6) | C(3)-C(4)-C(5) | 117.6(4) | C(18)-C(13)-C(14) | 118.4(4) |
| C(8)-C(11) | 1.544(6) | C(5)-C(4)-H(4) | 121.2 | C(13)-C(14)-H(14) | 119.5 |
| C(10)-H(10A) | 0.98 | C(4)-C(5)-H(5) | 120.4 | C(15)-C(14)-C(13) | 121.0(4) |
| C(10)-H(10B) | 0.98 | C(6)-C(5)-C(4) | 119.1(4) | C(15)-C(14)-H(14) | 119.5 |
| C(10)-H(10C) | 0.98 | C(6)-C(5)-H(5) | 120.4 | C(14)-C(15)-H(15) | 120.2 |
| C(11)-H(11) | 1 | C(1)-C(6)-S(1) | 119.6(4) | C(14)-C(15)-C(16) | 119.7(4) |
| C(11)-C(12) | 1.547(6) | C(1)-C(6)-C(5) | 122.1(4) | C(16)-C(15)-H(15) | 120.2 |
| C(11)-C(13) | 1.508(6) | C(5)-C(6)-S(1) | 118.1(3) | C(15)-C(16)-I(1) | 119.1(3) |
| C(12)-H(12A) | 0.99 | N(2)-C(7)-H(7A) | 111.6 | C(17)-C(16)-I(1) | 120.1(3) |
| C(12)-H(12B) | 0.99 | N(2)-C(7)-H(7B) | 111.6 | C(17)-C(16)-C(15) | 120.7(4) |
| C(13)-C(14) | 1.396(6) | N(2)-C(7)-C(8) | 101.0(4) | C(16)-C(17)-H(17) | 120.5 |
| C(13)-C(18) | 1.388(6) | H(7A)-C(7)-H(7B) | 109.4 | C(16)-C(17)-C(18) | 119.1(4) |
| C(14)-H(14) | 0.95 | C(8)-C(7)-H(7A) | 111.6 | C(18)-C(17)-H(17) | 120.5 |
| C(14)-C(15) | 1.384(6) | C(8)-C(7)-H(7B) | 111.6 | C(13)-C(18)-C(17) | 121.2(4) |
| C(15)-H(15) | 0.95 | C(7)-C(8)-H(8) | 108.2 | C(13)-C(18)-H(18) | 119.4 |
| C(15)-C(16) | 1.386(6) | C(7)-C(8)-C(11) | 103.9(4) | C(17)-C(18)-H(18) | 119.4 |

**Supplementary Table 4 | Anisotropic displacement parameters (Å$^2$ x 10$^3$). The anisotropic displacement factor exponent takes the form: $-2\pi^2[ h^2 a^{*2}U^{11} + ... + 2 hr k a^* b^* U^{12}]$**

| | $U^{11}$ | $U^{22}$ | $U^{33}$ | $U^{23}$ | $U^{13}$ | $U^{12}$ |
|---|---|---|---|---|---|---|
| I(1) | 39(1) | 16(1) | 18(1) | 9(1) | 6(1) | 4(1) |
| S(1) | 15(1) | 11(1) | 12(1) | 2(1) | 2(1) | 2(1) |
| O(1) | 20(2) | 25(2) | 25(2) | 10(2) | 4(2) | -2(2) |
| O(2) | 46(3) | 23(2) | 35(2) | -10(2) | 22(2) | -4(2) |

| | | | | | | |
|---|---|---|---|---|---|---|
| O(3) | 16(2) | 18(2) | 18(2) | 4(1) | 2(1) | 0(1) |
| O(4) | 23(2) | 12(2) | 19(2) | 1(1) | 4(1) | 6(1) |
| O(5) | 15(2) | 17(2) | 30(2) | -1(1) | 1(1) | 2(1) |
| O(6) | 22(2) | 12(2) | 37(2) | -7(2) | 2(2) | -1(2) |
| N(1) | 19(2) | 20(2) | 16(2) | 3(3) | 4(1) | 7(2) |
| N(2) | 19(2) | 5(2) | 12(2) | 0(1) | 1(1) | 1(1) |
| C(1) | 22(2) | 6(2) | 16(2) | 0(2) | 0(2) | -1(2) |
| C(2) | 23(3) | 11(2) | 14(2) | 5(2) | 2(2) | -4(2) |
| C(3) | 18(2) | 16(3) | 12(2) | 5(2) | 3(2) | 1(2) |
| C(4) | 19(3) | 17(2) | 13(2) | 0(2) | 2(2) | 1(2) |
| C(5) | 21(3) | 11(2) | 15(2) | 2(2) | 2(2) | -3(2) |
| C(6) | 12(2) | 13(2) | 13(2) | 3(2) | -1(2) | 2(2) |
| C(7) | 13(2) | 9(2) | 17(3) | -1(2) | 2(2) | 0(2) |
| C(8) | 11(2) | 11(2) | 10(2) | 3(2) | 2(2) | 1(2) |
| C(9) | 18(2) | 13(2) | 12(2) | -2(2) | 8(2) | 1(2) |
| C(10) | 30(3) | 23(3) | 39(3) | -12(3) | 6(2) | -11(3) |
| C(11) | 15(2) | 9(2) | 11(2) | -1(2) | 4(2) | 1(2) |
| C(12) | 21(3) | 8(2) | 14(2) | 2(2) | -3(2) | 0(2) |
| C(13) | 13(2) | 8(2) | 10(2) | -1(2) | -1(2) | 0(2) |
| C(14) | 16(2) | 7(2) | 14(2) | -1(2) | -1(2) | 2(2) |
| C(15) | 15(2) | 14(3) | 10(2) | -2(2) | 2(2) | -2(2) |
| C(16) | 23(3) | 12(2) | 10(2) | 2(2) | -3(2) | -2(2) |
| C(17) | 18(2) | 7(2) | 18(2) | 1(2) | -2(2) | 4(2) |
| C(18) | 16(2) | 13(3) | 11(3) | 0(2) | 4(2) | -1(2) |

**Supplementary Table 5 | Hydrogen coordinates (x $10^4$) and isotropic displacement parameters (Å$^2$x $10^3$).**

| | x | y | z | U(eq) |
|---|---|---|---|---|
| H(1) | 3156 | 8318 | 4857 | 18 |
| H(2) | 5999 | 8088 | 3838 | 20 |
| H(4) | 3659 | 4767 | 3460 | 20 |
| H(5) | 864 | 4984 | 4512 | 19 |
| H(7A) | 2746 | 8039 | 7023 | 16 |
| H(7B) | 4919 | 7537 | 6482 | 16 |
| H(8) | 3513 | 6637 | 8040 | 13 |
| H(10A) | 10695 | 8787 | 8247 | 46 |
| H(10B) | 9857 | 8369 | 9125 | 46 |
| H(10C) | 8904 | 9592 | 8746 | 46 |
| H(11) | 6587 | 5360 | 6992 | 14 |
| H(12A) | 3192 | 4947 | 6089 | 18 |
| H(12B) | 1319 | 4804 | 6812 | 18 |
| H(14) | 1980 | 4938 | 8479 | 15 |
| H(15) | 2032 | 3405 | 9418 | 15 |
| H(17) | 8255 | 2071 | 8483 | 18 |
| H(18) | 8248 | 3642 | 7559 | 16 |

#### 1.4. Enumerated DEL targets synthesized off-DNA

The following targets in this section were synthesized by multiple chemists and did not follow consistent protocols. A full synthetic protocol is given for each target compound to provide added clarity.

2-((3'-((2R,3R)-1-(2-(allylamino)thiazole-5-carbonyl)-2-(methylcarbamoyl)pyrrolidin-3-yl)-5-fluoro-[1,1'-biphenyl]-3-yl)oxy)acetic acid (**3**)

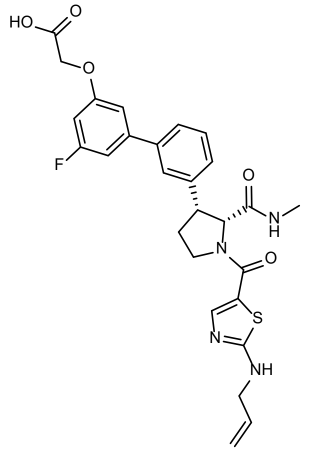

To a vial charged with **6c** (30 mg, 1 eq, 72 µmol), $NH_3MeCl$ (9.7 mg, 2.0 eq) and HATU (32.9 mg, 1.2 eq) were added. The vial was purged with $N_2$, before addition of anhydrous DMF (750 µL) and DIPEA (63 µL, 5 eq). The reaction was stirred overnight at rt, diluted with ethyl acetate and washed twice with water, then brine, before being dried over anhydrous $MgSO_4$, filtered, and removed solvent under reduced pressure. The crude *tert*-butyl (2*R*,3*R*)-3-(3-iodophenyl)-2-(methylcarbamoyl)pyrrolidine-1-carboxylate (29 mg, 68 µmol, 94%) was used directly in the next step without additional purification.

A vial was charged with *tert*-butyl (2*R*,3*R*)-3-(3-iodophenyl)-2-(methylcarbamoyl)pyrrolidine-1-carboxylate (27 mg, 63 µmol, 1 eq), 2-(3-borono-5-fluorophenoxy)acetic acid (16 mg, 75 µmol, 1.2 eq), $Pd(PPh_3)_4$ (7.3 mg, 6.3 µmol, 0.1 eq), and $Na_2CO_3$ (27 mg, 250 µmol, 4.0 eq). The vial was sealed and purged with $N_2$ before addition of water (0.5 mL), DMF (0.5 mL), and MeCN (0.5 mL) were added, and the reaction was stirred 80 °C for 1 hr under microwave irradiation. The crude solution was purified by preparative HPLC to afford 2-((3'-((2*R*,3*R*)-1-(*tert*-butoxycarbonyl)-2-(methylcarbamoyl)pyrrolidin-3-yl)-5-fluoro-[1,1'-biphenyl]-3-yl)oxy)acetic acid (5.9 mg, 12.6 µmol, 20%).

2-(Allylamino)thiazole-5-carboxylic acid (30 mg, 0.16 mmol, 1 eq), NHS (21 mg, 0.18 mmol, 1.1 eq) and EDC (34 mg, 0.18 mmol, 1.1 eq) was dissolved in THF and stirred at rt for 6 hr. The resulting solution was purified by flash column chromatography (0–100% EtOAC in hexanes) to afford 2,5-dioxopyrrolidin-1-yl 2-(allylamino)thiazole-5-carboxylate (29 mg, 0.10 mmol, 65%).

2-((3'-((2*R*,3*R*)-1-(*tert*-Butoxycarbonyl)-2-(methylcarbamoyl)pyrrolidin-3-yl)-5-fluoro-[1,1'-biphenyl]-3-yl)oxy)acetic acid (5 mg, 11 μmol, 1 eq) was dissolved in $CH_2Cl_2$ (1 mL), followed by addition of TFA (0.2 mL). The mixture was stirred at rt for 2 hr before removal of solvents under reduced pressure, then co-evaporated with toluene. To the residue was added 2,5-dioxopyrrolidin-1-yl 2-(allylamino)thiazole-5-carboxylate (4.46 mg, 15.9 μmol, 1.5 eq) before being dissolved in DMF (250 μL), followed by addition of DIPEA (5 μL, 3 eq). The mixture was stirred overnight, then was purified by preparative HPLC to afford **3** (1.5 mg, 2.75 μmol, 25%).

$^1H$ NMR (600 MHz, DMSO) δ 8.39 – 8.19 (m, 1.5H), 7.70 – 7.50 (m, 3.5H), 7.37 (t, *J* = 7.8 Hz, 1H), 7.23 (d, *J* = 8.0 Hz, 1H), 7.05 (d, *J* = 9.9 Hz, 1H), 7.01 (s, 1H), 6.71 (dd, *J* = 10.7, 2.7 Hz, 1H), 5.96 – 5.71 (m, 0.5H), 5.24 (d, *J* = 19.4 Hz, 0.5H), 5.13 (d, *J* = 11.6 Hz, 0.5H), 4.05 (t, *J* = 10.0 Hz, 0.6H), 4.00 – 3.76 (m, 3H), 3.76 – 3.58 (m, 0.6H), 2.79 – 2.63 (m, 0.5H), 2.33 – 2.07 (m, 3.5H). LRMS (ESI): calculated for $C_{27}H_{28}FN_4O_5S$ $[M + H]^+$, 539.18; found, 539.36.

(2*S*,4*S*)-4-(3-((*E*)-2-cyclohexylvinyl)phenoxy)-1-(3-ethynylbenzyl)-*N*-methylpyrrolidine-2-carboxamide (**27**)

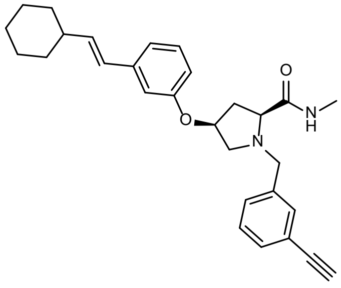

To a solution of (2*S*,4*S*)-4-(3-iodophenoxy)-2-(methoxycarbonyl)pyrrolidin-1-ium 2,2,2-trifluoroacetate (50 mg, 1 eq, 0.11 mmol) di-*tert*-butyl dicarbonate (28 mg, 1.2 eq, 0.13 mmol) in $CH_2Cl_2$ (1.0 mL) was added $NEt_3$ (13 mg, 18 μL, 1.2 eq, 0.13 mmol). The resulting mixture was stirred at rt for 16 h. The reaction was then concentrated, and the crude residue purified by column chromatography on silica gel (12 g, 0 to 70% EtOAc in Hexanes), affording the Boc-protected amine, **9aii**. To **9aii** (50 mg, 1 eq, 0.11 mmol) were added THF (3 mL) and LiOH (40 mg, 1.7 mL, 1 M, 15 eq, 1.7 mmol). The resulting reaction mixture was vigorously stirred for 16 h, then diluted with $H_2O$ (15 mL). The aqueous layer was acidified with 1N HCl to pH of ~2 and extracted with $CH_2Cl_2$ (3 x 20 mL). The combined organics were then dried over $Na_2SO_4$ and concentrated under reduced pressure to afford the crude acid. This material was dissolved in $CH_2Cl_2$ (2 mL) and treated with HATU (58 mg, 1.2 eq, 0.15 mmol) and DIPEA (49 mg, 66 µL, 3.0 eq, 0.38 mmol). The resulting reaction mixture was stirred for 15 min at rt and methylamine hydrochloride (13 mg, 1.5 eq, 0.19 mmol) was added. The reaction was stirred for 16 hr at rt. The $CH_2Cl_2$ was removed under reduced pressure and the crude residue was diluted in EtOAc (25 mL). The organics were washed with water (1 x 25 mL), 10% aq. NaOH (1 x 25 mL) and brine (1 x 25 mL), dried over $Na_2SO_4$, and concentrated under reduced pressure to afford the cure amide, which was purified by column chromatography on silica gel (4 g, 0 to 80% EtOAc in hexanes). The resulting material was then charged to a 5 mL µW vial equipped with a triangular magnetic stir bar. (*E*)-(2-cyclohexylvinyl)boronic acid (23 mg, 1.2 eq, 0.15 mmol), Pd-Xphos-G2 (19 mg, 0.20 eq, 25 µmol), and DMF (0.93 mL) were added, followed by aq. $K_3PO_4$ (0.10 g, 0.27 mL, 1.8 molar, 4 eq, 0.49 mmol). The resulting mixture was sparged with argon for 3 min, then the vial was sealed and heated to 60 °C for 1 hour in a µW reactor. The reaction mixture was diluted with EtOAc (40 mL) and washed with $H_2O$ (1 x 15 mL) then brine (2 x 15 mL). The organics were then dried over $Na_2SO_4$ and concentrated under reduced pressure. The crude residues were purified by column chromatography on silica gel (4 g, 0% to 90% EtOAc in hexanes) to afford the crude product as a colorless oil which was immediately dissolved in methanolic HCl (1.0 mL, 2 M, 2.0 mmol, 29 eq.) and incubated for 3 h, after which time volatiles were removed under reduced pressure to afford (2*S*,4*S*)-4-(3-((*E*)-2-cyclohexylvinyl)phenoxy)-*N*-methylpyrrolidine-2-carboxamide hydrochloride (26 mg, 70 µmol, 52% over five steps).

The above hydrochloride salt (26 mg, 1 eq., 71 µmol) was taken up in DCE (0.5 mL) and treated with 3-ethynylbenzaldehyde (11 mg, 1.2 eq, 86 µmol) and sodium triacetoxyhydroborate (23 mg, 1.5 eq, 0.11 mmol). The resulting mixture was stirred for 1 hr at rt, then concentrated under reduced pressure, re-dissolved in $CH_2Cl_2$ (25 mL), and washed with water (15 mL). The aqueous layer was then back-extracted with $CH_2Cl_2$ (2 x 25 mL) and the combined organics dried over $Na_2SO_4$ and concentrated under reduced pressure. The crude residue was purified by preparative HPLC to afford **27** (15 mg, 33 μmol, 46% over two steps) as a white solid.

$^{1}$H NMR (400 MHz, $CD_3OD$) δ 7.53 – 7.49 (m, 1H), 7.44 – 7.29 (m, 3H), 7.16 (t, *J* = 7.9 Hz, 1H), 6.95 – 6.90 (m, 1H), 6.83 (t, *J* = 2.0 Hz, 1H), 6.68 (dd, *J* = 8.1, 2.3 Hz, 1H), 6.31 (d, *J* = 15.9 Hz, 1H), 6.17 (dd, *J* = 16.0, 6.9 Hz, 1H), 4.92 – 4.87 (m, 1H), 3.82 (d, *J* = 12.9 Hz, 1H), 3.66 (d, *J* = 12.8 Hz, 1H), 3.49 (s, 1H), 3.32 – 3.24 (m, 2H), 2.77 (dd, *J* = 10.8, 4.4 Hz, 1H), 2.71 (s, 3H), 2.62 (ddd, *J* = 14.2, 10.3, 5.6 Hz, 1H), 2.18 – 2.02 (m, 2H), 1.87 – 1.65 (m, 5H), 1.44 – 1.14 (m, 5H). $^{13}$C NMR (151 MHz, DMSO) δ 173.11, 173.03, 157.82, 139.48, 139.23, 137.11, 132.57, 130.97, 130.15, 130.00, 129.00, 127.40, 122.09, 118.87, 114.34, 113.28, 84.03, 81.18, 75.58, 66.96, 58.89, 58.59, 40.89, 37.13, 32.86, 26.14, 26.03, 25.94, 25.90. HRMS (ESI): calculated for $C_{29}H_{35}N_2O_2$ $[M + H]^+$, 443.2693; found, 443.2689.

(2*R*,4*S*)-4-(3-((*E*)-2-cyclohexylvinyl)phenoxy)-1-(3-ethynylbenzyl)-*N*-methylpyrrolidine-2-carboxamide (**28**)

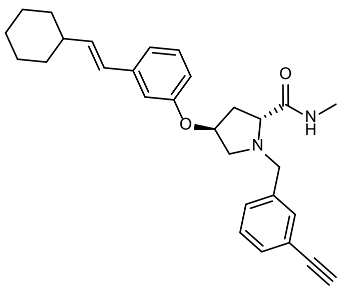

To a 1-dram vial containing (2*R*,4*S*)-1-(*tert*-butoxycarbonyl)-4-(3-iodophenoxy)pyrrolidine-2-carboxylic acid (17 mg, 39 μmol, 1 eq), methylamine hydrochloride (3.2 mg, 47 μmol, 1.2 eq) and HATU (18 mg, 47 μmol, 1.2 eq) were added. Additional $CH_2Cl_2$ (0.4 mL) was added, followed by DIPEA (34 μL, 200 μmol, 5 eq). The resulting yellow solution was stirred overnight, then diluted with $CH_2Cl_2$ and washed with water (3 x 1 mL) and brine (1 x 1 mL). The organic phase was dried over anhydrous $MgSO_4$, then concentrated under reduced pressure. The combined crude material from several reactions was pooled to provide crude acid (61 mg, 0.14 mmol), which was charged to a 2 mL μW vial. (*E*)-(2-cyclohexylvinyl)boronic acid (32 mg, 0.21 mmol, 1.5 eq) and XPhos-Pd-G2 (22 mg, 28 μmol, 0.2 eq) were added; the vial was then capped with a septum and purged with $N_2$ (3 times). Anhydrous DMF (2 mL) was added, followed by aq. $K_3PO_4$ (47 mL of a 1.8 M solution, 6.00 eq). The reaction was then heated to 60 °C for 50 mins in a μW reactor. The resulting dark brown suspension was transferred to a scintillation vial and concentrated under reduced pressure, then diluted with EtOAc and washed with water and brine. The organics were dried over $MgSO_4$, then concentrated under reduced pressure to provide crude material which was purified by flash column chromatography on silica gel (ethyl acetate/hexanes), then dissolved in $CH_2Cl_2$ (2 mL) and treated with TFA (0.4 mL), stirring at r.t. for 1.5 hr at which time LCMS showed complete consumption of starting material. Concentration under reduced pressure followed by azeotropic removal of residual TFA with toluene gave the free amine (20 mg, 4% over three steps). Half of this material (10 mgs, 0.03 mmol, 1.00 eq) was charged to a 1-dram vial containing 3-ethynylbenzaldehyde (5 mg, 0.04 mmol, 1.2 eq) and sodium acetoxyborohydride (10 mg, 0.06 mmol, 2.0 eq). The vial was purged with $N_2$, then charged with DCE (0.2 mL) and $NEt_3$ (4 µL, 0.03 mmol, 1.0 eq), yielding a yellow suspension which was stirred overnight. The reaction solution was then concentrated under reduced pressure, dissolved in DMSO and purified by preparative HPLC, affording **28** as a white solid (9 mg, 70%).

$^1H$ NMR (400 MHz, $CDCl_3$) δ 7.46 – 7.38 (m, 2H), 7.30 (t, *J* = 7.5 Hz, 1H), 7.20 – 7.10 (m, 2H), 6.94 (d, *J* = 7.6 Hz, 1H), 6.80 (t, *J* = 2.0 Hz, 1H), 6.64 (dd, *J* = 8.1, 2.5 Hz, 1H), 6.28 (d, *J* = 16.0 Hz, 1H), 6.14 (dd, *J* = 16.0, 6.8 Hz, 1H), 4.84 – 4.76 (m, 1H), 3.92 (d, *J* = 13.4 Hz, 1H), 3.70 – 3.55 (m, 2H), 3.41 (dd, *J* = 12.0, 4.9 Hz, 1H), 3.09 (s, 1H), 2.81 (d, *J* = 5.0 Hz, 3H), 2.78 – 2.69 (m, 1H), 2.59 – 2.48 (m, 1H), 2.23 – 2.04 (m, 2H), 1.85 – 1.72 (m, 4H), 1.72 – 1.62 (m, 1H), 1.30 (tt, *J* = 12.1, 3.1 Hz, 2H), 1.24 – 1.11 (m, 3H). $^{13}C$ NMR (101 MHz, $CDCl_3$) δ 173.8, 157.9, 139.8, 138.7, 132.1, 131.2, 129.5, 129.1, 128.6, 126.9, 122.4, 119.1, 113.6, 113.1, 83.4, 77.5, 77.2, 76.1, 67.0, 60.1, 58.9, 41.1, 37.6, 32.9, 26.2, 26.0, 25.8. HRMS (ESI): calculated for $C_{29}H_{35}N_2O_2$ $[M + H]^+$, 443.2693; found, 443.2697.

(2*S*,4*S*)-4-(4-((*E*)-2-cyclohexylvinyl)phenoxy)-1-(3-cyclopropyl-1-ethyl-1H-pyrazole-5-carbonyl)-*N*-methylpyrrolidine-2-carboxamide (**29**)

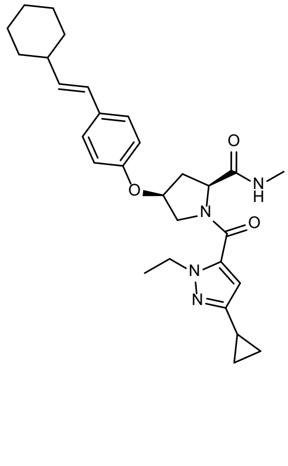

To a solution of (2*S*,4*S*)-4-(4-iodophenoxy)-2-(methoxycarbonyl)pyrrolidin-1-ium 2,2,2-trifluoroacetate (55 mg, 1 eq, 0.12 mmol) and di-*tert*-butyl dicarbonate (31 mg, 1.2 eq, 0.14 mmol) in $CH_2Cl_2$ (1.0 mL) was added $NEt_3$ (0.025 mL, 0.17 mmol). The resulting mixture was stirred at rt for 16 h, then concentrated under reduced pressure. The crude residue was purified by column chromatography on silica gel (12 g, 0 to 70% EtOAc in hexanes) to afford the desired product as a clear oil which was immediately dissolved in THF (3 mL) and treated with aq. LiOH (40 mg, 1.7 mL of a 1 M solution, 15 eq, 1.7 mmol). The resulting mixture was vigorously stirred 16 h, then diluted with water (15 mL) and acidified with 1 N aq. HCl to pH ~2. The aqueous mixture was extracted with $CH_2Cl_2$ (3 x 20 mL); organics were then combined dried over $Na_2SO_4$ and concentrated under reduced pressure to reveal the crude acid. This material was immediately dissolved in $CH_2Cl_2$ (2 mL) and treated with HATU (58 mg, 1.2 eq, 0.15 mmol) and DIPEA (49 mg, 66 µL, 3.0 eq, 0.38 mmol). The resulting mixture was stirred 15 min at rt, then methylamine hydrochloride (13 mg, 1.5 eq, 0.19 mmol) was added and the reaction stirred an additional 3 hr at rt. Volatiles were removed under reduced pressure and the crude residue diluted with EtOAc (25 mL), washed with water (25 mL), 10% aq. NaOH (25 mL), and brine (25 mL), then dried over $Na_2SO_4$. Volatiles were evaporated under reduced pressure and the residue purified by column chromatography on silica gel (4 g, EtOAc in hexanes, 0% to 80% EtOAc) to the amide. This material was then charged to a 5 mL µW vial equipped with a magnetic stir bar, followed by (*E*)-(2-cyclohexylvinyl)boronic acid (24 mg, 1.2 eq, 0.15 mmol), Palladium-Xphos (20 mg, 0.20 eq, 26 µmol), DMF (0.93 mL) and aq. $K_3PO_4$ (0.11 g, 0.29 mL of a 1.8 M solution, 4 eq, 0.51 mmol). The resulting mixture was sparged with argon for 3 min, then the vial was sealed, and the reaction mixture heated to 60 °C for 1 hr in a µW reactor. The mixture was diluted with EtOAc (40 mL), washed with $H_2O$ (1 x 15 mL) and brine (2 x 15 mL), then dried over $Na_2SO_4$. Volatiles were removed under reduced pressure and the crude residue purified by column chromatography on silica gel (EtOAc in hexanes gradient, 0% to 90%) to afford *tert*-butyl (2*S*,4*S*)-4-(4-((*E*)-2-cyclohexylvinyl)phenoxy)-2-(methylcarbamoyl)pyrrolidine-1-carboxylate, which was immediately dissolved in methanolic HCl (1.0 mL of a 2 M solution, 29 eq, 2.0 mmol) and incubated for 3 h, at which point volatiles were removed under reduced pressure to give (2*S*,4*S*)-4-(4-((*E*)-2-cyclohexylvinyl)phenoxy)-*N*-methylpyrrolidine-2-carboxamide hydrochloride (44 mg, 0.06 mmol, 50% over five steps).

A portion (4.7 mg, 1.2 eq, 26 μmol) of the above hydrochloride salt was treated with 3-cyclopropyl-1-ethyl-1H-pyrazole-5-carboxylic acid (4.7 mg, 1.2 eq, 26 μmol), HATU (10 mg, 1.2 eq, 26 μmol) and DIPEA (7.1 mg, 9.5 μL, 2.5 eq, 55 μmol) which had all been pre-stirred for 15 min in $CH_2Cl_2$ (3 mL). The resulting solution was stirred for 4 hr at rt, then diluted with $CH_2Cl_2$ (25 mL) and washed with water (1 x 25 mL), sat. aq. $NaHCO_3$ (1 x 25 mL) and brine (1 x 25 mL). The combined organics were dried over $Na_2SO_4$ and concentrated under reduced pressure, affording crude material which was purified by HPLC to afford **29** (8.8 mg, 18 μmol, 69%) as a white solid.

$^1$H NMR (400 MHz, $CD_3OD$) (~3:2 mixture of rotamers) δ 7.28 (d, *J* = 8.3 Hz, 2H), 6.82 (d, *J* = 8.4 Hz, 2H), 6.41 (s, 0.6H), 6.30 (d, *J* = 15.9 Hz, 1H), 6.07 (app.dd, *J* = 15.9, 7.0 Hz, 1.4H), 5.11 – 5.01 (m, 1H), 4.74 (dd, *J* = 9.4, 3.7 Hz, 0.6H), 4.58 (d, *J* = 8.8 Hz, 0.4H), 4.27 (dh, *J* = 27.8, 7.1 Hz, 2H), 4.10 (ddd, *J* = 29.6, 12.6, 5.2 Hz, 1H), 3.90 (dt, *J* = 11.7, 5.0 Hz, 1H), 2.79 (s, 2H), 2.72 (s, 1H), 2.58 (ddd, *J* = 14.0, 9.2, 5.1 Hz, 1H), 2.47 (s, 0.4H), 2.36 (dd, *J* = 10.4, 6.9 Hz, 0.6H), 2.12 (dtt, *J* = 10.8, 7.2, 3.4 Hz, 1H), 1.92 (dtt, *J* = 11.7, 8.3, 4.3 Hz, 1H), 1.77 (d, *J* = 3.1 Hz, 4H), 1.75 – 1.67 (m, 1H), 1.45 – 1.13 (m, 8H), 0.94 (dt, *J* = 9.5, 3.1 Hz, 2H), 0.76 – 0.62 (m, 2H). $^{13}$C NMR (151 MHz, DMSO) δ 171.61, 171.52, 171.25, 171.17, 161.69, 160.78, 156.23, 156.04, 152.51, 152.49, 136.25, 135.62, 134.78, 131.08, 131.03, 127.52, 127.49, 126.90, 126.87, 116.18, 116.00, 104.36, 102.35, 100.00, 75.34, 74.15, 61.29, 59.49, 59.45, 54.27, 52.59, 45.91, 45.44, 40.89, 40.54, 36.99, 34.80, 33.06, 26.30, 26.22, 26.16, 26.09, 25.99, 16.27, 16.03, 9.36, 9.31, 8.39, 8.33. MS (ESI): calculated for $C_{29}H_{39}N_4O_3$ $[M + H]^+$ 491.30; found, 491.42.

(2*S*,4*R*)-4-(4-((*E*)-2-cyclohexylvinyl)phenoxy)-1-(3-cyclopropyl-1-ethyl-1H-pyrazole-5-carbonyl)-*N*-methylpyrrolidine-2-carboxamide (**30**)

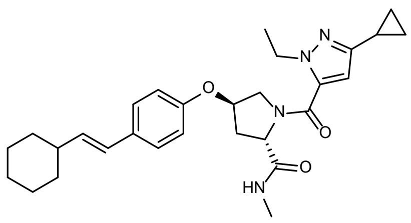

To a suspension of (2*S*,4*R*)-1-(*tert*-butoxycarbonyl)-4-(4-iodophenoxy)pyrrolidine-2-carboxylic acid (25 mg, 1 eq, 58 μmol) in $CH_2Cl_2$ (3.0 mL), was added DIPEA (22 mg, 30 μL, 3.0 eq, 0.17 mmol). The reaction was stirred for 15 min at rt before addition of methanamine hydrochloride (5.8 mg, 1.5 eq, 87 µmol) and continued stirring for 16 hr at rt. The volatiles were removed under reduced pressure and the crude residue was diluted in EtOAc (25 mL). The organic layer was washed with water (1 x 25 mL), 10% aq. NaOH (1 x 25 mL) and brine (1 x 25 mL), dried over $Na_2SO_4$ and concentrated under reduced pressure. The crude residue was purified by column chromatography (EtOAc/Hexanes gradient from 0% to 90% EtOAc, 4 g column) to afford the intermediate material *tert*-butyl (2*S*,4*R*)-4-(4-iodophenoxy)-2-(methylcarbamoyl)pyrrolidine-1-carboxylate which was then charged to a 5 mL µW vial equipped with a triangular magnetic stir bar. To this vial was charged (*E*)-(2-cyclohexylvinyl)boronic acid, Palladium-Xphos (9.5 mg, 0.20 eq, 12 µmol), DMF (0.5 mL) and 1.8 M aq. $K_3PO_4$ (77 mg, 0.20 mL, 6.0 eq, 0.36 mmol). The mixture was purged with Ar for 3 min. The vial was then sealed and heated to 60 °C for 1 hr using a µW reactor. The reaction mixture was diluted with EtOAc (40 mL) and washed with $H_2O$ (1 x 15mL) and brine (2 x 15 mL) before being dried over $Na_2SO_4$ and concentrated under reduced pressure. The crude residue was purified by column chromatography (EtOAc/Hexanes; 0% to 90% EtOAc, 4 g column) to afford *tert*-butyl (2*S*,4*R*)-4-(4-((*E*)-2-cyclohexylvinyl)phenoxy)-2-(methylcarbamoyl)pyrrolidine-1-carboxylate as a clear oil (20 mg, 47 µmol, 81% over 2 steps).

*tert*-butyl (2*S*,4*R*)-4-(4-((*E*)-2-cyclohexylvinyl)phenoxy)-2-(methylcarbamoyl)pyrrolidine-1-carboxylate (20 mg, 1 eq, 47 µmol) was dissolved in HCl (1.5 g, 1.0 mL, 2 M solution in MeOH, 43 eq, 2.0 mmol). The reaction was allowed to stand for 3 hr then volatiles were removed under reduced pressure to afford (2*S*,4*R*)-4-(4-((*E*)-2-cyclohexylvinyl)phenoxy)-*N*-methylpyrrolidine-2-carboxamide hydrochloride (quant.).

3-Cyclopropyl-1-ethyl-1H-pyrazole-5-carboxylic acid (10 mg, 1.2 eq, 56 µmol), and 2-(3H-[1,2,3]triazolo[4,5-*b*]pyridin-3-yl)-1,1,3,3-tetramethylisouronium hexafluorophosphate(V) (21 mg, 1.2 eq, 56 µmol) were dissolved in $CH_2Cl_2$ (1.0 mL) and DIPEA (15 mg, 20 µL, 2.5 eq, 0.12 mmol). The reaction was stirred for 15 min at rt before addition of (2*S*,4*R*)-4-(4-((*E*)-2-cyclohexylvinyl)phenoxy)-*N*-methylpyrrolidine-2-carboxamide hydrochloride (17 mg, 1 eq, 47 µmol) in $CH_2Cl_2$ (1.0 mL). The reaction was stirred for 16 hr at rt. The volatiles were removed under reduced pressure and the crude residue was diluted in EtOAc (25 mL). The organic layer was washed with water (1 x 25 mL), sat. aq. $NaHCO_3$ (1 x 25 mL) and brine (1 x 25 mL), dried over $Na_2SO_4$ and concentrated under reduced pressure. The crude residue was purified by preparative HPLC to afford **30** (16 mg, 33 µmol, 70%).

$^{1}$H NMR (400 MHz, $CD_3OD$) (~3:1 mixture of rotamers) δ 7.34 – 7.23 (m, 2H), 6.89 (d, *J* = 8.6 Hz, 0.5H), 6.85 – 6.78 (m, 1.5H), 6.30 (app.dd, *J* = 15.8, 8.0 Hz, 1H), 6.19 (s, 0.75H), 6.14 – 6.01 (m, 1.25H), 5.09 (s, 0.25H), 5.07 – 5.02 (m, 0.75 H), 4.70 (dd, *J* = 9.4, 7.7 Hz, 0.75H), 4.60 (t, *J* = 8.2 Hz, 0.25H), 4.34 – 4.11 (m, 2H), 4.09 – 4.02 (m, 0.25H), 3.96 – 3.81 (m, 1.75H), 2.80 (s, 2.25H), 2.65 – 2.56 (m, 1H), 2.55 (s, 0.75H), 2.25 (ddt, *J* = 13.6, 8.4, 4.2 Hz, 1H), 2.11 (dtd, *J* = 10.8, 7.2, 3.3 Hz, 1H), 1.89 (tt, *J* = 8.4, 5.0 Hz, 1H), 1.84 – 1.75 (m, 4H), 1.75 – 1.66 (m, 1H), 1.46 – 1.13 (m, 8H), 0.92 (td, *J* = 8.4, 2.5 Hz, 2H), 0.67 (dtt, *J* = 7.6, 5.2, 2.5 Hz, 2H). $^{13}$C NMR (101 MHz, $CD_3OD$) δ 172.84, 172.39, 162.48, 161.76, 155.76, 155.48, 153.95, 135.84, 135.49, 134.69, 134.61, 131.94, 131.85, 126.88, 126.49, 126.44, 115.46, 115.40, 103.39, 101.81, 75.51, 74.14, 60.61, 59.00, 54.96, 52.72, 45.25, 45.12, 41.13, 37.29, 35.04, 32.86, 25.89, 25.77, 25.03, 24.87, 14.87, 14.42, 8.24, 7.02, 7.00, 6.94. HRMS (ESI): calculated for $C_{29}H_{39}N_3O_3$ $[M + H]^+$, 491.3017; found, 491.3026.

(2*R*,4*R*)-4-(4-((*E*)-2-Cyclohexylvinyl)phenoxy)-1-(3-cyclopropyl-1-ethyl-1H-pyrazole-5-carbonyl)-*N*-methylpyrrolidine-2-carboxamide (**31**)

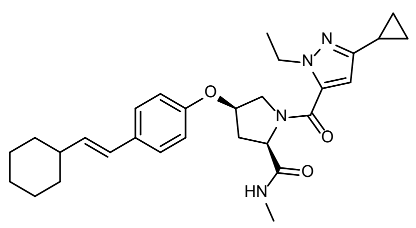

**10ci** (65 mg, 1 eq, 0.12 mmol), and 2-(3H-[1,2,3]triazolo[4,5-*b*]pyridin-3-yl)-1,1,3,3-tetramethylisouronium hexafluorophosphate(V) (53 mg, 1.2 eq, 0.14 mmol) were dissolved in $CH_2Cl_2$ (2 mL) and DIPEA (30 mg, 2.0 eq, 0.23 mmol). The reaction was stirred for 15 min at rt and methanamine hydrochloride (12 mg, 1.5 eq, 0.18 mmol) was added. The reaction was stirred for 16 hr at rt. The $CH_2Cl_2$ was removed under a stream of $N_2$, and the crude residue was diluted in EtOAc (25 mL). The organic layer was washed with water (1 x 25 mL), sat. aq. $NaHCO_3$ (1 x 25 mL) and brine (1 x 25 mL), dried over $Na_2SO_4$ and concentrated under reduced pressure. The crude residue was purified by flash chromatography (EtOAc/Hexanes gradient from 0% to 40% EtOAc, 4g column) to afford (9H-fluoren-9-yl)methyl (2*R*,4*R*)-

4-(4-iodophenoxy)-2-(methylcarbamoyl)pyrrolidine-1-carboxylate (60 mg, 0.11 mmol, 90%) as a white solid.

A 5 mL μW vial equipped with a triangular magnetic stir bar was charged with (9H-fluoren-9-yl)methyl (2*R*,4*R*)-4-(4-iodophenoxy)-2-(methylcarbamoyl)pyrrolidine-1-carboxylate (60 mg, 1 eq, 0.11 mmol), (*E*)-(2-cyclohexylvinyl)boronic acid (16 mg, 1 eq, 0.11 mmol), Palladium-Xphos (17 mg, 0.20 eq, 21 μmol), DMF (1.0 mL) and 1.8 M aq. $K_3PO_4$ (0.13 g, 0.35 mL, 6.0 eq, 0.63 mmol). The mixture was sparged with Ar for 3 min before heating to 60 °C for 1 hr using a μW reactor. The reaction mixture was diluted with EtOAc (40 mL) and washed with $H_2O$ (1 x 15mL) and brine (2 x 15 mL). The organic layer was dried over $Na_2SO_4$ and concentrated under reduced pressure. The crude residue was purified by flash chromatography (EtOAc/Hexanes gradient from 0% to 90% EtOAc, 4g column) to afford (9H-fluoren-9-yl)methyl (2*R*,4*R*)-4-(4-((*E*)-2-cyclohexylvinyl)phenoxy)-2-(methylcarbamoyl)pyrrolidine-1-carboxylate (26 mg, 47 μmol, 45%) as a light brown solid.

(9H-fluoren-9-yl)methyl (2*R*,4*R*)-4-(4-((*E*)-2-cyclohexylvinyl)phenoxy)-2-(methylcarbamoyl)pyrrolidine-1-carboxylate (26 mg, 1 eq, 47 μmol) was dissolved in DMF (0.52 mL) and piperidine (121 mg, 140 μL, 30 eq, 1.42 mmol) was added. The reaction mixture was allowed to stand for 45 min. *n*-Pentane (50 mL) was added and the volatiles were removed under reduced pressure. The process was repeated 2 additional times and the crude solid was purified by flash chromatography (0–10% MeOH [with 1% NH4OH] in $CH_2Cl_2$, 4 g silica column) to afford (2*R*,4*R*)-4-(4-((*E*)-2-cyclohexylvinyl)phenoxy)-*N*-methylpyrrolidine-2-carboxamide (12 mg, 37 μmol, 77%) as a white solid.

3-Cyclopropyl-1-ethyl-1H-pyrazole-5-carboxylic acid (16 mg, 1.2 eq, 91 μmol), and 2-(3H-[1,2,3]triazolo[4,5-*b*]pyridin-3-yl)-1,1,3,3-tetramethylisouronium hexafluorophosphate(V) (35 mg, 1.2 eq, 91 μmol) were dissolved in $CH_2Cl_2$ (2 mL) and DIPEA (25 mg, 33 μL, 2.5 eq, 0.19 mmol). The reaction was stirred for 15 min at rt and (2*R*,4*R*)-4-(4-((*E*)-2-cyclohexylvinyl)phenoxy)-*N*-methylpyrrolidine-2-carboxamide (25 mg, 1 eq, 76 μmol) was added. The reaction was stirred for 16 hr at rt. The $CH_2Cl_2$ was removed under a stream of $N_2$, and the crude residue was diluted in EtOAc (25 mL). The organic layer was washed with water (1 x 25 mL), sat. aq. $NaHCO_3$ (1 x 25 mL) and brine (1 x 25 mL), dried over

$Na_2SO_4$ and concentrated under reduced pressure. The crude residue was purified by preparative HPLC to afford **31** (10.5 mg, 58%).

$^1$H NMR (400 MHz, $CD_3OD$) (~3:2 mixture of rotamers) δ 7.28 (d, *J* = 8.3 Hz, 2H), 6.82 (d, *J* = 8.4 Hz, 2H), 6.41 (s, 0.6H), 6.30 (d, *J* = 15.9 Hz, 1H), 6.07 (app. dd, *J* = 15.9, 7.0 Hz, 1.4H), 5.12 – 4.99 (m, 1H), 4.74 (dd, *J* = 9.5, 3.7 Hz, 0.6H), 4.66 – 4.52 (m, 0.4H), 4.38 – 4.20 (m, 2H), 4.19 – 4.02 (m, 1H), 3.95 – 3.85 (m, 1H), 2.79 (s, 1.8H), 2.72 (s, 1.2H), 2.64 – 2.53 (m, 1H), 2.45 (d, *J* = 14.0 Hz, 0.4H), 2.35 (dt, *J* = 13.8, 3.6 Hz, 0.6H), 2.11 (dtt, *J* = 10.7, 6.9, 3.4 Hz, 1H), 2.00 – 1.85 (m, 1H), 1.85 – 1.75 (m, 4H), 1.75 – 1.66 (m, 1H), 1.45 – 1.15 (m, 8H), 0.99 – 0.89 (m, 2H), 0.77 – 0.62 (m, 2H). $^{13}$C NMR (101 MHz, $CD_3OD$) δ 172.54, 162.12, 161.65, 155.74, 153.80, 135.31, 134.55, 131.76, 126.80, 126.51, 115.36, 103.72, 102.01, 75.26, 73.92, 61.57, 59.59, 54.42, 52.57, 45.59, 45.37, 41.13, 36.54, 34.30, 32.87, 25.89, 25.78, 25.17, 14.85, 14.67, 8.27, 6.99. HRMS (ESI): calculated for $C_{29}H_{39}N_3O_3$ $[M + H]^+$, 491.3017; found, 491.3014.

(2*R*,4*S*)-4-(4-((*E*)-2-cyclohexylvinyl)phenoxy)-1-(3-(2,3-dihydro-1H-inden-1-yl)propanoyl)-*N*-methylpyrrolidine-2-carboxamide (**32**)

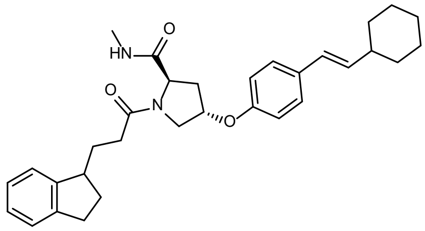

A solution of (2*R*,4*S*)-4-(4-iodophenoxy)-2-(methoxycarbonyl)pyrrolidin-1-ium 2,2,2-trifluoroacetate (80 mg, 1 eq, 0.17 mmol) and di-*tert*-butyl dicarbonate (45 mg, 1.2 eq, 0.21 mmol) in $CH_2Cl_2$ (1 mL) was added $NEt_3$ (21 mg, 1.2 eq, 0.21 mmol), and was stirred at rt for 16 hr. The reaction was concentrated, and the crude residue was purified by flash chromatography (0–70% EtOAc in hexanes, 12 g silica column) to afford 1-(*tert*-butyl) 2-methyl (2*R*,4*S*)-4-(4-iodophenoxy)pyrrolidine-1,2-dicarboxylate (52 mg, 0.12 mmol, 68%).

1-(*tert*-butyl) 2-methyl (2*R*,4*S*)-4-(4-iodophenoxy)pyrrolidine-1,2-dicarboxylate (50 mg, 1 eq, 0.11 mmol) was dissolved in THF:$H_2O$ (3 mL) and LiOH (40 mg, 15 eq, 1.7 mmol) was added. The reaction mixture was stirred vigorously for 16 hr, and the reaction mixture was diluted with $H_2O$ (15 mL). The aqueous layer was acidified with 1 N HCl to pH ~2 and extracted with $CH_2Cl_2$ (3 x 20 mL). The combined organics were dried over $Na_2SO_4$ and concentrated under reduced pressure to afford (2*R*,4*S*)-1-(*tert*-butoxycarbonyl)-4-(4-iodophenoxy)pyrrolidine-2-carboxylic acid (44 mg, 0.10 mmol, 93%) as a white solid.

(2*R*,4*S*)-1-(*tert*-butoxycarbonyl)-4-(4-iodophenoxy)pyrrolidine-2-carboxylic acid (43 mg, 1 eq, 99 µmol), and 2-(3H-[1,2,3]triazolo[4,5-*b*]pyridin-3-yl)-1,1,3,3-tetramethylisouronium hexafluorophosphate(V) (45 mg, 1.2 eq, 0.12 mmol) were dissolved in DMF (0.3 mL) and DIPEA (38 mg, 52 µL, 3.0 eq, 0.30 mmol). The reaction was stirred for 15 min at rt and methanamine hydrochloride (10 mg, 1.5 eq, 0.15 mmol) was added. The reaction was stirred for 16 hr at rt. The DMF was removed under a stream of $N_2$, and the crude residue was diluted in EtOAc (25 mL). The organic layer was washed with water (1 x 25 mL), sat. aq. $NaHCO_3$ (1 x 25 mL) and brine (1 x 25 mL), dried over $Na_2SO_4$ and concentrated under reduced pressure. The crude was purified by flash chromatography (0-40% EtOAc in hexanes, 4 g column) to afford *tert*-butyl (2*R*,4*S*)-4-(4-iodophenoxy)-2-(methylcarbamoyl)pyrrolidine-1-carboxylate (20 mg, 45 µmol, 45%).

A 5 mL µW vial equipped with a triangular magnetic stir bar was charged with *tert*-butyl (2*R*,4*S*)-4-(4-iodophenoxy)-2-(methylcarbamoyl)pyrrolidine-1-carboxylate (20 mg, 1 eq, 45 µmol), (*E*)-(2-cyclohexylvinyl)boronic acid (6.9 mg, 1 eq, 45 µmol), Palladium-Xphos (7.1 mg, 0.20 eq, 9.0 µmol), DMF (1.0 mL) and 1.8 M aq. $K_3PO_4$ (57 mg, 0.15 mL, 6.0 eq, 0.27 mmol). The mixture was purged with Ar for 3 min. The vial was then sealed and heated to 60 °C for 1 hr using a µW reactor. The reaction mixture was diluted with EtOAc (40 mL) and washed with $H_2O$ (1 x 15mL) and brine (2 x 15 mL). dried over $Na_2SO_4$ and concentrated under reduced pressure. The crude residue was purified by column chromatography (0–90% EtOAc in hexanes, 4 g column) to afford *tert*-butyl (2*R*,4*S*)-4-(4-((*E*)-2-cyclohexylvinyl)phenoxy)-2-(methylcarbamoyl)pyrrolidine-1-carboxylate (18 mg, 42 µmol, 94%) as a white solid.

*tert*-Butyl (2*R*,4*S*)-4-(4-((*E*)-2-cyclohexylvinyl)phenoxy)-2-(methylcarbamoyl)pyrrolidine-1-carboxylate (20 mg, 1 eq, 47 µmol) was dissolved in HCl (1.5 g, 1.0 mL, 2 M solution in MeOH, 43 eq, 2.0 mmol). The reaction was stood at rt for 3 hr and volatiles were removed under reduced pressure to afford (2*R*,4*S*)-4-(4-((*E*)-2-cyclohexylvinyl)phenoxy)-*N*-methylpyrrolidine-2-carboxamide (quant.)

3-(2,3-Dihydro-1H-inden-1-yl)propanoic acid (4.0 mg, 1 eq, 21 µmol), and 2-(3H-[1,2,3]triazolo[4,5-*b*]pyridin-3-yl)-1,1,3,3-tetramethylisouronium hexafluorophosphate(V) (9.6 mg, 1.2 eq, 25 µmol) were dissolved in $CH_2Cl_2$ (3 mL) and *N*-ethyl-*N*-isopropylpropan-2-amine (6.8 mg, 9.1 µL, 2.5 eq, 53 µmol). The reaction was stirred for 15 min at rt and (2*R*,4*S*)-4-(4-((*E*)-2-cyclohexylvinyl)phenoxy)-*N*-methylpyrrolidine-2-carboxamide hydrochloride (7.7 mg, 1 eq, 21 µmol) was added. The reaction was stirred for 16 hr at rt. The reaction was diluted with $CH_2Cl_2$ (25 mL). The organic layer was washed with water (1 x 25 mL), sat. aq. $NaHCO_3$ (1 x 25 mL) and brine (1 x 25 mL), dried over $Na_2SO_4$ and concentrated under reduced pressure. The crude residue was purified by preparative HPLC to afford **32** (5.1 mg, 10 µmol, 49%) as a white solid.

$^1$H NMR (400 MHz, $CD_3OD$) (~2:1 mixture of rotamers) δ 7.33 – 7.24 (m, 2H), 7.23 – 7.05 (m, 4H), 6.88 – 6.81 (m, 2H), 6.31 (dd, *J* = 15.9, 9.5 Hz, 1H), 6.13 – 6.03 (m, 1H), 5.09 (td, *J* = 3.9, 1.9 Hz, 0.8H), 5.03 – 4.99 (m, 0.2H), 4.53 (dt, *J* = 19.0, 8.1 Hz, 1H), 4.05 (dt, *J* = 12.8, 1.9 Hz, 0.2H), 3.92 – 3.80 (m, 1.6H), 3.75 – 3.68 (m, 0.2H), 3.20 – 3.03 (m, 1H), 3.00 – 2.71 (m, 5H), 2.69 – 2.00 (m, 7H), 1.87 – 1.76 (m, 4H), 1.76 – 1.54 (m, 3H), 1.45 – 1.14 (m, 5H). $^{13}$C NMR (101 MHz, $CD_3OD$) δ 173.52, 173.49, 155.70, 155.67, 146.29, 143.60, 143.54, 134.65, 134.62, 134.60, 131.85, 131.82, 126.94, 126.90, 126.83, 126.53, 126.51, 126.06, 125.78, 125.74, 123.99, 123.96, 123.19, 123.11, 115.48, 115.36, 75.59, 75.53, 58.96, 52.71, 44.09, 44.03, 41.15, 35.20, 35.15, 32.89, 32.87, 32.24, 32.07, 31.39, 31.34, 30.72, 30.68, 29.81, 25.89, 25.77, 24.96. HRMS (ESI): calculated for $C_{32}H_{41}N_2O_3$ $[M + H]^+$, 501.3122; found, 501.3127.

(2*R*,4*S*)-4-(4-((*E*)-2-cyclohexylvinyl)phenoxy)-1-(3-cyclopropyl-1-ethyl-1H-pyrazole-5-carbonyl)-*N*-methylpyrrolidine-2-carboxamide (**33**)

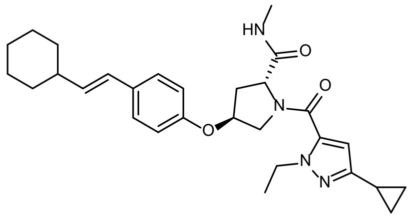

3-Cyclopropyl-1-ethyl-1H-pyrazole-5-carboxylic acid (4.5 mg, 1.2 eq, 25 μmol), and 2-(3H-[1,2,3]triazolo[4,5-*b*]pyridin-3-yl)-1,1,3,3-tetramethylisouronium hexafluorophosphate(V) (9.6 mg, 1.2 eq, 25 μmol) were dissolved in DMF (0.3 mL) and DIPEA (6.8 mg, 9.1 μL, 2.5 eq, 53 μmol). The reaction was stirred for 15 min at rt before the addition of (2*R*,4*S*)-4-(4-((*E*)-2-cyclohexylvinyl)phenoxy)-*N*-methylpyrrolidine-2-carboxamide hydrochloride (7.7 mg, 1 eq, 21 μmol) [*cf.* preparation of **32**]. The reaction was stirred for 16 hr at rt. The DMF was removed under a stream of $N_2$, and the crude residue was diluted in EtOAc (25 mL). The organic layer was washed with water (1 x 25 mL), sat. aq. $NaHCO_3$ (1 x 25 mL) and brine (1 x 25 mL), dried over $Na_2SO_4$ and concentrated under reduced pressure. The crude residue was purified by preparative HPLC to afford **33** (5.08 mg, 10.4 μmol, 49%) as a white solid.

$^1$H NMR (400 MHz, $CD_3OD$) (~3:1 mixture of rotamers) δ 7.31 (d, *J* = 8.6 Hz, 0.5H), 7.30 – 7.25 (m, 1.5H), 6.90 (d, *J* = 8.6 Hz, 0.5H), 6.86 – 6.78 (m, 1.5H), 6.31 (dd, *J* = 15.9, 7.9 Hz, 1H), 6.19 (s, 0.75H), 6.14 – 6.02 (m, 1.25H), 5.09 (t, *J* = 3.8 Hz, 0.25H), 5.05 (t, *J* = 3.8 Hz, 0.75H), 4.70 (dd, *J* = 9.4, 7.7 Hz, 0.75H), 4.60 (t, *J* = 8.2 Hz, 0.25H), 4.35 – 4.11 (m, 2H), 4.11 – 4.03 (m, 0.25H), 3.97 – 3.80 (m, 1.75H), 2.80 (s, 2.25H), 2.66 – 2.56 (m, 1H), 2.55 (s, 0.75H), 2.31 – 2.20 (m, 1H), 2.11 (dtd, *J* = 10.8, 7.3, 3.4 Hz, 1H), 1.98 – 1.85 (m, 1H), 1.85 – 1.75 (m, 4H), 1.75 – 1.67 (m, 1H), 1.45 – 1.17 (m, 8H), 0.97 – 0.85 (m, 2H), 0.73 – 0.61 (m, 2H). $^{13}$C NMR (101 MHz, $CD_3OD$) δ 172.84, 161.78, 155.48, 153.96, 135.50, 134.69, 131.94, 126.87, 126.44, 115.46, 103.37, 101.98, 75.51, 59.00, 54.95, 52.38, 41.14, 37.28, 35.05, 32.86, 25.88, 25.77, 25.01, 24.86, 14.86, 14.52, 8.22, 7.00, 6.98. HRMS (ESI): calculated for $C_{29}H_{39}N_3O_3$ [M + H]$^+$, 491.3017; found, 491.3024.

(6*R*,7*S*,8*S*)-4-((3-Chloro-4-fluorophenyl)sulfonyl)-7-(4-((*E*)-2-cyclohexylvinyl)phenyl)-8-(hydroxymethyl)-1,4-diazabicyclo[4.2.0]octan-2-one (**34**)

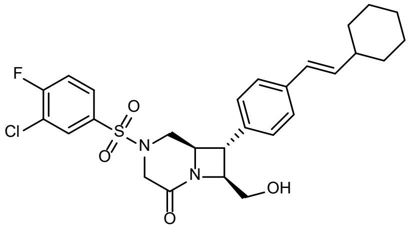

A 1-dram vial equipped with a magnetic stir bar was charged with (6*R*,7*S*,8*S*)-7-(4-bromophenyl)-8-(hydroxymethyl)-4-((2-nitrophenyl)sulfonyl)-1,4-diazabicyclo[4.2.0]octan-2-one (60 mg, 1 eq, 0.12 mmol). $CH_2Cl_2$ (1 mL) was added to give a yellow-tinted solution. 2,6-Lutidine (0.028 mL, 2.0 eq, 0.24 mmol) was added, followed by TBSOTf (0.042 mL, 1.5 eq, 0.18 mmol). The vial was blown out with nitrogen, tightly capped, stirred 10 hr, at which time the reaction was quenched by the addition of MeOH (0.1 mL) and allowed to stir for 30 min. Volatiles were evaporated under reduced pressure to reveal a viscous, opaque yellow oil which was dried under high vacuum overnight. To this crude material were added $Cs_2CO_3$ (0.20 g, 5.0 eq, 0.60 mmol) and DMF (1 mL), followed by 4-methoxythiophenol (0.059 mL, 4.0 eq, 0.48 mmol). The resulting yellow mixture was stirred for 2 hr, then transferred to a 100 mL round bottom flask (rinsing the reaction flask with ethyl acetate). Volatiles were removed under reduced pressure; DMF was removed by co-evaporation with heptane, affording a yellow residue, which was suspended in $CH_2Cl_2$ and washed with water. The organic portion was separated, dried over $Na_2SO_4$ and purified directly by flash chromatography (0–100% MeOH [containing 1% $NH_4OH$] in $CH_2Cl_2$, 4 g silico column), giving (6*R*,7*S*,8*S*)-7-(4-bromophenyl)-8-(((tert-butyldimethylsilyl)oxy)methyl)-1,4-diazabicyclo[4.2.0]octan-2-one (33 mg, 64% over two steps) as a pale yellow oil.

A 20 mL glass scintillation vial containing a magnetic stir bar and (6*R*,7*S*,8*S*)-7-(4-bromophenyl)-8-(((tert-butyldimethylsilyl)oxy)methyl)-1,4-diazabicyclo[4.2.0]octan-2-one (29 mg, 1.00 eq, 68 μmol) was charged with $CH_2Cl_2$ (0.75 mL), giving a colorless solution. 4-DMAP (0.075 mL of a 0.1 M solution in $CH_2Cl_2$, 0.11 eq, 7.5 μmol) and *N*,*N*-diisopropylethylamine (0.82 mL of a 0.1 M solution in $CH_2Cl_2$, 1.20 eq, 82 μmol) were then added, followed by 3-chloro-4-fluorobenzenesulfonyl chloride (0.75 mL of a 0.1 M solution in $CH_2Cl_2$, 1.10 eq, 75 μmol) positive $N_2$ pressure. The reaction vial was then capped, and the colorless reaction solution stirred for 2 hr. The reaction mixture was purified directly by flash chromatography (0–10% MeOH [containing 1% $NH_4OH$] in $CH_2Cl_2$, 4 g silica column) giving (6*R*,7*S*,8*S*)-7-(4-bromophenyl)-8-(((*tert*-butyldimethylsilyl)oxy)methyl)-4-((3-chloro-4-fluorophenyl)sulfonyl)-1,4-diazabicyclo[4.2.0]octan-2-one (40 mg, 95%) as a colorless solid.

A 2 mL μW vial containing a magnetic stir bar was charged with (*E*)-(2-cyclohexylvinyl)boronic acid (15 mg, 1.50 eq, 97 μmol) and Pd-Xphos-G2 (10 mg, 0.20 eq, 13 μmol). The vial was capped with a rubber septum and (6*R*,7*S*,8*S*)-7-(4-bromophenyl)-8-(((*tert*-butyldimethylsilyl)oxy)methyl)-4-((3-chloro-4-fluorophenyl)sulfonyl)-1,4-diazabicyclo[4.2.0]octan-2-one (40 mg, 1.00 eq, 65 µmol) was added as a solution in DMF (0.5 mL + 2 x 0.5 mL washes). $K_3PO_4$ (0.24 mL of a 1.6 M aq. solution, 6.00 eq, 0.39 mmol) was added, to afford an off-white cloudy suspension. The vial was then evacuated and backfilled with $N_2$ (5x), then heated to 60 °C for 1 hr under µW irradiation. The reaction mixture was concentrated under reduced pressure, then residual DMF was removed by azeotropic drying with heptane. The resulting residue was purified by flash chromatography (EtOAc in hexanes, 4 g silica column) to give (7*S*,8*S*)-8-(((*tert*-butyldimethylsilyl)oxy)methyl)-4-((3-chloro-4-fluorophenyl)sulfonyl)-7-(4-((*E*)-2-cyclohexylvinyl)phenyl)-1,4-diazabicyclo[4.2.0]octan-2-one (20 mg, 31 µmol, 48%).

This resulting material was dissolved in DMF (250 µL), followed by addition of water (3 µL, 5 eq, 155 µmol,). TAS-F (21 mg, 2.5 eq, 77.5 µmol) was added in one portion and the resulting solution was stirred at 50 °C for 40 min. Upon completion, the reaction mixture was purified by preparative HPLC to afford **34** (1.5 mg, 2.8 µmol, 9%).

$^1$H NMR (400 MHz, $CDCl_3$) δ 7.88 (dd, *J* = 6.6, 2.3 Hz, 1H), 7.70 (ddd, *J* = 8.6, 4.3, 2.3 Hz, 1H), 7.38 – 7.32 (m, 3H), 7.15 (d, *J* = 8.0 Hz, 2H), 6.33 (d, *J* = 15.9 Hz, 1H), 6.20 (dd, *J* = 16.0, 6.8 Hz, 1H), 4.82 – 4.66 (m, 2H), 4.25 (d, *J* = 16.8 Hz, 1H), 4.17 (dd, *J* = 11.6, 4.5 Hz, 1H), 4.10 – 4.00 (m, 1H), 3.97 (dd, *J* = 9.7, 3.4 Hz, 1H), 3.79 – 3.67 (m, 1H), 3.48 (t, *J* = 7.6 Hz, 1H), 3.41 (d, *J* = 16.8 Hz, 1H), 2.70 (t, *J* = 10.9 Hz, 1H), 2.19 – 2.07 (m, 1H), 1.85 – 1.74 (m, 4H), 1.69 (d, *J* = 12.6 Hz, 1H), 1.39 – 1.13 (m, 6H). $^{13}$C NMR (101 MHz, $CDCl_3$) δ 164.39, 138.28, 137.98, 134.74, 130.35, 127.98, 127.89, 127.10, 126.62, 126.32, 117.98, 117.76, 77.21, 74.73, 65.33, 63.08, 48.43, 47.59, 44.72, 41.17, 32.88, 26.15, 26.01. LRMS (ESI): calculated for $C_{27}H_{31}ClFN_2O_4S$ $[M + H]^+$, 533.17; found 533.35.

*N*-(((4*R*,5*R*)-8-((*E*)-2-Cyclohexylvinyl)-2-((*R*)-1-hydroxypropan-2-yl)-4-methyl-1,1-dioxido-2,3,4,5-tetrahydrobenzo[*b*][1,4,5]oxathiazocin-5-yl)methyl)-*N*-methyl-2-phenylcyclopent-1-ene-1-carboxamide (**35**)

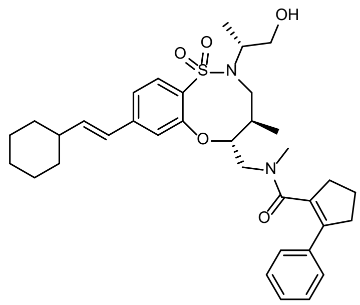

A 20 mL scintillation vial equipped with a magnetic stir bar was charged with (9H-fluoren-9-yl)methyl (((4*R*,5*R*)-8-bromo-2-((*R*)-1-hydroxypropan-2-yl)-4-methyl-1,1-dioxido-2,3,4,5-tetrahydrobenzo[*b*][1,4,5]oxathiazocin-5-yl)methyl)(methyl)carbamate (65.3 mg, 1.00 eq, 104 µmol), dry DMF (3.5 mL) and dry piperidine (0.16 g, 0.18 mL, 18 eq, 1.8 mmol) at rt. The clear, colorless solution was stirred for 20 min. It was then concentrated under reduced pressure to afford a white solid. The material was absorbed on silica gel and purified by flash chromatography (4–8% MeOH [containing 1% $NH_4OH$] in $CH_2Cl_2$) affording (4*R*,5*R*)-8-bromo-2-((*R*)-1-hydroxypropan-2-yl)-4-methyl-5-((methylamino)methyl)-2,3,4,5-tetrahydrobenzo[*b*][1,4,5]oxathiazocine 1,1-dioxide as a colorless oil (28.2 mg, 69.2 µmol, 67%).

A 2-dram scintillation vial equipped with a magnetic stir bar was charged with (4*R*,5*R*)-8-bromo-2-((*R*)-1-hydroxypropan-2-yl)-4-methyl-5-((methylamino)methyl)-2,3,4,5-tetrahydrobenzo[*b*][1,4,5]oxathiazocine 1,1-dioxide (28.2 mg, 1.00 eq, 69.2 µmol), dry DMF (0.69 mL), DIPEA (25 µL, 2.1 eq, 0.15 mmol), 2-phenylcyclopent-1-ene-1-carboxylic acid (95% wt, 15.5 mg, 1.13 eq, 78.2 µmol) and PyBOP (39.6 mg, 1.1 eq, 76.2 µmol) at rt. The clear, faint brown solution was stirred for 25 min. EtOAc and water were added, and the aqueous phase was extracted twice with EtOAc. The organic layers were washed with water and were then combined, dried over $Na_2SO_4$ and concentrated under reduced pressure. The crude material was purified by flash chromatography (0–8% MeOH in $CH_2Cl_2$), affording *N*-(((4*R*,5*R*)-8-bromo-2-((*R*)-1-hydroxypropan-2-yl)-4-methyl-1,1-dioxido-2,3,4,5-tetrahydrobenzo[*b*][1,4,5]oxathiazocin-5-yl)methyl)-*N*-methyl-2-phenylcyclopent-1-ene-1-carboxamide (35.3 mg, 61.1 µmol, 88%) as a colorless oil.

A 5 mL cone-shaped µW vial equipped with a magnetic stir bar was charged with *N*-(((4*R*,5*R*)-8-bromo-2-((*R*)-1-hydroxypropan-2-yl)-4-methyl-1,1-dioxido-2,3,4,5-tetrahydrobenzo[*b*][1,4,5]oxathiazocin-5-yl)methyl)-*N*-methyl-2-phenylcyclopent-1-ene-1-carboxamide (35.3 mg, 1.00 eq, 61.1 µmol) in dry DMF

(0.61 mL) at rt. (*E*)-(2-Cyclohexylvinyl)boronic acid (98% wt, 14.4 mg, 1.5 eq, 91.7 μmol), XPhos-Pd-G2 (9.6 mg, 20 mol%, 12 μmol) and aq. $K_3PO_4$ (1.8 M, 0.20 mL, 5.9 eq) were successively added and the biphasic mixture was sparged with Ar for 3 min. The vial was then sealed, and the reaction mixture was heated to 60 °C for 3 hr using μW irradiation. More XPhos-Pd-G2 (4.8 mg, 10 mol%, 6.1 μmol) and aq. $K_3PO_4$ (1.8 M, 0.10 mL, 0.18 mmol, 3.0 eq) were added after 2 hr. EtOAc was added, and the mixture was filtered over cotton. The filtrate was concentrated under reduced pressure and the obtained crude material was purified by flash chromatography (20–80% EtOAc in hexanes) affording the product as a yellow oil. The material was further purified by preparative HPLC to afford **35** (19.6 mg, 32.3 μmol, 53%) as a white foam.

$^1$H NMR (400 MHz, $CDCl_3$) (~3:2 mixture of rotamers) δ 7.73 (dd, *J* = 8.3, 6.1 Hz, 1H), 7.31 – 7.13 (m, 5H), 7.08 (ddd, *J* = 13.5, 8.2, 1.7 Hz, 1H), 6.95 (d, *J* = 1.7 Hz, 0.6H), 6.86 (d, *J* = 1.7 Hz, 0.4H), 6.36 (dd, *J* = 16.0, 6.8 Hz, 0.4H), 6.28 – 6.18 (m, 1.6H), 4.50 (td, *J* = 8.7, 2.6 Hz, 0.6H), 4.28 – 4.08 (m, 1H), 3.80 – 3.67 (m, 1H), 3.66 – 3.38 (m, 4H), 3.34 (dd, *J* = 15.8, 4.9 Hz, 0.4H), 3.23 (dd, *J* = 14.1, 2.6 Hz, 0.6H), 2.99 – 2.36 (m, 8.75H), 2.23 – 1.93 (m, 4.25H), 1.83 – 1.62 (m, 5.2H), 1.34 – 1.07 (m, 8.8H), 0.91 (d, *J* = 6.9 Hz, 1.75H), 0.61 (d, *J* = 6.9 Hz, 1.25H). $^{13}$C NMR (101 MHz, $CDCl_3$) δ 172.08, 171.69, 155.99, 155.34, 144.96, 144.30, 141.80, 141.15, 140.15, 139.88, 136.22, 136.16, 133.02, 132.91, 130.82, 130.51, 130.02, 129.92, 128.58, 127.98, 127.91, 126.78, 126.72, 125.66, 125.32, 122.27, 121.40, 120.77, 120.73, 85.10, 84.69, 77.36, 64.96, 64.86, 57.47, 57.34, 53.96, 49.76, 49.50, 49.33, 41.38, 37.42, 37.13, 36.92, 36.65, 36.44, 36.34, 36.17, 33.21, 32.79, 32.76, 32.67, 32.65, 26.18, 26.06, 22.61, 22.58, 15.97, 15.83, 15.75, 15.31. HRMS (ESI): calculated for $C_{35}H_{47}N_2O_5S$ $[M + H]^+$, 607.3200; found, 607.3219.

*N*-(((4*S*,5*S*)-8-((*E*)-2-Cyclohexylvinyl)-2-((*R*)-1-hydroxypropan-2-yl)-4-methyl-1,1-dioxido-2,3,4,5-tetrahydrobenzo[*b*][1,4,5]oxathiazocin-5-yl)methyl)-*N*-methyl-2-(3-methylthiophen-2-yl)cyclopropane-1-carboxamide (**36**)

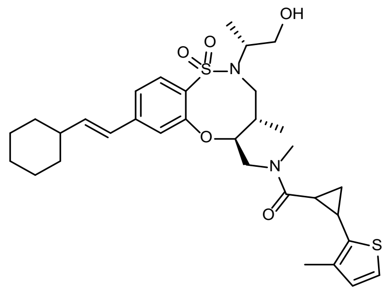

A 2 dram scintillation vial equipped with a magnetic stir bar was charged with (9H-fluoren-9-yl)methyl (((4*S*,5*S*)-8-bromo-2-((*R*)-1-hydroxypropan-2-yl)-4-methyl-1,1-dioxido-2,3,4,5-tetrahydrobenzo[*b*][1,4,5]oxathiazocin-5-yl)methyl)(methyl)carbamate (52.1 mg, 1.00 eq, 82.8 µmol), dry DMF (2.8 mL) and dry piperidine (0.14 mL, 17 eq, 1.4 mmol) at rt. The colorless solution was stirred for 40 min. It was then concentrated under reduced pressure and the obtained white solid was purified by flash chromatography (0–10% MeOH [containing 1% $NH_4OH$] in $CH_2Cl_2$) to afford (4*S*,5*S*)-8-bromo-2-((*R*)-1-hydroxypropan-2-yl)-4-methyl-5-((methylamino)methyl)-2,3,4,5-tetrahydrobenzo[*b*][1,4,5]oxathiazocine 1,1-dioxide (31.0 mg, 76.1 µmol, 92%) as a colorless oil.

A 2 dram scintillation vial equipped with a magnetic stir bar was charged with (4*S*,5*S*)-8-bromo-2-((*R*)-1-hydroxypropan-2-yl)-4-methyl-5-((methylamino)methyl)-2,3,4,5-tetrahydrobenzo[*b*][1,4,5]oxathiazocine 1,1-dioxide (31.0 mg, 1 eq, 76.1 µmol), *racemic* *trans*-2-(3-methylthiophen-2-yl)cyclopropane-1-carboxylic acid (0.22 M in DMF, 0.45 mL, 1.3 eq, 99 µmol), DIPEA (21 mg, 28 µL, 2.1 eq, 0.16 mmol) and PyBOP (0.22 M in DMF, 0.45 mL, 0.22 molar, 1.3 eq, 99 µmol) at r.t. The light brown solution was stirred for 50 min. EtOAc and water (each 30 mL) were added and the aqueous phase was extracted twice with EtOAc. The organic layers were washed two more times with water. They were combined, dried over $MgSO_4$ and concentrated under reduced pressure. Purification of the crude material by flash chromatography (0–10% MeOH [containing 1% $NH_4OH$] in $CH_2Cl_2$) afforded *N*-(((4*S*,5*S*)-8-bromo-2-((*R*)-1-hydroxypropan-2-yl)-4-methyl-1,1-dioxido-2,3,4,5-tetrahydrobenzo[*b*][1,4,5]oxathiazocin-5-yl)methyl)-*N*-methyl-2-(3-methylthiophen-2-yl)cyclopropane-1-carboxamide (34 mg, 0.060 mmol, 78%) as a mixture of both diastereomers and rotamers.

A 5 mL cone-shaped µW vial equipped with a magnetic stir bar was charged with *N*-(((4*S*,5*S*)-8-bromo-2-((*R*)-1-hydroxypropan-2-yl)-4-methyl-1,1-dioxido-2,3,4,5-tetrahydrobenzo[*b*][1,4,5]oxathiazocin-5-

yl)methyl)-*N*-methyl-2-(3-methylthiophen-2-yl)cyclopropane-1-carboxamide (34 mg, 1.0 eq, 59 µmol) in dry DMF (0.59 mL), (*E*)-(2-cyclohexylvinyl)boronic acid (14 mg, 1.5 eq, 89 µmol), XPhos-Pd-G2 (9.4 mg, 20 mol%, 12 µmol) and aq. $K_3PO_4$ (1.8 M, 0.10 mL, 3.0 eq, 0.18 mmol) at r.t. The reaction mixture was purged with Argon for two min and the vial was sealed. The mixture was heated to 60 °C for 1 hour using µW irradiation. More aq. $K_3PO_4$ (0.26 g, 0.10 mL, 1.8 molar, 3.0 eq, 0.18 mmol) was added after 30 min. EtOAc was added and the mixture was filtered over Celite. The filtrate was concentrated under reduced pressure and the crude material was purified by flash chromatography (25–75% EtOAc in hexanes) to afford a yellow oil, that was further purified by preparative HPLC to **36** (19.6 mg, 32.6 µmol, 55%) as a white solid and as a mixture of rotamers and diastereomers (ratio approximately 1:1:1:1, according to analysis by $^{1}$H NMR spectroscopy).

$^{1}$H NMR (400 MHz, $CD_3OD$) δ 7.72 (dt, *J* = 8.2, 2.1 Hz, 1H), 7.24 – 7.09 (m, 1H), 7.07 – 6.90 (m, 2H), 6.75 (dd, *J* = 12.6, 5.1 Hz, 1H), 6.43 – 6.14 (m, 2H), 4.72 – 4.63 (m, 0.5H), 4.58 (ddd, *J* = 10.7, 9.0, 2.9 Hz, 0.25H), 4.44 (ddd, *J* = 11.3, 8.9, 2.8 Hz, 0.25H), 4.28 (td, *J* = 15.0, 10.4 Hz, 0.5H), 4.18 – 4.05 (m, 0.5H), 3.85 – 3.49 (m, 5.75H), 3.46 (dd, *J* = 14.1, 2.8 Hz, 0.25H), 3.36 (dd, *J* = 13.9, 2.7 Hz, 0.25H), 3.22 (s, 0.75H), 3.08 (s, 0.75H), 2.86 (s, 0.75H), 2.77 (s, 1H), 2.58 – 2.38 (m, 1.25H), 2.35 – 2.03 (m, 5.25H), 2.02 – 1.92 (m, 0.5H), 1.86 – 1.63 (m, 5.5H), 1.50 – 1.03 (m, 10H), 0.95 – 0.90 (m, 2.25H), 0.86 (d, *J* = 6.9 Hz, 0.75H). $^{13}$C NMR (101 MHz, methanol-d$^{4}$) δ 174.66, 174.60, 174.46, 174.19, 157.18, 157.16, 157.01, 156.66, 145.78, 145.58, 145.37, 145.34, 142.20, 142.17, 141.70, 141.62, 138.53, 138.43, 138.37, 138.27, 136.32, 136.28, 136.21, 133.99, 133.64, 133.59, 133.51, 131.40, 131.31, 131.26, 130.47, 130.41, 130.21, 126.95, 126.91, 126.75, 126.66, 123.22, 123.15, 122.18, 122.16, 122.07, 122.00, 121.89, 121.66, 121.38, 86.34, 85.22, 85.13, 84.66, 65.73, 65.64, 65.52, 58.86, 58.51, 58.46, 58.38, 54.90, 54.61, 53.32, 51.28, 51.19, 51.04, 50.98, 49.71, 42.65, 42.58, 42.55, 42.42, 38.19, 38.02, 37.98, 37.92, 37.23, 36.69, 34.88, 34.35, 33.89, 33.81, 33.67, 33.64, 27.20, 27.14, 27.07, 27.02, 26.97, 24.69, 24.22, 23.96, 23.66, 21.30, 20.80, 20.40, 20.08, 18.62, 17.74, 17.66, 17.57, 15.92, 15.71, 15.27, 15.22, 15.10, 14.83, 14.10, 13.99, 13.88, 13.78. HRMS (ESI): calculated for $C_{32}H_{45}N_2O_5S_2$ $[M + H]^+$, 601.2764; found, 601.2778.

*N*-(((4*S*,5*S*)-8-(2-Chloro-5-(hydroxymethyl)phenyl)-2-((*R*)-1-hydroxypropan-2-yl)-4-methyl-1,1-dioxido-2,3,4,5-tetrahydrobenzo[*b*][1,4,5]oxathiazocin-5-yl)methyl)-6-ethyl-*N*-methyl-2-oxo-1,2-dihydropyridine-4-carboxamide (**37**)

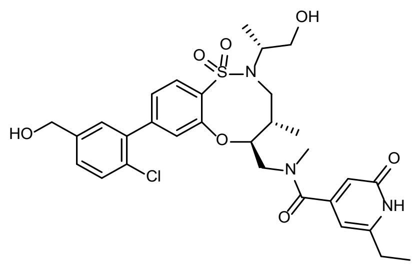

A 2 dram scintillation vial equipped with a magnetic stir bar was charged with (4*S*,5*S*)-8-bromo-2-((*R*)-1-hydroxypropan-2-yl)-4-methyl-5-((methylamino)methyl)-2,3,4,5-tetrahydrobenzo[*b*][1,4,5]oxathiazocine 1,1-dioxide [*cf*. synthesis of **36**] (30.8 mg, 1 eq, 75.6 µmol) at r.t. Dry DMF (0.76 mL), DIPEA (28 µL, 2.1 eq, 0.16 mmol), 6-ethyl-2-oxo-1,2-dihydropyridine-4-carboxylic acid (16.4 mg, 1.3 eq, 98.3 µmol) and PyBOP (51.2 mg, 1.3 eq, 98.3 µmol) were successively added and the obtained brown solution was stirred for 50 min at rt. EtOAc and water–brine (9:1) (each 30 mL) were added and the aqueous phase was extracted twice with EtOAc. The organic layers were washed two more times with water:brine (9:1). They were combined, dried over $Na_2SO_4$ and concentrated under reduced pressure to afford a yellow oil. The crude material was purified by flash chromatography (0–10% MeOH in $CH_2Cl_2$), then further purified by preparative HPLC to afford *N*-(((4*S*,5*S*)-8-bromo-2-((*R*)-1-hydroxypropan-2-yl)-4-methyl-1,1-dioxido-2,3,4,5-tetrahydrobenzo[*b*][1,4,5]oxathiazocin-5-yl)methyl)-6-ethyl-*N*-methyl-2-oxo-1,2-dihydropyridine-4-carboxamide (22.8 mg, 41.0 µmol, 54%) as a white foam.

A 5 mL cone-shaped µW vial equipped with a magnetic stir bar was charged with *N*-(((4*S*,5*S*)-8-bromo-2-((*R*)-1-hydroxypropan-2-yl)-4-methyl-1,1-dioxido-2,3,4,5-tetrahydrobenzo[*b*][1,4,5]oxathiazocin-5-yl)methyl)-6-ethyl-*N*-methyl-2-oxo-1,2-dihydropyridine-4-carboxamide (22.8 mg, 1.00 eq, 41.0 µmol) in dry DMF (0.41 mL) at rt. (2-chloro-5-(hydroxymethyl)phenyl)boronic acid (11.5 mg, 1.5 eq, 61.5 µmol), XPhos-Pd-G2 (6.5 mg, 20 mol%, 8.2 µmol) and aq. $K_3PO_4$ (1.8 M, 0.14 mL, 6.2 eq, 0.25 mmol) were successively added. The mixture was sparged with Ar for 3 min, the vial was sealed, and the reaction mixture was heated to 60 °C for 4 hr using µW irradiation. EtOAc was added and the mixture was filtered over cotton. The filtrate was concentrated under reduced pressure and the crude material was purified by flash chromatography (0–10% MeOH in $CH_2Cl_2$), then further purified by preparative HPLC to afford **37** (7.4 mg, 12 μmol, 29%) as a white foam.

$^1$H NMR (400 MHz, $CD_3OD$) (1:1 mixture of rotamers) δ 7.94 (d, *J* = 8.2 Hz, 1H), 7.52 – 7.30 (m, 4H), 7.28 (d, *J* = 1.8 Hz, 0.5H), 7.13 (d, *J* = 1.8 Hz, 0.5H), 6.41 (s, 0.5H), 6.10 (s, 0.5H), 6.01 (s, 0.5H), 5.85 (s, 0.5H), 4.63 (d, *J* = 21.1 Hz, 2H), 4.59 – 4.52 (m, 0.5H), 4.18 (dd, *J* = 14.2, 8.8 Hz, 0.5H), 3.81 – 3.55 (m, 6.5H), 3.05 (s, 1.5H), 2.93 (s, 1.5H), 2.58 (q, *J* = 7.6 Hz, 1H), 2.43 (q, *J* = 7.6 Hz, 1H), 2.34 (p, *J* = 7.5 Hz, 0.5H), 2.22 – 2.09 (m, 0.5H), 1.36 – 1.08 (m, 6.5H), 1.02 (d, *J* = 6.9 Hz, 1.5H), 0.83 (d, *J* = 6.9 Hz, 1.5H). $^{13}$C NMR (101 MHz, $CD_3OD$) δ 171.24, 171.16, 165.76, 165.55, 156.57, 156.08, 154.30, 154.27, 154.26, 151.19, 151.06, 146.68, 146.35, 142.71, 139.25, 139.20, 135.31, 135.20, 131.58, 131.21, 131.17, 130.88, 130.59, 130.24, 129.31, 129.26, 126.74, 126.28, 125.68, 125.43, 115.30, 114.84, 104.18, 103.39, 86.17, 84.85, 65.62, 65.49, 64.20, 64.17, 59.00, 58.57, 55.90, 52.36, 51.18, 50.91, 49.28, 39.11, 38.10, 37.94, 34.05, 27.17, 27.15, 27.12, 16.12, 15.90, 15.52, 14.84, 13.35, 13.05. HRMS (ESI): calculated for $C_{30}H_{37}ClN_3O_7S$ [M + H]$^+$, 618.2035; found, 618.2053

*N*-(((4*R*,5*R*)-8-((*E*)-2-Cyclohexylvinyl)-2-((*S*)-1-hydroxypropan-2-yl)-4-methyl-1,1-dioxido-2,3,4,5-tetrahydrobenzo[*b*][1,4,5]oxathiazocin-5-yl)methyl)-*N*-methyl-2-phenylcyclopent-1-ene-1-carboxamide (**38**)

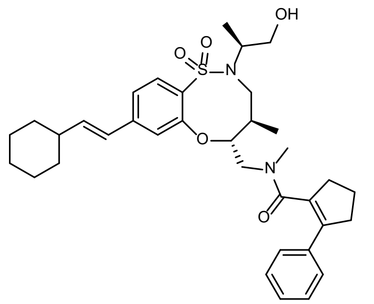

To a 0.5 mL μW vial was added (9H-fluoren-9-yl)methyl (((4*R*,5*R*)-8-bromo-2-((*S*)-1-hydroxypropan-2-yl)-4-methyl-1,1-dioxido-2,3,4,5-tetrahydrobenzo[*b*][1,4,5]oxathiazocin-5-yl)methyl)(methyl)carbamate (20 mg, 1 eq, 32 μmol), (*E*)-(2-cyclohexylvinyl)boronic acid (7.3 mg, 1.50 eq, 48 μmol) and Palladium-Xphos-G2 (5.0 mg, 0.20 eq, 6.4 μmol). The vial was equipped with a magnetic stir bar, capped with a rubber septum and charged with DMF (0.3 mL), giving, after light stirring, a colorless, homogenous solution. This solution was degassed by evacuating at high vacuum and backfilling with $N_2$ (3 x). 1.6 M aq. $K_3PO_4$ (0.12 mL, 6 eq, 0.19 mmol) was then added through the septum. A yellow color evolved quickly; the reaction vessel was heated under µW irradiation at 60 °C for 45 min. The mixture was allowed to reach rt before addition of piperidine (0.06 mL) and stirring for 6 hr. The mixture was then concentrated under reduced pressure and purified by flash chromatography (0–100% MeOH [containing 1% $NH_4OH$] in $CH_2Cl_2$, 4 g silica column) giving (4*R*,5*R*)-8-((*E*)-2-cyclohexylvinyl)-2-((*S*)-1-hydroxypropan-2-yl)-4-methyl-5-((methylamino)methyl)-2,3,4,5-tetrahydrobenzo[*b*][1,4,5]oxathiazocine 1,1-dioxide (15 mg) as a brown residue. The residue was charged to a 20 mL glass scintillation vial containing a magnetic stir bar, then treated with a pre-incubated (10 min) solution of 2-phenylcyclopent-1-ene-1-carboxylic acid (10 mg, 1.5 eq, 53 µmol), PyBOP (30 mg, 1.7 eq, 58 µmol) and DIPEA (15 µL, 2.50 eq, 86 µmol) in DMF (0.1 mL). The walls of the vial were rinsed with additional DMF (0.3 mL), and the resulting brown solution was stirred for 1.25 hr. The reaction solution was partitioned between EtOAc and water and the resulting layers separated. The aqueous phase was extracted with EtOAc (4 x 1 mL) and the combined organics dried over $Na_2SO_4$. The organics were decanted, concentrated under reduced pressure and purified by flash chromatography (0–100% MeOH [containing 1% $NH_4OH$] in $CH_2Cl_2$, 4 g silica column), followed by preparative HPLC to afford **38** (10.5 mg, 50%, 17 µmol) as a white solid.

$^1$H NMR (400 MHz, $CDCl_3$) 1 : 1 mixture of rotamers δ 7.79 (dd, *J* = 8.3, 4.3 Hz, 1H), 7.30 (t, *J* = 3.9 Hz, 3.7H), 7.26 – 7.22 (m, 0.8H), 7.15 (ddd, *J* = 15.8, 8.3, 1.7 Hz, 1H), 7.01 (d, *J* = 1.7 Hz, 0.5H), 6.96 (d, *J* = 1.7 Hz, 0.5H), 6.40 (dd, *J* = 16.0, 6.8 Hz, 0.5H), 6.29 – 6.19 (m, 1.5H), 4.70 (td, *J* = 8.8, 2.6 Hz, 0.5H), 4.46 – 4.35 (m, 1H), 3.95 – 3.75 (m, 2H), 3.65 – 3.56 (m, 1H), 3.56 – 3.43 (m, 1.5H), 3.43 – 3.20 (m, 2H), 3.20 – 3.12 (m, 0.5H), 3.06 – 2.69 (m, 4H), 2.67 (s, 1.4H), 2.63 (s, 1.5H), 2.62 – 2.53 (m, 0.6H), 2.21 – 2.07 (m, 2.5H), 2.06 – 1.92 (m, 1.5H), 1.87 – 1.73 (m, 4H), 1.73 – 1.64 (m, 1H), 1.64 – 1.57 (m, 0.5H), 1.38 – 1.10 (m, 8H), 0.95 (d, *J* = 6.8 Hz, 1.5H), 0.64 (d, *J* = 6.8 Hz, 1.5H). $^{13}$C NMR (101 MHz, $CDCl_3$) δ 171.89, 171.40, 154.38, 153.64, 144.70, 143.95, 141.78, 141.01, 140.21, 139.67, 136.18, 136.05, 132.88, 132.87, 131.49, 131.16, 129.54, 129.33, 128.43, 128.41, 127.84, 127.71, 126.58, 125.45, 125.07, 122.78, 121.87, 120.79, 84.16, 82.81, 77.20, 64.51, 64.48, 57.59, 57.47, 53.51, 49.35, 47.55, 47.10, 41.22, 41.20, 37.19, 37.02,

36.90, 36.44, 36.31, 36.19, 35.82, 32.65, 32.59, 32.51, 32.43, 26.02, 25.91, 25.89, 22.50, 22.43, 15.69, 15.62, 15.09, 14.53. HRMS (ESI): calculated for $C_{35}H_{47}N_2O_5S$ $[M + H]^+$, 607.3200; found, 607.3196.

*N*-(((4*R*,5*R*)-8-((*E*)-2-Cyclohexylvinyl)-2-((*S*)-1-hydroxypropan-2-yl)-4-methyl-1,1-dioxido-2,3,4,5-tetrahydrobenzo[*b*][1,4,5]oxathiazocin-5-yl)methyl)-2-(2,3-dihydrobenzofuran-3-yl)-*N*-methylacetamide (**39**)

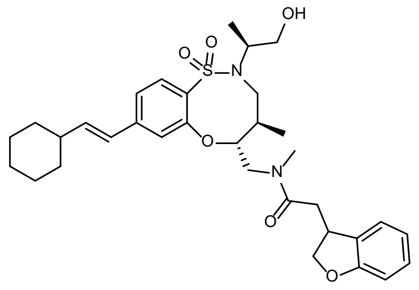

In a 2 mL microwave reaction vial, a solution of (9H-fluoren-9-yl)methyl (((4*R*,5*R*)-8-bromo-2-((*S*)-1-hydroxypropan-2-yl)-4-methyl-1,1-dioxido-2,3,4,5-tetrahydrobenzo[*b*][1,4,5]oxathiazocin-5-yl)methyl)(methyl)carbamate (36 mg, 1.0 eq, 0.057 mmol), XPhos-Pd-G2 (9 mg, 0.2 eq, 0.011 mmol) (*E*)-(2-cyclohexylvinyl)boronic acid (13.2 mg, 1.5 eq, 0.085 mmol) and 1.8 M aq. $K_3PO_4$ (95 µL, 3 eq, 0.171 mmol) in DMF (0.6 mL) was deaerated with Ar at rt for 2 min. The vial was sealed and heated under µW irradiation at 60 °C for 50 min. The solvent was evaporated under the reduced pressure, and the residue was diluted with EtOAc (50 mL), washed with water (10 mL), and brine (10 mL), then was dried over anhydrous. $Na_2SO_4$, was filtered and concentrated under reduced pressure. The residue was purified by flash chromatography (0–60% EtOAc in hexanes) to afford (9H-fluoren-9-yl)methyl (((4*R*,5*R*)-8-((*E*)-2-cyclohexylvinyl)-2-((*S*)-1-hydroxypropan-2-yl)-4-methyl-1,1-dioxido-2,3,4,5-tetrahydrobenzo[*b*][1,4,5]oxathiazocin-5-yl)methyl)(methyl)carbamate (29 mg, 0.044 mmol, 77%) as an off-white foam.

In a 4 mL vial, piperidine (50 µL) was added to a solution of (9H-fluoren-9-yl)methyl (((4*R*,5*R*)-8-((*E*)-2-cyclohexylvinyl)-2-((*S*)-1-hydroxypropan-2-yl)-4-methyl-1,1-dioxido-2,3,4,5-tetrahydrobenzo[*b*][1,4,5]oxathiazocin-5-yl)methyl)(methyl)carbamate (19 mg, 1.0 eq, 0.029 mmol) in DMF (1 mL) at rt and the reaction was stirred at rt for 30 min. The solvent was evaporated, and the residue was purified by flash chromatography (0–10% MeOH [with 1% $NH_4OH$] in $CH_2Cl_2$) to afford

(4*R*,5*R*)-8-((*E*)-2-cyclohexylvinyl)-2-((*S*)-1-hydroxypropan-2-yl)-4-methyl-5-((methylamino)methyl)-2,3,4,5-tetrahydrobenzo[*b*][1,4,5]oxathiazocine 1,1-dioxide (9 mg, 0.021 mmol, 71%) as an off-white foam.

In an 8 mL vial DIPEA (0.019 g, 2.1 eq, 0.15 mmol) was added to a solution of 2-(2,3-dihydrobenzofuran-3-yl)acetic acid (4 mg, 1.0 eq, 0.021 mmol), (4*R*,5*R*)-8-((*E*)-2-cyclohexylvinyl)-2-((*S*)-1-hydroxypropan-2-yl)-4-methyl-5-((methylamino)methyl)-2,3,4,5-tetrahydrobenzo[*b*][1,4,5]oxathiazocine 1,1-dioxide (8 mg, 1.0 eq, 0.058 mmol), and PyBOP (12 mg, 1.1 eq, 0.022 mmol) in DMF (1 mL) at rt and stirred for 30–45 min. The solvent was evaporated under reduced pressure and the residue was purified by flash chromatography (0–10% MeOH [with 1% $NH_4OH$] in $CH_2Cl_2$) to afford a clear oil, which was further purified by preparative HPLC to afford **39** (8 mg, 0.038 mmol, 65%) as an off-white solid, and as a mixture of rotamers and diastereomers (ratio approximately 1:1:1:1, according to analysis by $^1$H NMR spectroscopy).

1H NMR (400 MHz, $CDCl_3$) δ 7.81 (ddd, *J* = 8.0, 6.5, 1.4 Hz, 1H), 7.25 – 7.08 (m, 3H), 6.93 – 6.75 (m, 3H), 6.41 – 5.99 (m, 2H), 4.96 – 4.57 (m, 2H), 4.36 – 4.13 (m, 2H), 4.12 – 3.69 (m, 4H), 3.62 (dt, *J* = 13.1, 9.6 Hz, 1H), 3.56 – 3.44 (m, 1H), 3.38 (dt, *J* = 15.8, 5.9 Hz, 1H), 3.33 – 3.13 (m, 1H), 2.84 (s, 2H), 2.80 – 2.61 (m, 2H), 2.60 – 2.30 (m, 2H), 2.29 – 1.88 (m, 4H), 1.83 – 1.59 (m, 5H), 1.40 – 0.99 (m, 9H), 0.92 (dd, *J* = 23.0, 7.8 Hz, 3H). $^{13}$C NMR (101 MHz, $CDCl_3$) δ 171.94, 171.59, 162.68, 160.17, 160.15, 160.06, 154.84, 154.75, 153.98, 144.91, 144.86, 144.04, 144.01, 141.99, 141.88, 141.30, 131.81, 131.56, 129.92, 129.77, 129.73, 129.68, 129.62, 129.52, 128.78, 128.76, 128.72, 128.68, 125.46, 125.40, 125.37, 124.59, 124.50, 124.36, 122.80, 122.74, 121.87, 121.78, 120.69, 120.61, 120.53, 120.37, 109.94, 109.86, 109.82, 83.91, 83.70, 83.04, 82.66, 77.99, 77.89, 77.68, 77.57, 77.36, 64.75, 57.78, 57.71, 53.31, 50.86, 50.76, 47.90, 47.88, 47.65, 41.38, 41.32, 41.27, 40.12, 40.01, 38.83, 38.78, 38.48, 37.47, 37.39, 36.91, 36.86, 36.62, 35.91, 35.83, 32.79, 32.74, 32.65, 32.59, 31.58, 26.13, 26.07, 25.98, 15.81, 15.79, 14.89, 14.86. HRMS (ESI): calculated for $C_{33}H_{44}N_2O_6S$ $[M + H]^+$, 597.2998; found, 597.3000.

3-(Benzo[*d*][1,3]dioxol-5-yl)-*N*-(((4*R*,5*R*)-2-((*S*)-1-hydroxypropan-2-yl)-4-methyl-1,1-dioxido-8-(1-oxo-2,3,4,5-tetrahydro-1H-benzo[*c*]azepin-7-yl)-2,3,4,5-tetrahydrobenzo[*b*][1,4,5]oxathiazocin-5-yl)methyl)-*N*-methylpropiolamide (**40**)

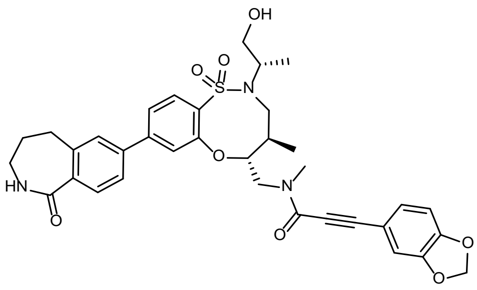

A 5 mL cone-shaped µW vial equipped with a magnetic stir bar was charged with (9H-fluoren-9-yl)methyl (((4*R*,5*R*)-8-bromo-2-((*S*)-1-hydroxypropan-2-yl)-4-methyl-1,1-dioxido-2,3,4,5-tetrahydrobenzo[*b*][1,4,5]oxathiazocin-5-yl)methyl)(methyl)carbamate (31.8 mg, 1 eq, 50.5 µmol), 7-(4,4,5,5-tetramethyl-1,3,2-dioxaborolan-2-yl)-2,3,4,5-tetrahydro-1*H*-benzo[*c*]azepin-1-one (22.9 mg, 95% *wt*, 75.8 µmol, 1.5 eq), XPhos Pd G3 (9.0 mg, 95% Wt, 0.010 mmol, 20 mol%), dry DMF (0.51 mL) and aq. $K_3PO_4$ (1.8 M, 84 µL, 0.15 mmol, 3.0 eq) at rt. The reaction mixture was purged with Ar for 3 min. The vial was then sealed, and the mixture was heated to 60 °C for 2 hr using µW irradiation. EtOAc was added and the mixture was filtered over cotton. The filtrate was concentrated under reduced pressure and the obtained crude material was purified by flash chromatography (0–20% MeOH [with 1% $NH_4OH$] in $CH_2Cl_2$), affording 7-((4*R*,5*R*)-2-((*S*)-1-hydroxypropan-2-yl)-4-methyl-5-((methylamino)methyl)-1,1-dioxido-2,3,4,5-tetrahydrobenzo[*b*][1,4,5]oxathiazocin-8-yl)-2,3,4,5-tetrahydro-1H-benzo[*c*]azepin-1-one (4.8 mg, 9.8 µmol, 19%) as a cloudy oil.

A 2 dram scintillation vial equipped with a magnetic stir bar was charged with 7-((4*R*,5*R*)-2-((*S*)-1-hydroxypropan-2-yl)-4-methyl-5-((methylamino)methyl)-1,1-dioxido-2,3,4,5-tetrahydrobenzo[*b*][1,4,5]oxathiazocin-8-yl)-2,3,4,5-tetrahydro-1H-benzo[*c*]azepin-1-one (4.8 mg, 9.8 µmol, 1.0 eq) at rt. A solution of 3-(benzo[*d*][1,3]dioxol-5-yl)propiolic acid (2.2 mg, 12 µmol, 1.2 eq), COMU (5.2 mg, 12 µmol, 1.2 eq) and DIPEA (3.5 µL, 20 µmol, 2.0 eq) in dry DMF (0.24 mL) was then added and the resulting orange solution was stirred for 14 min. Water and EtOAc were added, and the aqueous phase was extracted twice with EtOAc. The organic layers were combined, dried over $MgSO_4$ and concentrated under reduced pressure. The material was then purified by preparative HPLC to afford **40**(4.0 mg, 6.1 µmol, 62%) as a pale yellow oil.

$^{1}H$ NMR (400 MHz, $CD_3OD$) (~3:2 ratio of rotamers) δ 7.94 (d, *J* = 0.8 Hz, 0.4H), 7.92 (d, *J* = 0.8 Hz, 0.6H), 7.70 (d, *J* = 1.1 Hz, 1H), 7.65 (dd, *J* = 7.9, 1.8 Hz, 1H), 7.57 (t, *J* = 2.0 Hz, 0.6H), 7.55 (t, *J* = 2.0 Hz, 0.4H), 7.52 (d, *J* = 1.8 Hz, 0.6H), 7.47 (d, *J* = 8.0 Hz, 0.6H), 7.42 (d, *J* = 1.8 Hz, 0.6H), 7.38 (d, *J* = 1.8 Hz, 0.4H), 7.19 (dd, *J* = 8.1, 1.6 Hz, 0.4H), 7.06 (d, *J* = 1.6 Hz, 0.4H), 6.87 (d, *J* = 8.1 Hz, 0.4H), 6.83 (dd, *J* = 8.1, 1.6 Hz, 0.6H), 6.77 (d, *J* = 8.0 Hz, 0.6H), 6.68 (d, *J* = 1.5 Hz, 0.6H), 6.02 (s, 2H), 4.83 – 4.75 (m, 1H), 4.44 (ddd, *J* = 24.7, 14.2, 10.5 Hz, 1H), 3.87 – 3.56 (m, 5.4H), 3.37 – 3.33 (m, 0.6H), 3.16 (s, 1.75H), 3.10 (t, *J* = 6.6 Hz, 0.8H), 2.95 (t, *J* = 7.1 Hz, 0.8H), 2.89 (t, *J* = 6.6 Hz, 1.2H), 2.87 (s, 1.25H), 2.84 – 2.68 (m, 1.2H), 2.44 – 2.26 (m, 1H), 2.14 – 2.05 (m, 0.8H), 2.00 – 1.86 (m, 1.2H), 1.27 (dd, *J* = 6.7, 1.8 Hz, 3H), 1.00 (d, *J* = 6.9 Hz, 1.25H), 0.96 (d, *J* = 7.0 Hz, 1.75H). $^{13}C$ NMR (101 MHz, $CD_3OD$) δ 175.46, 175.07, 157.88, 157.62, 157.10, 156.90, 151.55, 151.51, 149.38, 149.20, 147.17, 146.81, 142.91, 142.58, 140.87, 140.78, 136.40, 136.23, 134.93, 134.92, 134.92, 131.16, 131.11, 130.26, 130.16, 129.19, 128.86, 128.34, 126.83, 126.53, 123.76, 123.64, 123.33, 122.89, 114.19, 113.72, 112.76, 112.72, 109.87, 109.74, 103.38, 93.99, 93.57, 84.80, 84.10, 81.35, 80.88, 65.57, 58.50, 58.38, 56.34, 51.14, 51.09, 40.36, 40.22, 38.16, 38.05, 37.47, 33.11, 31.68, 31.59, 31.23, 31.20, 15.87, 15.85, 14.95, 14.81. HRMS (ESI): calculated for $C_{35}H_{38}N_3O_8S$ $[M + H]^+$, 660.2374; found, 660.2408.

3-(Benzo[*d*][1,3]dioxol-5-yl)-*N*-(((4*R*,5*R*)-8-bromo-2-((*R*)-1-hydroxypropan-2-yl)-4-methyl-1,1-dioxido-2,3,4,5-tetrahydrobenzo[*b*][1,4,5]oxathiazocin-5-yl)methyl)-*N*-methylpropiolamide (**41**)

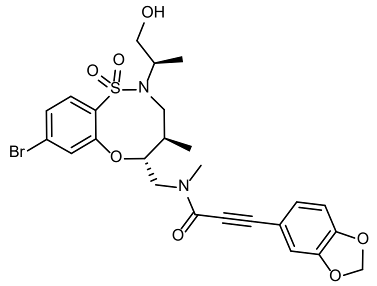

A 2 dram scintillation vial equipped with a magnetic stir bar was charged with (4*R*,5*R*)-8-bromo-2-((*R*)-1-hydroxypropan-2-yl)-4-methyl-5-((methylamino)methyl)-2,3,4,5-tetrahydrobenzo[*b*][1,4,5]oxathiazocine 1,1-dioxide [*cf*. synthesis of **35**] (8.0 mg, 1.0 eq, 20 μmol), dry DMF (0.20 mL), DIPEA (5.3 mg, 7.1 μL, 2.1 eq, 41 μmol), 3-(benzo[*d*][1,3]dioxol-5-yl)propiolic acid (4.8 mg, 1.3 eq, 25 μmol) and PyBOP (13 mg, 1.3 eq, 25 μmol) under $N_2$ atmosphere and at rt. The faint yellow reaction mixture was stirred for 35 min. The sample was then concentrated under reduced pressure and purified by preparative HPLC (optimal gradient: 30–70% MeCN in $H_2O$) to afford **41** (7.3 mg, 13 μmol, 64%) as a white solid.

$^{1}H$ NMR (400 MHz, $CDCl_3$) (~3:1 ratio of rotamers) δ 7.76 (dd, *J* = 8.5, 2.2 Hz, 1H), 7.39 – 7.27 (m, 2H), 7.13 (ddd, *J* = 8.1, 3.9, 1.6 Hz, 1H), 7.00 (dd, *J* = 4.3, 1.6 Hz, 1H), 6.80 (dd, *J* = 8.0, 2.3 Hz, 1H), 6.02 (s, 2H), 4.79 – 4.61 (m, 1H), 4.45 (dd, *J* = 14.0, 9.9 Hz, 0.7H), 4.27 (dd, *J* = 14.6, 9.7 Hz, 0.3H), 3.89 – 3.46 (m, 5.3H), 3.18 (dd, *J* = 14.0, 2.7 Hz, 0.7H), 3.12 (s, 2.1H), 2.87 (s, 0.9H), 2.41 – 2.22 (m, 2H), 1.24 (t, *J* = 7.0 Hz, 3H), 0.96 (t, *J* = 7.6 Hz, 3H). $^{13}C$ NMR (101 MHz, $CDCl_3$) δ 156.16, 156.05, 155.71, 155.49, 149.75, 147.81, 147.73, 132.24, 131.83, 131.32, 131.16, 128.25, 128.00, 127.71, 127.52, 127.13, 127.08, 113.54, 113.51, 112.32, 112.16, 108.91, 108.83, 101.86, 101.83, 92.41, 91.60, 84.21, 83.64, 80.62, 80.23, 64.88, 56.58, 56.44, 54.89, 49.50, 48.78, 48.55, 37.34, 37.06, 36.85, 33.17, 15.95, 15.92, 15.30, 15.19. HRMS (ESI): calculated for $C_{25}H_{28}{}^{81}BrN_2O_7S$ $[M + H]^+$, 581.0775 found, 581.0788.

3-(Benzo[*d*][1,3]dioxol-5-yl)-*N*-(((4*R*,5*R*)-8-bromo-2-((*S*)-1-hydroxypropan-2-yl)-4-methyl-1,1-dioxido-2,3,4,5-tetrahydrobenzo[*b*][1,4,5]oxathiazocin-5-yl)methyl)-*N*-methylpropiolamide (**42**)

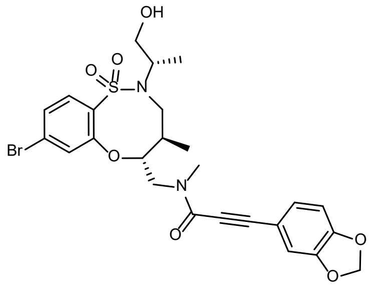

(9H-Fluoren-9-yl)methyl (((4*R*,5*R*)-8-bromo-2-((*S*)-1-hydroxypropan-2-yl)-4-methyl-1,1-dioxido-2,3,4,5-tetrahydrobenzo[*b*][1,4,5]oxathiazocin-5-yl)methyl)(methyl)carbamate (60 mg, 1 eq, 95 μmol) was charged to a 1-dram glass vial. DMF (0.56 mL) and piperidine (0.14 mL) were then added and the colorless solution was stirred at rt for 45 min. The reaction solution was concentrated under reduced pressure, then azeotropically dried with heptane (4 x 50 mL) to provide the crude product as a white powder, a portion of which was carried on without further purification.

In a 1-dram vial containing a magnetic stir bar were combined 3-(benzo[*d*][1,3]dioxol-5-yl)propiolic acid (15 mg, 1.30 eq, 77 μmol) and PyBOP (40 mg, 1.30 eq, 77 μmol). DMF (0.2 mL) was added, giving a light brown solution. Addition of DIPEA (26 µL, 2.50 eq, 0.15 mmol) then afforded a thick, off-white slurry. After stirring for 5 min at rt, (4*R*,5*R*)-8-bromo-2-((*S*)-1-hydroxypropan-2-yl)-4-methyl-5-((methylamino)methyl)-2,3,4,5-tetrahydrobenzo[*b*][1,4,5]oxathiazocine 1,1-dioxide (24 mg, 1.00 eq, 59 µmol) was added and the walls of the flask were rinsed with DMF (0.8 mL), giving a brown solution. The reaction solution was stirred for 10 min before dilution with EtOAc and water, giving a biphasic mixture. Layers were separated, and the aqueous phase was extracted with EtOAc (3 x 1 mL); the combined organics were then dried over $Na_2SO_4$, decanted and concentrated under reduced pressure to reveal a yellow oil. Purification by flash chromatography (0–100% MeOH [with 1% $NH_4OH$] in $CH_2Cl_2$, 4 g silica column), followed by (0–100% EtOAc in hexanes) followed by preparative HPLC afforded **42** (8.7 mg, 15 µmol, 26%) as a white solid.

$^1H$ NMR (400 MHz, $CDCl_3$) (~2:1 ratio of rotamers) δ 7.77 (dd, *J* = 8.8, 2.9 Hz, 1H), 7.40 – 7.27 (m, 2H), 7.13 (ddd, *J* = 8.1, 2.7, 1.6 Hz, 1H), 7.00 (dd, *J* = 5.7, 1.6 Hz, 1H), 6.80 (dd, *J* = 8.1, 5.5 Hz, 1H), 6.02 (s, 2H), 4.82 – 4.74 (m, 1H), 4.54 (dd, *J* = 14.0, 10.2 Hz, 0.7H), 4.36 (dd, *J* = 14.6, 10.0 Hz, 0.3H), 3.97 – 3.81 (m, 2H), 3.67 – 3.36 (m, 3.3H), 3.13 (dd, *J* = 14.1, 2.8 Hz, 0.7H), 3.10 (s, 2H), 2.83 (s, 1H), 2.33 – 2.18 (m, 1H), 1.22 (dd, *J* = 6.8, 5.5 Hz, 3H), 0.97 (dd, *J* = 13.8, 6.9 Hz, 3H). $^{13}C$ NMR (101 MHz, $CDCl_3$) δ 156.24, 155.55, 154.72, 154.32, 149.59, 147.52, 147.46, 133.11, 132.86, 130.42, 128.11, 128.05, 127.95, 127.77, 127.48, 127.01, 126.89, 113.02, 112.92, 112.05, 111.95, 108.70, 108.61, 101.63, 92.77, 92.09, 83.65, 82.92, 79.96, 79.73, 77.20, 64.36, 64.24, 57.53, 57.41, 54.59, 48.38, 48.27, 36.69, 36.59, 36.47, 32.72, 15.39, 14.65, 14.43. HRMS (ESI): calculated for $C_{25}H_{28}{}^{81}BrN_2O_7S$ $[M + H]^+$, 581.0775 found, 581.0797.

3-(Benzo[*d*][1,3]dioxol-5-yl)-1-((8*R*,9*R*,10*S*)-9-(4-bromophenyl)-10-(hydroxymethyl)-1,6-diazabicyclo[6.2.0]decan-6-yl)prop-2-yn-1-one (**43**)

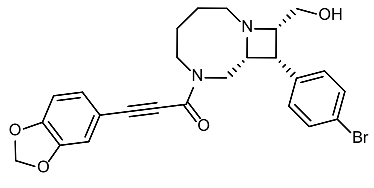

A 1-dram scintillation vial equipped with a magnetic stir bar was charged with ((8*R*,9*R*,10*S*)-9-(4-bromophenyl)-6-((4-nitrophenyl)sulfonyl)-1,6-diazabicyclo[6.2.0]decan-10-yl)methanol (8.5 mg, 1 eq, 17 µmol). Dry DMF (0.28 mL), $K_2CO_3$ (12 mg, 5.0 eq, 83 µmol) and 4-methoxythiophenol (9.6 mg, 8.4 µL, 97% Wt, 4.0 eq, 67 µmol) were successively added and the yellow suspension was heated to 50 °C and stirred for 50 min. The obtained yellow/orange suspension was concentrated under reduced pressure. EtOAc (30 mL) and 0.3 M aq. HCl (30 mL) were added, phases separated, and aqueous portion was extracted twice with EtOAc (2 x 30 mL). The aqueous layer was then basified with 40% aq. KOH solution and extracted EtOAc (3 x 30 mL). The organic layers were combined, dried over $Na_2SO_4$, filtered and solvent was removed under reduced pressure to afford crude ((8*R*,9*R*,10*S*)-9-(4-bromophenyl)-1,6-diazabicyclo[6.2.0]decan-10-yl)methanol as a faintly yellow, grey solid. To this solid was added dry DMF (0.34 mL), DIPEA (4.5 mg, 6.1 µL, 2.1 eq, 35 µmol), 3-(benzo[*d*][1,3]dioxol-5-yl)propiolic acid (3.5 mg, 1.1 eq, 18 µmol) and COMU (8.1 mg, 97%wt, 1.1 eq, 18 µmol) at rt. The orange/brown solution was stirred for 50 min, then a drop of water and DMSO were added, and the mixture was filtered prior to purification by preparative HPLC to afford **43** (3.64 mg, 17 µmol, 44%) as a faint yellow oil.

$^1$H NMR (400 MHz, $CD_3OD$) (~55:45 ratio of rotamers) δ 7.53 – 7.40 (m, 4H), 7.12 (dd, *J* = 8.0, 1.6 Hz, 0.45H), 7.03 (dd, *J* = 8.0, 1.6 Hz, 0.55H), 7.00 (d, *J* = 1.6 Hz, 0.45H), 6.91 (d, *J* = 1.6 Hz, 0.55H), 6.88 (dd, *J* = 8.0, 5.3 Hz, 1H), 6.04 (d, *J* = 6.3 Hz, 2H), 4.30 (ddd, *J* = 14.9, 6.7, 3.4 Hz, 0.45H), 4.03 (ddd, *J* = 13.8, 6.5, 4.0 Hz, 0.55H), 3.76 – 3.54 (m, 3.75H), 3.50 – 3.32 (m, 5.25H), 3.22 – 3.09 (m, 1.55H), 3.01 – 2.93 (m, 0.45H), 2.41 – 2.28 (m, 1H), 2.11 (ddd, *J* = 14.6, 7.0, 3.5 Hz, 0.45H), 1.94 – 1.57 (m, 3.55H). $^{13}$C NMR (101 MHz, $CD_3OD$) δ 156.54, 156.19, 151.47, 151.46, 151.45, 149.38, 136.79, 136.71, 133.83, 132.25, 132.09, 129.16, 129.01, 121.95, 121.84, 114.26, 114.16, 112.76, 112.69, 109.86, 109.84, 103.38, 103.36, 93.03, 92.57, 81.13, 80.86, 69.86, 69.56, 68.07, 67.19, 62.01, 61.91, 60.06, 59.72, 54.14, 52.39, 50.39, 43.97, 43.68, 29.91, 28.29, 28.03, 27.58. HRMS (ESI): calculated for $C_{25}H_{26}{}^{81}BrN_2O_4$ $[M + H]^+$, 499.1050 found, 499.1056.

3-(Benzo[*d*][1,3]dioxol-5-yl)-1-((8*S*,9*S*,10*S*)-9-(4-bromophenyl)-10-(hydroxymethyl)-1,6-diazabicyclo[6.2.0]decan-6-yl)prop-2-yn-1-one (**44**)

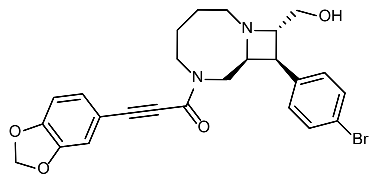

A 20 mL scintillation vial equipped with a magnetic stir bar was charged with ((8*S*,9*S*,10*S*)-9-(4-bromophenyl)-6-((2-nitrophenyl)sulfonyl)-1,6-diazabicyclo[6.2.0]decan-10-yl)methanol (49 mg, 1.0 eq, 96 μmol), dry DMF (1.6 mL), 4-methoxy-benzenethiol (54 mg, 47 μL, 4 eq, 0.38 mmol) and $K_2CO_3$ (66 mg, 5 eq, 0.48 mmol) at rt. The suspension was heated to 50 °C and stirred for 80 min. The obtained yellow suspension was concentrated under reduced pressure to afford a yellow solid. EtOAc (30 mL) and 0.3 M aq. HCl (30 mL) were added, phases separated, and the aqueous phase was extracted twice with EtOAc. The aqueous portion then had pH adjusted to 13–14 with 40% aq. KOH solution and was extracted three times with EtOAc. The organic layers were combined, dried over $Na_2SO_4$ and concentrated under reduced pressure to afford crude ((8*S*,9*S*,10*S*)-9-(4-bromophenyl)-1,6-diazabicyclo[6.2.0]decan-10-yl)methanol (27 mg, 83 μmol) as a pale-yellow solid. To this residue was added dry DMF (1.6 mL), 3-(benzo[*d*][1,3]dioxol-5-yl)propiolic acid (19 mg, 1.2 eq, 0.10 mmol), DIPEA (21 mg, 2 eq, 0.17 mmol) and COMU (39 mg, 1.1 eq, 91 μmol) at rt and the mixture was stirred for 30 min. The mixture was concentrated under reduced pressure, then purified by preparative HPLC to afford **44** (10 mg, 20 μmol, 24 %) as a colorless oil.

$^1$H NMR (400 MHz, $CD_3OD$) (~55:45 ratio of rotamers) δ 7.54 (dd, *J* = 33.9, 8.5 Hz, 2H), 7.34 (t, *J* = 8.1 Hz, 2H), 7.13 (dd, *J* = 8.1, 1.6 Hz, 0.55H), 7.00 (d, *J* = 1.6 Hz, 0.55H), 6.94 – 6.89 (m, 0.9H), 6.87 (d, *J* = 8.0 Hz, 0.55H), 6.83 (t, *J* = 1.0 Hz, 0.45H), 6.07 (s, 0.9H), 6.03 (s, 1.1H), 4.68 – 4.55 (m, 1.55H), 4.47 (dt, *J* = 8.0, 4.1 Hz, 0.45H), 4.30 (dt, *J* = 15.5, 4.8 Hz, 0.56H), 4.23 – 3.66 (m, 5H), 3.57 – 3.25 (m, 3H), 3.19 (ddd, *J* = 14.4, 8.2, 3.6 Hz, 0.45H), 2.11 – 1.60 (m, 4H). $^{13}$C NMR (101 MHz, $CD_3OD$) δ 156.77, 156.20, 151.66, 151.58, 149.38, 135.07, 134.37, 133.25, 133.15, 131.68, 131.48, 129.39, 129.03, 123.05, 122.98, 113.84, 113.71, 112.80, 112.66, 109.89, 109.88, 103.46, 103.41, 93.31, 93.25, 80.61, 80.34, 71.21, 70.87, 68.79, 67.42, 60.22, 59.82, 50.24, 46.03, 45.16, 40.67, 40.07, 27.55, 26.11, 24.86, 23.46. HRMS (ESI): calculated for $C_{25}H_{26}{}^{81}BrN_2O_4$ $[M + H]^+$, 499.1050 found, 499.1059.

(2*R*,4*R*)-*N*-Methyl-1-((4-sulfamoylphenyl)sulfonyl)-4-(3-(1,2,3,6-tetrahydropyridin-4-yl)phenoxy)pyrrolidine-2-carboxamide (**45**)

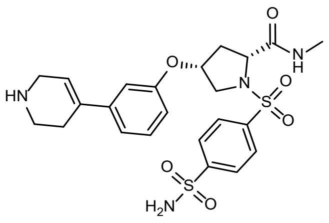

(2*R*,4*R*)-4-(3-Iodophenoxy)-2-(methoxycarbonyl)pyrrolidin-1-ium 2,2,2-trifluoroacetate (43 mg, 93 μmol, 1 eq) and 4-sulfamoylbenzenesulfonyl chloride (2.9 mg, 0.11 mmol, 1.2 eq) were dissolved in anhydrous $CH_2Cl_2$ (500 μL, 0.2 M) in a 1-dram vial. $NEt_3$ (39 μL, 0.28 mmol, 3 eq) was added and the reaction was allowed to stir at rt for 4 h. The crude reaction mixture was washed with 1:1 mixture of water:brine (1x), the organic phase was concentrated under reduced pressure, and the crude product was dissolved in THF (2 mL), followed by addition of 1 N aq. LiOH solution (1 mL). The reaction was stirred overnight, then was acidified to pH 2 with dilute aq. HCl and extracted with $CH_2Cl_2$ five times. The pooled organics were dried over anhydrous $MgSO_4$ and concentrated under reduced pressure. The residue was used for the next step without further purification.

To a 1-dram vial containing crude (2*R*,4*R*)-4-(3-iodophenoxy)-1-((4-sulfamoylphenyl)sulfonyl)pyrrolidine-2-carboxylic acid (26 mg, 47 μmol, 1 eq), $NH_3MeCl$ (6.4 mg, 94 μmol, 2 eq) and HATU (27 mg, 71 μmol, 1.5 eq) was added anhydrous DMF (450 μL), followed by DIPEA (41 μL, 0.24 mmol, 5 eq). The resulting yellow solution was stirred overnight at rt. The solution was diluted with $CH_2Cl_2$ and washed with water (3 x 1 mL), followed by brine (1 x 1 mL). The organic phase was dried over anhydrous $MgSO_4$ and concentrated under reduced pressure. The resulting crude mixture was added to a 2 mL μW vial and 4-(4,4,5,5-tetramethyl-1,3,2-dioxaborolan-2-yl)-1,2,3,6-tetrahydropyridine (15 mg, 71 μmol, 1.5 eq) and XPhos-Pd-G2 (7.4 mg, 9.4 μmol, 0.2 eq) was added. The vial was capped with a septum and purged 3 times with $N_2$ gas. Anhydrous DMF (0.5 mL) was added, followed by 1.8 M aq. $K_3PO_4$ (0.16 mL, 6 eq). The reaction was stirred at 60 °C for 2 hr under μW irradiation. The resulting dark brown suspension was diluted with DMSO, filtered through a syringe filter, and purified by preparative to afford **45** (3.9 mg, 7.5 μmol, 16%) as a white solid.

$^1H$ NMR (400 MHz, $CD_3OD$) δ 8.55 (s, 1H), 8.12 (dd, *J* = 16.7, 2.6 Hz, 4H), 7.27 (t, *J* = 8.0 Hz, 1H), 7.07 (dd, *J* = 7.7, 1.7 Hz, 1H), 6.87 (t, *J* = 2.1 Hz, 1H), 6.80 (dd, *J* = 8.2, 2.4 Hz, 1H), 6.13 (t, *J* = 1.8 Hz, 1H), 4.91 (t, *J* = 4.1 Hz, 1H), 4.27 (dd, *J* = 9.9, 1.7 Hz, 1H), 3.87 – 3.73 (m, 3H), 3.56 (dd, *J* = 11.5, 4.5 Hz, 1H), 3.41 (t, *J* = 6.1 Hz, 2H), 2.83 (s, 3H), 2.77 – 2.69 (m, 2H), 2.37 (d, *J* = 14.1 Hz, 1H), 1.91 (ddd, *J* = 14.2, 9.9, 4.2 Hz, 1H). $^{13}C$ NMR (101 MHz, $CD_3OD$) δ 174.24, 158.09, 149.94, 142.38, 140.35, 136.73, 130.90, 129.86, 128.42, 119.53, 118.19, 116.10, 114.10, 76.88, 62.96, 55.81, 43.63, 42.31, 36.61, 26.76, 25.23. HRMS (ESI): calculated for $C_{23}H_{29}N_4O_6S_2$ $[M + H]^+$, 521.1523 found, 521.1536.

(*R*)-5-((2-Acetyl-8-(3'-((dimethylamino)methyl)-4'-fluoro-[1,1'-biphenyl]-4-yl)-2,6-diazaspiro[3.4]octan-6-yl)methyl)furan-2-sulfonamide (**46**)

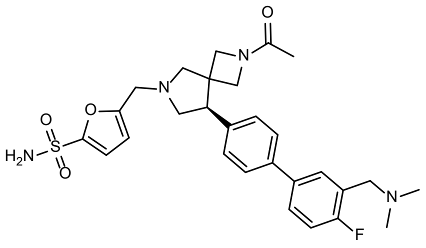

*tert*-Butyl (*R*)-8-(4-iodophenyl)-2-((2-nitrophenyl)sulfonyl)-2,6-diazaspiro[3.4]octane-6-carboxylate (30 mg, 55 μmol, 1 eq) and $K_2CO_3$ (14 mg, 0.1 mmol, 2 eq) was suspended in anhydrous DMF (400 μL). The resulting suspension was cooled to 0 °C in an ice bath, followed by addition of thiophenol (5.7 μL, 55 μmol, 1.1 eq). The resulting mixture was stirred for 1 hr at 0 °C, then diluted with EtOAc and filtered through a pad of celite. The filtrate was concentrated under reduced pressure and used for the next step without further purification.

The crude mixture from previous reaction (55 μmol, 1 eq) was dissolved in $CH_2Cl_2$ (0.5 mL), followed by addition of $Ac_2O$ (10 μL, 0.11 mmol, 2 eq) and $NEt_3$ (15.0 μL, 0.11 μmol, 2 eq). The solution was stirred at rt for 2 hr and concentrated under reduced pressure. This crude mixture was added to a 2 mL μW vial and (3-((dimethylamino)methyl)-4-fluorophenyl)boronic acid (16 mg, 83 μmol, 1.5 eq) and XPhos-Pd-G2 (8.7 mg, 11 μmol, 0.2 eq) were added. The vial was capped with a septum and purged 3 times with $N_2$ gas. Anhydrous DMF (550 μL) was added, followed by 1.8 M aq. $K_3PO_4$ (0.18 mL, 330 μmol, 6 eq). The reaction was stirred at 60 °C for 120 min under μW irradiation, then the solution was washed with brine twice and product was purified by flash chromatography (MeOH [with 1% $NH_4OH$] in $CH_2Cl_2$) to afford *tert*-butyl (*R*)-2-acetyl-8-(3'-((dimethylamino)methyl)-4'-fluoro-[1,1'-biphenyl]-4-yl)-2,6-diazaspiro[3.4]octane-6-carboxylate (15 mg, 31 μmol, 57% over 3 steps).

*tert*-butyl (*R*)-2-acetyl-8-(3'-((dimethylamino)methyl)-4'-fluoro-[1,1'-biphenyl]-4-yl)-2,6-diazaspiro[3.4]octane-6-carboxylate (15 mg, 31 μmol, 1 eq) was dissolved in $CH_2Cl_2$ (0.5 mL), and TFA (100 μL) was added. After 1.5 hr at rt the solution was concentrated under reduced pressure and to the resulting crude residue was added 5-formylfuran-2-sulfonamide (6.0 mg, 34 μmol, 1.1 eq) and sodium acetoxyborohydride (13 mg, 62 μmol, 2 eq). Anhydrous DCE (300 μL) and anhydrous DMF (100 μL) were added, followed by $NEt_3$ (4.3 μL, 31 μmol, 1 eq). The reaction was stirred at rt for 1 hr and concentrated under reduced pressure. The residue was diluted with DMSO and purified directly by preparative HPLC to afford (*R*)-5-((2-acetyl-8-(3'-((dimethylamino)methyl)-4'-fluoro-[1,1'-biphenyl]-4-yl)-2,6-diazaspiro[3.4]octan-6-yl)methyl)furan-2-sulfonamide (2.2 mg, 4 μmol, 13% over 2 steps).

$^1H$ NMR (400 MHz, $CD_3OD$) (~1:1 ratio of rotamers) δ 7.78 (ddq, *J* = 7.5, 4.9, 2.6 Hz, 2H), 7.72 – 7.58 (m, 2H), 7.45 – 7.25 (m, 3H), 6.93 (d, *J* = 3.4 Hz, 1H), 6.49 (d, *J* = 3.4 Hz, 1H), 4.29 (d, *J* = 8.8 Hz, 2H), 4.21 (d, *J* = 2.2 Hz, 1H), 3.98 (s, 1H), 3.93 – 3.81 (m, 2.5H), 3.67 (d, *J* = 9.2 Hz, 0.5H), 3.58 (d, *J* = 10.4 Hz, 0.5H), 3.52 – 3.44 (m, 1.5H), 3.30 – 3.22 (m, 1.5H), 3.19 (d, *J* = 9.8 Hz, 0.5H), 3.13 (d, *J* = 9.8 Hz, 0.5H), 3.07 (d, *J* = 9.9 Hz, 0.5H), 2.93 (ddd, *J* = 20.5, 9.7, 7.6 Hz, 1H), 2.80 (d, *J* = 6.4 Hz, 6H), 1.80 (s, 1.6H), 1.61 (s, 1.4H). $^{13}C$ NMR (101 MHz, $CD_3OD$) δ 155.42, 151.27, 128.83, 126.95, 116.18, 115.96, 113.86, 109.56, 64.93, 64.66, 62.52, 59.90, 59.13, 54.52, 54.27, 51.61, 51.49, 50.36, 43.53, 43.35, 42.28, 42.23, 17.07. LRMS (ESI): calculated for $C_{28}H_{34}FN_4O_4S$ $[M + H]^+$, 541.23 found, 541.26.

(2*S*,3*S*)-3-(3'-(Ethylcarbamoyl)-[1,1'-biphenyl]-4-yl)-*N*-methyl-1-(4-sulfamoylbenzyl)pyrrolidine-2-carboxamide (**47**)

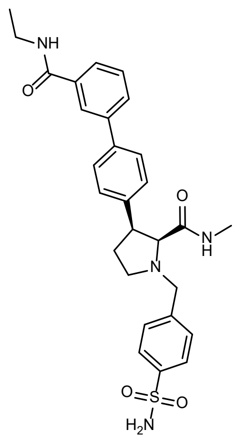

A 20 mL vial was charged with (2S,3S)-1-(tert-butoxycarbonyl)-3-(4-iodophenyl)pyrrolidine-2-carboxylic acid (20.9 mg, 1.00 eq, 50.0 μmol) and HATU (22.8 mg, 1.20 eq, 60.0 μmol). DMF (5 mL) was added followed by $NEt_3$ (15.2 mg, 20.9 μL, 3.00 eq, 150 μmol) and solution was stirred for 15 min. Methanamine, HCl (6.75 mg, 2.00 eq, 100 μmol) was then added as a solid and reaction was further stirred for 3 hr. The solution was diluted with $CH_2Cl_2$ (50 mL), washed with 1 M aq. citric acid (2 x 50 mL), dried over $MgSO_4$, filtered, and solvent removed under reduced pressure. *tert*-Butyl (2*S*,3*S*)-3-(4-iodophenyl)-2-(methylcarbamoyl)pyrrolidine-1-carboxylate (89%) was isolated after flash chromatography (0–20% MeOH in $CH_2Cl_2$) as a yellow oil.

*tert*-Butyl (2*S*,3*S*)-3-(4-iodophenyl)-2-(methylcarbamoyl)pyrrolidine-1-carboxylate (21.5 mg, 1.00 eq, 50.0 μmol) was dissolved by $CH_2Cl_2$ (5 mL). TFA (114 mg, 77.0 μL, 20.0 eq, 1.00 mmol) was added via syringe and reaction stirred for 30 min. Volatiles were removed under reduced pressure, the residue was dissolved in $CH_2Cl_2$ (50 mL) and washed with 1M $K_2CO_3$ ( 3 x 50 mL). Aqueous layers were combined and extracted with $CH_2Cl_2$ (3 x 100 mL). Organic portions were combined, dried over $Na_2SO_4$, filtered, and solvent was removed under reduced pressure to afford crude (2*S*,3*S*)-3-(4-iodophenyl)-*N*-methylpyrrolidine-2-carboxamide, which was used directly in the next step.

Crude (2*S*,3*S*)-3-(4-iodophenyl)-*N*-methylpyrrolidine-2-carboxamide (30.4 mg, 1.00 eq, 92.0 μmol), 4-formylbenzenesulfonamide (25.6 mg, 1.50 eq, 138 μmol), and sodium triacetoxyborohydride (97.5 mg, 5.00 eq, 460 μmol) were stirred in DCE (8 mL) for 1.5 hr at rt. The reaction mixture was diluted with $CH_2Cl_2$ (50 mL) and washed with saturated aq. $NaHCO_3$ (3 x 50 mL). Aqueous layers were combined and extracted with $CH_2Cl_2$ (3 x 100 mL). Organic portions were combined, dried over $MgSO_4$, filtered, and volatiles removed under reduced pressure prior to purification by preparative HPLC to afford (2*S*,3*S*)-3-(4-iodophenyl)-*N*-methyl-1-(4-sulfamoylbenzyl)pyrrolidine-2-carboxamide (22 mg, 43 µmol, 47% over two steps) as an off-white solid.

A 5 mL µW vial was charged with (2*S*,3*S*)-3-(4-iodophenyl)-*N*-methyl-1-(4-sulfamoylbenzyl)pyrrolidine-2-carboxamide (12.5 mg, 1.00 eq, 25.0 µmol), (3-(ethylcarbamoyl)phenyl)boronic acid (7.24 mg, 1.50 eq, 37.5 µmol), and Palladium-Xphos (3.93 mg, 0.20 eq, 5.00 µmol). DMF was added followed by 1.8 M aq. $K_3PO_4$ (15.9 mg, 41.7 µL, 3.00 eq, 75.0 µmol). The vial was capped and sparged with $N_2$ for 5 min before heating at 60 °C for 1 hr under µW irradiation. After cooing the solution was diluted with 1:4 MeOH:$CH_2Cl_2$ (20 mL), filtered through celite and rinsed with flushing 1:4 MeOH:$CH_2Cl_2$ (5 x 50 mL). Volatiles were removed under reduced pressure and crude product was purified by preparative HPLC to afford **47** (7.4 mg, 14.3 µmol, 57%).

$^1$H NMR (400 MHz, DMSO) δ 8.57 (t, *J* = 5.5 Hz, 1H), 8.10 (t, *J* = 1.8 Hz, 1H), 7.83 – 7.77 (m, 4H), 7.61 (dd, *J* = 8.4, 2.0 Hz, 4H), 7.59 – 7.51 (m, 2H), 7.34 (d, *J* = 7.8 Hz, 4H), 3.90 (d, *J* = 13.7 Hz, 1H), 3.70 – 3.57 (m, 2H), 3.46 (d, *J* = 9.3 Hz, 1H), 3.10 (ddd, *J* = 9.0, 6.5, 3.2 Hz, 1H), 2.58 – 2.52 (m, 1H), 2.33 (d, *J* = 4.7 Hz, 3H), 2.16 (tq, *J* = 7.9, 3.3 Hz, 2H), 1.16 (t, *J* = 7.2 Hz, 3H). $^{13}$C NMR (101 MHz, DMSO) δ 170.79, 165.82, 142.88, 142.75, 140.64, 139.96, 137.40, 135.30, 129.08, 128.99, 128.94, 128.91, 126.09, 125.95, 125.53, 124.99, 71.70, 57.67, 52.25, 46.85, 34.07, 31.19, 25.18, 14.81. HRMS (ESI): calculated for $C_{28}H_{33}N_4O_4S$ $[M + H]^+$, 522.2217 found, 521.2228.

(2*R*,3*S*)-3-(3'-Ethoxy-[1,1'-biphenyl]-3-yl)-*N*-methyl-1-((5-sulfamoylfuran-3-yl)methyl)pyrrolidine-2-carboxamide (**48**)

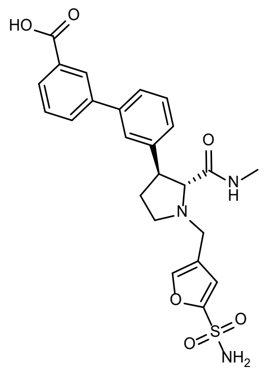

A 20 mL vial was charged with (2*R*,3*S*)-1-(*tert*-butoxycarbonyl)-3-(3-iodophenyl)pyrrolidine-2-carboxylic acid (83.4 mg, 1 eq, 200 μmol) and HATU (91.3 mg, 1.2 eq, 240 μmol). DMF was added followed by $NEt_3$ (101 mg, 139 μL, 5 eq, 1.00 mmol) and the solution was stirred for 15 min. Methanamine, HCl (40.5 mg, 3 eq, 600 μmol) was then added as a solid and reaction was further stirred 3 hr. Solvent was removed under reduced pressure, residue was dissolved as far as possible in $CH_2Cl_2$ (100 mL), washed with 1 M aq. citric acid (2 x 50 mL), dried over $MgSO_4$, filtered, and solvent removed under reduced pressure. *tert*-Butyl (2*R*,3*S*)-3-(3-iodophenyl)-2-(methylcarbamoyl)pyrrolidine-1-carboxylate (63 mg, 146 μmol, 73%) was obtained as a yellow oil after flash chromatography (0–20% MeOH in $CH_2Cl_2$).

A 20 mL vial was charged with *tert*-butyl (2*S*,3*S*)-3-(4-iodophenyl)-2-(methylcarbamoyl)pyrrolidine-1-carboxylate (22 mg, 1 eq, 50 μmol) followed by $CH_2Cl_2$ . TFA (0.11 g, 77 μL, 20 eq, 1.0 mmol) was added by syringe and the mixture was stirred for 30 min. Solvent and TFA were then removed under reduced pressure. Crude residue was taken up in $CH_2Cl_2$ (50 mL) and washed 1M $K_2CO_3$ (3 x 50 mL). Combined aqueous layers were extracted with $CH_2Cl_2$ (3 x 100 mL). Organic portions were combined and dried over $Na_2SO_4$, filtered, and (2*R*,3*S*)-3-(3-iodophenyl)-*N*-methylpyrrolidine-2-carboxamide was isolated under reduced pressure in sufficient purity to carry to next step.

A 20 mL vial was charged with (2*R*,3*S*)-3-(3-iodophenyl)-*N*-methylpyrrolidine-2-carboxamide (132 mg, 1.00 eq, 400 μmol), 4-formylfuran-2-sulfonamide (105 mg, 1.50 eq, 600 μmol), and sodium triacetoxyborohydride (424 mg, 5.00 eq, 2.00 mmol). DCE (10 mL) was added, and the reaction was stirred for 1.5 hr at rt. The reaction solution was diluted with $CH_2Cl_2$ (50 mL) and washed with saturated $NaHCO_3$ (3 x 50 mL). Combined aqueous portions were extracted with $CH_2Cl_2$ (3 x 100 mL). Then combined organics were dried over $MgSO_4$, filtered, and solvent removed under reduced pressure. (2*R*,3*S*)-3-(3-Iodophenyl)-*N*-methyl-1-

((5-sulfamoylfuran-3-yl)methyl)pyrrolidine-2-carboxamide (145 mg, 296 μmol, 74% over two steps) was isolated as a white solid following purification by preparative HPLC.

A 5 mL μW vial was charged with (2*R*,3*S*)-3-(3-iodophenyl)-*N*-methyl-1-((5-sulfamoylfuran-3-yl)methyl)pyrrolidine-2-carboxamide (7.3 mg, 1.00 eq, 15 μmol), 3-boronobenzoic acid (3.1 mg, 1.25 eq, 19 μmol), and Palladium-Xphos (2.4 mg, 0.20 eq, 3.0 μmol). DMF (2.5 mL) was added followed by 1.8 M aq. $K_3PO_4$ (16 mg, 42 μL, 5.00 eq, 75 μmol). The vial was capped and sparged with $N_2$ for 5 min, then the solution was heated at 60 °C for 1 hr under μW irradiation. The solution was diluted with 1:4 MeOH:$CH_2Cl_2$ (20 mL), filtered through celite and rinsed with 1:4 MeOH:$CH_2Cl_2$ (5 x 50 mL). Solvents were removed under reduced pressure and product was purified by preparative HPLC to afford **48** (3.3 mg, 6.9 μmol, 46%) as an off-white solid.

$^1$H NMR (400 MHz, DMSO) δ 8.18 (t, *J* = 1.8 Hz, 1H), 7.95 (dt, *J* = 7.7, 1.4 Hz, 1H), 7.92 – 7.84 (m, 3H), 7.70 (s, 2H), 7.59 (t, *J* = 7.7 Hz, 1H), 7.56 – 7.51 (m, 2H), 7.43 (t, *J* = 7.6 Hz, 1H), 7.28 (dt, *J* = 7.6, 1.4 Hz, 1H), 7.10 (d, *J* = 1.0 Hz, 1H), 3.63 (dd, *J* = 13.7, 1.2 Hz, 1H), 3.47 (d, *J* = 13.7 Hz, 1H), 3.40 (dt, *J* = 9.2, 6.8 Hz, 1H), 3.19 (d, *J* = 7.4 Hz, 1H), 3.09 (ddd, *J* = 9.3, 7.6, 4.4 Hz, 1H), 2.70 – 2.59 (m, 4H), 2.35 – 2.22 (m, 1H), 1.97 – 1.87 (m, 1H). $^{13}$C NMR (101 MHz, DMSO) δ 172.38, 167.55, 151.98, 144.59, 143.36, 140.45, 139.52, 132.69, 130.70, 129.14, 128.23, 127.35, 126.81, 126.06, 124.83, 123.74, 114.24, 74.64, 52.10, 49.22, 47.80, 32.14, 25.57. HRMS (ESI): calculated for $C_{24}H_{26}N_3O_6S$ [M + H]$^+$, 484.1537 found, 484.1552.

(2*R*,3*S*)-3-(3'-Ethoxy-[1,1'-biphenyl]-4-yl)-*N*-methyl-1-((5-sulfamoylfuran-3-yl)methyl)pyrrolidine-2-carboxamide (**49**)

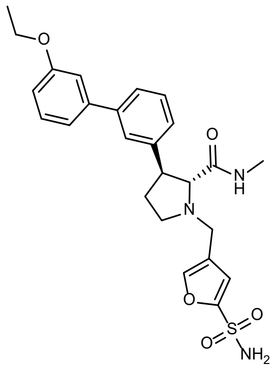

A 5 mL µW vial was charged with (2*R*,3*S*)-3-(3-iodophenyl)-*N*-methyl-1-((5-sulfamoylfuran-3-yl)methyl)pyrrolidine-2-carboxamide [*cf.* synthesis of **48**] (7.3 mg, 1.00 eq, 15 µmol), (3-ethoxyphenyl)boronic acid (3.1 mg, 1.25 eq, 19 µmol), and Palladium-Xphos (2.4 mg, 0.20 eq, 3.0 µmol). DMF (2.5 mL) was added followed by 1.8 M aq. $K_3PO_4$ (16 mg, 42 µL, 5.00 eq, 75 µmol). The vial was capped and sparged with $N_2$ for 5 min. Solution was placed in a µwave reactor and µwaved at 60 °C for 1 hour. Solution was diluted with 1:4 MeOH:$CH_2Cl_2$ (20 mL), filtered through celite and rinsed with flushing 1:4 MeOH:$CH_2Cl_2$ (5 x 50 mL). Solvents were removed under reduced pressure, and product was purified by preparative HPLC to **49** (3.7 mg, 7.7 µmol, 51%) as an off-white solid.

$^1$H NMR (400 MHz, DMSO) δ 7.93 – 7.83 (m, 2H), 7.69 (s, 2H), 7.53 – 7.47 (m, 2H), 7.38 (td, *J* = 7.8, 3.6 Hz, 2H), 7.24 (dt, *J* = 7.6, 1.5 Hz, 1H), 7.20 (dt, *J* = 7.9, 1.1 Hz, 1H), 7.16 (t, *J* = 2.1 Hz, 1H), 7.10 (d, *J* = 1.0 Hz, 1H), 6.94 (ddd, *J* = 8.3, 2.5, 0.9 Hz, 1H), 4.11 (q, *J* = 7.0 Hz, 2H), 3.63 (dd, *J* = 13.6, 1.2 Hz, 1H), 3.46 (d, *J* = 13.7 Hz, 1H), 3.37 (dt, *J* = 9.3, 6.7 Hz, 1H), 3.17 (d, *J* = 7.4 Hz, 1H), 3.09 (ddd, *J* = 9.2, 7.6, 4.3 Hz, 1H), 2.70 – 2.63 (m, 1H), 2.62 (d, *J* = 4.8 Hz, 3H), 2.27 (ddt, *J* = 12.5, 9.2, 7.6 Hz, 1H), 1.96 – 1.86 (m, 1H), 1.37 (t, *J* = 6.9 Hz, 3H). $^{13}$C NMR (101 MHz, DMSO) δ 172.41, 158.99, 151.96, 144.37, 143.34, 141.82, 140.15, 129.97, 128.90, 126.55, 126.08, 124.80, 123.72, 119.01, 114.23, 113.31, 112.85, 74.62, 63.03, 52.08, 49.21, 47.80, 32.15, 25.55, 14.69. HRMS (ESI): calculated for $C_{25}H_{30}N_3O_5S$ $[M + H]^+$, 484.1901 found, 484.1957.

(2*S*,3*R*)-3-(3'-Ethoxy-[1,1'-biphenyl]-4-yl)-*N*-methyl-1-((5-sulfamoylfuran-3-yl)methyl)azetidine-2-carboxamide (**50**)

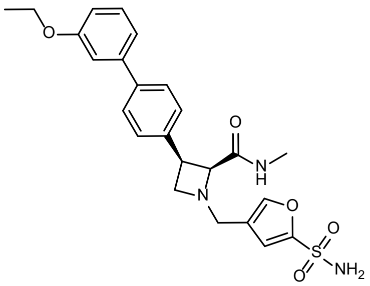

(2*S*,3*R*)-3-(3'-Ethoxy-[1,1'-biphenyl]-4-yl)-*N*-methyl-1-((5-sulfamoylfuran-3-yl)methyl)azetidine-2-carboxamide (7.2 mg, 15.4 µmol, 77%, final step) was prepared as a white solid following an analogous synthesis to **48**. Intermediate yields are provided below:

*tert*-Butyl (2*S*,3*R*)-3-(4-iodophenyl)-2-(methylcarbamoyl)azetidine-1-carboxylate (88%) as yellow oil.

(2*S*,3*R*)-3-(4-Iodophenyl)-*N*-methyl-1-((4-sulfamoylfuran-2-yl)methyl)azetidine-2-carboxamide (66% over 2 steps).

$^1$H NMR (400 MHz, DMSO) δ 7.91 (s, 1H), 7.71 (s, 2H), 7.66 (q, *J* = 4.7 Hz, 1H), 7.59 – 7.54 (m, 2H), 7.49 – 7.44 (m, 2H), 7.36 (t, *J* = 7.9 Hz, 1H), 7.28 (d, *J* = 1.0 Hz, 1H), 7.20 (dt, *J* = 7.8, 1.2 Hz, 1H), 7.17 – 7.13 (m, 1H), 6.91 (ddd, *J* = 8.3, 2.6, 0.9 Hz, 1H), 4.10 (q, *J* = 7.0 Hz, 2H), 3.97 (d, *J* = 9.1 Hz, 1H), 3.82 (td, *J* = 8.4, 3.2 Hz, 1H), 3.63 (dd, *J* = 13.2, 1.2 Hz, 1H), 3.45 (d, *J* = 13.1 Hz, 1H), 3.36 (t, *J* = 7.7 Hz, 1H), 2.35 (d, *J* = 4.8 Hz, 3H), 1.36 (t, *J* = 7.0 Hz, 3H). $^{13}$C NMR (101 MHz, DMSO) δ 169.20, 159.01, 151.99, 143.54, 141.45, 139.20, 138.20, 129.96, 128.73, 126.10, 122.67, 118.73, 114.41, 113.20, 112.52, 69.66, 63.00, 55.72, 50.98, 48.60, 37.86, 25.08, 14.69. HRMS (ESI): calculated for $C_{24}H_{28}N_3O_5S$ $[M + H]^+$, 470.1744 found, 470.1806.

(2*S*,3*S*)-3-(5'-Chloro-2'-methyl-[1,1'-biphenyl]-3-yl)-*N*-methyl-1-((5-sulfamoylfuran-3-yl)methyl)piperidine-2-carboxamide (**51**)

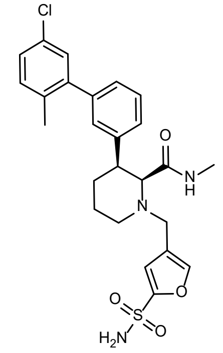

An 8 mL vial was charged with (2*S*,3*S*)-1-(((9H-fluoren-9-yl)methoxy)carbonyl)-3-(3-iodophenyl)piperidine-2-carboxylic acid (14 mg, 1.00 eq, 25 µmol) and HATU (10 mg, 1.10 eq, 28 µmol). $CH_2Cl_2$ (2 mL) was added followed by DIPEA (3.6 mg, 4.8 µL, 1.10 eq, 28 µmol) and the solution was stirred for 15 min. Methanamine, HCl (0.85 mg, 1.10 eq, 28 µmol) was then added as a solid and reaction was further stirred 2 hr. The solution was diluted with $CH_2Cl_2$ (50 mL), washed with 1 M aq. citric acid (2 x 50 mL), dried over $MgSO_4$, filtered, and solvent removed under reduced pressure. Crude product was purified by flash chromatography

(0–20% MeOH in $CH_2Cl_2$) to afford (9*H*-fluoren-9-yl)methyl (2*S*,3*S*)-3-(3-iodophenyl)-2-(methylcarbamoyl)piperidine-1-carboxylate (13 mg, 23 µmol, 93%) as a yellow solid.

An 8 mL vial was charged with (9H-fluoren-9-yl)methyl (2*S*,3*S*)-3-(3-iodophenyl)-2-(methylcarbamoyl)piperidine-1-carboxylate (11 mg, 1 eq, 20 µmol) followed by DMF (2 mL). Piperidine (17 mg, 20 µL, 10 eq, 0.20 mmol) was added and the reaction stirred for 30 min. Solvent was removed under reduced pressure, and product purified by preparative HPLC to afford (2*S*,3*S*)-3-(3-iodophenyl)-*N*-methylpiperidine-2-carboxamide (5.3 mg, 15 µmol, 77%) as a colorless oil.

An 8 mL vial was charged with (2*S*,3*S*)-3-(3-iodophenyl)-*N*-methylpiperidine-2-carboxamide (5.2 mg, 1 eq, 15 µmol), 4-formylfuran-2-sulfonamide (3.9 mg, 1.5 eq, 23 µmol), and sodium triacetoxyborohydride (32 mg, 10 eq, 0.15 mmol). DCE (3 mL) was added, and the reaction stirred for 1.5 hr at rt. The solution was then diluted with $CH_2Cl_2$ (50 mL) and washed with saturated $NaHCO_3$ (3 x 50 mL). Combined aqueous washes were extracted with $CH_2Cl_2$ (3 x 100 mL). Organic portions were combined, dried over $MgSO_4$, filtered, and solvent removed under reduced pressure. Product was purified by preparative HPLC to afford (2*S*,3*S*)-3-(3-iodophenyl)-*N*-methyl-1-((5-sulfamoylfuran-3-yl)methyl)piperidine-2-carboxamide (6.2 mg, 12 µmol, 82%) as a white solid.

A 5 mL µW vial was charged with (2*S*,3*S*)-3-(3-iodophenyl)-*N*-methyl-1-((5-sulfamoylfuran-3-yl)methyl)piperidine-2-carboxamide (5.0 mg, 1 eq, 10 µmol), (5-chloro-2-methylphenyl)boronic acid (1.9 mg, 1.1 eq, 11 µmol), and Pd-Xphos (1.6 mg, 0.2 eq, 2.0 µmol). DMF (2 mL) was added followed by 1.8 M aq. $K_3PO_4$ (11 mg, 28 µL, 5 eq, 50 µmol). The vial was capped and sparged with $N_2$ for 5 min. The reaction was heated at 60 °C for 1 hr. After cooing the solution was diluted with 1:4 MeOH:$CH_2Cl_2$ (20 mL), filtered through celite and rinsed with flushing 1:4 MeOH:$CH_2Cl_2$ (5 x 50 mL). Volatiles were removed under reduced pressure and crude product was purified by preparative HPLC to afford **51** (2.4 mg, 5 µmol, 48%) as an off-white solid.

$^1$H NMR (400 MHz, DMSO) δ 7.71 (d, *J* = 1.0 Hz, 1H), 7.64 (s, 2H), 7.39 (q, *J* = 4.5 Hz, 1H), 7.30 – 7.24 (m, 3H), 7.19 (dt, *J* = 7.8, 1.5 Hz, 1H), 7.10 (dq, *J* = 10.0, 1.4 Hz, 3H), 6.81 (d, *J* = 1.0 Hz, 1H), 3.47 – 3.38 (m, 2H), 3.25 – 3.20 (m, 1H), 3.09 – 2.97 (m, 2H), 2.24 (d, *J* = 4.5 Hz, 3H), 2.11 (s, 3H), 1.74 – 1.67 (m, 1H), 1.56 – 1.45 (m, 2H). $^{13}$C NMR (101 MHz, DMSO) δ 170.80, 152.02, 143.49, 143.37, 142.68, 139.39,

133.92, 132.08, 130.14, 128.83, 128.28, 127.83, 127.29, 126.94, 126.79, 124.06, 113.78, 66.21, 49.17, 45.15, 44.49, 25.07, 24.62, 24.19, 19.54. HRMS (ESI): calculated for $C_{25}H_{29}ClN_3O_4S$ $[M + H]^+$, 502.1562 found, 502.1635.

(2*R*,4*S*)-1-((6,7-Dihydro-5H-pyrrolo[1,2-a]imidazol-2-yl)methyl)-*N*-methyl-4-((4'-sulfamoyl-[1,1'-biphenyl]-3-yl)oxy)pyrrolidine-2-carboxamide (**52**)

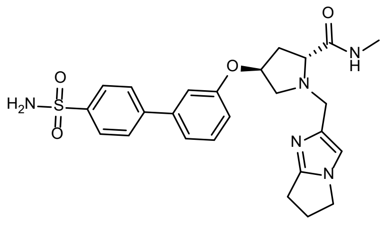

To a 1-dram vial containing (2*R*,4*S*)-1-(*tert*-butoxycarbonyl)-4-(3-iodophenoxy)pyrrolidine-2-carboxylic acid (23 mg, 53 μmol, 1 eq), $NH_3MeCl$ (4.3 mg, 64 μmol, 1.2 eq) and HATU (24 mg, 64 μmol, 1.2 eq) were added. Anhydrous $CH_2Cl_2$ (400 μL) was added, followed by addition of DIPEA (46 μL, 0.27 mmol, 5 eq). The resulting yellow solution was stirred overnight and then diluted with $CH_2Cl_2$ and washed with water (3 x 1 mL), followed by brine (1mL). The organic phase was dried over anhydrous $MgSO_4$ and concentrated under reduced pressure to around 2 mL volume. To the resulting crude mixture was added TFA (300 μL) and after 1.5 hr the solution was concentrated under reduced pressure, then co-evaporated with toluene (1 mL). This crude mixture was used without further purification.

Crude (2*R*,4*S*)-4-(3-iodophenoxy)-*N*-methylpyrrolidine-2-carboxamide (53 μmol, 1 eq), 6,7-dihydro-5H-pyrrolo[1,2-*a*]imidazole-2-carbaldehyde (8.7 mg, 64 μmol, 1.2 eq) and sodium triacetoxyborohydride (22 mg, 0.11 mmol, 2 eq) were dissolved in DCE (500 μL), followed by addition of $NEt_3$ (7.4 μL, 53 μmol, 1 eq). The mixture was stirred at rt overnight, then was washed with sat. aq. $NaHCO_3$ twice and brine once. The combined organic portions were dried over anhydrous $MgSO_4$ and concentrated under reduced pressure. To the resulting residue was added (4-sulfamoylphenyl)boronic acid (16 mg, 80 μmol, 1.5 eq) and XPhos-Pd-G2 (8.3 mg, 11 μmol, 0.2 eq). The vial was capped with a septum and purged 3 times with $N_2$ gas. Then anhydrous DMF (2 mL) was added, followed by 1.8 M aq. $K_3PO_4$ (0.18 mL, 318 μmol, 6 eq). The reaction was stirred at 60 °C for 50 min under μW irradiation, then the resulting solution was diluted with DMSO and purified by preparative HPLC to afford **52** (3.4 mg, 6.9 μmol, 13%) as a white solid.

$^1$H NMR (400 MHz, $CD_3OD$) δ 8.00 – 7.92 (m, 2H), 7.81 – 7.73 (m, 2H), 7.38 (t, *J* = 8.0 Hz, 1H), 7.28 – 7.21 (m, 1H), 7.13 (t, *J* = 2.1 Hz, 1H), 6.94 – 6.88 (m, 2H), 4.95 (dt, *J* = 5.4, 2.8 Hz, 1H), 3.96 (t, *J* = 7.1 Hz, 2H), 3.75 (d, *J* = 13.4 Hz, 1H), 3.63 – 3.41 (m, 3H), 2.88 – 2.77 (m, 3H), 2.75 (s, 3H), 2.57 (p, *J* = 7.4 Hz, 2H), 2.46 – 2.36 (m, 1H), 2.15 (ddd, *J* = 13.6, 9.0, 5.9 Hz, 1H). $^{13}$C NMR (101 MHz, $CD_3OD$) δ 176.86, 159.32, 155.88, 145.92, 143.95, 142.73, 142.41, 131.30, 128.53, 127.73, 121.02, 116.24, 115.61, 115.12, 77.44, 67.31, 59.14, 53.32, 46.17, 38.80, 26.97, 26.05, 23.78. HRMS (ESI): calculated for $C_{25}H_{30}N_5O_4S$ $[M + H]^+$, 496.2013 found, 496.2009.

(*R*)-4'-(6-Acetyl-2-(3-methyl-5,6,7,8-tetrahydroimidazo[1,5-*a*]pyridine-1-carbonyl)-2,6-diazaspiro[3.4]octan-8-yl)-[1,1'-biphenyl]-3-sulfonamide (**53**)

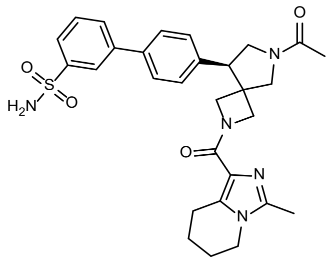

A 1-dram scintillation vial equipped with a magnetic stir bar was charged with **19** (40 mg, 67 µmol, 1.0 eq) and $K_2CO_3$ (18 mg, 130 µmol, 2.0 eq) were dissolved with anhydrous DMF (600 µL, 0.1 M), cooled with an ice-water bath, then thiophenol (7.6 µL, 73 µmol 1.1 eq) was added in one portion. The resulting mixture was stirred for 1 hr in the ice-water bath, then diluted with EtOAc and filtered through a pad of celite. The filtrate was concentrated under reduced pressure, and to the residue was added 3-methyl-5,6,7,8-tetrahydroimidazo[1,5-*a*]pyridine-1-carboxylic acid hydrochloride (17 mg, 80 µmol, 1.2 eq) and HATU (31 mg, 80 µmol, 1.2 eq). DMF (1 mL, 65 mM) and DIPEA (29 µL, 0.17 mmol, 2.5 eq) were added and the resulting yellow solution was stirred overnight. The reaction was diluted with EtOAc and washed three times with a 50% brine solution. The organic phase was dried over anhydrous $MgSO_4$, filtered, and concentrated under reduced pressure. Flash chromatography (0–50% EtOAc in hexanes) afforded *tert*-butyl (*R*)-8-(4-iodophenyl)-2-(3-methyl-5,6,7,8-tetrahydroimidazo[1,5-*a*]pyridine-1-carbonyl)-2,6-diazaspiro[3.4]octane-6-carboxylate (19 mg, 33 µmol, 49% over 2 steps).

To a solution of *tert*-butyl (*R*)-8-(4-iodophenyl)-2-(3-methyl-5,6,7,8-tetrahydroimidazo[1,5-*a*]pyridine-1-carbonyl)-2,6-diazaspiro[3.4]octane-6-carboxylate (19 mg, 33 µmol, 1 eq) in $CH_2Cl_2$ (400 µL), TFA (100 µL) was added, and the resulting solution was stirred for 1 hr at rt. The solution was concentrated under reduced pressure, and co-evaporated with MeOH three times. To the residue was added in $CH_2Cl_2$ (350 µL), DIPEA (17 µL, 99 µmol, 3 eq) and $Ac_2O$ (3.7 µL, 40 µmol, 1.2 eq). After stirring for 5 hr at rt the solution was diluted with $CH_2Cl_2$, washed 3 times with water and solvent was removed into a 2–5 mL µW vial. 3-(4,4,5,5-Tetramethyl-1,3,2-dioxaborolan-2-yl)benzenesulfonamide (14 mg, 49 µmol, 1.5 eq) and XPhos-Pd-G2 (5.2 mg, 6.6 µmol, 0.2 eq) were added, the vial capped with a septum and purged 3 times with $N_2$ gas. Anhydrous DMF (0.5 mL) was added, followed by 1.8 M aq. $K_3PO_4$ (0.12 mL, 215 µmol, 6.5 eq). The reaction was stirred at 60 °C for 50 min under microwave irradiation. Upon cooling the dark brown suspension was diluted with water and extracted five times with $CH_2Cl_2$. The organic phase was dried over $MgSO_4$, and solvents were removed under reduced pressure. The residue was purified by preparative HPLC to afford **53** (4.9 mg, 8.9 µmol, 27% over 3 steps) as a colorless oil.

$^1$H NMR (400 MHz, $CD_3OD$) δ 8.14 (s, 1H), 7.86 (dd, *J* = 14.6, 7.8 Hz, 2H), 7.71 – 7.64 (m, 2H), 7.64 – 7.57 (m, 1H), 7.47 – 7.38 (m, 2H), 4.63 – 4.49 (m, 1H), 4.43 – 3.71 (m, 10H), 3.62 (dt, *J* = 32.3, 7.0 Hz, 1H), 2.94 – 2.74 (m, 2H), 2.26 (d, *J* = 27.4 Hz, 3H), 2.15 (d, *J* = 7.3 Hz, 3H), 1.88 (d, *J* = 23.7 Hz, 2H), 1.75 – 1.53 (m, 2H). $^{13}$C NMR (101 MHz, $CD_3OD$) δ 170.90, 165.41, 144.42, 143.46, 141.25, 137.67, 137.35, 135.09, 130.02, 129.28, 128.67, 128.60, 126.99, 124.52, 123.96, 54.73, 50.72, 48.52, 42.52, 22.15, 20.70, 20.52, 19.04, 10.96. HRMS (ESI): calculated for $C_{29}H_{34}N_5O_4S$ $[M + H]^+$, 548.2326 found, 548.2425.

4'-((8*S*)-6-Acetyl-2-(2-(3-cyanophenoxy)propanoyl)-2,6-diazaspiro[3.4]octan-8-yl)-[1,1'-biphenyl]-4-sulfonamide (**54**)

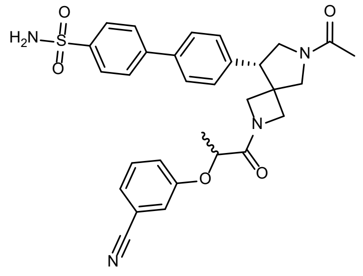

**20** (20 mg, 33 µmol, 1 eq) and $K_2CO_3$ (9.2 mg, 67 µmol, 2 eq) were suspended in anhydrous DMF (300 µL) and cooled to 0 °C in an ice bath, followed by addition of thiophenol (3.8 µL, 37 µmol, 1.1 eq). The resulting mixture was stirred for 1 hr at 0 °C, then diluted with EtOAc and filtered through a pad of celite. The filtrate was concentrated under reduced pressure and 2-(3-cyanophenoxy)propanoic acid (7.6 mg, 40 µmol, 1.2 eq) and PyBOP (21 mg, 40 µmol, 1.2 eq) were added. The vial was purged with $N_2$, followed by addition of DMF (300 µL) to dissolve the reactants. DIPEA (17 µL, 99 µmol, 3 eq) was added, and the solution was stirred at rt overnight. The solution was diluted with EtOAc and washed with water (1 mL) 3 times, followed by brine (1 mL). Combined organic portions were dried over anhydrous $MgSO_4$ and concentrated under reduced pressure to approx 2 mL volume. The product was purified by flash chromatography (EtOAc in hexanes) to afford *tert*-butyl (8*S*)-2-(2-(3-cyanophenoxy)propanoyl)-8-(4-iodophenyl)-2,6-diazaspiro[3.4]octane-6-carboxylate (7.8 mg, 13 µmol, 40% over 2 steps).

*tert*-Butyl (8*S*)-2-(2-(3-cyanophenoxy)propanoyl)-8-(4-iodophenyl)-2,6-diazaspiro[3.4]octane-6-carboxylate (7.0 mg, 12 µmol, 1 eq) was dissolved in $CH_2Cl_2$ (0.5 mL), and TFA (100 µL) was added. After 1.5 hr the solution was concentrated under reduced pressure, then co-evaporated to constant mass with toluene (1 mL). The resulting crude mixture was dissolved in $CH_2Cl_2$ (0.5 mL), followed by addition of $Ac_2O$ (1.2 µL, 13 µmol, 1.05 eq), $NEt_3$ (5.0 µL, 36 µmol, 3 eq) and pyridine (0.97 µL, 12 µmol, 1.0 eq). The solution was stirred at rt overnight and concentrated under reduced pressure. The resulting residue was added to a 2 mL µW vial along with (4-sulfamoylphenyl)boronic acid (3.6 mg, 18 µmol, 1.5 eq) and XPhos-Pd-G2 (1.9 mg, 2.4 µmol, 0.2 eq). The vial was capped with a septum and purged 3 times with $N_2$. Anhydrous DMF (2 mL) was added, followed by addition of 1.8 M aq. $K_3PO_4$ (40 µL, 72 µmol, 6 eq) and the reaction was stirred at 60 °C for 2 hr under µW irradiation. The resulting solution was diluted with DMSO and purified by preparative HPLC to afford **54** (3.5 mg, 6.2 µmol, 52%) as a white solid.

$^1H$ NMR (400 MHz, $CD_3OD$) δ 8.05 – 7.93 (m, 2H), 7.85 – 7.69 (m, 3H), 7.62 – 7.55 (m, 1H), 7.55 – 7.40 (m, 1H), 7.41 – 7.18 (m, 3H), 7.16 – 6.95 (m, 2H), 4.78 – 3.40 (m, 10H), 2.18 – 2.06 (m, 3H), 1.49 (dd, *J* = 6.7, 5.6 Hz, 1H), 1.44 – 1.36 (m, 2H), 1.20 (dd, *J* = 6.7, 5.3 Hz, 1H). HRMS (ESI): calculated for $C_{30}H_{31}N_4O_5S$ $[M + H]^+$, 559.2010 found, 559.2082.

3-(3'-((2S,3R)-2-(Methylcarbamoyl)-1-(4-sulfamoylbenzoyl)pyrrolidin-3-yl)-[1,1'-biphenyl]-3-yl)propanoic acid (**55**)

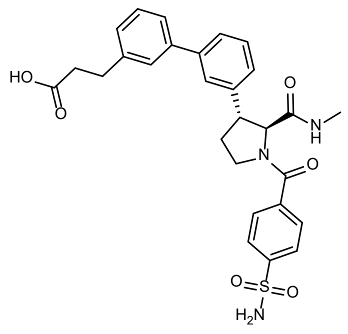

A 20 mL vial was charged with (2*S*,3*R*)-1-(*tert*-butoxycarbonyl)-3-(3-iodophenyl)pyrrolidine-2-carboxylic acid (209 mg, 1.00 eq, 500 μmol) and HATU (228 mg, 1.20 eq, 600 μmol). DMF (5 mL) was added followed by $NEt_3$ (126 mg, 174 μL, 2.50 eq, 1.25 mmol) and the solution was stirred for 15 min. Methanamine, HCl (67.5 mg, 2.00 eq, 1.00 mmol) was then added as a solid and the reaction was further stirred for 3 hr. The mixture was concentrated under reduced pressure and the resulting residue was dissolved as far as possible in $CH_2Cl_2$ (100 mL), washed with 1 M aq. citric acid (2 x 100 mL), dried over $MgSO_4$, filtered, and had solvent removed under reduced pressure. Crude product was purified by flash chromatography (0–20% MeOH in $CH_2Cl_2$) to afford *tert*-butyl (2*S*,3*R*)-3-(3-iodophenyl)-2-(methylcarbamoyl)pyrrolidine-1-carboxylate (153 mg, 355 μmol, 71%) as a yellow oil.

A 20 mL vial was charged with *tert*-butyl (2*S*,3*R*)-3-(3-iodophenyl)-2-(methylcarbamoyl)pyrrolidine-1-carboxylate (116 mg, 1.00 eq, 270 μmol) followed by $CH_2Cl_2$ (10 mL). TFA (308 mg, 208 μL, 10.0 eq, 2.70 mmol) was added via syringe and reaction stirred for 30 min. Volatiles were removed under reduced pressure. Crude was taken up in $CH_2Cl_2$ (50 mL) and washed 1 M $K_2CO_3$ (3 x 50 mL). Aqueous portions were combined and extracted with $CH_2Cl_2$ (3 x 100 mL). Organics were then combined, dried over $Na_2SO_4$, filtered, and the residue was used directly in the next step without further purification.

A 20 mL vial was charged with 4-sulfamoylbenzoic acid (7.04 mg, 1.05 eq, 35.0 μmol) and CDI (5.94 mg, 1.10 eq, 36.6 μmol). DMF (5 mL) was added, and the solution was stirred for 15 min. (2*S*,3*R*)-3-(3-iodophenyl)-*N*-methylpyrrolidine-2-carboxamide (11.0 mg, 1.00 eq, 33.3 μmol) was then added and the reaction was further stirred for 3 hr. The mixture was diluted with EtOAc (50 mL), washed with 0.5 M aq.

citric acid (2 x 50 mL), dried over $MgSO_4$, filtered, and solvent was removed under reduced pressure. Purification was performed by flash chromatography (0–20% MeOH in $CH_2Cl_2$) to afford (*2S,3R*)-3-(3-iodophenyl)-*N*-methyl-1-(4-sulfamoylbenzoyl)pyrrolidine-2-carboxamide (42%) as a yellow solid.

A µW vial was charged with (2*S*,3*R*)-3-(3-iodophenyl)-*N*-methyl-1-(4-sulfamoylbenzoyl)pyrrolidine-2-carboxamide (26 mg, 1.00 eq, 50 µmol), 3-(3-boronophenyl)propanoic acid (12 mg, 1.25 eq, 63 µmol), and Palladium-Xphos (7.9 mg, 0.20 eq, 10 µmol). DMF (5 mL) was added followed by 1.8 M aq. $K_3PO_4$ (53 mg, 0.14 mL, 5.00 eq, 0.25 mmol). The vial was capped and sparged with $N_2$ for 5 min. The reaction was heated at 60 °C for 1 hr by µW irradiation. Once cooled the solution was diluted with 1:4 MeOH:$CH_2Cl_2$ (20 mL), filtered through celite and rinsed with 1:4 MeOH:$CH_2Cl_2$ (5 x 50 mL). Solvent was removed under reduced pressure, and crude product was purified by preparative HPLC to afford **55** (20 mg, 37 µmol, 74%) as a white solid.

$^1$H NMR (400 MHz, DMSO) δ 8.00 – 7.88 (m, 2H), 7.80 – 7.68 (m, 2H), 7.66 – 7.12 (m, 10H), 4.37 (dd, *J* = 98.1, 6.5 Hz, 1H), 3.93 – 3.43 (m, 4H), 2.89 (t, *J* = 7.6 Hz, 2H), 2.61 – 2.57 (m, 3H), 2.37 – 2.21 (m, 2H), 2.16 – 2.04 (m, 1H). $^{13}$C NMR (101 MHz, DMSO) δ 173.92, 171.01, 170.72, 167.40, 145.28, 144.78, 141.72, 141.63, 140.58, 140.20, 139.36, 129.09, 128.85, 127.92, 127.43, 127.20, 126.89, 126.39, 126.09, 125.77, 125.67, 125.45, 125.35, 124.51, 68.04, 66.55, 49.99, 49.59, 48.28, 46.54, 35.61, 33.36, 30.60, 29.01, 28.80, 28.69, 25.67, 25.46. HRMS (ESI): calculated for $C_{28}H_{30}N_3O_6S$ $[M + H]^+$, 536.1850 found, 536.1871.

4-Cyano-*N*-(((4*S*,5*R*)-2-((*R*)-1-hydroxypropan-2-yl)-4-methyl-1,1-dioxido-8-(3-sulfamoylphenyl)-2,3,4,5-tetrahydrobenzo[*b*][1,4,5]oxathiazocin-5-yl)methyl)-*N*-methylbenzenesulfonamide (**56**)

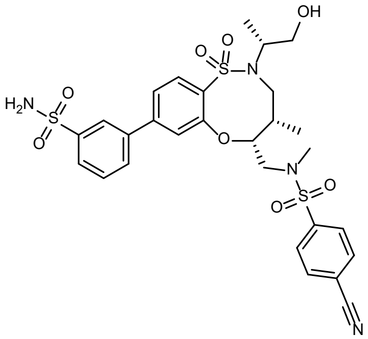

A 5 mL cone-shaped µW vial equipped with a magnetic stir bar was charged with (9H-fluoren-9-yl)methyl (((4*S*,5*R*)-8-bromo-2-((*R*)-1-hydroxypropan-2-yl)-4-methyl-1,1-dioxido-2,3,4,5-tetrahydrobenzo[*b*][1,4,5]oxathiazocin-5-yl)methyl)(methyl)carbamate (44.1 mg, 1.00 eq, 70.0 µmol), (3-sulfamoylphenyl)boronic acid (21.1 mg, 1.5 eq, 105 µmol), XPhos-Pd-G2 (11.0 mg, 0.20 eq, 14.0 µmol), dry DMF (0.70 mL) and aq. $K_3PO_4$ (1.8 M, 0.12 mL, 0.22 mmol, 3.0 eq). The mixture was purged with Ar for 2 min, the vial was sealed, and the reaction mixture was heated to 60 °C for 75 min using µW irradiation. The reaction mixture was concentrated under reduced pressure and then purified by flash chromatography (0–20% MeOH [with 1% $NH_4OH$] in $CH_2Cl_2$) to afford 3-((4*S*,5*R*)-2-((*R*)-1-hydroxypropan-2-yl)-4-methyl-5-((methylamino)methyl)-1,1-dioxido-2,3,4,5-tetrahydrobenzo[*b*][1,4,5]oxathiazocin-8-yl)benzenesulfonamide (29.5 mg, 61.0 µmol, 87%) as a white solid.

A 1 dram scintillation vial equipped with a magnetic stir bar was charged with 3-((4*S*,5*R*)-2-((*R*)-1-hydroxypropan-2-yl)-4-methyl-5-((methylamino)methyl)-1,1-dioxido-2,3,4,5-tetrahydrobenzo[*b*][1,4,5]oxathiazocin-8-yl)benzenesulfonamide (10 mM in THF, 1.0 mL, 1.0 eq, 10 µmol), $NEt_3$ (2.0 mg, 2.8 µL, 2.0 eq, 20 µmol), 4-dimethylaminopyridine (0.1 mg, 0.1 eq, 1 µmol) and 4-cyanobenzenesulfonyl chloride (2.4 mg, 1.20 eq, 12 µmol). The reaction mixture was stirred for 40 min. $CH_2Cl_2$ and silica gel were added, and the suspension was concentrated under reduced pressure. The material was purified by flash chromatography (0–10% MeOH in $CH_2Cl_2$) affording **56** (5.1 mg, 7.9 µmol, 78%) as a colorless oil.

$^1$H NMR (400 MHz, $CD_3OD$) δ 8.19 (t, *J* = 1.8 Hz, 1H), 8.07 – 7.80 (m, 7H), 7.71 – 7.60 (m, 2H), 7.57 (dd, *J* = 8.2, 1.8 Hz, 1H), 4.68 (dt, *J* = 8.2, 3.8 Hz, 1H), 4.12 (dd, *J* = 15.8, 10.0 Hz, 1H), 4.01 (dq, *J* = 13.9, 6.7 Hz, 1H), 3.84 (dd, *J* = 14.0, 8.5 Hz, 1H), 3.59 – 3.47 (m, 2H), 3.38 (dd, *J* = 15.7, 5.4 Hz, 1H), 3.11 (dd, *J* = 14.0, 3.7 Hz, 1H), 2.75 (s, 3H), 2.32 – 2.20 (m, 1H), 1.23 (d, *J* = 6.9 Hz, 3H), 1.04 (d, *J* = 6.9 Hz, 3H). $^{13}$C NMR (101 MHz, $CD_3OD$) δ 158.29, 146.05, 145.91, 142.26, 141.12, 138.49, 134.35, 132.00, 130.88, 129.75, 129.57, 126.95, 125.81, 124.67, 124.00, 118.37, 117.60, 84.74, 66.03, 58.71, 53.63, 36.33, 36.25, 15.91, 12.65. HRMS (ESI): calculated for $C_{28}H_{33}N_4O_8S_3$ $[M + H]^+$, 649.1455 found, 649.1464.

5-(Furan-2-yl)-*N*-(((4*S*,5*R*)-2-((*S*)-1-hydroxypropan-2-yl)-4-methyl-1,1-dioxido-8-(4-sulfamoylphenyl)-2,3,4,5-tetrahydrobenzo[*b*][1,4,5]oxathiazocin-5-yl)methyl)-*N*-methylisoxazole-4-carboxamide (**57**)

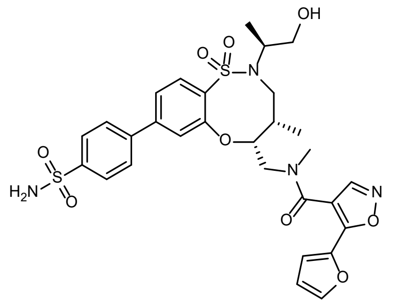

A 5 mL cone-shaped µW vial equipped with magnetic stir bar was charged with (9H-fluoren-9-yl)methyl (((4*S*,5*R*)-8-bromo-2-((*S*)-1-hydroxypropan-2-yl)-4-methyl-1,1-dioxido-2,3,4,5-tetrahydrobenzo[*b*][1,4,5]oxathiazocin-5-yl)methyl)(methyl)carbamate (52.5 mg, 1.00 eq, 83.4 µmol), (4-sulfamoylphenyl)boronic acid (25.1 mg, 1.5 eq, 125 µmol), XPhos-Pd-G2 (13.1 mg, 0.20 eq, 16.7 µmol), aq. $K_3PO_4$ (1.8 M, 0.14 mL, 0.25 mmol, 3.0 eq) and dry DMF (0.83 mL). The reaction mixture was sparged with Ar for 3 min, the vial was sealed, and the reaction mixture was heated to 60 °C for 80 min using µW irradiation. The orange reaction mixture was concentrated under reduced pressure and absorbed on silica gel. The material was then purified by flash chromatography (0–20% MeOH [with 1% $NH_4OH$] in $CH_2Cl_2$) to afford 4-((4*S*,5*R*)-2-((*S*)-1-hydroxypropan-2-yl)-4-methyl-5-((methylamino)methyl)-1,1-dioxido-2,3,4,5-tetrahydrobenzo[*b*][1,4,5]oxathiazocin-8-yl)benzenesulfonamide (14.8 mg, 30.6 µmol, 37%) as a cloudy white oil.

A 2 dram scintillation vial equipped with a magnetic stir bar was charged with 4-((4*S*,5*R*)-2-((*S*)-1-hydroxypropan-2-yl)-4-methyl-5-((methylamino)methyl)-1,1-dioxido-2,3,4,5-tetrahydrobenzo[*b*][1,4,5]oxathiazocin-8-yl)benzenesulfonamide (14.8 mg, 1.00 eq, 30.6 µmol) at rt. 5-(Furan-2-yl)isoxazole-4-carboxylic acid (0.22 M in DMF, 0.17 mL, 1.2 eq, 37 µmol), DIPEA (11.2 µL, 2.1 eq, 64.3 µmol) and PyBOP (0.22 M in DMF, 0.17 mL, 1.2 eq, 37 µmol) were successively added and the brown solution was stirred for 1 hr. EtOAc and water (each ca 30 mL) were added and the aqueous phase was extracted twice with EtOAc. The organic layers were washed two more times with water, then they were combined, dried over $Na_2SO_4$ and concentrated under reduced pressure to afford a brown oil.

The crude material was purified by preparative HPLC to afford **57** (7.9 mg, 12 µmol, 40%) as a white powder.

$^{1}H$ NMR (400 MHz, $CD_3OD$) δ 8.75 (s, 0.45H), 8.45 (s, 0.55H), 8.07 – 7.83 (m, 4H), 7.76 (d, *J* = 8.2 Hz, 1H), 7.62 – 7.53 (m, 2.55H), 7.40 (s, 0.45H), 7.07 (dd, *J* = 22.4, 3.5 Hz, 1H), 6.59 – 6.49 (m, 1H), 4.94 – 4.88 (m, 0.55H), 4.79 – 4.71 (m, 0.45H), 4.17 (dd, *J* = 15.9, 8.3 Hz, 0.55H), 4.06 (dd, *J* = 14.0, 4.6 Hz, 0.55H), 3.94 (dd, *J* = 14.1, 8.5 Hz, 2H), 3.88 – 3.82 (m, 0.45H), 3.82 – 3.71 (m, 1H), 3.63 – 3.52 (m, 1H), 3.52 – 3.44 (m, 0.55H), 3.36 (dd, *J* = 15.3, 3.1 Hz, 0.45H), 3.23 – 3.12 (m, 2H), 3.08 (s, 1.55H), 2.42 – 2.30 (m, 0.55H), 2.27 – 2.17 (m, 0.45H), 1.17 (d, *J* = 7.1 Hz, 1.7H), 1.13 (d, *J* = 7.1 Hz, 3H), 0.85 (d, *J* = 7.1 Hz, 1.3H). $^{13}C$ NMR (101 MHz, $CD_3OD$) δ 165.43, 164.92, 159.81, 159.19, 157.63, 156.99, 151.01, 150.82, 147.09, 146.94, 146.44, 146.27, 145.02, 145.01, 144.99, 143.73, 143.66, 143.19, 143.05, 138.58, 129.70, 129.06, 128.78, 127.89, 124.43, 124.24, 123.70, 123.60, 114.70, 114.53, 113.38, 113.23, 110.71, 86.08, 85.16, 65.27, 65.12, 59.06, 58.85, 52.63, 49.82, 49.00, 48.79, 39.25, 36.75, 36.17, 34.10, 16.89, 13.68, 13.53. HRMS (ESI): calculated for $C_{29}H_{33}N_4O_9S_2$ $[M + H]^+$, 645.1683 found, 645.1699.

*N*-(((4*S*,5*S*)-2-((*R*)-1-Hydroxypropan-2-yl)-4-methyl-1,1-dioxido-8-(4-sulfamoylphenyl)-2,3,4,5-tetrahydrobenzo[*b*][1,4,5]oxathiazocin-5-yl)methyl)-*N*-methyl-5-(methylsulfonyl)furan-3-carboxamide (**58**)

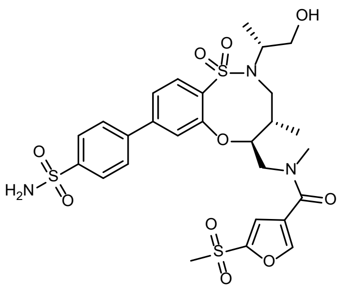

A 2-dram scintillation vial equipped with a magnetic stir bar was charged with (4*S*,5*S*)-8-bromo-2-((*R*)-1-hydroxypropan-2-yl)-4-methyl-5-((methylamino)methyl)-2,3,4,5-tetrahydrobenzo[*b*][1,4,5]oxathiazocine 1,1-dioxide [*cf*. synthesis of **36**] (29.2 mg, 1 eq, 71.7 µmol), dry DMF (0.72 mL), DIPEA (26 µL, 2.1 eq, 0.15 mmol), 5-(methylsulfonyl)furan-3-carboxylic acid (15.0 mg, 1.1 eq, 78.9 µmol) and PyBOP (41.0 mg, 1.1 eq, 78.9 µmol). The light brown solution was stirred for 18 min at rt. EtOAc and water–brine (2:1) were added and the aqueous phase was extracted twice with EtOAc. The organic layers were washed with more water–brine (2:1) and water. The organic layers were then combined and dried over $Na_2SO_4$.

Concentration under reduced pressure afforded the crude material, which was purified by flash chromatography (0–10% MeOH in $CH_2Cl_2$) to afford *N*-(((4*S*,5*S*)-8-bromo-2-((*R*)-1-hydroxypropan-2-yl)-4-methyl-1,1-dioxido-2,3,4,5-tetrahydrobenzo[*b*][1,4,5]oxathiazocin-5-yl)methyl)-*N*-methyl-5-(methylsulfonyl)furan-3-carboxamide (32.5 mg, 56.1 μmol, 78%) as a colorless oil.

A 5 mL cone-shaped μW vial equipped with a magnetic stir bar was charged with *N*-(((4*S*,5*S*)-8-bromo-2-((*R*)-1-hydroxypropan-2-yl)-4-methyl-1,1-dioxido-2,3,4,5-tetrahydrobenzo[*b*][1,4,5]oxathiazocin-5-yl)methyl)-*N*-methyl-5-(methylsulfonyl)furan-3-carboxamide (32.4 mg, 1.00 eq, 55.9 μmol) in dry DMF (0.56 mL) at rt. (4-Sulfamoylphenyl)boronic acid (16.9 mg, 1.5 eq, 83.9 μmol), XPhos-Pd-G2 (8.8 mg, 20 mol%, 11 μmol) and 1.8 M aq. $K_3PO_4$ (0.19 mL, 6.0 eq, 0.34 mmol) were successively added and the reaction mixture was purged with Ar for 3 min. The vial was sealed, and the reaction mixture was heated to 60 °C for 1 hr using μW irradiation. EtOAc was added to the dark orange, cloudy solution and the mixture was filtered over cotton. It was concentrated under reduced pressure to afford an orange oil. The material was purified by flash chromatography (0–10% MeOH in $CH_2Cl_2$) affording a yellow solid. The material was further purified by preparative HPLC to give **58** (23.8 mg, 36.3 μmol, 65 %) as a white foam.

$^1H$ NMR (400 MHz, $CDCl_3$) δ 8.34 (s, 0.45H), 8.03 – 7.75 (m, 4.55H), 7.65 (d, *J* = 8.2 Hz, 1H), 7.58 (td, *J* = 8.6, 1.8 Hz, 1H), 7.46 (s, 0.45H), 7.31 (d, *J* = 1.9 Hz, 0.55H), 7.20 (d, *J* = 1.8 Hz, 0.45H), 7.14 (s, 0.55H), 4.91 – 4.85 (m, 0.55H), 4.77 – 4.65 (m, 0.45H), 4.59 – 4.47 (m, 0.55H), 4.15 (dd, *J* = 14.8, 11.0 Hz, 0.45H), 3.86 – 3.73 (m, 2H), 3.73 – 3.55 (m, 3.45H), 3.43 (dd, *J* = 14.1, 2.5 Hz, 0.55H), 3.25 (s, 1.35H), 3.11 (s, 1.65H), 3.07 (s, 1.65H), 2.89 (s, 1.35H), 2.42 – 2.26 (m, 1H), 1.27 (d, *J* = 6.6 Hz, 3H), 0.98 (d, *J* = 6.9 Hz, 1.65H), 0.91 (d, *J* = 6.9 Hz, 1.35H). $^{13}C$ NMR (101 MHz, $CDCl_3$) δ 165.87, 165.42, 157.28, 156.61, 151.91, 151.46, 149.96, 149.30, 146.49, 145.72, 145.03, 144.93, 143.50, 143.04, 135.69, 135.53, 131.35, 131.30, 128.93, 128.42, 127.98, 124.20, 123.96, 123.84, 123.66, 123.39, 122.87, 118.37, 118.01, 85.13, 84.48, 65.62, 65.51, 58.59, 55.99, 52.98, 51.26, 43.44, 43.41, 38.47, 37.90, 37.87, 34.10, 15.94, 15.86, 15.05, 14.70. HRMS (ESI): calculated for $C_{27}H_{34}N_3O_{10}S_3$ $[M + H]^+$, 656.1401 found, 656.1414.

5-(Furan-2-yl)-*N*-(((4*R*,5*R*)-2-((*S*)-1-hydroxypropan-2-yl)-4-methyl-1,1-dioxido-8-(4-sulfamoylphenyl)-2,3,4,5-tetrahydrobenzo[*b*][1,4,5]oxathiazocin-5-yl)methyl)-*N*-methylisoxazole-4-carboxamide (**59**)

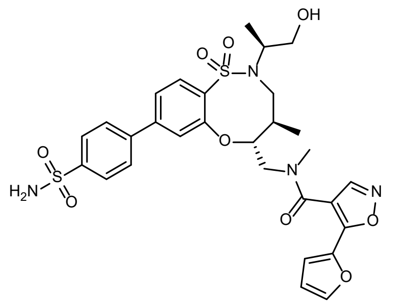

In a µW vial were combined (9H-fluoren-9-yl)methyl (((4*S*,5*R*)-8-bromo-2-((*S*)-1-hydroxypropan-2-yl)-4-methyl-1,1-dioxido-2,3,4,5-tetrahydrobenzo[*b*][1,4,5]oxathiazocin-5-yl)methyl)(methyl)carbamate (49 mg, 1 eq, 78 µmol), (4-sulfamoylphenyl)boronic acid (23 mg, 1.5 eq, 0.12 mmol), XPhos-Pd-G2 (12 mg, 0.20 eq, 16 µmol) and DMF (0.85 mL) to give an off-white cloudy mixture. The vial was capped with a rubber septum and 1.8 M aq. $K_3PO_4$ (0.13 mL, 3.0 eq, 0.23 mmol) was added. The mixture was degassed by evacuating at high vacuum and backfilling with $N_2$ five times; the vial was then capped and heated to 60 °C for 1 hr under µW irradiation. Piperidine (0.2 mL) was added, and the orange reaction mixture stirred at rt 30 min. The vial was then opened, and the reaction mixture transferred to a 100 mL round bottom flask, rinsing with EtOAc and water. Volatiles were removed under reduced pressure and the crude residue was purified by flash chromatography (0–30% MeOH [with 1% $NH_4OH$] in $CH_2Cl_2$), giving 4-((4*R*,5*R*)-2-((*S*)-1-hydroxypropan-2-yl)-4-methyl-5-((methylamino)methyl)-1,1-dioxido-2,3,4,5-tetrahydrobenzo[*b*][1,4,5]oxathiazocin-8-yl)benzenesulfonamide (28 mg, 58 µmol, 74%) as a colorless residue.

In a 20 mL glass scintillation vial 4-((4*R*,5*R*)-2-((*S*)-1-hydroxypropan-2-yl)-4-methyl-5-((methylamino)methyl)-1,1-dioxido-2,3,4,5-tetrahydrobenzo[*b*][1,4,5]oxathiazocin-8-yl)benzenesulfonamide (28 mg, 58 µmol, 1 eq) was stirred with DIPEA (0.021 mL, 2.1 eq, 0.12 mmol), 5-(furan-2-yl)isoxazole-4-carboxylic acid (12 mg, 1.20 eq, 69 µmol) and PyBOP (36 mg, 1.20 eq, 69 µmol) in DMF (0.2 mL). Additional DMF (0.4 mL) was added, giving a dark brown solution, the vial was sealed under $N_2$, and the reaction was stirred for 3 hr. The reaction was then quenched by diluting with additional DMF (0.6 mL), filtered through a micron filter and purified by preparative HPLC to give **59** (8.7 mg, 13.5 µmol, 23%) as a colorless solid.

$^{1}H$ NMR (400 MHz, $CD_3OD$, 330 K) δ 8.16 – 7.64 (m, 7H), 7.52 (dd, *J* = 8.3, 1.8 Hz, 1H), 7.42 – 7.17 (m, 2H), 6.61 (dd, *J* = 3.7, 1.7 Hz, 1H), 4.84 (td, *J* = 9.8, 2.7 Hz, 1H), 4.45 – 4.31 (m, 1H), 3.86 – 3.55 (m, 6H), 3.15 (s, 3H), 2.41 – 2.27 (m, 1H), 1.27 (d, *J* = 6.7 Hz, 3H), 0.96 (d, *J* = 6.9 Hz, 3H). $^{13}C$ NMR (151 MHz, DMSO) δ 163.36, 155.73, 144.39, 144.34, 141.39, 134.65, 129.92, 127.91, 126.66, 123.06, 122.18, 113.06, 63.90, 57.42, 50.22, 49.07, 40.51, 36.51, 16.24, 15.17. LRMS (ESI): calculated for $C_{29}H_{33}N_4O_9S_2$ $[M + H]^+$, 645.17 found, 645.23.

(*R*,*E*)-4-(4-(2-Cyclohexylvinyl)phenyl)-2-((3,5-dichlorophenyl)sulfonyl)-3-(2-hydroxyethyl)-*N*-methyl-2,3-dihydro-1H-pyrrolo[3,4-*c*]pyridine-6-carboxamide (**60**)

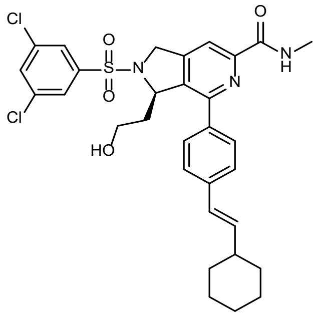

A 5 mL μW vial was charged with ethyl (*R*)-4-(4-bromophenyl)-2-((*R*)-tert-butylsulfinyl)-3-(2-hydroxyethyl)-2,3-dihydro-1H-pyrrolo[3,4-*c*]pyridine-6-carboxylate (39.6 mg, 1.00 eq, 80.0 μmol), (*E*)-(2-cyclohexylvinyl)boronic acid (19.7 mg, 1.60 eq, 128 μmol), and Palladium-Xphos (12.6 mg, 0.20 eq, 16.0 μmol). DMF (2.0 mL) was then added to the vial followed by 1.8 M aq. $K_3PO_4$ (84.9 mg, 222 μL, 5.00 eq, 400 μmol). The reaction vial was sparged with $N_2$ and capped before heating at 60 °C for 75 min. After the solution had cooled, DMF was removed under reduced pressure and the crude material was taken up in $CH_2Cl_2$ (40 mL). The organic layer was washed $H_2O$ (40 mL) and brine (40 mL). After removal of solvent under reduced pressure the crude material was purified by flash chromatography (0–100% EtOAc in hexanes). Ethyl (*R*)-2-((*S*)-*tert*-butylsulfinyl)-4-(4-((*E*)-2-cyclohexylvinyl) phenyl)-3-(2-hydroxyethyl)-2,3-dihydro-1H-pyrrolo[3,4-*c*]pyridine-6-carboxylate (35 mg, 66.4 μmol, 83%) was isolated as a yellow solid.

Ethyl (*R*)-2-((*R*)-*tert*-butylsulfinyl)-4-(4-((*E*)-2-cyclohexylvinyl)phenyl)-3-(2-hydroxyethyl)-2,3-dihydro-1H-pyrrolo[3,4-*c*]pyridine-6-carboxylate (40.0 mg, 1.00 eq, 76.2 μmol) was added to an 8 mL vial and dissolved in EtOH (1.5 mL). HCl (13.9 mg, 95.3 μL, 4 M in 1,4-dioxane, 5.00 eq, 381 μmol) was then added and the solution was stirred at rt for 30 min. Volatiles were removed under reduced pressure, the residue was suspended in $Et_2O$ (30 mL), filtered, and washed with $Et_2O$ (2 x 30 mL). Ethyl (*R*,*E*)-4-(4-(2-cyclohexylvinyl)phenyl)-3-(2-hydroxyethyl)-2,3-dihydro-1H-pyrrolo[3,4-*c*]pyridine-6-carboxylate, HCl (34.8 mg, 76.2 µmol, 100%) was collected as an off-white solid.

An 8 mL vial was charged with 3,5-dichlorobenzenesulfonyl chloride (28 mg, 1.50 eq, 114 µmol) and a small stir bar. The vial was then capped with a septum and purged with $N_2$. Dry $CH_2Cl_2$ (4 mL) was added, and the solution was chilled to 0 °C in an ice-water bath. In a separate vial, ethyl (*R,E*)-4-(4-(2-cyclohexylvinyl)phenyl)-3-(2-hydroxyethyl)-2,3-dihydro-1H-pyrrolo[3,4-c]pyridine-6-carboxylate, HCl (34.8 mg, 1.00 eq, 76.2 µmol) was dissolved in a solution of dry $CH_2Cl_2$ (2 mL) and $NEt_3$ (116 mg, 159 µL, 15.0 eq, 1.14 mmol). This solution was then added dropwise to the reaction vial over the course of 5 min and the reaction was stirred for 18 hr with gradual warming from 0 °C to rt. The reaction solution was diluted with $CH_2Cl_2$ (40 mL) and washed with 0.5 M aq. citric acid (2 x 40 mL). The organic layer was dried over $MgSO_4$, filtered, and the solvent was removed under reduced pressure. The residue was then purified by flash chromatography (0–100% EtOAC in hexanes) to afford ethyl (*R*,*E*)-4-(4-(2-cyclohexylvinyl)phenyl)-2-((3,5-dichlorophenyl)sulfonyl)-3-(2-hydroxyethyl)-2,3-dihydro-1*H*-pyrrolo[3,4-*c*]pyridine-6-carboxylate (46 mg, 73.2 µmol, 96%) as a yellow powder.

Ethyl (*R*,*E*)-4-(4-(2-cyclohexylvinyl)phenyl)-2-((3,5-dichlorophenyl)sulfonyl)-3-(2-hydroxyethyl)-2,3-dihydro-1H-pyrrolo[3,4-*c*]pyridine-6-carboxylate (31.5 mg, 1.00 eq, 50.0 µmol) was weighed into an 8 mL vial containing a small stir bar, dissolved in THF (2 mL), and chilled to 0 °C in an ice-water bath. A solution of LiOH (24.0 mg, 20.0 eq, 1.00 mmol) in water (2 mL) was added drop-wise over 5 min, and the reaction was stirred for 18 hr with gradual warming from 0 °C to rt. The basic solution was acidified using a 0.5 M aq. citric acid, diluted with water (30 mL), then extracted with $Et_2O$ (3 x 50 mL). Combined organic portions were dried over $MgSO_4$, filtered, and (*R,E*)-4-(4-(2-cyclohexylvinyl)phenyl)-2-((3,5-dichlorophenyl)sulfonyl)-3-(2-hydroxyethyl)-2,3-dihydro-1*H*-pyrrolo[3,4-*c*]pyridine-6-carboxylic acid was isolated under reduced pressure. To the resulting residue was added HATU (22.8 mg, 1.20 eq, 60.0 µmol, $CH_2Cl_2$ (3 mL), and $NEt_3$ (15.2 mg, 20.9 µL, 3.00 eq, 150 µmol). The mixture was stirred for 15 min at rt before addition of methanamine, HCl (6.75 mg, 2.00 eq, 100 µmol) and continued stirring for 2 hr. The solution was diluted with $CH_2Cl_2$ (50 mL), washed with 0.5 M aq. citric acid (2 x 50 mL), and dried over $MgSO_4$. After filtration solvent was removed under reduced pressure, and the residue was purified by preparative HPLC to afford **60** (24.5 mg, 40 µmol, 80% over two steps) as an off-white solid.

$^{1}$H NMR (400 MHz, DMSO) δ 8.60 (q, *J* = 4.8 Hz, 1H), 7.99 (d, *J* = 1.8 Hz, 2H), 7.88 – 7.82 (m, 2H), 7.73 (d, *J* = 8.4 Hz, 2H), 7.54 (d, *J* = 8.4 Hz, 2H), 6.48 (d, *J* = 16.1 Hz, 1H), 6.40 (dd, *J* = 16.1, 6.4 Hz, 1H), 5.63 – 5.55 (m, 1H), 5.13 (d, *J* = 17.4 Hz, 1H), 4.76 (dt, *J* = 17.3, 1.3 Hz, 1H), 4.36 (t, *J* = 5.0 Hz, 1H), 3.46 – 3.36 (m, 1H), 3.30 – 3.23 (m, 1H), 2.83 (d, *J* = 4.9 Hz, 3H), 2.19 (tdt, *J* = 10.2, 6.5, 3.4 Hz, 1H), 1.85 – 1.71 (m, 4H), 1.70 – 1.54 (m, 3H), 1.37 – 1.15 (m, 6H). $^{13}$C NMR (101 MHz, DMSO) δ 163.85, 150.85, 149.59, 149.52, 139.72, 138.28, 137.72, 135.71, 135.42, 135.36, 132.91, 128.61, 126.58, 126.10, 125.84, 114.69, 62.65, 57.04, 52.36, 40.56, 35.78, 32.37, 32.33, 26.06, 25.67, 25.49. HRMS (ESI): calculated for $C_{31}H_{34}Cl_2N_3O_4S$ $[M + H]^+$, 614.1642 found, 614.1695.

((8*R*,9*R*,10*S*)-9-(4-((*E*)-2-Cyclohexylvinyl)phenyl)-6-((2-methoxyphenyl)sulfonyl)-1,6-diazabicyclo[6.2.0]decan-10-yl)methanol (**61**)

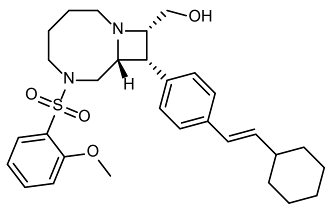

A 1-dram scintillation vial equipped with a magnetic stir bar was charged with ((8*R*,9*R*,10*S*)-9-(4-bromophenyl)-6-((2-nitrophenyl)sulfonyl)-1,6-diazabicyclo[6.2.0]decan-10-yl)methanol (12.5 mg, 24.5 µmol, 1 eq), dry DMF (0.3 mL), $K_2CO_3$ (16.9 mg, 122 µmol, 5 eq) and 4-methoxybenzenethiol (12.4 µL, 98 µmol, 4 eq). The yellow suspension was heated to 50 °C and stirred for 50 min. The obtained yellow/orange suspension was concentrated under reduced pressure. EtOAc (10 mL) and 0.3 M aq. HCl (10 mL) were added, and the aqueous phase was extracted twice with EtOAc. The aqueous layer was then basified with 40% aq. KOH solution and extracted three times with EtOAc. The organic layers were combined and dried over $Na_2SO_4$. After filtration the resulting solution was concentrated under reduced pressure and the crude ((8*R*,9*R*,10*S*)-9-(4-bromophenyl)-1,6-diazabicyclo[6.2.0]decan-10-yl)methanol (24.5 µmol, 1 eq) was dissolved in $CH_2Cl_2$ (0.5 mL), followed by addition of $NEt_3$ (6.83 µL, 49 µmol, 2.0

eq). 2-Methoxybenzenesulfonyl chloride (5.57 mg, 27.0 µmol, 1.1 eq) was added and the reaction was stirred for 1 hr before concentrating under reduced pressure. The resulting residue was added to a 2 mL µW vial along with (*E*)-(2-cyclohexylvinyl)boronic acid (4.5 mg, 29 µmol, 1.2 eq) and XPhos-Pd-G2 (3.9 mg, 4.9 µmol, 0.2 eq). The vial was capped with a septum and purged 3 times with $N_2$ gas and vacuum. Anhydrous DMF (0.5 mL) was added, followed by addition of 1.8 M aq. $K_3PO_4$ (0.16 mL, 294 µmol, 12 eq). The reaction was stirred at 60 °C for 2 hr under µW irradiation, then the resulting dark brown suspension was filtered through celite. Solvents were removed under reduced pressure, and target compound was purified by flash chromatography (30–60% EtOAc in hexanes) followed by further purification by preparative HPLC to afford **61** (2.3 mg, 4.4 µmol, 18% over 3 steps) as a white solid.

$^1$H NMR (400 MHz, $CD_3OD$) δ 7.77 (dd, *J* = 7.8, 1.7 Hz, 1H), 7.53 (ddd, *J* = 8.8, 7.5, 1.7 Hz, 1H), 7.40 (d, *J* = 8.1 Hz, 2H), 7.32 (d, *J* = 8.0 Hz, 2H), 7.07 (d, *J* = 8.4 Hz, 1H), 7.02 (t, *J* = 7.6 Hz, 1H), 6.36 (d, *J* = 16.0 Hz, 1H), 6.21 (dd, *J* = 16.0, 6.9 Hz, 1H), 3.74 – 3.40 (m, 6H), 3.53 (s, 3H), 3.16 (td, *J* = 9.2, 4.7 Hz, 1H), 2.97 – 2.86 (m, 2H), 2.50 (ddd, *J* = 12.2, 7.0, 3.2 Hz, 1H), 2.14 (dtd, *J* = 11.0, 7.3, 3.4 Hz, 1H), 2.02 – 1.91 (m, 1H), 1.88 – 1.77 (m, 6H), 1.74 – 1.60 (m, 2H), 1.45 – 1.17 (m, 6H). $^{13}$C NMR (101 MHz, $CD_3OD$) δ 156.53, 136.78, 136.10, 134.64, 134.35, 131.00, 130.87, 126.84, 126.75, 125.03, 119.75, 112.17, 69.43, 61.10, 58.55, 54.52, 52.75, 49.47, 42.23, 41.16, 32.76, 28.35, 25.88, 25.76. HRMS (ESI): calculated for $C_{30}H_{41}N_2O_4S$ [M + H]$^+$, 525.2782 found, 525.2802.

(2*R*,4*S*)-4-(3-((*E*)-2-Cyclohexylvinyl)phenoxy)-1-((3,5-dichlorophenyl)sulfonyl)-*N*-methylpyrrolidine-2-carboxamide (**62**)

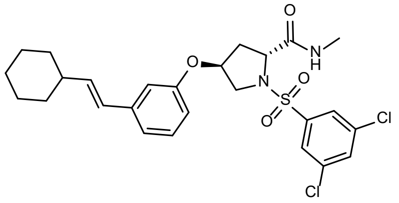

To a 1-dram vial containing (2*R*,4*S*)-1-(tert-butoxycarbonyl)-4-(3-iodophenoxy)pyrrolidine-2-carboxylic acid (17 mg, 39 µmol), $NH_3MeCl$ (3.2 mg, 47 µmol, 1.2 eq) and HATU (18 mg, 47 µmol, 1.2 eq) anhydrous $CH_2Cl_2$ (400 µL) was added, followed by DIPEA (34 µL, 200 µmol, 5 eq). The resulting yellow solution was stirred overnight at rt, diluted with $CH_2Cl_2$ and washed with water (1 mL) 3 times and brine (1 mL). The combined organic portions were dried over anhydrous $MgSO_4$, filtered, and concentrated under reduced pressure. The resulting crude *tert*-butyl (2*R*,4*S*)-4-(3-iodophenoxy)-2-(methylcarbamoyl)pyrrolidine-1-carboxylate was combined with analogous material from additional reactions and used for subsequent steps without further purification.

To a 2 mL μW vial was added crude *tert*-butyl (2*R*,4*S*)-4-(3-iodophenoxy)-2-(methylcarbamoyl)pyrrolidine-1-carboxylate (0.14 mmol, 1 eq), (*E*)-(2-cyclohexylvinyl)boronic acid (32 mg, 0.21 mmol, 1.5 eq) and XPhos-Pd-G2 (22 mg, 28 μmol, 0.2 eq). The vial was capped with a septum and purged 3 times with $N_2$ gas. Anhydrous DMF (2 mL) was added, followed by 1.8 M aq. $K_3PO_4$ (0.47 mL, 0.84 mmol, 6 eq). The reaction was stirred at 60 °C for 50 min under μW irradiation, then the resulting dark brown suspension was transferred to a scintillation vial and concentrated under reduced pressure. The mixture was diluted with EtOAc, followed by washing with water 3 times and brine. The combined organic portions were dried over anhydrous $MgSO_4$ and concentrated under reduced pressure. Purification by flash chromatography (50–70% EtOAc in hexanes) afforded *tert*-butyl (2*R*,4*S*)-4-(3-((*E*)-2-cyclohexylvinyl)phenoxy)-2-(methylcarbamoyl)pyrrolidine-1-carboxylate (53 mg, 0.12 mmol, 88% over 2 steps).

*tert*-Butyl (2*R*,4*S*)-4-(3-((*E*)-2-cyclohexylvinyl)phenoxy)-2-(methylcarbamoyl)pyrrolidine-1-carboxylate (53 mg, 0.12 mmol, 1 eq) was dissolved in $CH_2Cl_2$ (2 mL) before addition of TFA (400 μL) with stirring. After 1.5 hr the solution was concentrated under reduced pressure, and co-evaporated with toluene (2 mL). A portion of the resulting crude mixture was used in subsequent steps.

To a 1-dram vial charged with crude (2*R*,4*S*)-4-(3-((*E*)-2-cyclohexylvinyl)phenoxy)-*N*-methylpyrrolidine-2-carboxamide (0.03 mmol, 1 eq), 3,5-dichlorobenzenesulfonyl chloride (9 mg, 0.04 mmol, 1.2 eq) was added. This mixture was dissolved in anhydrous $CH_2Cl_2$ (200 μL) and chilled to 0 °C with an ice bath. $NEt_3$ (0.01 mL, 0.09 mmol, 3 eq) was added and the reaction was allowed to warm to rt with stirring overnight. The resulting solution was concentrated under reduced pressure, dissolved in DMSO, and directly purified by preparative HPLC to afford **62** (9 mg, 17 μmol, 60%) as a white solid.

$^1$H NMR (400 MHz, $CDCl_3$) δ 7.67 (d, *J* = 1.8 Hz, 2H), 7.52 (t, *J* = 1.9 Hz, 1H), 7.11 (td, *J* = 7.6, 1.1 Hz, 1H), 6.90 (dt, *J* = 7.6, 1.2 Hz, 1H), 6.70 (q, *J* = 5.1 Hz, 1H), 6.32 – 6.21 (m, 3H), 6.11 (dd, *J* = 16.0, 6.9 Hz, 1H), 4.77 – 4.70 (m, 1H), 4.31 (t, *J* = 8.3 Hz, 1H), 3.81 (dt, *J* = 13.0, 1.6 Hz, 1H), 3.75 (dd, *J* = 13.0,

3.4 Hz, 1H), 2.91 (d, *J* = 4.9 Hz, 3H), 2.52 (ddt, *J* = 14.2, 8.0, 2.0 Hz, 1H), 2.33 (ddd, *J* = 13.7, 8.7, 4.5 Hz, 1H), 2.14 (dtt, *J* = 10.7, 6.9, 3.5 Hz, 1H), 1.86 – 1.75 (m, 4H), 1.74 – 1.66 (m, 1H), 1.37 – 1.18 (m, 5H). $^{13}$C NMR (101 MHz, $CDCl_3$) δ 170.91, 155.86, 139.93, 138.98, 137.71, 136.20, 133.30, 129.47, 126.70, 126.46, 120.01, 112.66, 112.13, 77.20, 74.75, 62.04, 55.00, 41.16, 37.07, 32.97, 32.95, 26.45, 26.14, 26.02. HRMS (ESI): calculated for $C_{26}H_{31}ClN_2O_4S$ [M + H]$^+$, 537.1376 found, 537.1381.

(2*S*,4*S*)-4-(4-((*E*)-2-Cyclohexylvinyl)phenoxy)-1-(3-ethynylbenzyl)-*N*-methylpyrrolidine-2-carboxamide (**63**)

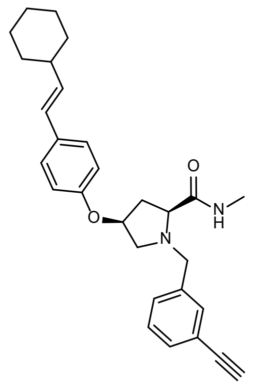

To a solution of (2*S*,4*S*)-4-(4-((*E*)-2-cyclohexylvinyl)phenoxy)-*N*-methylpyrrolidine-2-carboxamide hydrochloride [*cf.* synthesis of **29**] (15 mg, 1 eq, 41 μmol) in DCE (2 mL) was added 3-ethynylbenzaldehyde (6.4 mg, 1.2 eq, 49 μmol) and sodium triacetoxyhydroborate (13 mg, 1.5 eq, 62 μmol). The reaction mixture was stirred for 16 hr at rt and concentrated under reduced pressure. The crude residue was dissolved in $CH_2Cl_2$ (25 mL) and was washed with $H_2O$ (15 mL). The aqueous layer was back-extracted with $CH_2Cl_2$ (2 x 25 mL), and the combined organics were dried over $Na_2SO_4$ and concentrated under reduced pressure. The crude residue was purified by preparative to afford **63** (10.5 mg, 58%) as a white solid.

$^1$H NMR (400 MHz, $CD_3OD$) δ 7.51 (d, *J* = 1.7 Hz, 1H), 7.40 (ddt, *J* = 10.9, 7.7, 1.6 Hz, 2H), 7.33 (t, *J* = 7.6 Hz, 1H), 7.27 – 7.22 (m, 2H), 6.80 – 6.74 (m, 2H), 6.28 (dd, *J* = 16.0, 1.1 Hz, 1H), 6.04 (dd, *J* = 16.0, 7.0 Hz, 1H), 4.88 (t, *J* = 5.1 Hz, 1H), 3.82 (d, *J* = 12.9 Hz, 1H), 3.67 (d, *J* = 12.9 Hz, 1H), 3.49 (s, 1H), 3.32 – 3.25 (m, 2H), 2.78 (dd, *J* = 10.8, 4.4 Hz, 1H), 2.71 (s, 3H), 2.62 (ddd, *J* = 14.2, 10.3, 5.6 Hz, 1H), 2.15 – 2.03 (m, 2H), 1.79 (ddq, *J* = 9.1, 7.0, 3.5 Hz, 4H), 1.70 (ddt, *J* = 12.4, 3.5, 1.6 Hz, 1H), 1.45 – 1.11 (m, 6H). $^{13}$C NMR (151 MHz, DMSO) δ 173.04, 156.58, 139.19, 134.55, 132.57, 130.96, 130.63, 130.15,

129.00, 127.48, 126.90, 122.08, 115.86, 84.02, 81.20, 75.64, 66.98, 58.83, 58.55, 40.89, 37.13, 33.05, 26.15, 26.03, 25.98. HRMS (ESI): calculated for $C_{29}H_{35}N_2O_2$ $[M + H]^+$, 443.2693 found, 443.2696.

### 2. Synthesis: on-DNA

#### 2.1. General protocols

##### 2.1.1. Ethanol precipitation

To a reaction of a given volume (100% v/v), aq. NaCl (5 M, 10% v/v) and EtOH (300% v/v) were added. The reaction vessel was briefly vortexed and incubated on dry ice for 30 to 60 min. After centrifugation (10–20 min at 3750 rcf), the supernatant was removed by decanting into another vessel. To wash the pellet, 80% aq. EtOH (100-200% v/v) was added. The mixture was vortexed for approx 2 min and chilled again on dry ice for approx 10 min. After centrifugation (10–20 min at 3750 rcf), the supernatant was removed by decanting into another vessel. The washing procedure was repeated with 100% EtOH (100-200% v/v) and the obtained pellet was dried under vacuum.

#### 2.2. Optimization prior to library synthesis

##### 2.2.1. Attachment of skeletons to headpiece

###### 2.2.1.1. Amide formation

A small number of skeletons were used for development of effective skeleton-linkage protocols. **Supplementary Table 6** shows the effect of skeleton and DMTMM concentration, as well as organic solvent choice and reaction concentration. Blank data cells indicate that no peak/mass in the LCMS data could confidently be assigned to the noted species, in some cases due to a failed injection at the LCMS. General trends highlighted include:

1) Higher molar equivalents of skeleton relative to AOP-HP tended to afford greater conversion to desired product.
2) Higher organic content and/or higher dilution tended to afford greater conversion to desired product.

3) Higher molar equivalents of DMTMM relative to AOP-HP tended to afford more of the undesired acetylated AOP-HP product, which could be further augmented, though not to a dramatic extent, by a lower relative loading of skeleton.

A more focused set of reactions were performed for one enantiomer of each pair from the *meta*-iodo β-arylated Fmoc-protected amino acids. A summary of these experiments can be found in **Supplementary Table 7**, and the associated learnings could be summarized as follows:

1) The order of skeleton reactivity towards acylation follows azetidine>pyrrolidine>piperidine.
2) *trans*-Configured azetidines and pyrrolidines give marginally improved conversion to desired product than their *cis*-diastereomers, with no preference between *cis*- and *trans*-piperidines.
3) High organic content affords greater conversion to desired product, regardless of ring size.

Further focusing of conditions with %organic solvent mixtures of >70%, as shown in **Supplementary Table 8**, led to conditions generally affording >80%AUC product for all ring sizes and relative stereochemistry.

**Supplementary Table 6 | Initial acylation conditions tested for skeleton attachment to AOP-HP**

| Skeleton | Conc, DMF, mM | Eq to AOP-HP | DMTMM solvent | DMTMM eq rel to AOP-HP | % org | DNA conc, mM | % AOP-HP | % P | % DMT-add | % Ac-add |
|---|---|---|---|---|---|---|---|---|---|---|
| Fmoc Pro | 100 | 100 | water | 100 | 33 | 0.33 | | 81 | | 9 |
| | | | | 10 | 33 | 0.33 | | 97 | | 3 |
| | | | MeOH | 100 | 66 | 0.33 | | 75 | | 15 |
| | | | | 10 | 66 | 0.33 | | 100 | | 0 |
| | | | DMSO | 100 | 66 | 0.33 | | 100 | | 0 |
| | | | | 10 | 66 | 0.33 | | 89 | | 3 |
| | | 10 | water | 100 | 5 | 0.48 | | 0 | | 100 |
| | | | | 10 | 5 | 0.48 | | 50 | | 50 |
| | | | MeOH | 100 | 52 | 0.48 | | 0 | | 100 |
| | | | | 10 | 52 | 0.48 | 60 | 40 | | |
| | | | DMSO | 100 | 52 | 0.48 | | 38 | 10 | |
| | | | | 10 | 52 | 0.48 | | | | |
| | 10 | 100 | water | 100 | 83 | 0.08 | | 74 | | 8 |
| | | | | 10 | 83 | 0.08 | | 100 | | |
| | | | MeOH | 100 | 92 | 0.08 | | 50 | | 44 |
| | | | | 10 | 92 | 0.08 | 49 | 51 | | 0 |
| | | | DMSO | 100 | 92 | 0.08 | | 66 | | 25 |
| | | | | 10 | 92 | 0.08 | | 90 | | 6 |

| | | | | | | | | | | |
|---|---|---|---|---|---|---|---|---|---|---|
| | | 10 | water | 100 | 33 | 0.33 | | 66 | | 20 |
| | | | | 10 | 33 | 0.33 | | 50 | | 50 |
| | | | MeOH | 100 | 66 | 0.33 | | 47 | 10 | 31 |
| | | | | 10 | 66 | 0.33 | 100 | | | |
| | | | DMSO | 100 | 66 | 0.33 | | | | 5 |
| | | | | 10 | 66 | 0.33 | | | | |
| Fmoc β-arylPro | 100 | 100 | water | 100 | 33 | 0.33 | | 55 | 6 | 31 |
| | | | | 10 | 33 | 0.33 | | 97 | | 3 |
| | | | MeOH | 100 | 66 | 0.33 | | 48 | 5 | 41 |
| | | | | 10 | 66 | 0.33 | | 97 | | 3 |
| | | | DMSO | 100 | 66 | 0.33 | | 89 | | 8 |
| | | | | 10 | 66 | 0.33 | | 97 | | 3 |
| | | 10 | water | 100 | 5 | 0.48 | | | | 60 |
| | | | | 10 | 5 | 0.48 | 69 | 14 | | 13 |
| | | | MeOH | 100 | 52 | 0.48 | | | | |
| | | | | 10 | 52 | 0.48 | 64 | | | 27 |
| | | | DMSO | 100 | 52 | 0.48 | | | | 39 |
| | | | | 10 | 52 | 0.48 | 55 | | 17 | 28 |
| | 10 | 100 | water | 100 | 83 | 0.08 | 96 | | | |
| | | | | 10 | 83 | 0.08 | | 100 | | |
| | | | MeOH | 100 | 92 | 0.08 | | 100 | | |
| | | | | 10 | 92 | 0.08 | 20 | 62 | | 10 |
| | | | DMSO | 100 | 92 | 0.08 | | 100 | | |
| | | | | 10 | 92 | 0.08 | | 89 | | 7 |
| | | 10 | water | 100 | 33 | 0.33 | | 62 | 7 | 19 |
| | | | | 10 | 33 | 0.33 | 59 | 33 | | 5 |
| | | | MeOH | 100 | 66 | 0.33 | | 17 | 8 | 37 |
| | | | | 10 | 66 | 0.33 | 96 | | | 4 |
| | | | DMSO | 100 | 66 | 0.33 | | | | 9 |
| | | | | 10 | 66 | 0.33 | | 90 | | 10 |

**Supplementary Table 7 | Screening of acylation conditions for linkage of *meta*-iodo β-arylated Fmoc-protected amino acids to AOP-HP**

| Skeleton (100 mM, DMF, 50 eq) | DMTMM solvent | DMTMM eq | DMTMM conc, mM | % org | final [DNA], mM | % AUC_product |
|---|---|---|---|---|---|---|
| *cis*-Aze | DMSO | 50 | 100 | 50 | 0.50 | 70 |
| | | | 50 | 60 | 0.40 | 85 |
| | | 30 | 100 | 44 | 0.56 | 57 |
| | | | 50 | 52 | 0.48 | 82 |
| | water | 50 | 100 | 25 | 0.50 | 27 |
| | | | 50 | 20 | 0.40 | 8 |
| | | 30 | 100 | 28 | 0.56 | 20 |
| | | | 50 | 24 | 0.48 | 24 |
| *trans*-Aze | DMSO | 50 | 100 | 50 | 0.50 | 90 |
| | | | 50 | 60 | 0.40 | 94 |
| | | 30 | 100 | 44 | 0.56 | 90 |
| | | | 50 | 52 | 0.48 | 96 |
| | water | 50 | 100 | 25 | 0.50 | 45 |
| | | | 50 | 20 | 0.40 | 12 |
| | | 30 | 100 | 28 | 0.56 | 48 |
| | | | 50 | 24 | 0.48 | 9 |

| | | | | | | |
|---|---|---|---|---|---|---|
| *cis*-Pyr | DMSO | 50 | 100 | 50 | 0.50 | 10 |
| | | | 50 | 60 | 0.40 | 30 |
| | | 30 | 100 | 44 | 0.56 | 9 |
| | | | 50 | 52 | 0.48 | 13 |
| | water | 50 | 100 | 25 | 0.50 | |
| | | | 50 | 20 | 0.40 | 0 |
| | | 30 | 100 | 28 | 0.56 | 0 |
| | | | 50 | 24 | 0.48 | 0 |
| *trans*-Pyr | DMSO | 50 | 100 | 50 | 0.50 | 42 |
| | | | 50 | 60 | 0.40 | 65 |
| | | 30 | 100 | 44 | 0.56 | 33 |
| | | | 50 | 52 | 0.48 | 57 |
| | water | 50 | 100 | 25 | 0.50 | 5 |
| | | | 50 | 20 | 0.40 | 4 |
| | | 30 | 100 | 28 | 0.56 | |
| | | | 50 | 24 | 0.48 | 5 |
| *cis*-Pip | DMSO | 50 | 100 | 50 | 0.50 | 10 |
| | | | 50 | 60 | 0.40 | 30 |
| | | 30 | 100 | 44 | 0.56 | 9 |
| | | | 50 | 52 | 0.48 | 13 |
| | water | 50 | 100 | 25 | 0.50 | 0 |
| | | | 50 | 20 | 0.40 | 0 |
| | | 30 | 100 | 28 | 0.56 | 4 |
| | | | 50 | 24 | 0.48 | 0 |
| *trans*-Pip | DMSO | 50 | 100 | 50 | 0.50 | 18 |
| | | | 50 | 60 | 0.40 | 33 |
| | | 30 | 100 | 44 | 0.56 | 14 |
| | | | 50 | 52 | 0.48 | 18 |
| | water | 50 | 100 | 25 | 0.50 | 4 |
| | | | 50 | 20 | 0.40 | 5 |
| | | 30 | 100 | 28 | 0.56 | 7 |
| | | | 50 | 24 | 0.48 | 5 |

**Supplementary Table 8 | Focused screening of acylation conditions for linkage of *meta*-iodo β-arylated Fmoc-protected amino acids to AOP-HP**

| Skeleton (10 mM, DMF, 50 eq) | [DNA], mM | DMTMM solvent (50 mM, 50 eq) | % org | % AUC_product | % AUC_AOP-HP | % AUC_Ac-AOP-HP |
|---|---|---|---|---|---|---|
| *cis*-Aze | 0.14 | DMSO | 86 | 90 | 2 | 0 |
| | | water | 71 | 94 | 0 | 0 |
| *trans*-Aze | | DMSO | 86 | 99 | 0 | 0 |
| | | water | 71 | 92 | 0 | 0 |
| *cis*-Pyr | | DMSO | 86 | 44 | 38 | 3 |
| | | water | 71 | 72 | 16 | 5 |
| *trans*-Pyr | | DMSO | 86 | 98 | 2 | 0 |
| | | water | 71 | 100 | 0 | 0 |
| *cis*-Pip | | DMSO | 86 | 63 | 28 | 8 |
| | | water | 71 | 87 | 10 | 3 |
| *trans*-Pip | | DMSO | 86 | 89 | 9 | 2 |

| | | | | | | |
|---|---|---|---|---|---|---|
| | | water | 71 | 83 | 13 | 4 |

**2.2.1.2. Carbamylation**

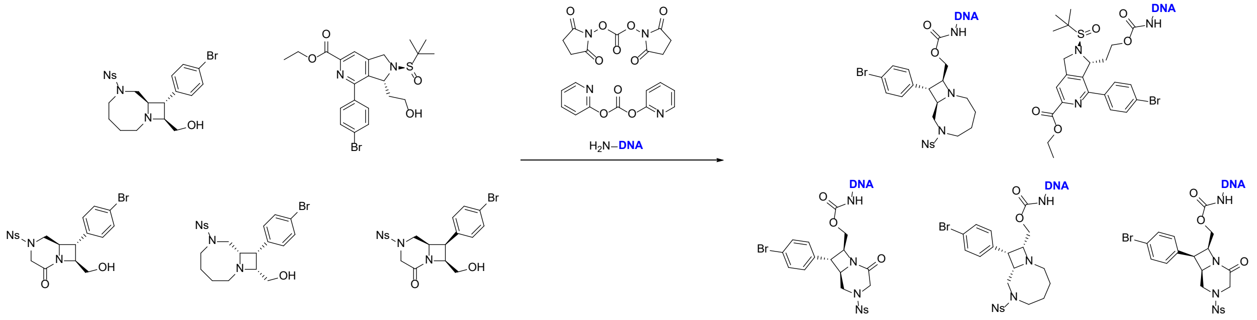

**Supplementary Scheme 4 | Initial DNA-carbamate formation reactions**

An initial experiment sought to compare the efficiency of carbamate formation following skeleton incubation with two different activating dicarbonates (**Supplementary Scheme 4**). One of the two dicarbonates (0.5 µL 100 mM, dry DMSO, 1 eq) was mixed with a skeleton (1 µL, 50 mM, dry DMSO, 1 eq) and $NEt_3$ (0.5 µL, 100 mM, dry DMSO). Each reaction was set up in triplicate with the first being heated at 40 °C for 1 hr, the second being heated at 40 °C for 4 hr, and the third being incubated at rt for 24 hr. After incubation as indicated, AOP-HP (1 µL, 1 mM solution in 0.1 M PB pH 8, 1 nmol, *assumed* 0.02 eq) was added. After 24 hr at rt each reaction underwent EtOH precipitation and analysis by LCMS.

Analysis of the resulting LCMS data showed that a longer preactivation period was beneficial, DSC was more effective than dipyridin-2-yl carbonate in leading to product formation, but in most cases, unreacted SM was the result.

Then the reaction sequence represented in **Supplementary Scheme 4** was repeated as a two-step sequence. First, activated carbonates were prepared by mixing a skeleton (1 eq) in dry DCM (0.1 M), to which was added $NEt_3$ (3 eq) followed by DSC (3 eq, 0.25 M solution in dry MeCN). The resulting solutions were stirred until LCMS indicated conversion to desired activated carbonate (typically 24 hr), then reactions mixtures were purified by flash chromatography (4 g silica; 0–100% EtOAc in heptane). Isolated active carbonates were assumed to be 80% pure and at 80% recovery before dissoution in DMF (50 mM).

To AOP-headpiece (1 eq, 0.5 mM in 0.1 M PB pH 8) was added an active carbonate (5–10 eq, 50 mM in DMF) and additional DMF was added to ensure organic content of the reaction was 50%. Solutions were incubated at rt overnight and then underwent EtOH precipitation to afford crude products of typically >80 %AUC with respect to desired product. Example outcomes can be seen in **Supplementary Fig. 1**.

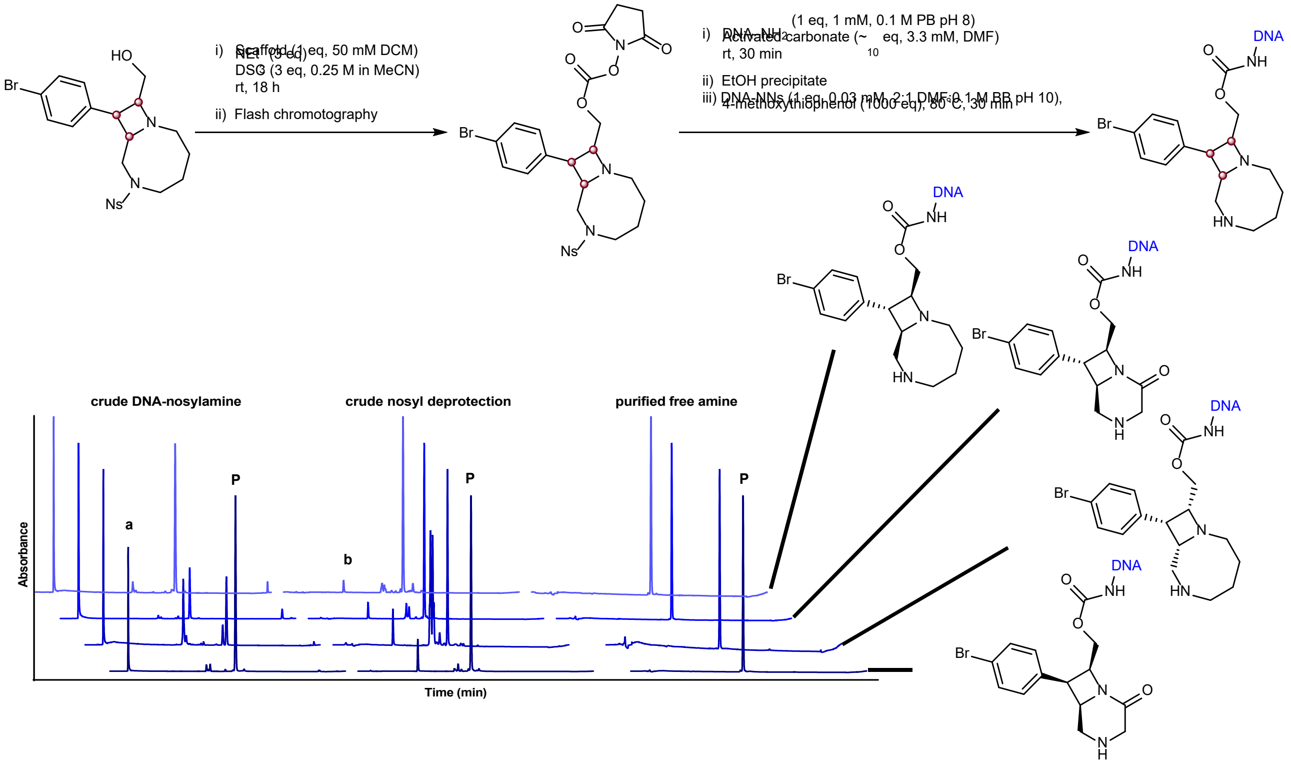

**Supplementary Fig. 1 | Optimized process for carbamylation and solution-phase nosyl deprotection**. Example traces showing UV absorbance at 260 nm during ion-pairing chromatography for four example bicyclic azetidine skeletons. a: excess non-DNA linked skeleton, b: residual 4-methoxy thiophenol following EtOH precipitation after nosyl deprotection, P – the desired product of each step.

##### 2.2.2. Nosyl deprotection

An operationally facile nosyl deprotection was investigated using the skeleton-DNA construct shown in **Supplementary Scheme 5**.

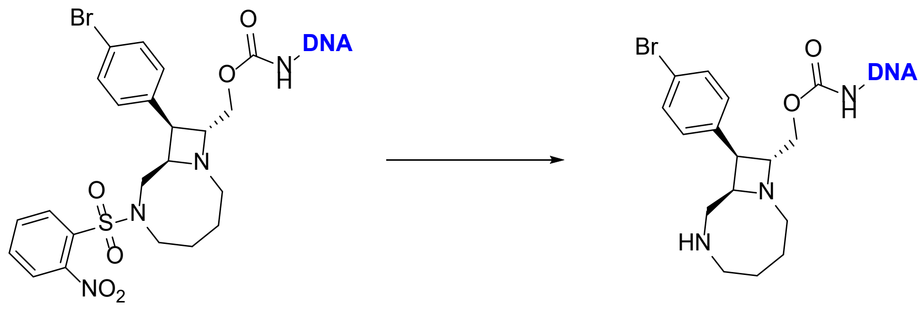

**Supplementary Scheme 5 | Model skeleton employed for optimization of a nosyl deprotection**

A set of four thiols were used for initial tests (mercaptoacetic acid, mecaptoethanol, 4-methoxy thiophenol, and SiliaMetS thiol). All thiols were dissolved/suspended in DMF to a concentration of 100 mM, and 10/50/100/500/1000 eq was added a nosyl-amine-DNA construct (0.5 nmol in 5 µL of 0.1 M bicarbonate buffer pH 10.4). Samples were briefly vortexed, centrifuged and stood at rt for either 1 hr or 24 hr before analyzing by LCMS. At the 1 hr time point only 4-methoxythiophenol afforded >10% conversion at 1000 eq. At the 24 hr time point the following observations could be made:

| | |
|---|---|
| Mercaptoacetic acid | up to 13%AUC deprotection (at 1000 eq) |
| Mercaptoethanol | deprotection levels between 7–41%AUC observed (10–1000 eq) |
| 4-methoxythiophenol | deprotection levels between 11–75%AUC observed (10–1000 eq) |
| SiliaMetS Thiol | minimal levels of deprotection could be observed, the extent of which appeared unrelated to ‘stoichiometry’ |

Following the observation that 4-methoxythiophenol led to the highest levels of deprotection we next investigated a set of temperature and time combinations as outlined in **Supplementary Table 9**. Identifying that suitable loading of 4-methoxythiophenol at elevated temperatures could afford a clean reaction profile in relatively short reaction times.

**Supplementary Table 9 | Optimization of a nosyl deprotection**

| temp, °C | time, hr | 4-methoxythiophenol (100 mM, DMF), eq | %AUC_amine | %AUC_nosyl amine |
|---|---|---|---|---|
| 21 | 70 | 500 | 36 | 64 |
| | | 1000 | 73 | 27 |
| | | 2000 | 70 | 30 |

| | | | | |
|---|---|---|---|---|
| 40 | 22 | 500 | 40 | 60 |
| | | 1000 | 79 | 21 |
| | | 2000 | 83 | 17 |
| 60 | 5 | 500 | 67 | 33 |
| | | 1000 | 92 | 8 |
| | | 2000 | 94 | 6 |
| 80 | 1 | 500 | 78 | 22 |
| | | 1000 | 86 | 14 |
| | | 2000 | 96 | 4 |

##### 2.2.3. Suzuki coupling

A screen was performed using four model skeletons at a single concentration in a single buffer with a single concentration and number of molar equivalents of coupling partner. The skeleton electronic properties varied slightly, but not dramatically – we sought to ensure reaction conditions gave consistent outcomes without major influence from small differences in electronic effects. To that end we screened four palladium catalysts (all dissolved to 1 mM in degassed MeCN), with two boronic acid pinacol esters which were dissolved in either EtOH, MeCN, or DMA. Reactions were run at three different temperature-time combinations in heat-sealed Eppendorf twin.tec PCR plates (skirted; 951020619), then LCMS analysis was performed, and the resulting data was analyzed at the level of individual samples as well as on aggregate for each property of interest (**Supplementary Fig. 2**).

In total, 288 reactions were performed, of which 11 afforded complex mixtures; all involving 4-(4,4,5,5-tetramethyl-1,3,2-dioxaborolan-2-yl)-1H-indole dissolved in EtOH, and employing either $PdCl_2$(dppf).$CH_2Cl_2$ or $PdCl_2$(dippf) as catalyst, with 8 of those 11 reactions being from the 60 °C/5 hr conditions set. Of the four catalysts tested, $PdCl_2$(dppf).$CH_2Cl_2$ afforded the highest conversion to desired product at all temperatures and in all co-solvents relative to matched reaction sets. In addition, $PdCl_2$(dppf).$CH_2Cl_2$ usage had the least variable outcomes across a range of temperature and time combinations. We also found that MeCN was the preferred solvent for this reaction in combination with any of the catalysts tested.

**a**

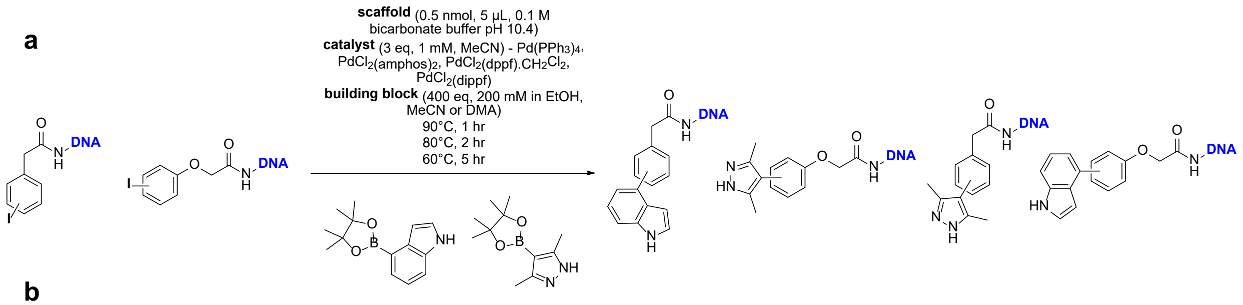

**b**

| Catalyst | Average %AUC yields at given temperatures with various co-solvents | | | | | | | | | | | | | | | |
|---|---|---|---|---|---|---|---|---|---|---|---|---|---|---|---|---|
| solvent | all | | | | EtOH | | | | MeCN | | | | DMA | | | |
| temp, °C | all | 90 | 80 | 60 | all | 90 | 80 | 60 | all | 90 | 80 | 60 | all | 90 | 80 | 60 |
| $Pd(PPh_3)_4$ | 48 | 61 | 43 | 40 | 41 | 44 | 51 | 29 | 58 | 81 | 43 | 48 | 45 | 58 | 35 | 43 |
| $PdCl_2(amphos)_2$ | 25 | 32 | 26 | 17 | 13 | 9 | 22 | 9 | 32 | 42 | 35 | 19 | 28 | 41 | 22 | 21 |
| $PdCl_2(dppf).CH_2Cl_2$ | 67 | 77 | 58 | 65 | 55 | 65 | 58 | 45 | 81 | 93 | 64 | 84 | 67 | 74 | 59 | 67 |
| $PdCl_2(dppf)$ | 46 | 65 | 43 | 29 | 28 | 46 | 24 | 13 | 66 | 85 | 69 | 43 | 43 | 65 | 37 | 31 |

**Supplementary Fig. 2 | Optimization of Suzuki coupling conditions. a**, Schematic overview of reaction conditions under investigation, representing 288 individual reactions. **b**, table showing mean %AUC yields for each reaction series, corrected for residual non-DNA species with absorbance at 260 nm, and treating all reactions equally (i.e., a reaction where %AUC w.r.t product was noted as "0", such as due to ambiguous deconvoluted mass, would not be discounted from the averaging). Conditional formatting was applied to specified temperature columns corresponding to average yield, where each of these cells is the mean %AUC of desired product from 8 reactions. The conditional formatting applied to "temp: all" columns corresponds to the standard deviation across all temperature/time combinations (as % of the mean value for the given solvent).

We also investigated the performance of Suzuki couplings on skeletons bearing free amines, using multiple stereoisomers of several skeleton series (**Supplementary Fig. 3**). For these test reactions three boronic acids/esters that had previously displayed consistently good performance with related substrates (with %AUC product in the 80–100% range and minimal unknown species evident in the LCMS data). We found that, as free amines, these skeletons displayed suboptimal performance, with low conversions to desired product and high variability dependent on either the boronic acid/ester, the skeleton series, or skeleton stereochemistry.

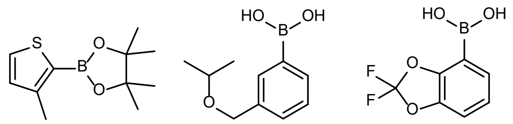

x y z

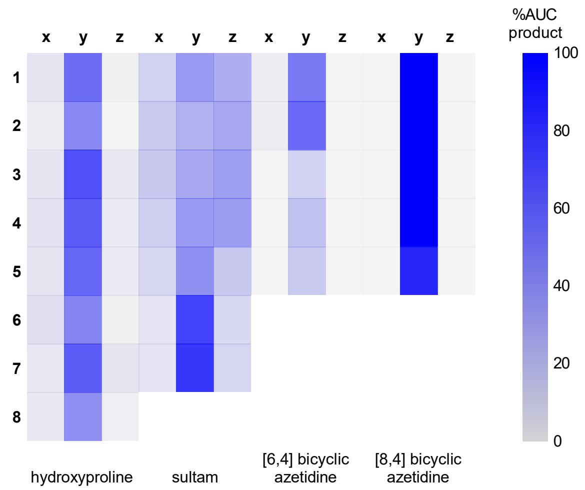

**Supplementary Fig. 3 | A heat map representation of %AUC product for a subset of test Suzuki coupling reactions with a subset of skeletons bearing free amines**

##### 2.2.4. Skeleton validation experiments

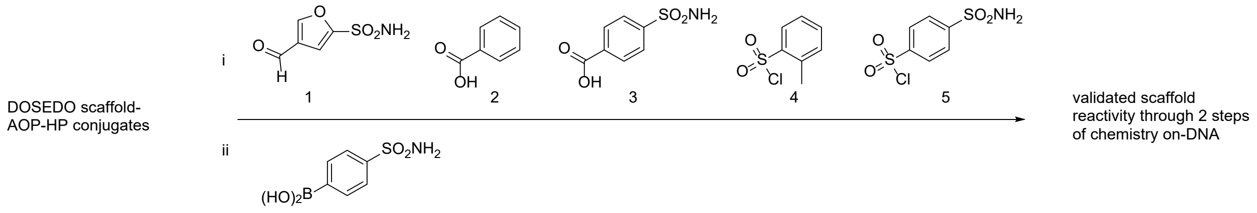

**Supplementary Scheme 6 | Overview of skeleton validation reactions performed**

All skeletons to be included in the DOSEDO library (after attachment to AOP-HP, amine deprotection, purification by micropreparative HPLC, desalting by ultrafiltration and normalization of concentration in nuclease free water) required a general assessment of reactivity and stability under all conditions to be employed in the DEL synthesis. Each skeleton was first assessed for purity by LCMS (**Supplementary Section 8.1.**), then reacted with five different amine capping building blocks followed by a single Suzuki coupling of each resulting product, under the following conditions:

**<u>Reductive amination conditions</u>:**

Skeleton-AOP-HP (2 µL, 0.9 nmol, ~0.45 mM, 1 eq, water), acetate buffer (8 µL, 1 M, pH 5) and aldehyde (2 µL, 200 mM, 444 eq, MeCN) were added to a 96-well plate (Eppendorf twin.tech lo-bind), mixing the components by three aspirate-dispense cycles. After 5 min of standing at rt $NaCNBH_3$ (1.5 µL, 333 eq, 200 mM, water) was added. The plate was sealed with heat-sealing foil, vortexed briefly, spun down, and incubated at 22 °C for 18 hr.

**<u>Acylation conditions</u>:**

DMTMM (1 µL, 100 mM, 111 eq, water, freshly prepared) and carboxylic acid (2 µL, 50 mM, 111 eq, DMF) were added to a 96-well plate (Eppendorf twin.tech, lo-bind), mixing the components by three aspirate-dispense cycles. After 5 min of standing at rt skeleton-AOP-HP (2 µL, 0.9 nmol, ~0.45 mM, 1 eq, water) was added. The plate was sealed with heat-sealing foil, vortexed briefly, spun down, and incubated at 22 °C for 18 hr.

**<u>Sulfonylation conditions:</u>**

Skeleton-AOP-HP (2 µL, 0.9 nmol, ~0.45 mM, 1 eq, water) was diluted with phosphate buffer (3 µL, 0.1M, pH 8) in a 96-well plate (Eppendorf twin.tech, lo-bind) before the addition of sulfonylchloride (1 µL, 200 mM, 222 eq, MeCN) and mixing the components by three aspirate-dispense cycles. The plate was sealed with heat-sealing foil, vortexed briefly, spun down, and incubated at 22 °C for 18 hr.

**<u>Amine capping common work-up:</u>**

Piperidine (2 µL, 10% solution in nuclease free water), was added to every reaction. After 2 min at rt each reaction was purified by EtOH precipitation, inverting supernatants into a pad of paper towels, and drying final pellets in air for several h. Pellets were dissolved in 10 µL of nuclease free water, 7 µL (~0.5 nmol) of which was transferred to a new plate for subsequent Suzuki coupling; allowing the water to evaporate overnight. The residual 3 µL was diluted with 50 µL of water and analyzed by ion-pairing LCMS

**<u>Suzuki coupling conditions:</u>**

N-capped-skeleton-AOP-HP-conjugate (0.5 nmol) was dissolved in 5 µL of 0.1 M bicarb buffer pH 10 (for the aryl iodides), or 5 µL of 0.1 M phosphate buffer pH 8 (for the aryl bromides). The added boronic acid/ester (1.5 µL, 200 mM, 600 eq1:1 mixture of MeCN and EtOH), followed by $PdCl_2(dppf).CH_2CL_2$ (1 µL [ArI] or 2.5 µL [ArBr], 2 eq [ArI] or 5 eq [ArBr], 1 mM in MeCN). Plate was sealed with heat-sealing foil, vortexed briefly, spun down, and incubated at 90 °C for 2 hr before cooling to 4 °C. Each well was treated with diethylthiocarbamic acid sodium salt (2 µL, 100 mM, water), plate sealed again, vortexed, spun down briefly, and stood at rt for approx 1 hr. Each reaction was EtOH precipitated by addition of 5 M NaCl (1 µL) and cold EtOH (50 µL) per well, then chilled on dry ice for ~20 min, centrifuged at 3750 rcf for 15 min before removing supernatant by inversion into a pad of fresh paper towels). Another portion of cold EtOH (100 µL) was added to each well, chilled on dry ice for ~20 min, spun down at 3750 rcf for 15 min before removing supernatant by inversion into a pad of fresh paper towels. The plate was then dried in air and the pellets were dissolved in water (50 µL) sealed thoroughly, vortexed briefly and spun down at 1000 rcf for 1 min before performing LCMS analysis.

##### 2.2.5. Building block validation experiments

###### 2.2.5.1. Amine capping reactions

Proline was attached to AOP-HP and purified following the general procedures in **Sections 2.3.1.1** and **2.3.2.1**. AOP-HP (1 µmol) and Fmoc-proline (100 eq) were used, resulting in proline-AOP-HP (637 nmol [Nanodrop], 64%) after acylation, Fmoc deprotection and purification. An LCMS trace following purification can be found in **Supplementary Fig. 4**. The proline-AOP-HP was subjected to acylation, reductive amination and sulfonylation conditions according to the below noted quantities and orders of addition.

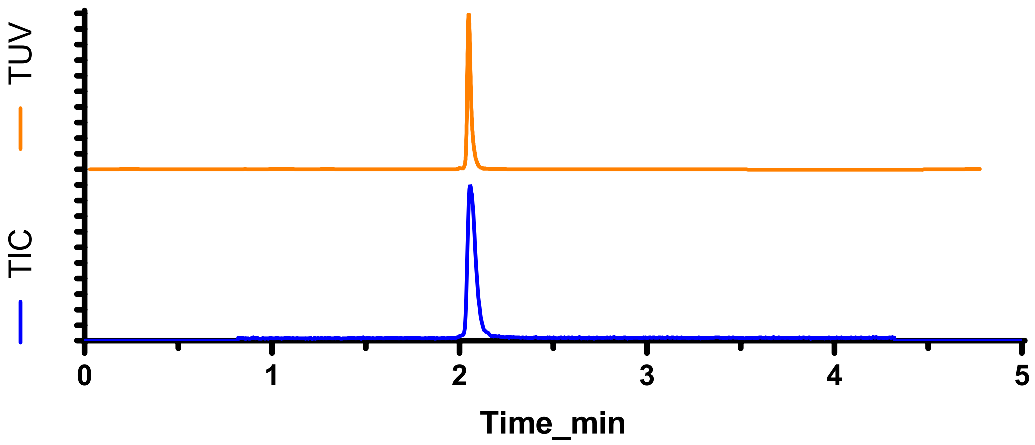

**Supplementary Fig. 4 | Isolated proline-AOP-HP used for amine capping reaction validation**

**<u>Reductive amination conditions:</u>**

In 96-well plates proline-AOP-HP (2 µL, 0.5 nmol, 0.25 mM, 1 eq, 1 mM pH 8 PB) was diluted in acetate buffer (8 µL, 1 M, pH 5) before the addition of aldehyde (2 µL, 400 nmol, 200 mM, 800 eq, MeCN). The mixture was mixed by pipetting 3 times and stood for <5 min before the addition of $NaCNBH_3$ (1.5 µL, 300 nmol, 200 mM, 600 eq, water). The plate was heat-sealed, vortexed briefly, spun down and incubated at 25 °C for 16 hr.

**<u>Acylation conditions:</u>**

In 96-well plates freshly prepared DMTMM (0.5 µL, 50 nmol, 100 mM, 100 eq) and carboxylic acid (2 µL, 50 mM, 50 nmol, 100 eq, DMF) were spun to the bottom of each well. Proline-AOP-HP (2 µL, 0.5 nmol, 0.25 mM, 1 eq, 1 mM pH 8 PB) was added, the plate heat-sealed, vortexed briefly, spun down and incubated at 25 °C for 16 hr.

**<u>Sulfonylation conditions:</u>**

In 96-well plates proline-AOP-HP (2 µL, 0.5 nmol, 0.25 mM, 1 eq, 1 mM pH 8 PB) was diluted with PB (3 µL, 0.1 M, pH 8) before the addition of sulfonylchloride (1 µL, 200 nmol, 200 mM, 400 eq, MeCN). The plate was heat-sealed, vortexed briefly, spun down and incubated at 25 °C for 16 hr.

All three reaction types were worked-up in parallel according to the below protocol:

To each well piperidine was added (5 µL, 20%v/v in water), and plates were spun down. After <5 min the reactions underwent EtOH precipitation, and dried pellets were dissolved in 40 µL of nuclease free water for LCMS analysis.

**2.2.5.2. Suzuki coupling**

All boronic acids and esters available were assessed for their performance in Suzuki coupling under conditions optimized for representative ArBr and ArI-bearing skeletons of the planned DOSEDO library.

The on-DNA skeletons employed for these assessments (provided as smiles strings) were: O=C(OC[C@H]1[C@H](C2=CC=C(Br)C=C2)[C@@]3([H])CN(C(C)=O)CC(N13)=O)NC[3H] and O=C([C@H]1N(C(C)=O)CC[C@@H]1C2=CC=CC(I)=C2)NC[3H]. In both cases the smiles strings utilize a tritium atom to indicate the DNA attachment point. Also note that the amines of both skeletons were acetylated. The relevant skeleton-AOP-HP constructs were prepared according to general procedures outlined in **Supplementary Sections 2.3.1.1** and **2.3.2.1**, with LCMS data for isolated constructs shown in **Supplementary Fig. 5**.

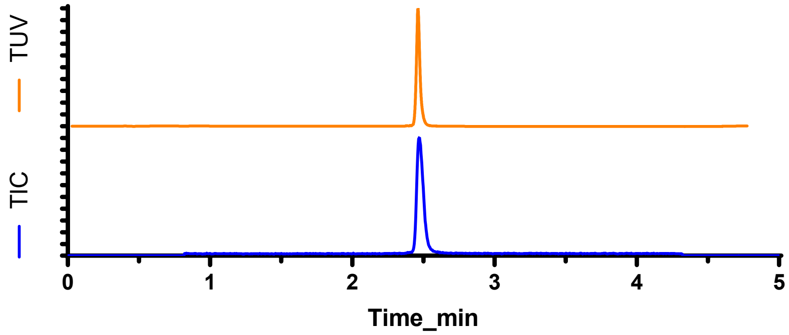

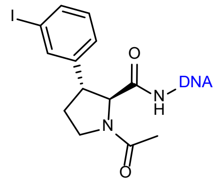

1

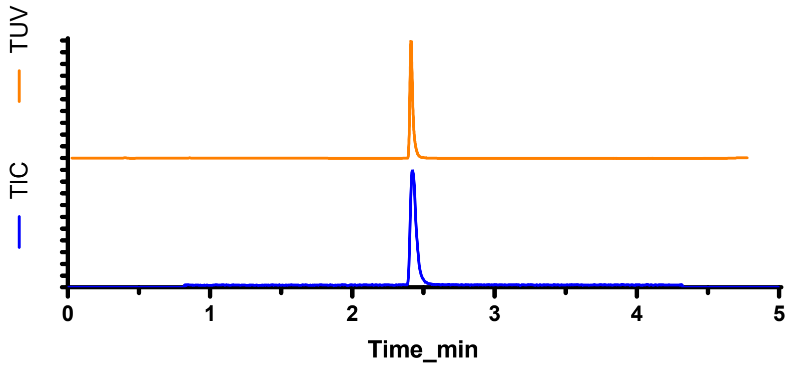

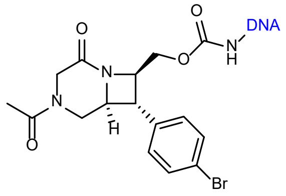

2

**Supplementary Fig. 5 | LCMS analysis of the example ArI and ArBr skeletons used for Suzuki coupling building block validation**

In 96-well plates, a relevant skeleton-AOP-HP construct (1 µL, 0.5 nmol, 0.5 mM, nuclease free water) was diluted with bicarbonate buffer (4 µL, pH 10, 0.1 M) for the ArI-containing construct, or phosphate buffer (4 µL, pH 8, 0.1 M) for the ArBr-containing construct. This was followed by the addition of boronic acid/ester (1 µL, 200 nmol, 200 mM, 400 eq, 1:1 MeCN:EtOH) and $PdCl_2$(dppf) (1 µL, 1 nmol, 2 eq, 1 mM, MeCN, freshly prepared). Plates were heat-sealed, vortexed briefly, spun down and heated to 90 °C for 1 hr before cooling to 4 °C. Each well was treated with diethylthiocarbamic acid sodium salt, (1 µL, 100 eq w.r.t. Pd, 100 mM, water). Plates were heat-sealed again, vortexed gently, spun down and stood at rt overnight. Reactions were EtOH precipitated, pellets dried in air and dissolved in water (50 µL) before analysis by LCMS.

ArBr couplings were less efficient than expected, with only ~20% meeting the criteria of >70% conversion to product). Generally, the non-product peaks corresponded to unreacted starting material and protodebromination. To increase the ArBr Suzuki validation rate, a second set of validation reactions were performed, adapting the conditions above to 667 eq of boronic acid/ester, 5 eq of $PdCl_2$(dppf).$CH_2Cl_2$ with heating at 90 °C for 2 h. This adaptation greatly improved the overlap between ArI and ArBr couplings which met criteria for inclusion in the final library build.

#### 2.3. Library synthesis

##### 2.3.1. Attachment of skeletons to headpiece

###### 2.3.1.1. General acylation protocol

Part 1: An appropriate 'skeleton' [Fmoc, Boc or Ns protected amine also bearing a carboxylic acid moiety] (1 eq) was dissolved to either 10 or 50 mM in DMF and mixed with a freshly prepared solution of DMTMM (0.9 eq, 100 mM, $H_2O$). The mixture was vortexed briefly, centrifuged, and stood at rt for approx 2 min.

Part 2: A solution of AOP-HP (0.03–0.07 eq, 1 mM in 0.1 M Ph 8 PB) was added to the intermediate mixture from part 1, vortexed briefly, centrifuged, and stood at rt until LCMS analysis indicated completion of reaction (typically 30 min – 20 hr). Skeletons were deprotected as outlined in **Supplementary Section 2.3.2**, then purified by microprep-HPLC.

###### 2.3.1.2. General carbamoylation protocol

Part 1: An appropriate 'skeleton' [Fmoc or Ns protected amine with a free hydroxyl group] (10 – 20 mg, 15-25 μmol, 1 eq) was dissolved in dry $CH_2Cl_2$ to a concentration of 50 mM. To the resulting solution $NEt_3$ (3 eq) was charged, followed by a 0.25 M solution of DSC (3 eq) in dry MeCN. The resulting solution was stirred overnight at rt in a 1-dram vial. Crude reaction mixture was loaded directly onto a 4 g prepacked silica column and eluted with heptane/EtOAc mixtures (to remove residual DSC and NHS). The major UV-absorbing species was collected, and in all cases recovery and purity was assumed to be 80%. The activated carbamates were dissolved in DMF to a 50 mM concentration.

Part 2: AOP-HP (1 mM in 0.1 M PB pH 8, 1 eq) was diluted with DMF (2.5 volumes relative to AOP-HP) before the addition of an activated carbamate (50 mM, 10 eq), such that the final reaction mixture contained 75% DMF by volume. Reactions were briefly mixed by benchtop vortexer and centrifuged.

Upon indication of completion by LCMS analysis (10 min – 1 hr) the reactions underwent EtOH precipitation and crude product amines were deprotected according to the protocols in **Section 2.3.2**. Following deprotection the desired products were purified by microprep-HPLC.

##### 2.3.2. Deprotection steps

###### 2.3.2.1. Fmoc

For a pellet of Fmoc-protected amine-DNA conjugate (derived from an EtOH precipitation): The pellet was treated with piperidine (10% in water), vortexed briefly, centrifuged, and stood at room temp for 10–30 min. For a crude reaction mixture containing Fmoc-protected amine-DNA conjugate (100%v/v): A 20% solution of piperidine in water (100%v/v) was added, vortexed briefly, centrifuged, and stood at rt for 10–30 min.

Another EtOH precipitation was performed, and the resulting pellet was dissolved in nuclease-free water at a concentration appropriate for microprep-HPLC purification (no more than 120 nmol per 100 µL; otherwise, UV detector signal saturation was highly likely). If any particulates presented from the solution these were removed by centrifugal filtration using Millipore ULTRAFREE-MC centrifugal filter units 0.22 µm GV DURAPORE (according to manufacturer guidelines) prior to microprep-HPLC purification.

###### 2.3.2.2. Nosyl

A pellet of Ns-protected amine-DNA conjugate (derived from an EtOH precipitation) was dissolved in bicarbonate buffer (0.1 M, pH 10) to a concentration of approx 0.1 mM with respect to the crude DNA-conjugate. A solution of 4-methyoxythiophenol (100 mM in DMF, 1000 eq) was then added and the mixture was heated at 80 °C for 30–60 min (when LCMS indicated complete reaction) with vigorous stirring by magnet. The mixture was cooled, then performed EtOH precipitation. The pellet was dissolved as much as possible in an appropriate volume of nuclease-free water for micro prep-HPLC purification (no more than 120 nmol per 100 µL; otherwise, UV detector signal saturation was highly likely). If any particulates presented from the solution these were removed by centrifugal filtration (as above) prior to microprep-HPLC purification.

##### 2.3.3. Tandem ligation of forward primer binding site and cycle1_tag

Target structures:

HP FWD primer binding site Cy1_tag

**5′ TGACTCCC AGGCCTGCTTCAATGTCGGATAGTG NNNNNNNCT 3′**

**3′ ACTGAG GGTCCGGACGAAGTTACAGCCTATC ACNNNNNNN 5′**

Skeleton-AOP-HP conjugates that were previously purified by microprep-HPLC followed by ultrafiltration had their concentrations normalized to 0.45 mM with nuclease free water; guided by photospectrometry. In a 2 mL deep well plate (Costar 2 mL Assay Block (3960, 96-well, square V-bottom, Sterile, Rnase/Dnase free), purified skeleton-AOP-HP conjugates (100 nmol; 0.45 mM; 222 µL; 1.15 eq) were added.

Forward primer binding site (86 nmol; 0.86 mM; 100 µL; 1.0 eq) was added as an annealed duplex in nuclease free water, followed by the appropriate skeleton tag/cycle 1 tag duplex (174 nmol; 1 mM; 174 µL; 2.0 eq) in nuclease free water. A freshly prepared ligation mixture* (300 µL) was then added. The deep-well plate was heat-sealed, gently vortexed, quickly centrifuged and stood at rt for 16 hr.

A 0.5 µL sample was taken from each reaction, diluted in 80 µL of water and analyzed alongside primer binding site duplex and ultralow weight DNA ladder (Invitrogen, 10488096) on 4% agarose E-gel (Invitrogen, G401004). All ligations were observed to be complete; with no detectable FWD primer binding site remaining and a solid band at ~40–45 base pairs relative to the ladder.

*Ligation mixture was composed of: 375 µL of 30 Weiss Units T4 ligase/µL (Invitrogen, EL0013), 3 mL of nuclease-free water (Ambion, AM9932) and 20 mL of 10x ligation buffer (Invitrogen).

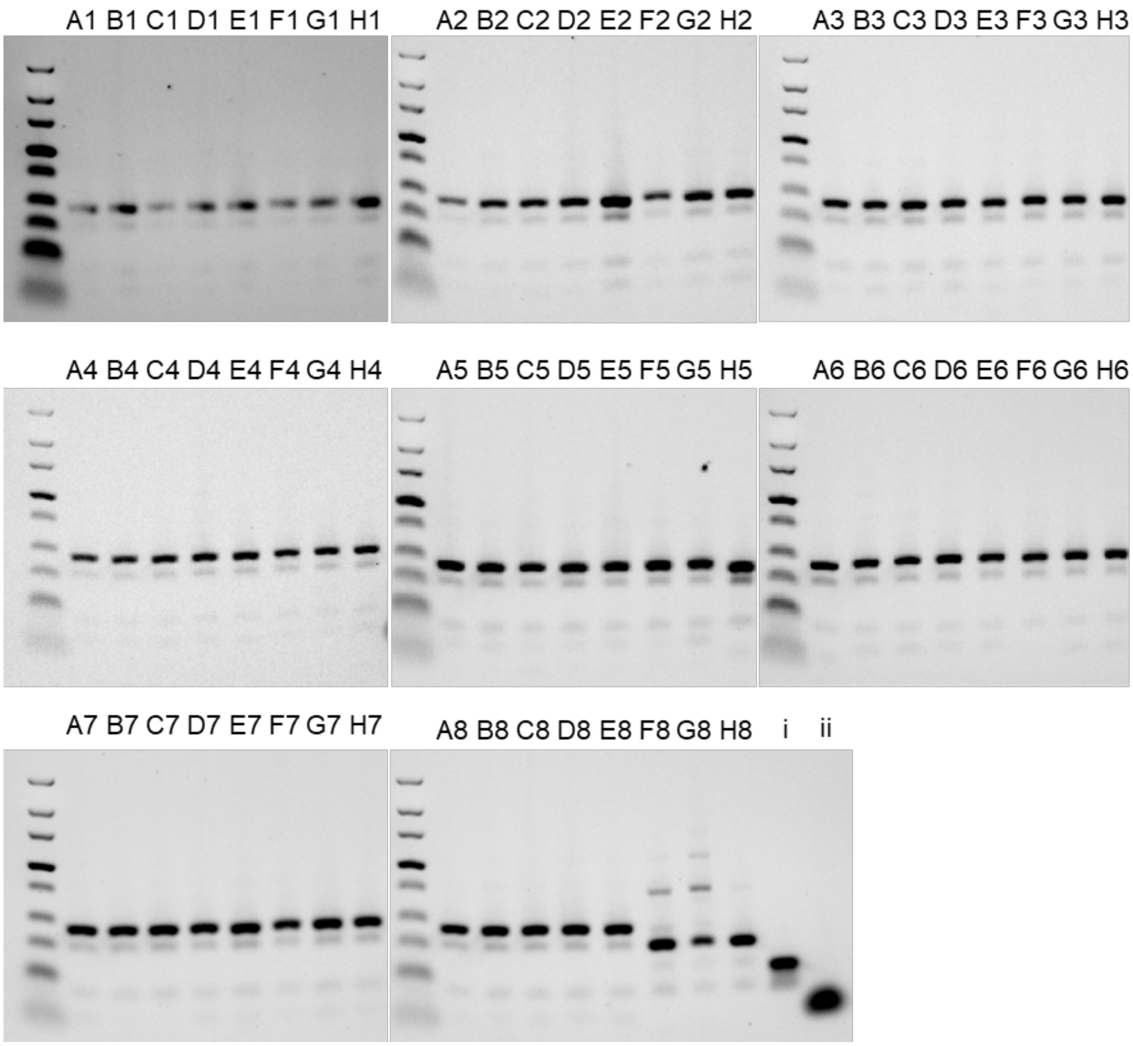

**Supplementary Fig. 6 | Analysis of tandem FWD primer binding site and Cy1_tag ligation products by 4% agarose electrophoresis**. F8–H8: no skeleton-headpiece construct present; i: FWD primer binding site duplex; ii: unmodified AOP-HP.

##### 2.3.4. Cycle 2 encoding tag ligations

Target structures:

HP FWD primer binding site Cy1_tag Cy2_tag

**5′ TGACTCCC AGGCCTGCTTCAATGTCGGATAGTG NNNNNNNCT NNNNNNNAC 3′**

**3′ ACTGAG GGTCCGGACGAAGTTACAGCCTATC ACNNNNNNN GANNNNNNN 5′**

Following ligation of FWD primer binding site and Cy1_tags, 174 nmol (2 eq relative to FWD primer binding site) of potentially ligation-competent species could be present in the mixture for each skeleton; therefore, >2 eq of Cy2_tag were required skeleton.

40 ArI-containing skeletons and 21 ArBr-containing skeletons were to be pooled separately for downstream synthetic steps.

40 x 86 nmol (maximum quantity of each encoded ArI) = 3.44 µmol of DNA

split between 300 wells = 11.5 nmol per well; requiring 25 nmol of Cy2 tag (2.2 eq) per ligation

21 x 86 nmol (maximum quantity of each encoded ArBr) = 1.801 µmol of DNA

split between 300 wells = 6 nmol per well; requiring 13 nmol of Cy2 tag (2.2 eq) per ligation

NOTE: initially planned to perform 300 Cy2_tag ligations, however available stocks of Cy2_tags were unexpectedly depleted, so 295 ligations were performed instead.

The two encoded skeleton pools were concentrated and desalted by ultrafiltration, then dissolved in nuclease free water to an assumed concentration of 0.9 mM with respect to pooled encoded skeleton constructs (3.8 mL and 2 mL for ArI and ArBr pools respectively). These 0.9 mM solutions were treated as 1.0 mM solutions for downstream processing to allow a 10% excess volume for pipetting purposes. A fresh ligation mixture was prepared comprising: 300 µL of 30 Weiss Units T4 ligase/µL, 3000 µL of nuclease free water and 15000 µL of 10x ligation buffer. Ligations were performed in Eppendorf Deepwell plates (951032808) with the indicated volumes of components per well:

ArI: 12 µL DNA (assuming ~1.0 mM wrt DNA conjugate of interest), 25 µL tag (1 mM, ~2.2 eq), ligation mixture (30 µL) as prepared above.

ArBr: 6 µL DNA (assuming ~1.0 mM wrt DNA conjugate of interest), 13 µL tag (1 mM, ~2.2 eq), ligation mixture (16 µL) as prepared above.

After plates were heat-sealed, mixed by benchtop vortexer and spun down briefly, the ligations were stored for 16 hr at 22 °C. A 1.0 µL sample was taken from each ligation and, in a fresh 96-well plate, was diluted in 45 µL (for ArI-pooled constructs) or 32 µL (for ArBr-containing constructs) of nuclease-free water. Differential dilutions were performed to maintain equivalent absolute concentration with respect to total DNA in each sample. From these sample plates each row of 12 ligations was pooled into a single well (5 μL taken from each of the diluted ligations) to allow for a whole plate of ligations to be analyzed on a single 10-lane e-gel. Of these pooled mixtures, 2 μL was mixed with 18 μL of e-gel loading buffer (Invitrogen, 10482055) for analysis.

All crude ligations (after sampling) underwent EtOH precipitation (6 μL of 5 M NaCl, followed by 400 μL EtOH, and incubation at -20 °C for 2 days). After centrifugation at 3750 rcf for 20 min and removal of supernatant, an additional 400 μL of cold 80% aq. EtOH was added to the pellets. The plates heat-sealed, mixed by benchtop vortexer, spun down quickly and chilled at -20 °C overnight. Plates were centrifuged again at 3750 rcf for 20 min, supernatant was discarded, and plates were dried in air.

Concurrent with EtOH precipitation ligation performance was analyzed for both sets of ArI and ArBr plates. A feint band under the desired 'product' band, co-running with the starting material, was present in all lanes, suggestive of residual Cy1_tag ligation product. However, this had been observed numerous times in other experiments [data not reported] and is believed to be the product of a single strand ligation (due to these species not being competent in further ligations, and LCMS analysis of un-pooled ligations indicating a deconvoluted mass concurrent with ssDNA ligation). To further assess this possibility, denaturing PAGE was also performed (15% PAGE with 6 M urea). Denaturing PAGE showed additional faint bands when compared to the native agarose gel images (**Supplementary Fig. 8**), but the [assumed] ssDNA ligation product still could not be clearly resolved from the starting material. After denaturing PAGE, the bands were quantified using Bio-Rad Image Lab to show ~1-3% of the faint band (either SM or ssDNA ligation). 3% starting material, in the worst case, would suggest that a single ligation in the pool of 12 contained ~36% unligated material. To assess the variability, a lane containing ~3% of the faint band was used for further study (**Supplementary Fig. 7** ArI-1 row H [lane 8]). Each ligation from this pool was run individually on 4% agarose gel, imaged, and quantified as for denaturing PAGE, to show very consistent ligations, with generally only ~2% of this faint band present (**Supplementary Fig. 8**).

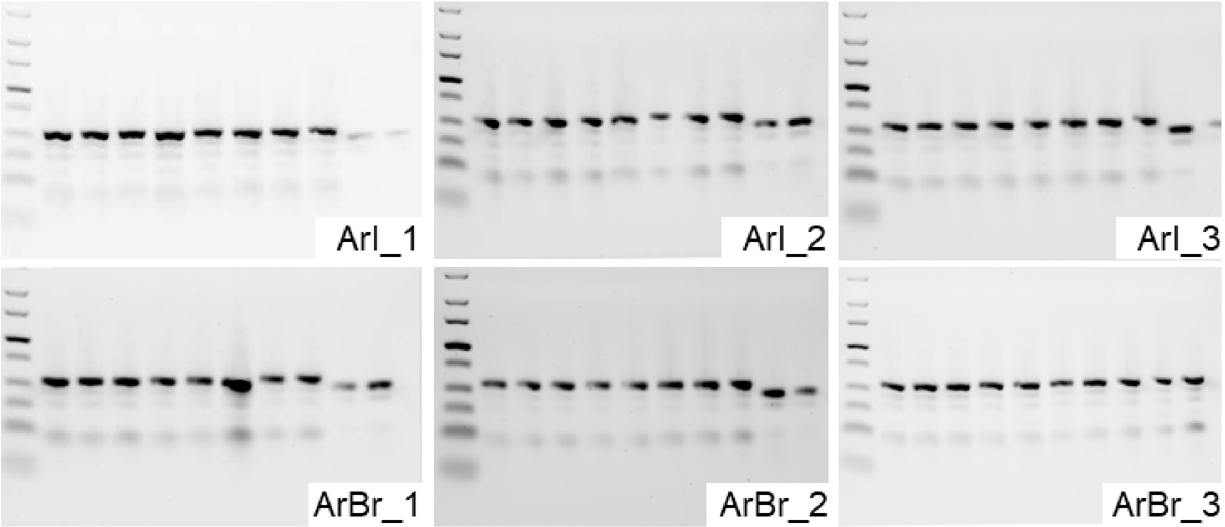

**Supplementary Fig. 7 | Analysis of Cy2_tag ligations by 4% agarose e-gel.** One lane contains a pooled sample from a single row of a 96-well plate. The gel image labelled as "ArBr_3" also contains the remaining 7 pooled ligations from ArI-plate4 in lane 9 and the remaining 7 pooled ligations from ArBr-plate4 in lane 10. Otherwise, lanes 9 and 10 of each gel both contain the product of tandem Cy1_tag and FWD primer binding site ligations for comparison.

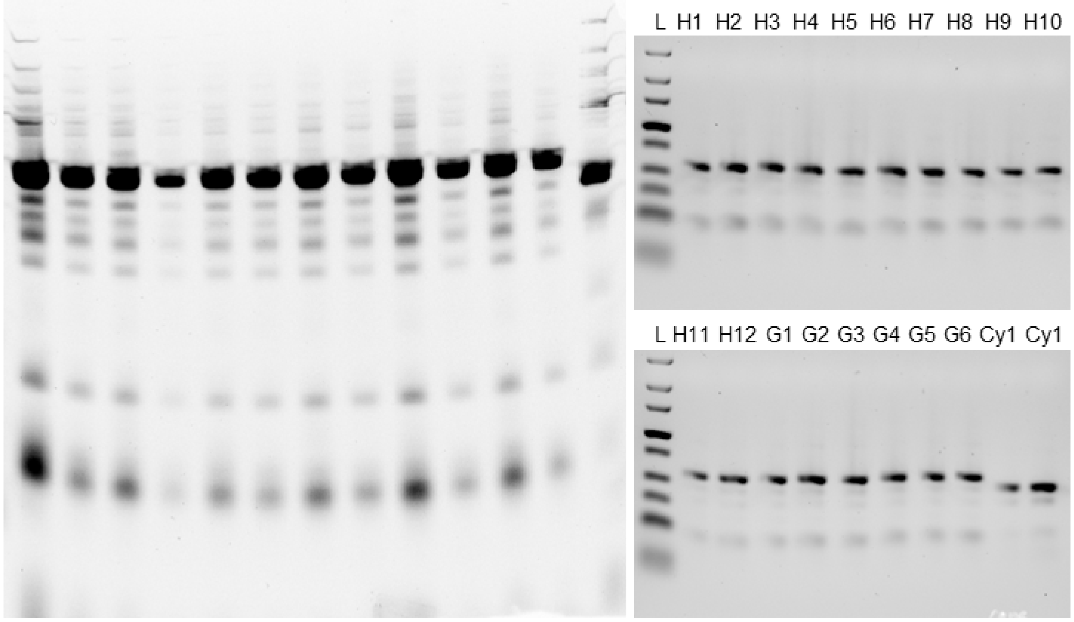

**Supplementary Fig. 8 | Examples of pooled samples denaturing PAGE and subsequent individual sample analysis by native 4% agarose electrophoresis for cycle 2 tag ligations.** Left: Denaturing

PAGE of ArI_1 (rows A-H) and ArI_2 (rows A-D) [lanes 1-12], lane 13 = product of CY1_tag ligation. Right: 4% agarose e-gel analysis of individual ligations from the ArI_1 plate, row H and some examples of row G (pooled samples of which are shown in lanes 8 and 7 of left denaturing PAGE gel respectively).

##### 2.3.5. Amine capping reactions

Following ligation of the Cy2_tags, EtOH precipitation, and thorough drying of pellets in air, amine-capping reactions were performed; at two different scales for the compounds derived from ArI and ArBr containing skeletons.

Acylation conditions (ArI):

DMTMM (10.3 µL, 100 mM, water, freshly prepared, 90 eq) and carboxylic acid (23 µL, 50 mM, DMF, 100 eq) were mixed in a 96-well twin.tech plate, then after approx 5 min DNA-amine (11.5 µL, 11.5 nmol, ~1 mM, 0.1 M pH 8 PB, 1 eq) was added. The plate was heat-sealed, gently vortexed, then spun down before incubating at 22 °C for 16 hr.

Reductive amination conditions (ArI):

DNA-amine (11.5 µL, 11.5 nmol, ~1 mM, 0.1 M pH 8 PB, 1 eq) was diluted with acetate buffer (58.5 µL, 1 M, pH 5) in a 96-well twin.tech plate, and aldehyde (23 µL, 200 mM, MeCN, 400 eq) was then added. After approx 5 min $NaCNBH_3$ (17.3 µL, 200 mM, $H_2O$, 300 eq) was added. The plate was heat-sealed, gently vortexed, then spun down before incubating at 22 °C for 16 hr.

Sulfonylation conditions (ArI):

To a 96-well twin.tech plate was added DNA-amine (11.5 µL, 11.5 nmol, ~1 mM, 0.1 M pH 8 PB, 1 eq), phosphate buffer (23 µL, 0.1 M, pH 8) and sulfonylchloride (11.5 µL, 200 mM, MeCN, 200 eq). The plate was heat-sealed, gently vortexed, then spun down before incubating at 22 °C for 16 hr.

Common work-up (ArI):

Piperidine (20 µL, 10%v/v in $H_2O$) was added to each well, then after approx 2 min each reaction was pooled and concentrated by ultrafiltration (10 kDa MWCO). After 4 hr at 3750 rcf membranes were washed twice with 15 mL of nuclease free water, spinning at 3750 rcf for 1 hr at each wash.

Acylation conditions (ArBr):

DMTMM (5.4 µL, 100 mM, water, freshly prepared, 90 eq) and carboxylic acid (12 µL, 50 mM, DMF, 100 eq) were mixed in a 96-well twin.tech plate, then after approx 5 min DNA-amine (6 µL, 6 nmol, ~1 mM, 0.1 M pH 8 PB, 1 eq) was added. The plate was heat-sealed, gently vortexed, then spun down before incubating at 22 °C for 16 hr.

Reductive amination conditions (ArBr):

DNA-amine (6 µL, 6 nmol, ~1 mM, 0.1 M pH 8 PB, 1 eq) was diluted with acetate buffer (34 µL, 1 M, pH 5) in a 96-well twin.tech plate, and aldehyde (12 µL, 200 mM, MeCN, 400 eq) was then added. After approx 5 min $NaCNBH_3$ (9 µL, 200 mM, $H_2O$, 300 eq) was added. The plate was heat-sealed, gently vortexed, then spun down before incubating at 22 °C for 16 hr.

Sulfonylation conditions (ArBr):

To a 96-well twin.tech plate was added DNA-amine (6 µL, 6 nmol, ~1 mM, 0.1 M pH 8 PB, 1 eq), phosphate buffer (12 µL, 0.1 M, pH 8) and sulfonylchloride (6 µL, 200 mM, MeCN, 200 eq). The plate was heat-sealed, gently vortexed, then spun down before incubating at 22 °C for 16 hr.

Common work-up ArBr:

Piperidine (10 µL, 10%v/v in $H_2O$) was added to each well, then after approx 2 min each reaction was pooled and concentrated by ultrafiltration (10 kDa MWCO). After 4 hr at 3750 rcf membranes were washed twice with 15 mL of nuclease free water, spinning at 3750 rcf for 1 hr at each wash.

Following the common work up procedures two pools were obtained which contained some insoluble yellow particulates. These was filtered by centrifugation (Millipore ULTRAFREE-MC centrifugal filter units 0.22 µm GV DURAPORE). EtOH precipitation was then performed and resulting DNA pellets were dried under high vacuum for 1 hr before dissolution in nuclease free water (7.57 mL [ArI pool] and 3.97 mL [ArBr pool], ~0.5 mM [with 10% excess to enable pipetting]).

##### 2.3.6. Cycle 3 encoding tag ligations

Target structures:

```
HP FWD primer binding site Cy1_tag Cy2_tag Cy3_tag
5′  TGACTCCC AGGCCTGCTTCAATGTCGGATAGTG NNNNNNNCT NNNNNNNAC NNNNNNNTT  3′
3′  ACTGAG GGTCCGGACGAAGTTACAGCCTATC ACNNNNNNN GANNNNNNN TGNNNNNNN  5′
```

207 ligations were required for Cy3_tag/Suzuki coupling, including the encoding of null reactions (i.e., without boronic acid or ester) in the presence of Pd-catalyst and heating, or in the absence of Pd-catalyst without heating.

ArI: 3.44 µmol of DNA (assuming no loss of material at any stage of synthesis), split between 207 wells = 16.6 nmol per well; requiring 38 nmol of Cy3_tag (2.3 eq)

ArBr: 1.801 µmol of DNA (assuming no loss of material at any stage of synthesis), split between 207 wells = 8.7 nmol per well; requiring 20 nmol of Cy3_tag (2.3 eq).

A fresh ligation mixture was prepared comprising: 300 µL of 30 Weiss Units T4 ligase/µL, 3000 µL of nuclease free water and 15000 µL of 10x ligation buffer. Ligations were performed in Eppendorf Deepwell plates (951032808) with the indicated volumes of components per well:

ArI: DNA (33.2 µL, 16.6 nmol, ~1.0 mM wrt DNA conjugate of interest), Cy3_tag (38 µL, 38 nmol, 1 mM, ~2.3 eq), ligation mixture (35 µL) as prepared above.

ArBr: DNA (17.4 µL, 8.7 nmol, ~1.0 mM wrt DNA conjugate of interest), Cy3_tag (20 µL, 20 nmol, 1 mM, ~2.3 eq), ligation mixture (19 µL) as prepared above.

After plates were heat-sealed, mixed by benchtop vortexer and spun down briefly, the ligations were stored for 16 hr at 22 °C. A 1.0 µL sample was taken from each ligation and, in a fresh 96-well plate, was diluted in 45 µL (for ArI-pooled constructs) or 32 µL (for ArBr-containing constructs) of nuclease-free water.

Samples were taken for ligation efficiency analysis by electrophoresis as described in Section 2.3.4 (gel images shown below in **Supplementary Fig. 9**).

All crude ligations were EtOH precipitated by the addition of 5 M NaCl (15 µL), followed by EtOH (400 µL), then storing at -20° for 2 days. Two washes with 80% EtOH (400 µL) in water were performed, and the pellets were dried thoroughly in air.

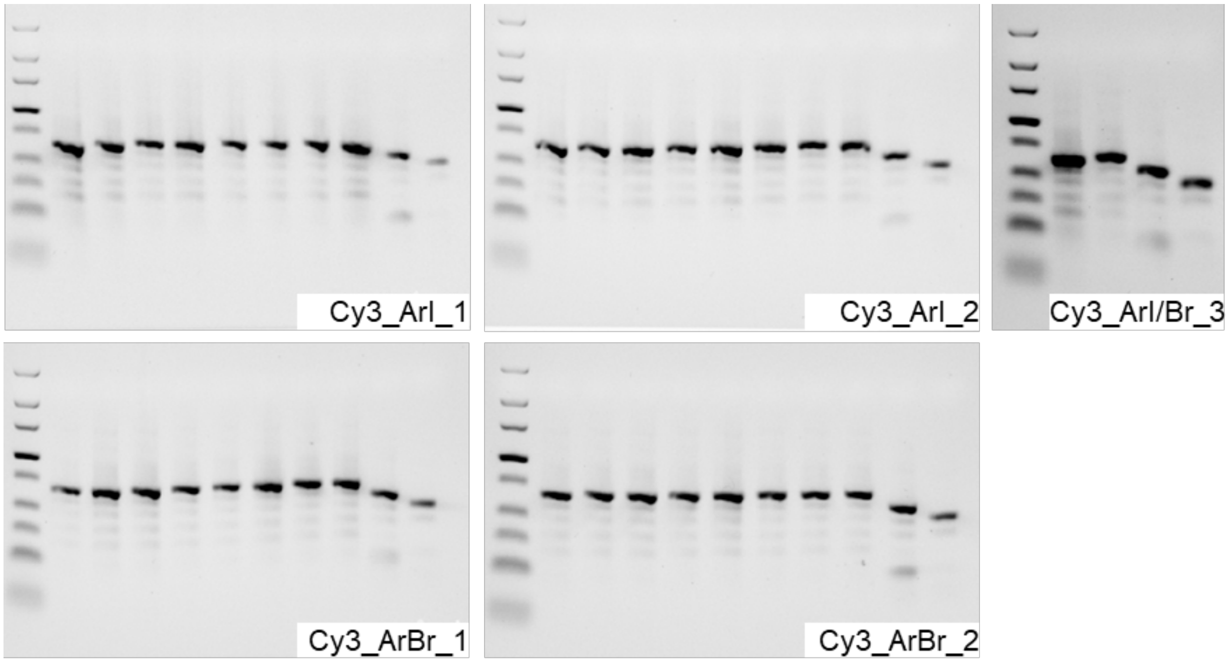

**Supplementary Fig. 9 | Analysis of Cy3_tag ligations by 4% agarose e-gel.** One lane contains a pooled sample from a single row of a standard 96-well plate. The gel image labelled as "Cy3_ArI/ArBr_3" contains the pooled ligations from ArI-plate3 in lane 1 and the pooled ligations from ArBr-plate3 in lane 2. Otherwise, the penultimate and final lanes shown on the right of each gel image contain the product of Cy2_tag ligation and product of tandem Cy1_tag and FWD primer binding site ligations, respectively, for comparison.

##### 2.3.7. Suzuki coupling

Product pellets derived from Cy3_tag ligations outlined in Section 2.3.6 were dissolved in the reaction buffers below (in their existing deepwell plates) and spun down before transferring to Eppendorf twin.tec plates and adding the other components. For stoichiometry/amount calculation, 100% recovery across all steps of synthesis up to this point was assumed. After addition of all components plates were sealed with heat-sealing foils, mixed by benchtop vortexer, spun down and heated to 90 °C for 1 hr (ArI) or 2 hr (ArBr), then cooled to 4 °C.

<u>ArI:</u>

DNA (83 µL, 16.6 nmol, 0.2 mM, 0.1 M bicarb buffer pH 10)

Boronic ester/acid (33.2 µL, 200 eq, 100 mM, 1:1 MeCN:EtOH)

$PdCl_2$(dppf) (33.2 µL, 2 eq, 1 mM, MeCN)

<u>ArBr:</u>

DNA(43.5 µL, 8.7 nmol, 0.2 mM, 0.1 M phosphate buffer pH 8)

Boronic ester/acid (29 µL, 333 eq, 100 mM, 1:1 MeCN:EtOH)

$PdCl_2$(dppf) (43.5 µL, 5 eq, 1 mM, MeCN)

To each well was added diethylthiocarbamic acid sodium salt (30 µL, 200 mM, water), plates were sealed, vortexed, and quickly spun down before standing at rt for 16 hr. All reactions were pooled (still maintaining ArI and ArBr-derived constructs as separate pools) before performing the following steps:

1) Performed centrifugation at 3750 rcf for 2 hr and collected supernatant

2) Washed pellet with water, vortexed briefly, performed centrifugation again at 3750 rcf for 1 hr then collected supernatant again

3) EtOH precipitation (added 5% v/v 5M NaCl, then 300%v/v EtOH, chilled in dry ice for 1 hr, centrifugation at 3750 rcf for 30 min at 4 °C and removed supernatant)

4) Washed pellet with cold 80% EtOH in water, chilled on dry ice for 1 hr, performed centrifugation at 3750 rcf for 30 min at 4 °C, removed supernatant and dried pellet under high vacuum.

5) Dissolved dried DNA pellet (still as separate ArI and ArBr-derived pools) in 1.5 mL of water and centrifuged at 14000 rcf, collecting supernatant. Pellet was then treated with 400 µL water, mixed vigorously by benchtop vortexer, and centrifuged at 14000 rcf again, combining supernatant with that of step 4.

The two pooled product solutions were diluted to 3.44 mL (iodides) and 1.80 mL (bromides) with nuclease free water to afford 1 mM solutions with respect to the desired encoded constructs; assuming no loss of material during library synthesis.

##### 2.4. Small scale library closure

Prior to large scale completion of the DOSEDO library (i.e., pooling the ArI and ArBr-derived sub-libraries, ligation of a REV primer binding site, and purification), a small-scale test was first performed as outlined below.

Target structures ("|" indicating ligation sites):

HP FWDprimerBS Cy1_tag Cy2_tag Cy3_tag Cy4-UMI-REVprimerBS

**5′TGACTCCC|AGGCCTGCTTCAATGTCGGATAGTG|NNNNNNNCT|NNNNNNNAC|NNNNNNNTT|CCGTGCTTGA**

**NNNNNNNNNNNNCTGATGGAGGTAGAAGCCGC 3′**

**3′ACTGAG|GGTCCGGACGAAGTTACAGCCTATC|ACNNNNNNN|GANNNNNNN|TGNNNNNNN|AAGGCACGAACT**

**5′**

ArI-derived pool (3.44 µL, 3.44 nmol) and ArBr-derived pool (1.80 µL, 1.80 nmol) were mixed in a 1.5 mL lo-bind tube (Eppendorf). To the pooled library (5.24 µL, 5.24 nmol) nuclease free water was added (5 µL), 10x ligation buffer (9 µL, Invitrogen), and annealed Cy4-UMI-REVprimerBS duplex [see above] (13.1 µL, 1 mM, 2.5 eq). Then was added 30 U/µL T4 DNA ligase (0.5 µL, Invitrogen), the mixture was vortexed, quickly spun down and stood at rt for 16 hr. A 0.5 µL sample of the crude ligation was diluted in 70 µL nuclease free water and analyzed alongside the pooled ligation samples from during library synthesis (**Supplementary Fig. 11**).

A 28 mm internal diameter Model 491 prep cell gel tube assembly was cast with 4% agarose in 1X TBE buffer to 9.25 cm and left to cure over 2 hr with circulating room temp water through the cold finger at ~100 mL/min. The prep cell was assembled according to manufacturer's instructions. After assembly, and using 1x TBE as running buffer, sample was loaded in 1x Novex TBE high density loading buffer with xylene cyanol and bromophenol blue as dyes. Electrophoresis was performed at 12 W constant power. Fraction collection was started after approx 1 hr (when the faster-running dye reached the end of the gel column), with a flow rate of approx 1 mL/min and 2 min fractionation. Each fraction was analyzed by 4% agarose e-gel (4 µL samples of each fraction diluted in 16 µL of e-gel loading buffer). Acceptably clean product containing fractions (25–30) were pooled before concentration and desalting by ultrafiltration, washing twice with nuclease free water. The purified desalted concentrate (250 µL) was collected and quantified by Qbit and NanoDrop indicating a yield of 1.0–1.7 nmol (19–37% assuming quantitative recovery through all steps of synthesis and purification).

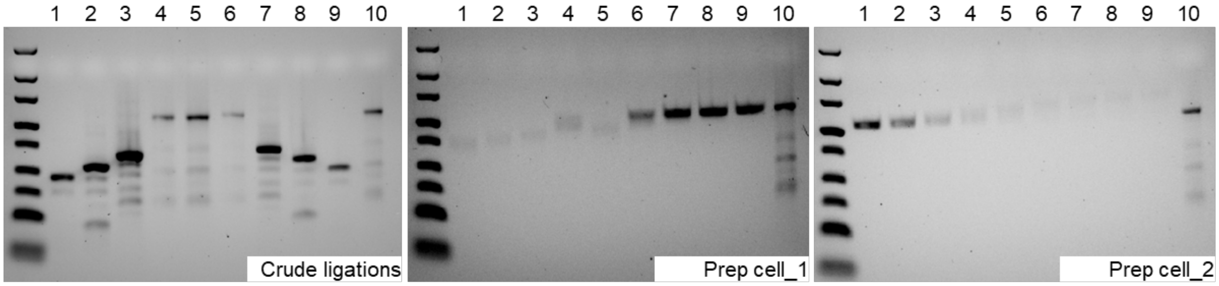

**Supplementary Fig. 10 |** Crude ligations: lanes 1/9, 2/8 and 3/7 show the crude products of Cy1_tag, Cy2_tag and Cy3_tag ligations. Lanes 4, 5, 6 and 10 show the crude product of Cy4-UMI-REVprimerBS ligation. Prep cell_1 and Prep cell_2 show individual fraction contents after prep cell purification alongside the crude Cy4-UMI-REVprimerBS duplex ligation product in Lane 10. Prep cell_1 lanes 6, 7, 8 and 9 plus Prep cell_2 lanes 1, 2 and 3 are fractions 25 – 30, which were pooled for further work (note: lanes 1 and 2 of Prep cell_2 gel are the same samples [at a slightly lower concentration] as those in lanes 8 and 9 of Prep Cell_1 gel image).

##### 2.5. Large scale library closure

Target structures ("|" indicating ligation sites):

HP FWDprimerBS Cy1_tag Cy2_tag Cy3_tag Cy4-UMI-REVprimerBS

**5′ TGACTCCC | AGGCCTGCTTCAATGTCGGATAGTG | NNNNNNNCT | NNNNNNNAC | NNNNNNNTT | CCGTGCTTGA NNNNNNNNNNNNCTGATGGAGGTAGAAGCCGC 3′**

**3′ ACTGAG | GGTCCGGACGAAGTTACAGCCTATC | ACNNNNNNN | GANNNNNNN | TGNNNNNNN | AAGGCACGAACT 5′**

Based on input sequence analysis of the small scale DOSEDO_v1 library closure, the assumed 1:1 pooling of ArI- and ArBr-derived sub-libraries afforded a 1.31:1 (ArI:ArBr) ratio of sequences at the individual skeleton level (on average). To correct for the imperfect pooling of sub-libraries 407.5 µL of the previously assumed 1 mM crude ArBr sub-library sample was pooled with 592.5 µL of the ArI sub-library (also previously assumed to be 1 mM) to give 1 mL of a new pool. Based on there being 40 ArI skeletons and 21 ArBr skeletons included in the DOSEDO library, this suggests a 1.31-fold correction, as desired.

Assuming the 1 mL pool of sub-libraries was at a concentration of 1 mM (as initially intended), annealed closing duplexes (2.5 mL, 1 mM, 2.5 eq) were added followed by 30 U/µL T4 DNA ligase (100 µL, ~3 Weiss units per nmol template DNA, Invitrogen) in 10 x ligation buffer (3 mL, Invitrogen). The mixture was vortexed, quickly centrifuged and stood at rt until analysis by 4% agarose e-gel electrophoresis indicated complete ligation (approx 1 hr). EtOH precipitation was performed, with pellets being dried under high vacuum before dissolving in minimal nuclease free water.

A 28 mm internal diameter Model 491 prep cell gel tube assembly was cast with 4% agarose in 1X TBE buffer to 9.0 cm and left to set over 2 hr with circulating room temp water through the cold finger at ~100 mL/min. a thin layer of 1:1 water:IPA was overlaid on the gel, which was stood overnight without further buffer recirculation through the cold finger. The prep cell was assembled according to manufacturer's instructions. After assembly, and using 1x TBE as running buffer, sample was loaded in Novex TBE high density loading buffer (5x, LC6678, Invitrogen) with xylene cyanol and bromophenol blue as dyes. Electrophoresis was performed at 12 W constant power. Fraction collection was started after approx 1 hr (when the faster-running dye reached the end of the gel column), with a flow rate of approx 1 mL/min and 2 min fractionation. Total electrophoresis run time was 4.5 hr. fractions were analyzed by 4% agarose e-gel. Acceptably clean product containing fractions (94–120) were pooled before concentration and desalting by ultrafiltration, washing twice with nuclease free water. The purified and desalted concentrate was collected and quantified by Qbit and NanoDrop indicating a yield of 167–247 nmol (17-25% assuming quantitative recovery through all steps of synthesis and purification).

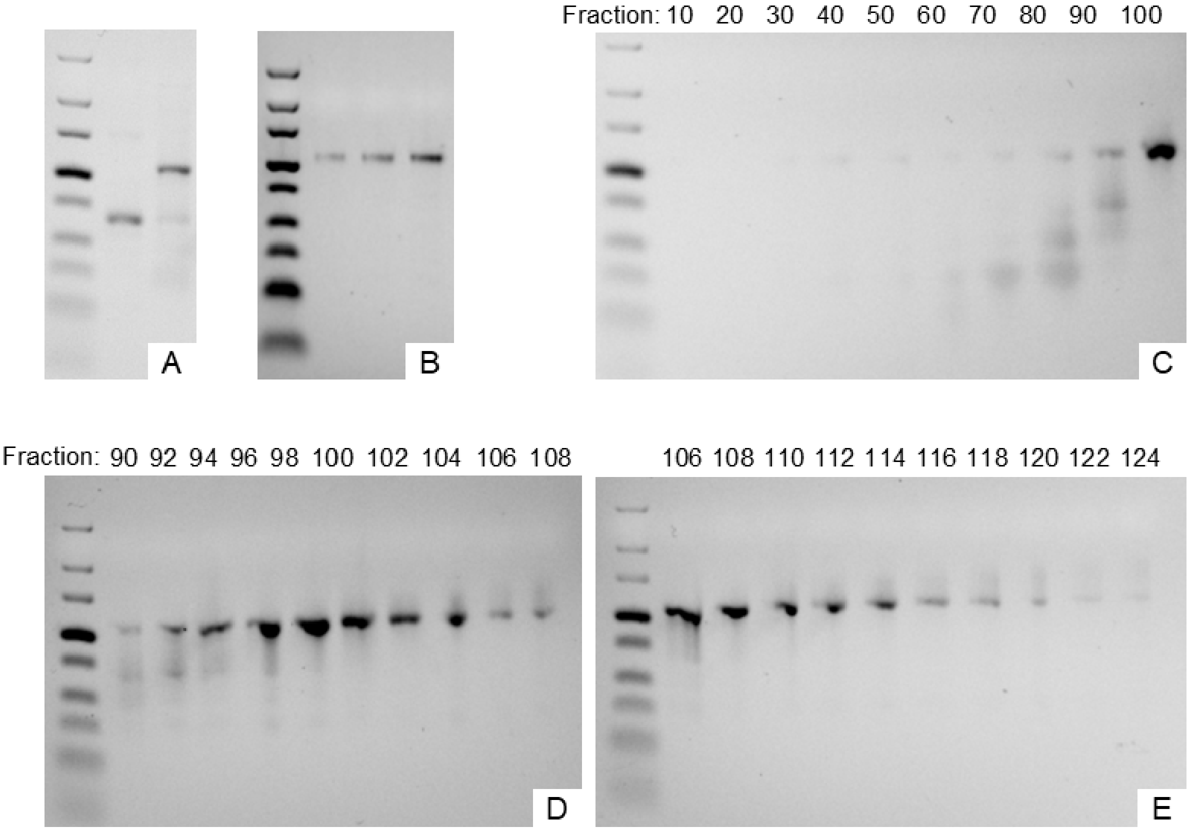

**Supplementary Fig. 11 | 4% agarose e-gel analysis of ligation samples and fractions from prep cell purification.** A: lane 1 – crude product of Cy3_tag ligation, lane 2 – crude product of Cy4-UMI-REVprimerBS ligation; B: increasing concentration of the prep cell purified closed DOSEDO library (pooled fractions); C: Assessment of every $10^{th}$ fraction during prep cell purification of the closed DOSEDO library; D and E: Assessment of every $2^{nd}$ fraction for the region containing desired DOSEDO library product.

#### 3. Preparation of next generation sequencing libraries – Supplementary Figures

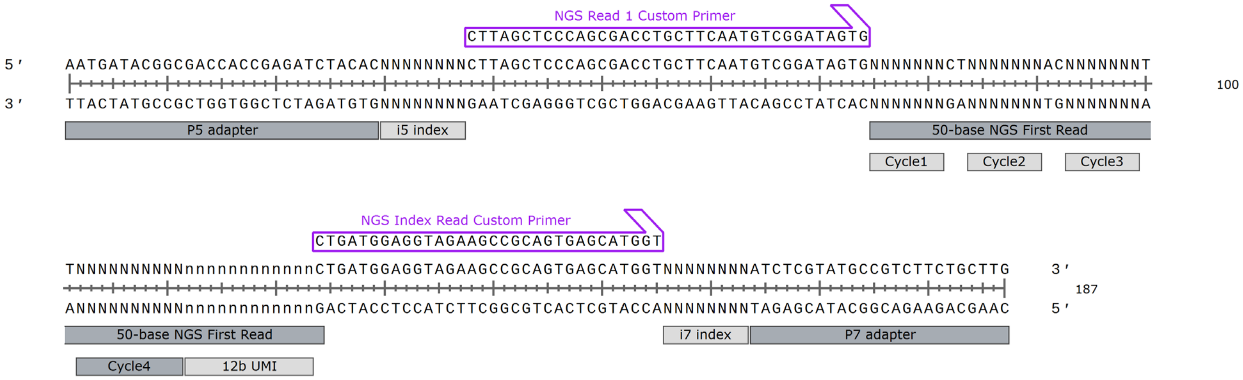

**Supplementary Fig. 12 | Overview of DOSEDO coding scheme as prepared for NGS**. Prepared with SnapGene Viewer.

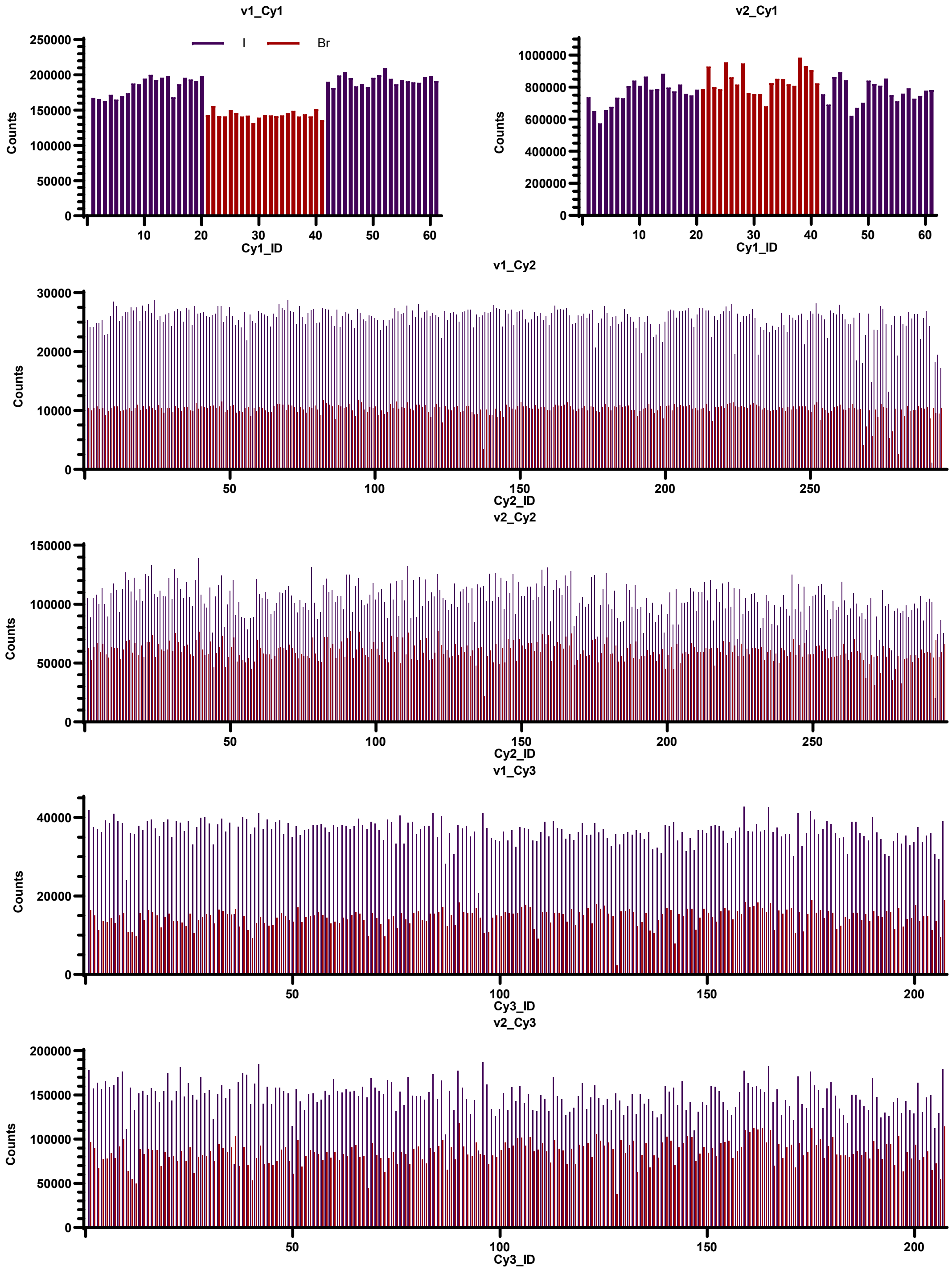

**Supplementary Fig. 13 | DOSEDO v1 and v2 input analysis aggregating on cycle 1, 2 or 3 tags.**

Ordered as plated during synthesis.

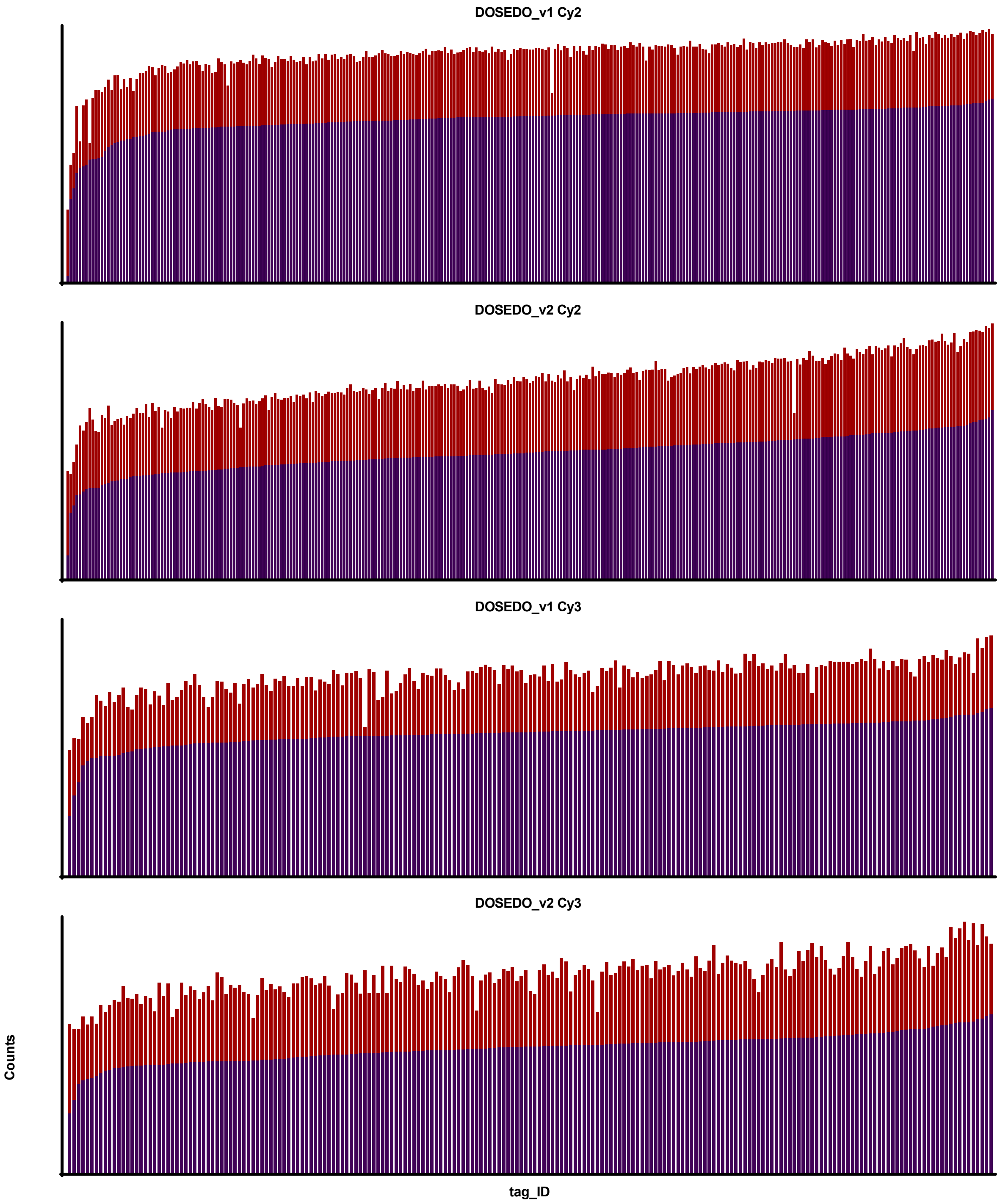

**Supplementary Fig. 14 | Comparison of DOSEDO v1 and v2 cycle 2 and 3 tag ligation product representation.** Sorted from least to most represented in the iodide-derived sub-library (purple) with corresponding aggregated count for the bromide-derived sub-library stacked (orange)

### 4. DEL screening – Supplementary Figures

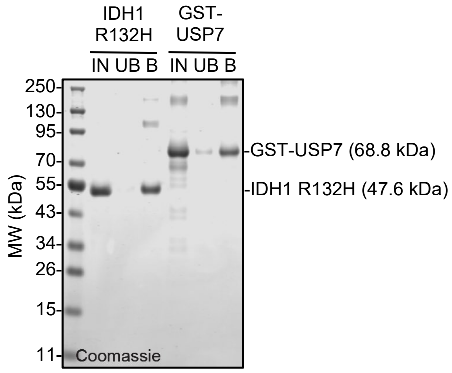

**Supplementary Fig. 15 | SDS-PAGE gel to access protein immobilization to the beads during the DEL screen.** IDH1 R132H and GST-USP7 fully bound the beads with little unbound. IN: input, 1.2 μM protein; UB: unbound, protein remaining in the supernatant after incubation of protein and beads; B: bound, beads after the screen heated and treated with NuPAGE™ LDS Sample Buffer to release the protein that bound the beads. All lanes are comparable.

### 5. Assay details – Supplementary Figures and Tables

**Supplementary Table 10 | Summary of carbonic anhydrase inhibition data obtained according to method highlighted above**

| cpd_id | beads_ppm | caix_ppm | er | er_lb | IC50, nM |
|---|---|---|---|---|---|
| **59** | 2 | 2248 | 1020 | 504 | 16 |
| **45** | 0 | 11 | 70 | 9 | 26 |
| **46** | 2 | 0 | 0 | 0 | 279 |
| **52** | 3 | 0 | 0 | 0 | 64 |
| **54** | 0 | 9 | 58 | 8 | 34 |
| **56** | 0 | 207 | 1348 | 198 | 2718 |
| **57** | 0 | 264 | 1716 | 253 | 31 |
| **58** | 0 | 922 | 5997 | 886 | 17 |
| **53** | 4 | 0 | 0 | 0 | > 10000 |

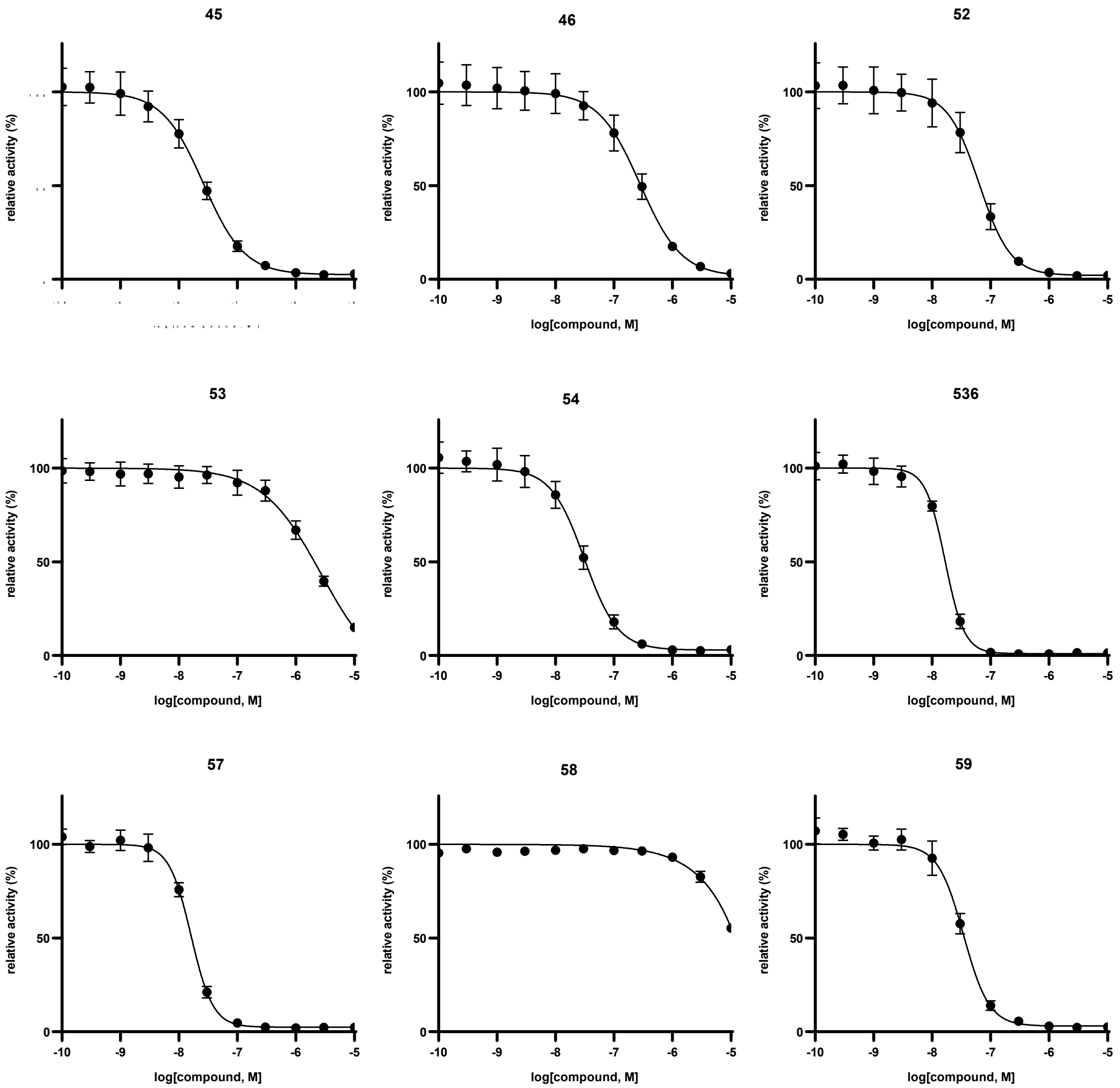

**Supplementary Fig. 16 | Carbonic anhydrase dose-response curves generated as described above**

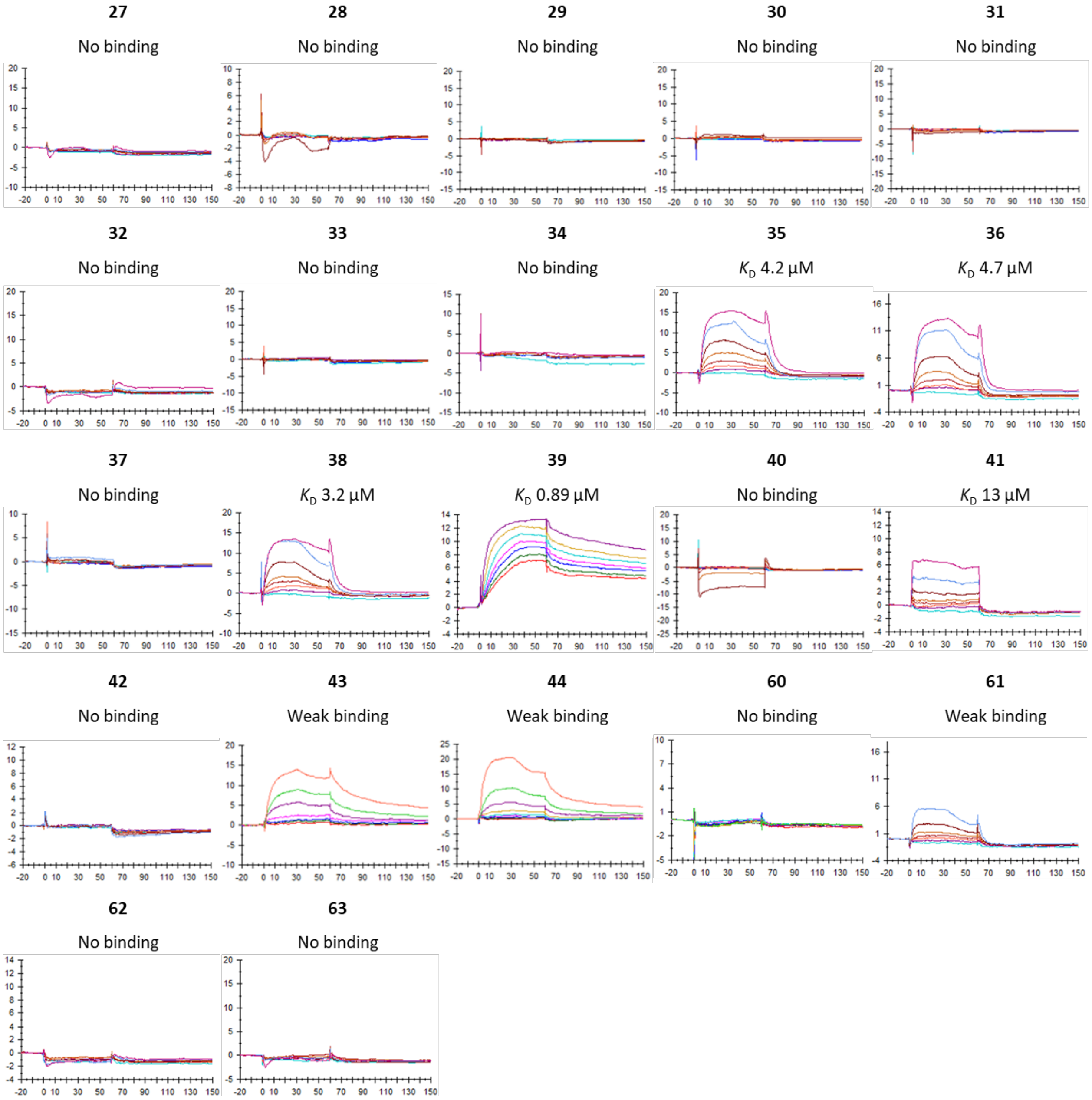

**Supplementary Fig. 17 | SPR sensograms of off-DNA DEL compounds binding to biotinylated IDH1 R132H**. The binding results were shown with $K_D$ values, weak binding (indicating signs of binding but KD >20 µM) or no binding at the tested concentrations.

**3 binding to GST-tagged USP7** **3 binding to GST**

$K_D$ 7.7 µM $K_D$ 10 µM

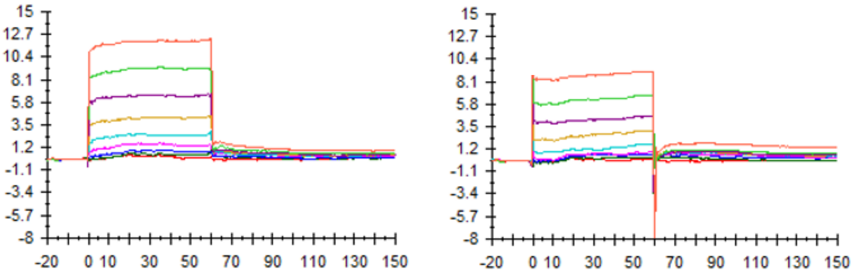

**Supplementary Fig. 18 | SPR sensograms of compound 3 binding to GST-tagged USP7 and GST with KDs of 7.7 µM and 10 µM respectively**

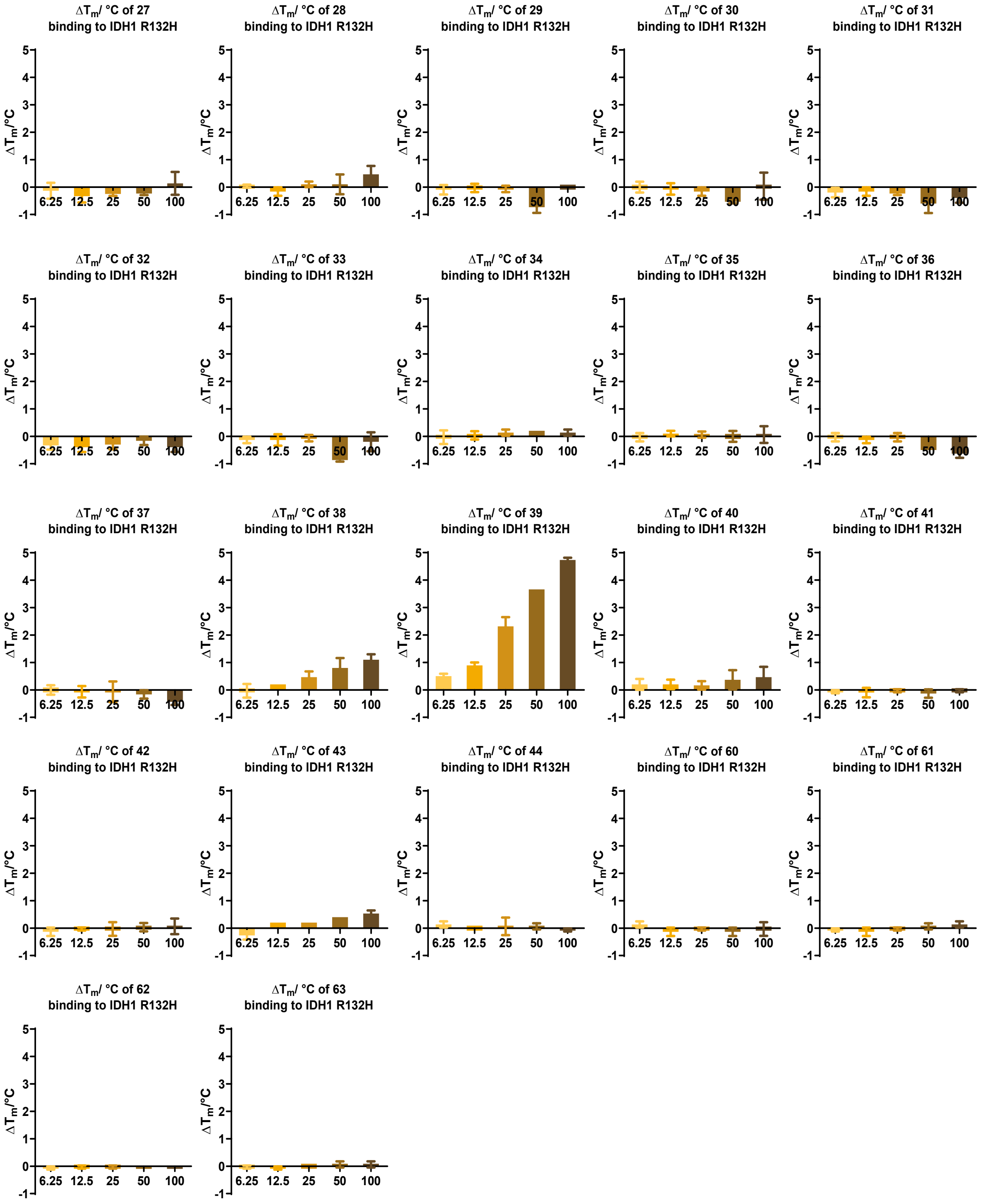

**Supplementary Fig. 19 | DSF results for off-DNA DOSEDO DEL compounds with IDH1 R132H in 50 mM Tris-HCl, pH 7.4.** x-axis is concentration of names compound in all cases, in μM. Compound **39** induces thermal stabilization of IDH1 R132H in a dose dependent manner, up to 4.7 °C at 100 μM. All other compounds showed no or weak thermal stabilization of IDH1 R132H. Data are mean ± SD, n= 3 technical replicates.

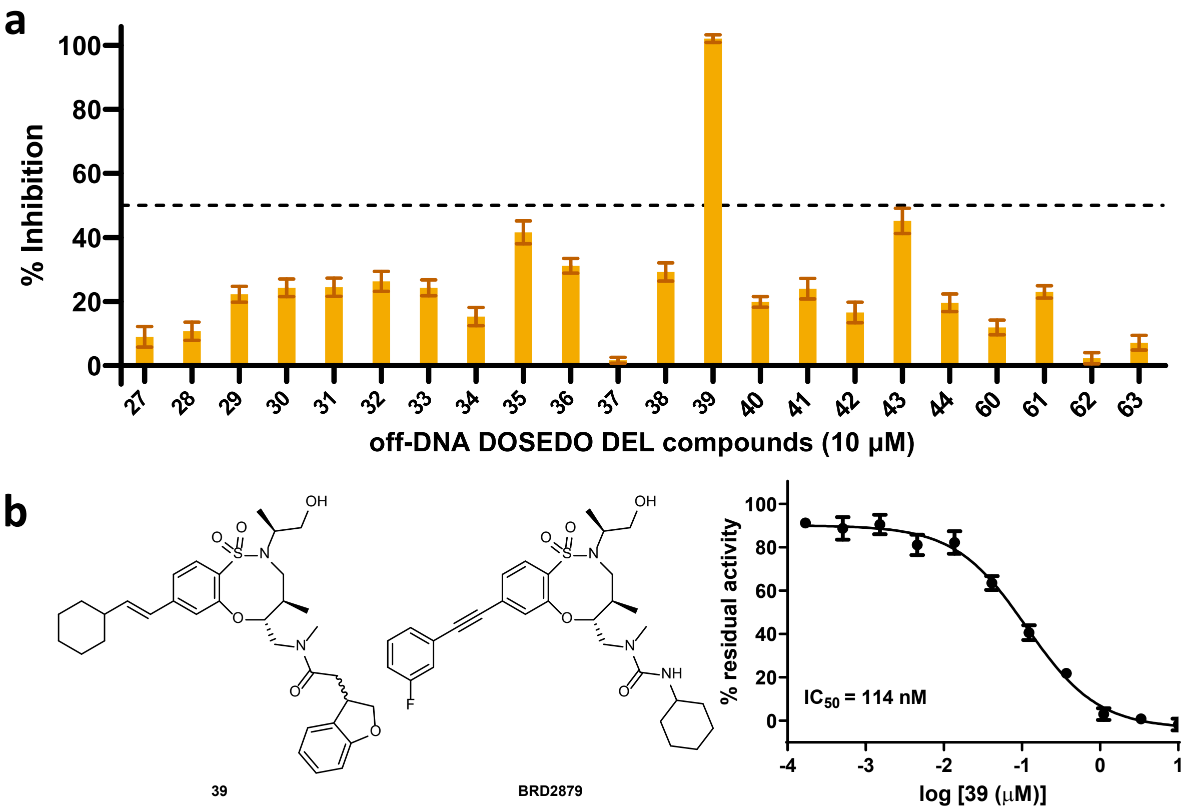

**Supplementary Fig. 20 | Summary of IDH1 R132H inhibition assays. a.** % inhibition of IDH1 R132H catalyzed conversion of 2OG to 2HG by off-DNA DOSEDO DEL compounds, as measured by absorbance assays. Compound **39** fully inhibits IDH1 R132H at 10 µM while all other compounds have an IC50 of >10 µM. All compounds have been tested for $IC_{50}$ with 3-fold serial dilution from 10 µM to confirm their weak (except 514) inhibitory activity at all concentrations below 10 µM. Data are mean ± SEM, n= 9 consisting of 3 technical replicates from 3 independent replicates; **b.** Structure of compound **39** and **BRD2879** alongside $IC_{50}$ of compound **39** against IDH1 R132H, as measured by absorbance assays. Data are mean ± SD, n= 3 technical replicates.

### 6. In silico – Supplementary Figures

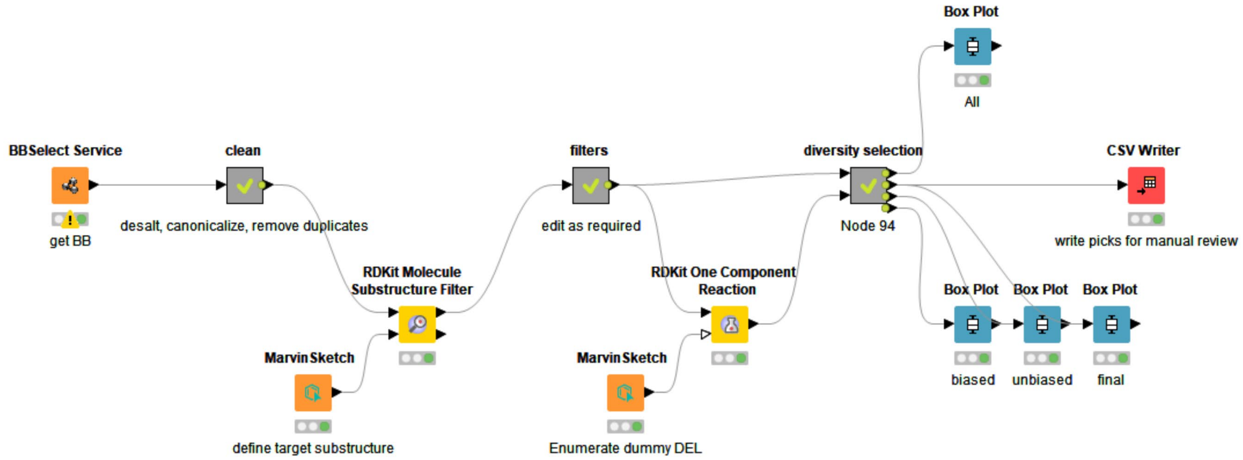

**Supplementary Fig. 21 | Overview of building block selection tool**. “BBSelect Service” is a tool internal to Novartis allowing for rapid searching of multiple curated building block catalogs, internally available materials, as well as commercial building block sources.

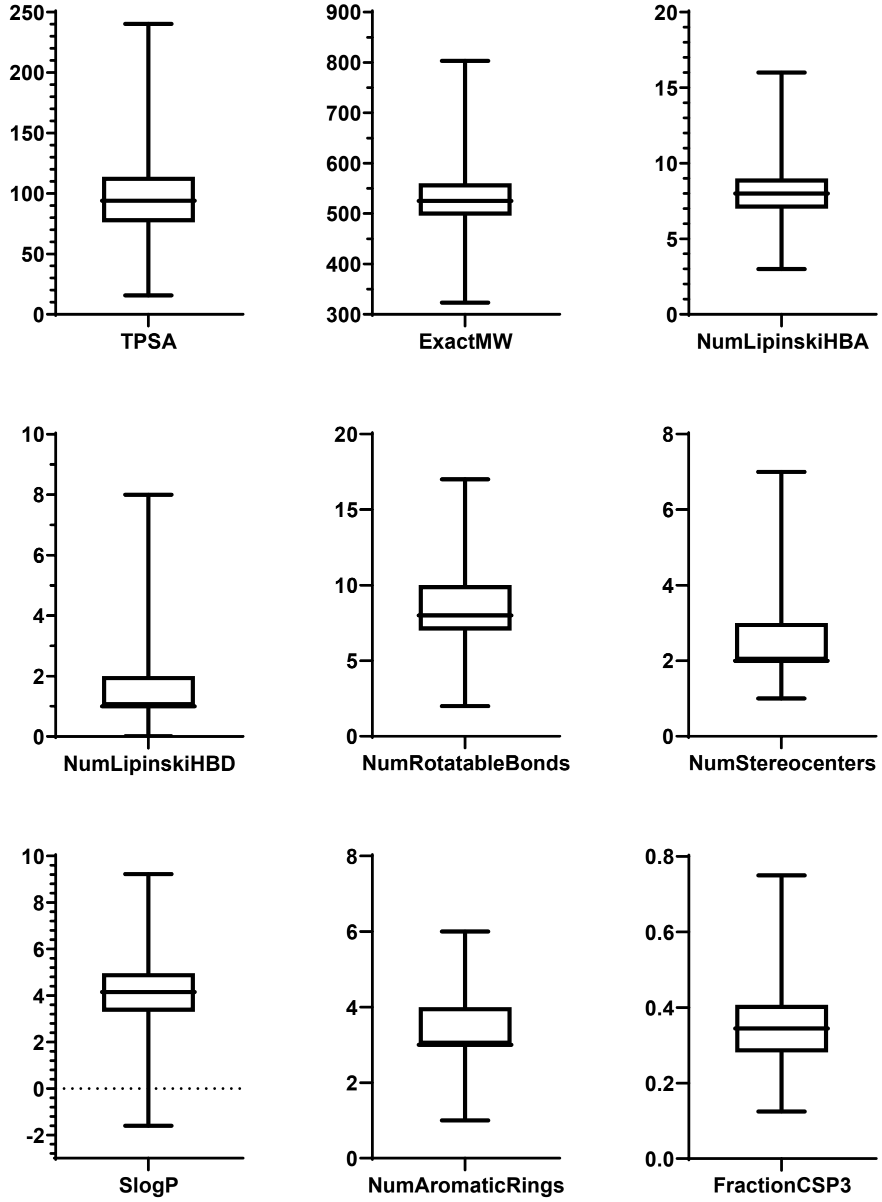

**Supplementary Fig. 22 | Box and whisker plots for selected calculated properties of the entire enumerated DOSEDO library**. Central bars show the median value, the box encapsulates the median 50% of compounds, and the whiskers show the upper and lower quartiles up to and including the maximum and minimum values for each property.

### 7. Small molecule NMR data

#### 7.1. β-arylated cyclic amino acids

(2*R*,3*S*)-1-(((9H-fluoren-9-yl)methoxy)carbonyl)-3-(3-iodophenyl)azetidine-2-carboxylic acid (**6a**)

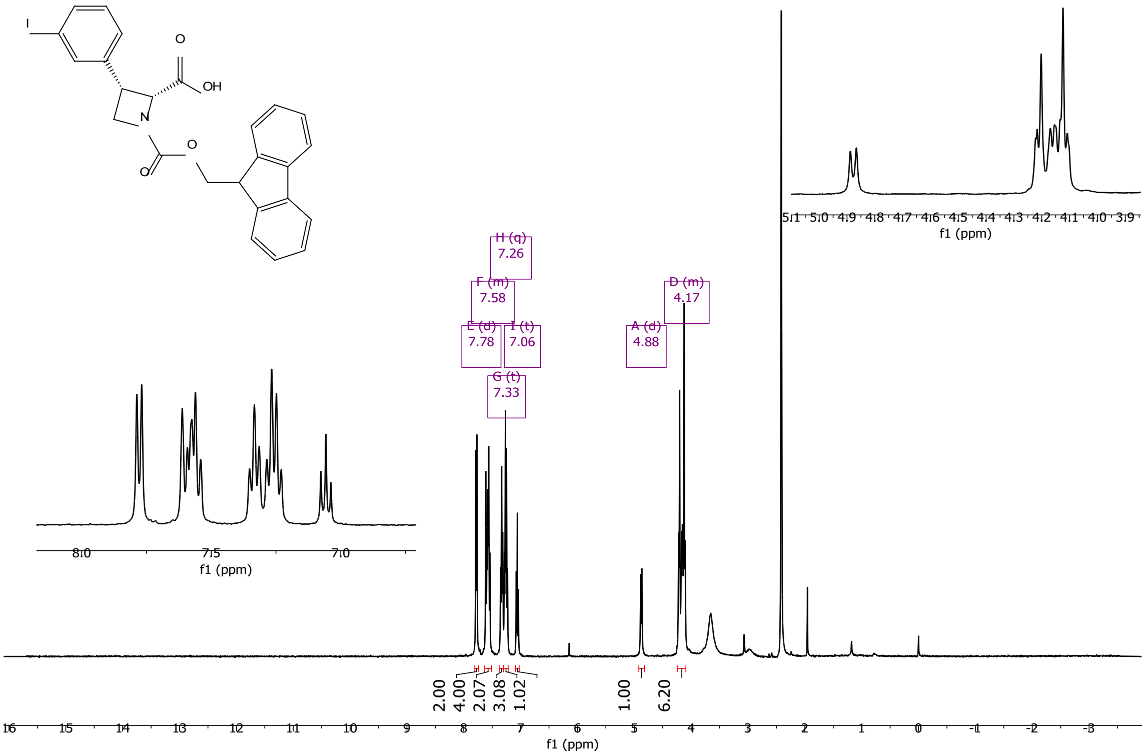

1, 2, 3 and 4 are spectra collected at 25, 40, 60 and 80°C, respectively:

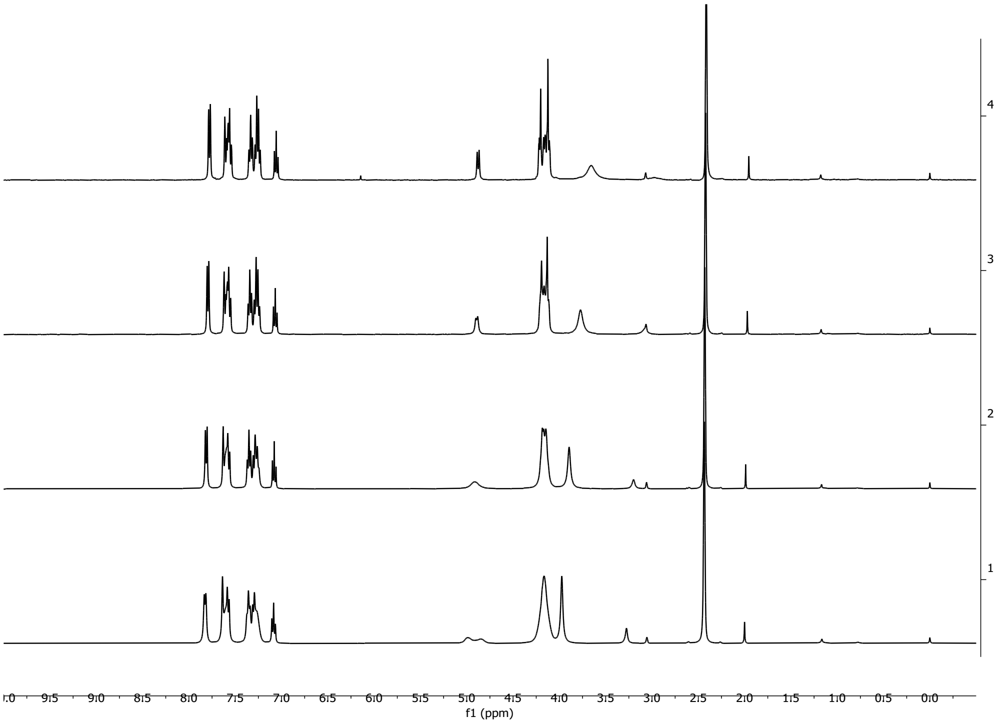

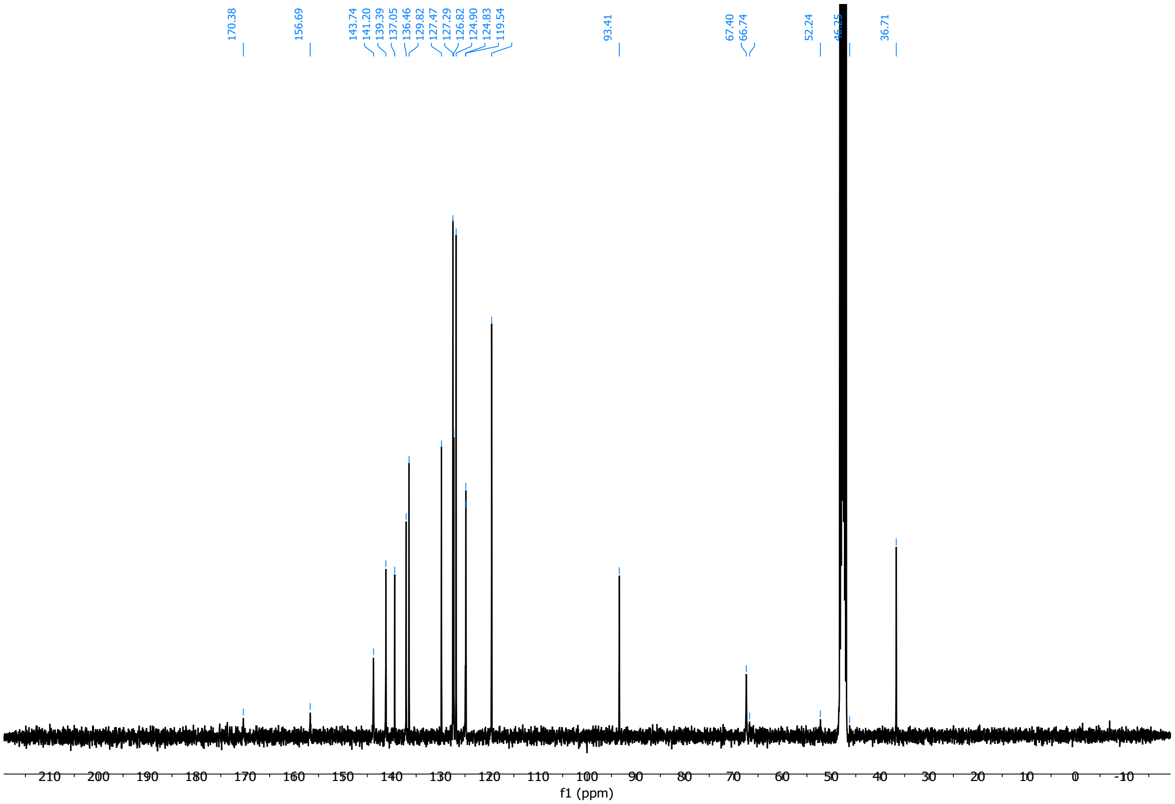

(2*S*,3*R*)-1-(((9H-fluoren-9-yl)methoxy)carbonyl)-3-(3-iodophenyl)azetidine-2-carboxylic acid (**6b**)

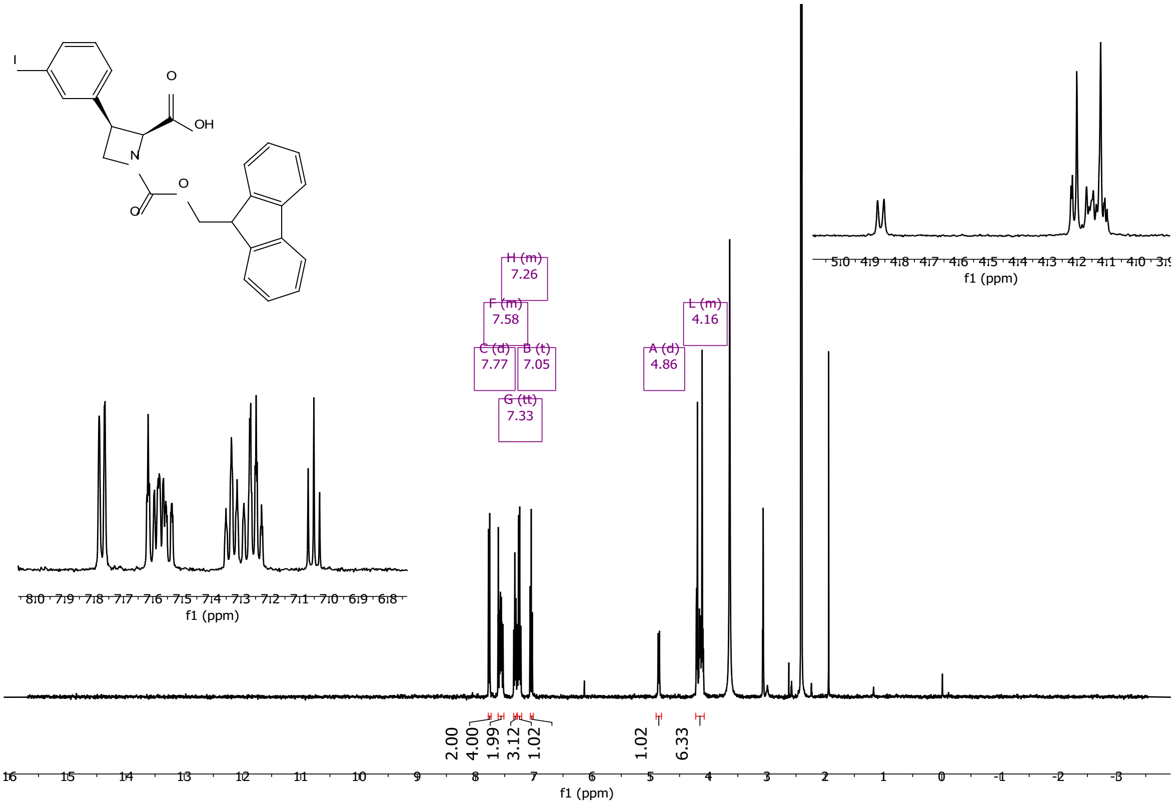

1, 2, 3 and 4 are spectra collected at 25, 40, 60 and 80°C, respectively:

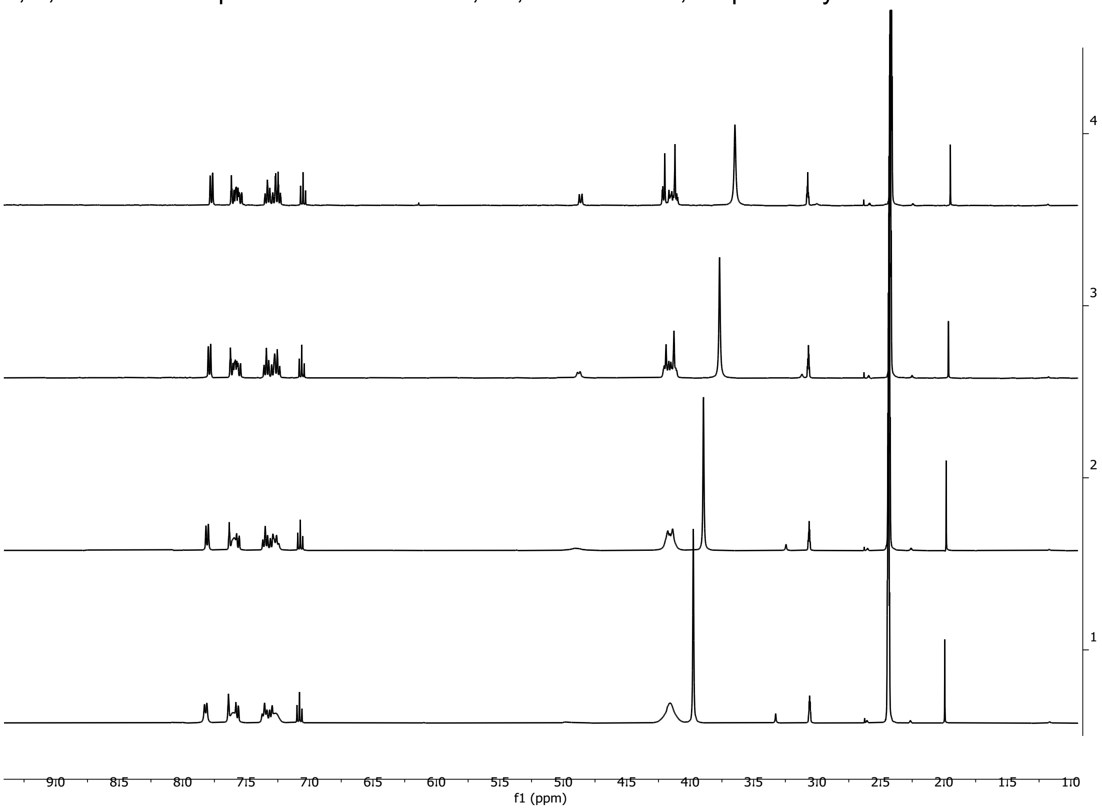

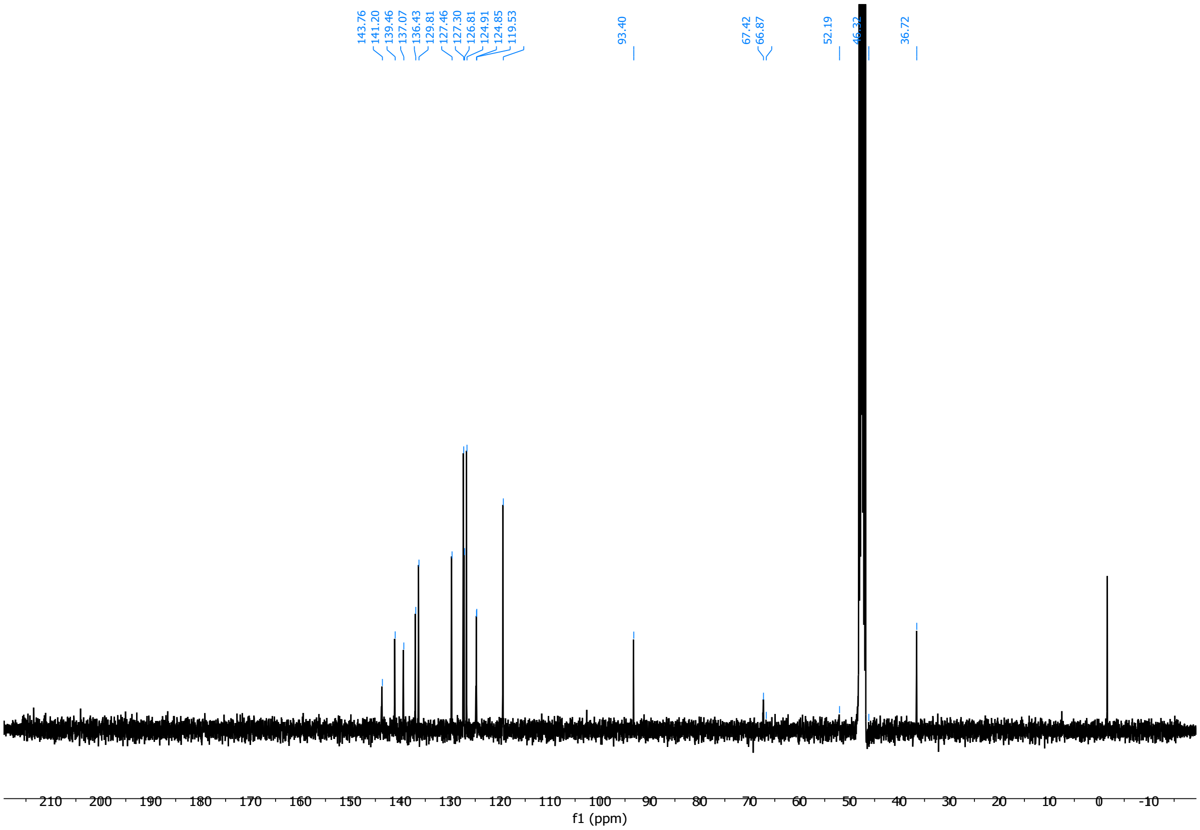

(2*R*,3*R*)-1-(((9H-fluoren-9-yl)methoxy)carbonyl)-3-(3-iodophenyl)pyrrolidine-2-carboxylic acid (**6c**)

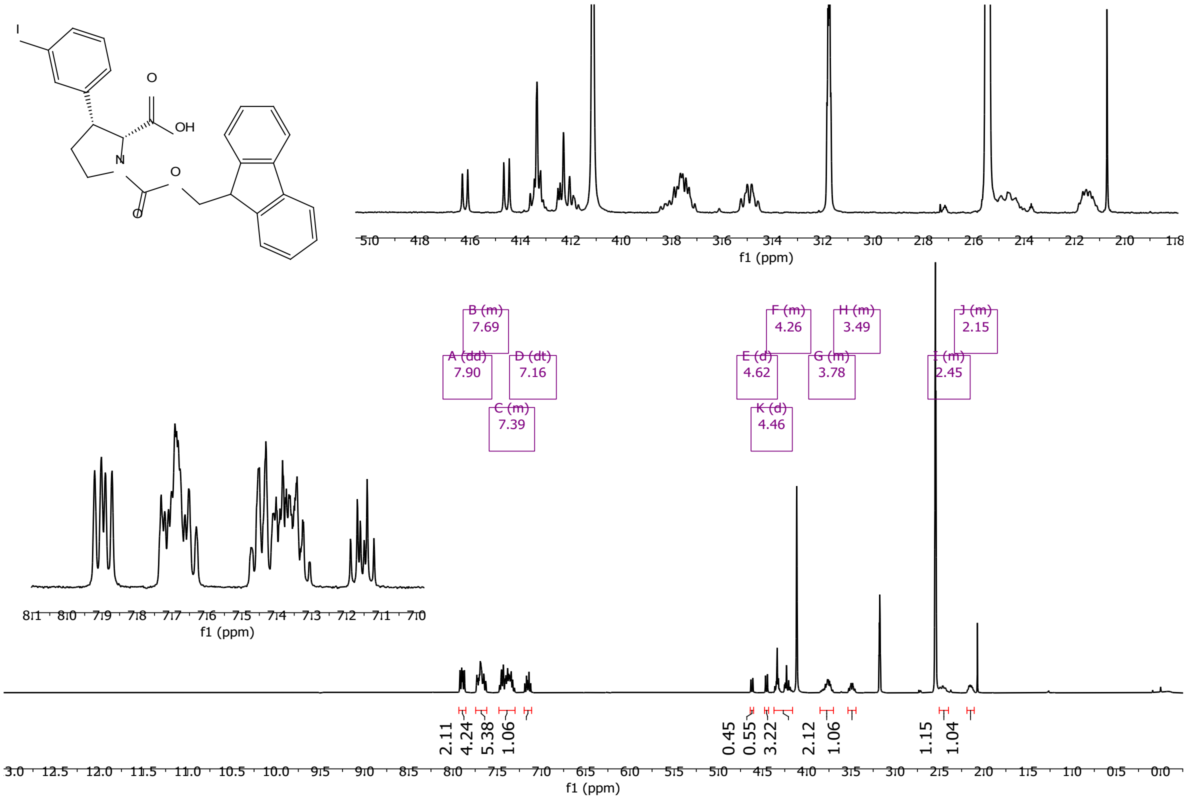

1, 2, 3 and 4 are spectra collected at 25, 40, 60 and 80°C, respectively:

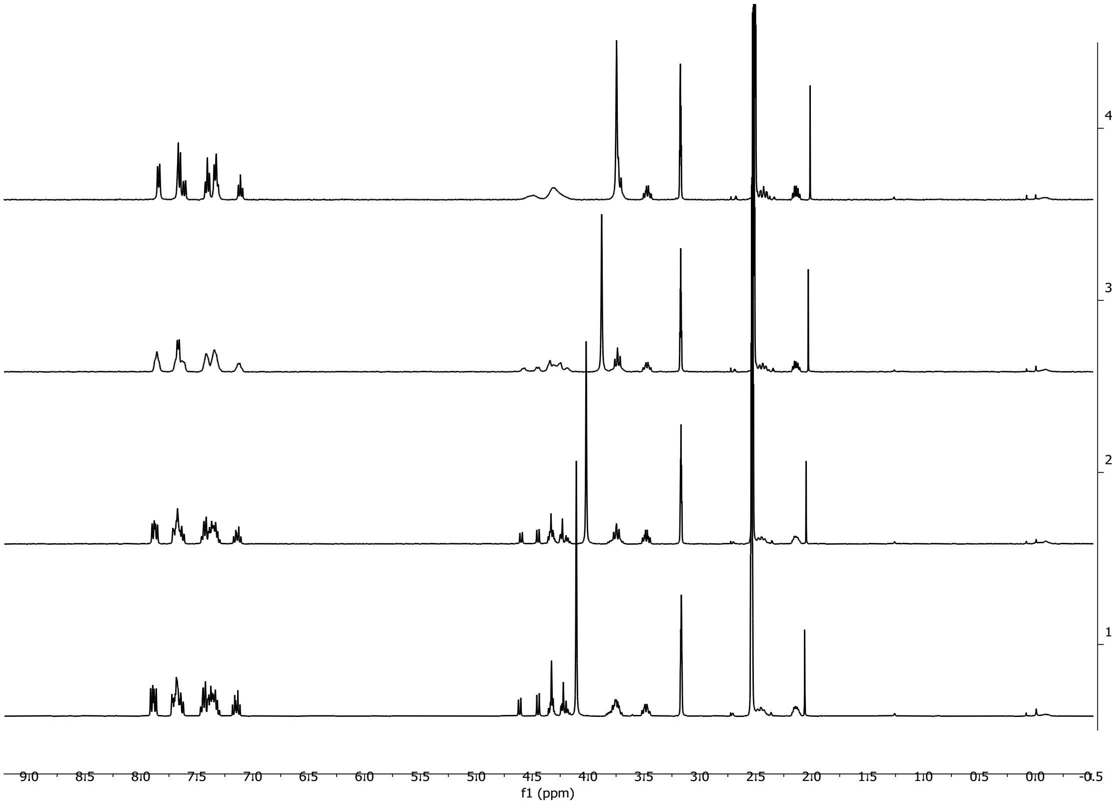

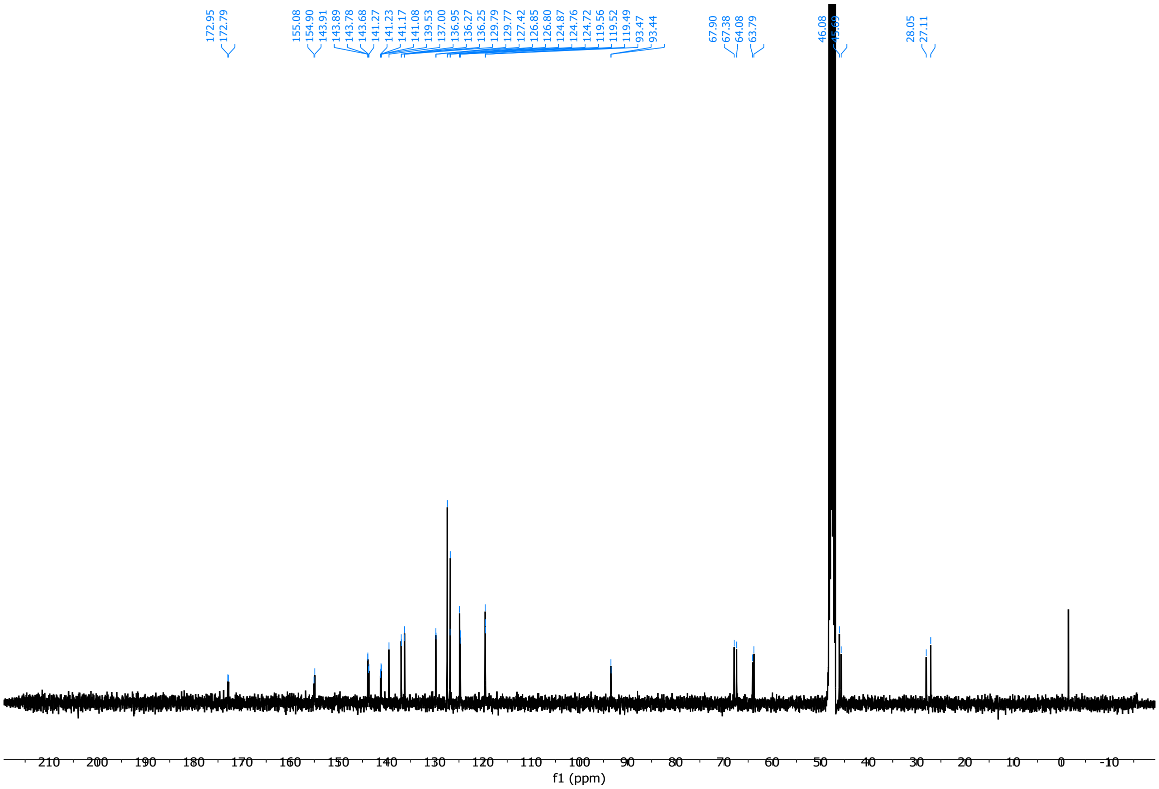

(2*S*,3*S*)-1-(((9H-fluoren-9-yl)methoxy)carbonyl)-3-(3-iodophenyl)pyrrolidine-2-carboxylic acid (**6d**)

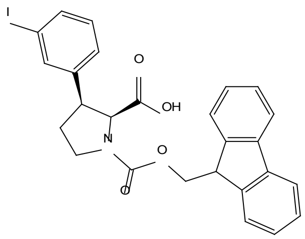

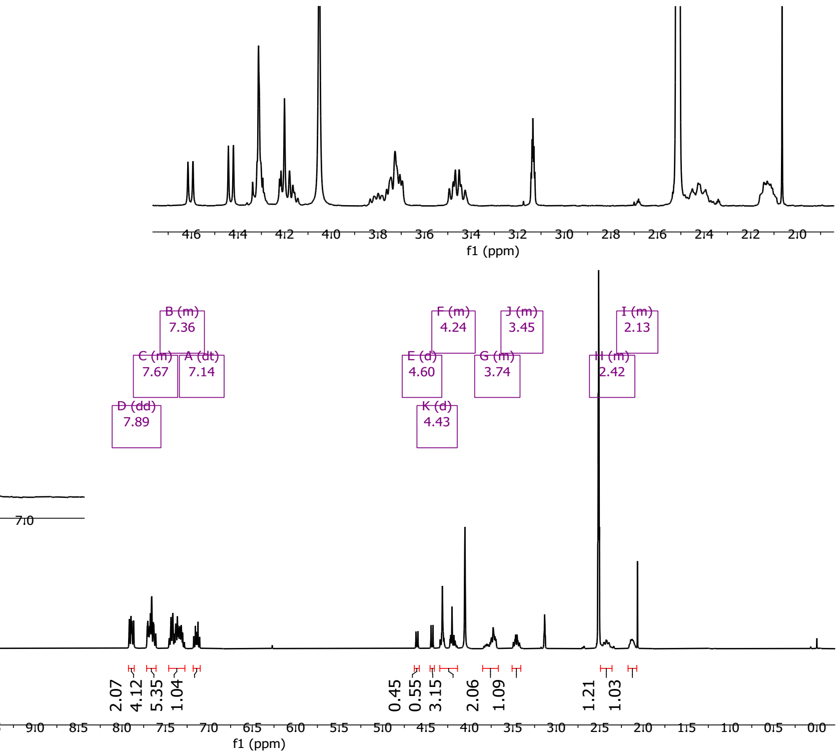

1, 2, 3 and 4 are spectra collected at 25, 40, 60 and 80°C, respectively:

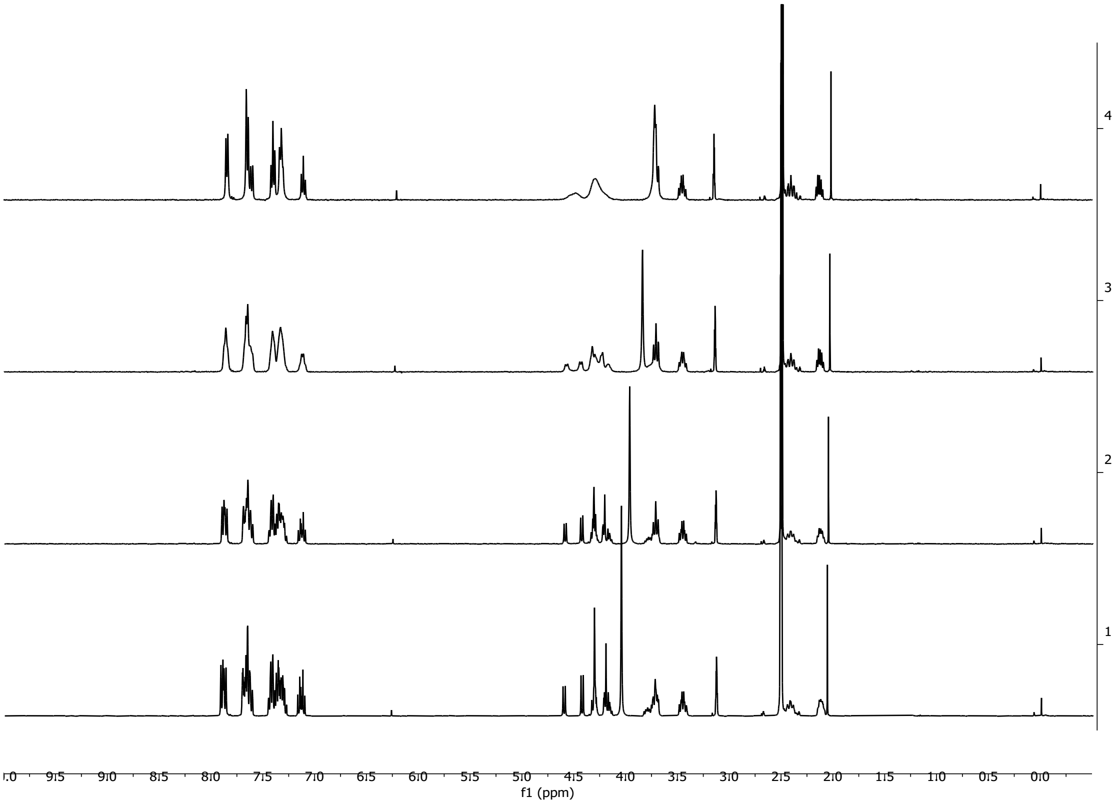

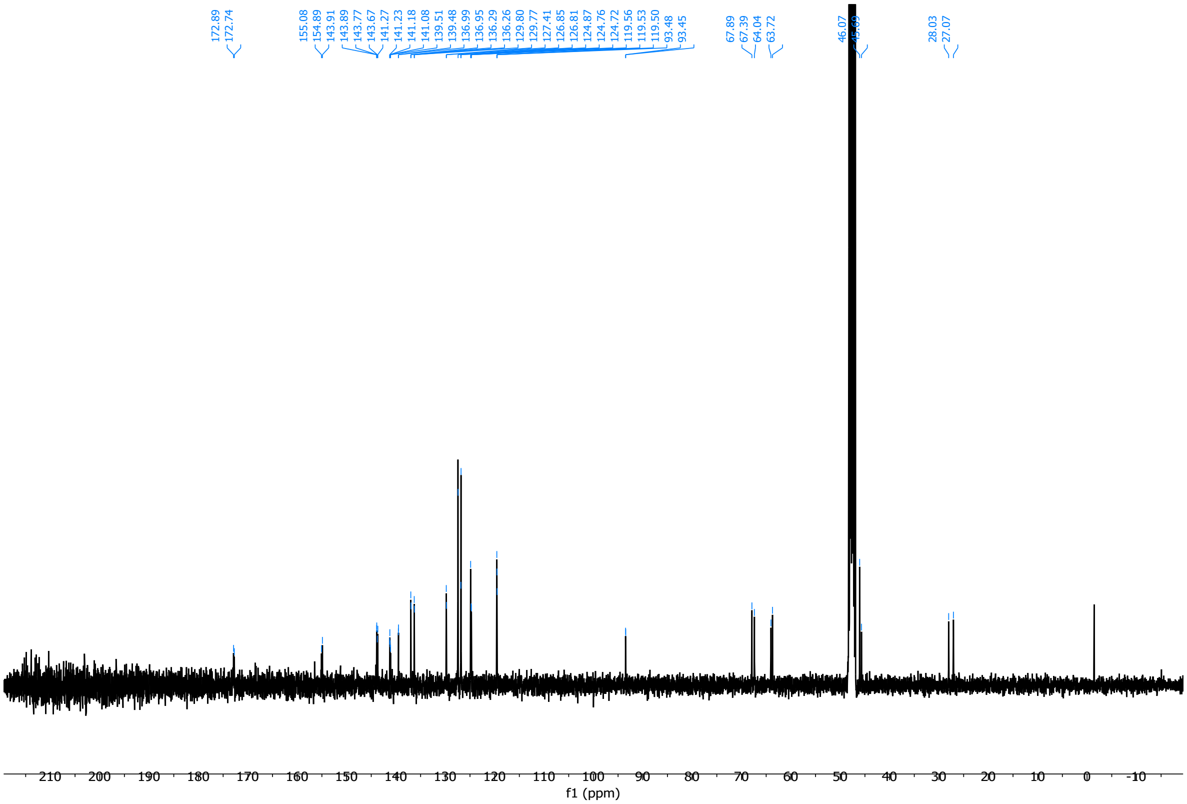

(2*R*,3*R*)-1-(((9H-fluoren-9-yl)methoxy)carbonyl)-3-(3-iodophenyl)piperidine-2-carboxylic acid (**6e**)

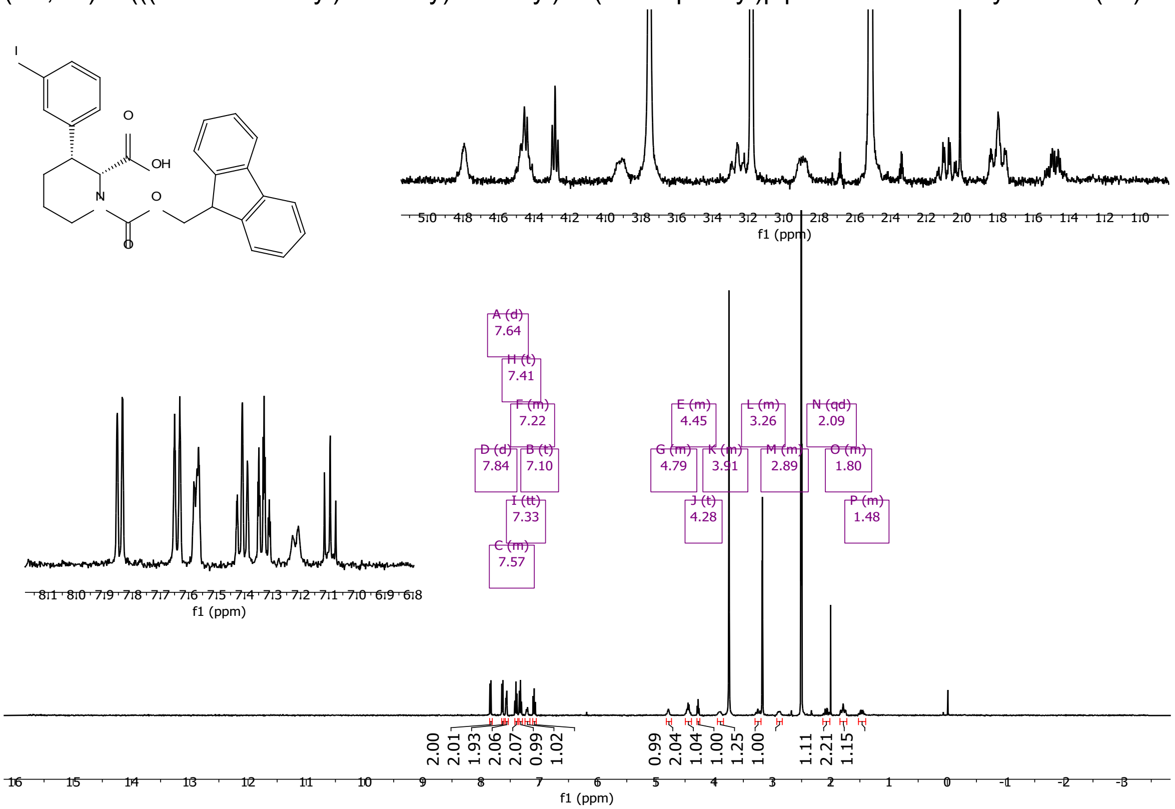

1, 2, 3 and 4 are spectra collected at 25, 40, 60 and 80°C, respectively:

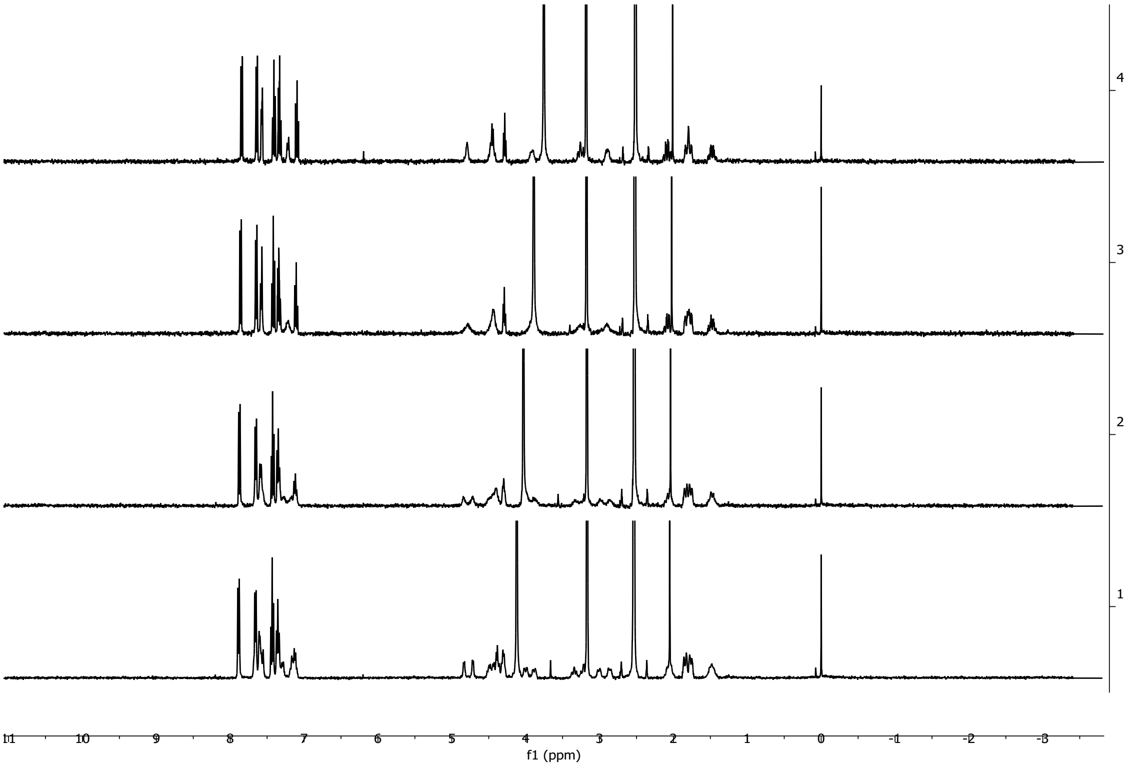

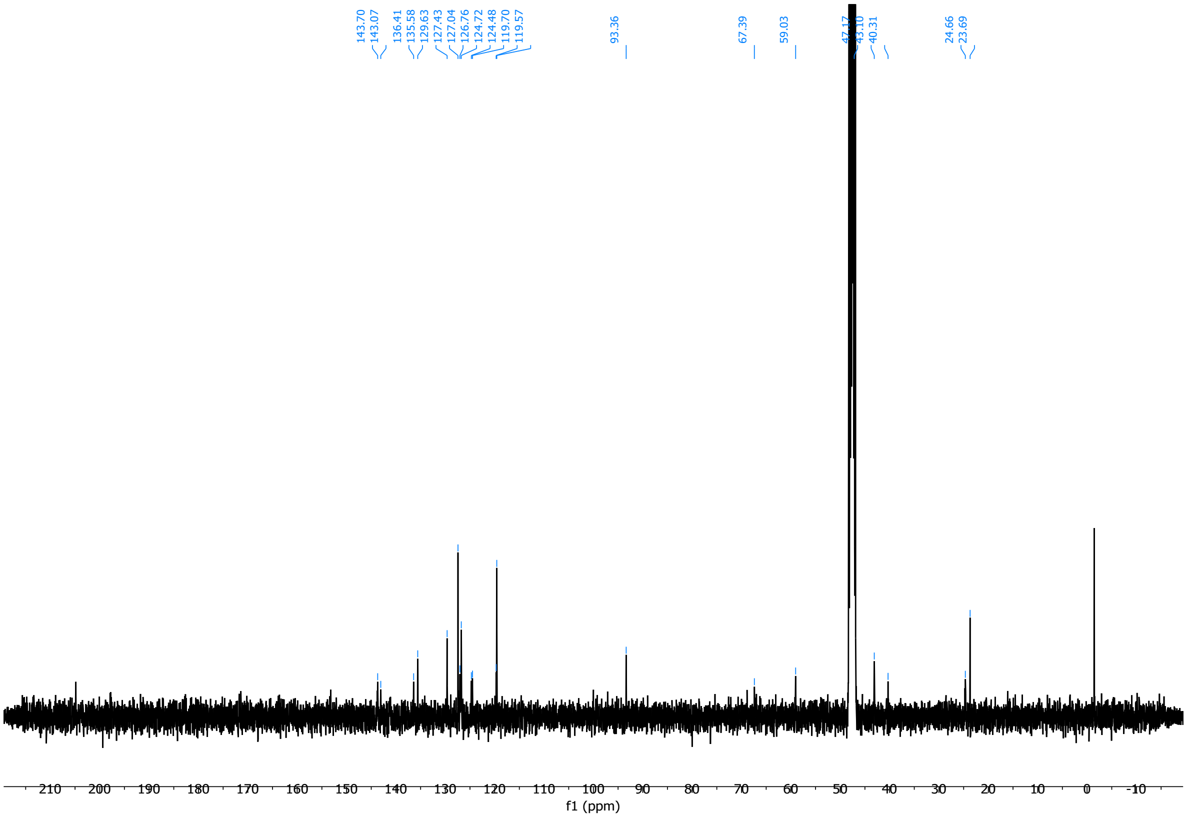

Overlay of HSQC and HMBC (rt in CD$_3$OD):

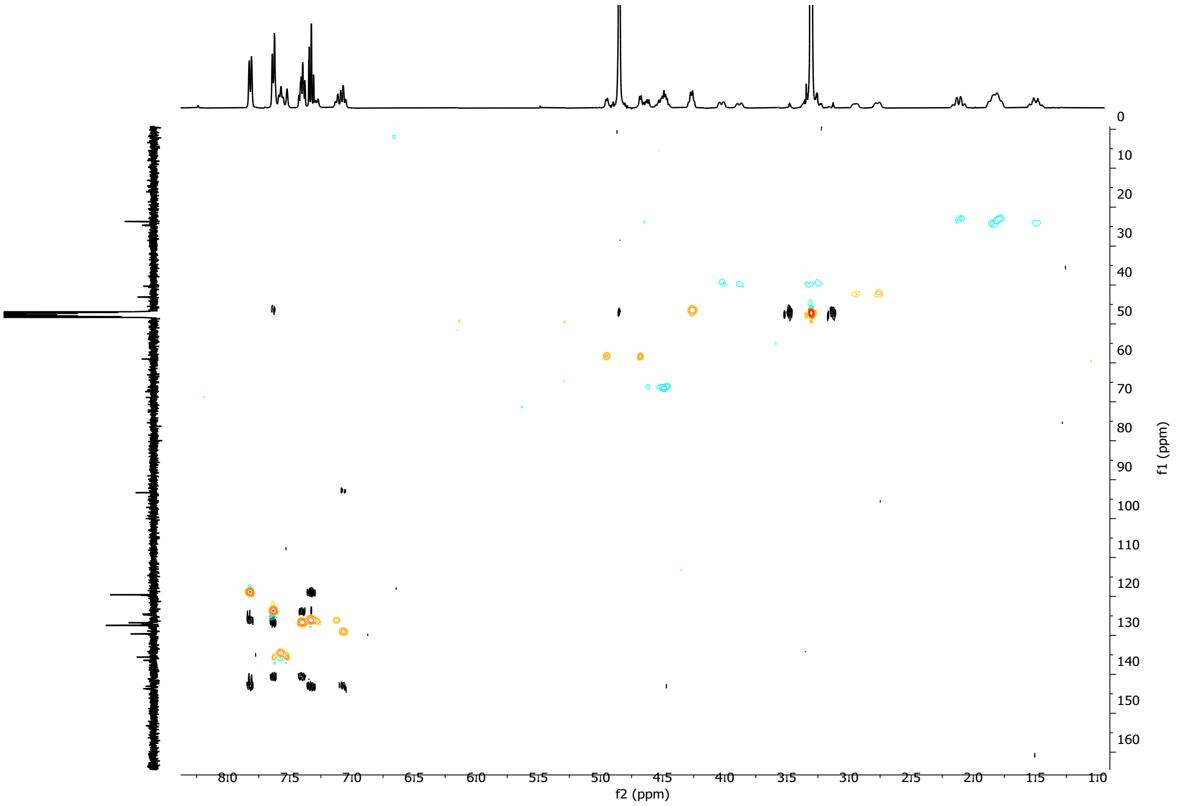

(2*S*,3*S*)-1-(((9H-fluoren-9-yl)methoxy)carbonyl)-3-(3-iodophenyl)piperidine-2-carboxylic acid (**6f**)

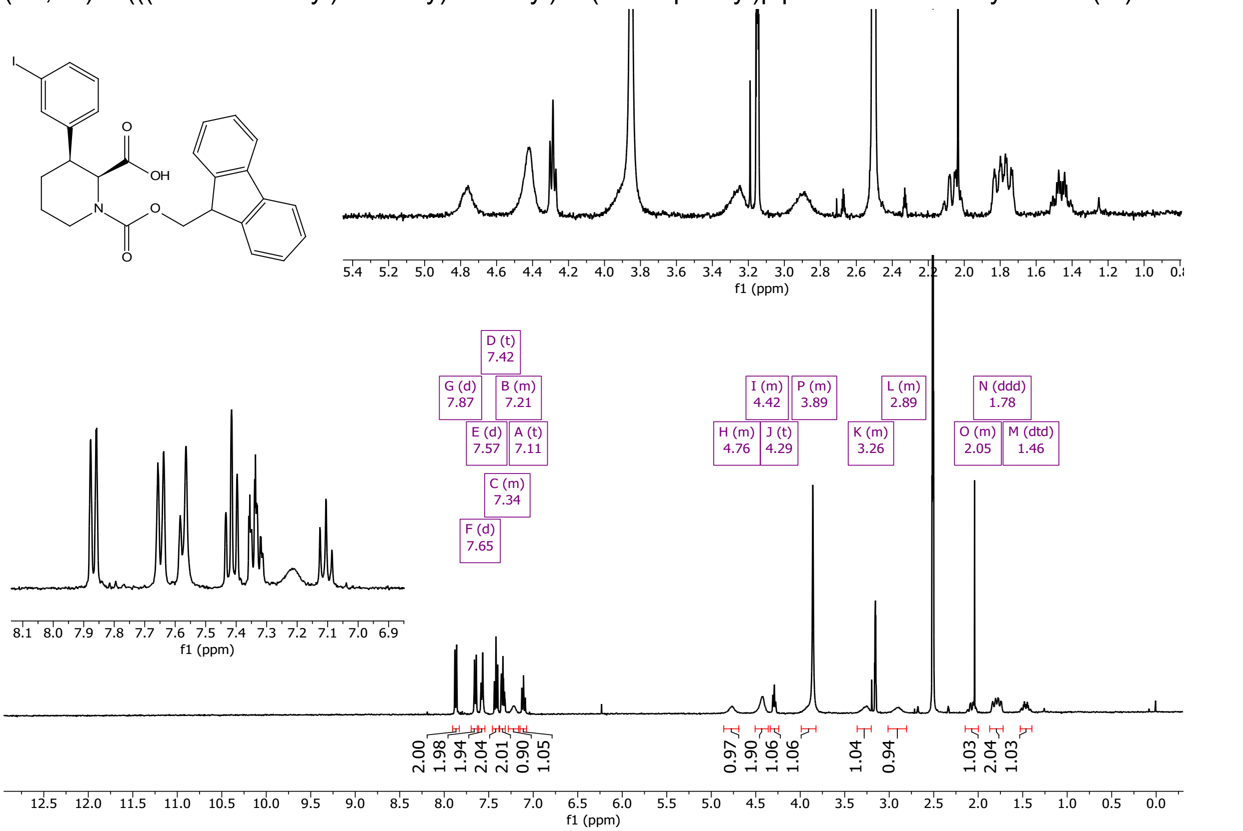

1, 2, 3, 4 and 5 are spectra collected at 25, 40, 60, 80, and then again at 25°C in sequence, respectively:

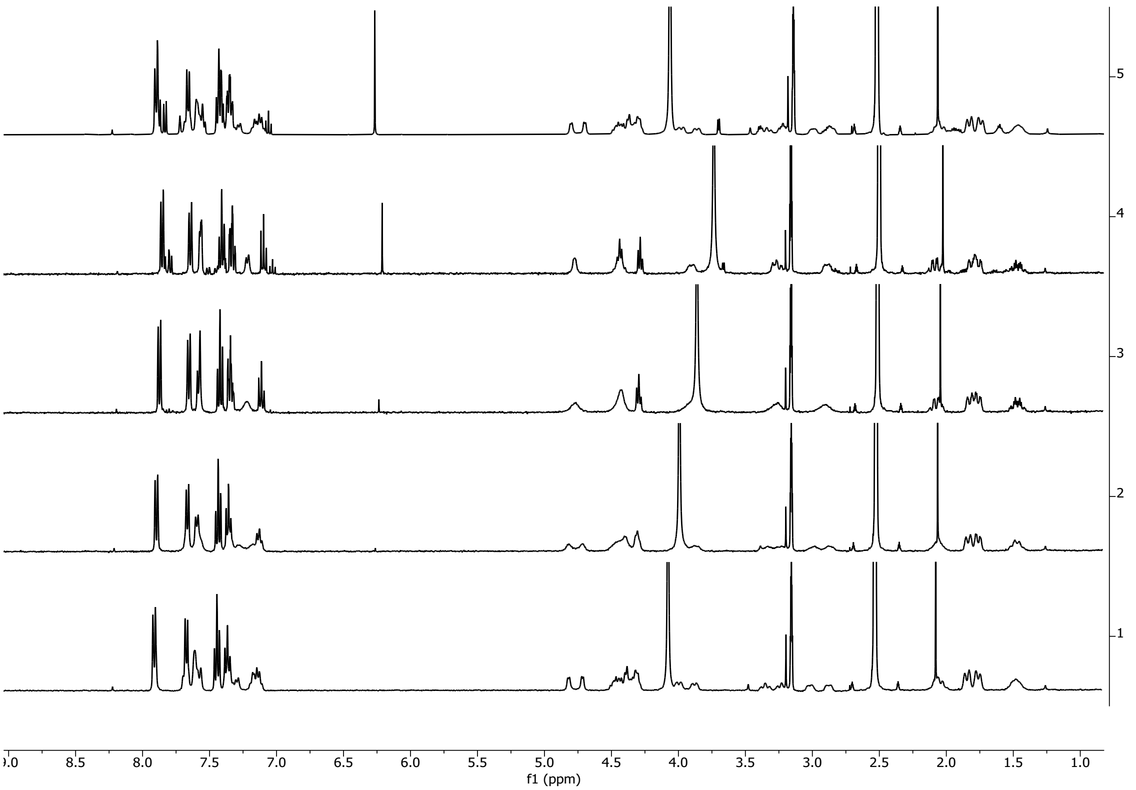

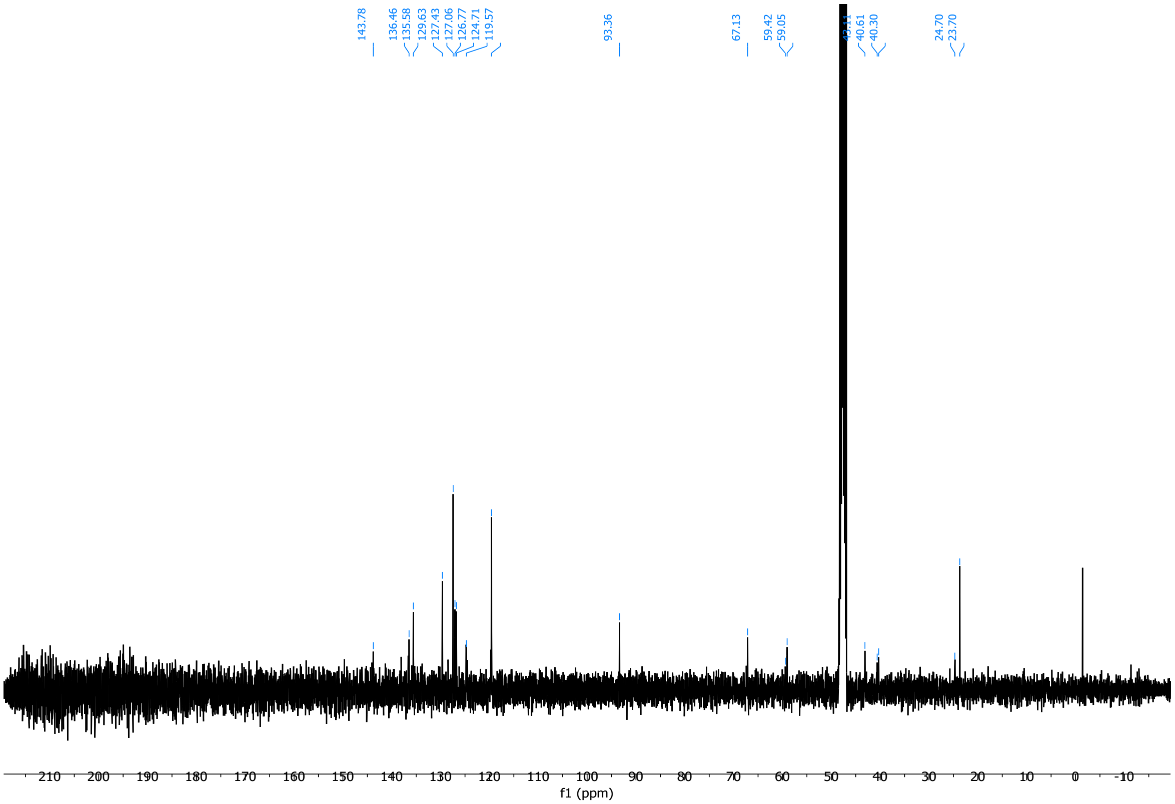

Overlay of HSQC and HMBC (rt in $CD_3OD$):

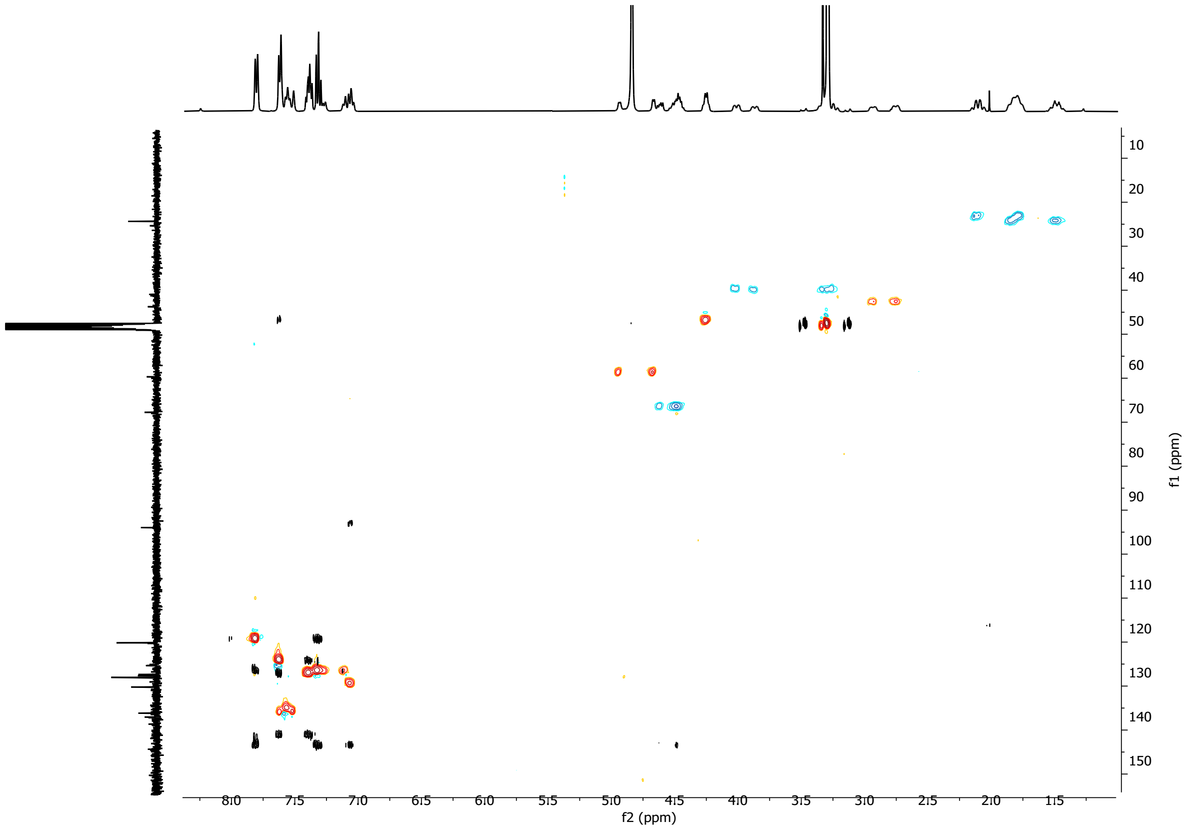

(2*S*,3*S*)-1-(((9H-fluoren-9-yl)methoxy)carbonyl)-3-(3-iodophenyl)azetidine-2-carboxylic acid (**7a**)

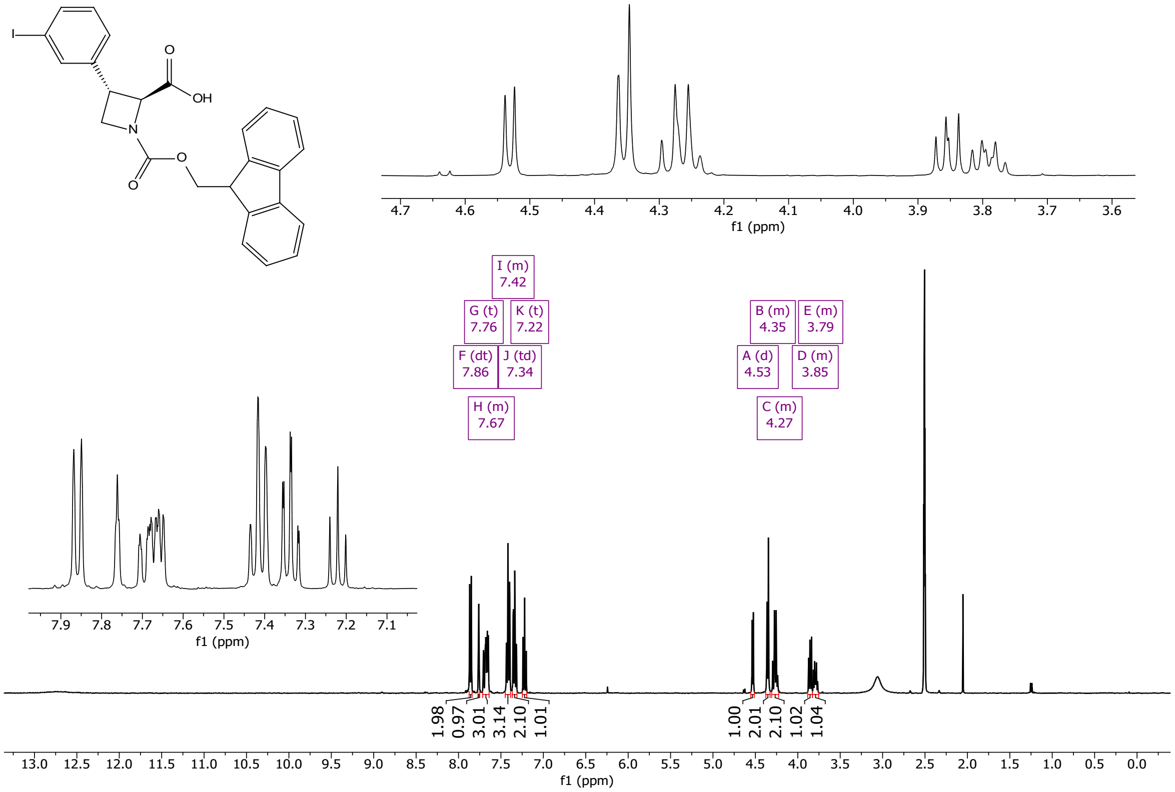

1, 2, 3 and 4 are spectra collected at 25, 40, 60 and 80°C, respectively:

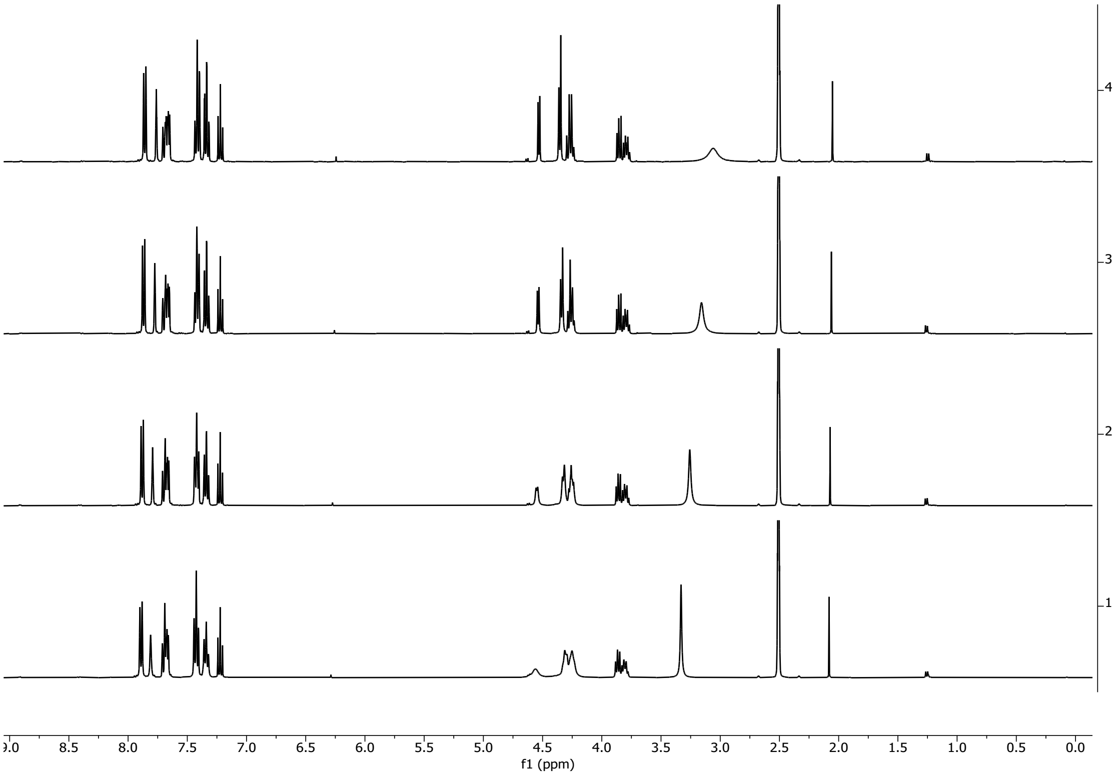

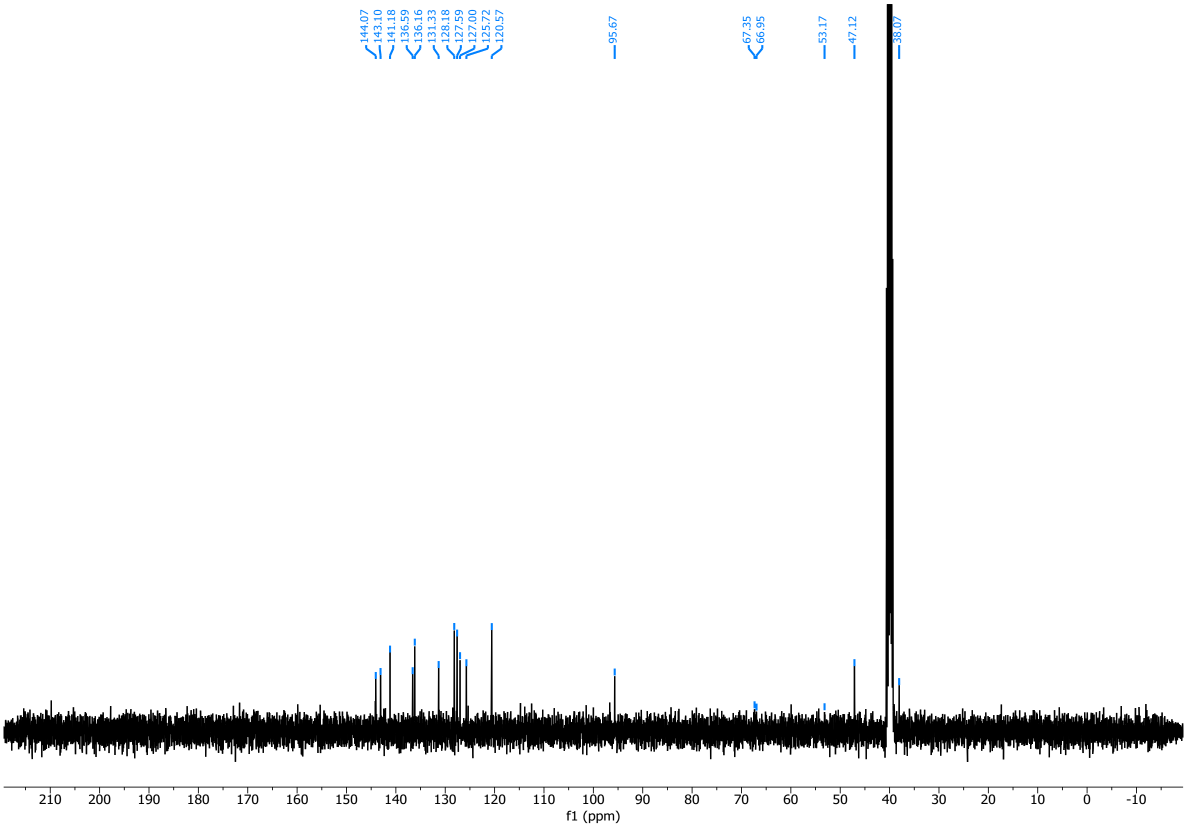

Overlay of HSQC and HMBC (rt in DMSO):

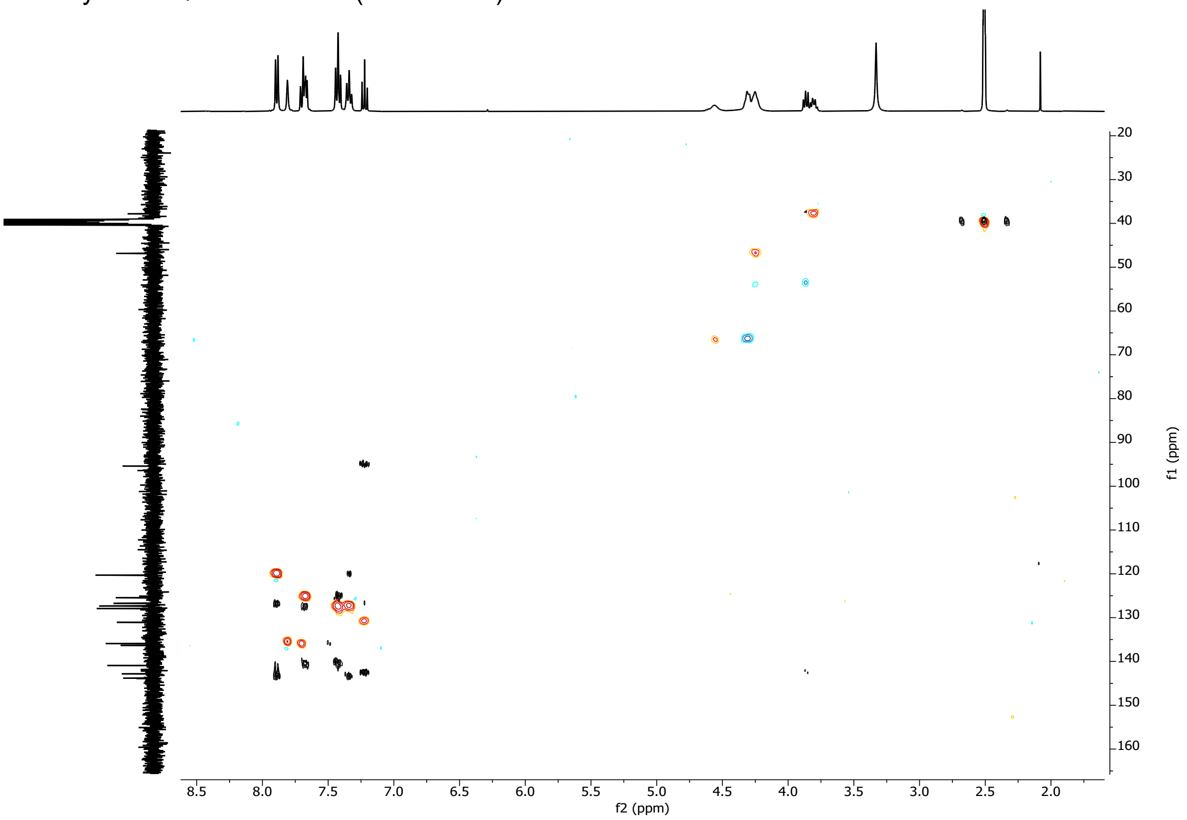

(2*R*,3*R*)-1-(((9H-fluoren-9-yl)methoxy)carbonyl)-3-(3-iodophenyl)azetidine-2-carboxylic acid (**7b**)

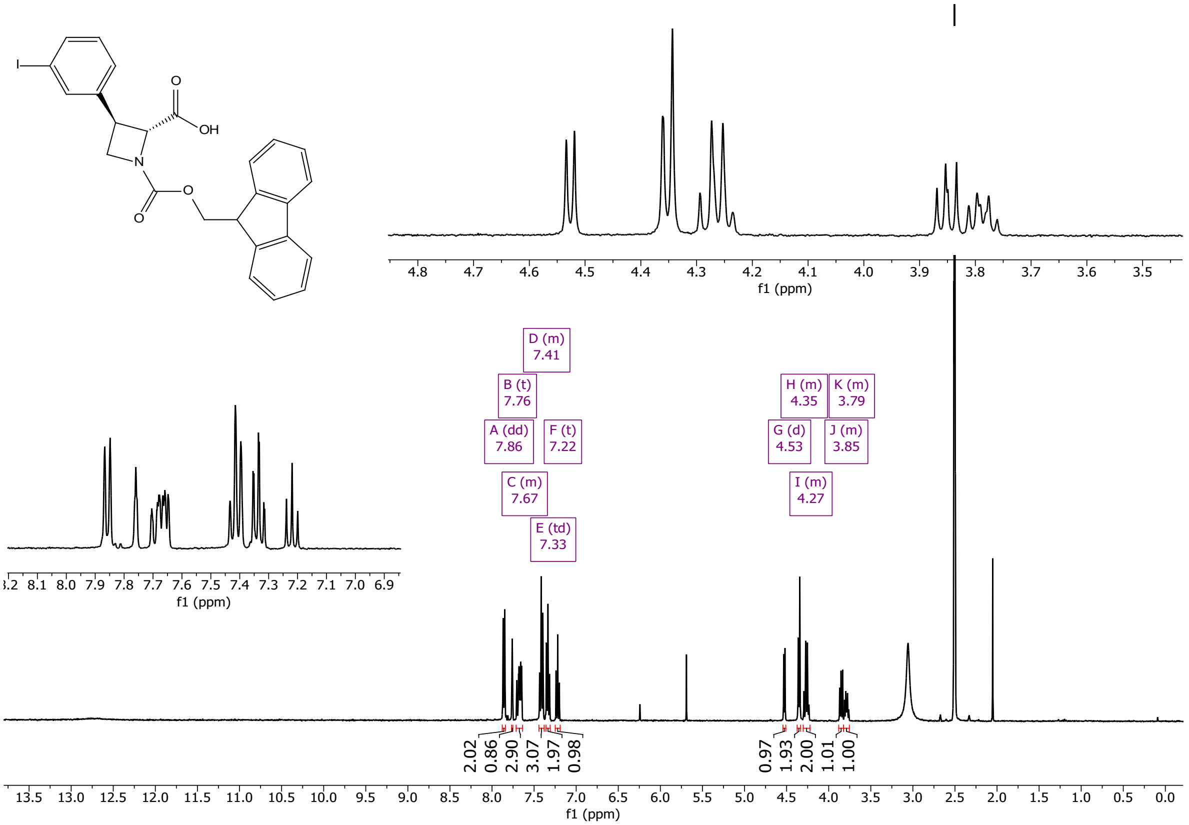

1, 2, 3 and 4 are spectra collected at 25, 40, 60 and 80°C, respectively:

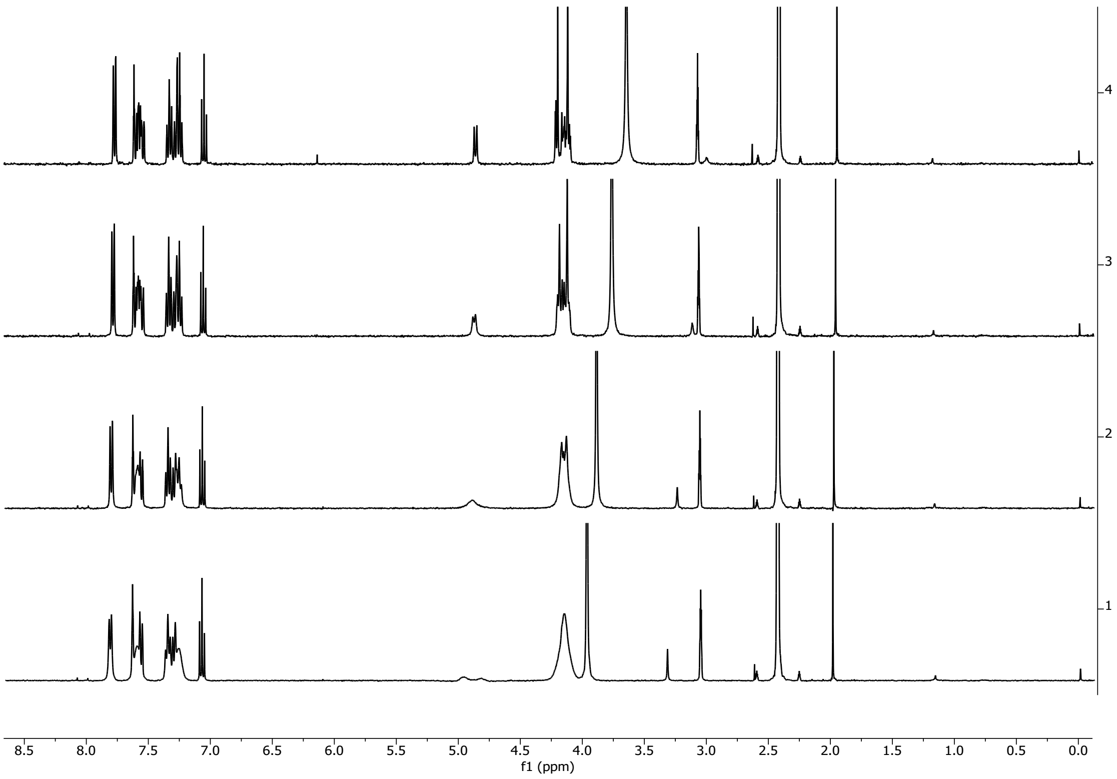

Overlay of HSQC and HMBC (rt in DMSO):

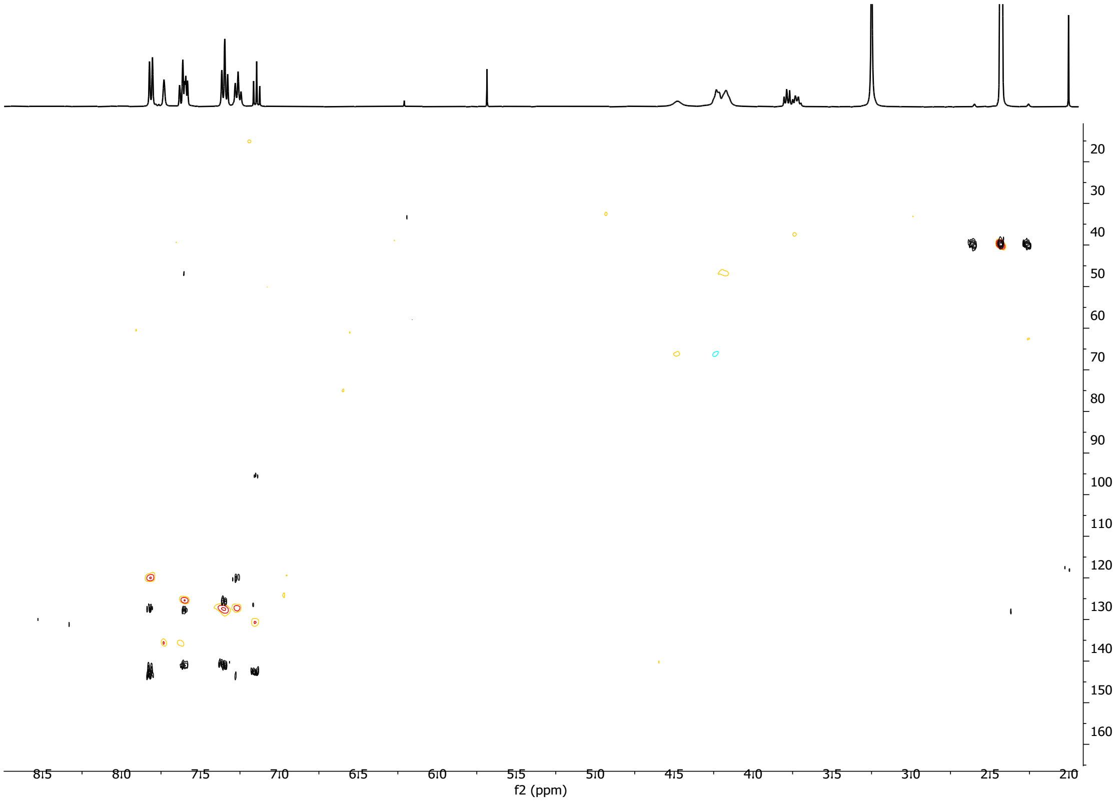

(2*S*,3*R*)-1-(((9H-fluoren-9-yl)methoxy)carbonyl)-3-(3-iodophenyl)pyrrolidine-2-carboxylic acid (**7c**)

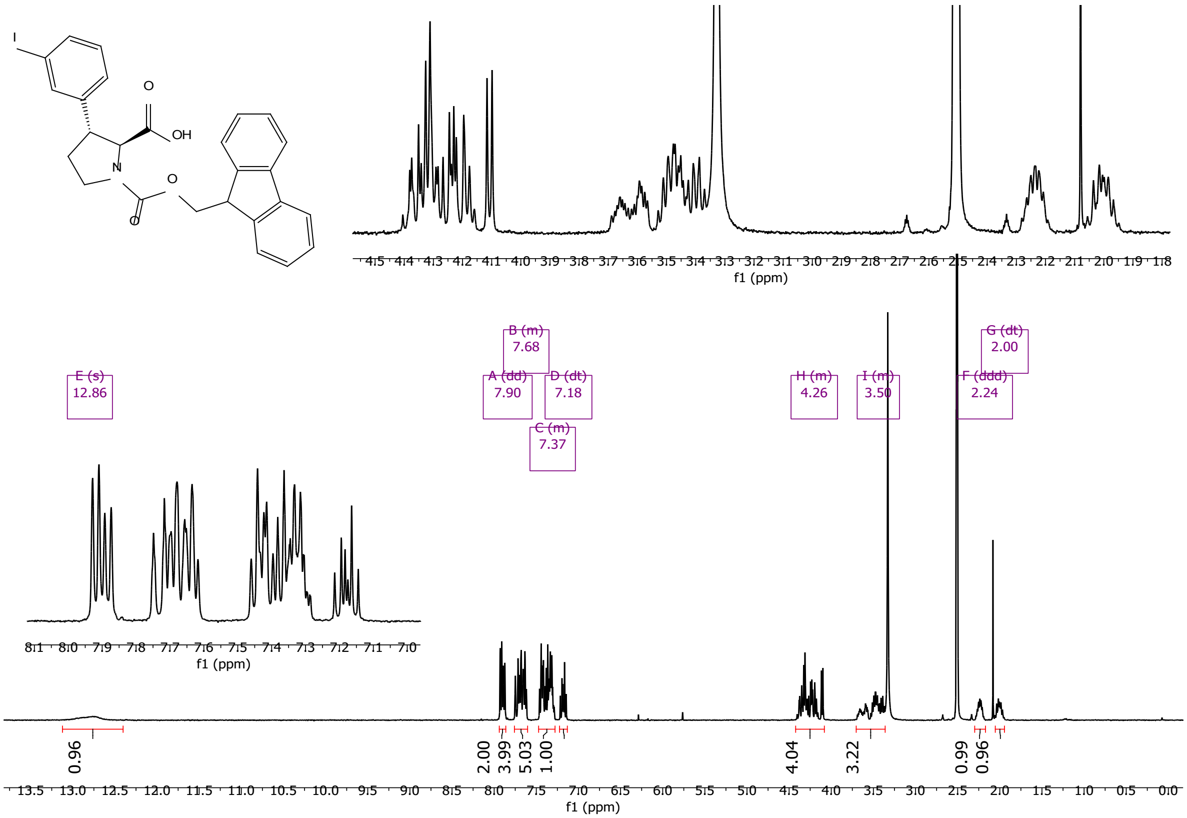

1, 2, 3 and 4 are spectra collected at 25, 40, 60 and 80°C, respectively:

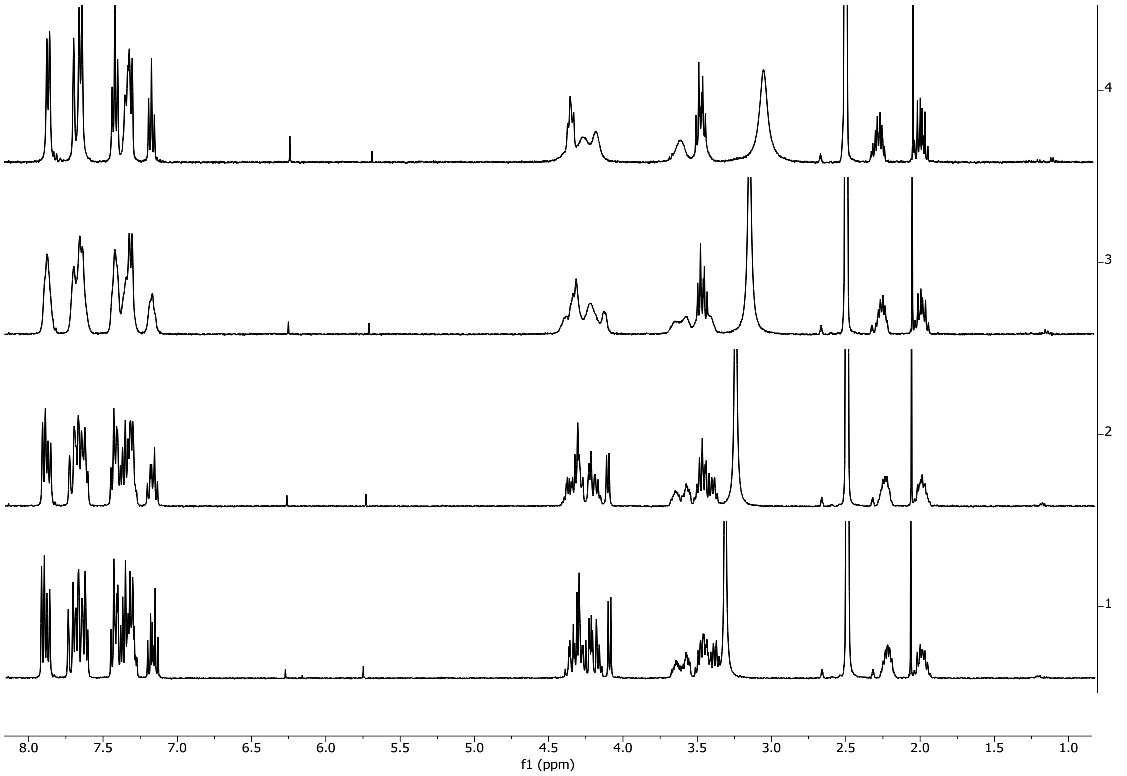

Overlay of HSQC and HMBC (rt in DMSO):

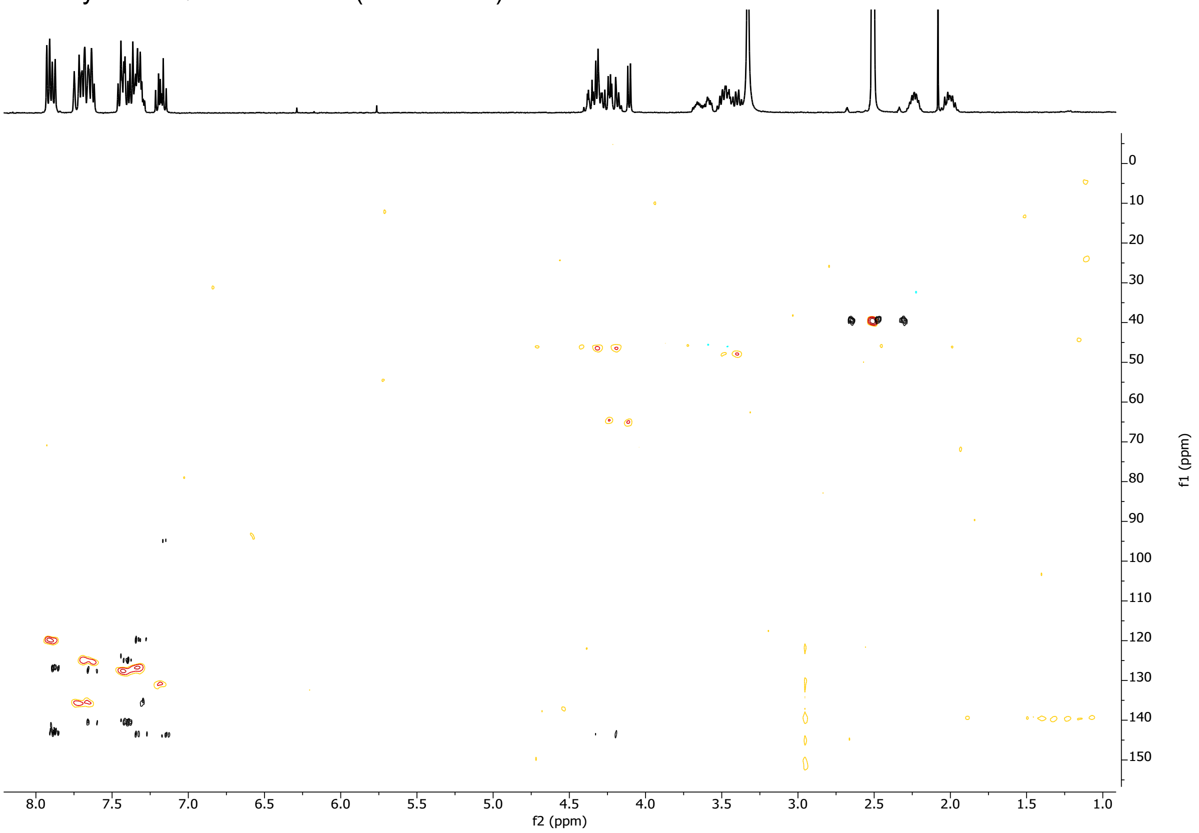

(2*R*,3*S*)-1-(((9H-fluoren-9-yl)methoxy)carbonyl)-3-(3-iodophenyl)pyrrolidine-2-carboxylic acid (**7d**)

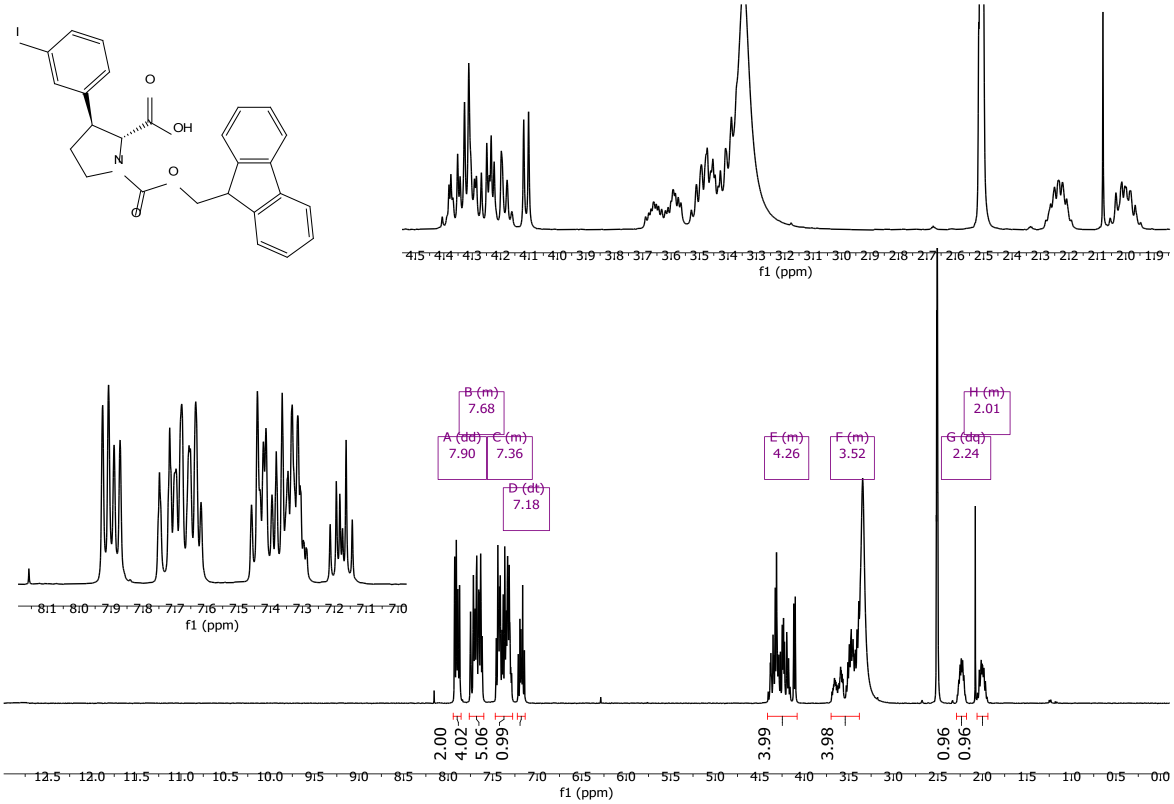

1, 2, 3 and 4 are spectra collected at 25, 40, 60 and 80°C, respectively:

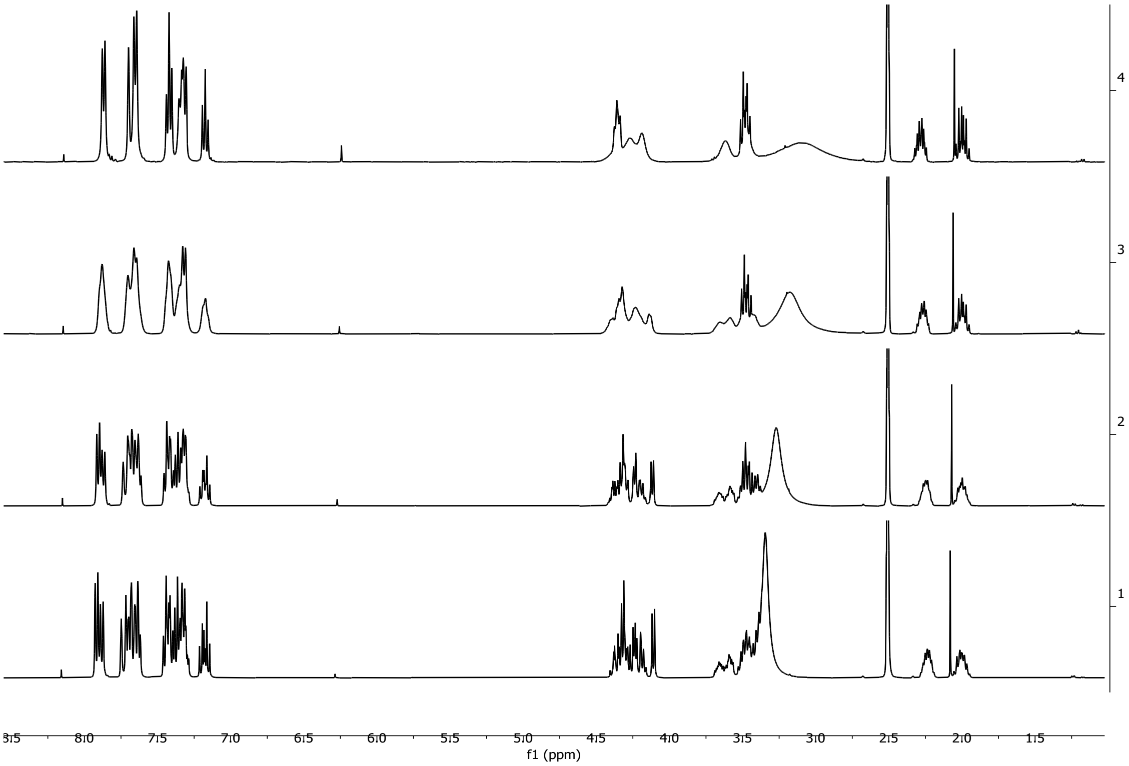

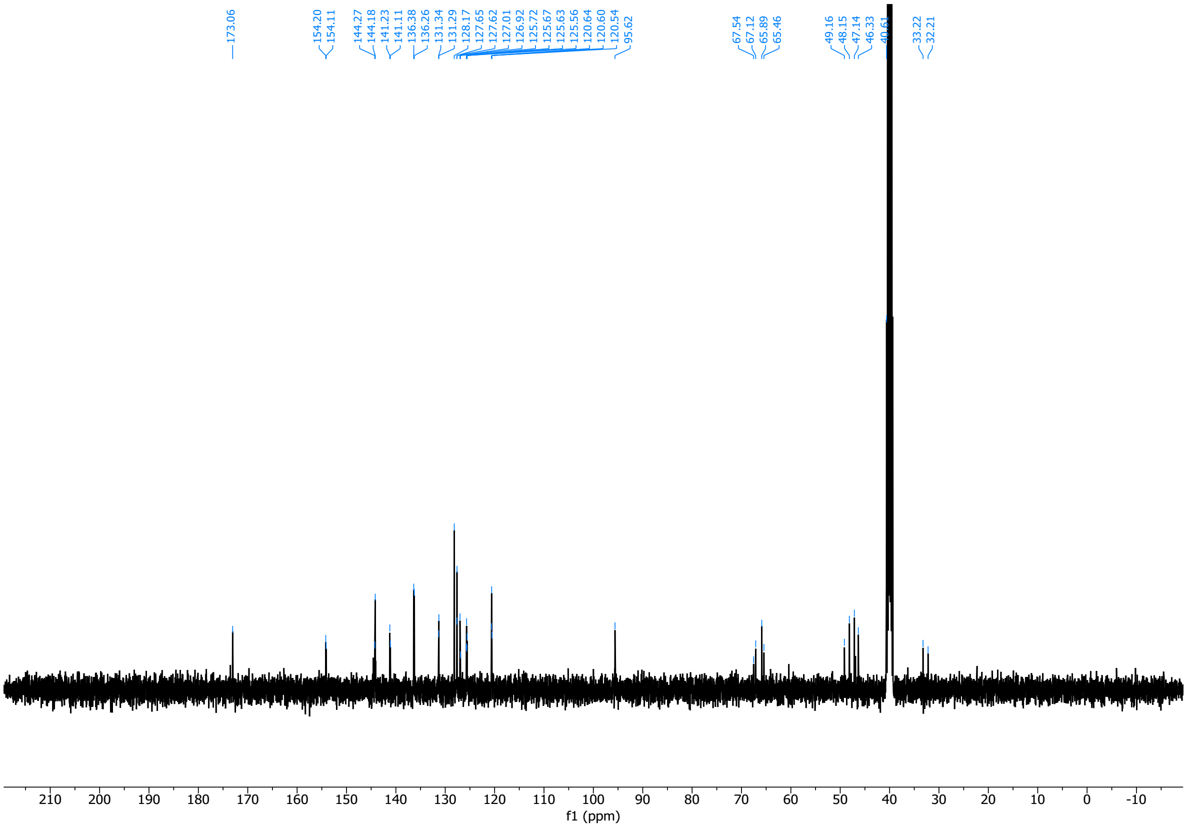

Overlay of HSQC and HMBC in DMSO:

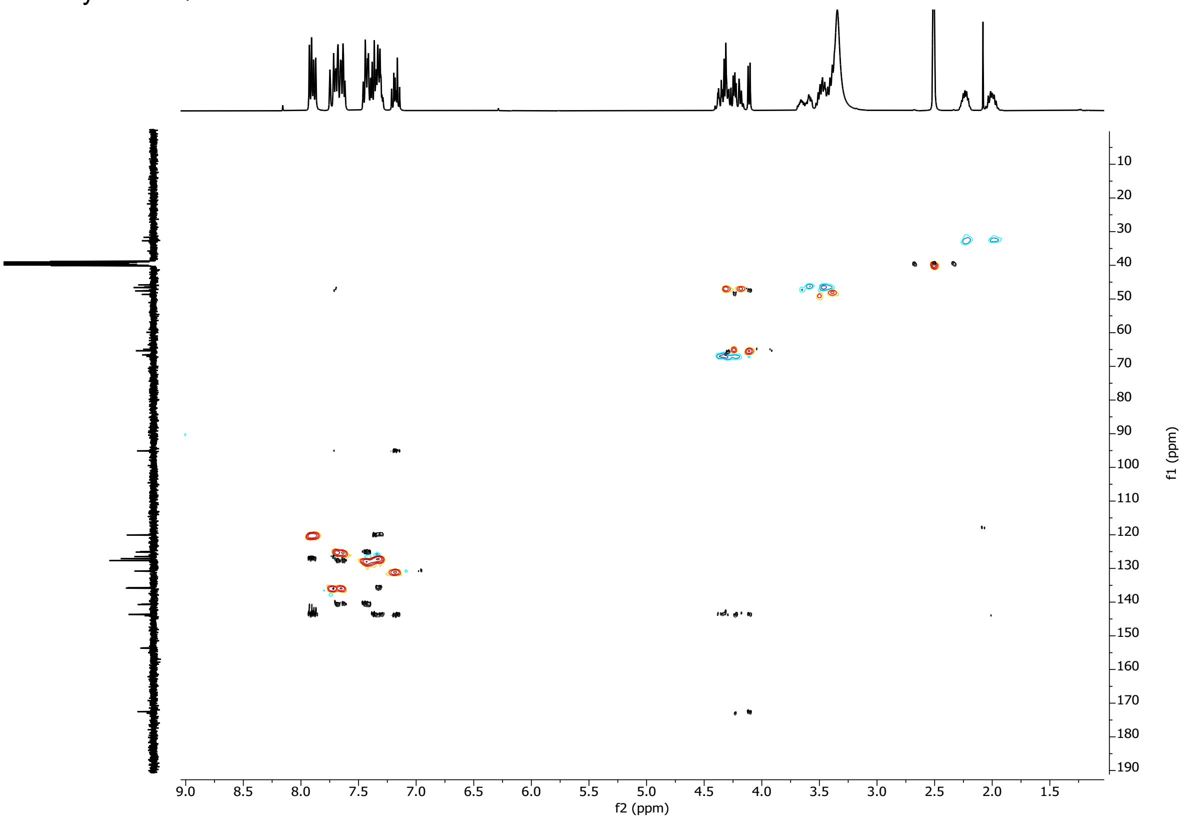

(2*S*,3*S*)-1-(((9H-fluoren-9-yl)methoxy)carbonyl)-3-(3-iodophenyl)piperidine-2-carboxylic acid (**7e**)

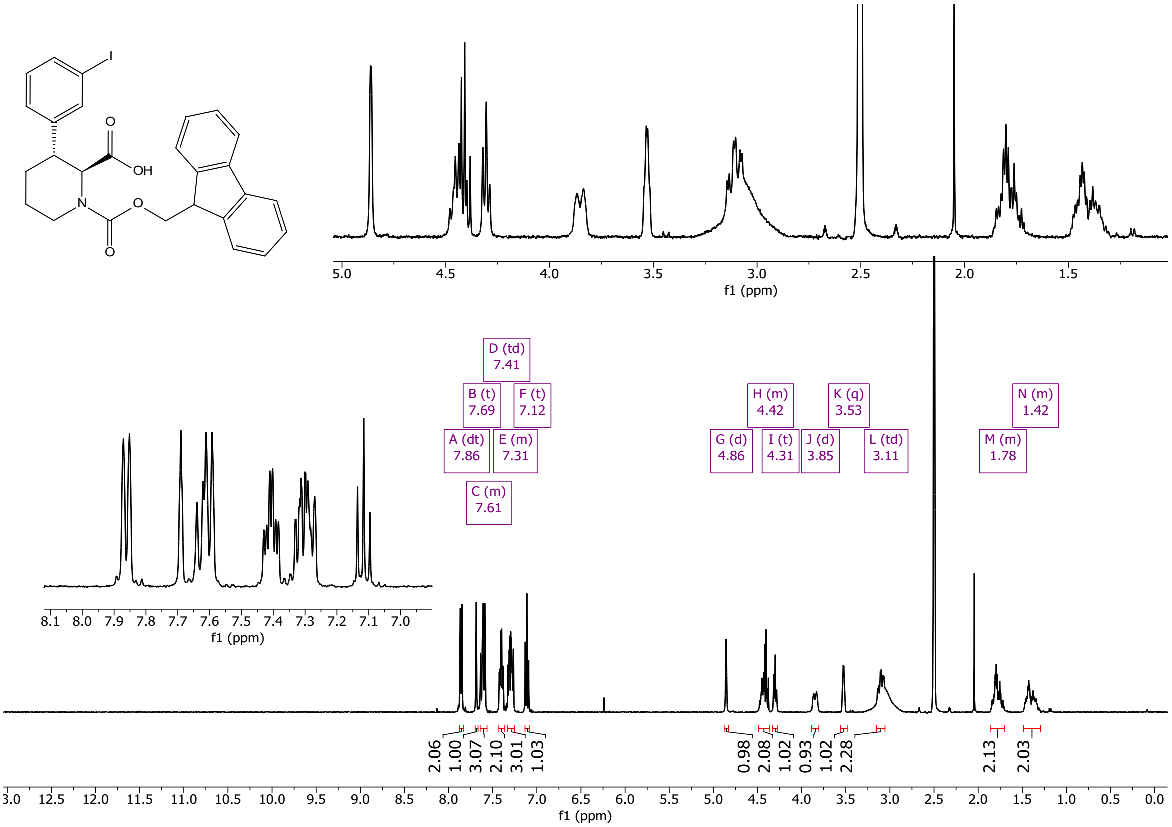

1, 2, 3 and 4 are spectra collected at 25, 40, 60 and 80°C, respectively:

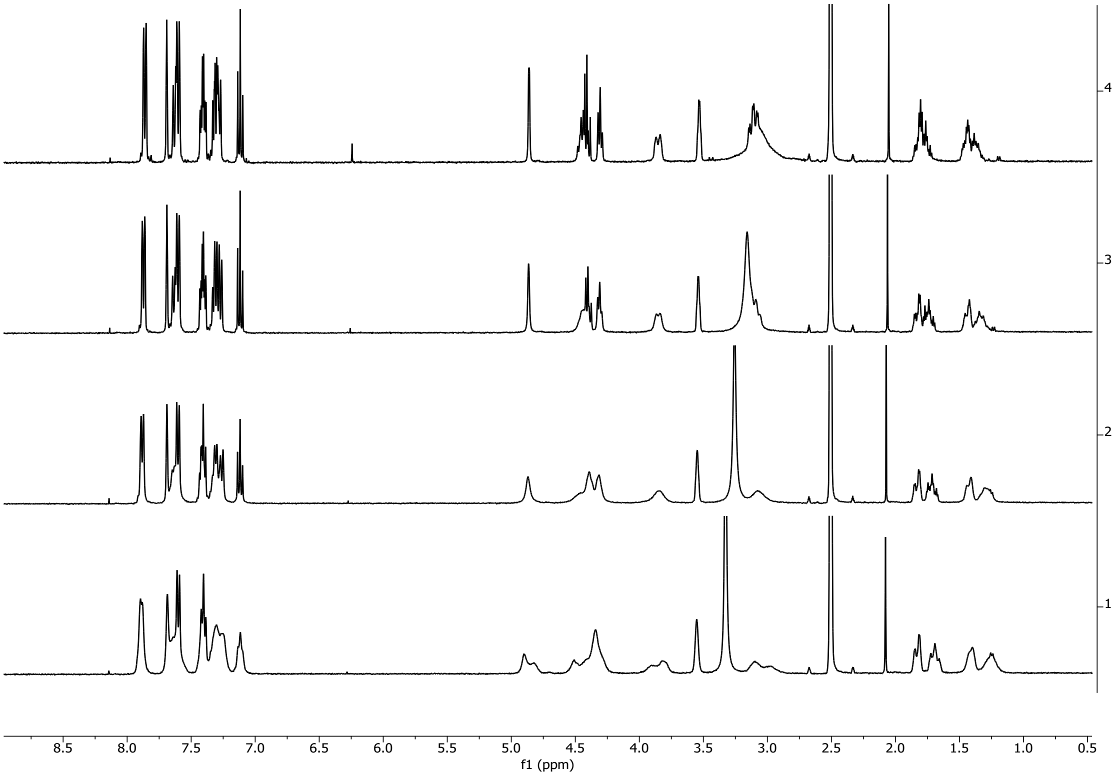

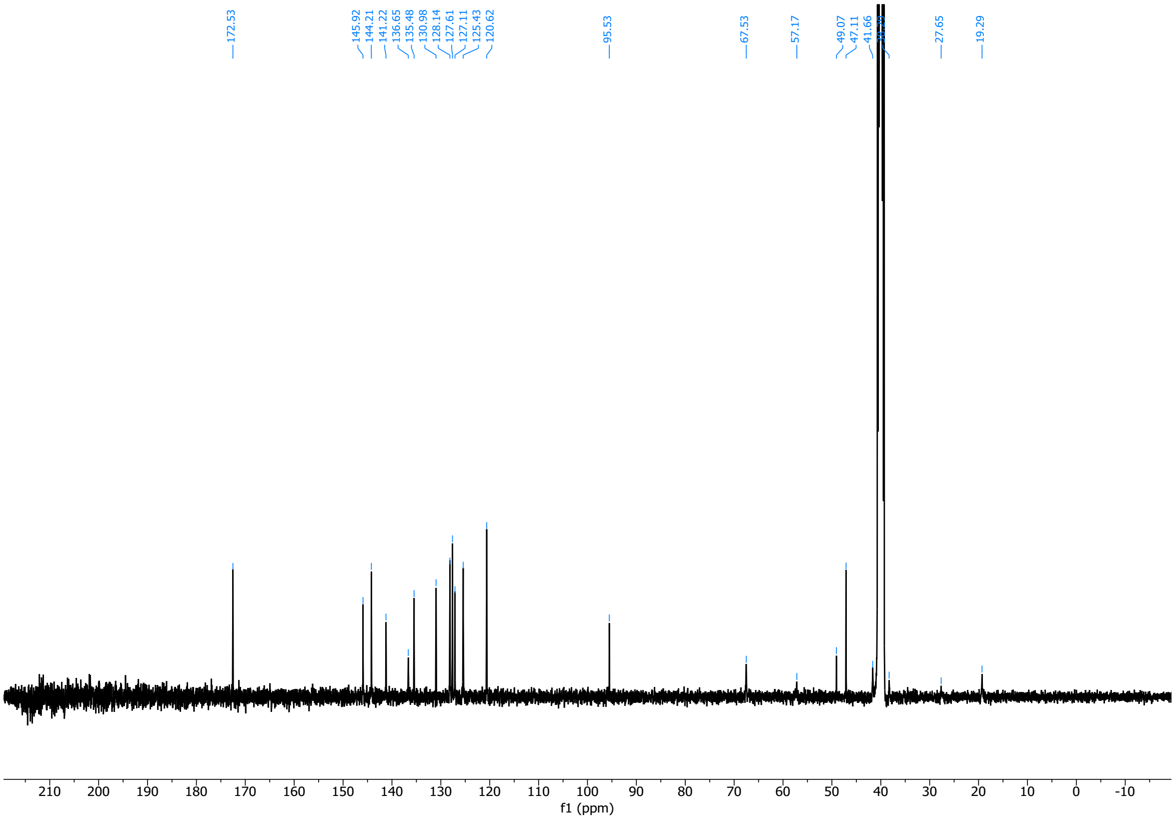

(2*R*,3*S*)-1-(((9H-fluoren-9-yl)methoxy)carbonyl)-3-(3-iodophenyl)piperidine-2-carboxylic acid (**7f**)

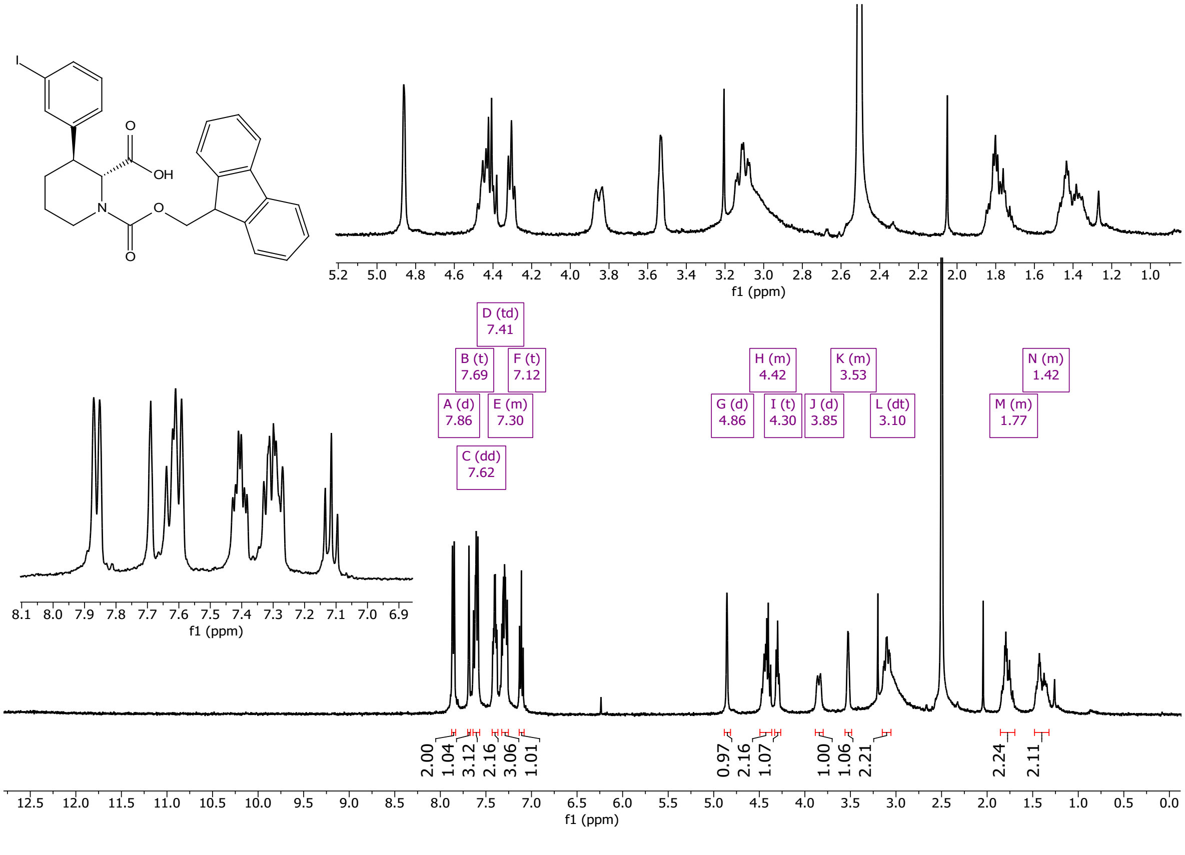

1, 2, 3 and 4 are spectra collected at 25, 40, 60 and 80°C, respectively:

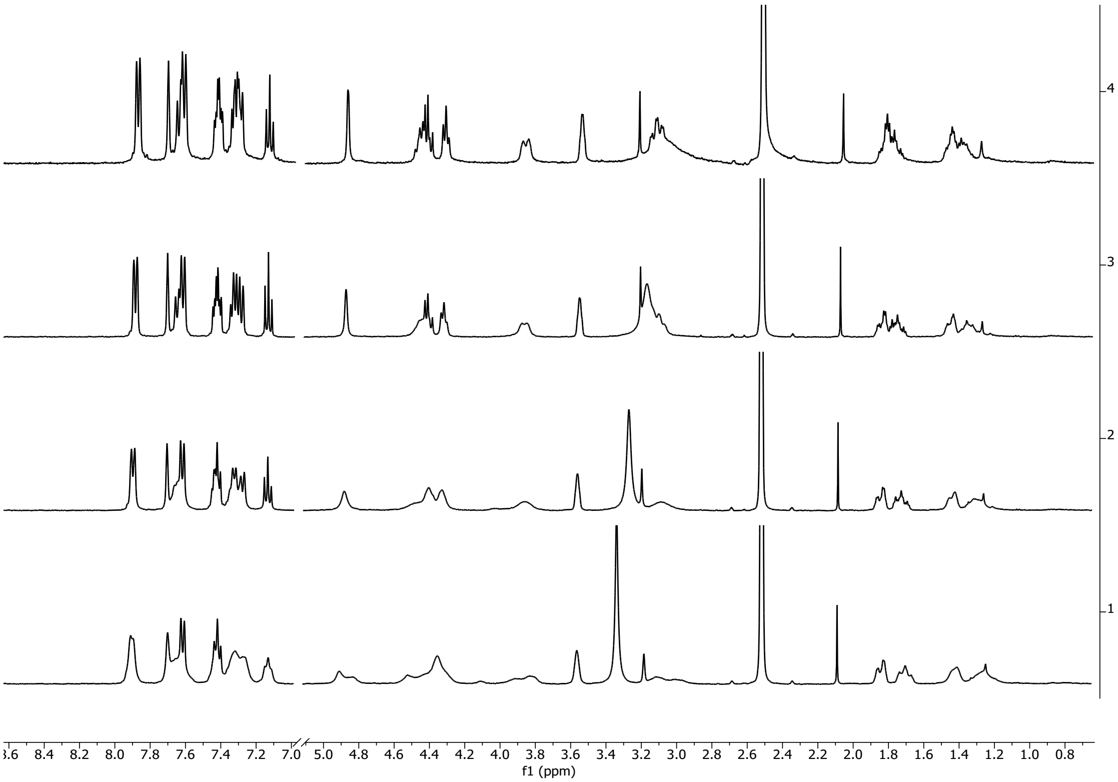

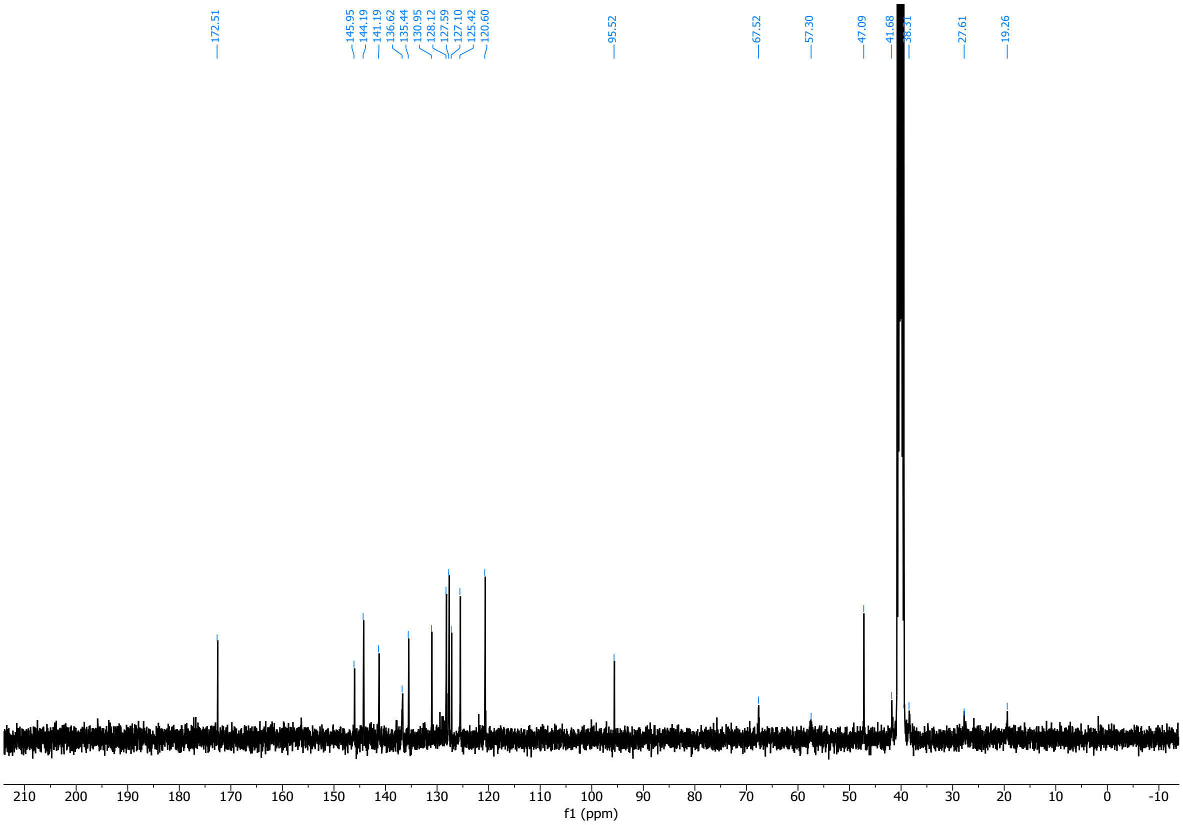

#### 7.2. Hydroxyproline-derived skeletons

(2*S*,4*S*)-1-tert-butyl 2-methyl 4-(4-iodophenoxy)pyrrolidine-1,2-dicarboxylate (**9ai**)

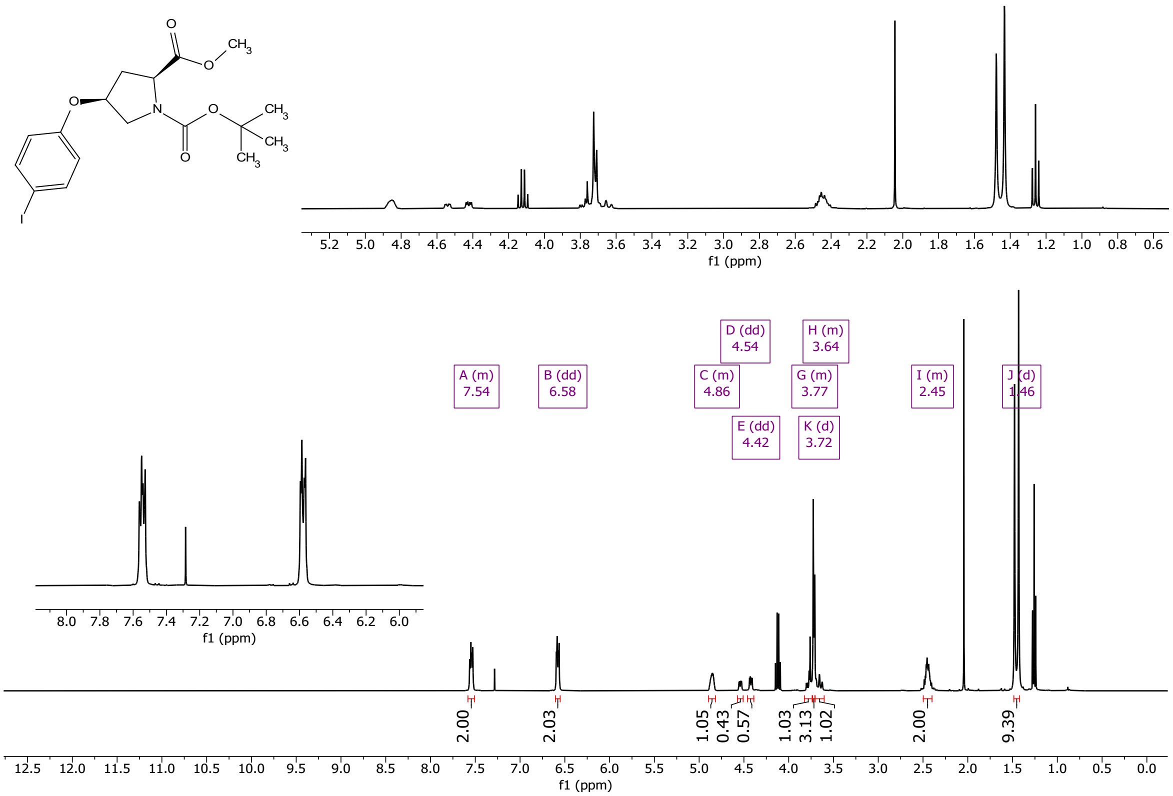

2988 (2*S*,4*R*)-1-tert-butyl 2-methyl 4-(4-iodophenoxy)pyrrolidine-1,2-dicarboxylate (**9bi**)

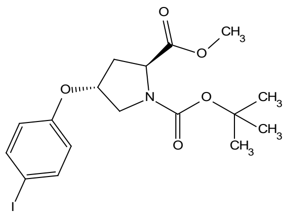

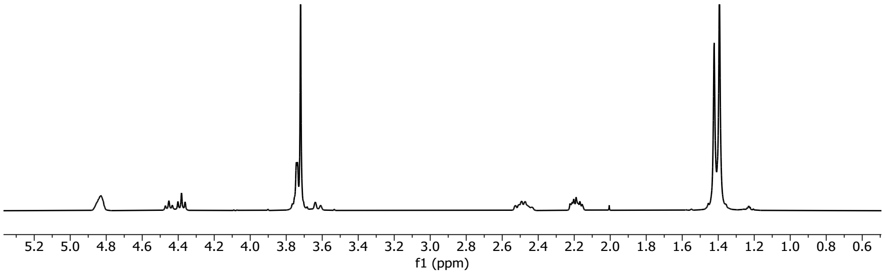

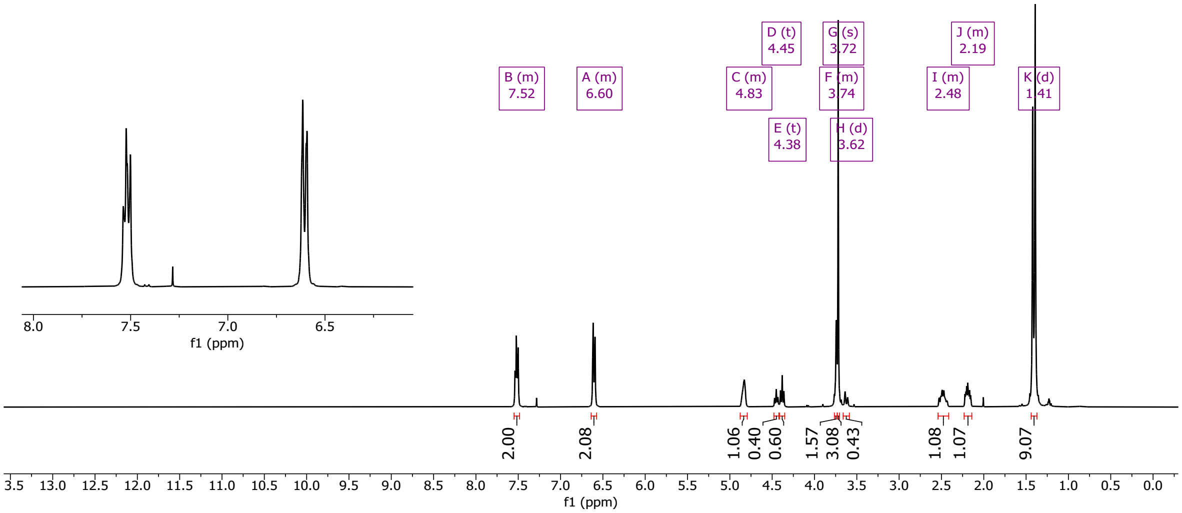

2989

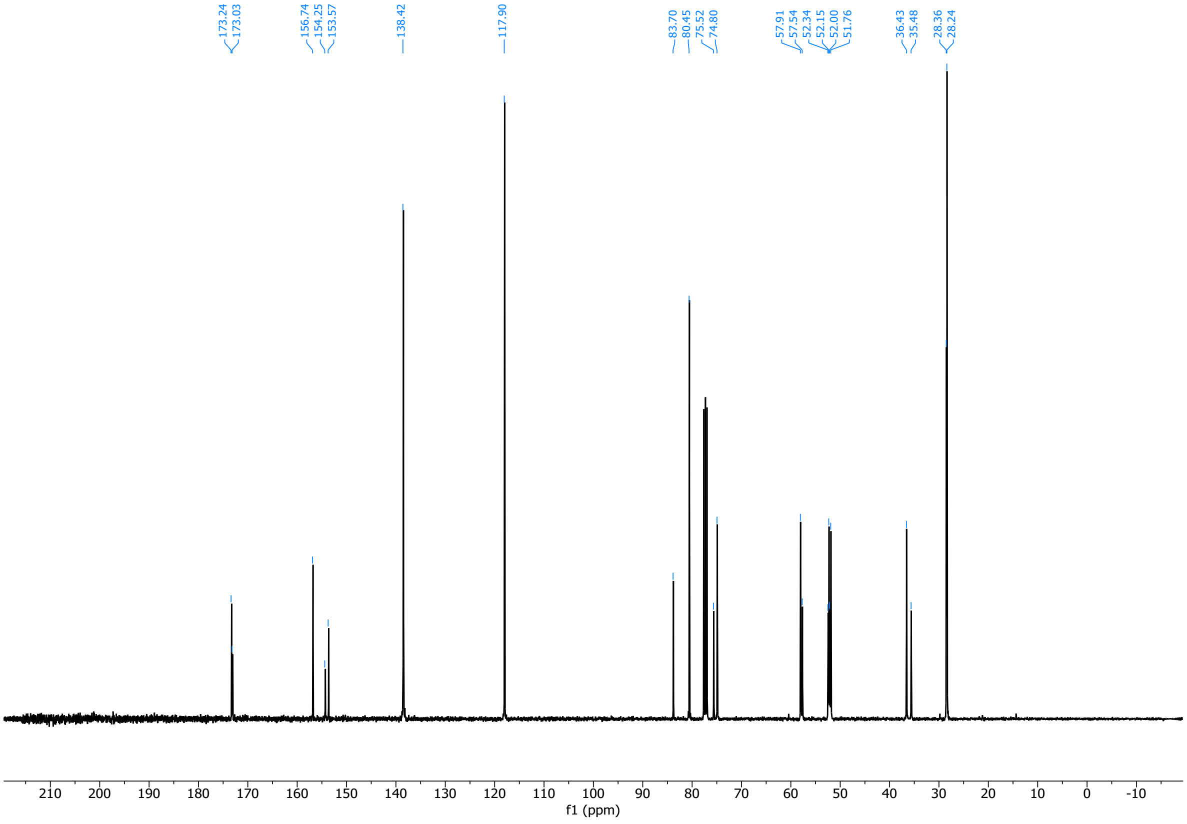

2990

(2*R*,4*R*)-1-tert-butyl 2-methyl 4-(4-iodophenoxy)pyrrolidine-1,2-dicarboxylate (**9ci**)

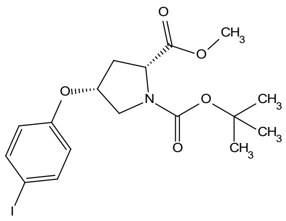

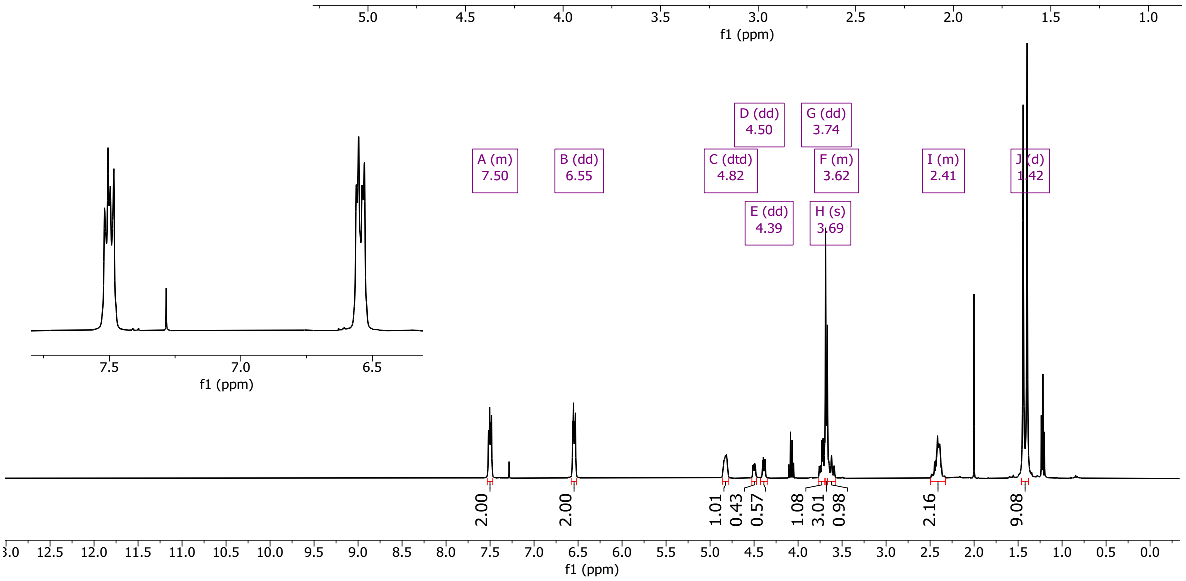

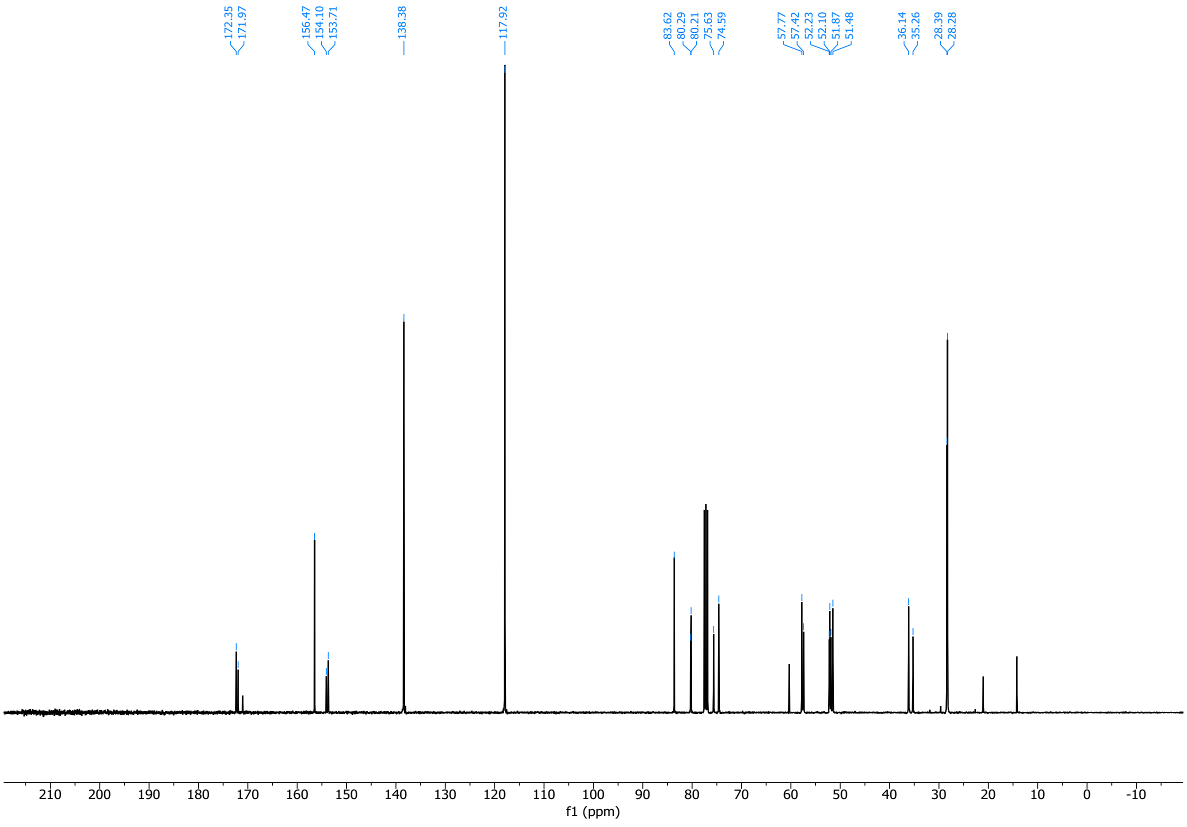

(2*R*,4*S*)-1-tert-butyl 2-methyl 4-(4-iodophenoxy)pyrrolidine-1,2-dicarboxylate (**9di**)

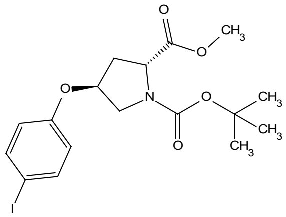

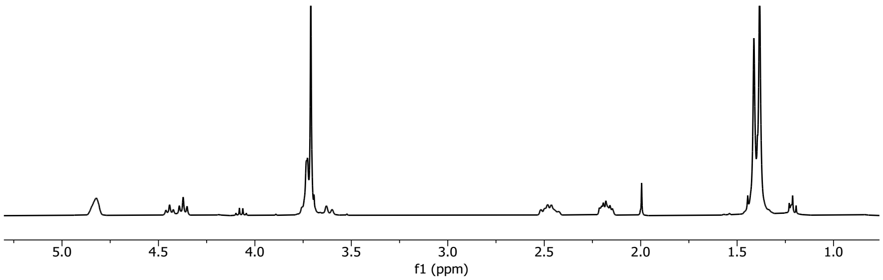

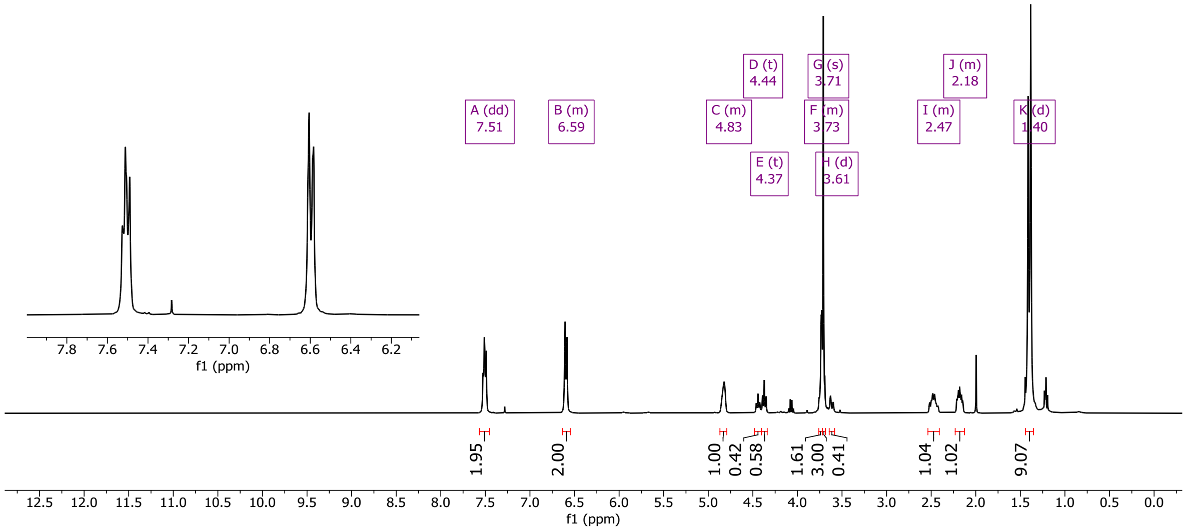

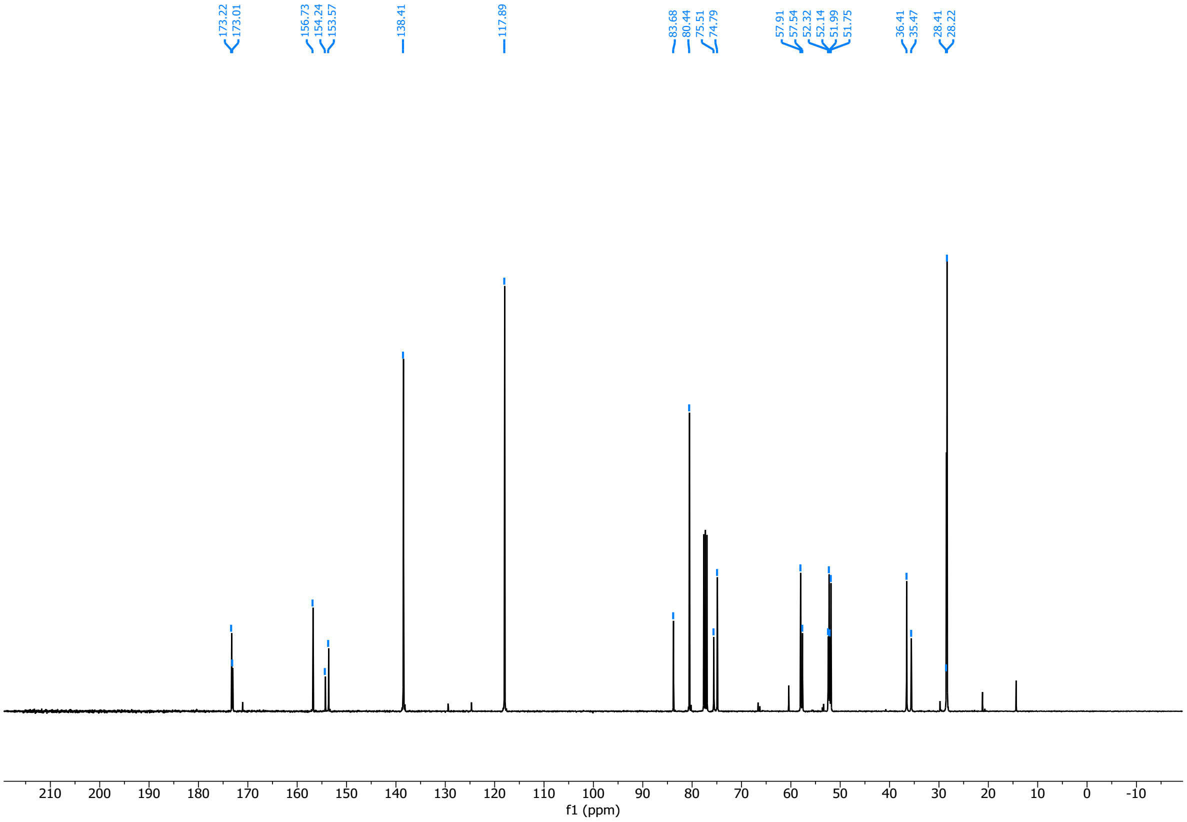

2997 (2*S*,4*S*)-1-tert-butyl 2-methyl 4-(3-iodophenoxy)pyrrolidine-1,2-dicarboxylate (**9aii**)

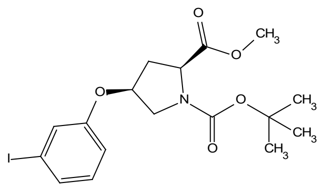

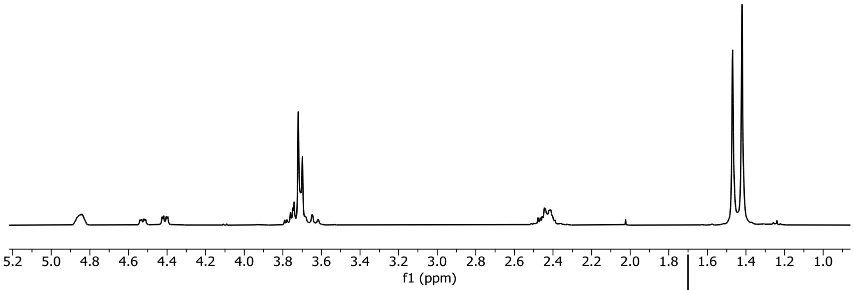

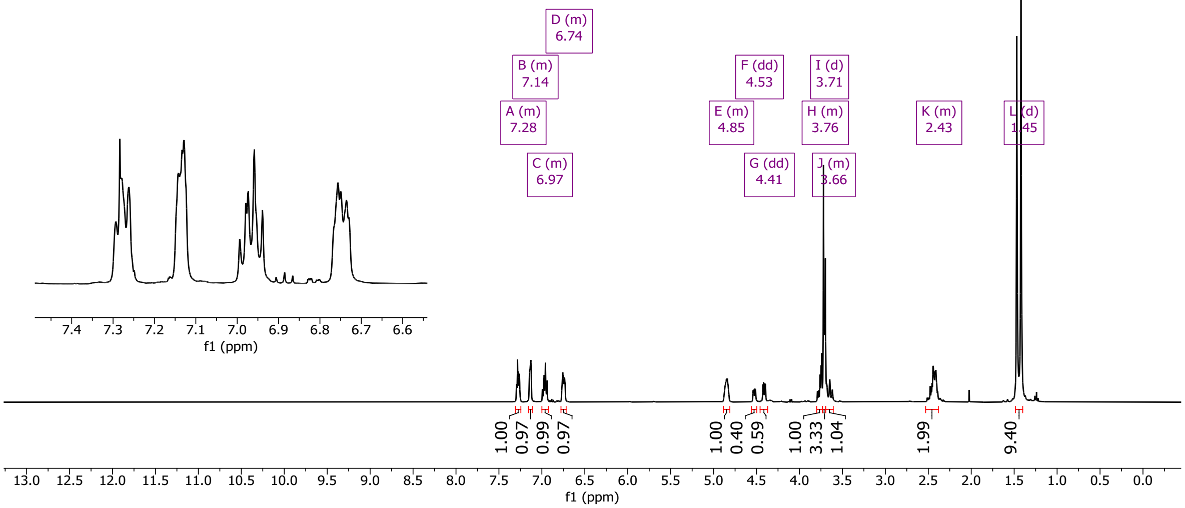

2998

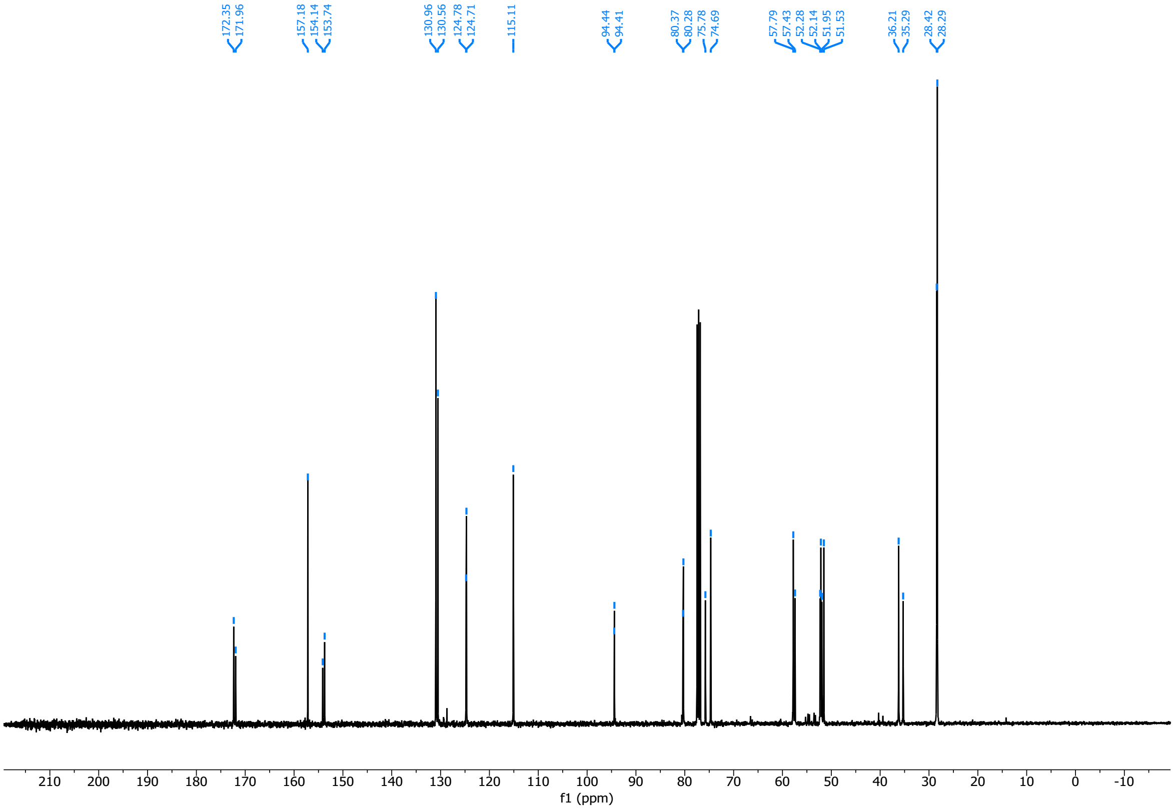

2999

3000 (2*S*,4*R*)-1-tert-butyl 2-methyl 4-(3-iodophenoxy)pyrrolidine-1,2-dicarboxylate (**9bii**)

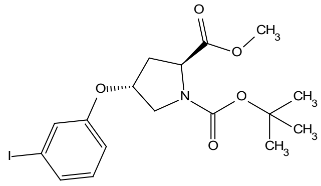

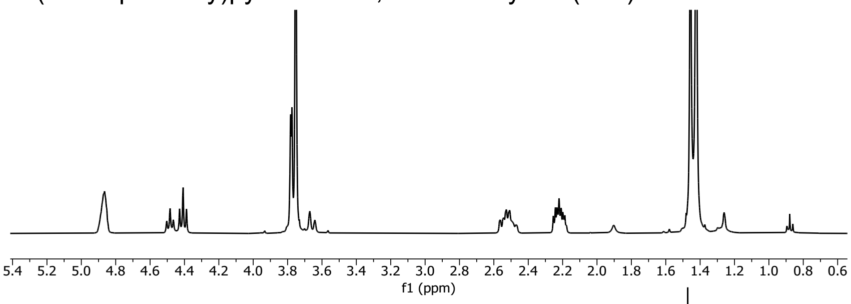

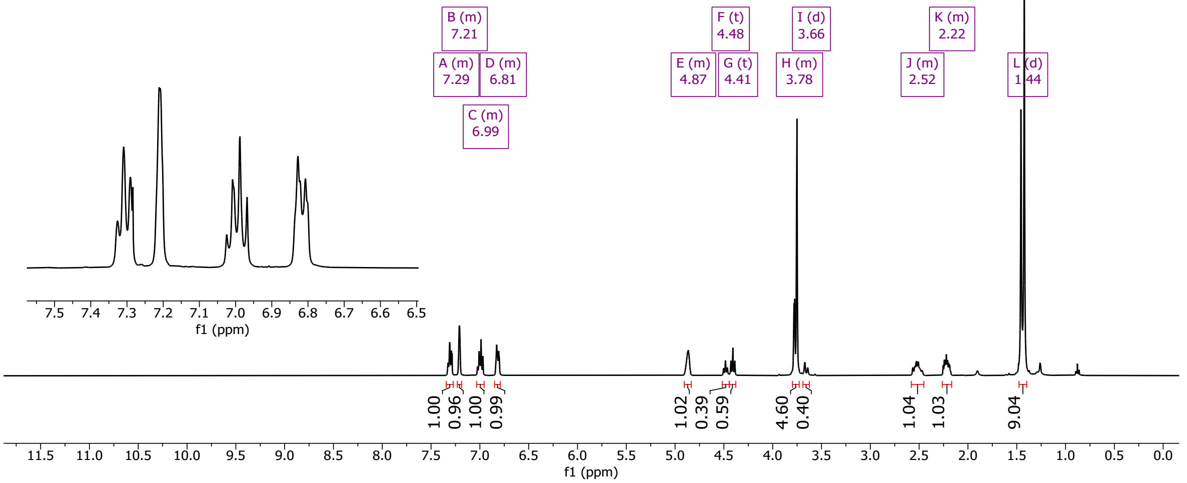

3001

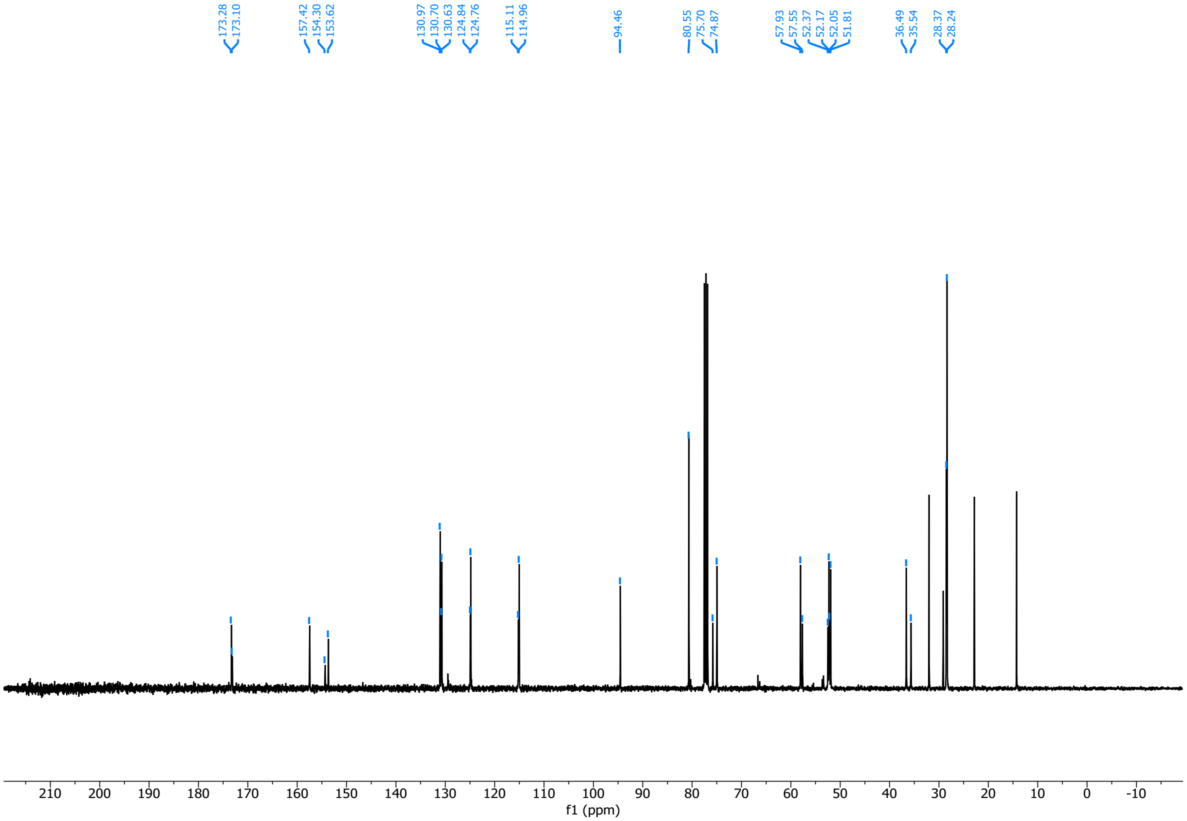

3002

3003 (2*R*,4*R*)-1-tert-butyl 2-methyl 4-(3-iodophenoxy)pyrrolidine-1,2-dicarboxylate (**9cii**)

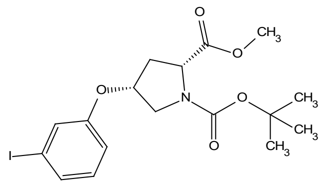

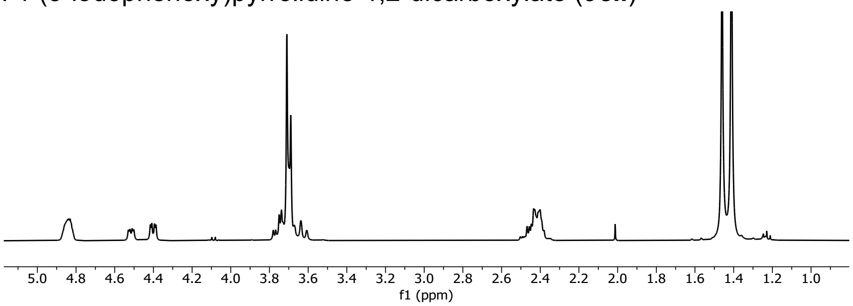

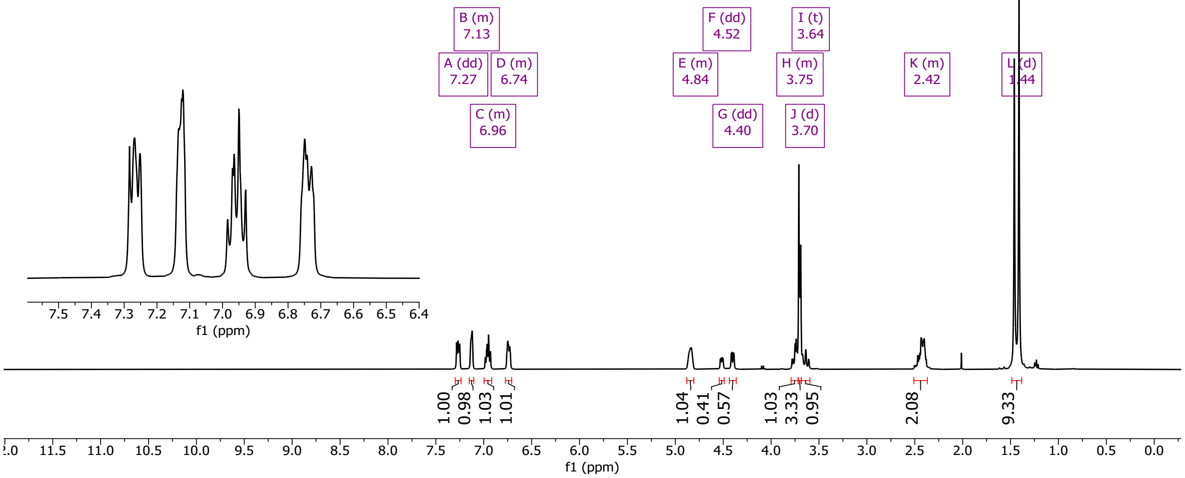

3004

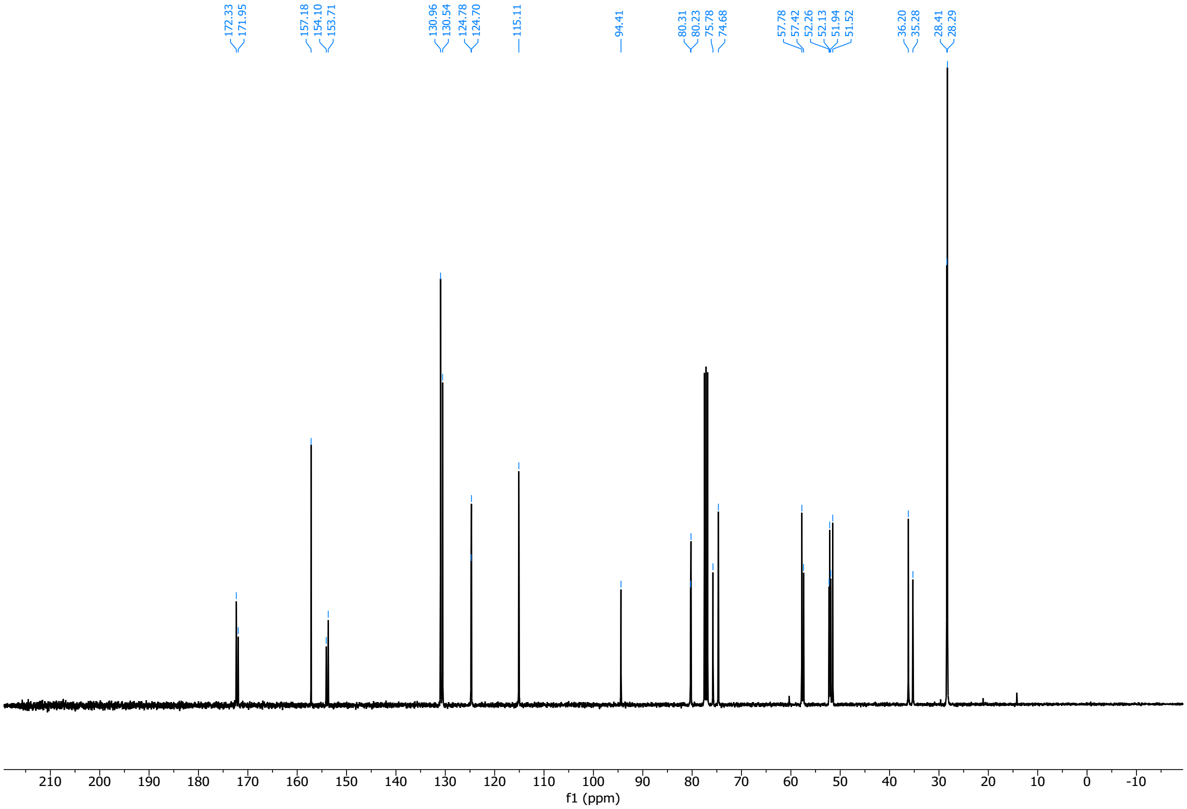

3005

3006 (2*R*,4*S*)-1-tert-butyl 2-methyl 4-(3-iodophenoxy)pyrrolidine-1,2-dicarboxylate (**9dii**)

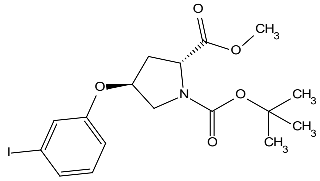

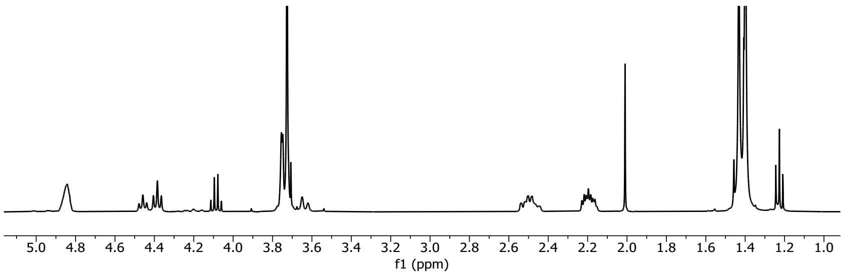

3007

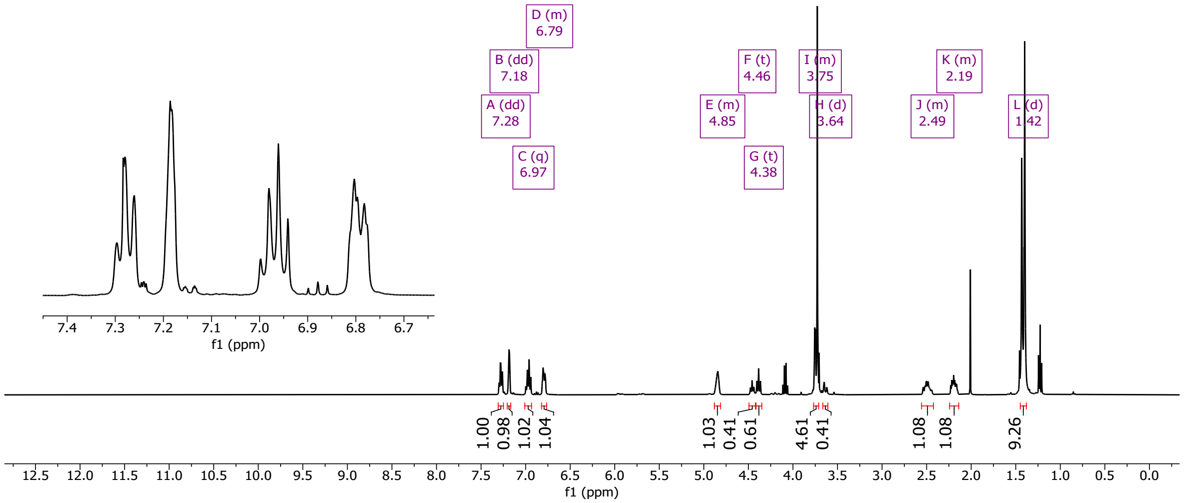

3008

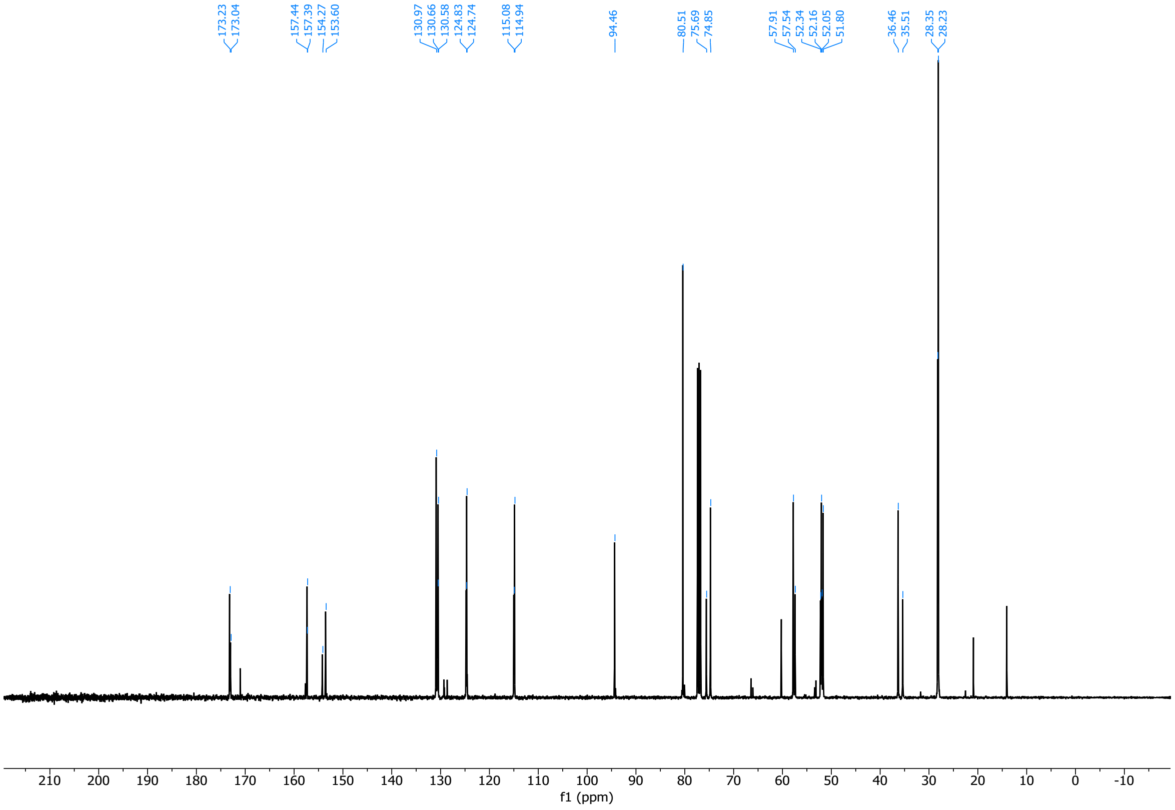

(2*S*,4*S*)-1-(tert-butoxycarbonyl)-4-((4-iodobenzyl)oxy)pyrrolidine-2-carboxylic acid (**13ai**)

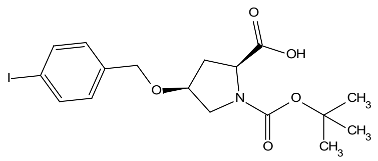

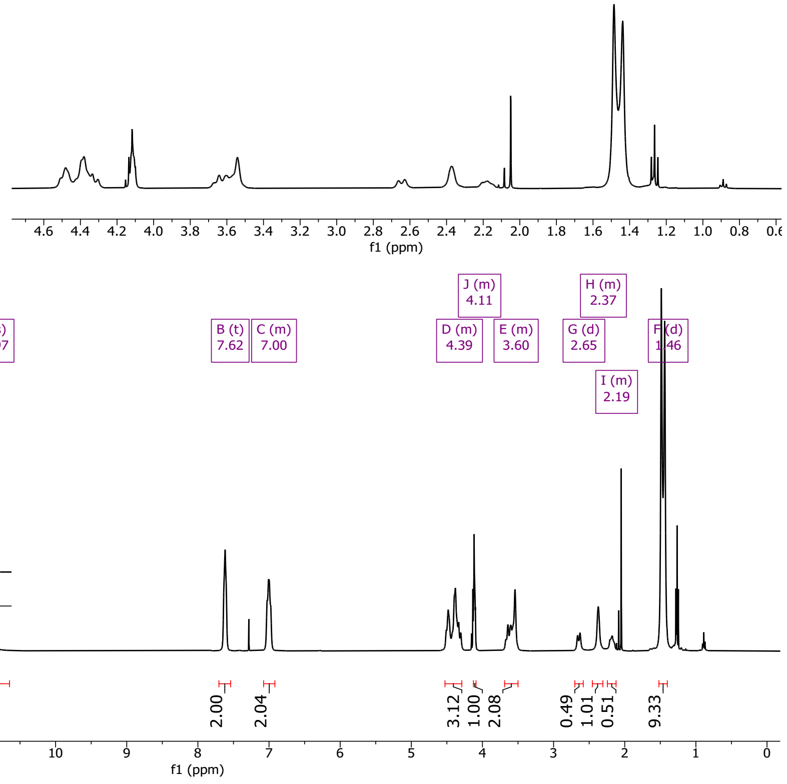

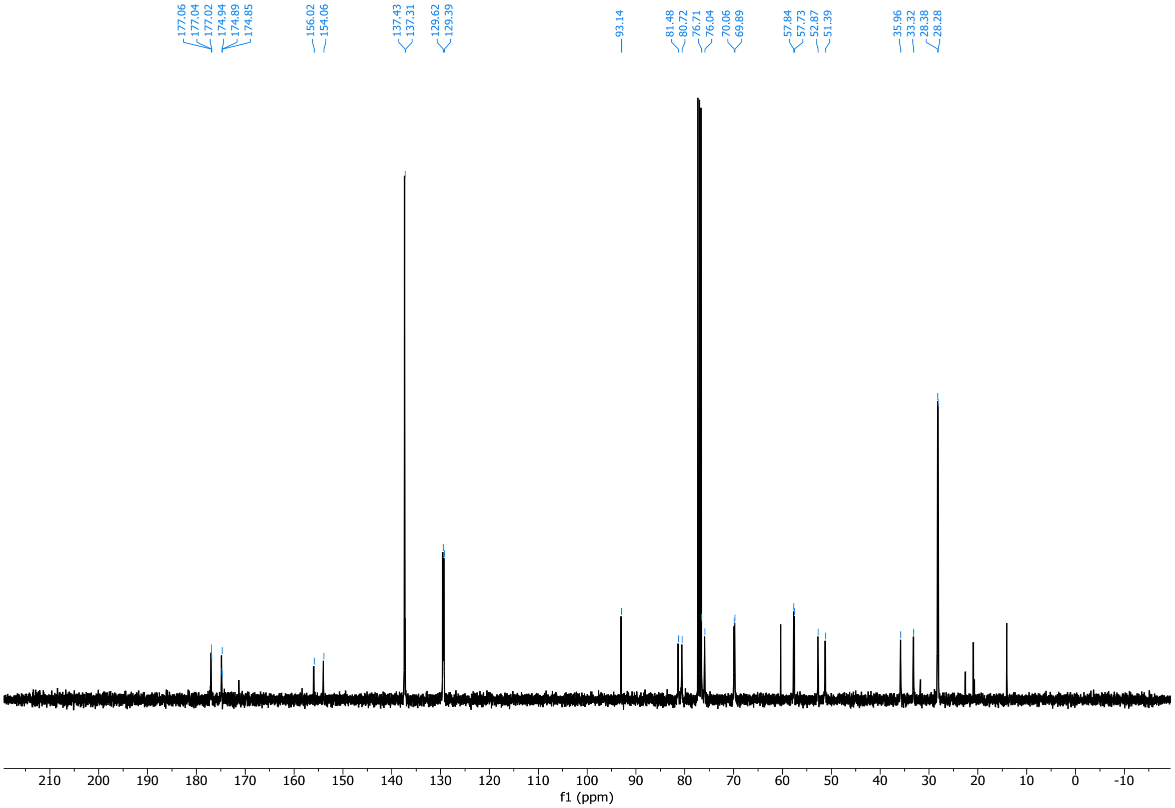

3012 (2*S*,4*R*)-1-(tert-butoxycarbonyl)-4-((4-iodobenzyl)oxy)pyrrolidine-2-carboxylic acid (**13bi**)

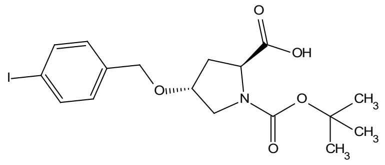

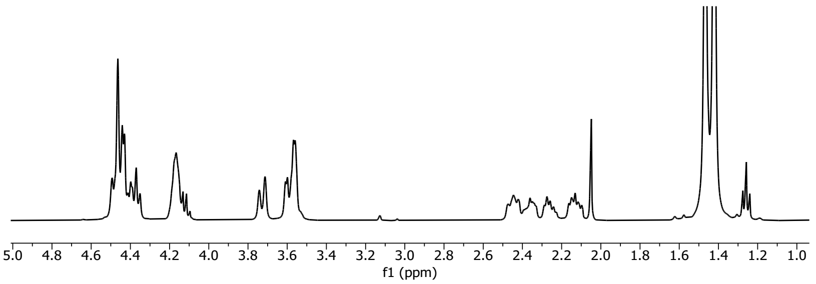

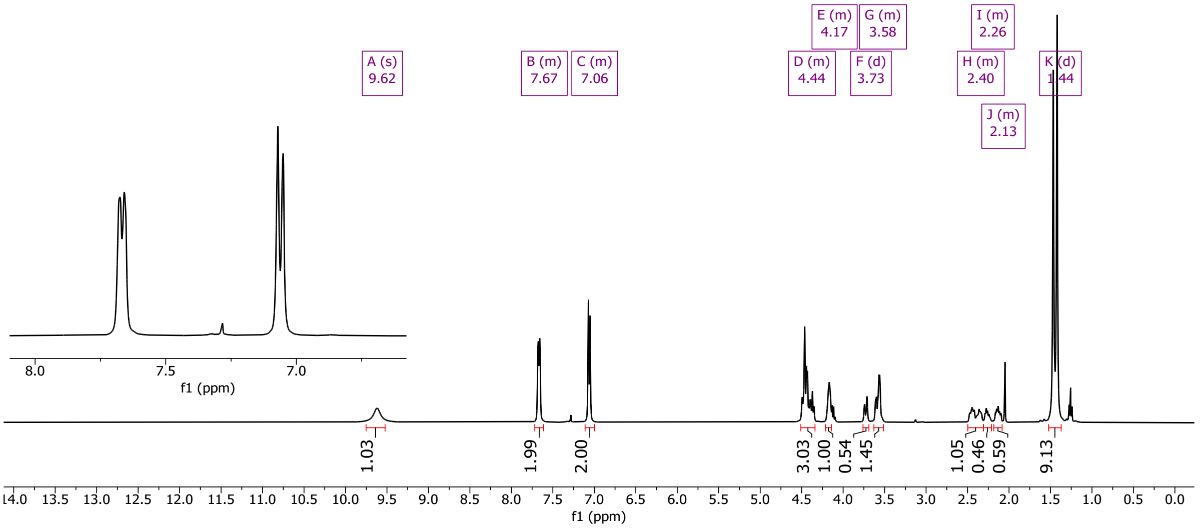

3013

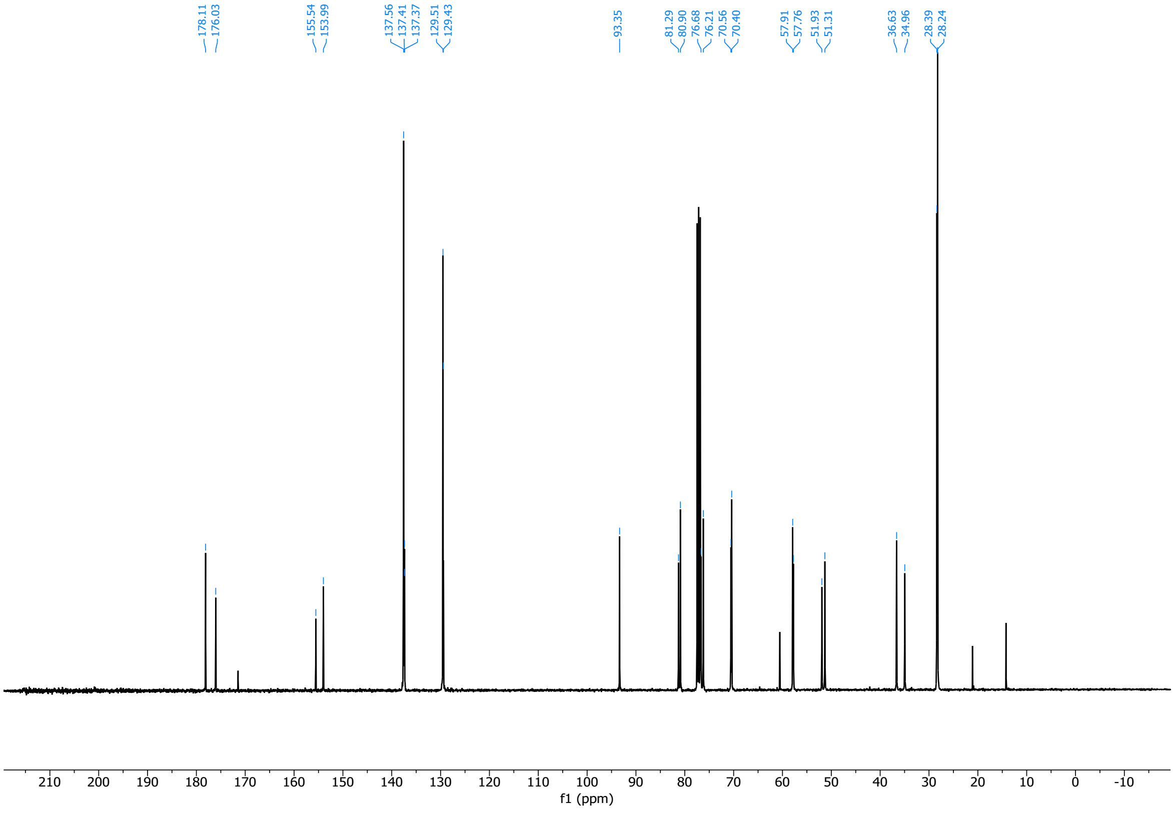

3014

3015 (2*R*,4*R*)-1-(tert-butoxycarbonyl)-4-((4-iodobenzyl)oxy)pyrrolidine-2-carboxylic acid (**13ci**)

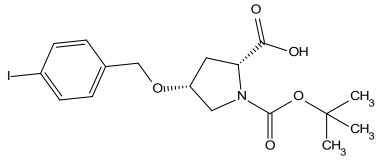

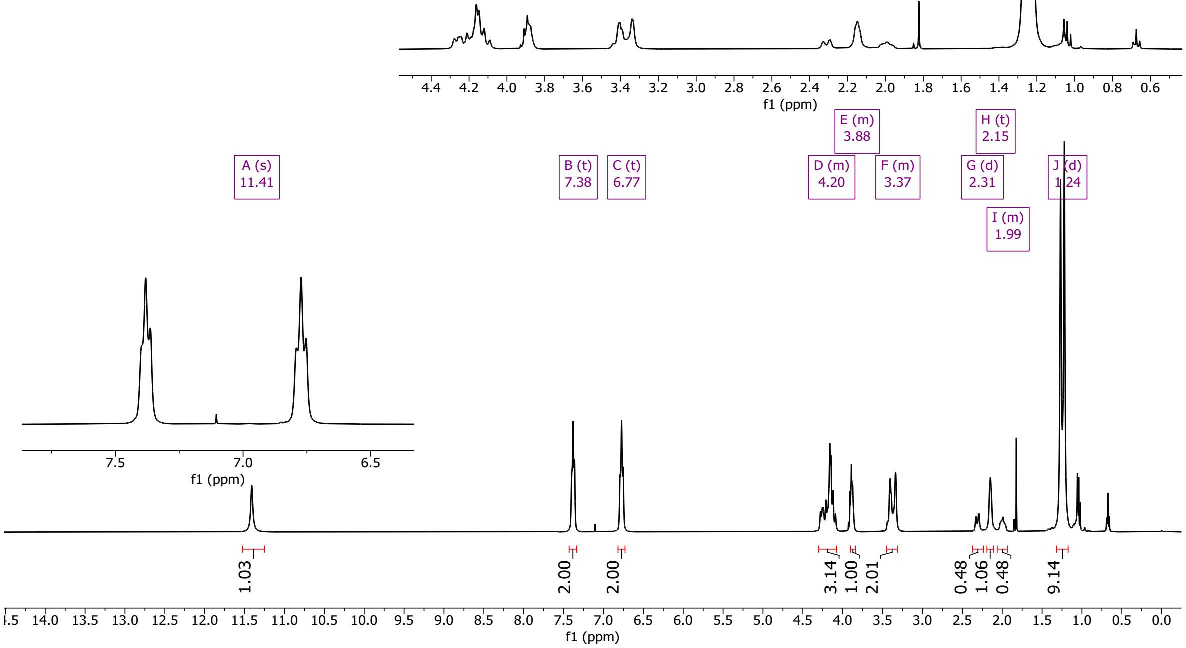

3016

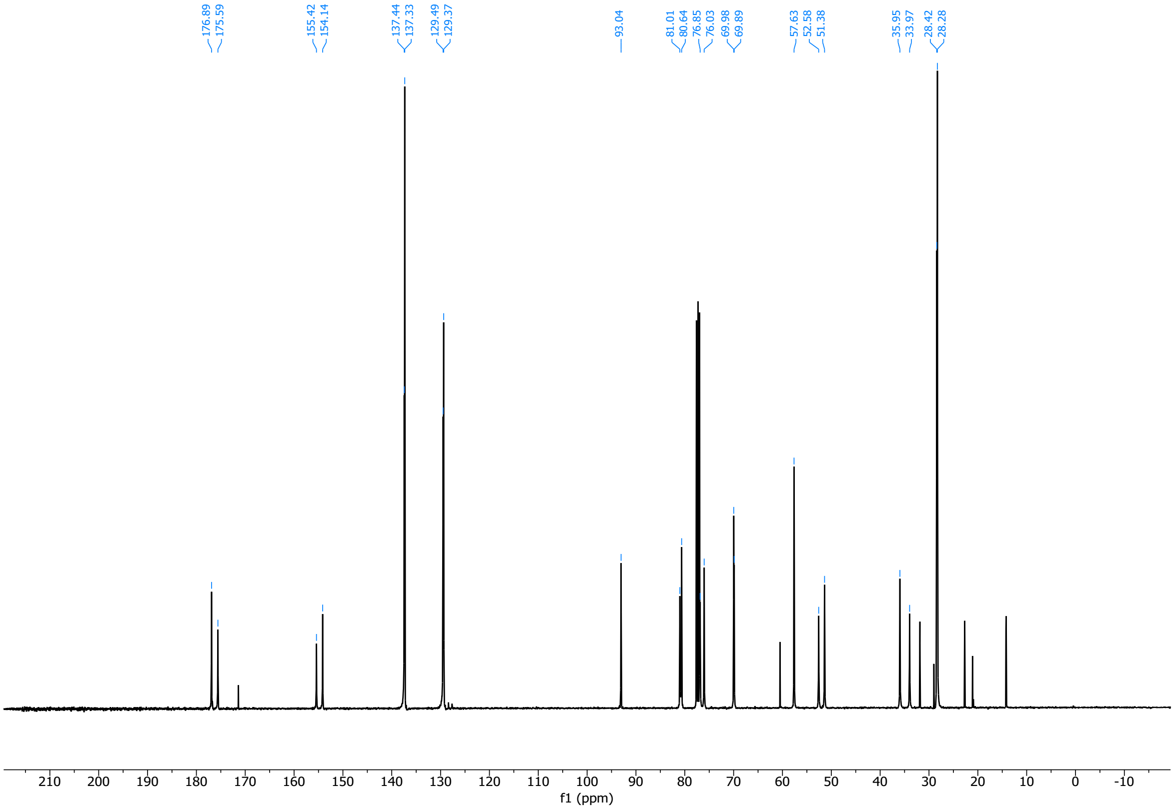

3017

(2*R*,4*S*)-1-(tert-butoxycarbonyl)-4-((4-iodobenzyl)oxy)pyrrolidine-2-carboxylic acid (**13di**)

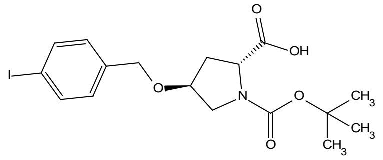

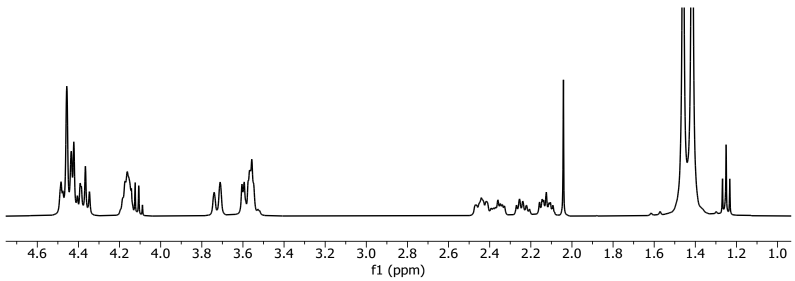

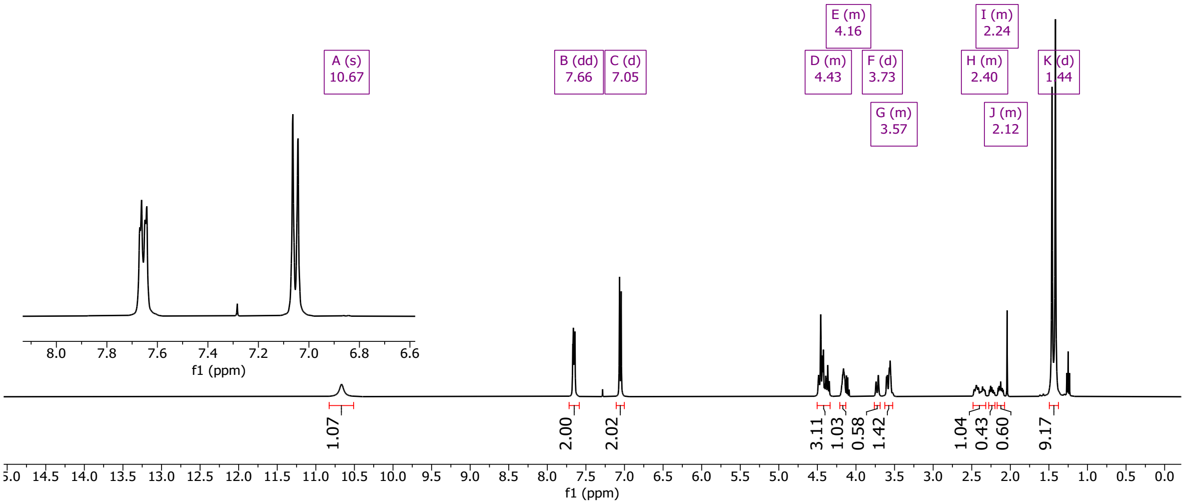

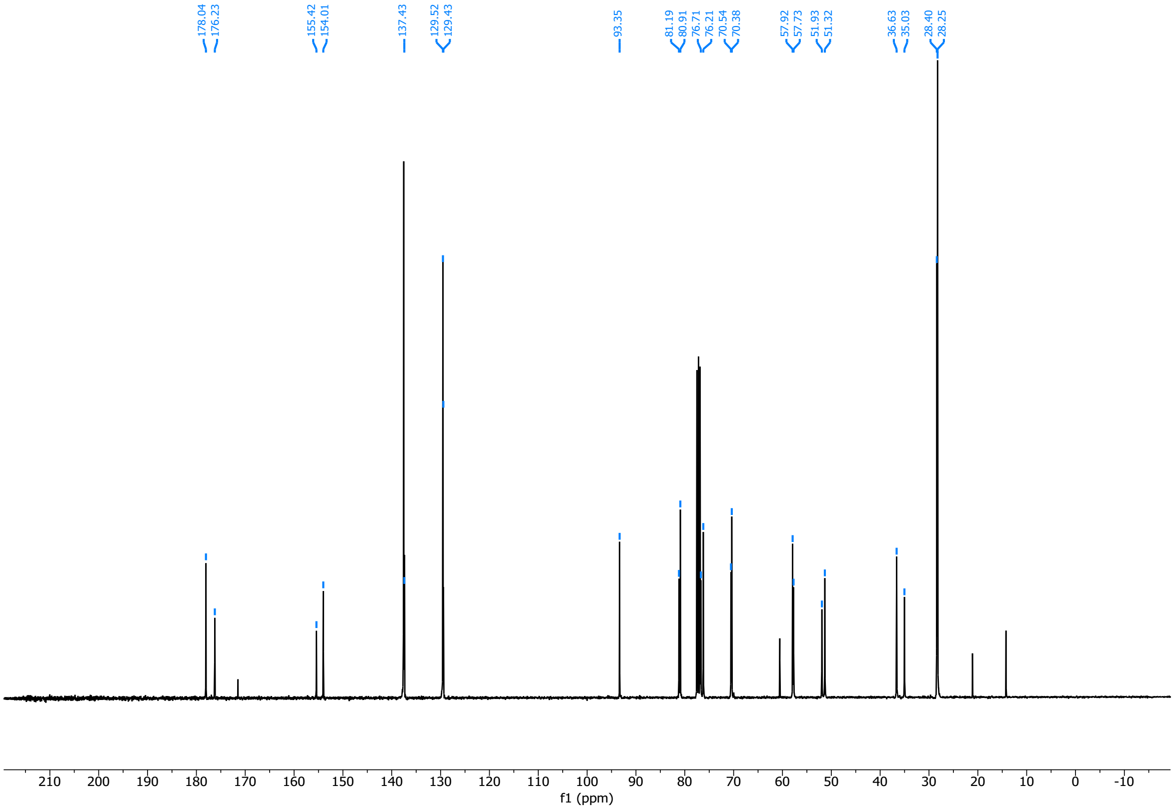

(2*R*,4*S*)-1-(*tert*-butoxycarbonyl)-4-((3-iodobenzyl)oxy)pyrrolidine-2-carboxylic acid (**14aii**)

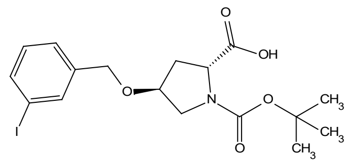

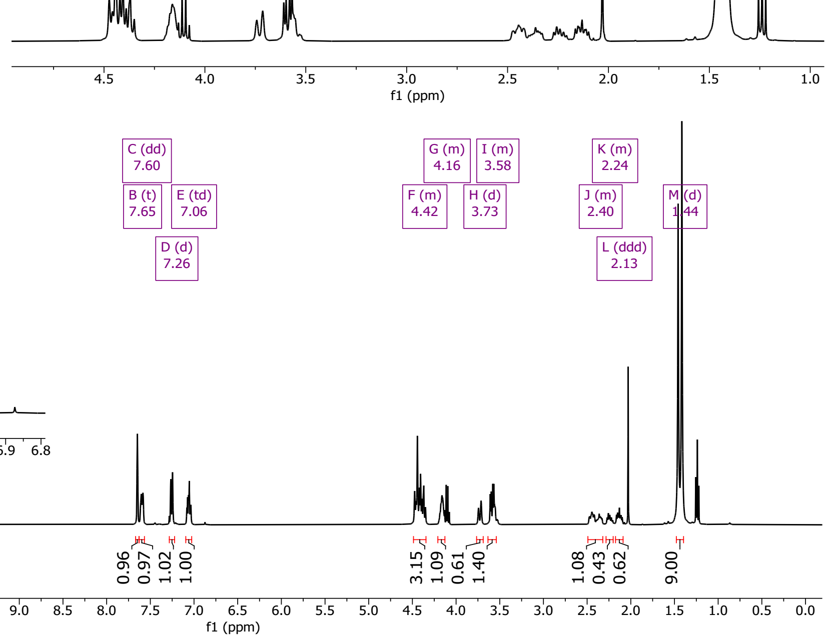

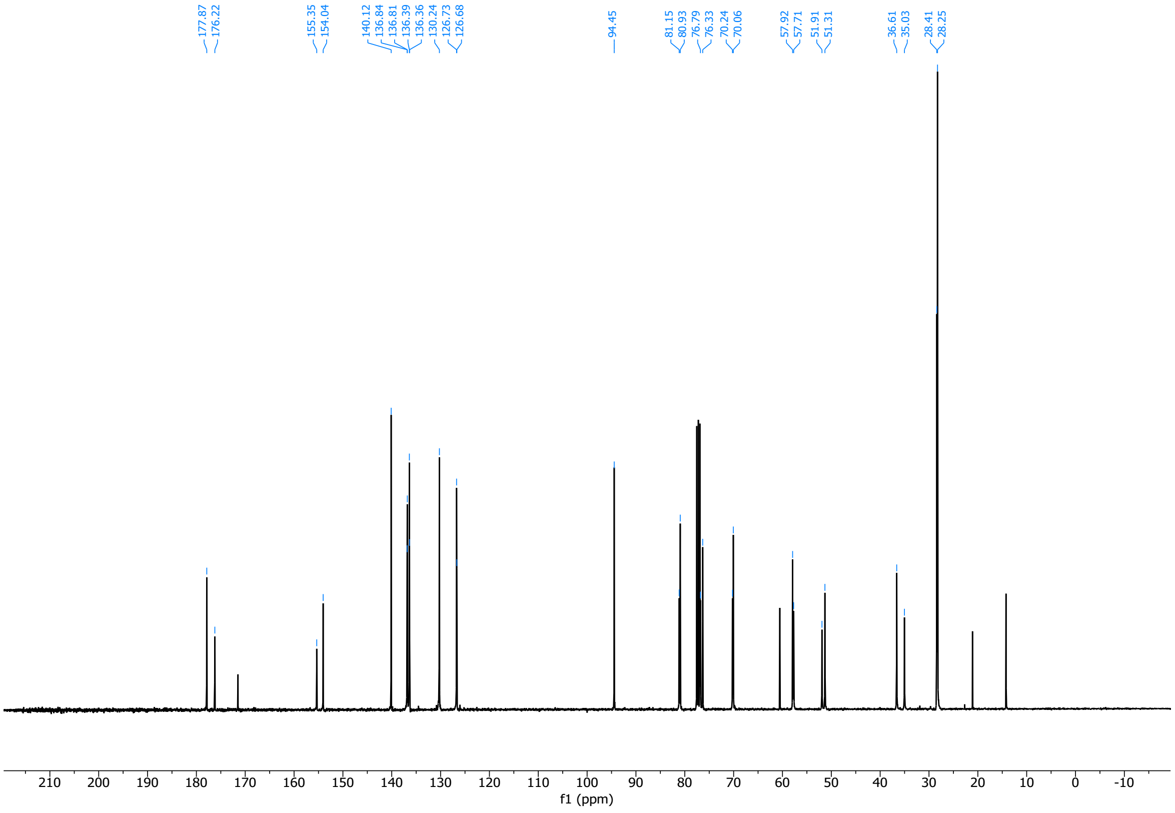

(2*S*,4*S*)-1-(*tert*-butoxycarbonyl)-4-((3-iodobenzyl)oxy)pyrrolidine-2-carboxylic acid (**14bii**)

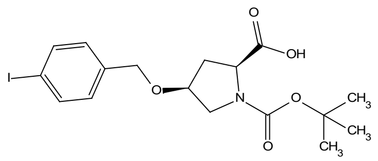

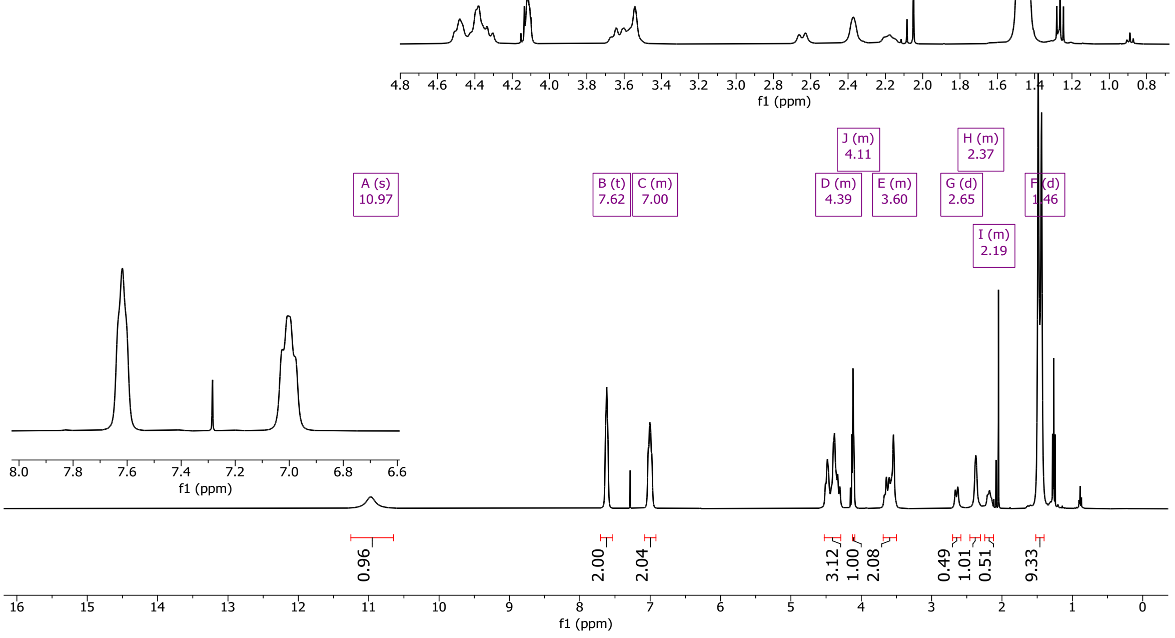

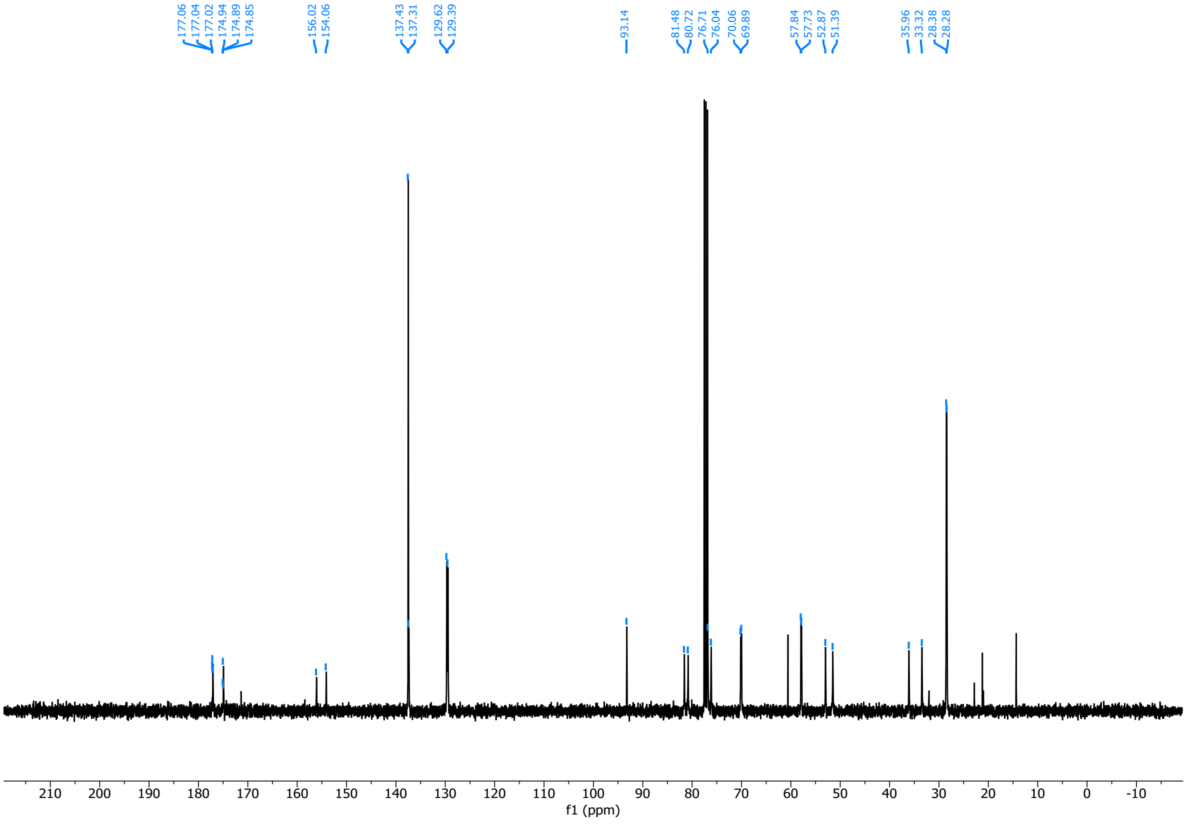

3027 (2*S*,4*R*)-1-(*tert*-butoxycarbonyl)-4-((3-iodobenzyl)oxy)pyrrolidine-2-carboxylic acid (**14cii**)

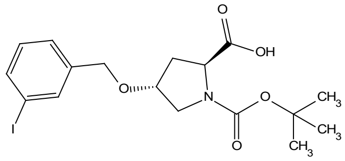

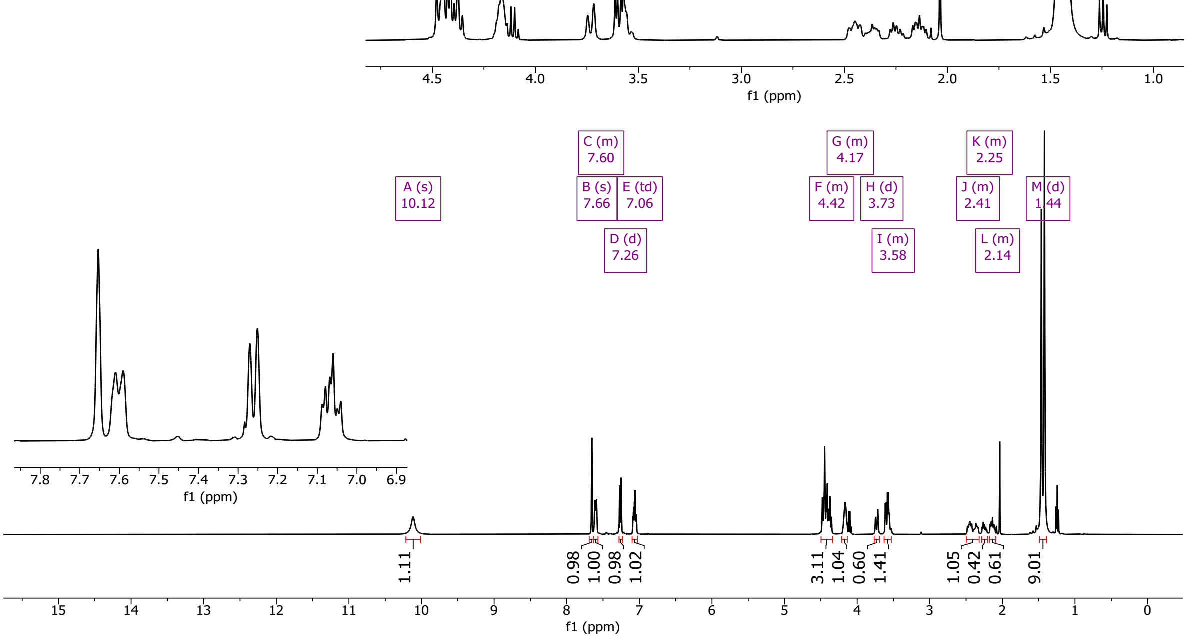

3028

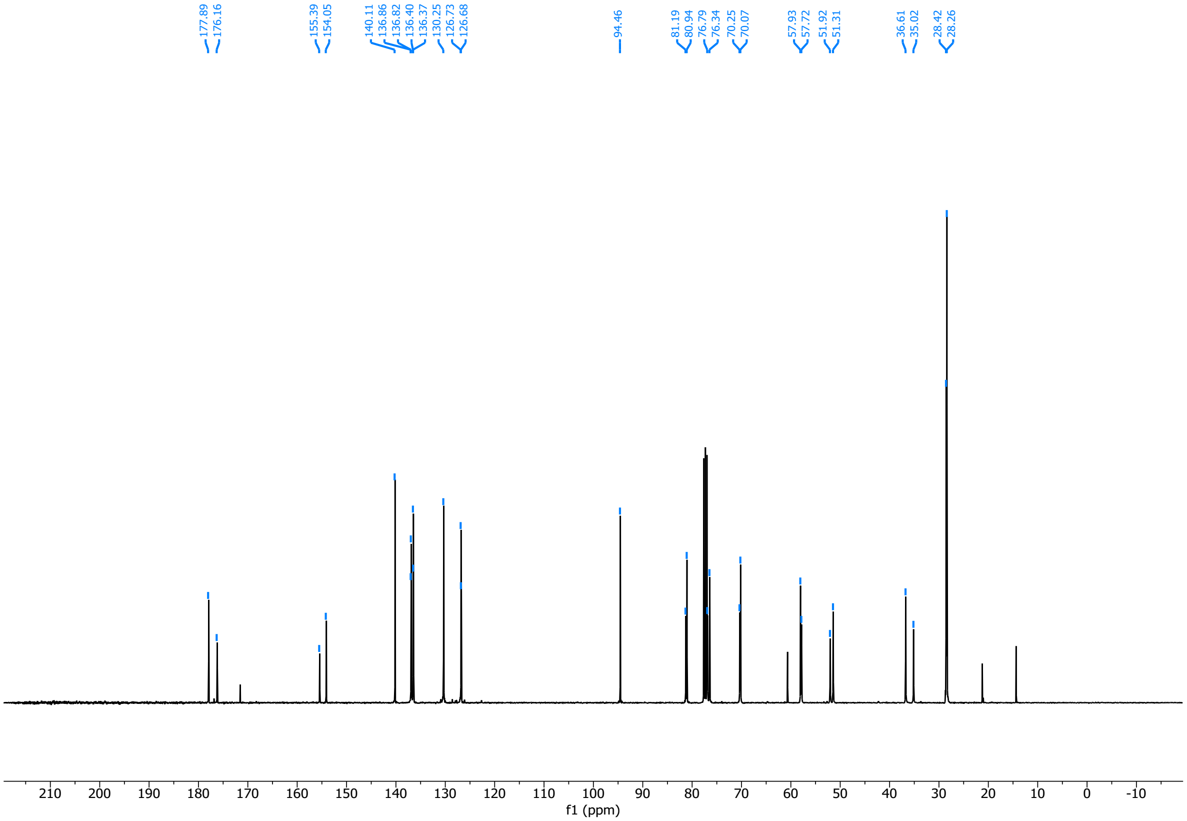

3029

3030 (2*R*,4*R*)-1-(*tert*-butoxycarbonyl)-4-((3-iodobenzyl)oxy)pyrrolidine-2-carboxylic acid (**14dii**)

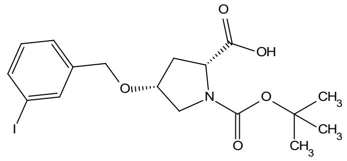

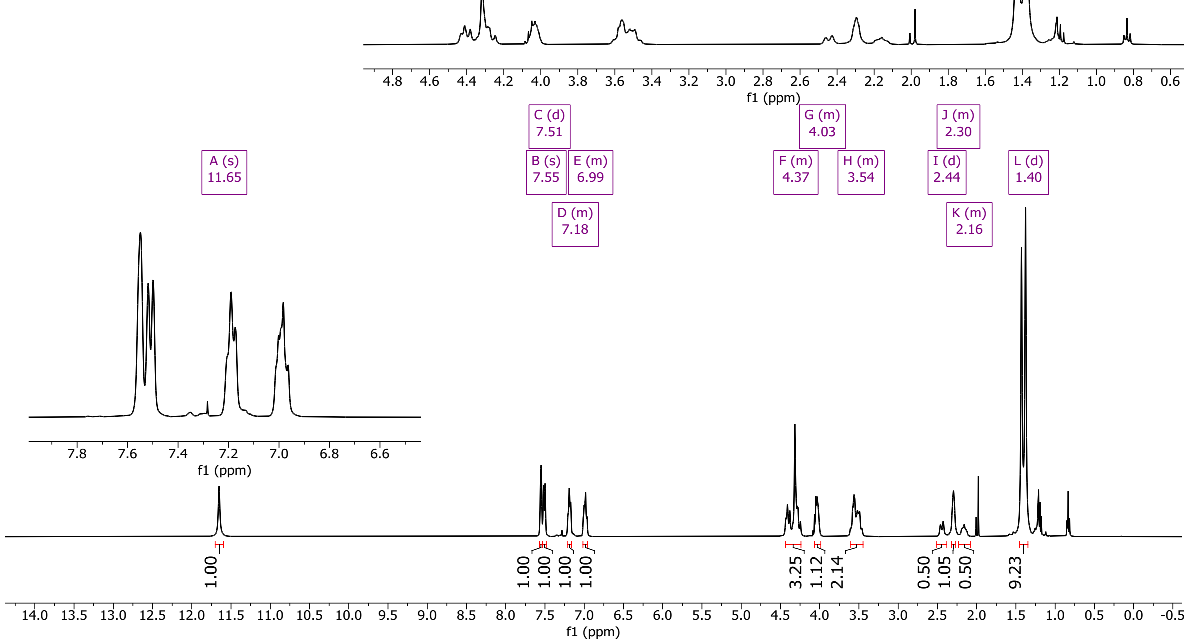

3031

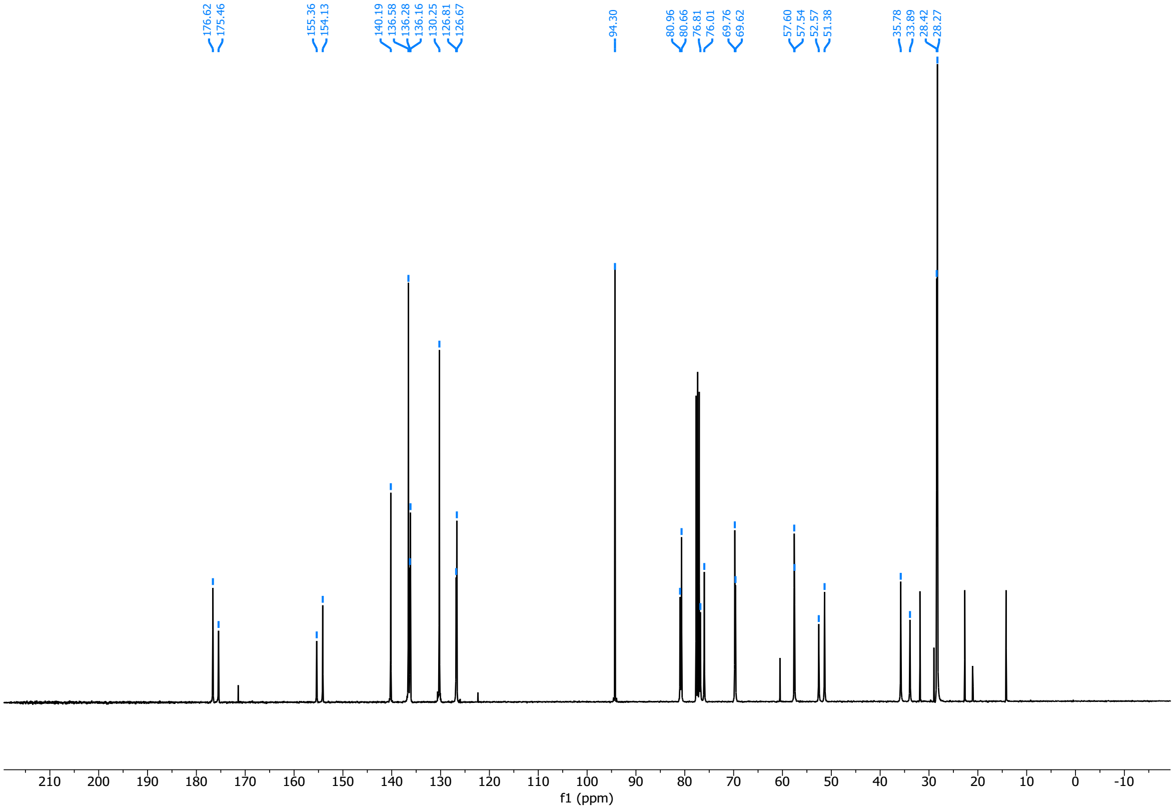

3032

(2*S*,4*S*)-1-(((9H-fluoren-9-yl)methoxy)carbonyl)-4-(4-iodophenoxy)pyrrolidine-2-carboxylic acid (**10ai**)

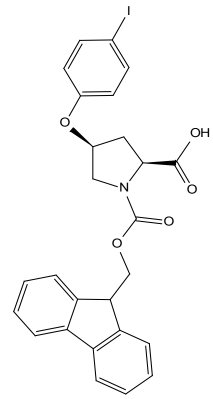

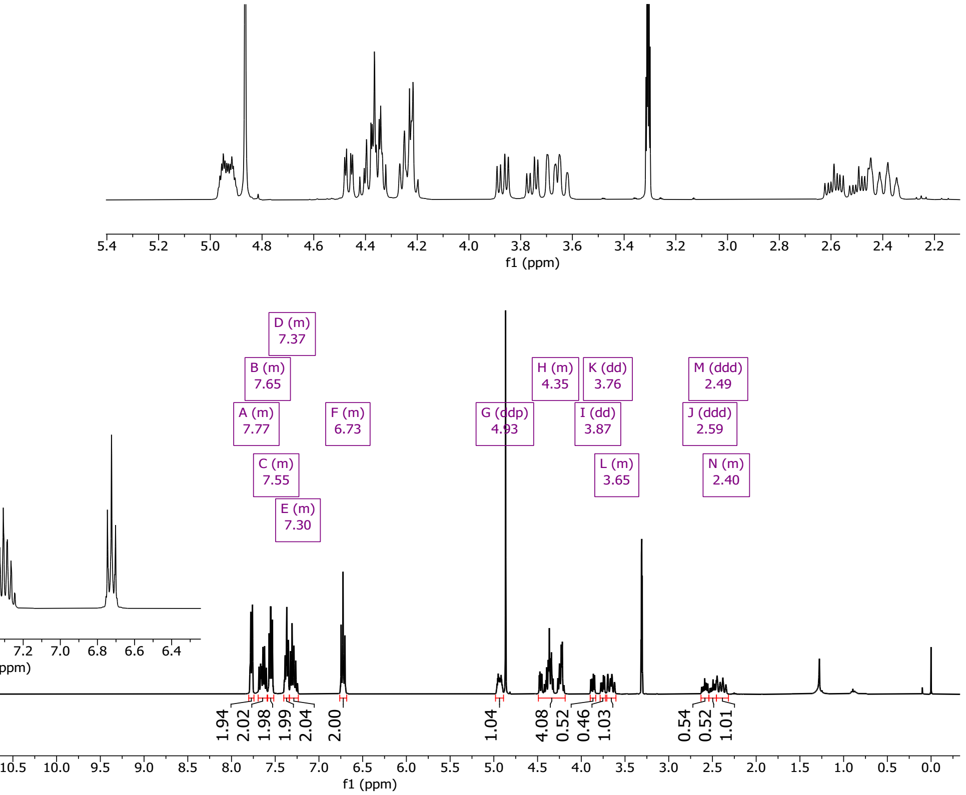

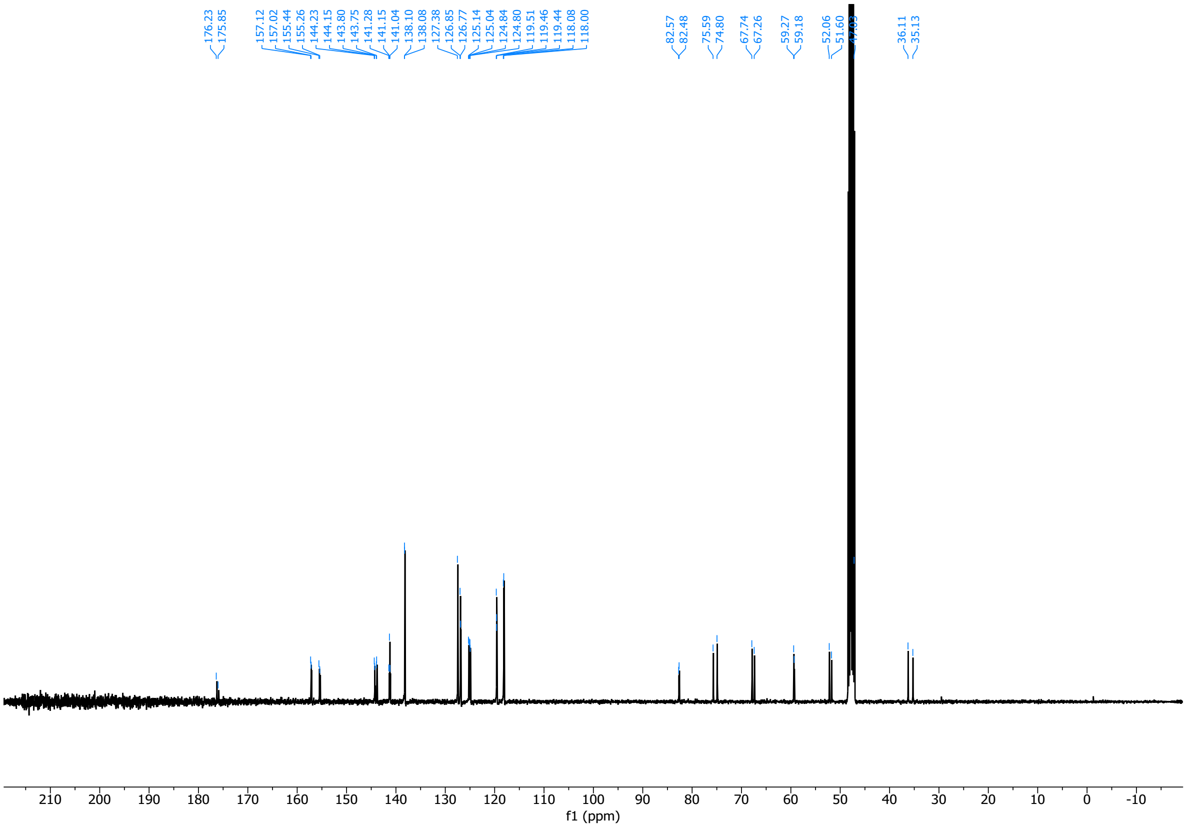

3036 (2*S*,4*R*)-1-(((9H-fluoren-9-yl)methoxy)carbonyl)-4-(4-iodophenoxy)pyrrolidine-2-carboxylic acid (**10bi**)

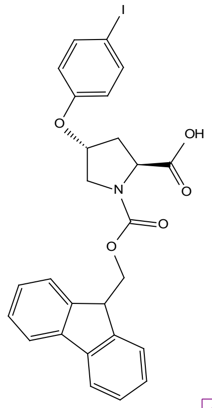

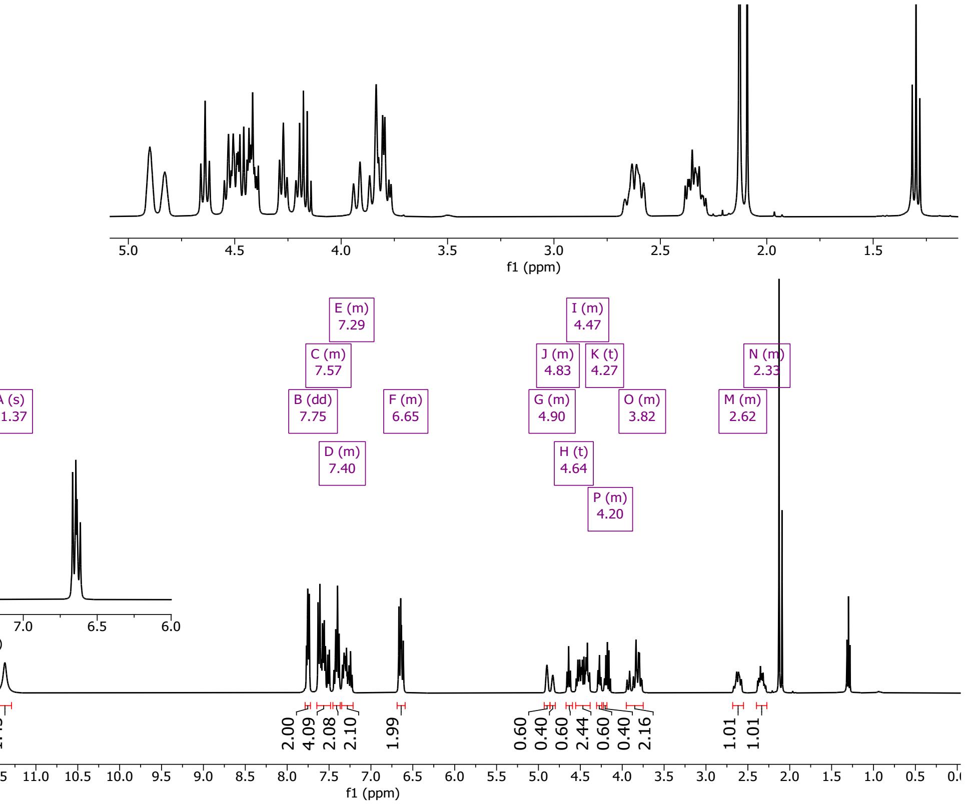

3037

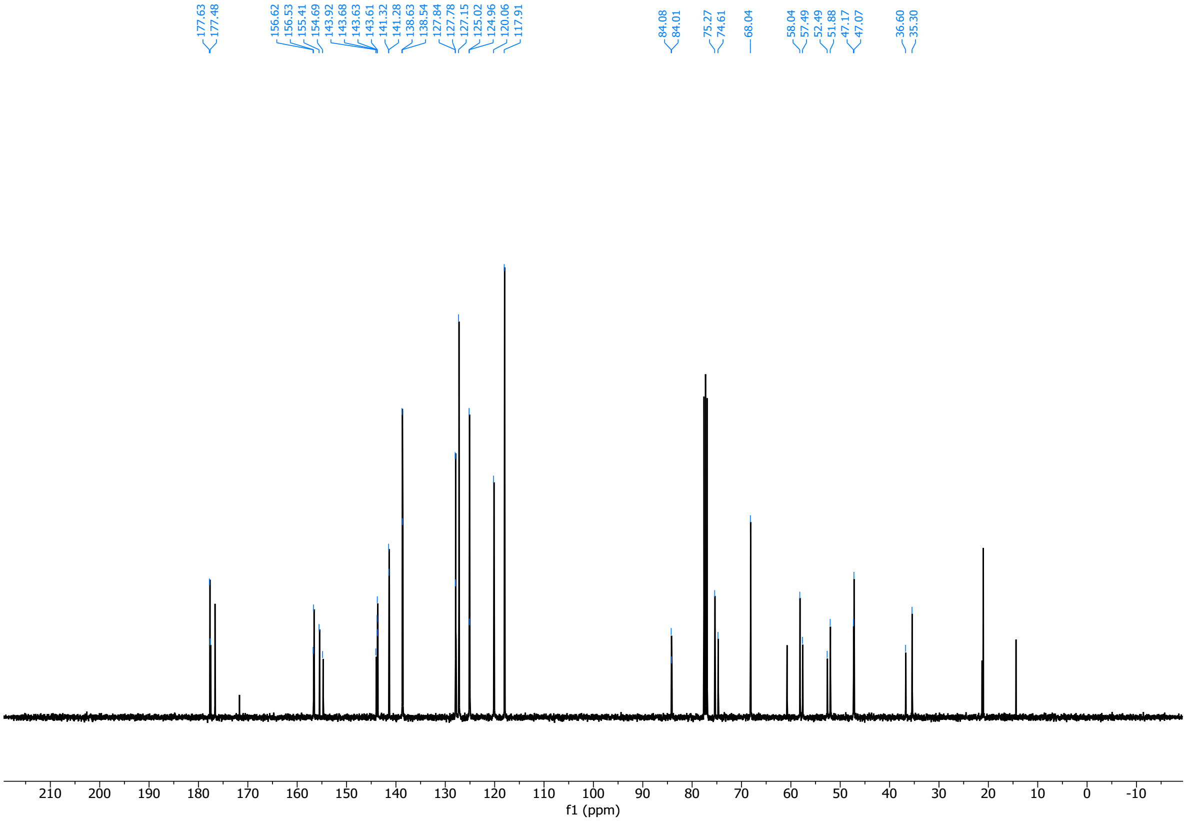

3038

(2*R*,4*R*)-1-(((9H-fluoren-9-yl)methoxy)carbonyl)-4-(4-iodophenoxy)pyrrolidine-2-carboxylic acid (**10ci**)

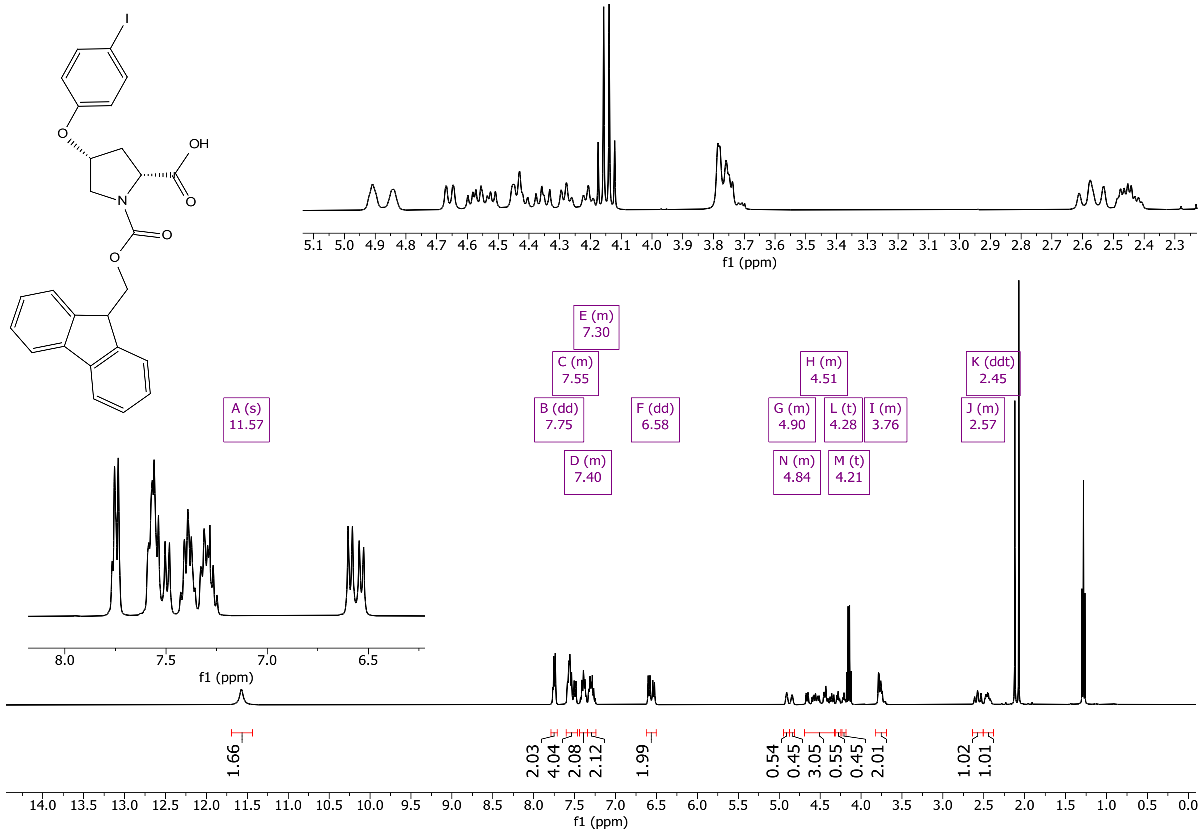

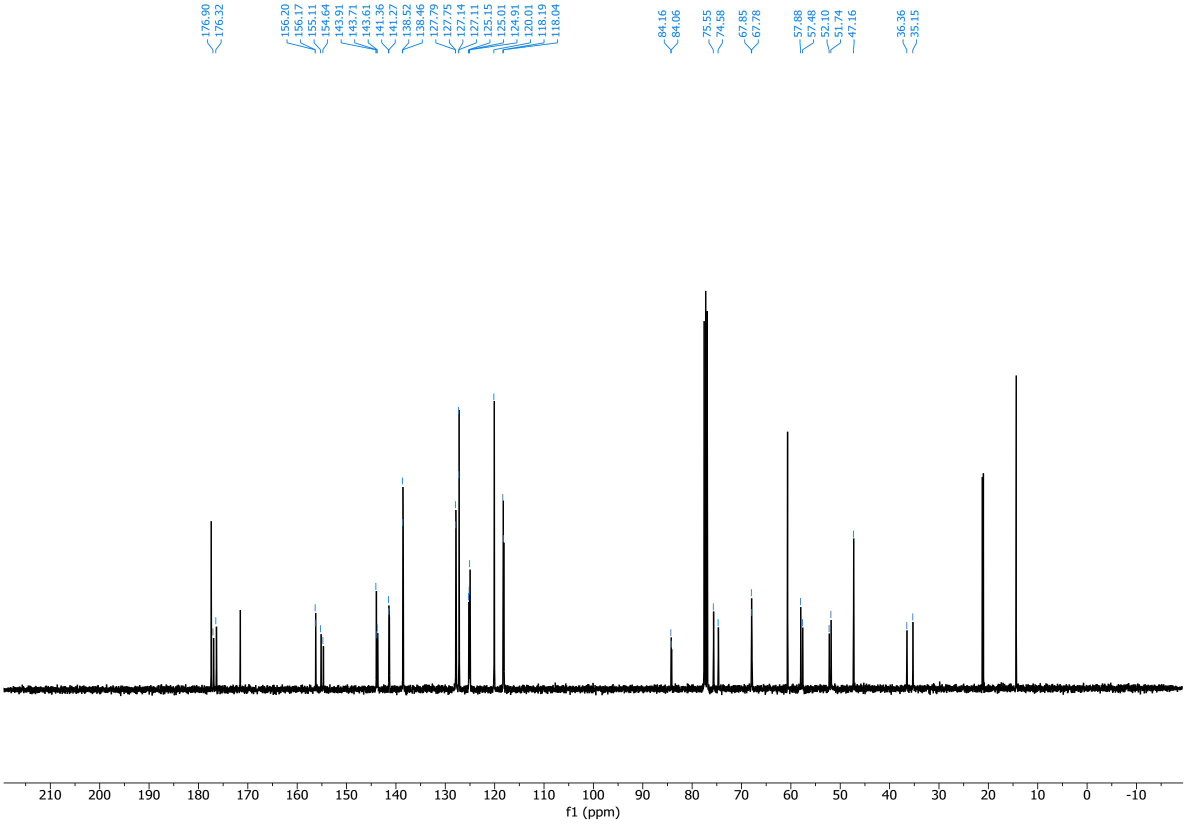

(2*R*,4*S*)-1-(((9H-fluoren-9-yl)methoxy)carbonyl)-4-(4-iodophenoxy)pyrrolidine-2-carboxylic acid (**10di**)

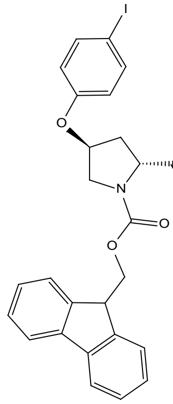

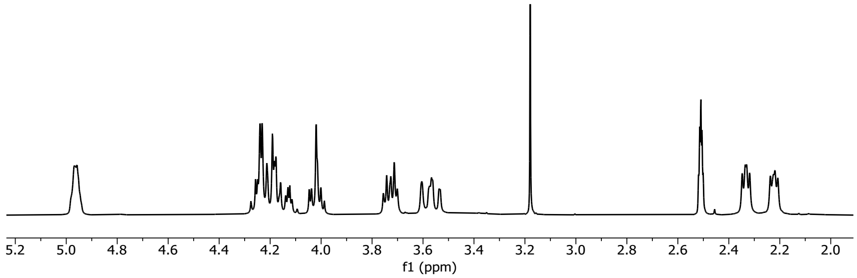

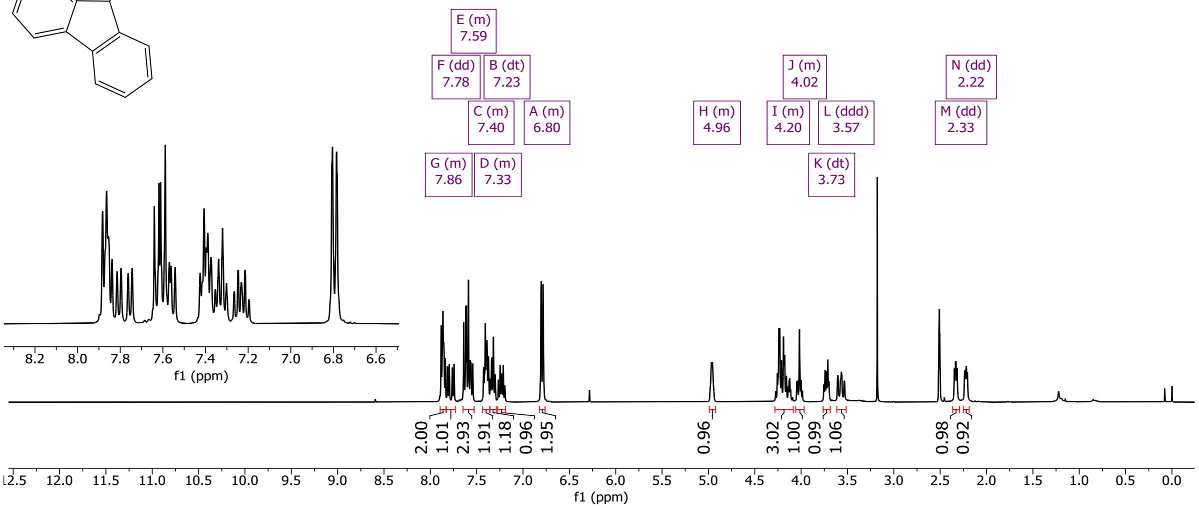

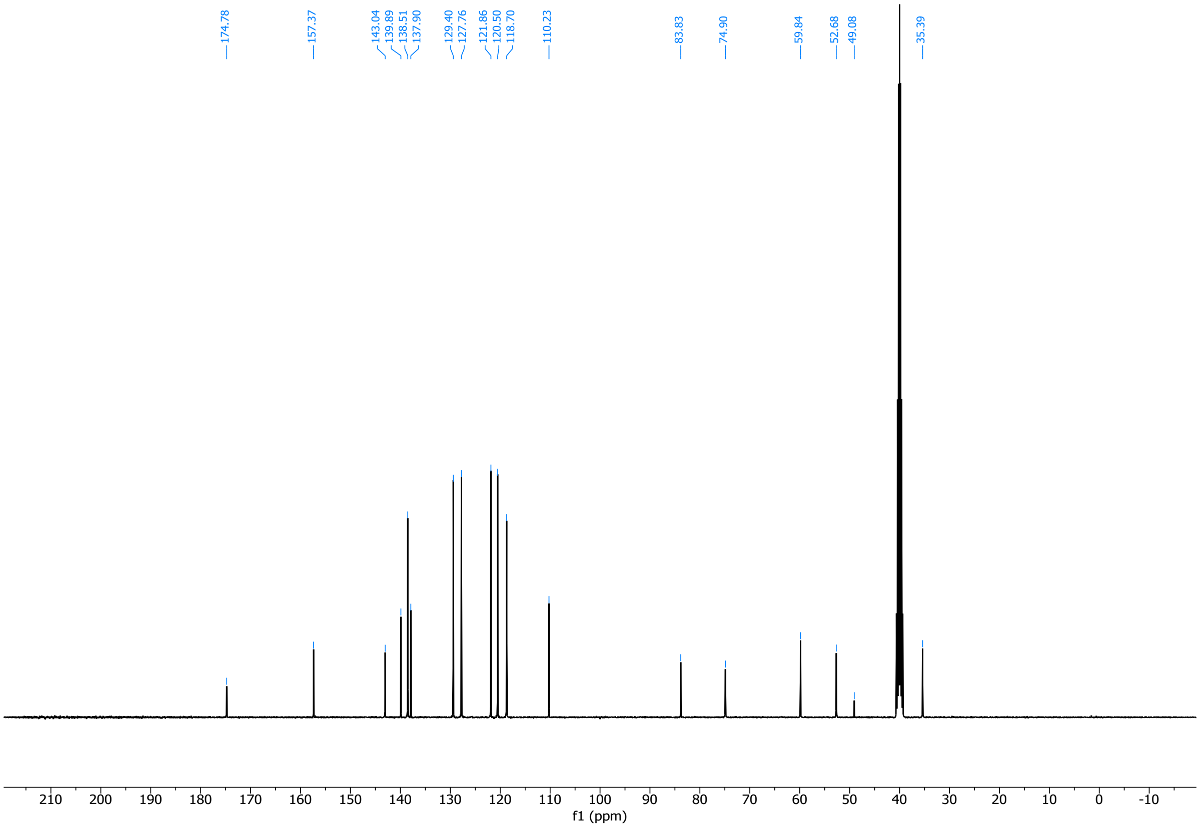

(2*R*,4*S*)-1-(((9H-fluoren-9-yl)methoxy)carbonyl)-4-(4-iodophenoxy)pyrrolidine-2-carboxylic acid (**10aii**)

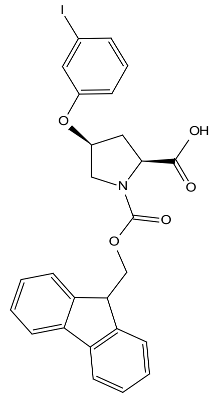

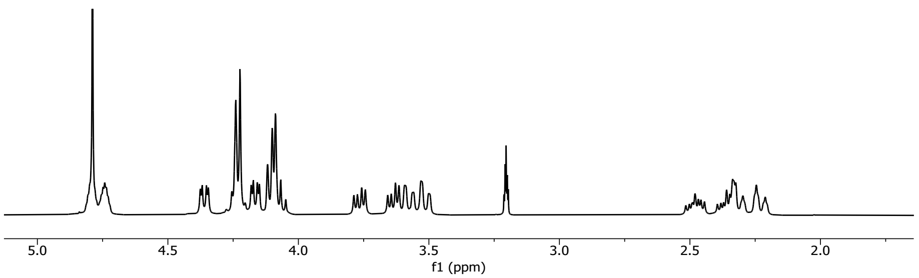

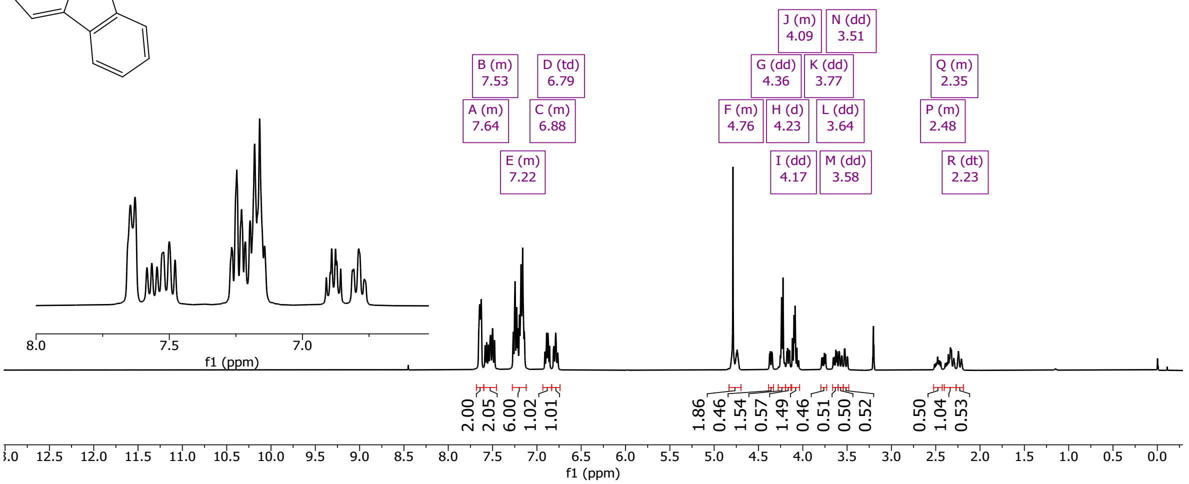

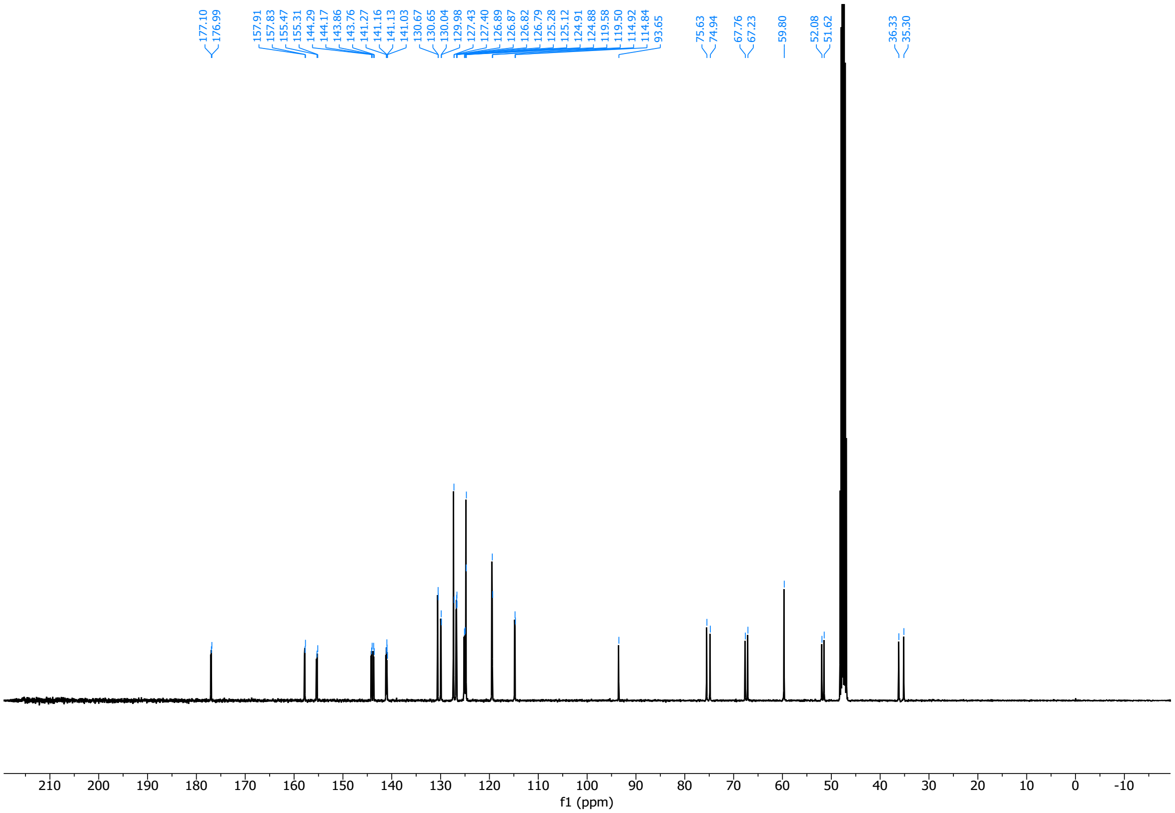

(2*S*,4*R*)-1-(((9H-fluoren-9-yl)methoxy)carbonyl)-4-(3-iodophenoxy)pyrrolidine-2-carboxylic acid (**10bii**)

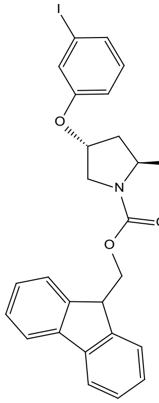

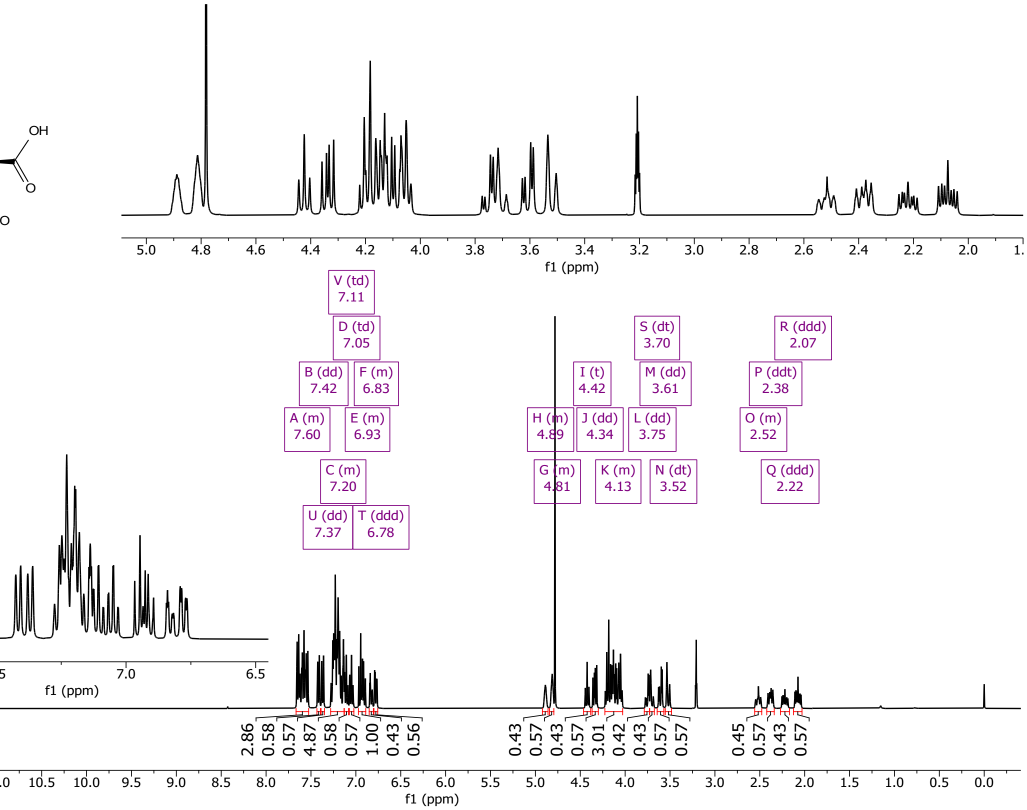

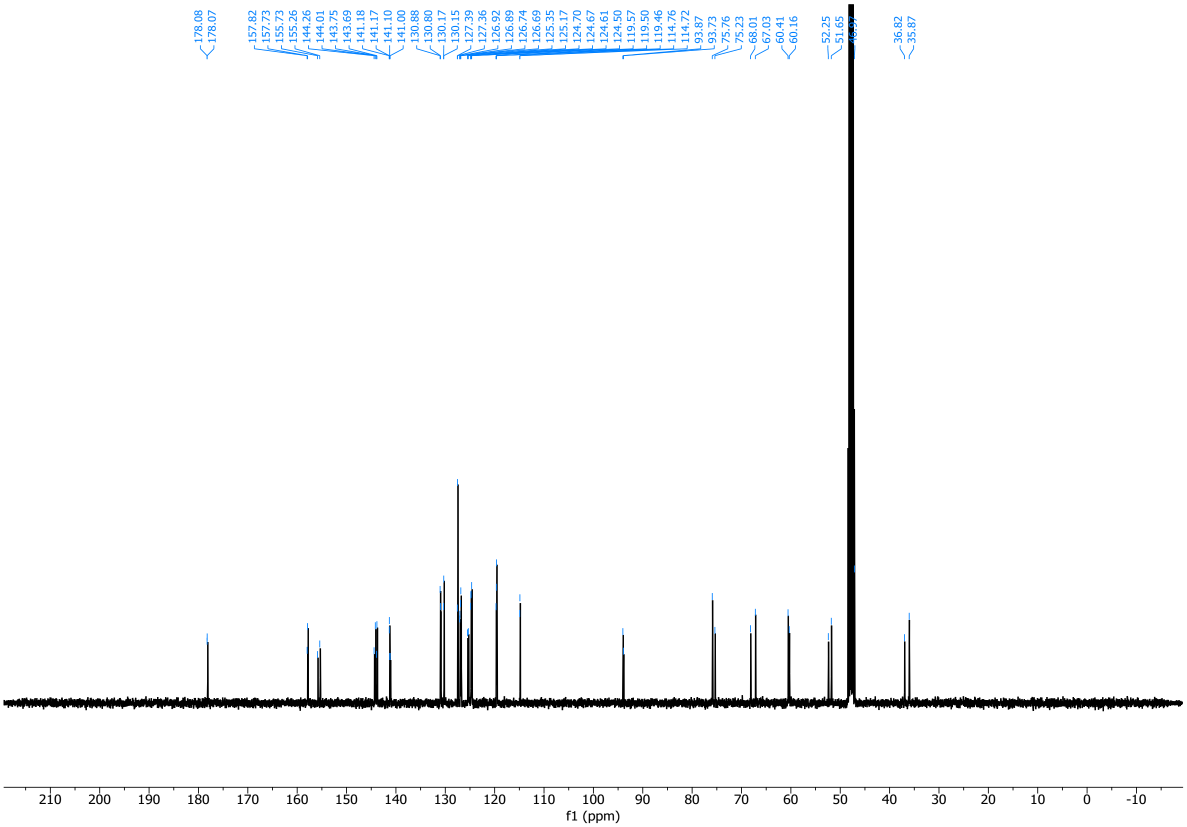

(2*R*,4*R*)-1-(((9H-fluoren-9-yl)methoxy)carbonyl)-4-(3-iodophenoxy)pyrrolidine-2-carboxylic acid (**10cii**)

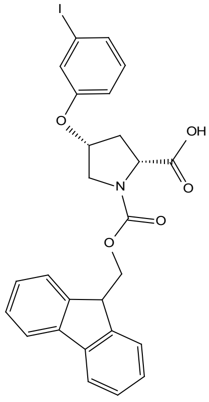

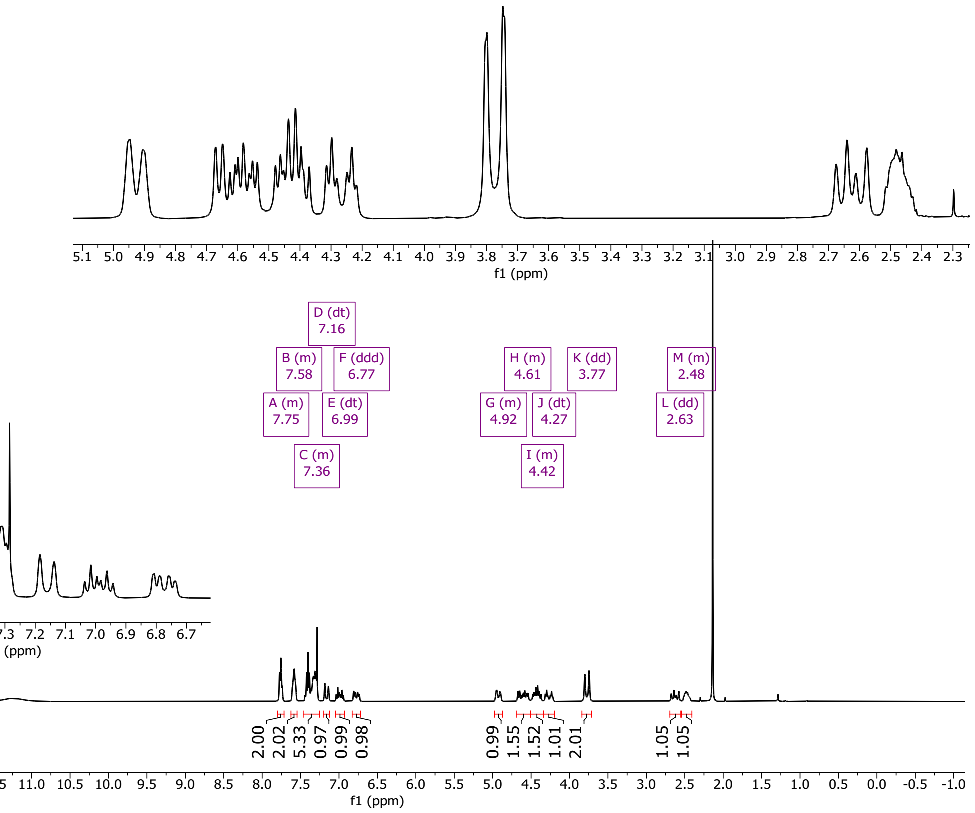

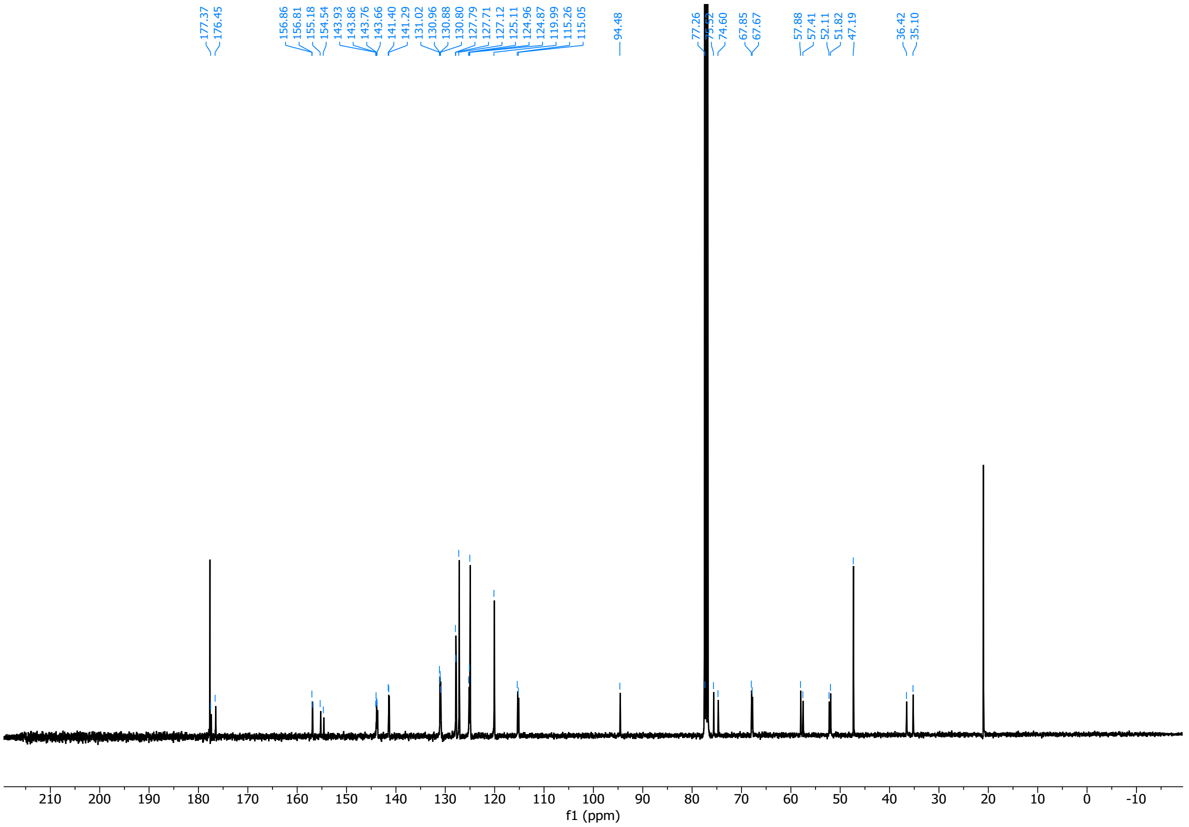

3054 (2*R*,4*S*)-1-(((9H-fluoren-9-yl)methoxy)carbonyl)-4-(3-iodophenoxy)pyrrolidine-2-carboxylic acid (**10dii**)

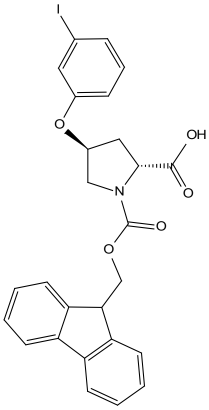

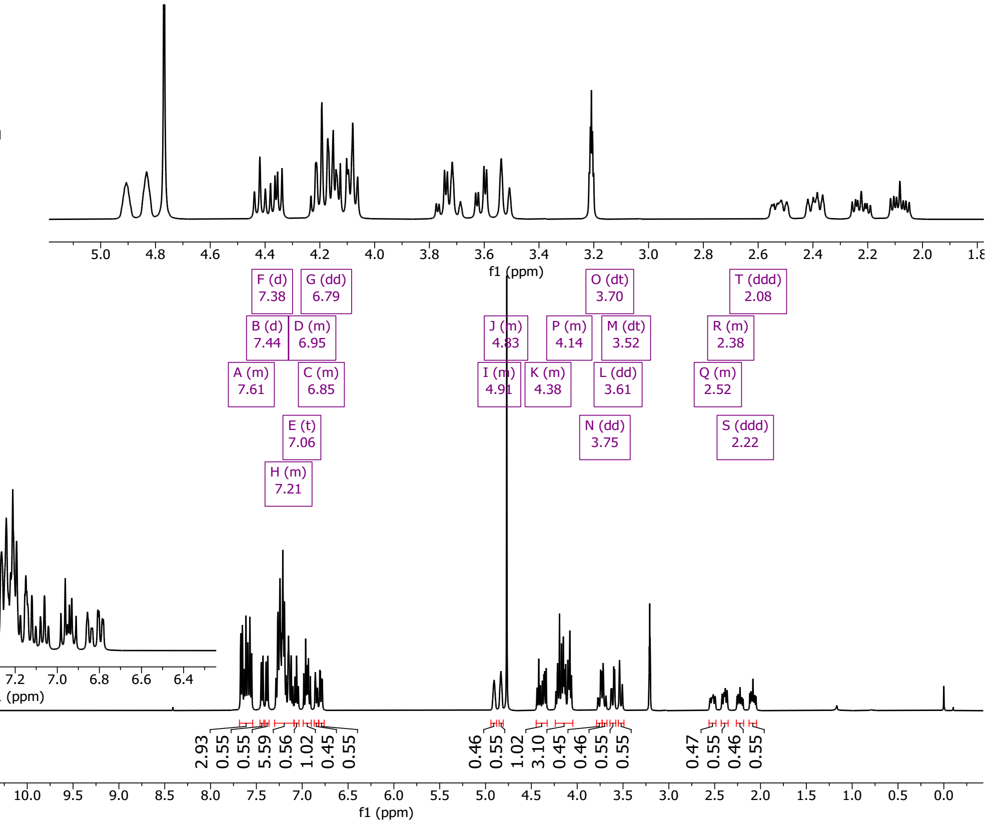

3055

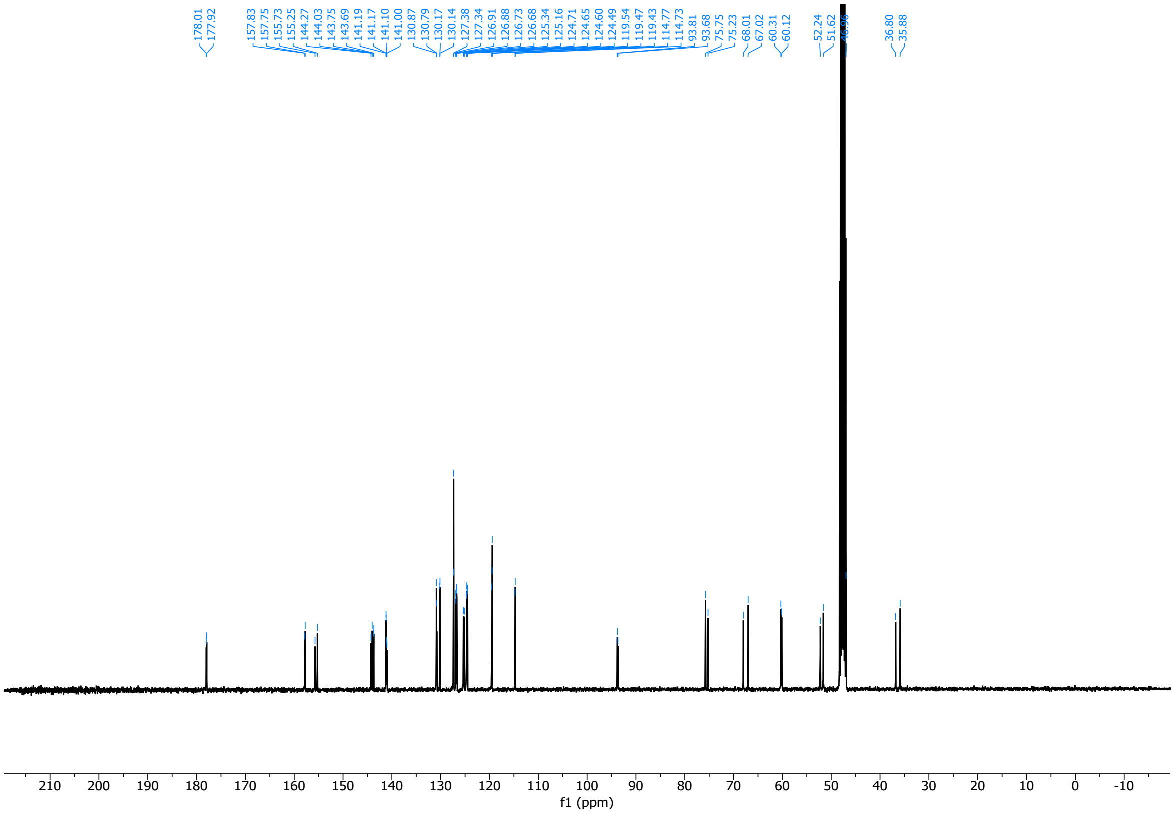

3056

3057 (2*S*,4*S*)-1-(((9H-fluoren-9-yl)methoxy)carbonyl)-4-((4-iodobenzyl)oxy)pyrrolidine-2-carboxylic acid (**14ai**)

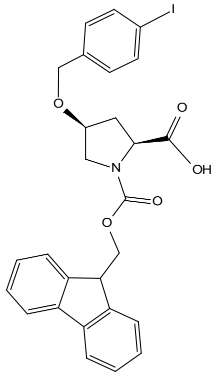

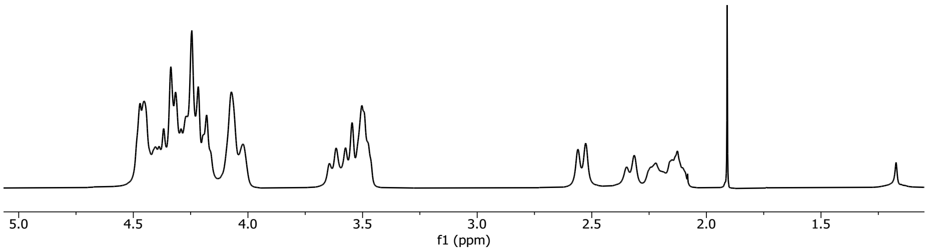

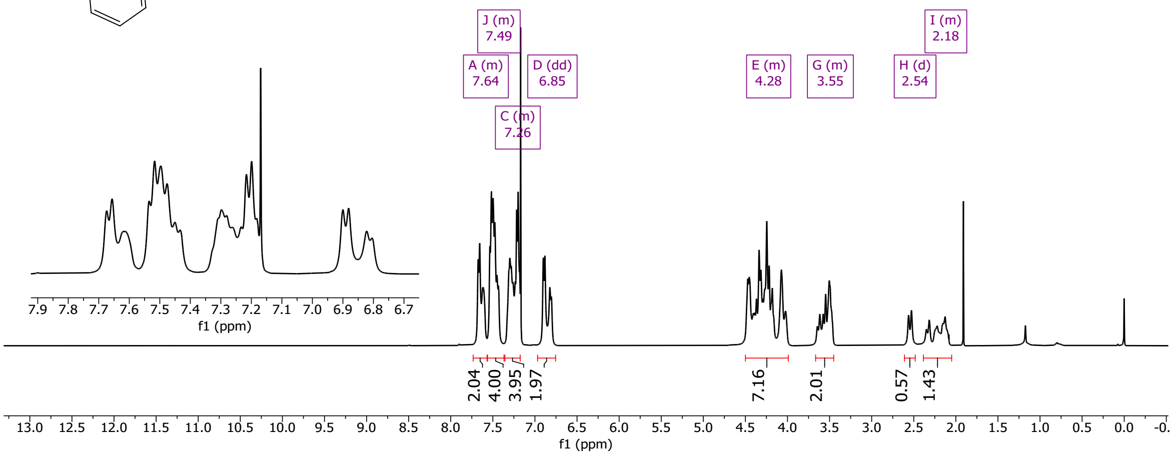

3058

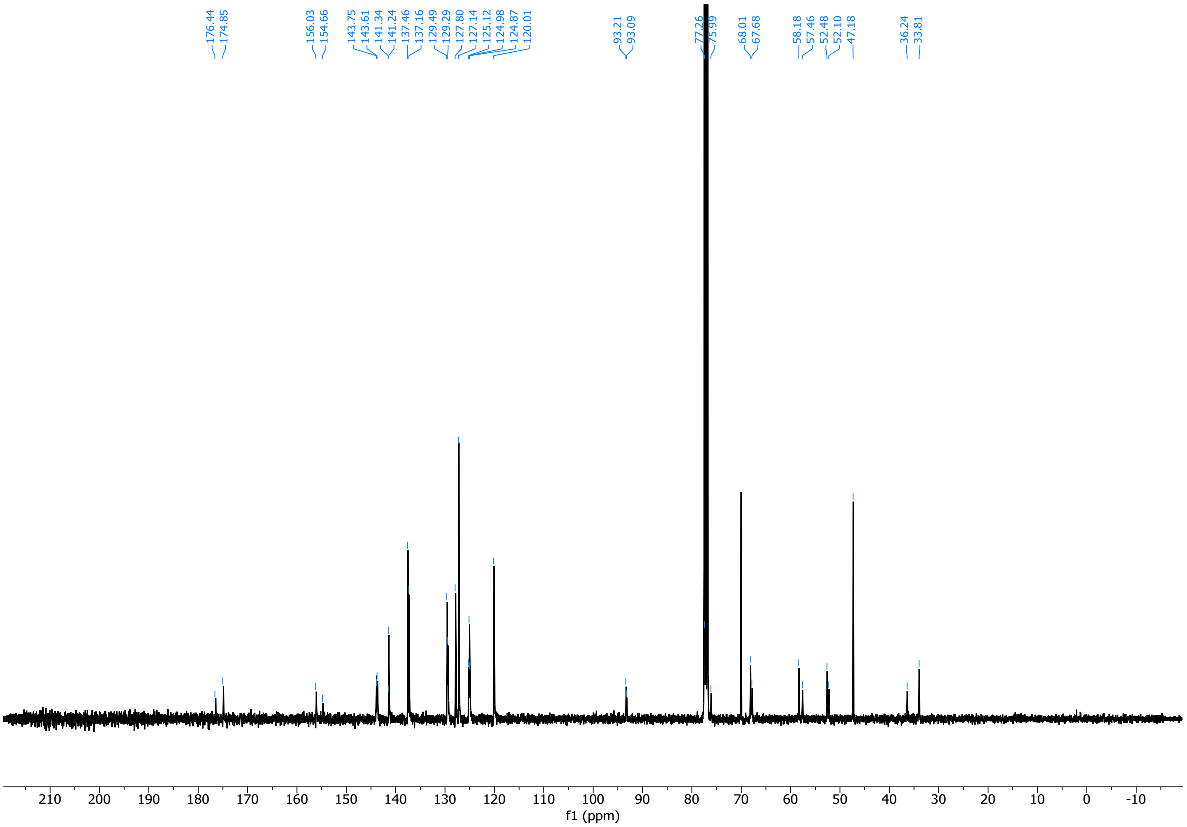

3059

(2*S*,4*R*)-1-(((9H-fluoren-9-yl)methoxy)carbonyl)-4-((4-iodobenzyl)oxy)pyrrolidine-2-carboxylic acid (**14bi**)

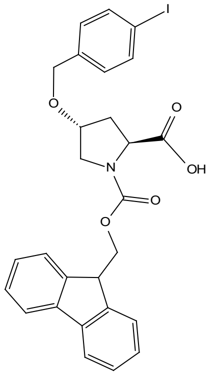

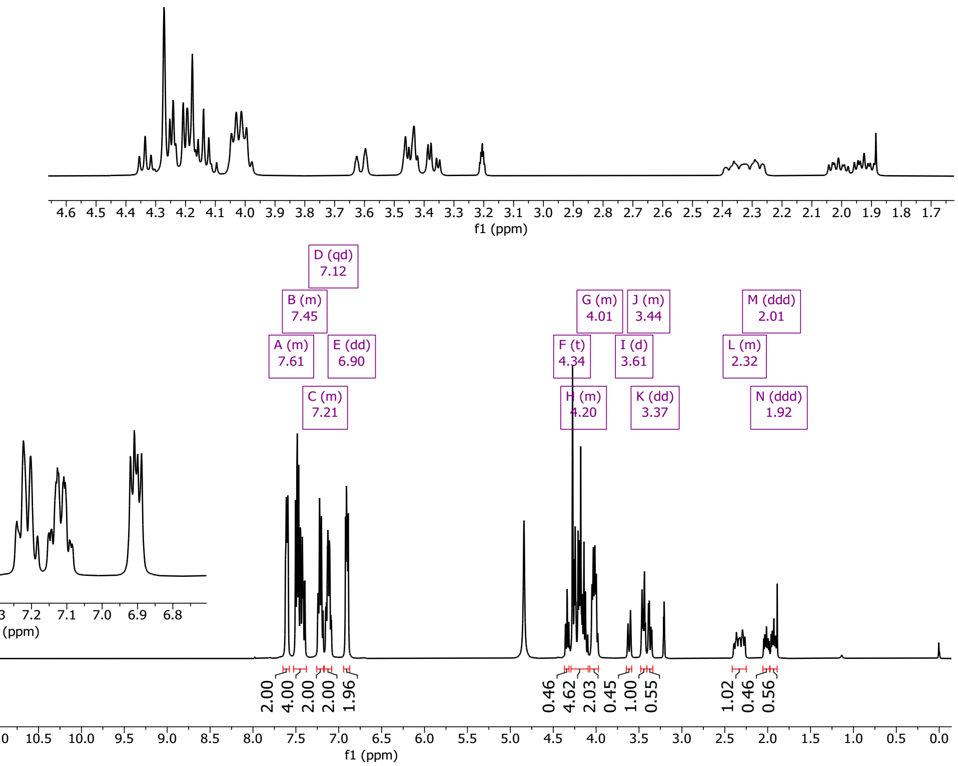

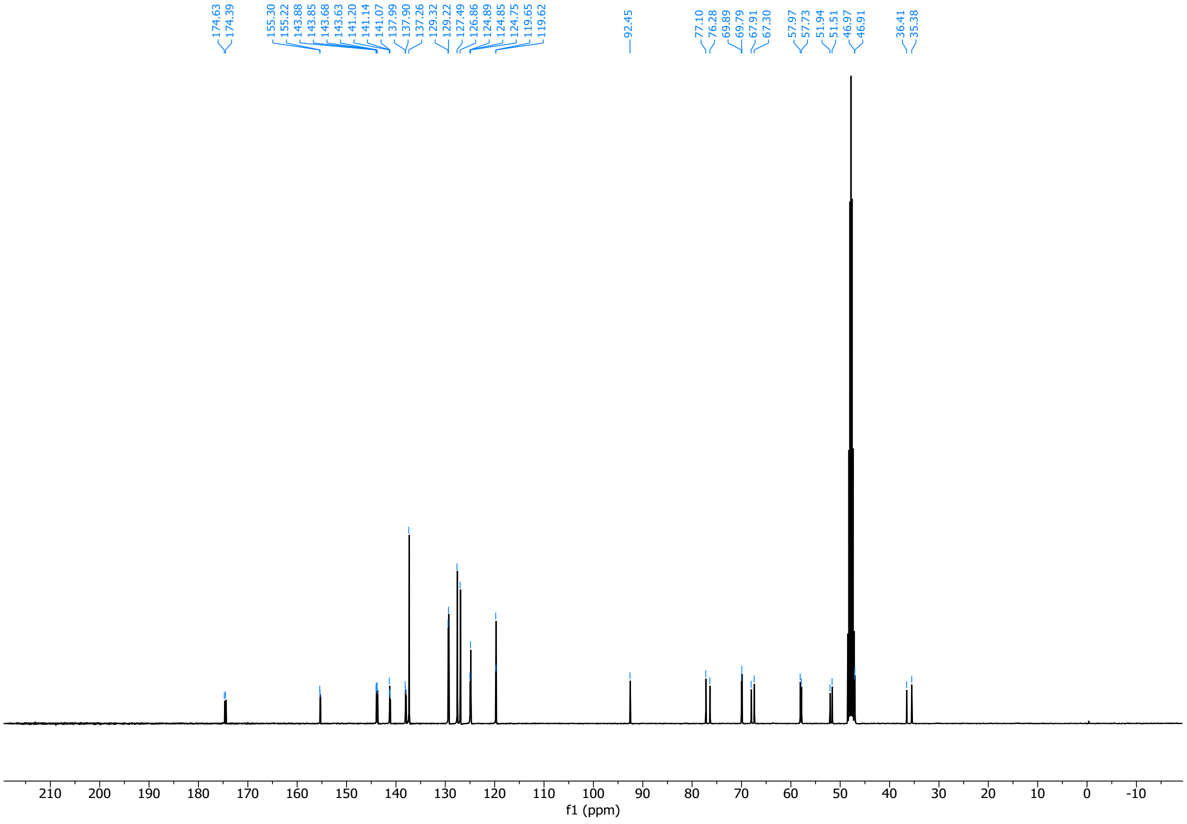

3063 (2*R*,4*R*)-1-(((9H-fluoren-9-yl)methoxy)carbonyl)-4-((4-iodobenzyl)oxy)pyrrolidine-2-carboxylic acid (**14ci**)

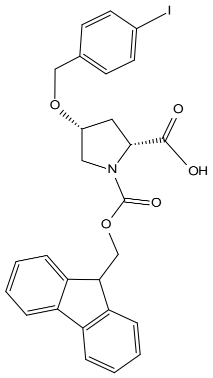

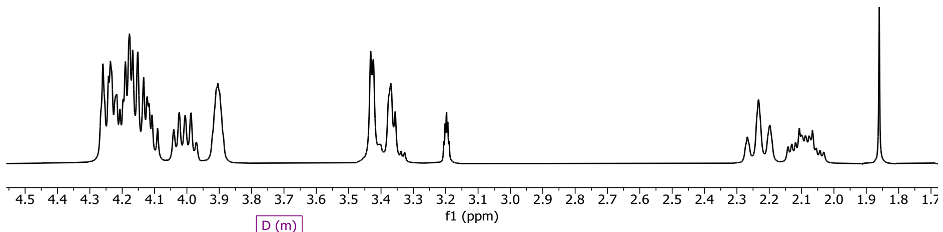

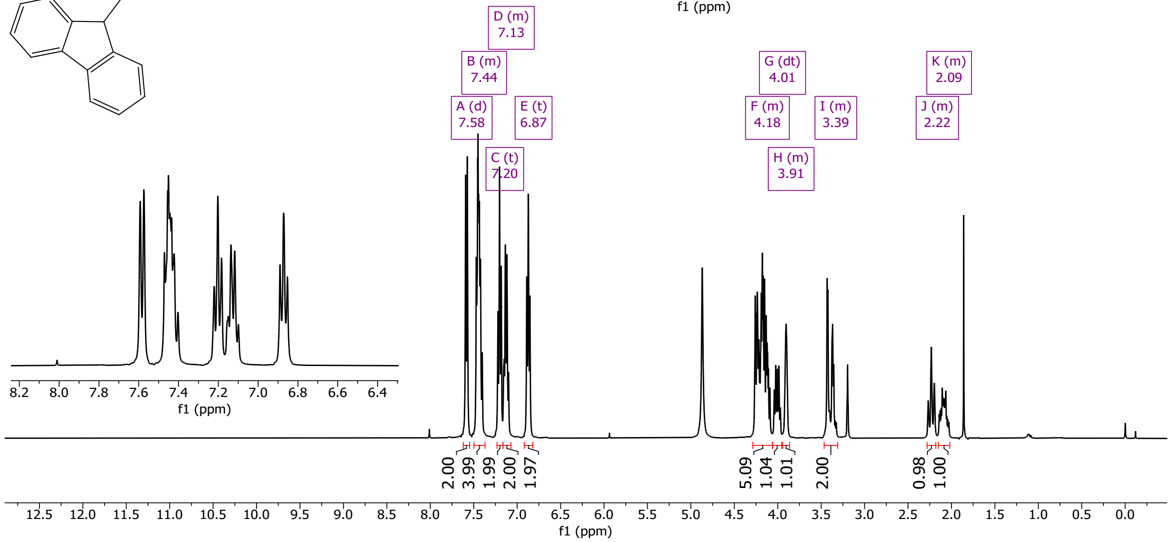

3064

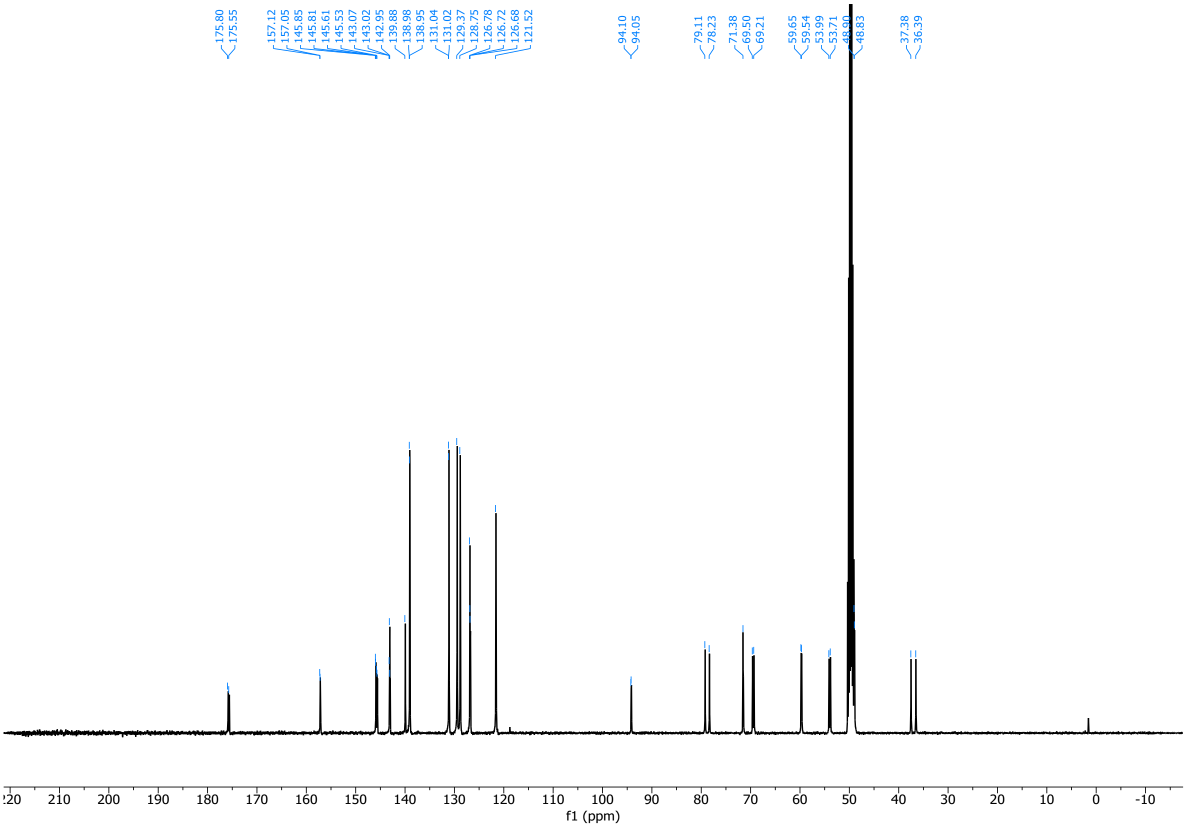

3065

3066 (2*R*,4*S*)-1-(((9H-fluoren-9-yl)methoxy)carbonyl)-4-((4-iodobenzyl)oxy)pyrrolidine-2-carboxylic acid (**14di**)

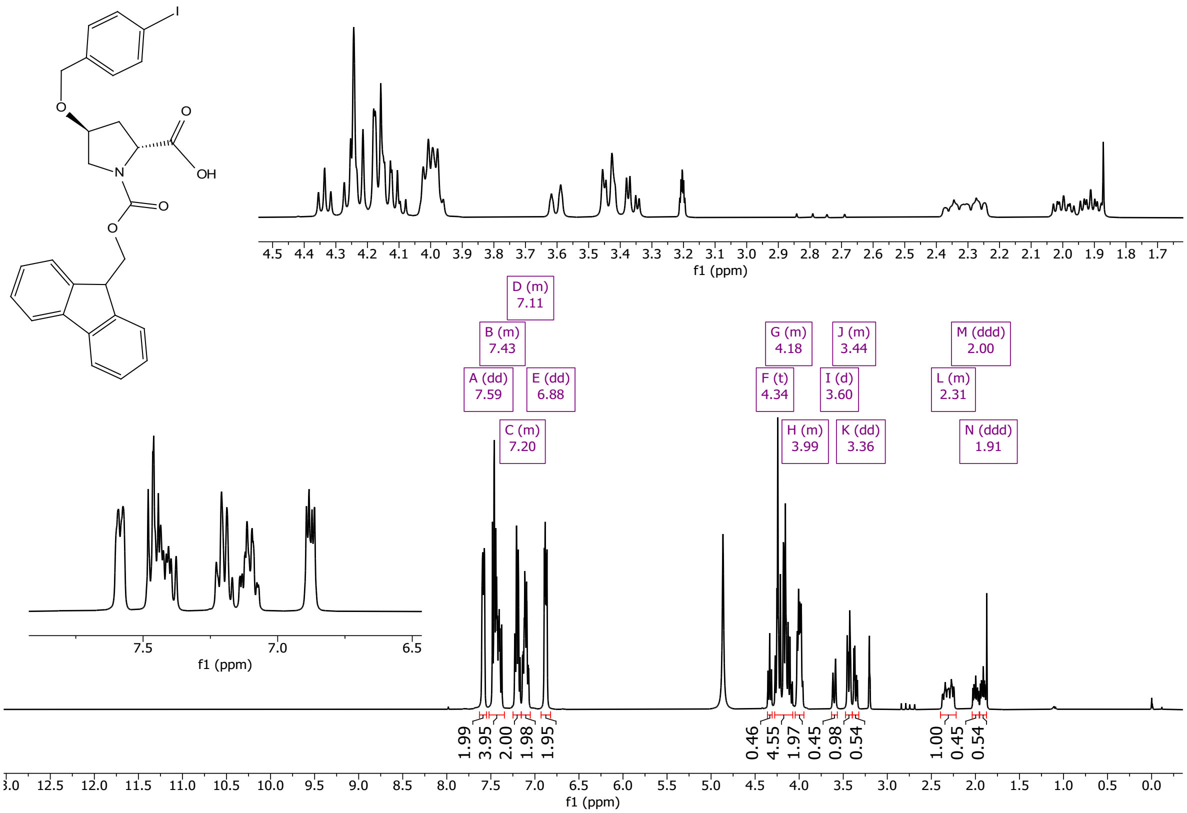

3067

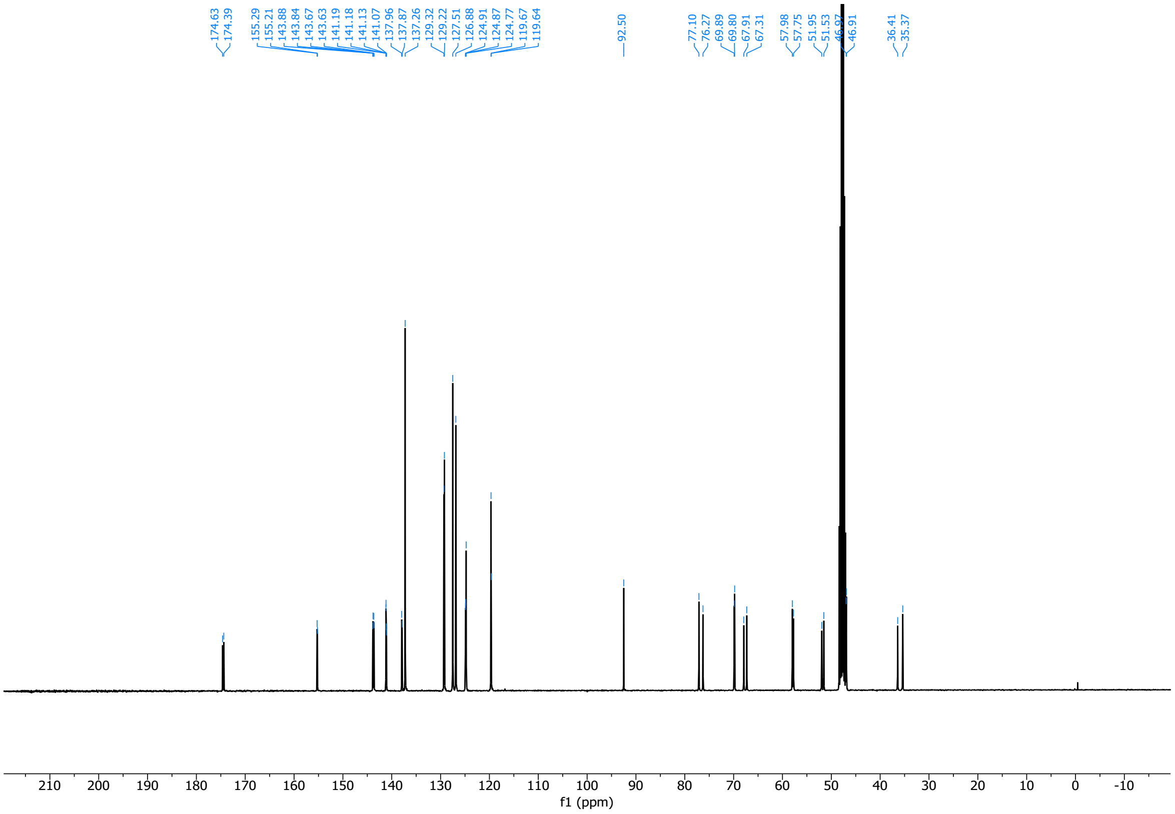

3068

(2*S*,4*S*)-1-(((9H-fluoren-9-yl)methoxy)carbonyl)-4-((3-iodobenzyl)oxy)pyrrolidine-2-carboxylic acid (**14aii**)

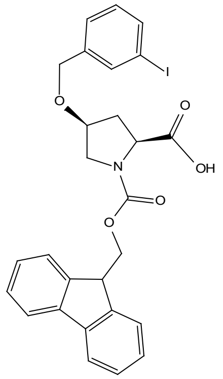

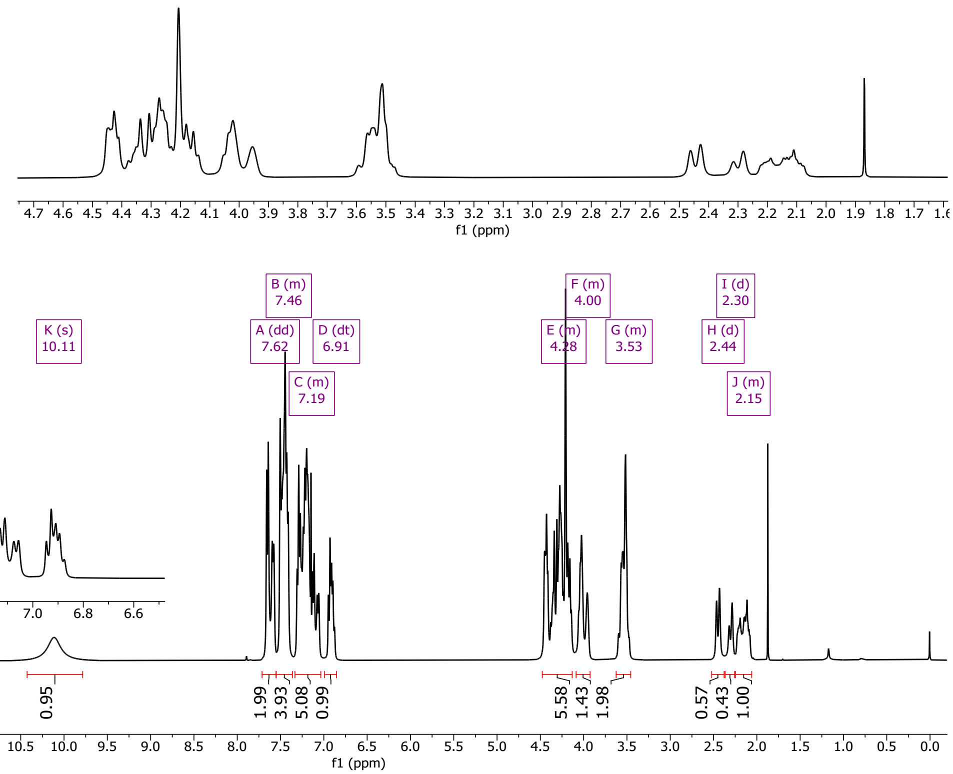

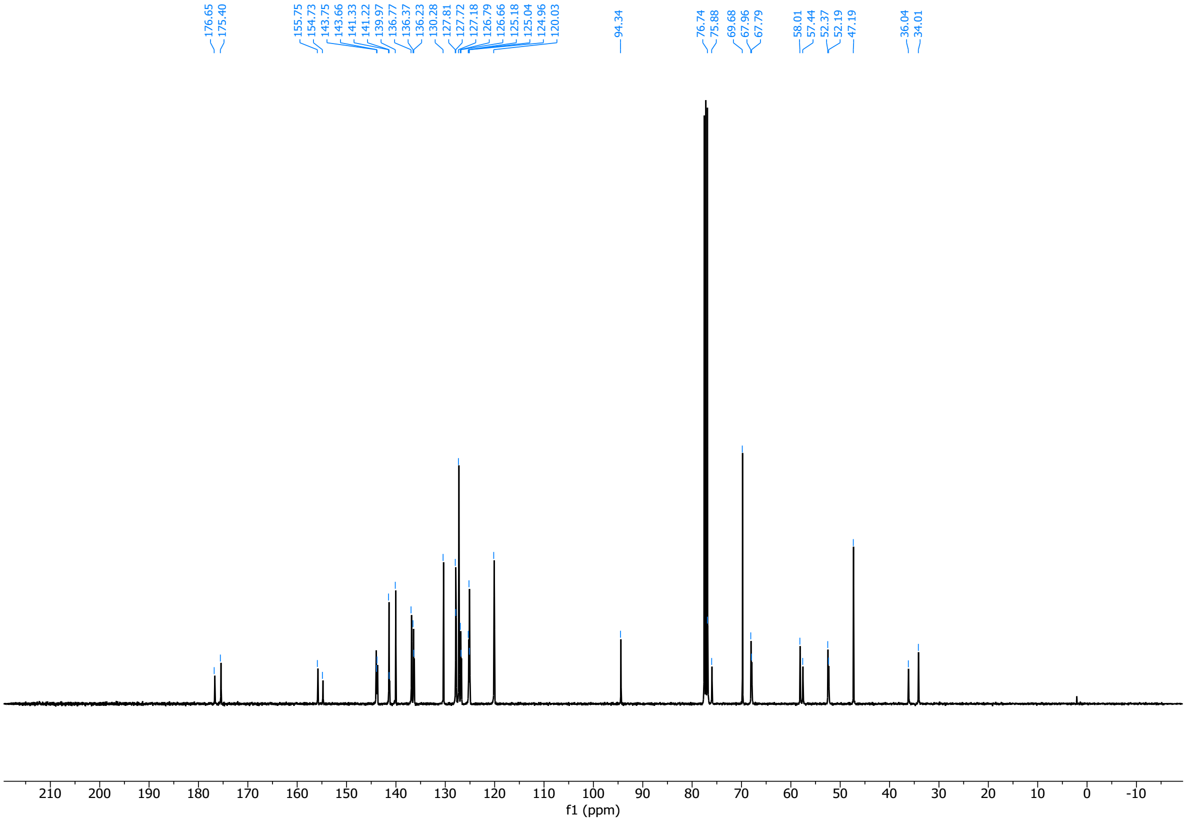

(2*S*,4*R*)-1-(((9H-fluoren-9-yl)methoxy)carbonyl)-4-((3-iodobenzyl)oxy)pyrrolidine-2-carboxylic acid (**14bii**)

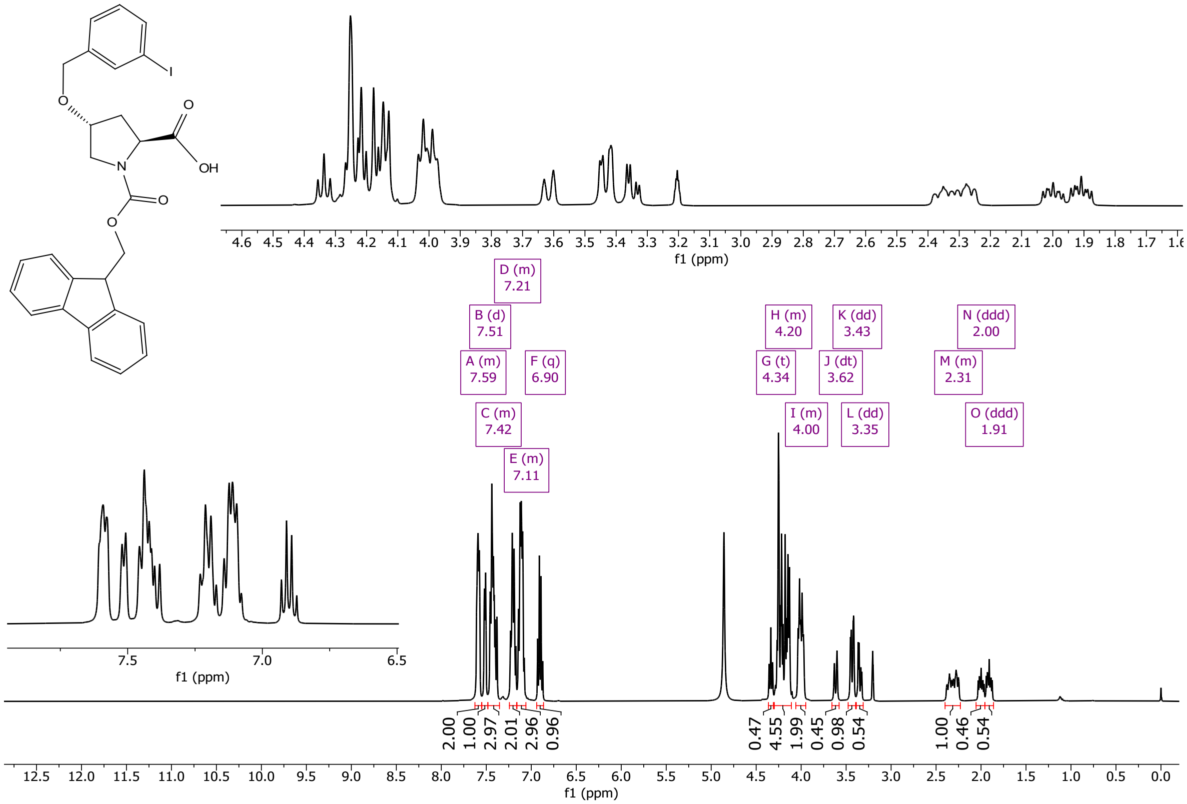

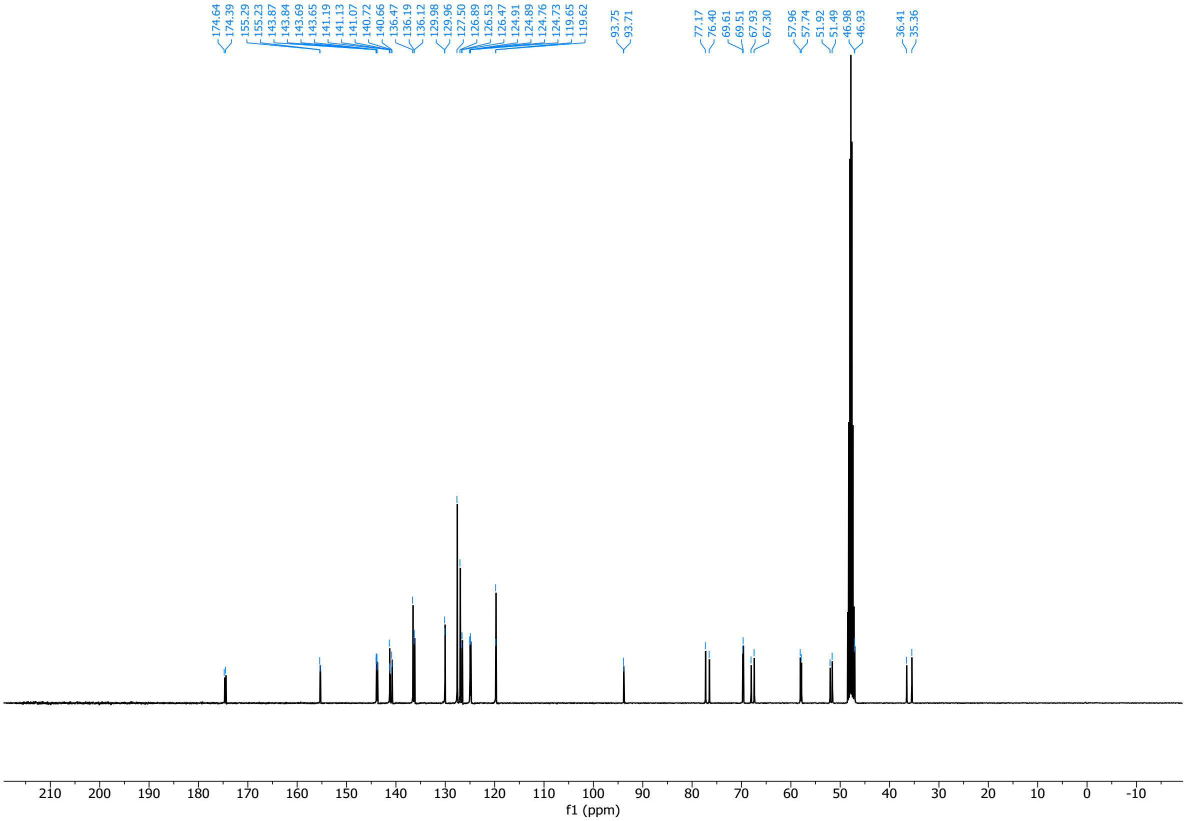

(2*R*,4*R*)-1-(((9H-fluoren-9-yl)methoxy)carbonyl)-4-((3-iodobenzyl)oxy)pyrrolidine-2-carboxylic acid (**14cii**)

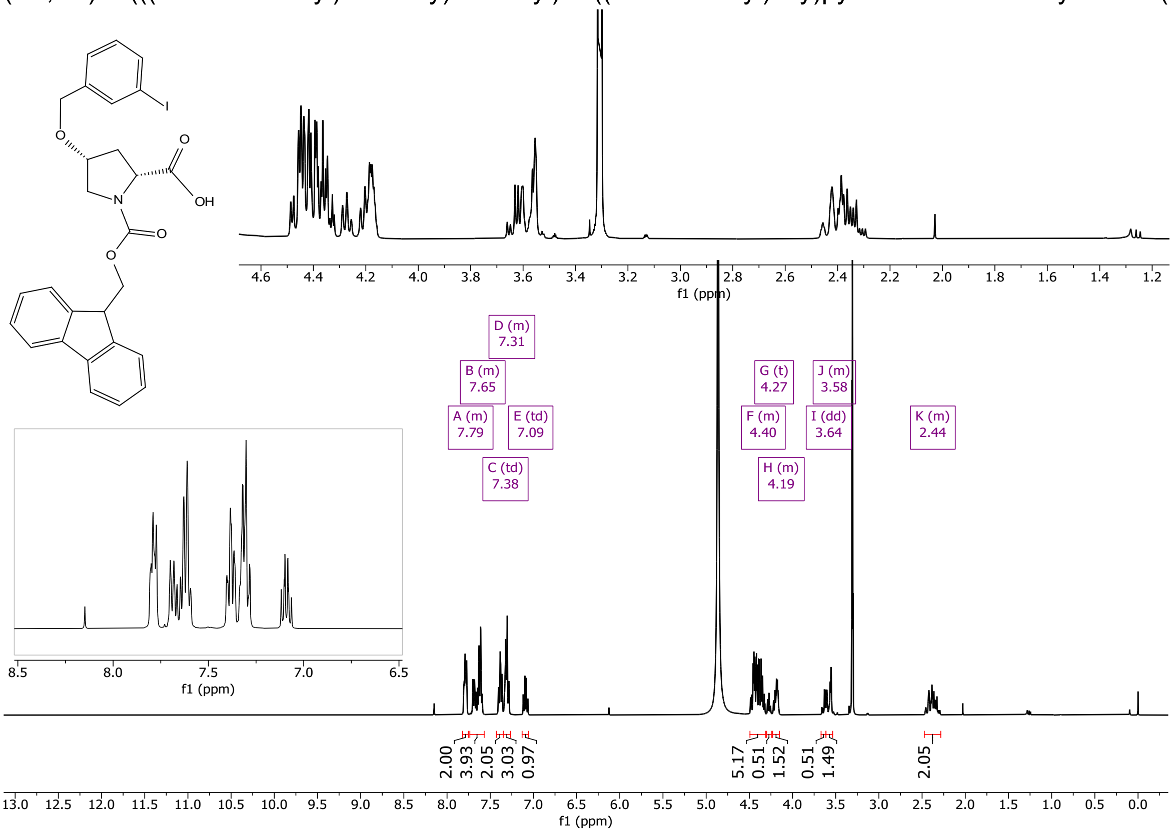

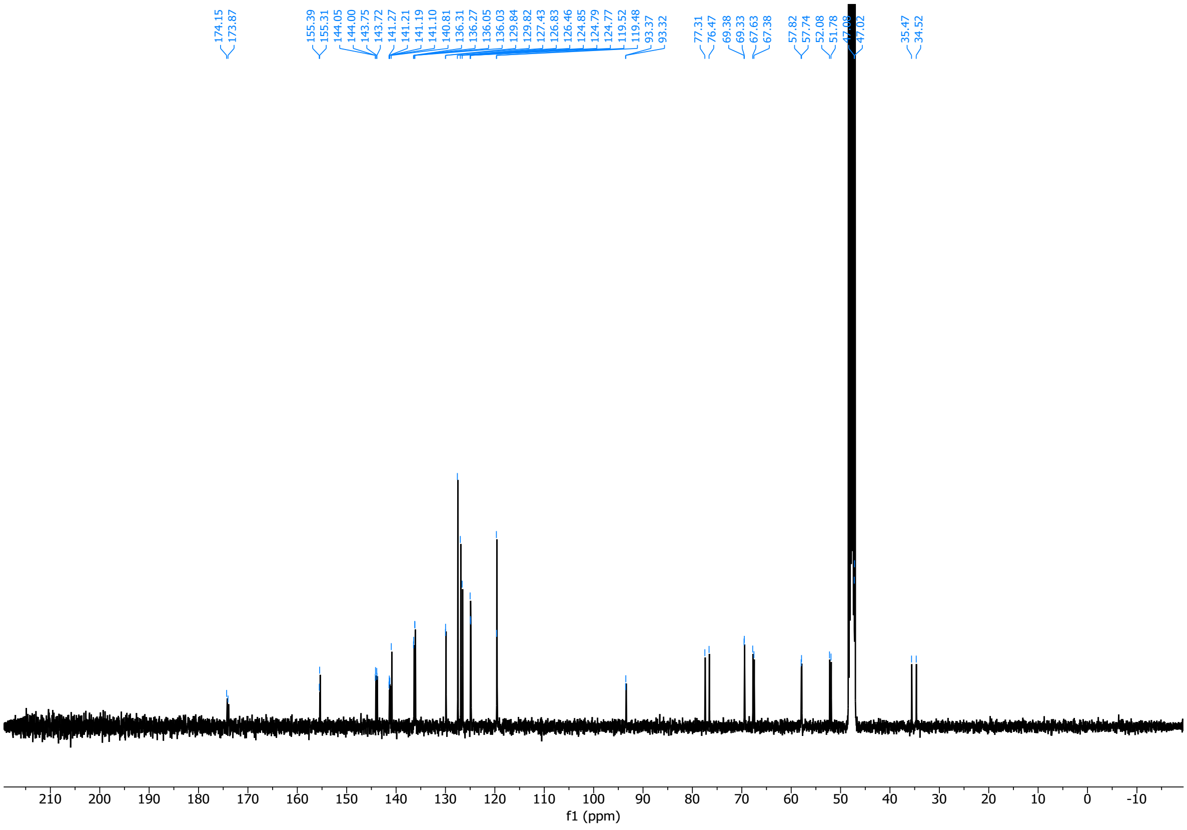

3078 (2*R*,4*S*)-1-(((9H-fluoren-9-yl)methoxy)carbonyl)-4-((3-iodobenzyl)oxy)pyrrolidine-2-carboxylic acid (**14dii**)

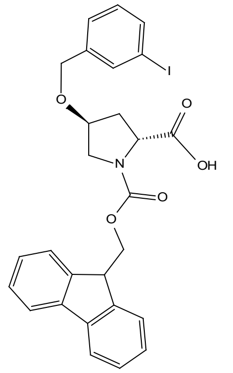

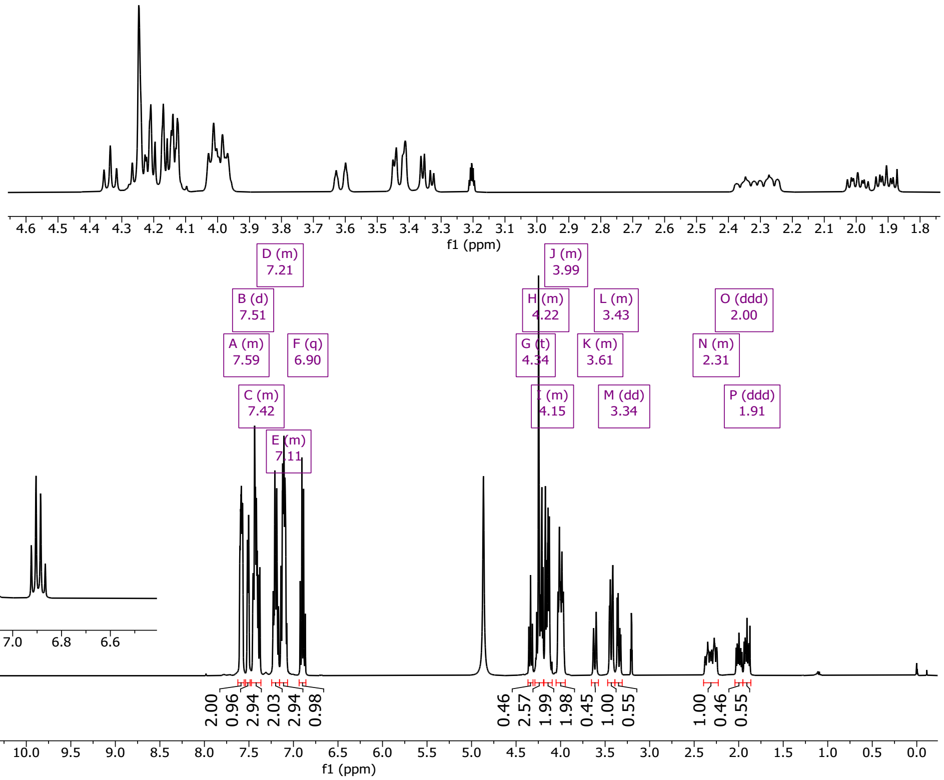

3079

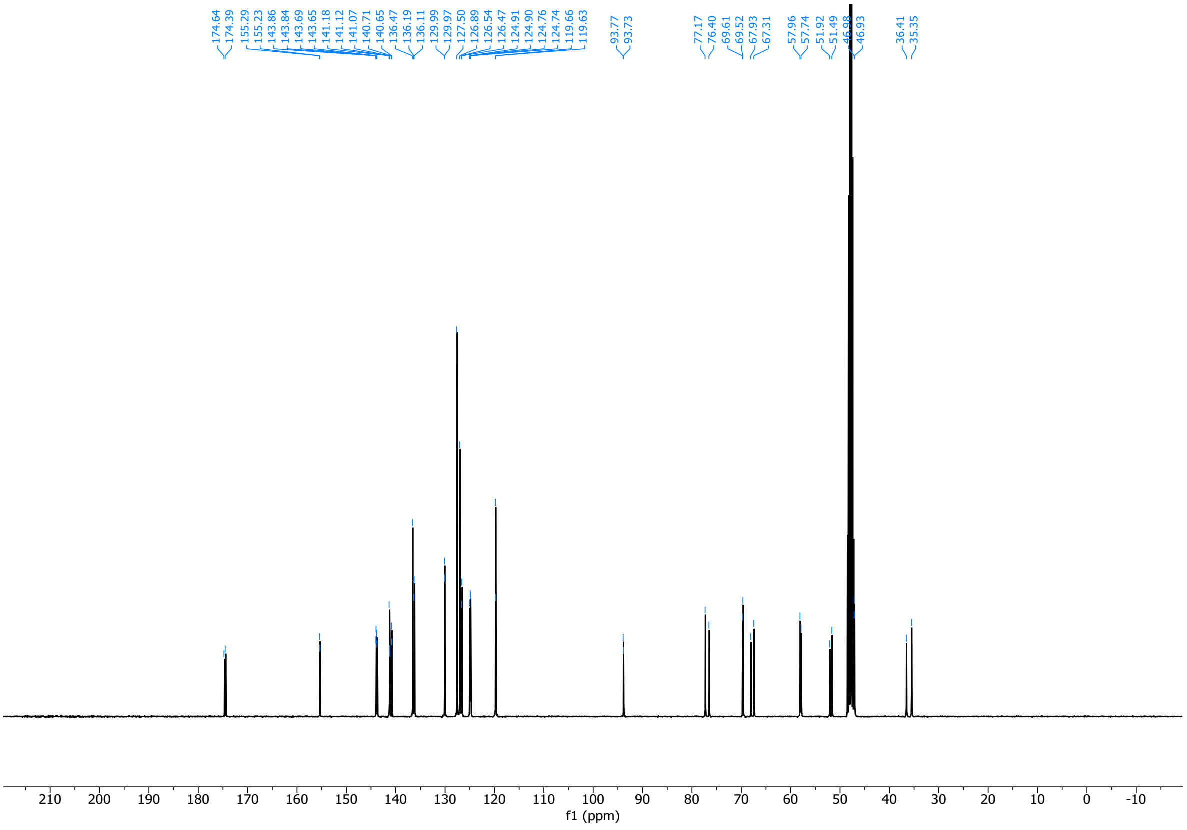

3080

#### 7.3. Spiro-azetdine skeletons

(*R*)-4-benzyl-3-((3*S*,4*R*)-1-benzyl-4-(4-iodophenyl)pyrrolidine-3-carbonyl)oxazolidin-2-one (**15**)

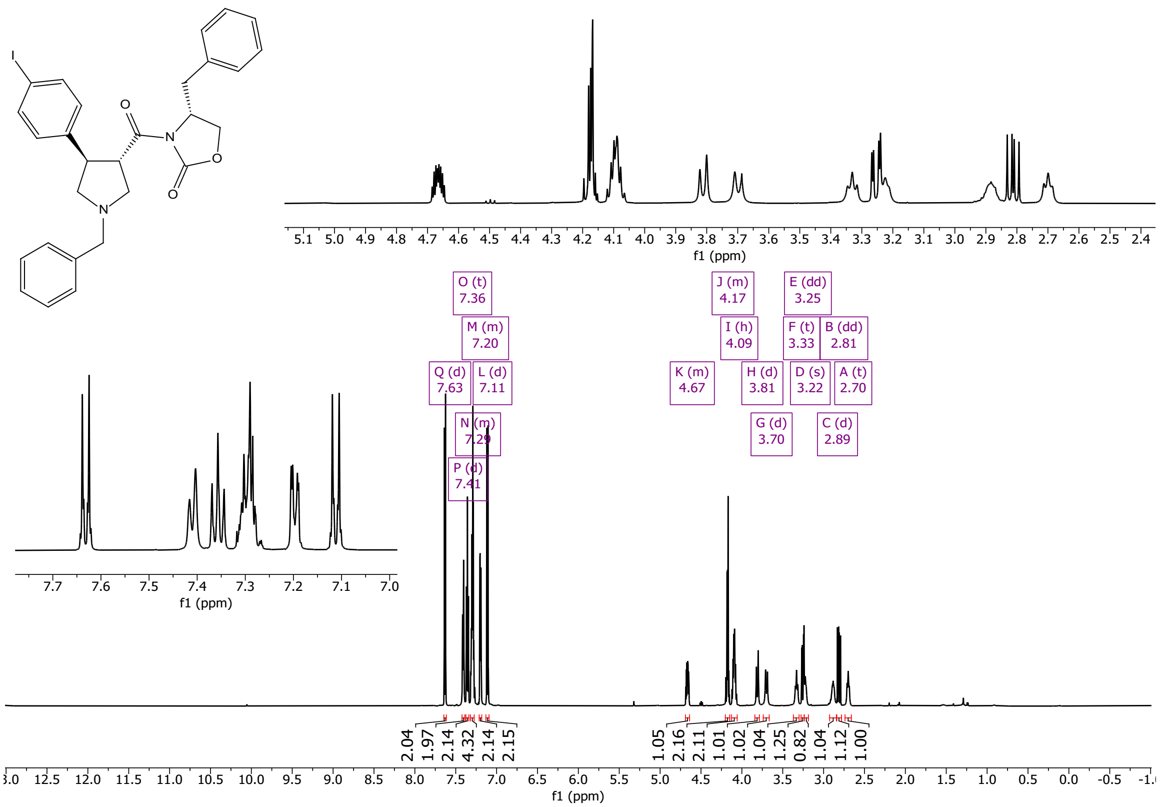

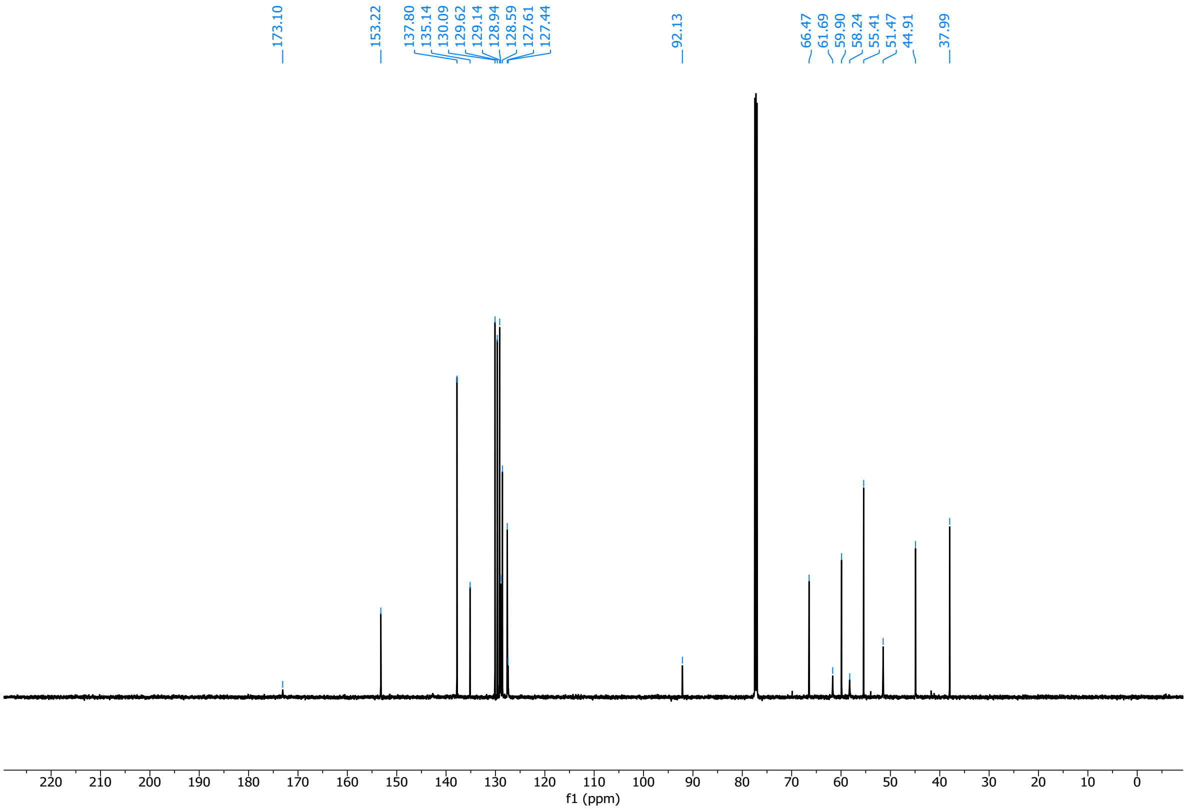

3085 (*R*)-4-benzyl-3-((3*R*,4*S*)-1-benzyl-4-(4-iodophenyl)pyrrolidine-3-carbonyl)oxazolidin-2-one (**16**)

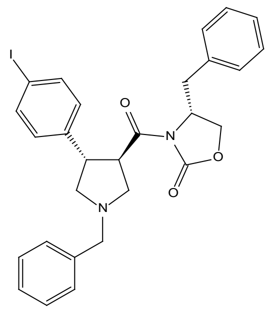

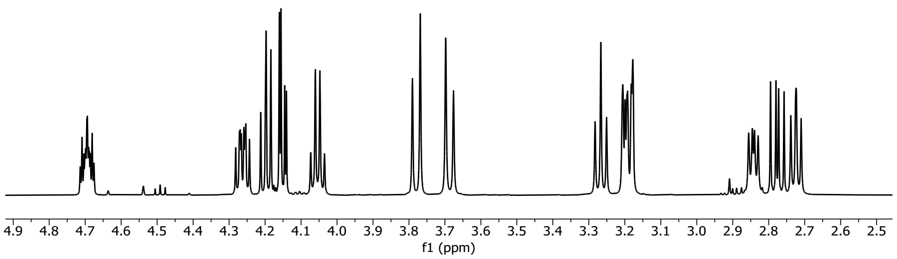

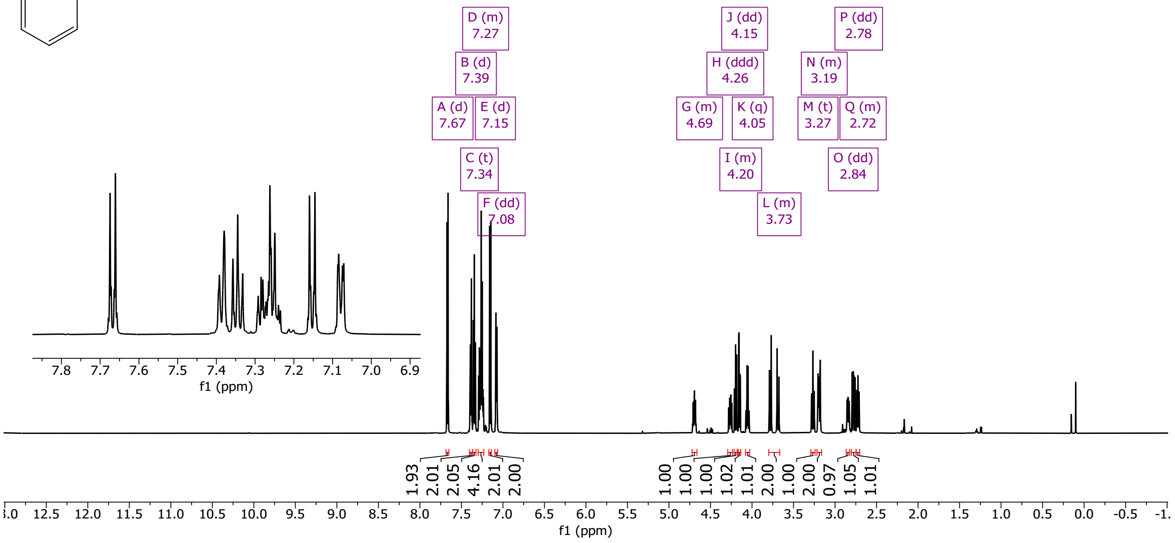

3086

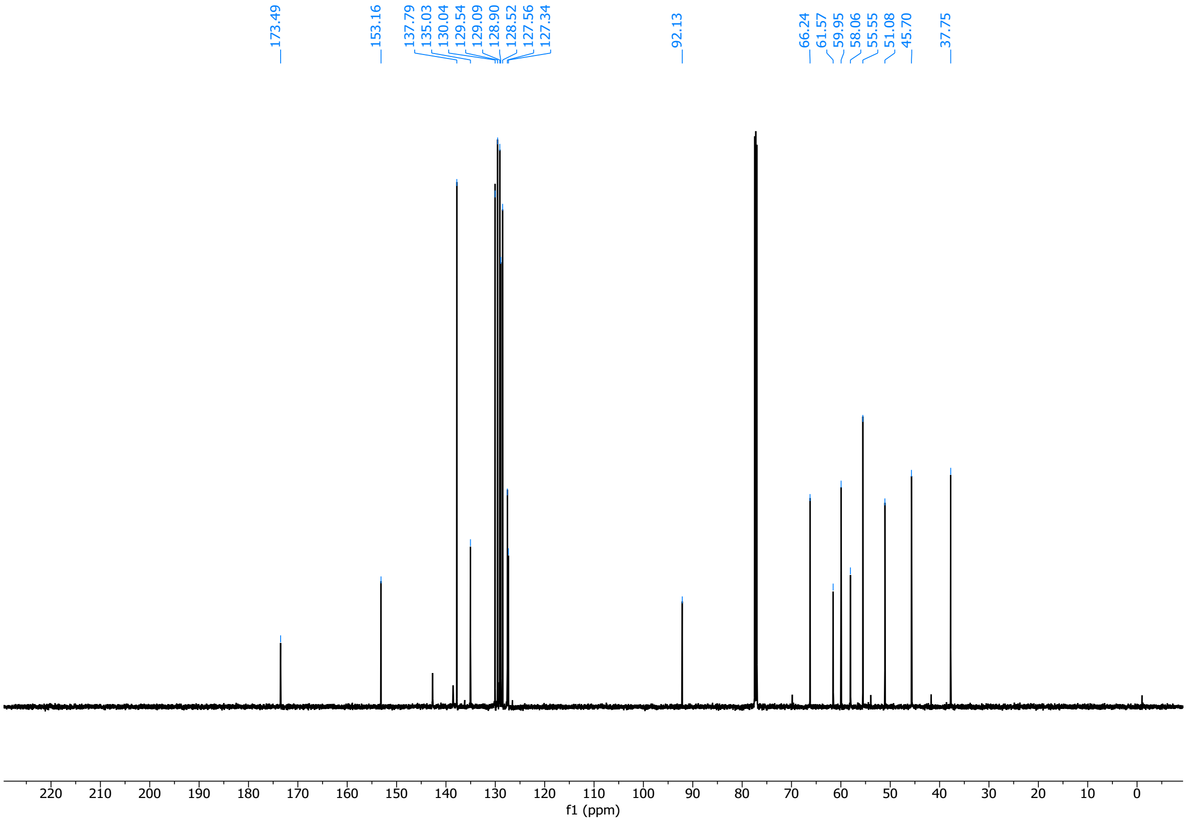

3087

3088 1-(*tert*-butyl) 3-methyl (3*R*,4*S*)-4-(4-iodophenyl)pyrrolidine-1,3-dicarboxylate (**17**)

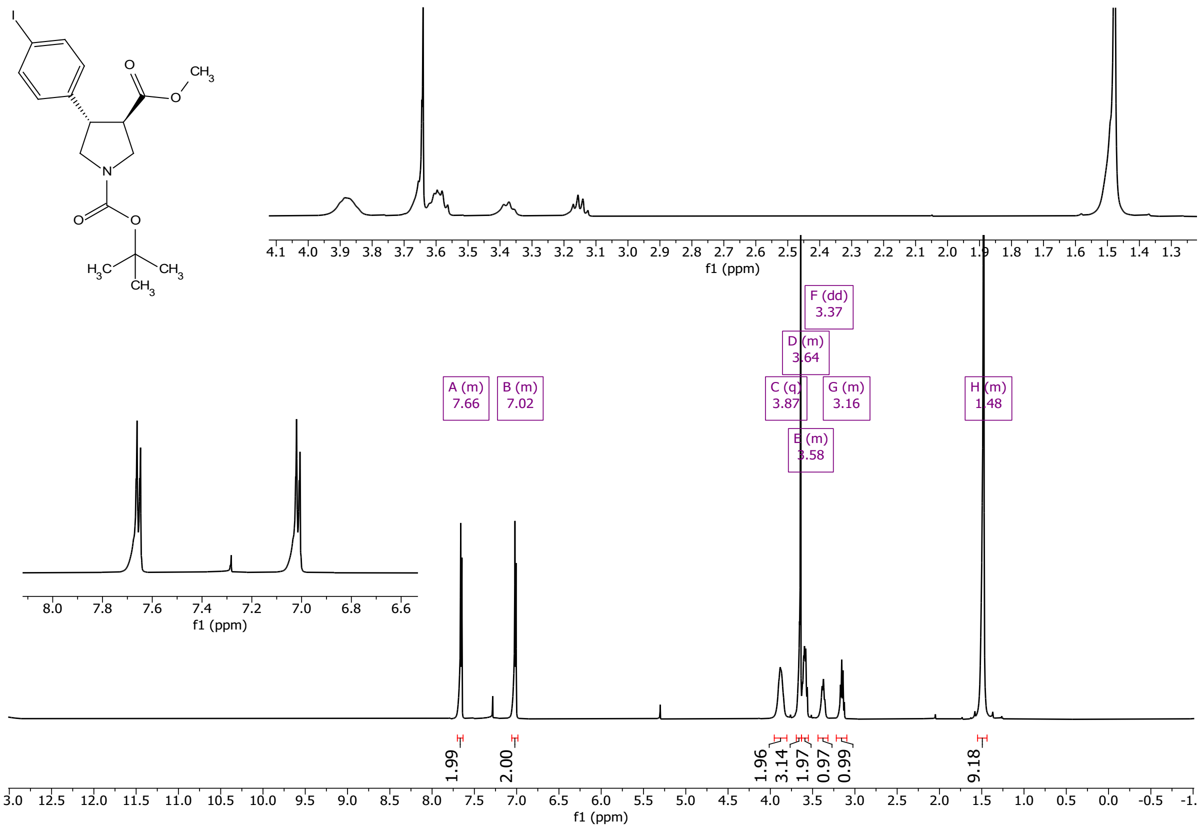

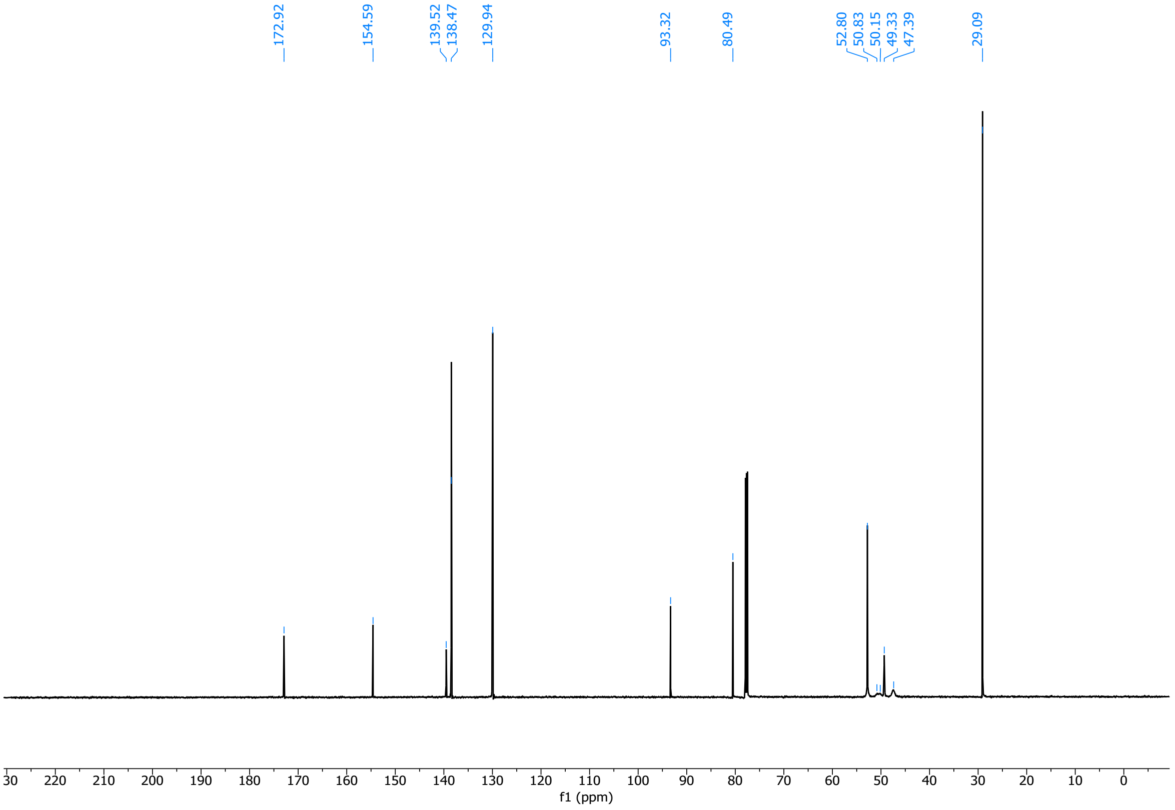

1-(*tert*-butyl) 3-methyl (3*S*,4*R*)-4-(4-iodophenyl)pyrrolidine-1,3-dicarboxylate (**18**)

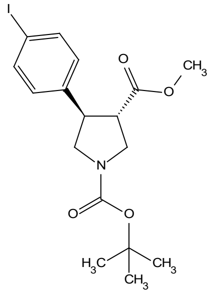

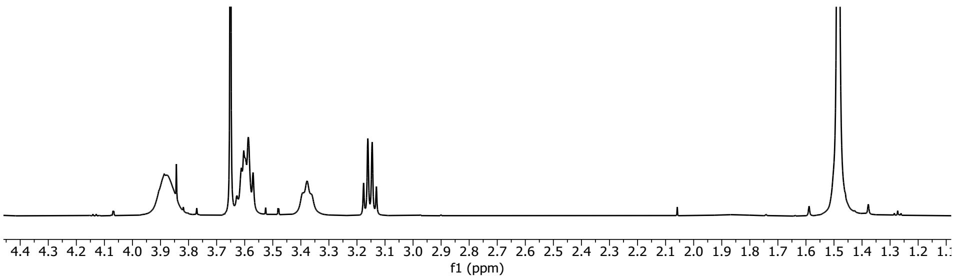

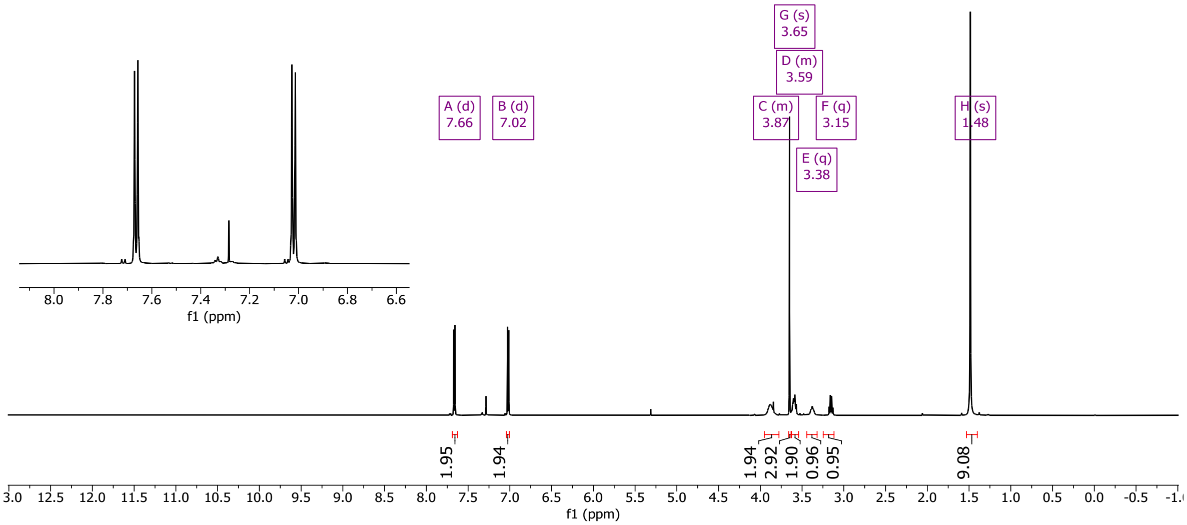

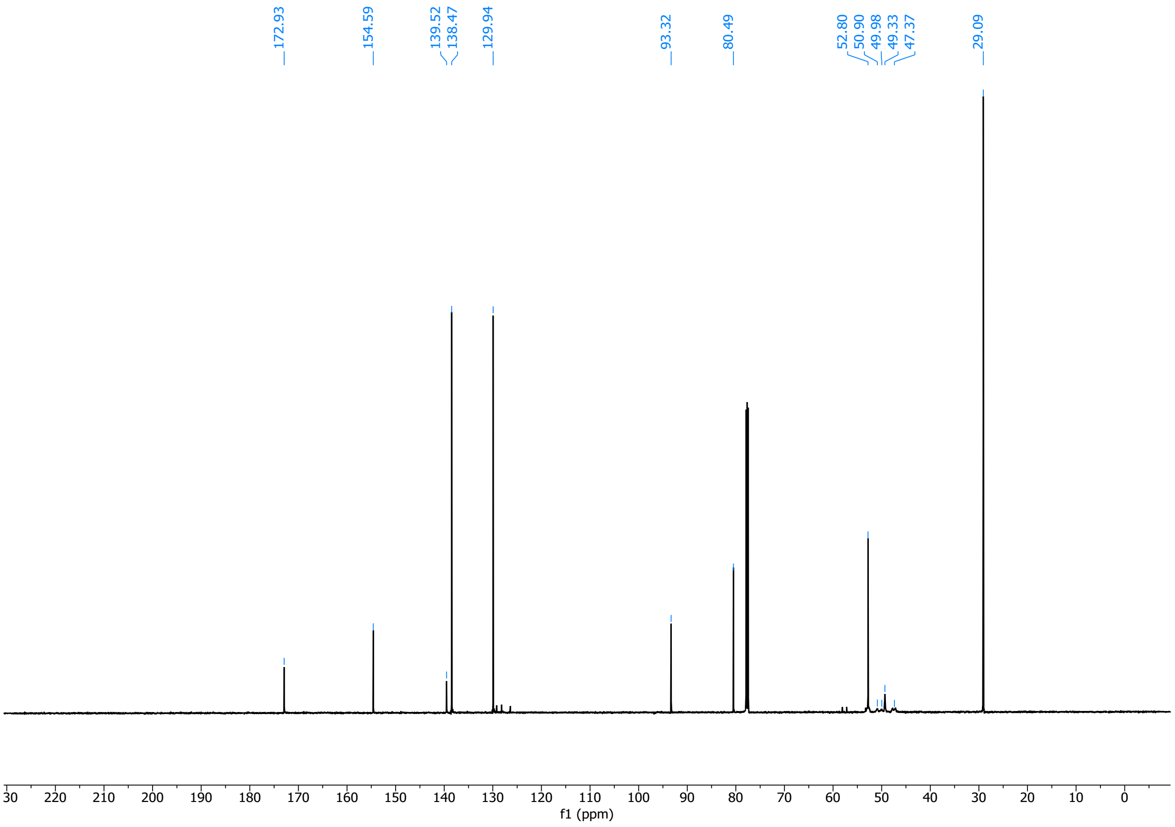

*tert*-butyl (*R*)-8-(4-iodophenyl)-2-((4-nitrophenyl)sulfonyl)-2,6-diazaspiro[3.4]octane-6-carboxylate (**19**)

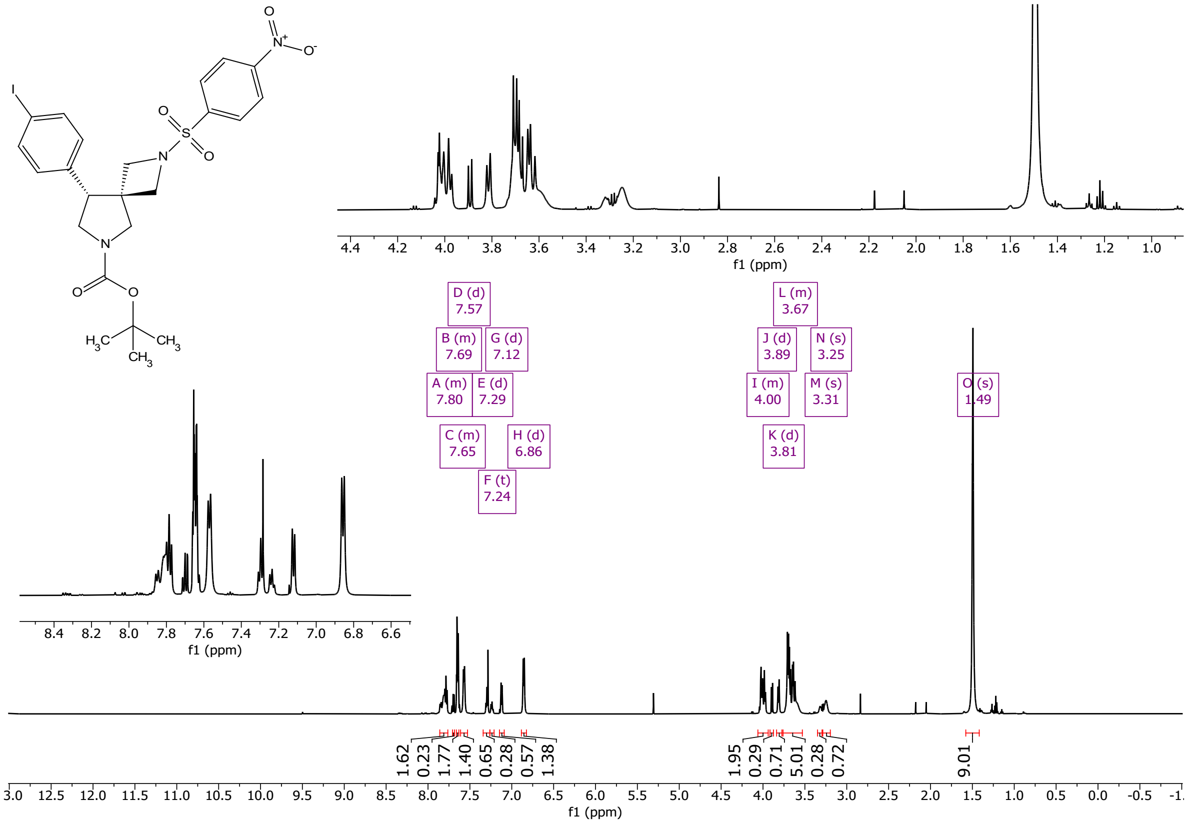

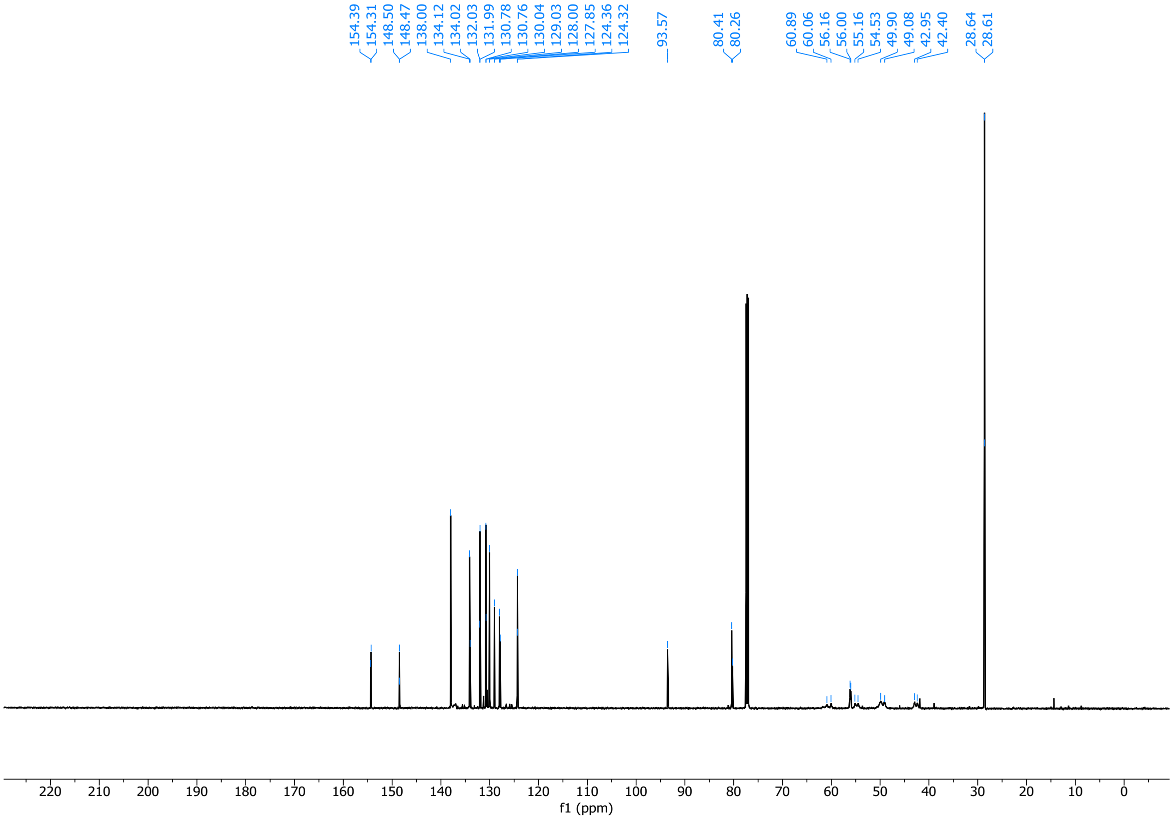

*tert*-butyl (*S*)-8-(4-iodophenyl)-2-((4-nitrophenyl)sulfonyl)-2,6-diazaspiro[3.4]octane-6-carboxylate (**20**)

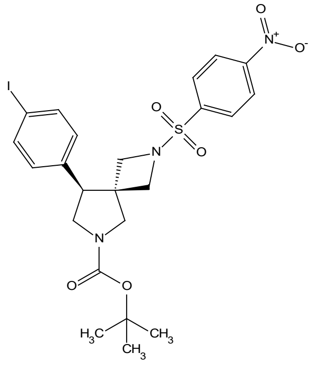

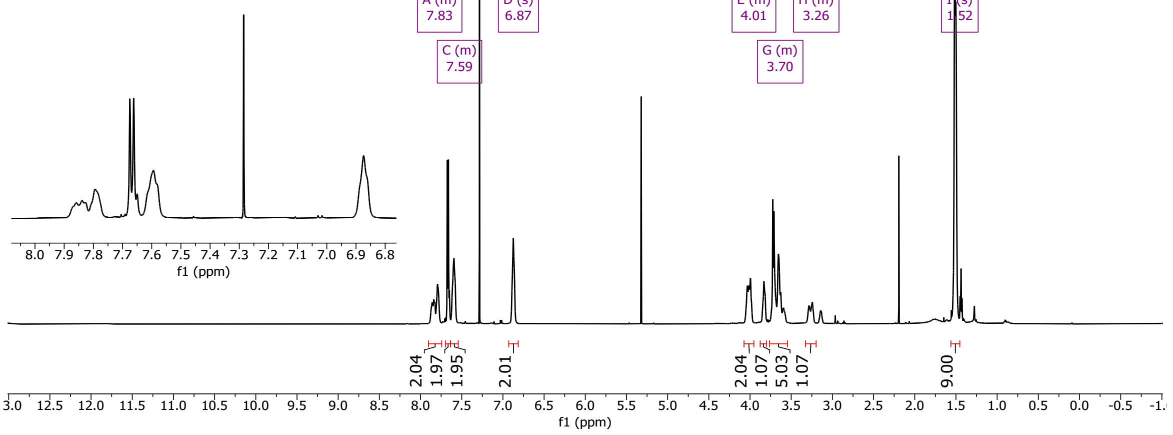

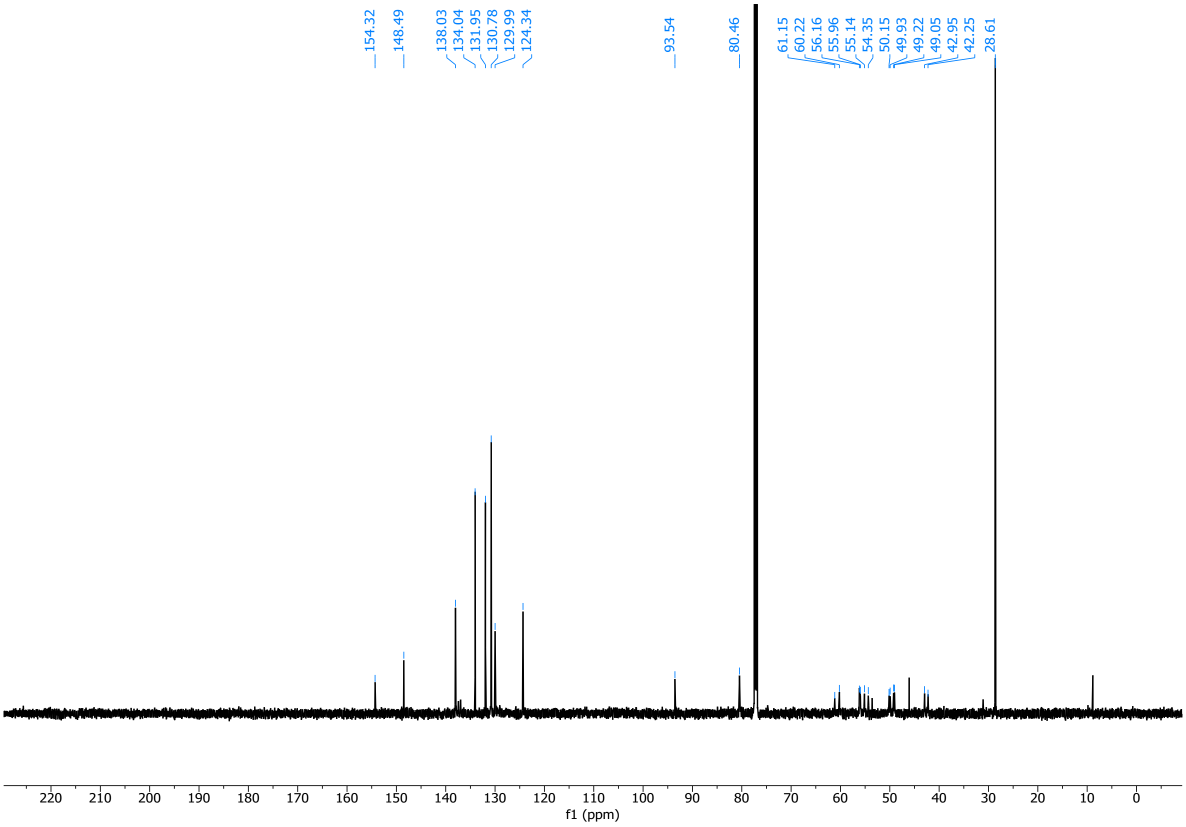

(*R*)-5-(8-(4-iodophenyl)-2-((4-nitrophenyl)sulfonyl)-2,6-diazaspiro[3.4]octan-6-yl)-5-oxopentanoic acid (**21**)

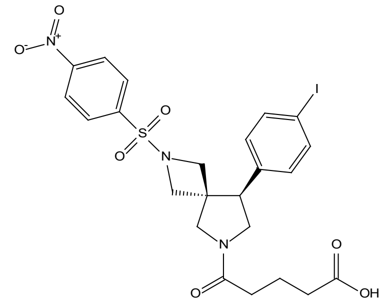

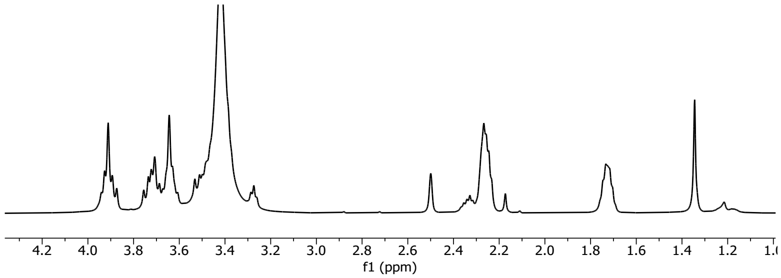

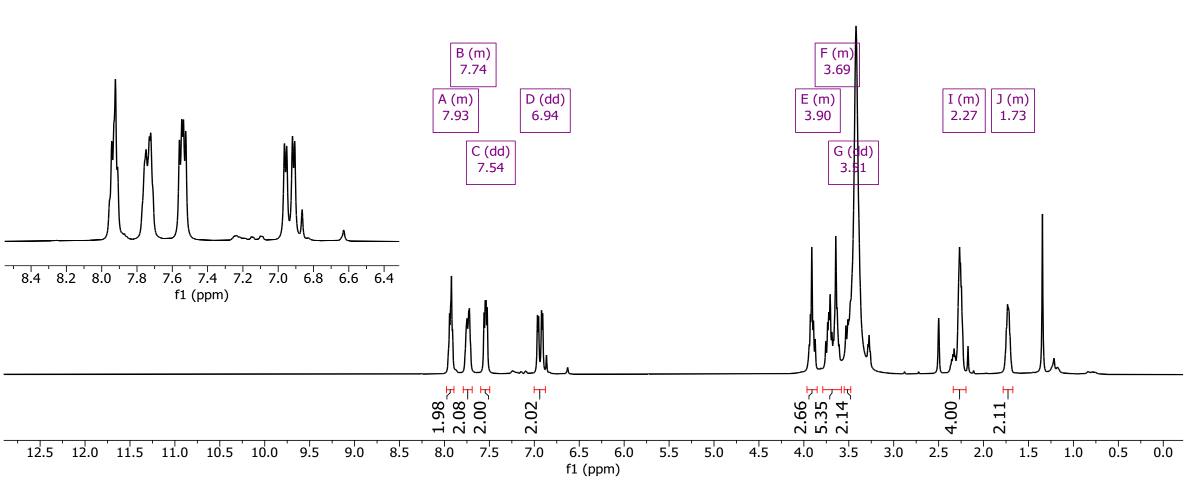

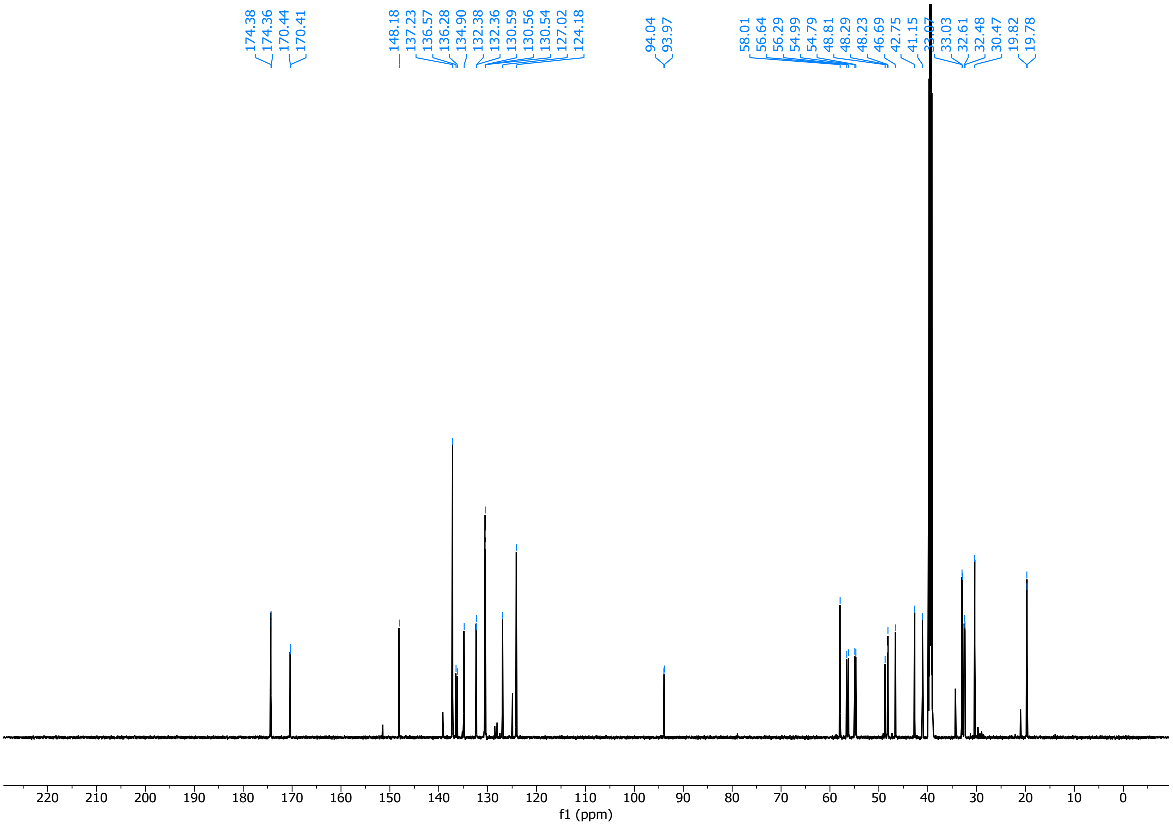

(*S*)-5-(8-(4-iodophenyl)-2-((4-nitrophenyl)sulfonyl)-2,6-diazaspiro[3.4]octan-6-yl)-5-oxopentanoic acid (**22**)

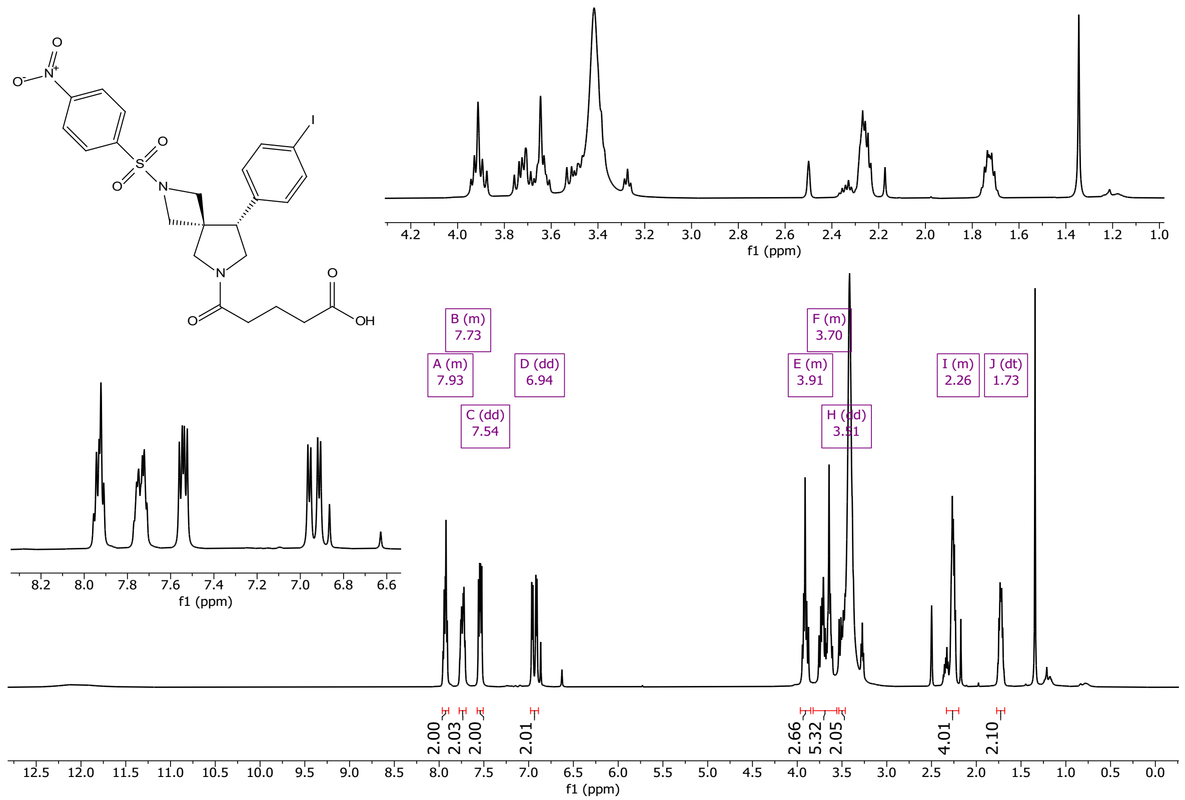

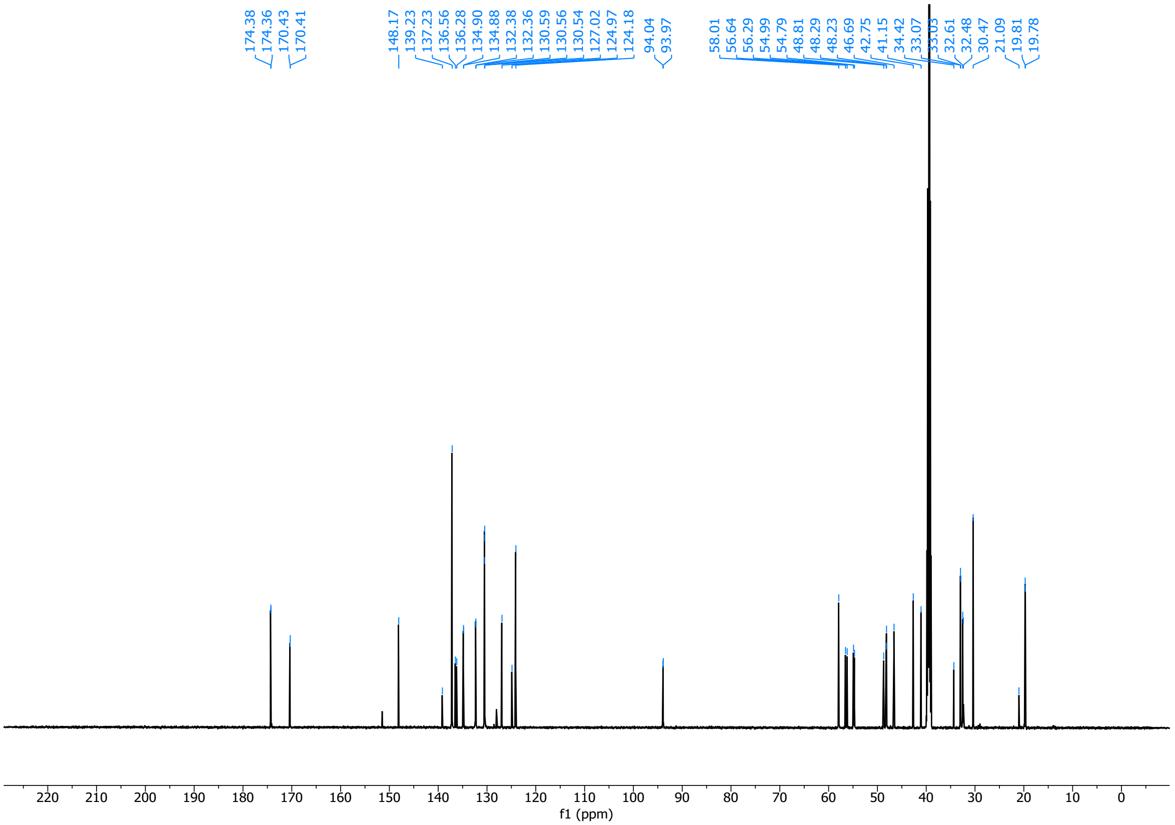

(*R*)-5-(6-(((9H-fluoren-9-yl)methoxy)carbonyl)-8-(4-iodophenyl)-2,6-diazaspiro[3.4]octan-2-yl)-5-oxopentanoic acid (**23**)

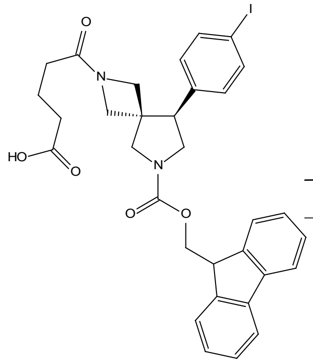

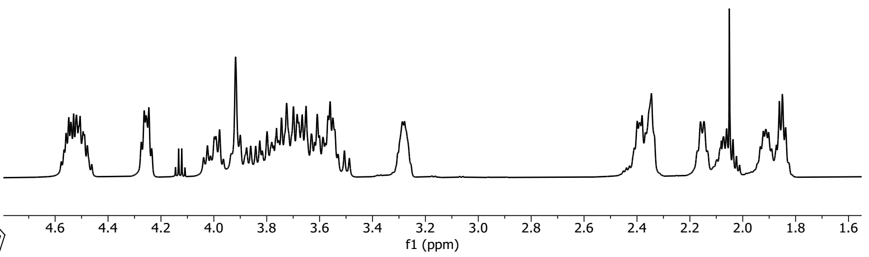

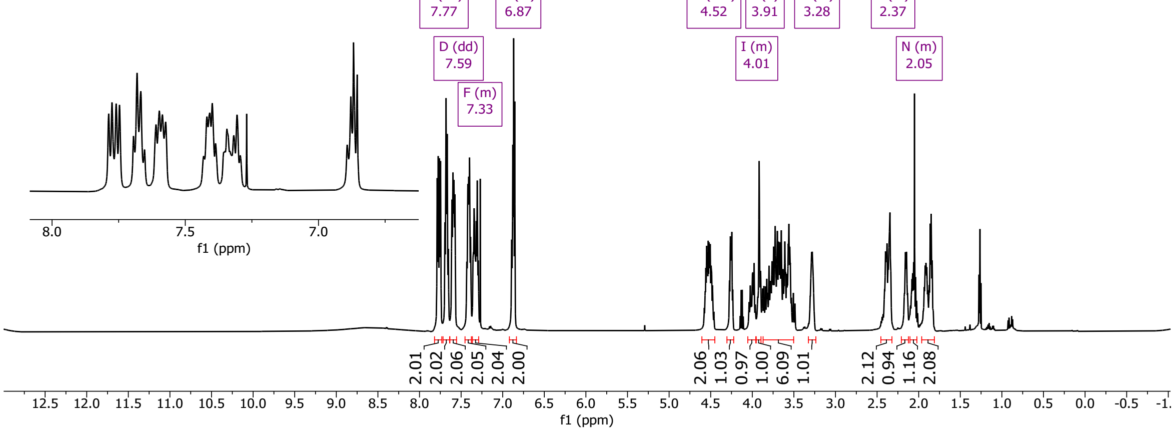

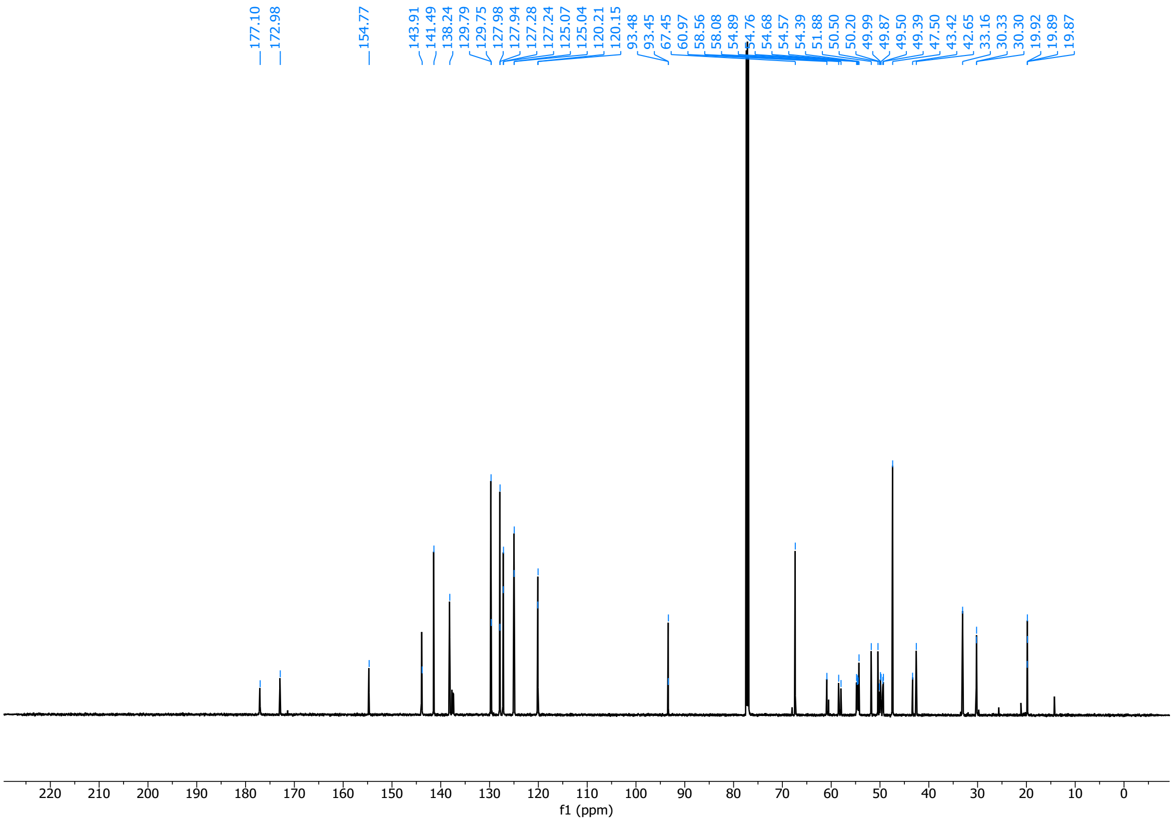

(*S*)-5-(6-(((9H-fluoren-9-yl)methoxy)carbonyl)-8-(4-iodophenyl)-2,6-diazaspiro[3.4]octan-2-yl)-5-oxopentanoic acid (**24**)

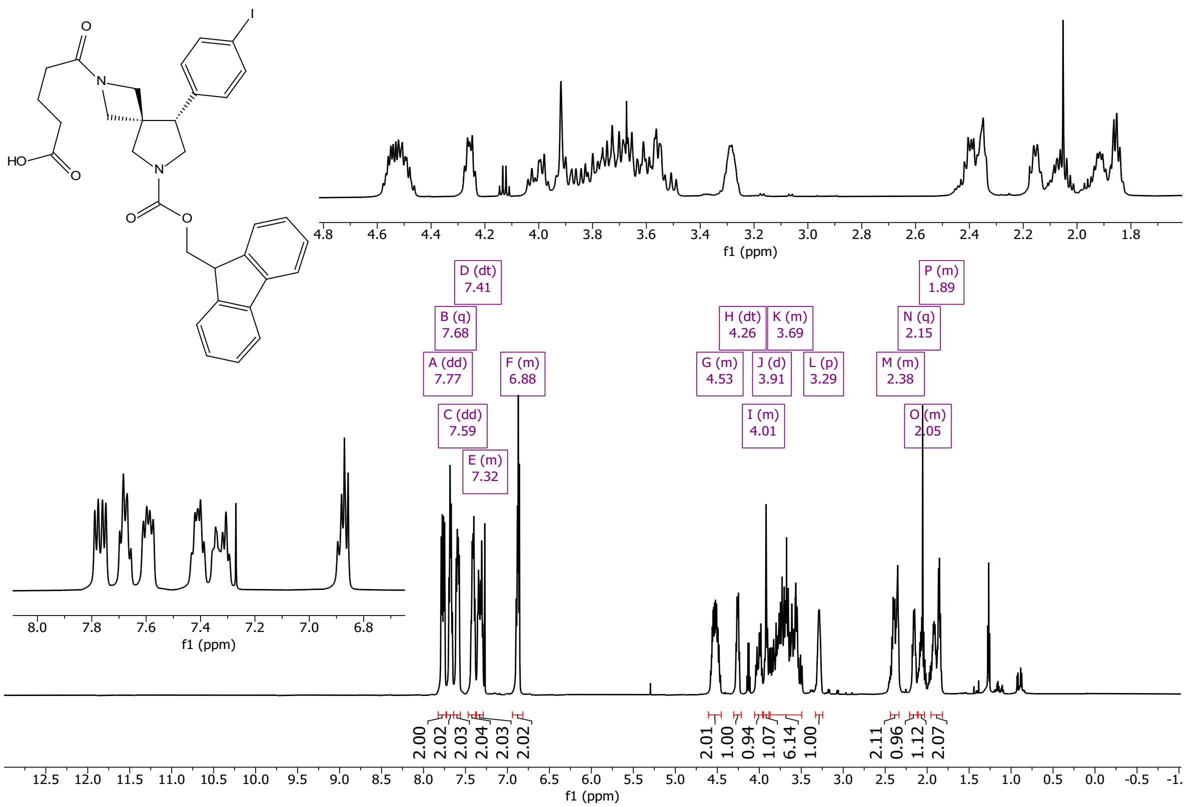

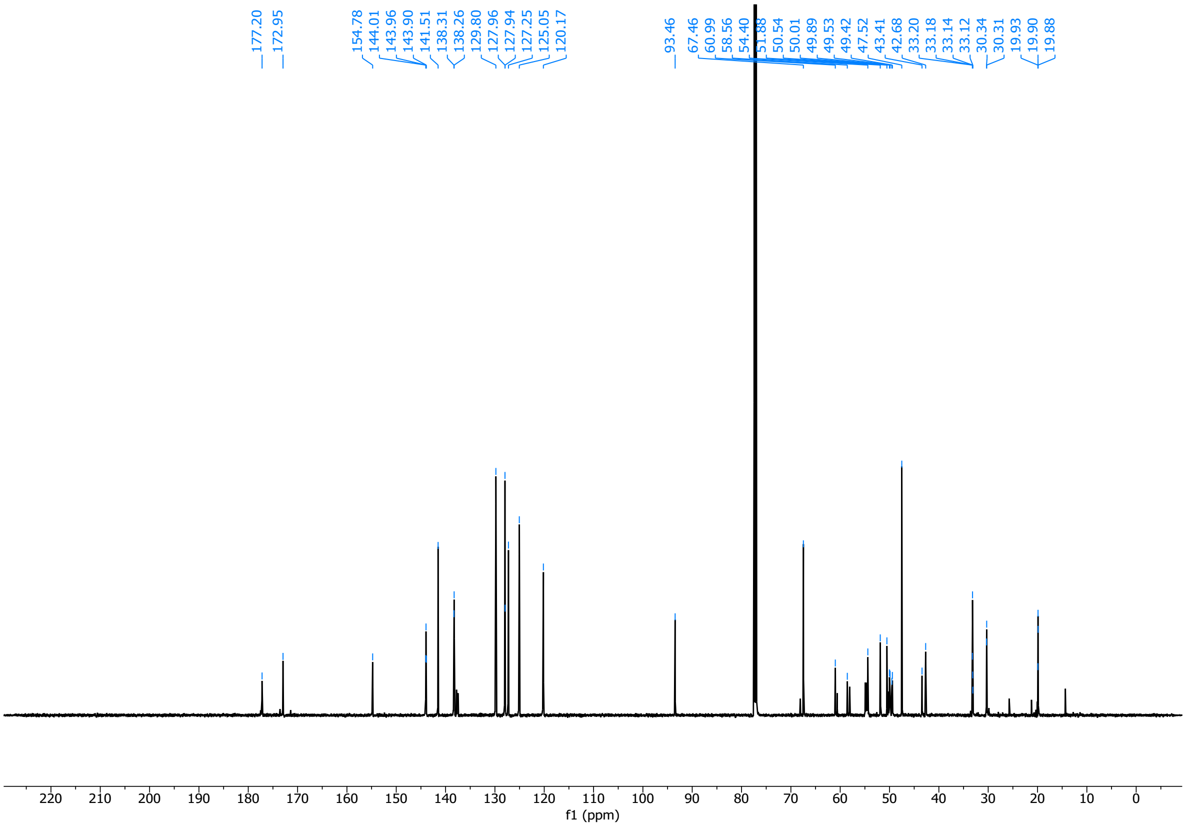

#### 7.4. Enumerated DEL compounds off-DNA

(2*S*,4*S*)-4-(3-((*E*)-2-cyclohexylvinyl)phenoxy)-1-(3-ethynylbenzyl)-*N*-methylpyrrolidine-2-carboxamide (**27**)

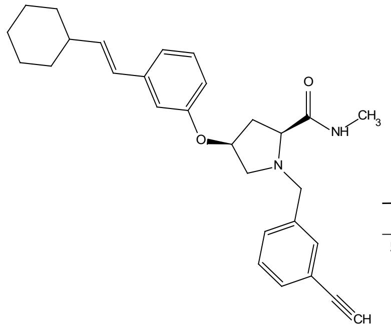

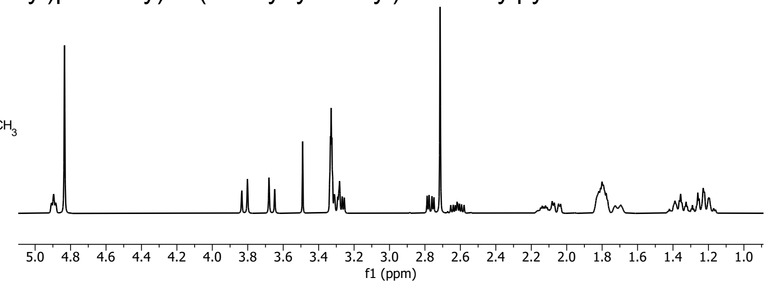

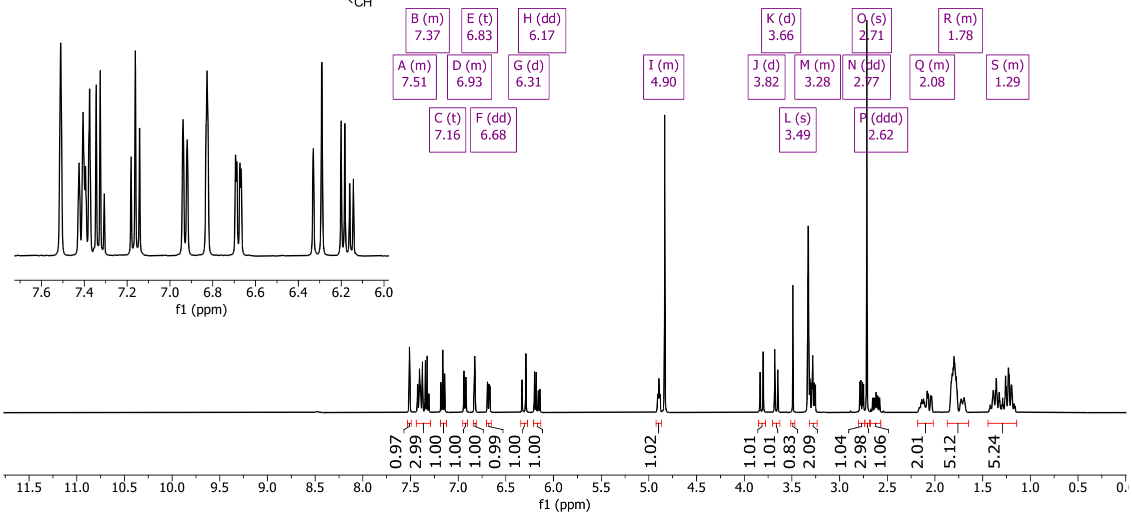

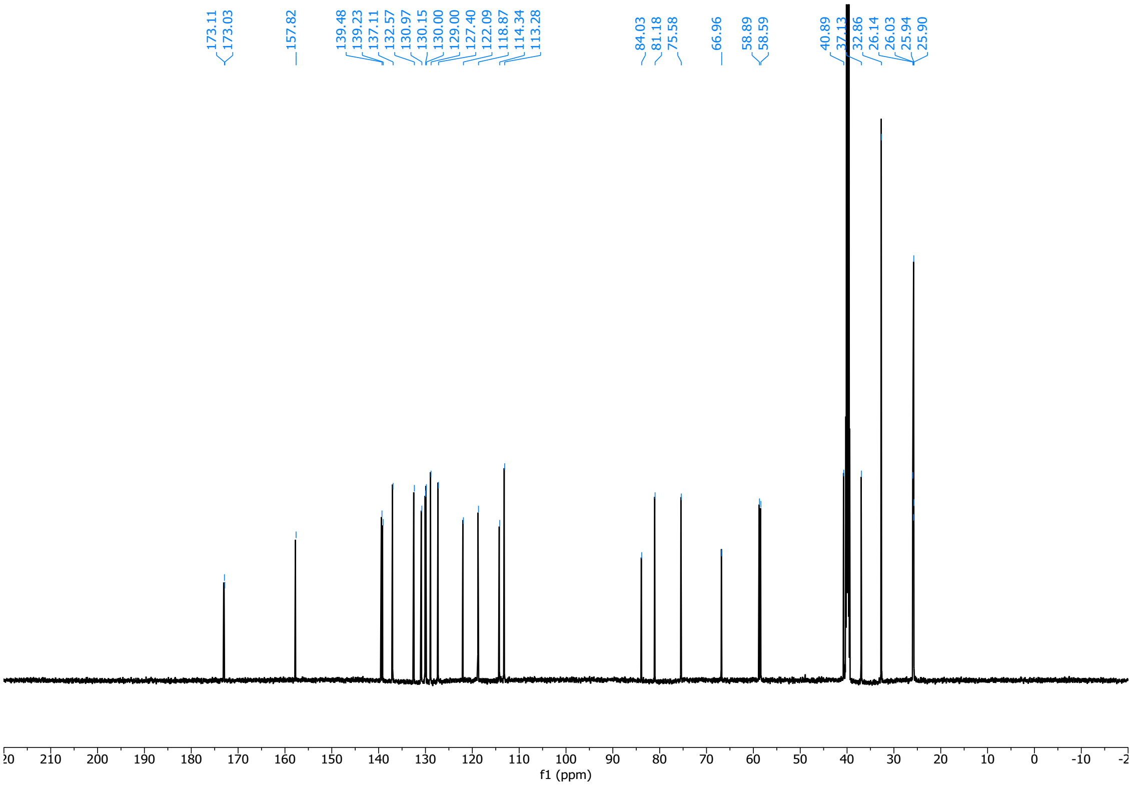

(2*R*,4*S*)-4-(3-((*E*)-2-cyclohexylvinyl)phenoxy)-1-(3-ethynylbenzyl)-*N*-methylpyrrolidine-2-carboxamide (**28**)

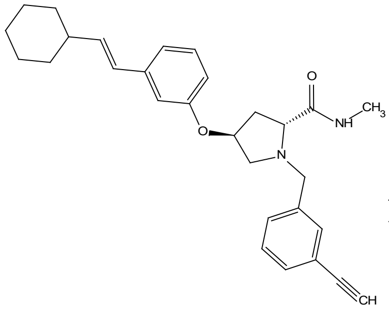

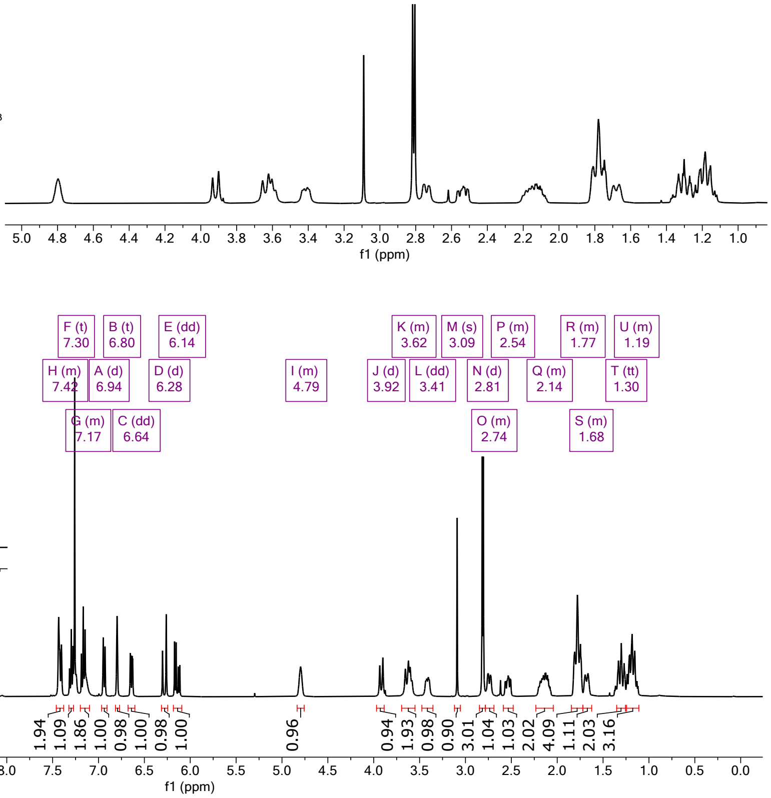

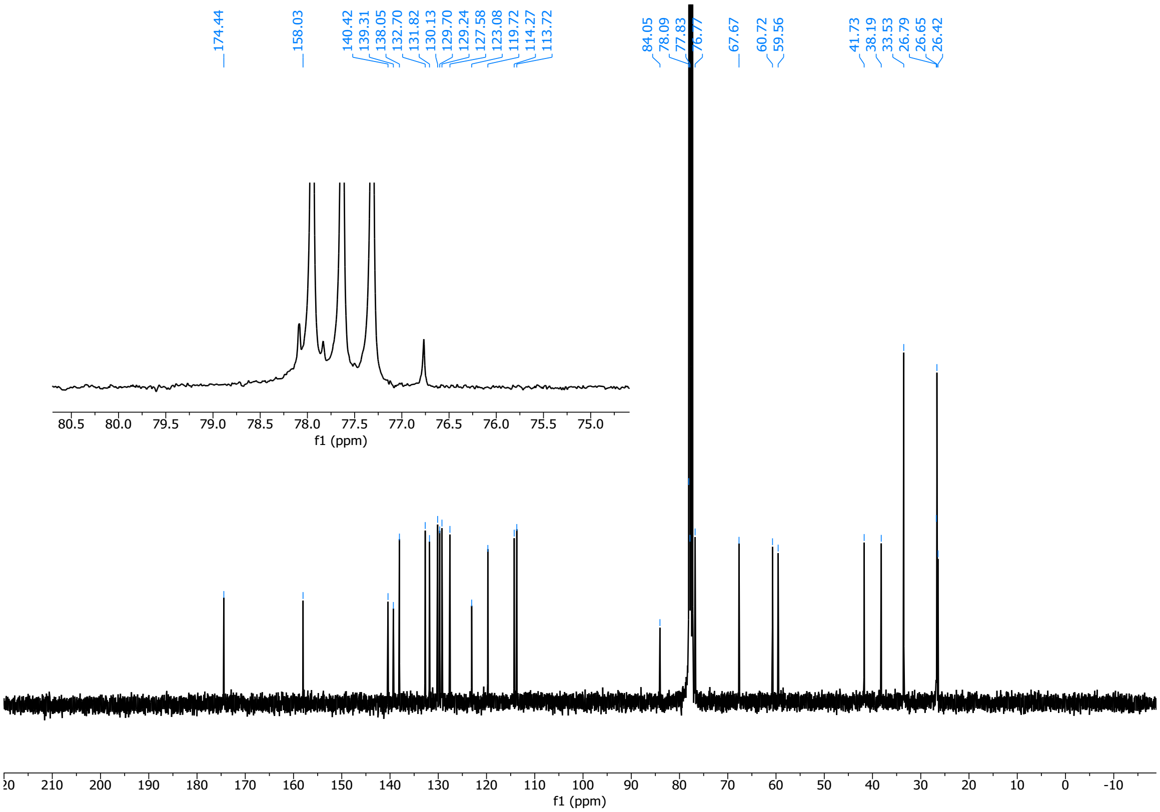

(2*S*,4*S*)-4-(4-((*E*)-2-cyclohexylvinyl)phenoxy)-1-(3-cyclopropyl-1-ethyl-1H-pyrazole-5-carbonyl)-*N*-methylpyrrolidine-2-carboxamide (**29**)

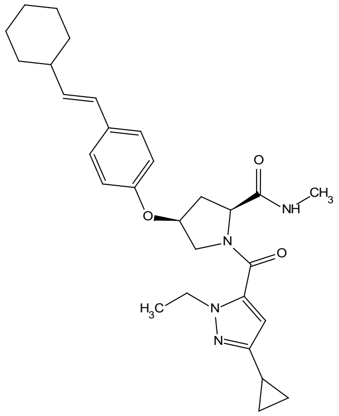

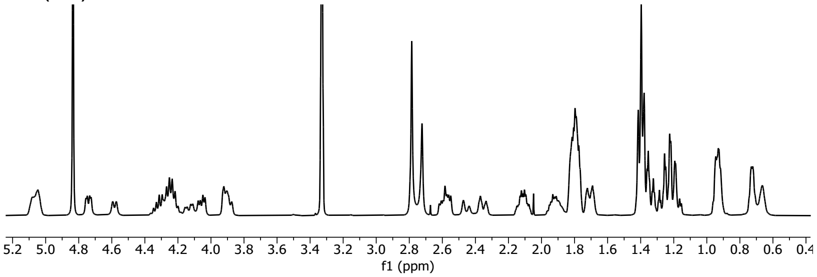

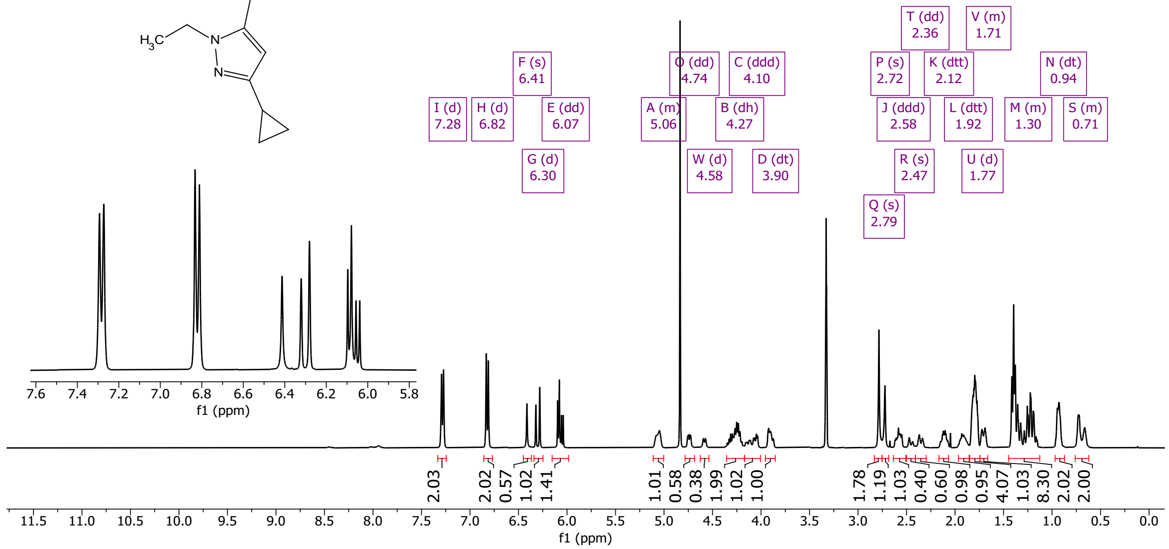

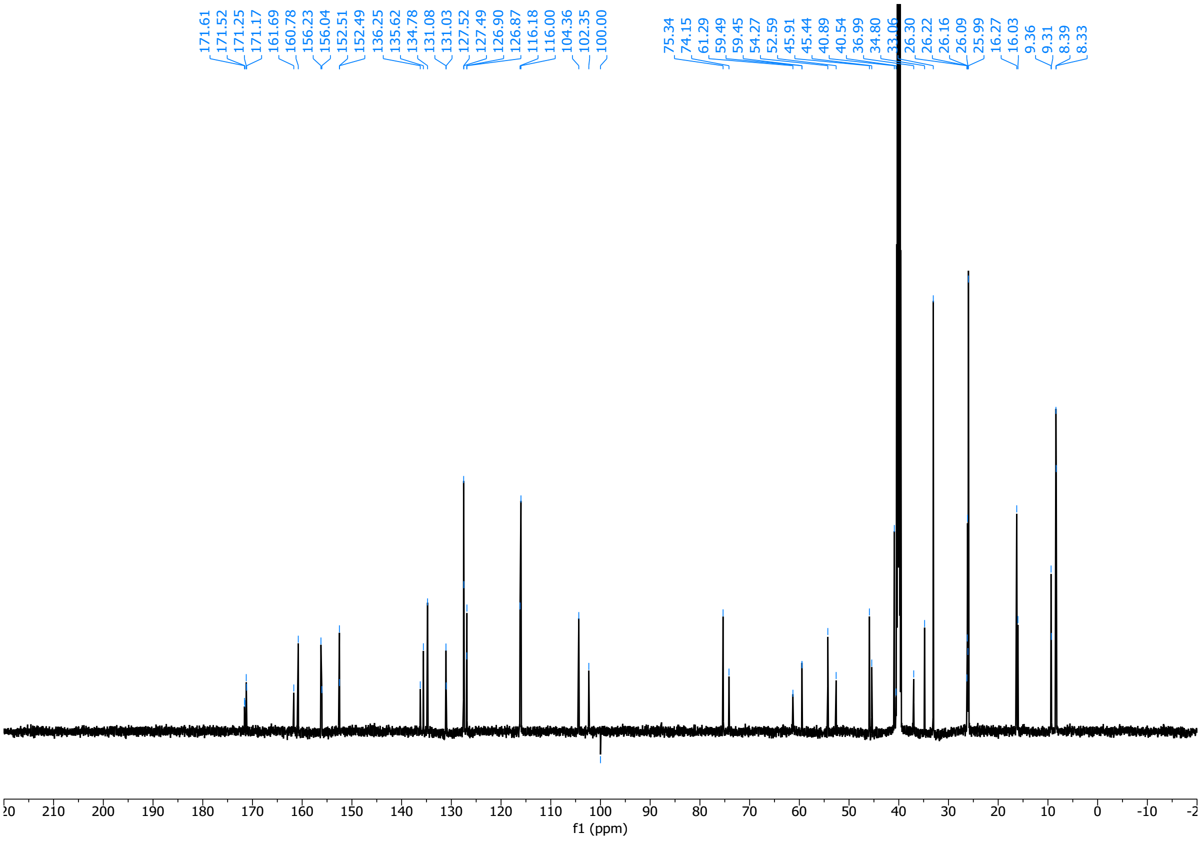

(2*S*,4*R*)-4-(4-((*E*)-2-cyclohexylvinyl)phenoxy)-1-(3-cyclopropyl-1-ethyl-1H-pyrazole-5-carbonyl)-*N*-methylpyrrolidine-2-carboxamide (**30**)

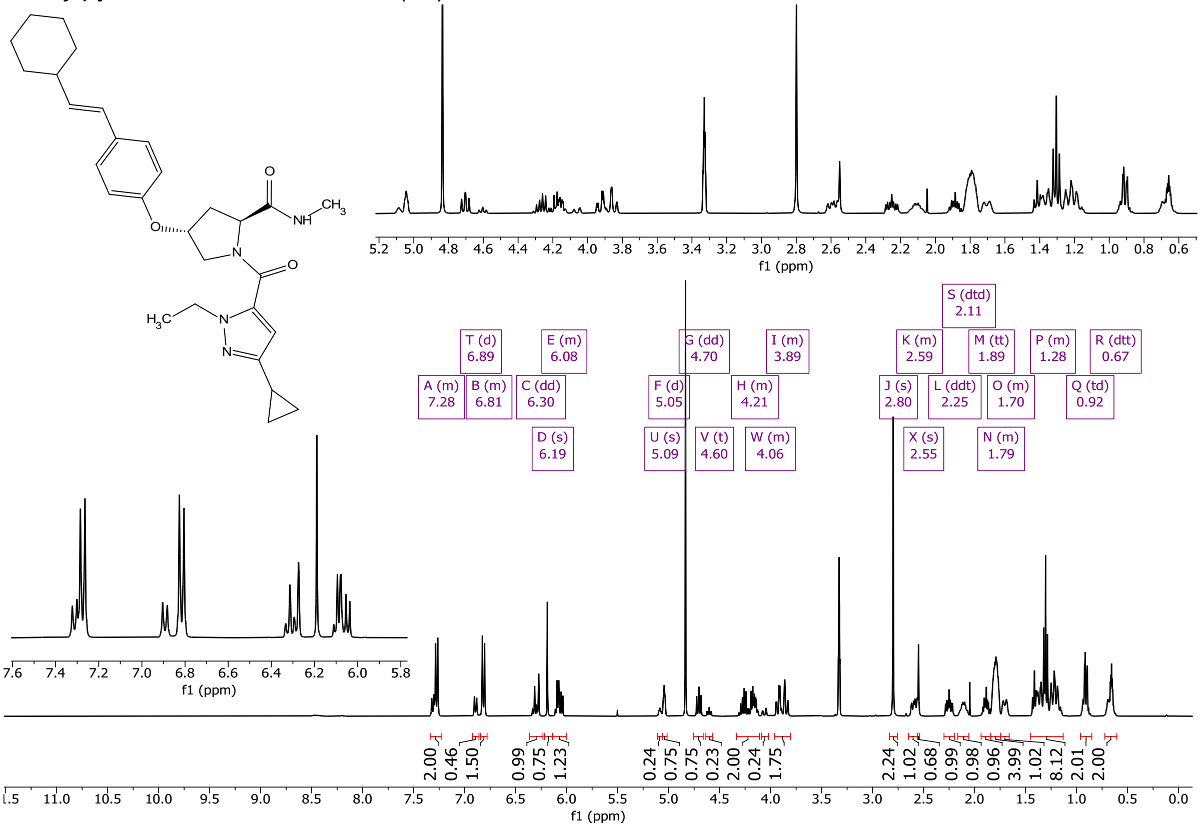

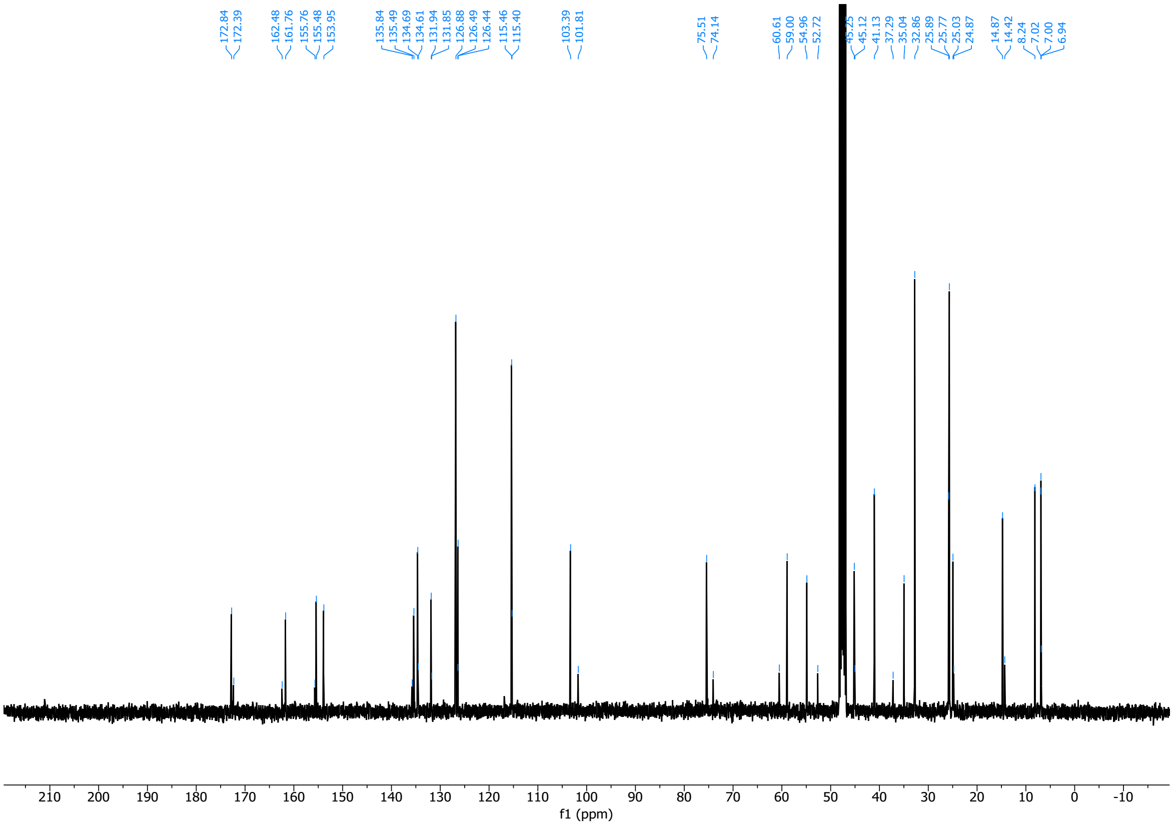

(2*R*,4*R*)-4-(4-((*E*)-2-cyclohexylvinyl)phenoxy)-1-(3-cyclopropyl-1-ethyl-1H-pyrazole-5-carbonyl)-*N*-methylpyrrolidine-2-carboxamide (**31**)

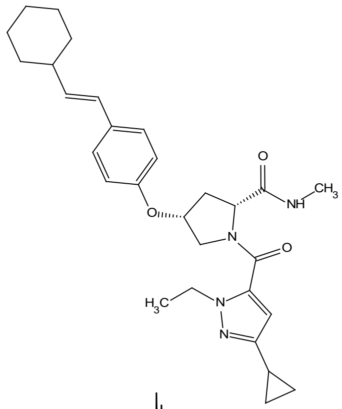

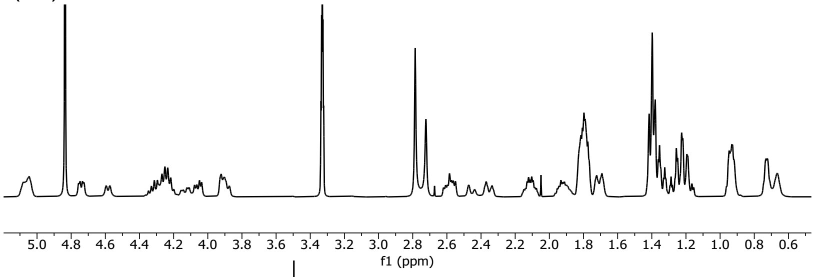

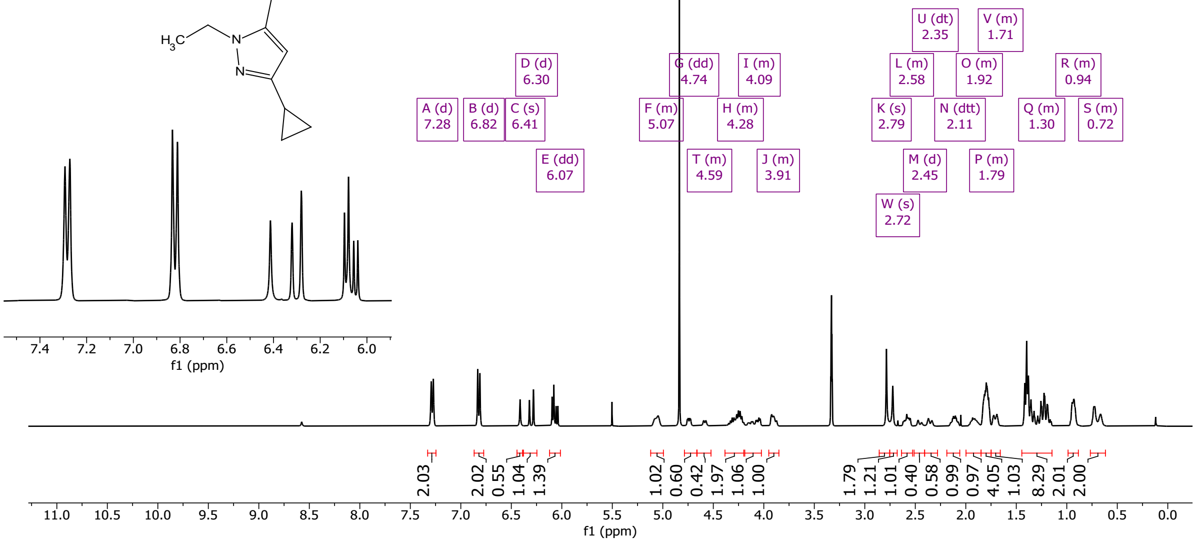

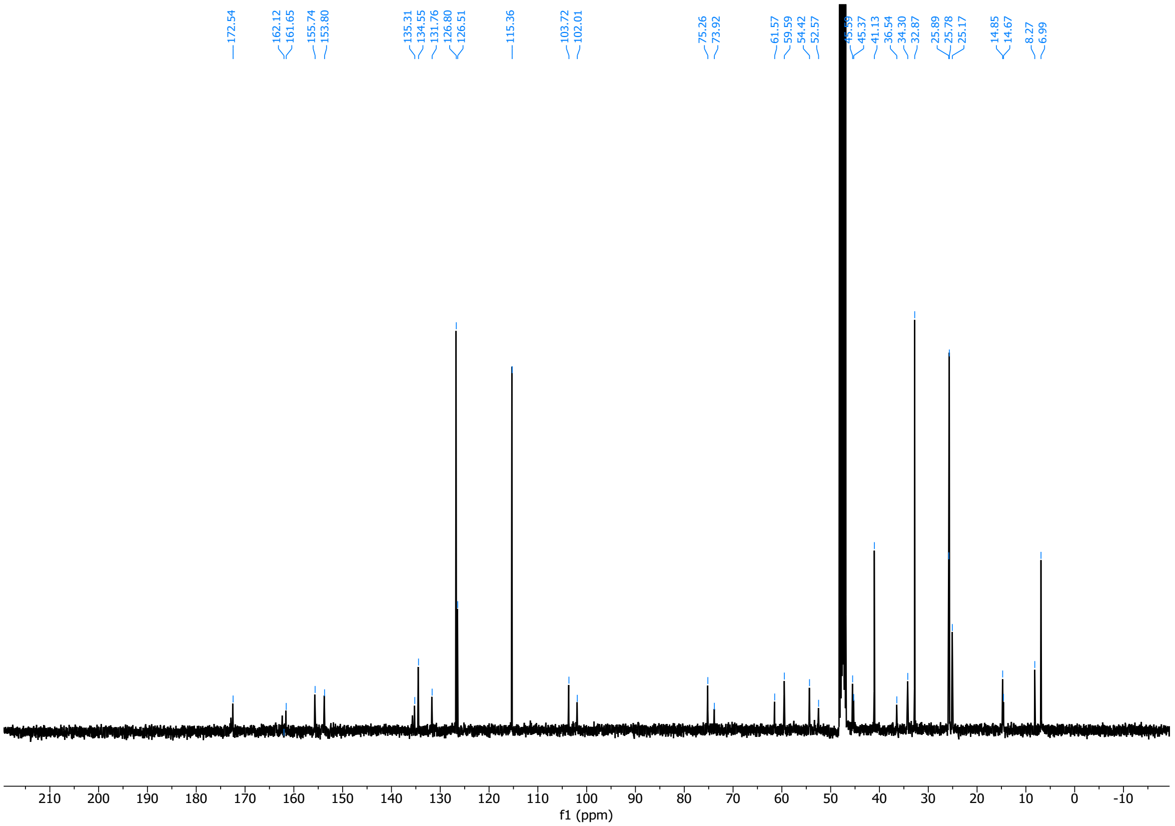

(2*R*,4*S*)-4-(4-((*E*)-2-cyclohexylvinyl)phenoxy)-1-(3-(2,3-dihydro-1H-inden-1-yl)propanoyl)-*N*-methylpyrrolidine-2-carboxamide (**32**)

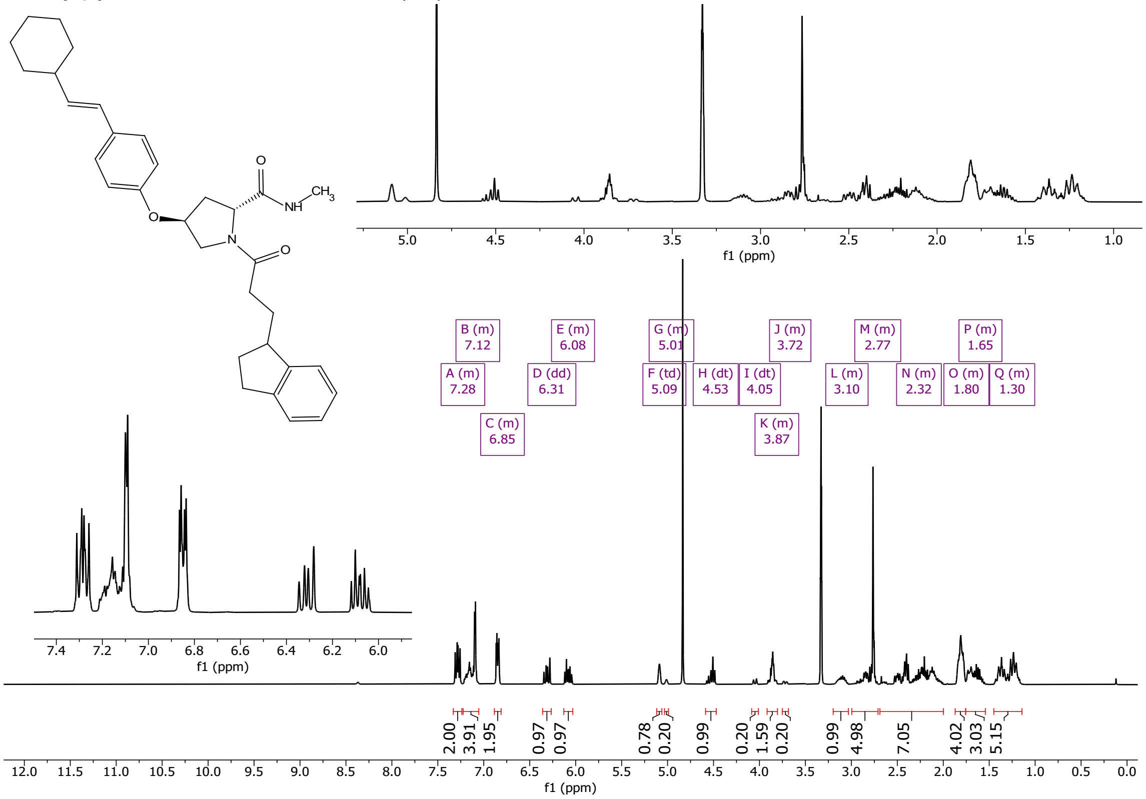

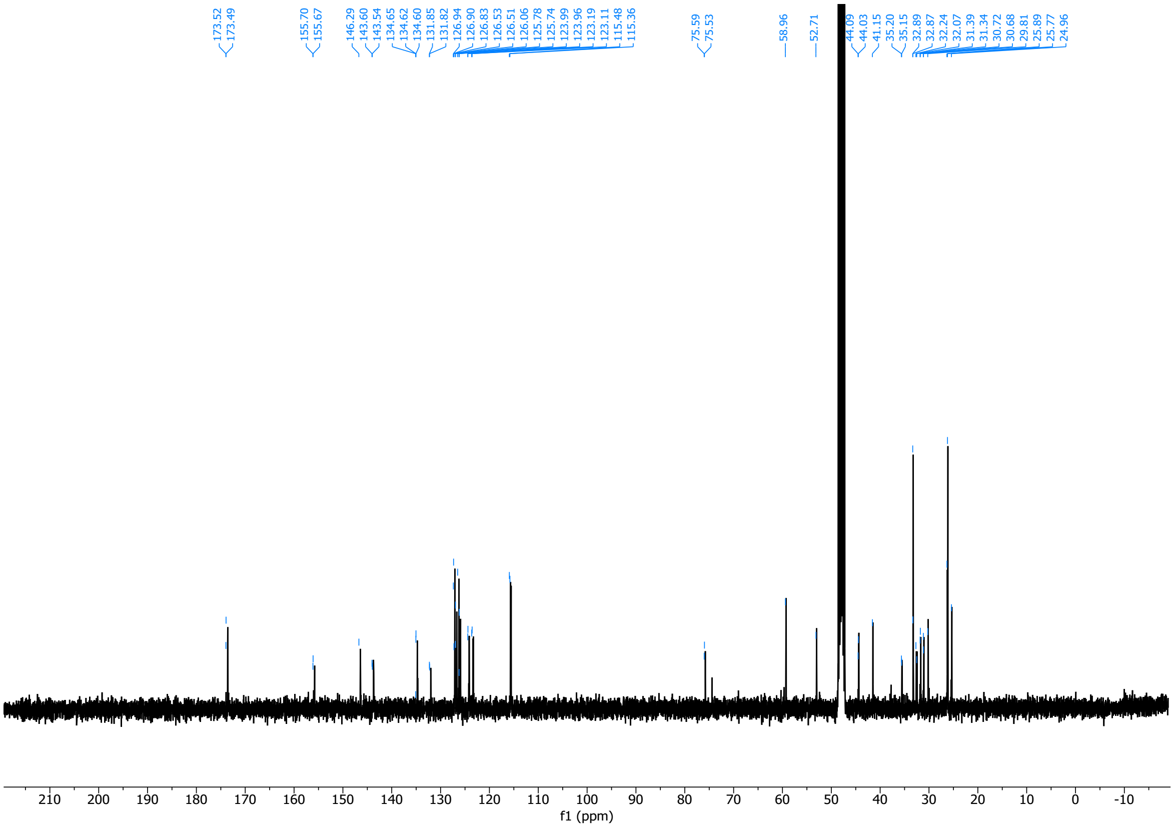

(2*R*,4*S*)-4-(4-((*E*)-2-cyclohexylvinyl)phenoxy)-1-(3-cyclopropyl-1-ethyl-1H-pyrazole-5-carbonyl)-*N*-methylpyrrolidine-2-carboxamide (**33**)

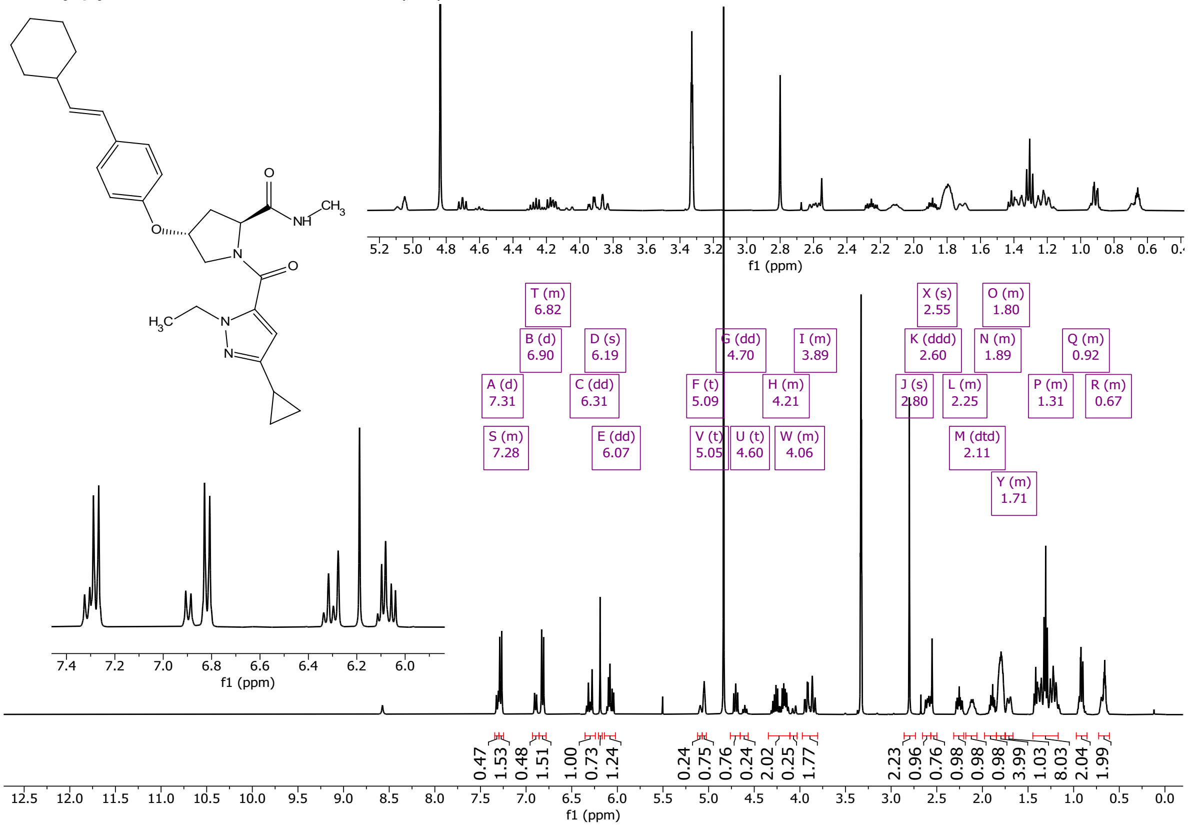

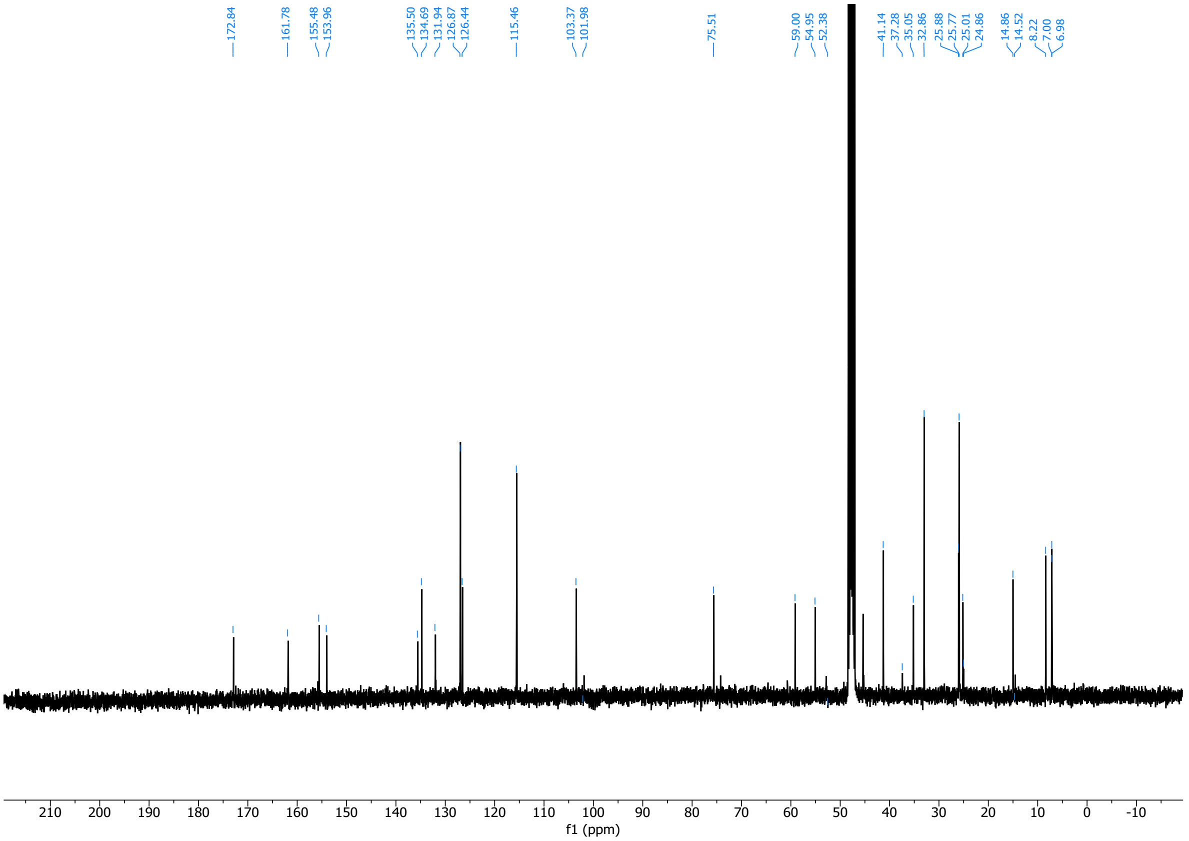

(6*R*,7*S*,8*S*)-4-((3-chloro-4-fluorophenyl)sulfonyl)-7-(4-((*E*)-2-cyclohexylvinyl)phenyl)-8-(hydroxymethyl)-1,4-diazabicyclo[4.2.0]octan-2-one (**34**)

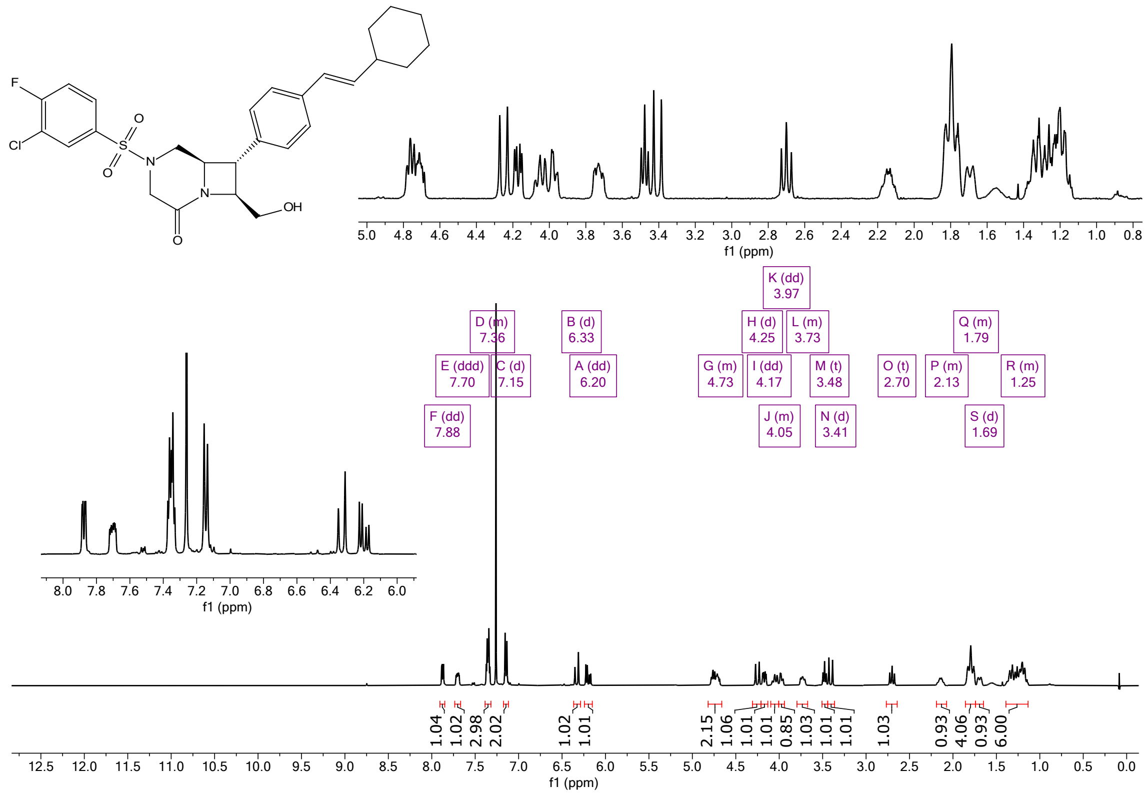

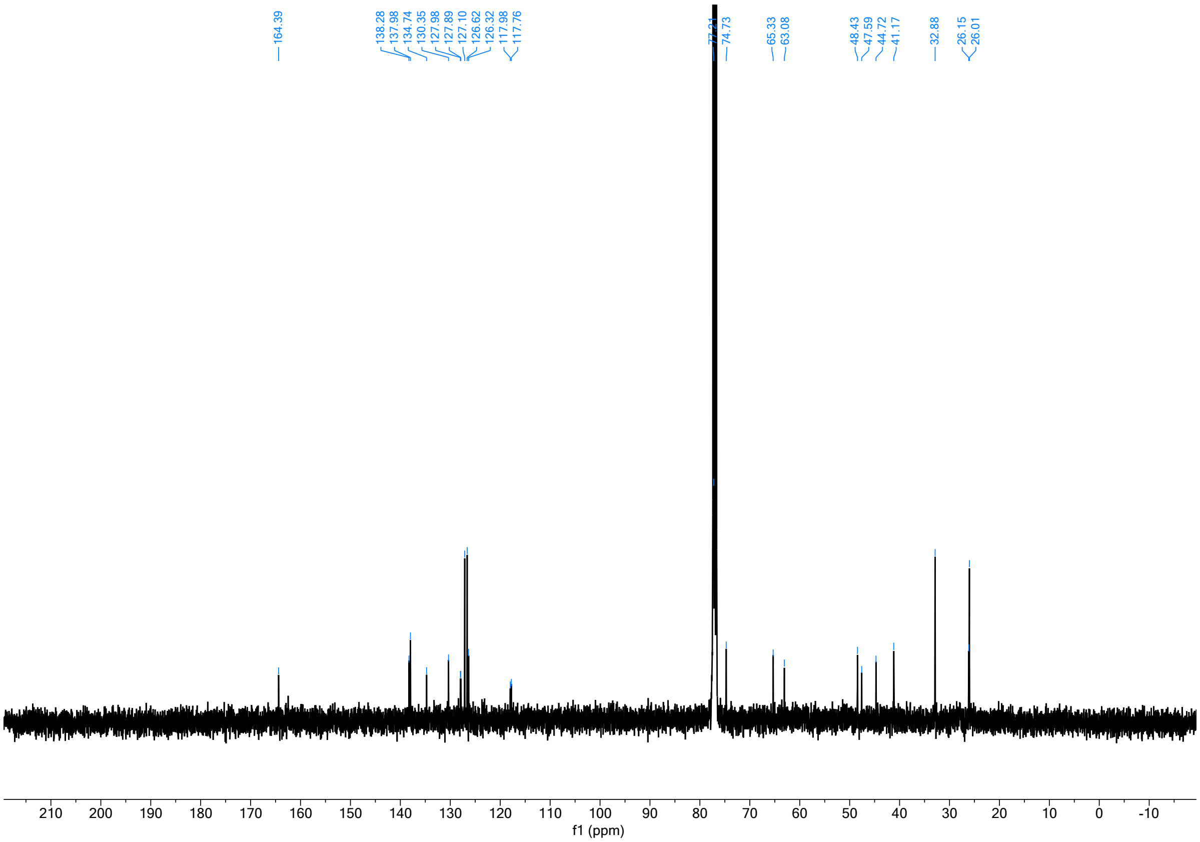
164.39
138.28
137.98
134.74
130.35
127.98
127.89
127.10
126.62
126.32
117.98
117.76
74.73
65.33
63.08
48.43
47.59
44.72
41.17
32.88
26.15
26.01
210 200 190 180 170 160 150 140 130 120 110 100 90 80 70 60 50 40 30 20 10 0 -10
f1 (ppm)

3147

*N*-(((4*R*,5*R*)-8-((*E*)-2-cyclohexylvinyl)-2-((*R*)-1-hydroxypropan-2-yl)-4-methyl-1,1-dioxido-2,3,4,5-tetrahydrobenzo[*b*][1,4,5]oxathiazocin-5-yl)methyl)-*N*-methyl-2-phenylcyclopent-1-ene-1-carboxamide (**35**)

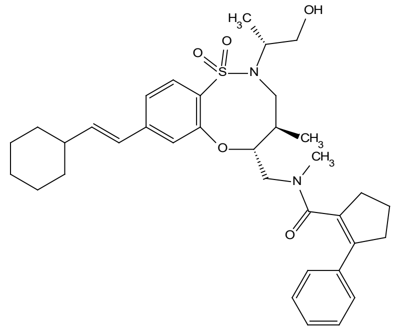

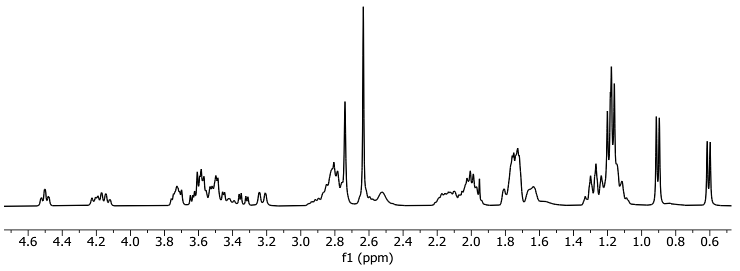

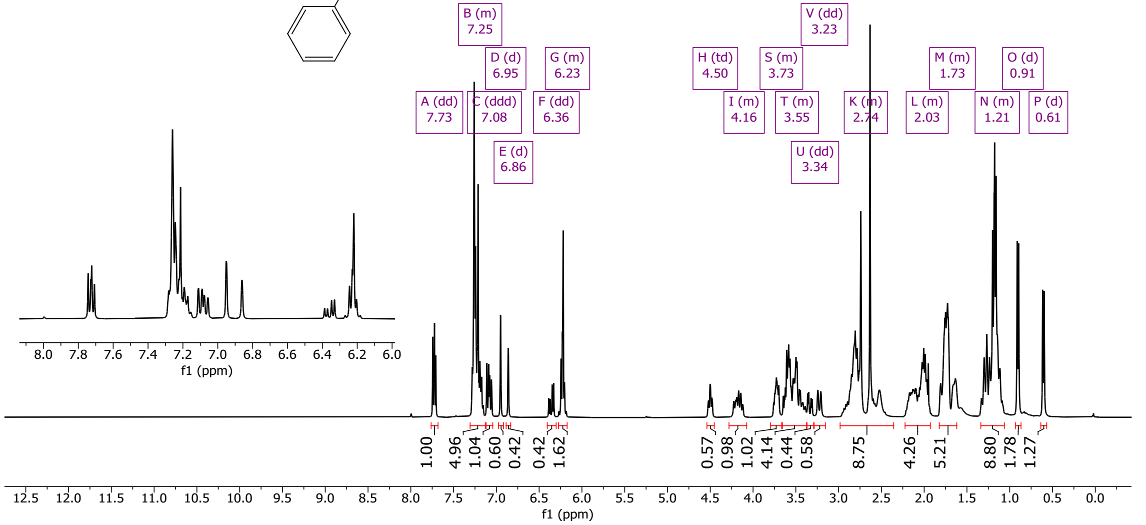

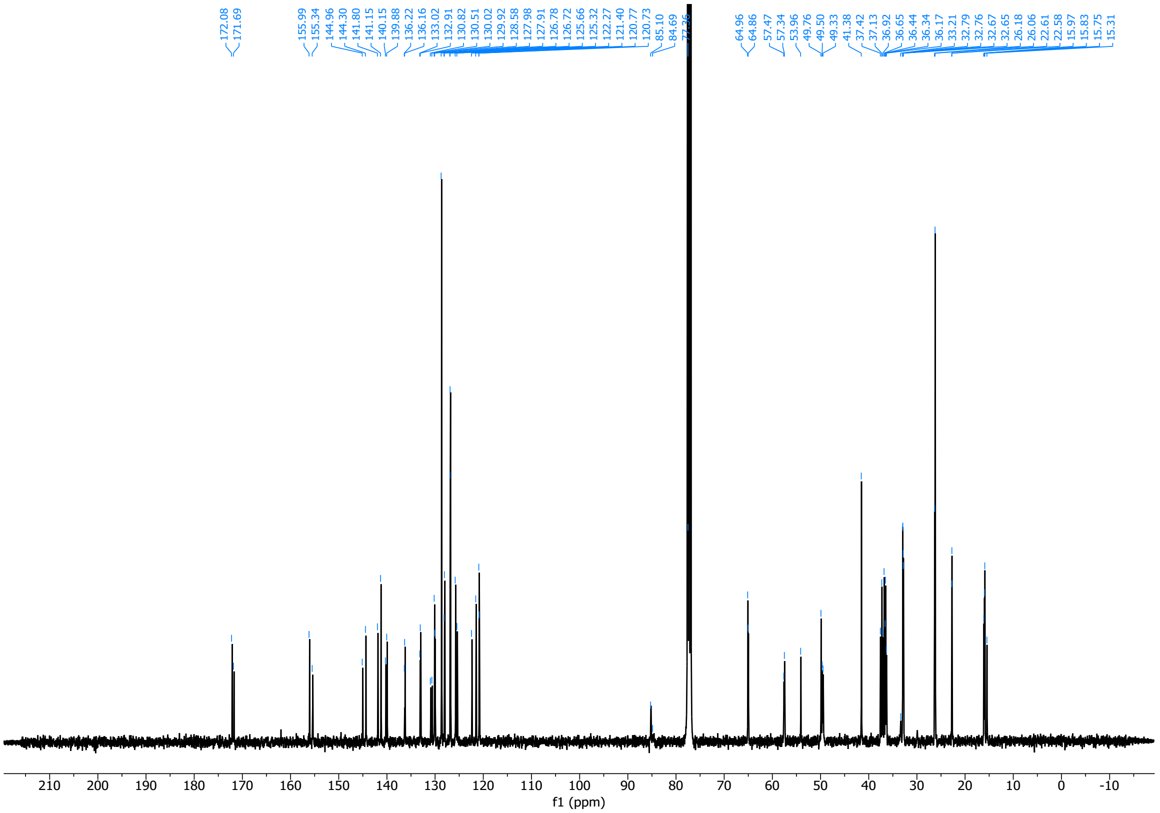

*N*-(((4*S*,5*S*)-8-((*E*)-2-cyclohexylvinyl)-2-((*R*)-1-hydroxypropan-2-yl)-4-methyl-1,1-dioxido-2,3,4,5-tetrahydrobenzo[*b*][1,4,5]oxathiazocin-5-yl)methyl)-*N*-methyl-2-(3-methylthiophen-2-yl)cyclopropane-1-carboxamide (**36**)

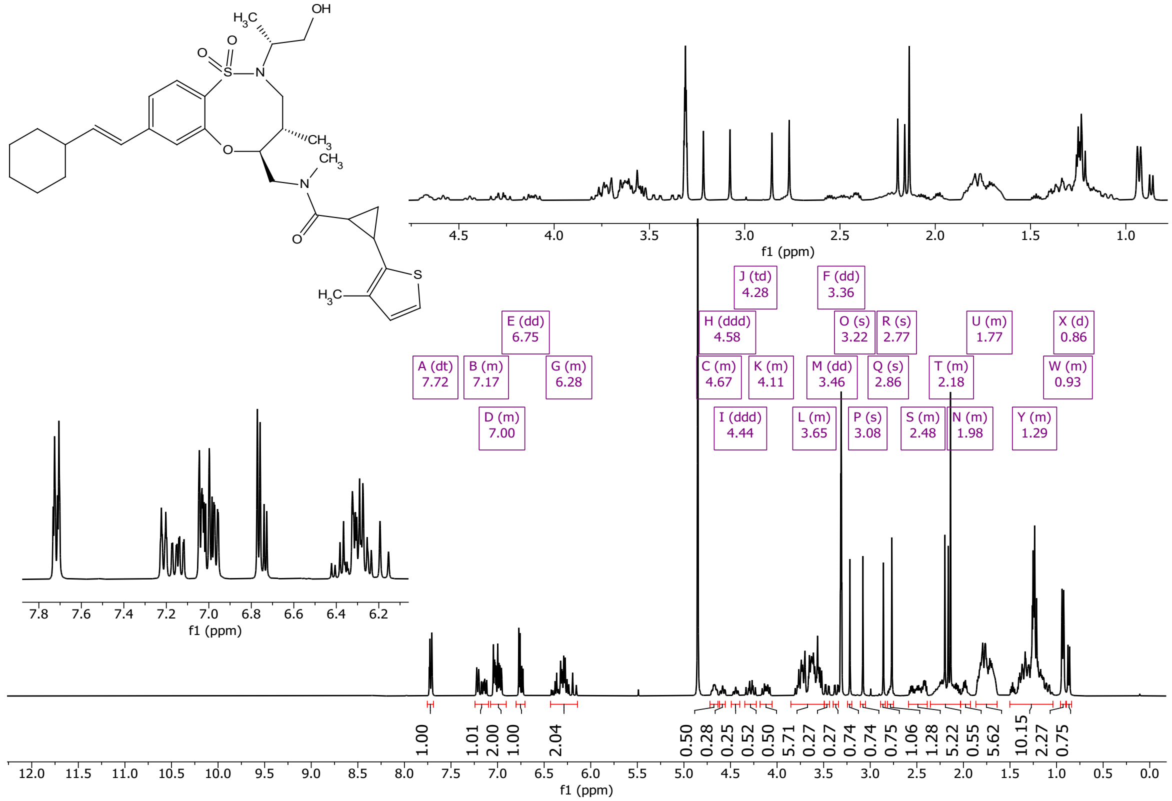

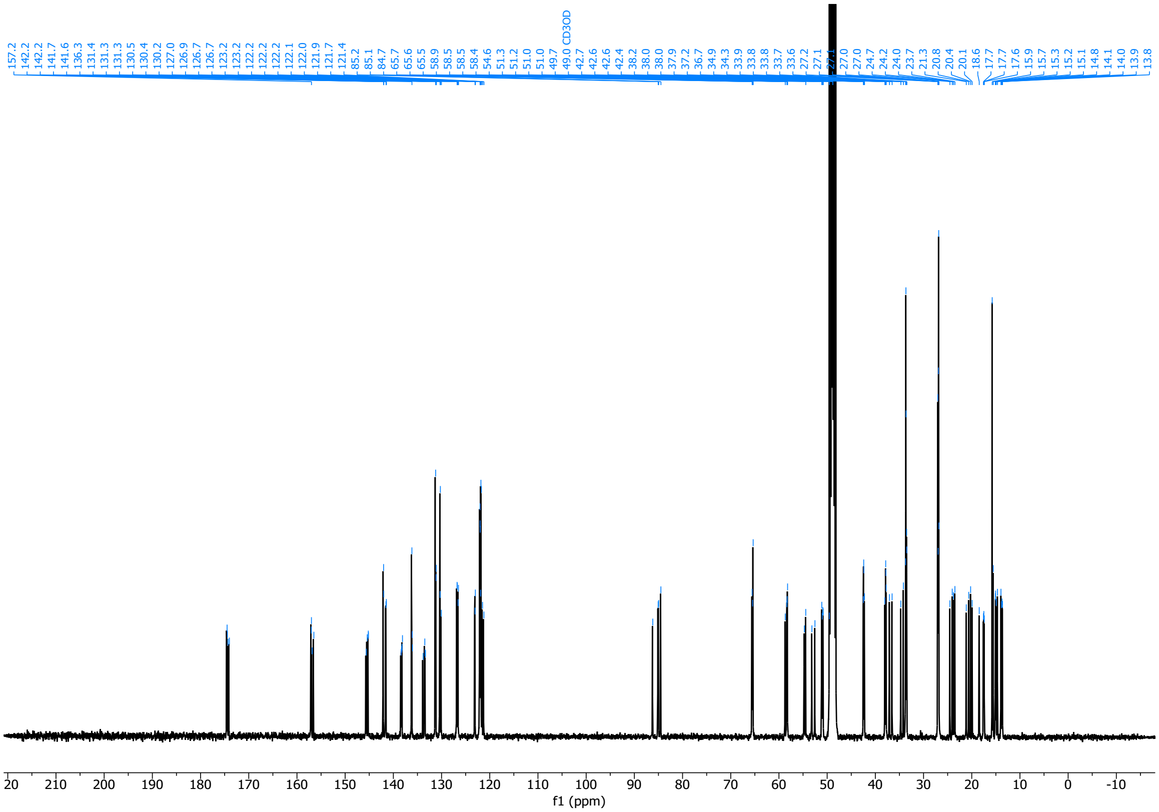

*N*-(((4*S*,5*S*)-8-(2-chloro-5-(hydroxymethyl)phenyl)-2-((*R*)-1-hydroxypropan-2-yl)-4-methyl-1,1-dioxido-2,3,4,5-tetrahydrobenzo[*b*][1,4,5]oxathiazocin-5-yl)methyl)-6-ethyl-*N*-methyl-2-oxo-1,2-dihydropyridine-4-carboxamide (**37**)

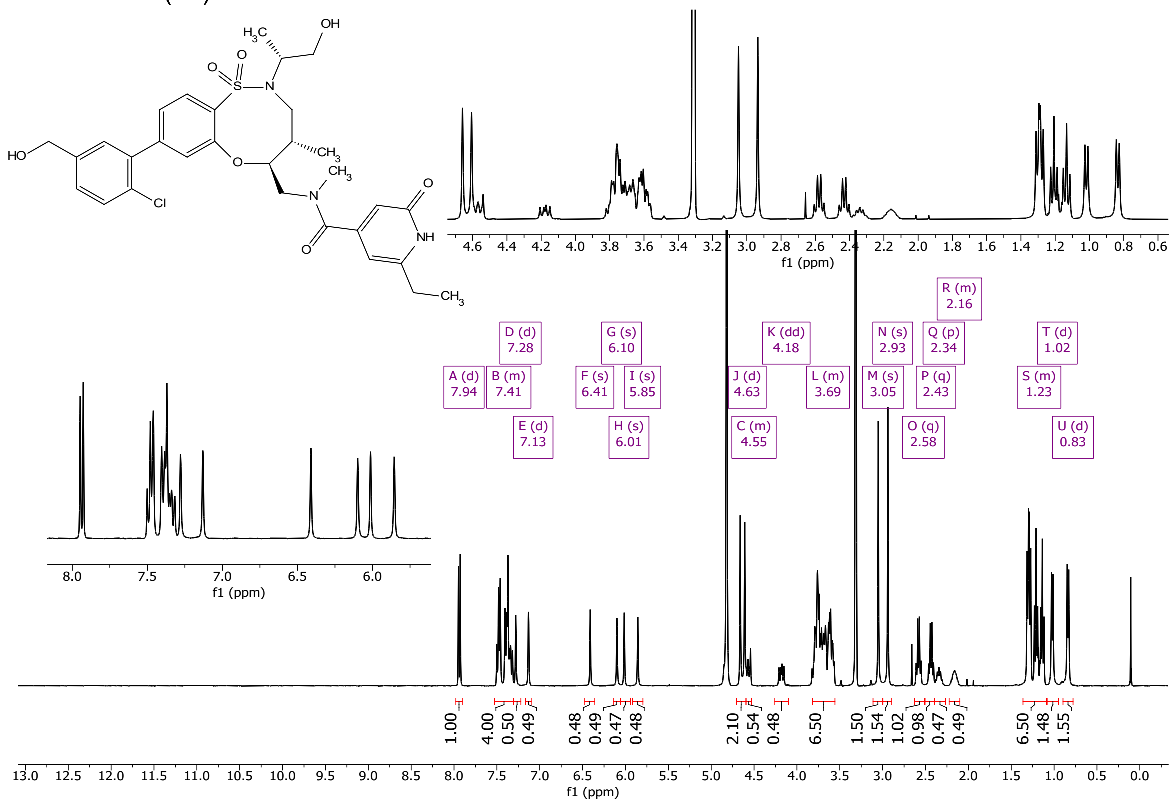

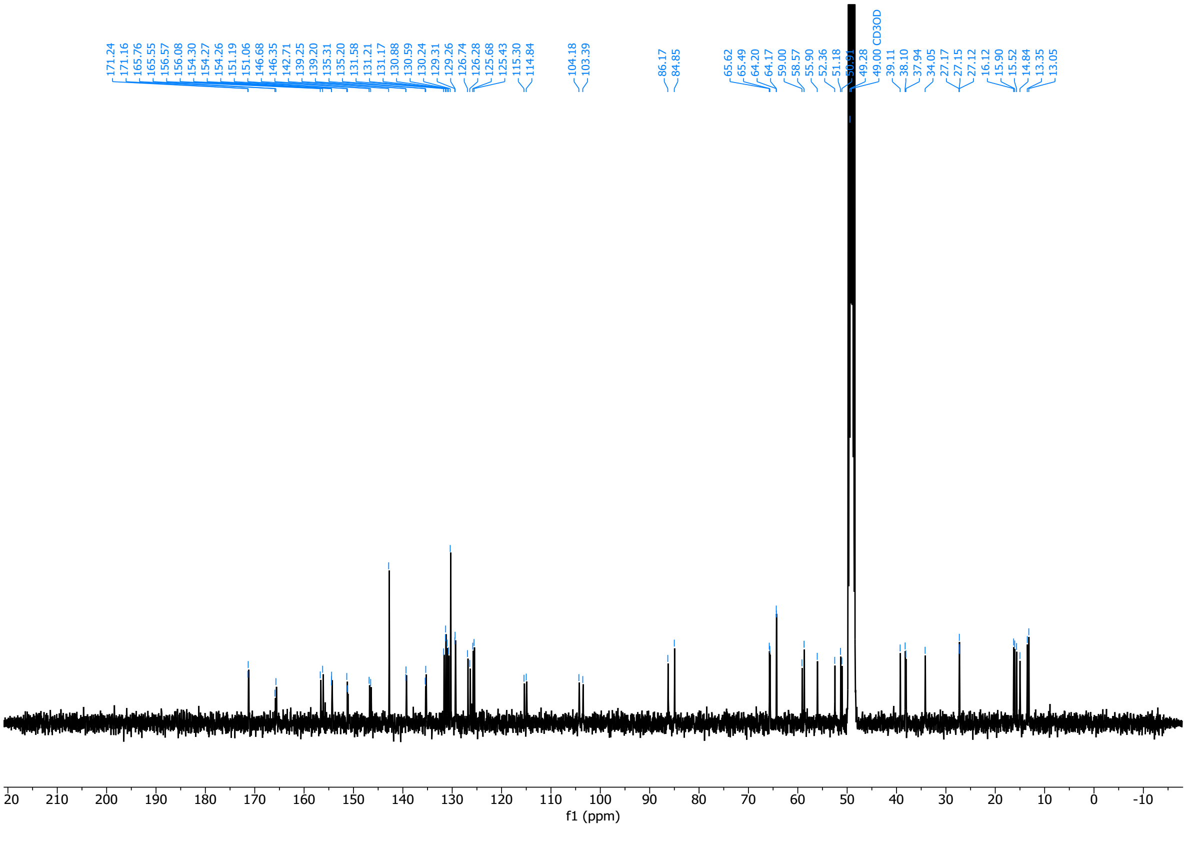

*N*-(((4*R*,5*R*)-8-((*E*)-2-cyclohexylvinyl)-2-((*S*)-1-hydroxypropan-2-yl)-4-methyl-1,1-dioxido-2,3,4,5-tetrahydrobenzo[*b*][1,4,5]oxathiazocin-5-yl)methyl)-*N*-methyl-2-phenylcyclopent-1-ene-1-carboxamide (**38**)

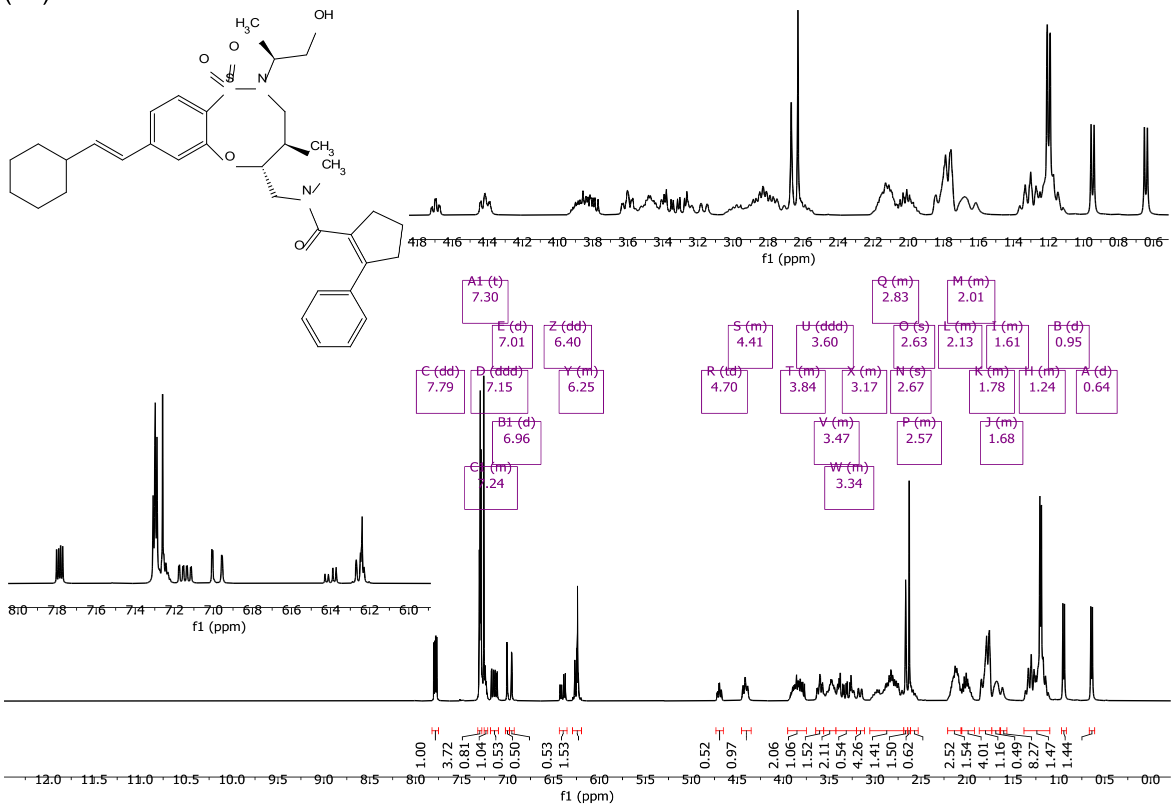

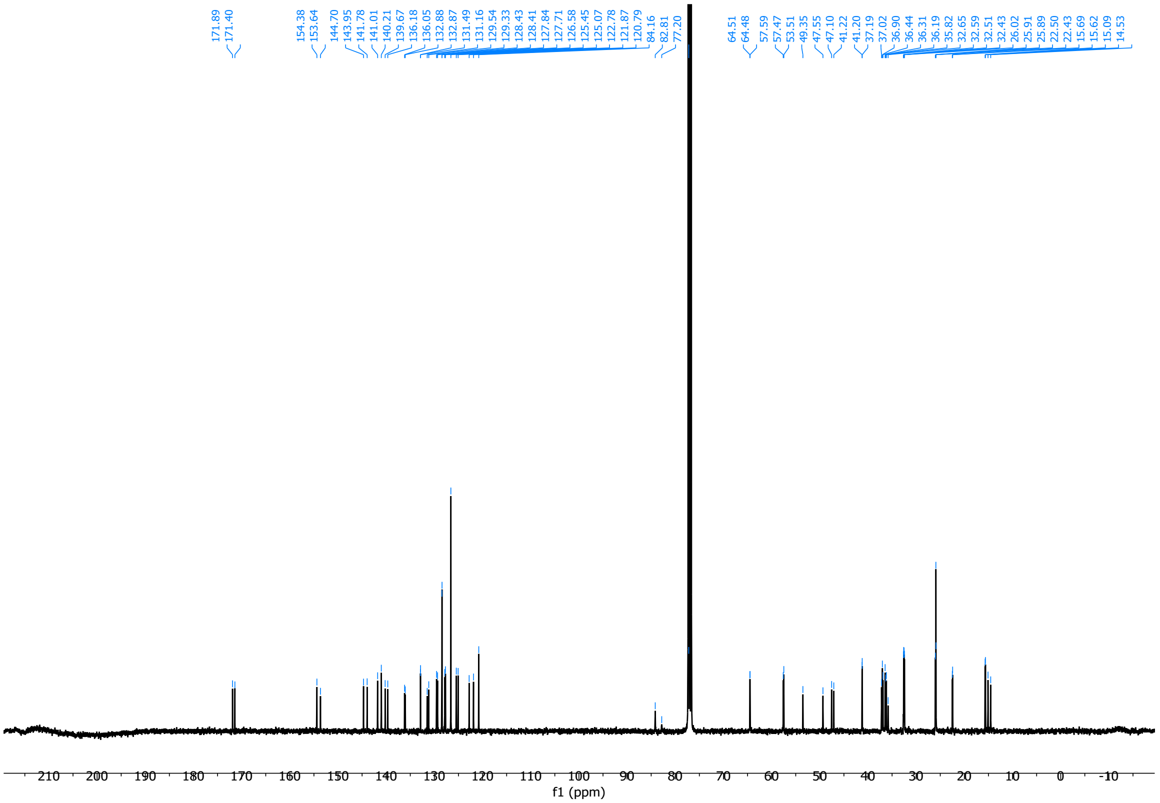

*N*-(((4*R*,5*R*)-8-((*E*)-2-cyclohexylvinyl)-2-((*S*)-1-hydroxypropan-2-yl)-4-methyl-1,1-dioxido-2,3,4,5-tetrahydrobenzo[*b*][1,4,5]oxathiazocin-5-yl)methyl)-2-(2,3-dihydrobenzofuran-3-yl)-*N*-methylacetamide (**39**)

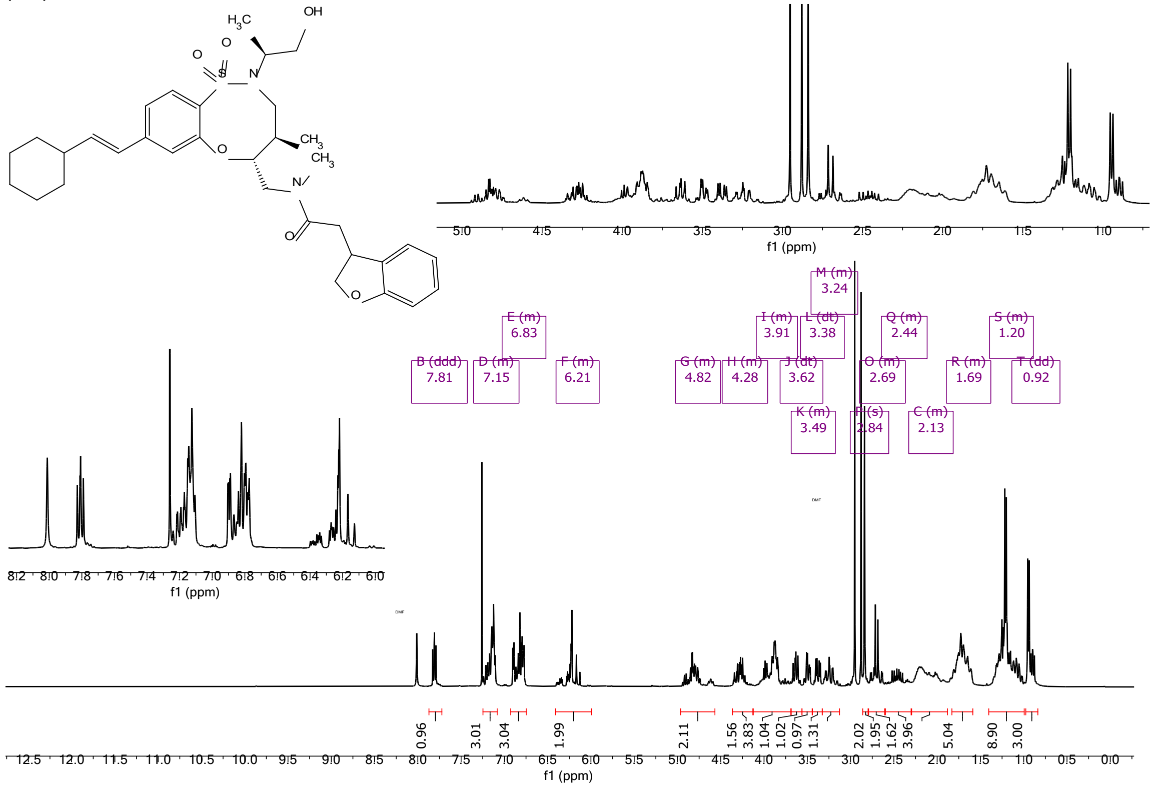

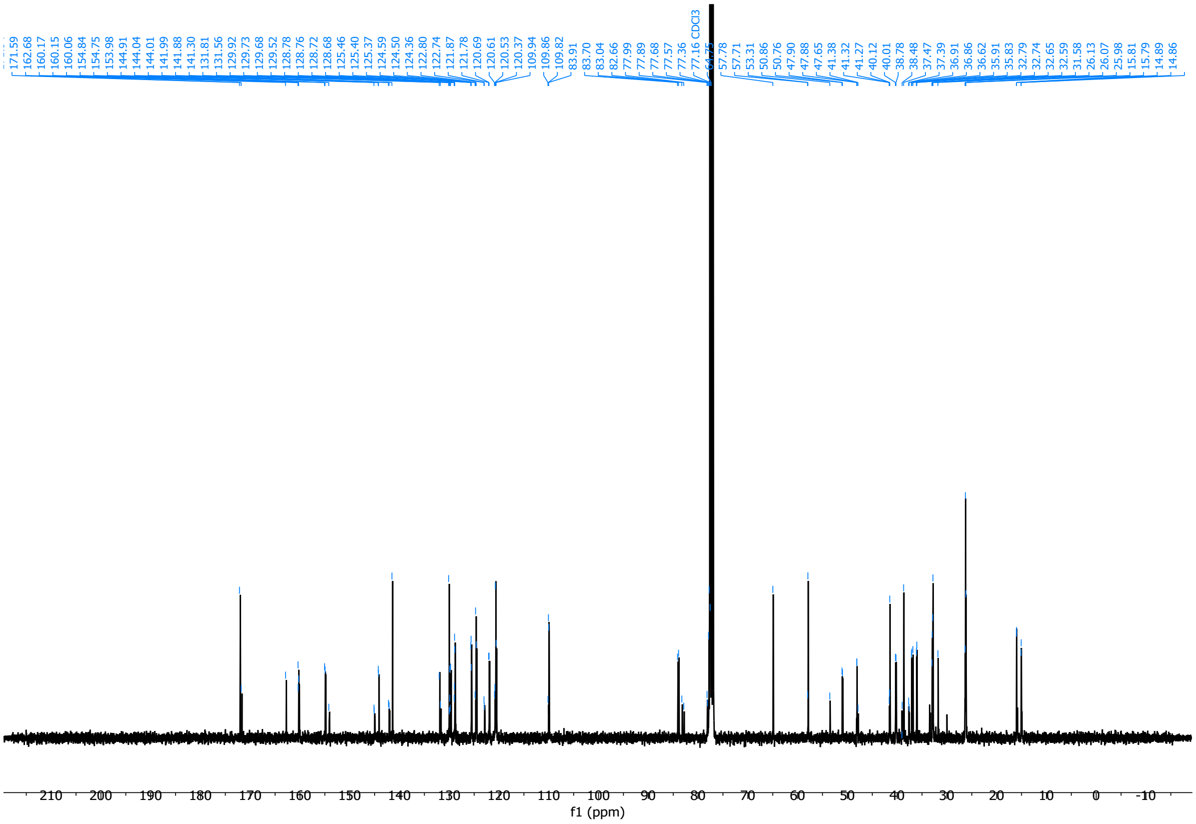

3-(benzo[*d*][1,3]dioxol-5-yl)-*N*-(((4*R*,5*R*)-2-((*S*)-1-hydroxypropan-2-yl)-4-methyl-1,1-dioxido-8-(1-oxo-2,3,4,5-tetrahydro-1H-benzo[*c*]azepin-7-yl)-2,3,4,5-tetrahydrobenzo[*b*][1,4,5]oxathiazocin-5-yl)methyl)-*N*-methylpropiolamide (**40**)

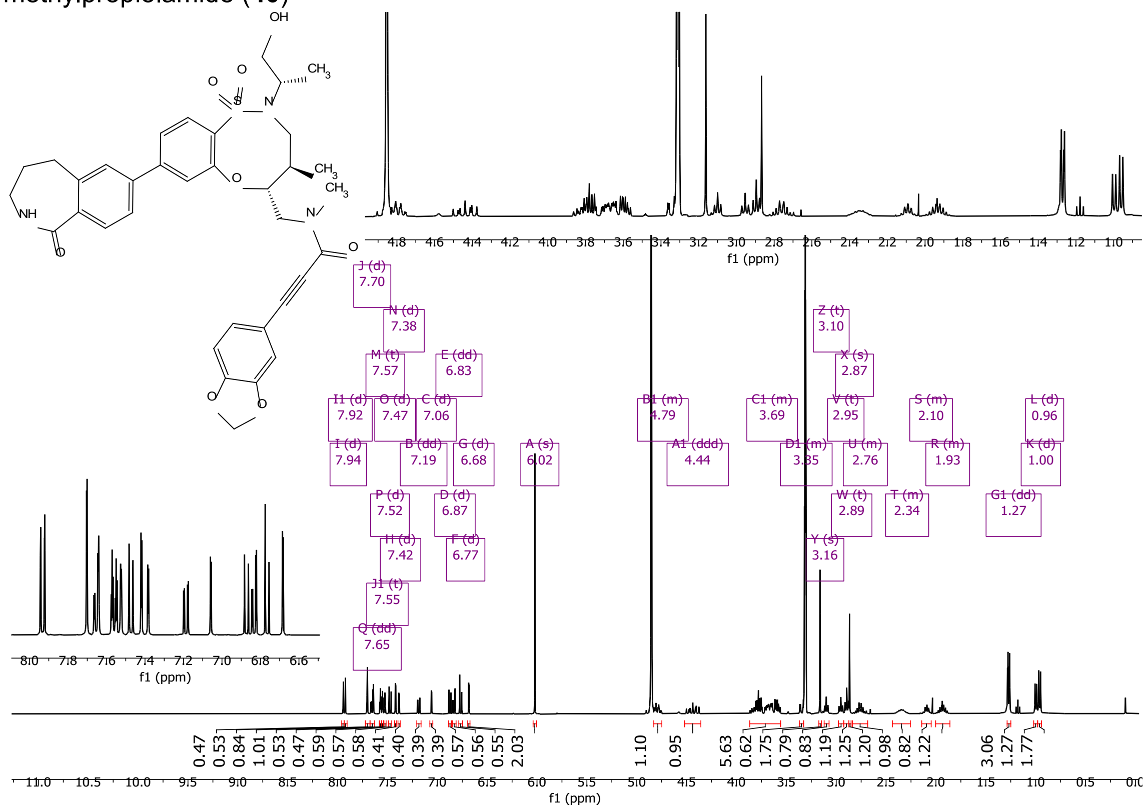

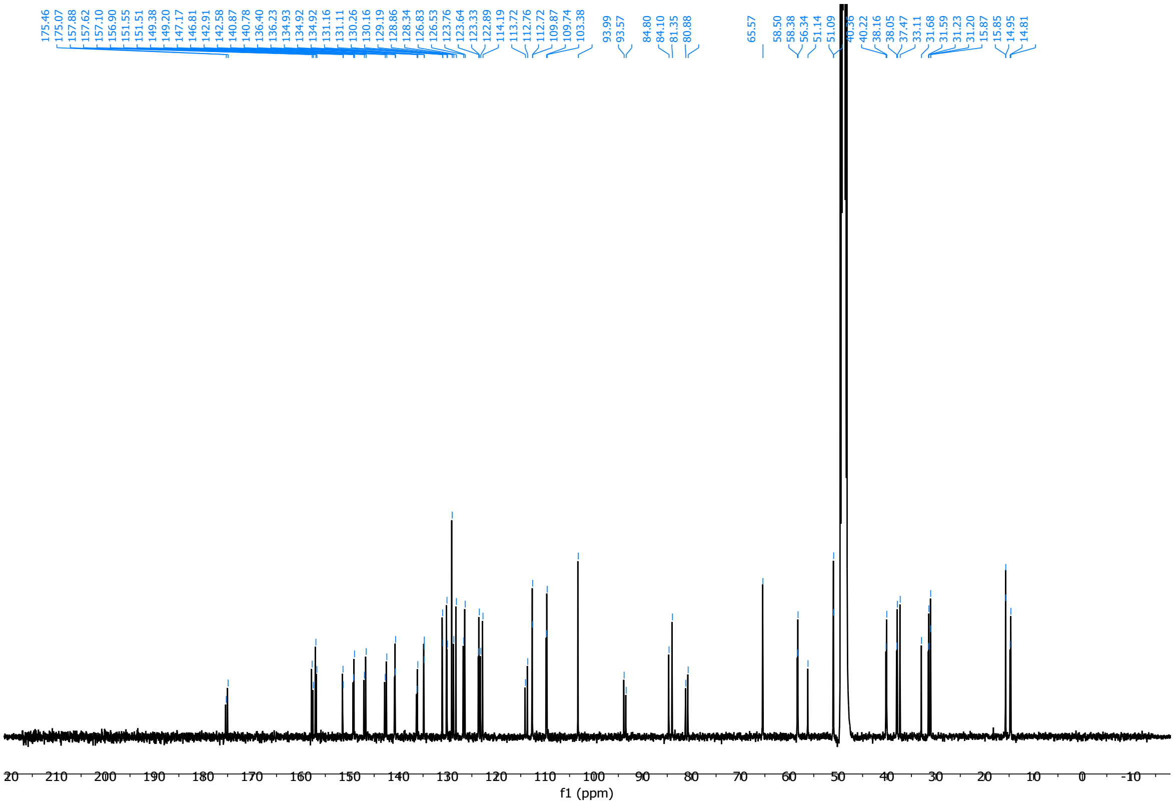

3-(Benzo[*d*][1,3]dioxol-5-yl)-*N*-(((4*R*,5*R*)-8-bromo-2-((*R*)-1-hydroxypropan-2-yl)-4-methyl-1,1-dioxido-2,3,4,5-tetrahydrobenzo[*b*][1,4,5]oxathiazocin-5-yl)methyl)-*N*-methylpropiolamide (**41**)

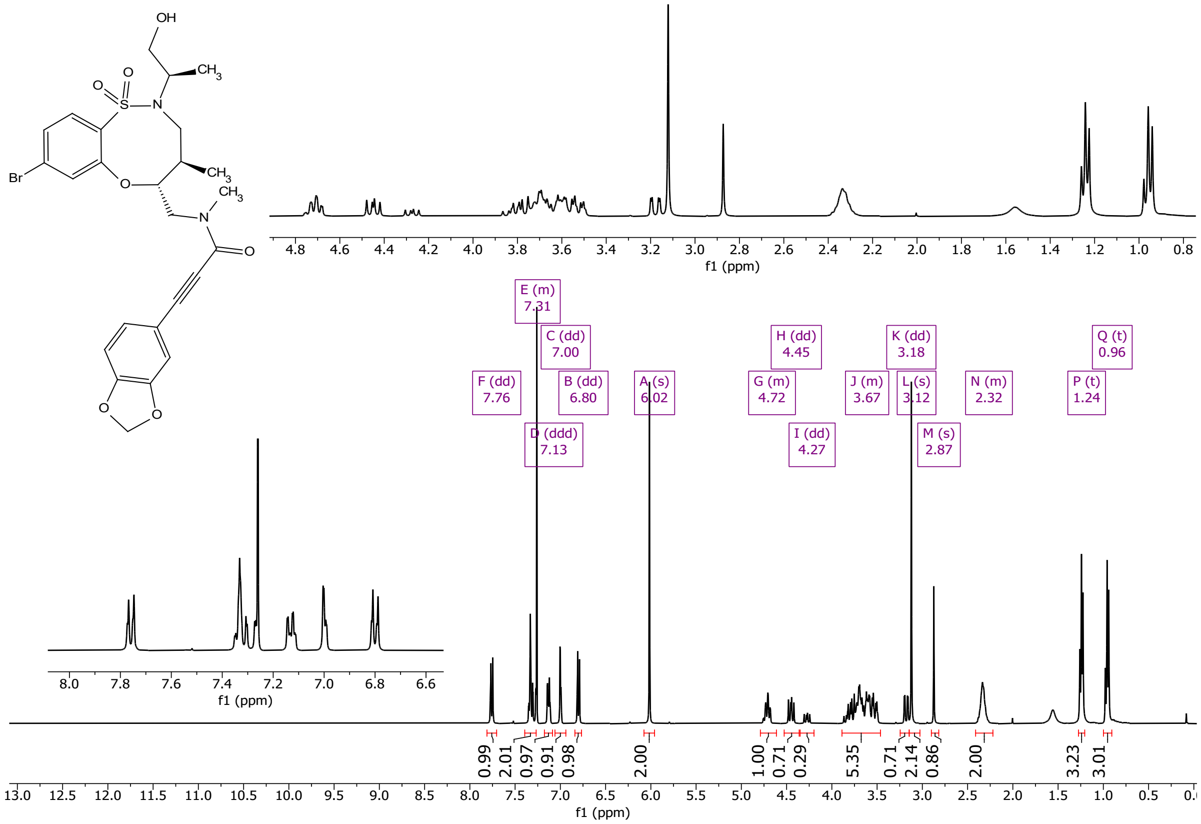

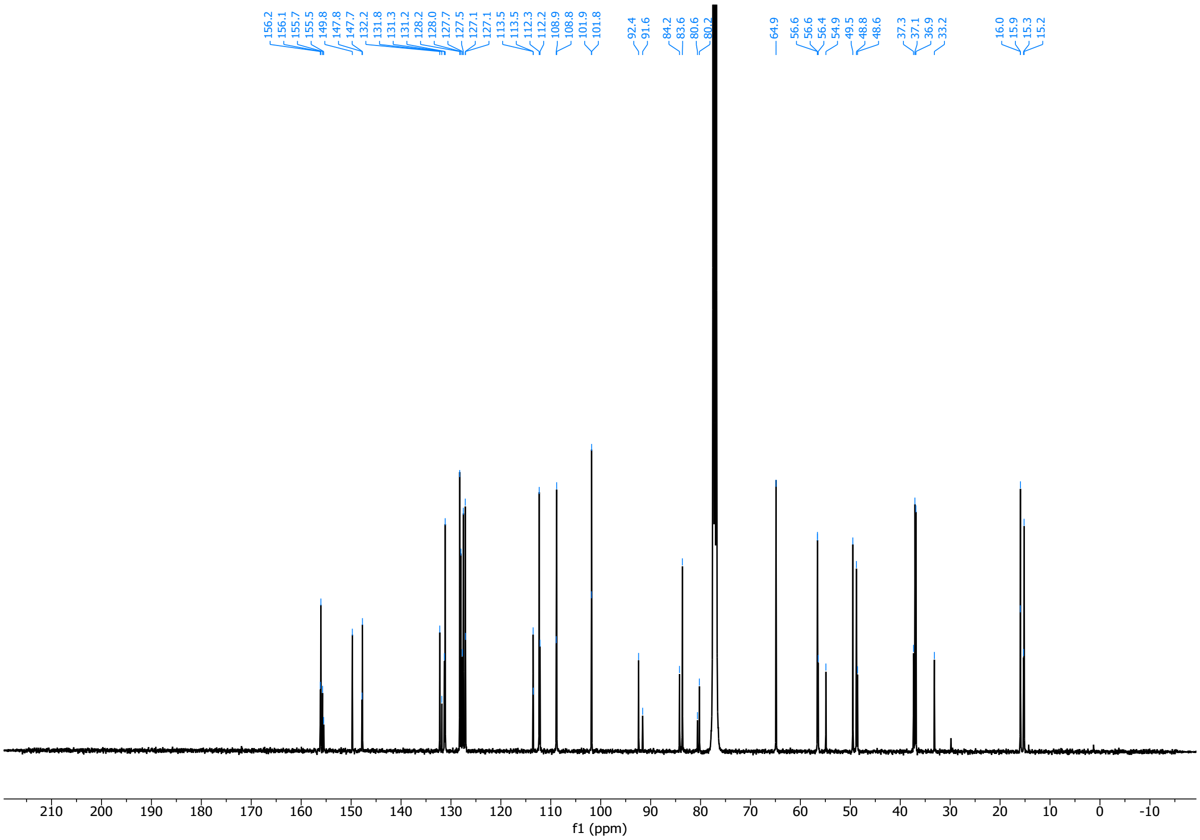
156.2
156.1
155.7
155.5
149.8
147.8
147.7
132.2
131.8
131.3
131.2
128.2
128.0
127.7
127.5
127.1
127.1
113.5
113.5
112.3
112.2
108.9
108.8
101.9
101.8
92.4
91.6
84.2
83.6
80.6
80.2
64.9
56.6
56.6
56.4
54.9
49.5
48.8
48.6
37.3
37.1
36.9
33.2
16.0
15.9
15.3
15.2
f1 (ppm)

3181

3-(Benzo[*d*][1,3]dioxol-5-yl)-*N*-(((4*R*,5*R*)-8-bromo-2-((*S*)-1-hydroxypropan-2-yl)-4-methyl-1,1-dioxido-2,3,4,5-tetrahydrobenzo[*b*][1,4,5]oxathiazocin-5-yl)methyl)-*N*-methylpropiolamide (**42**)

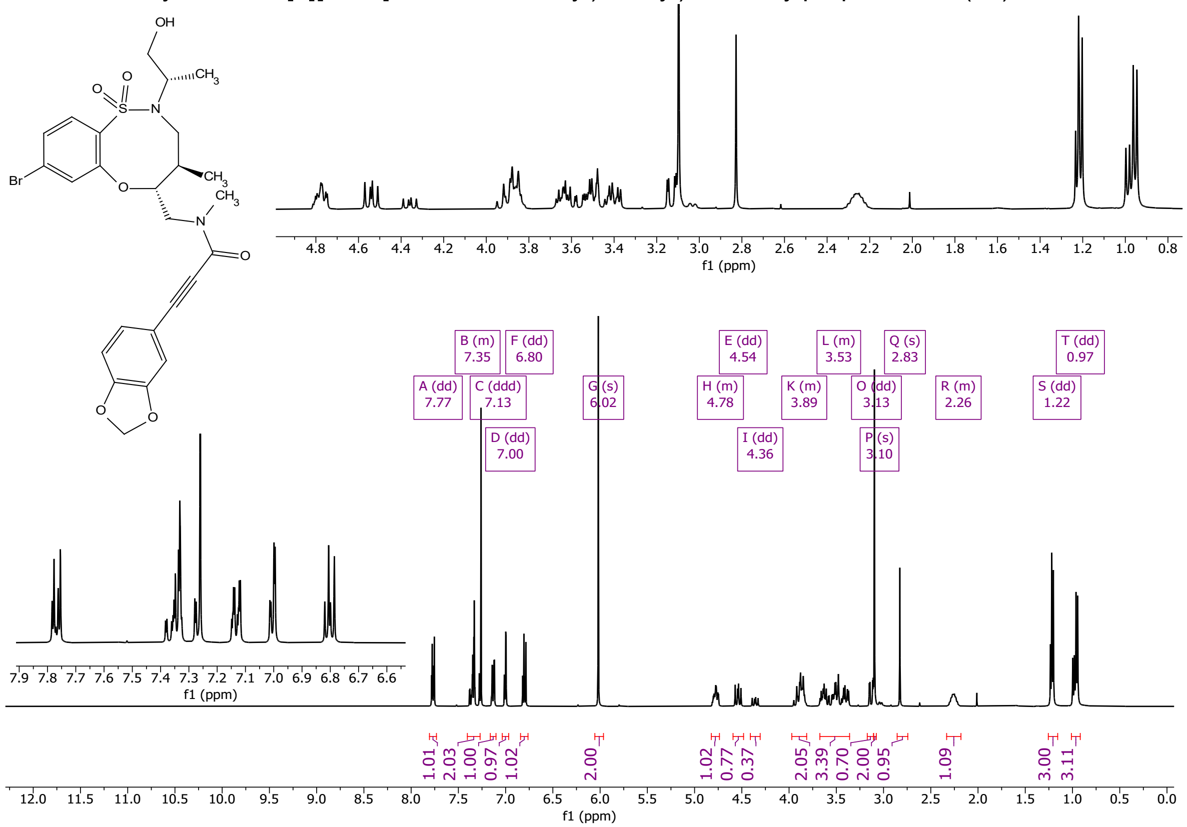

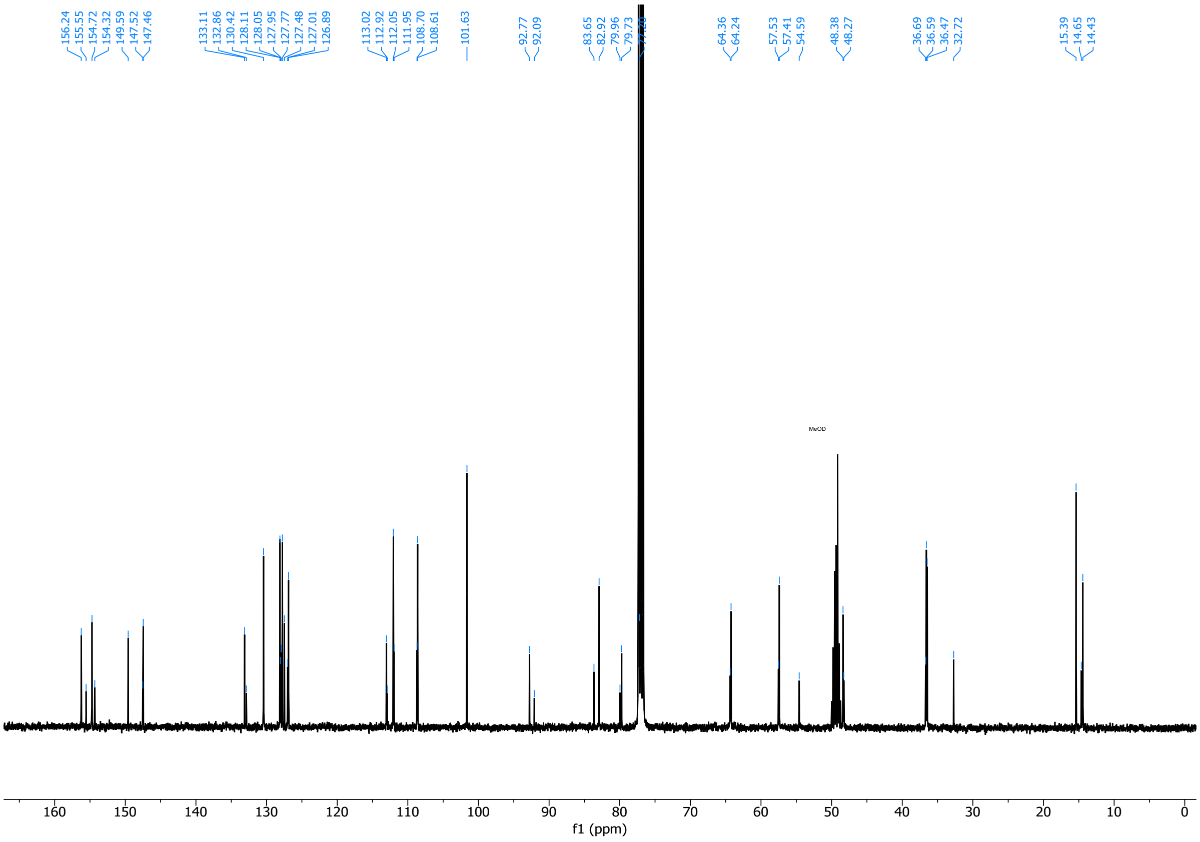
156.24
155.55
154.72
154.32
149.59
147.52
147.46
133.11
132.86
130.42
128.11
128.05
127.95
127.77
127.48
127.01
126.89
113.02
112.92
112.05
111.95
108.70
108.61
101.63
92.77
92.09
83.65
82.92
79.96
79.73
64.36
64.24
57.53
57.41
54.59
48.38
48.27
36.69
36.59
36.47
32.72
15.39
14.65
14.43
MeOD
160
150
140
130
120
110
100
90
80
70
60
50
40
30
20
10
0
f1 (ppm)

3185

3-(Benzo[*d*][1,3]dioxol-5-yl)-1-((8*R*,9*R*,10*S*)-9-(4-bromophenyl)-10-(hydroxymethyl)-1,6-diazabicyclo[6.2.0]decan-6-yl)prop-2-yn-1-one (**43**)

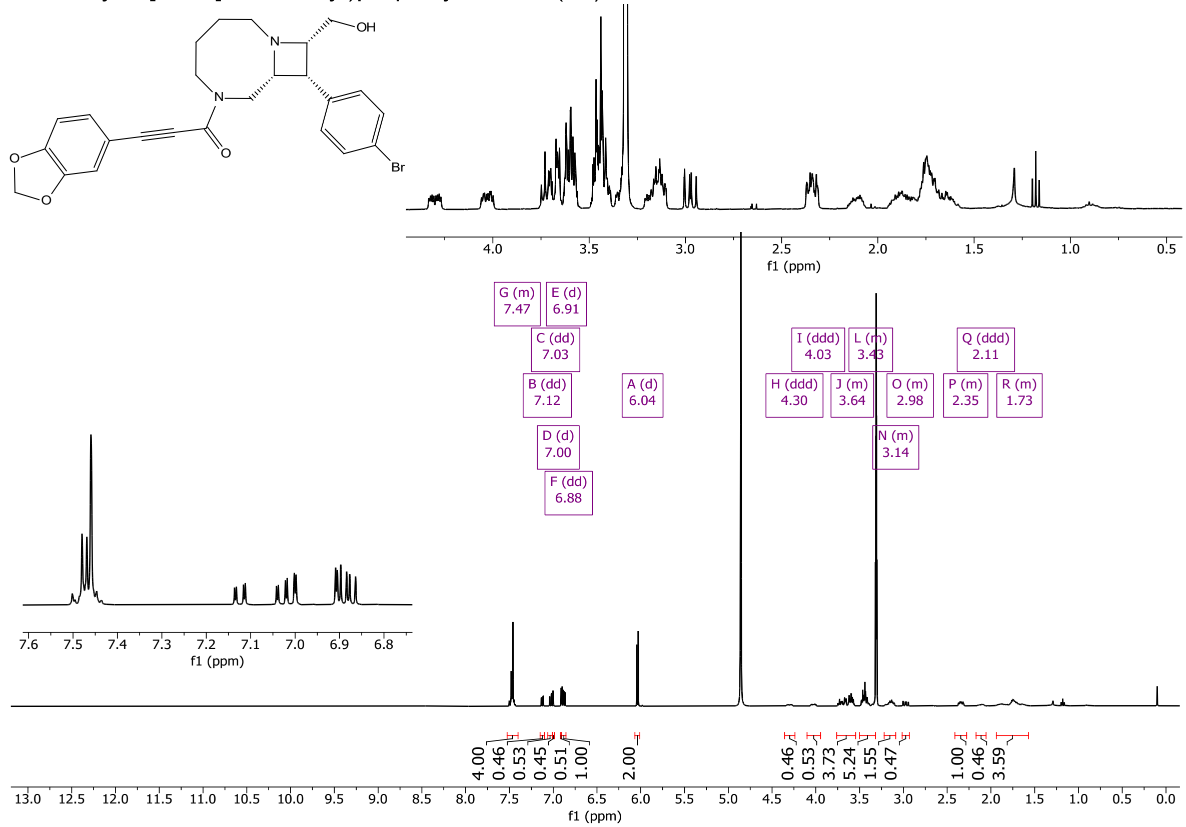

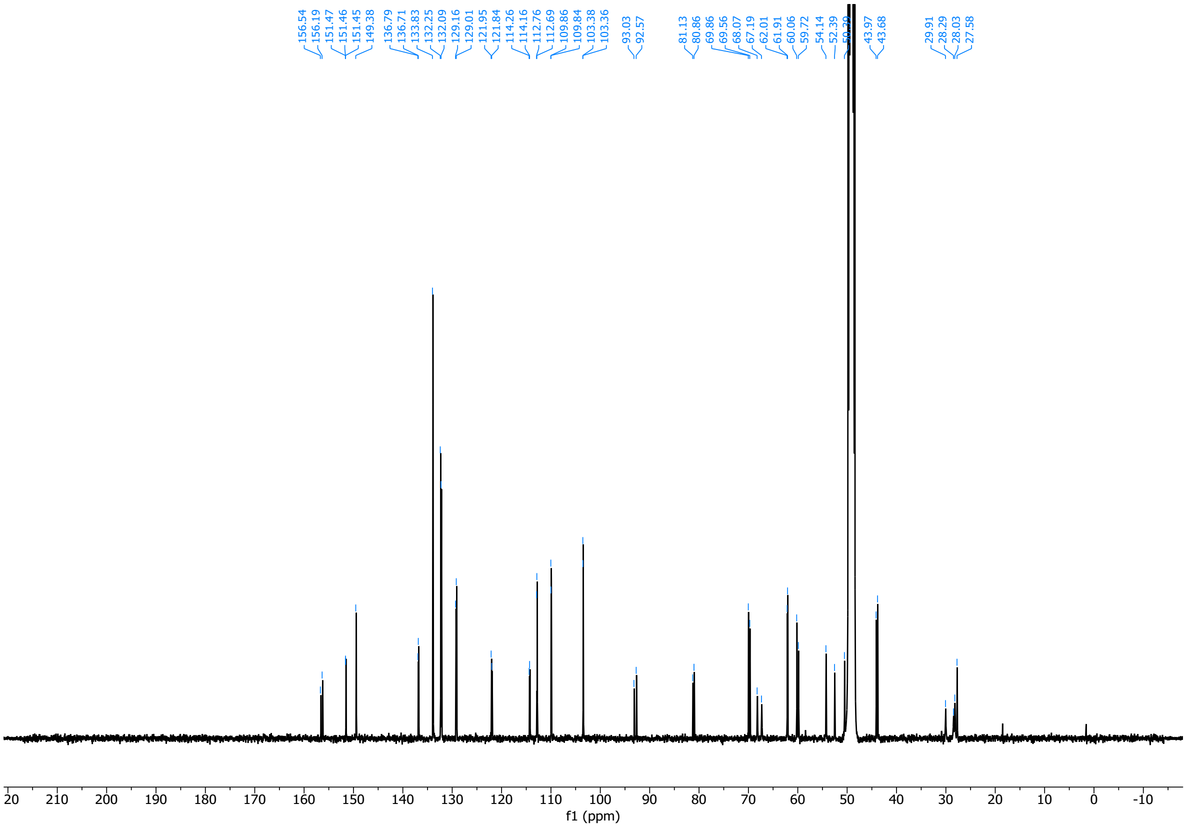

3-(Benzo[*d*][1,3]dioxol-5-yl)-1-((8*S*,9*S*,10*S*)-9-(4-bromophenyl)-10-(hydroxymethyl)-1,6-diazabicyclo[6.2.0]decan-6-yl)prop-2-yn-1-one (**44**)

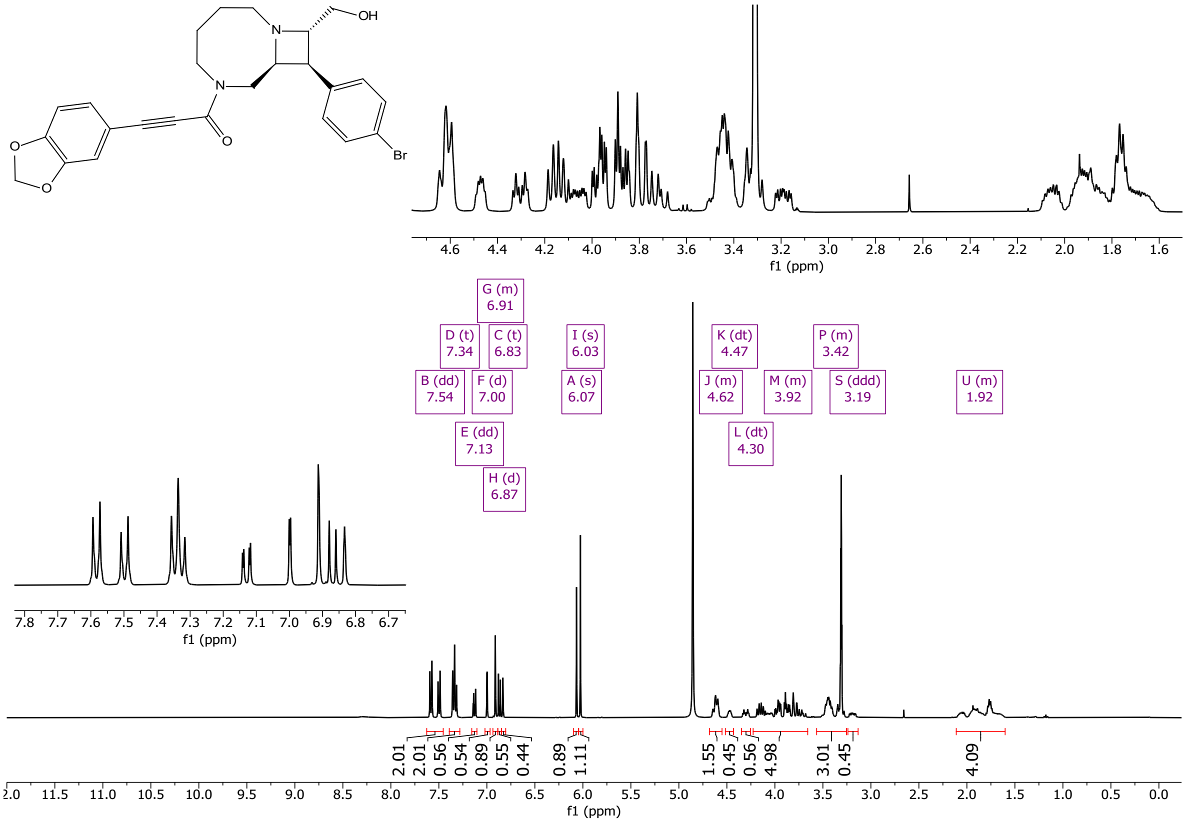

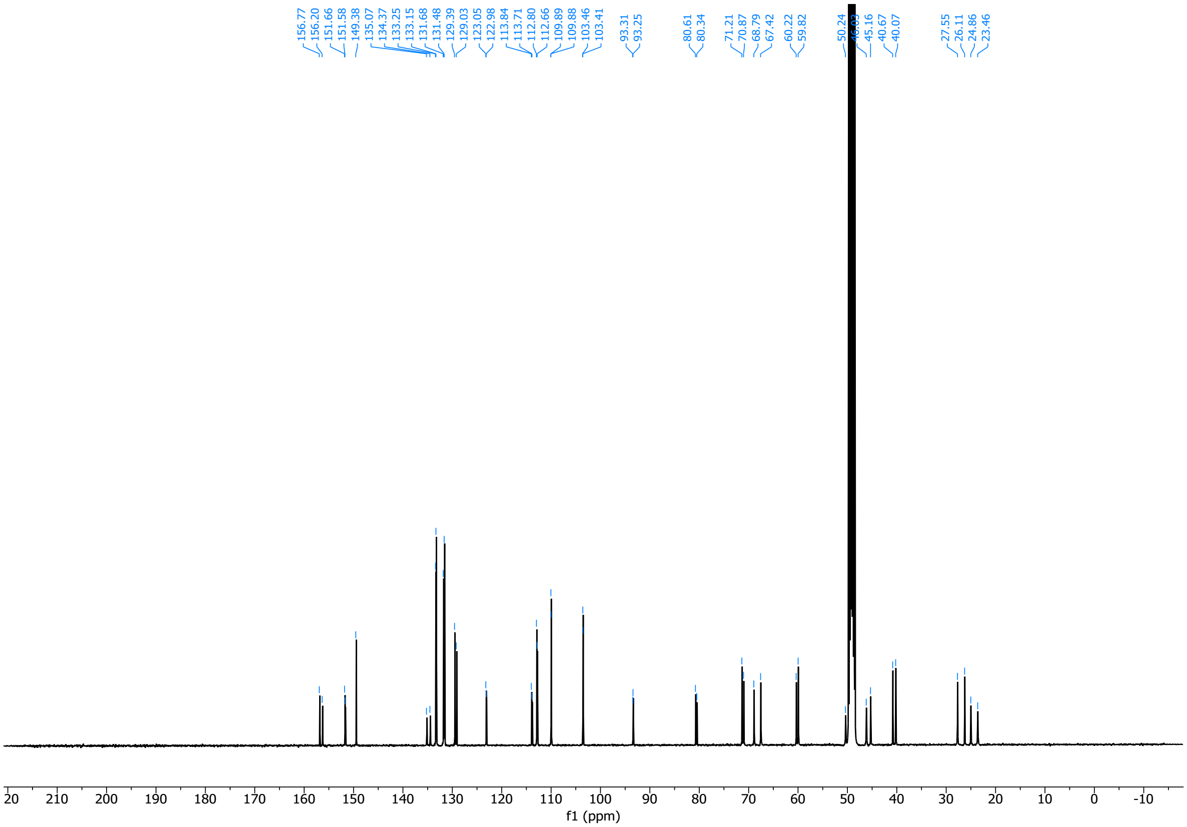

(2*R*,4*R*)-*N*-Methyl-1-((4-sulfamoylphenyl)sulfonyl)-4-(3-(1,2,3,6-tetrahydropyridin-4-yl)phenoxy)pyrrolidine-2-carboxamide (**45**)

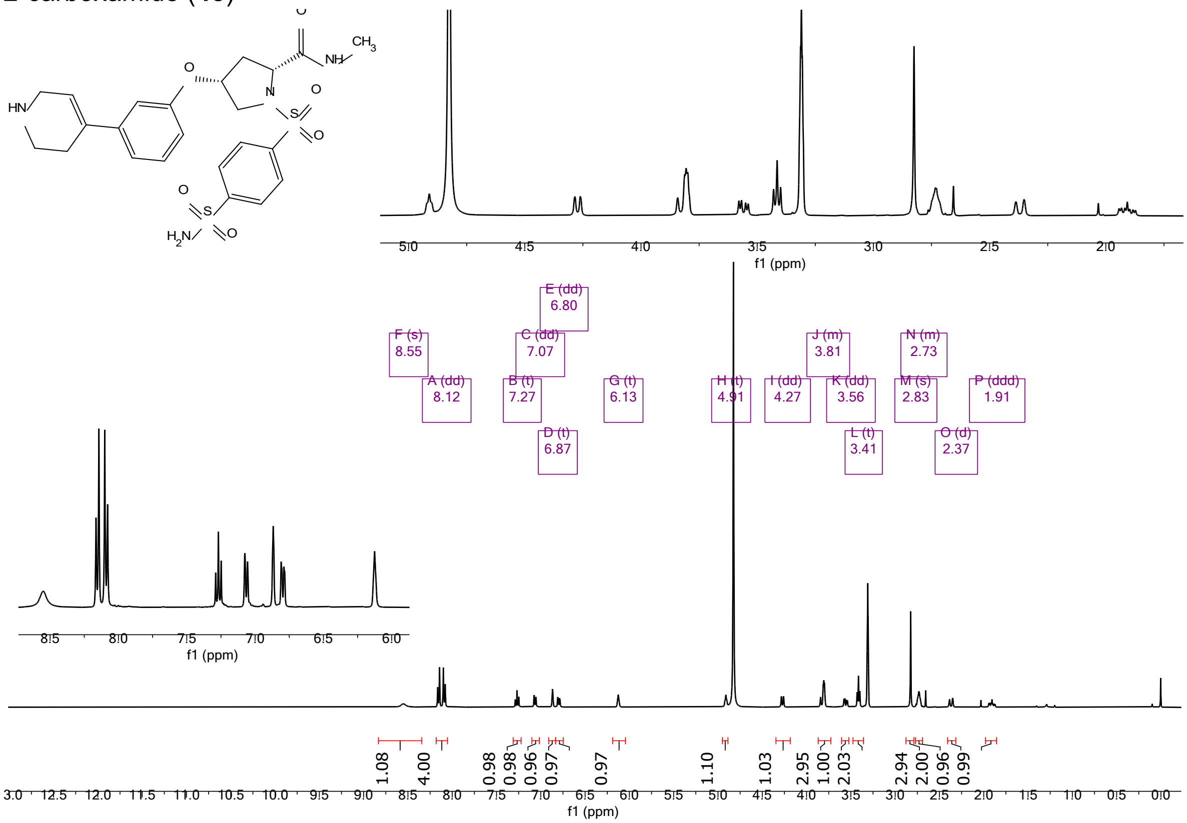

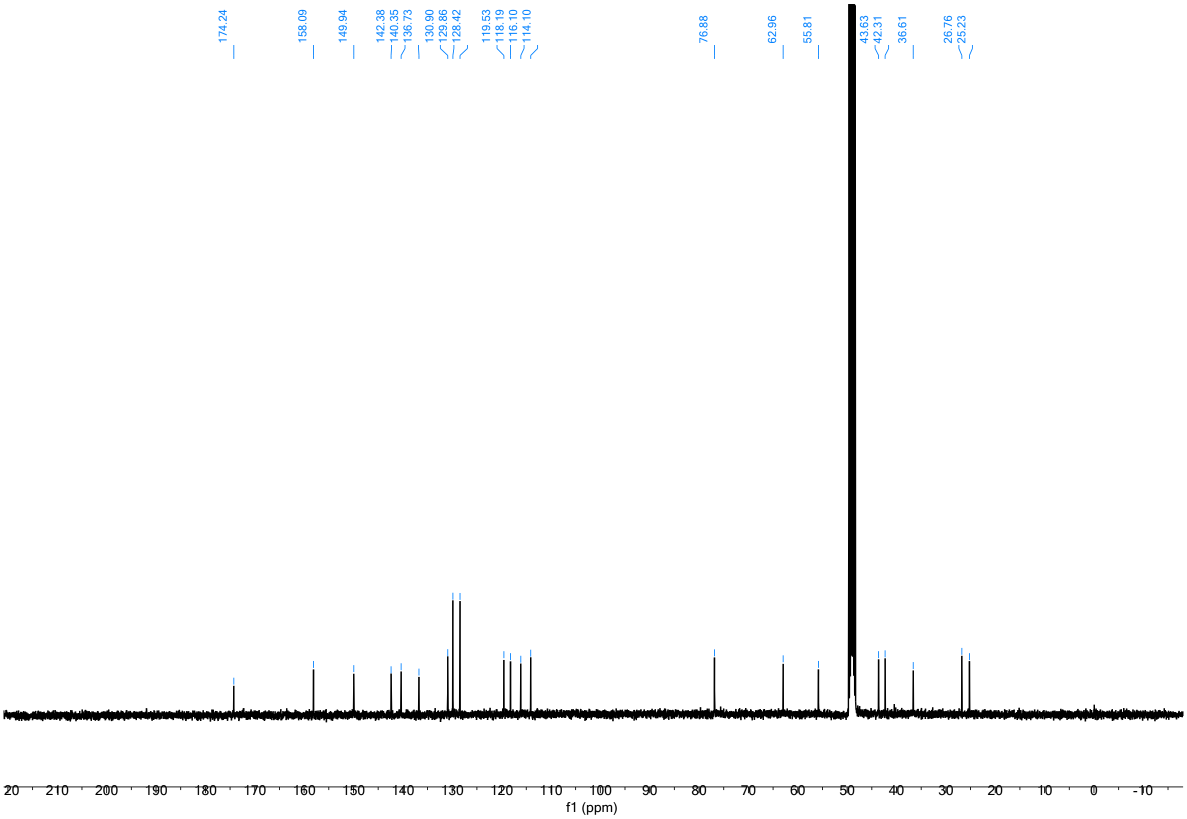

(*R*)-5-((2-acetyl-8-(3'-((dimethylamino)methyl)-4'-fluoro-[1,1'-biphenyl]-4-yl)-2,6-diazaspiro[3.4]octan-6-yl)methyl)furan-2-sulfonamide (**46**)

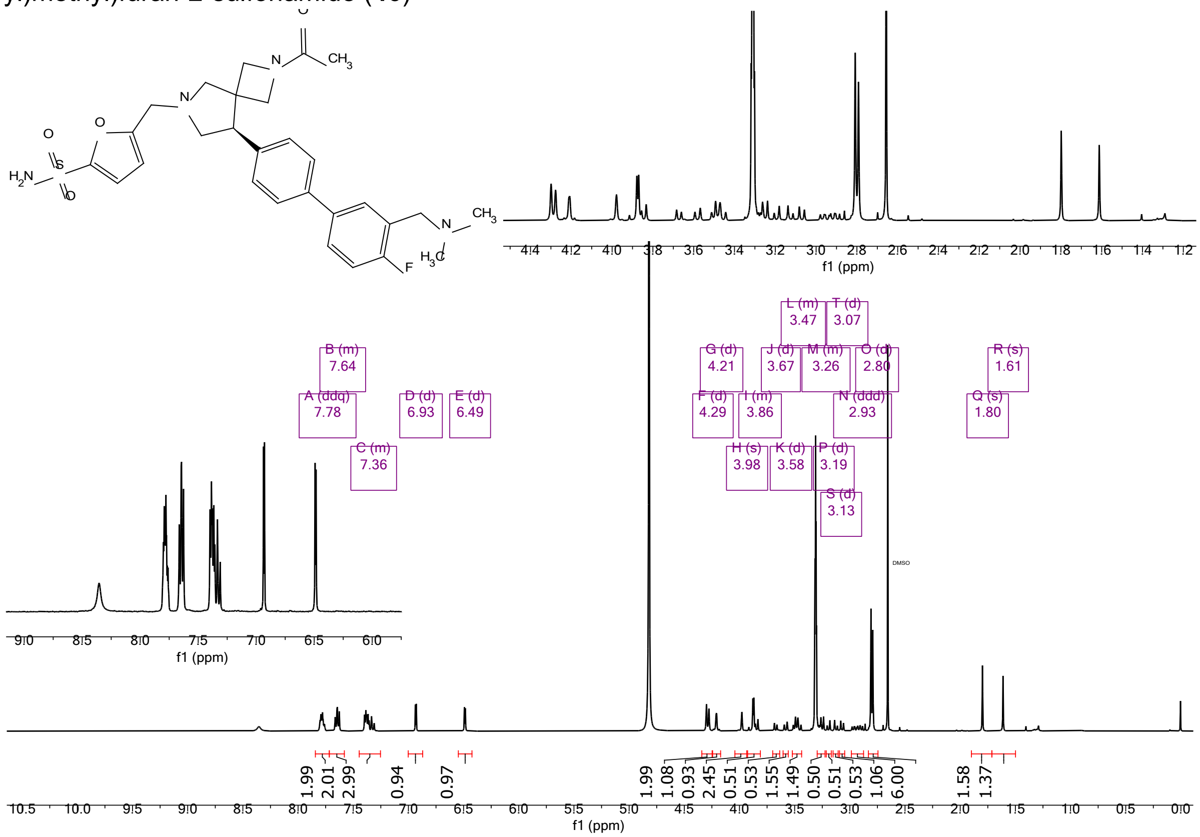

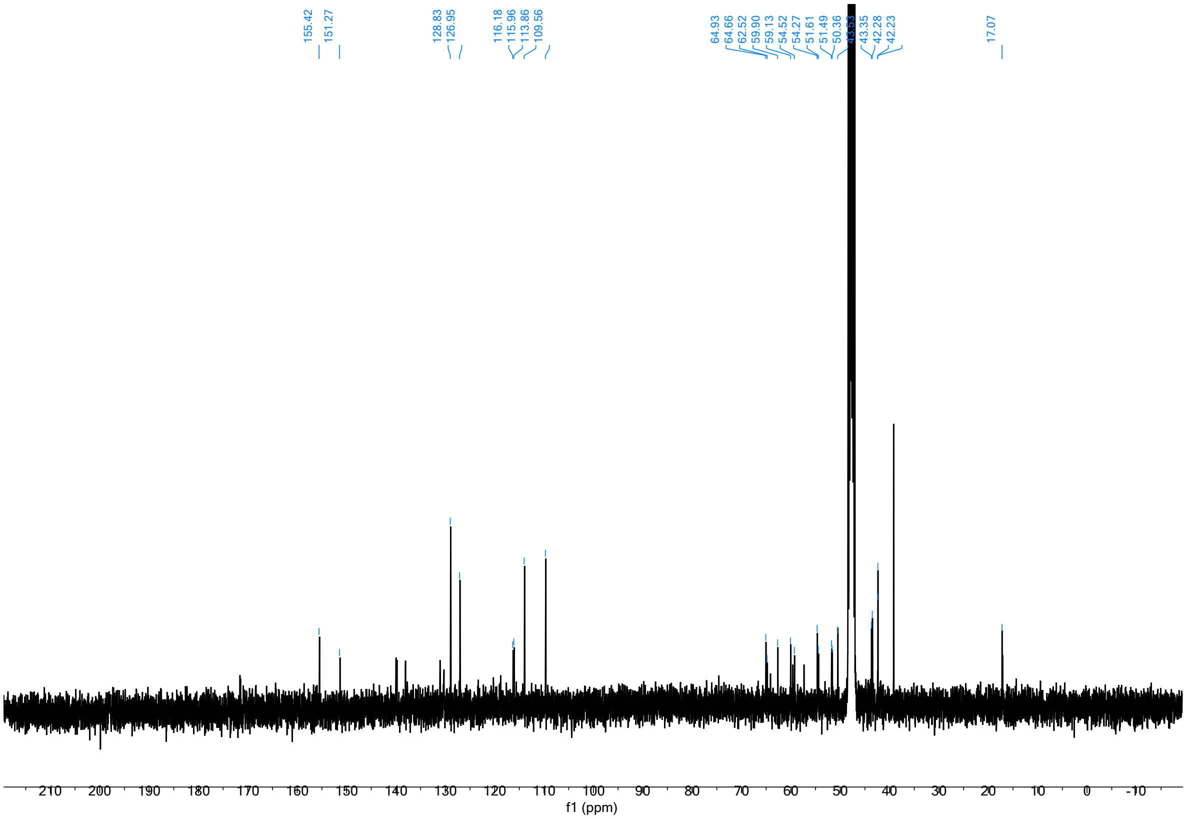

(2*S*,3*S*)-3-(3'-(ethylcarbamoyl)-[1,1'-biphenyl]-4-yl)-*N*-methyl-1-(4-sulfamoylbenzyl)pyrrolidine-2-carboxamide (**47**)

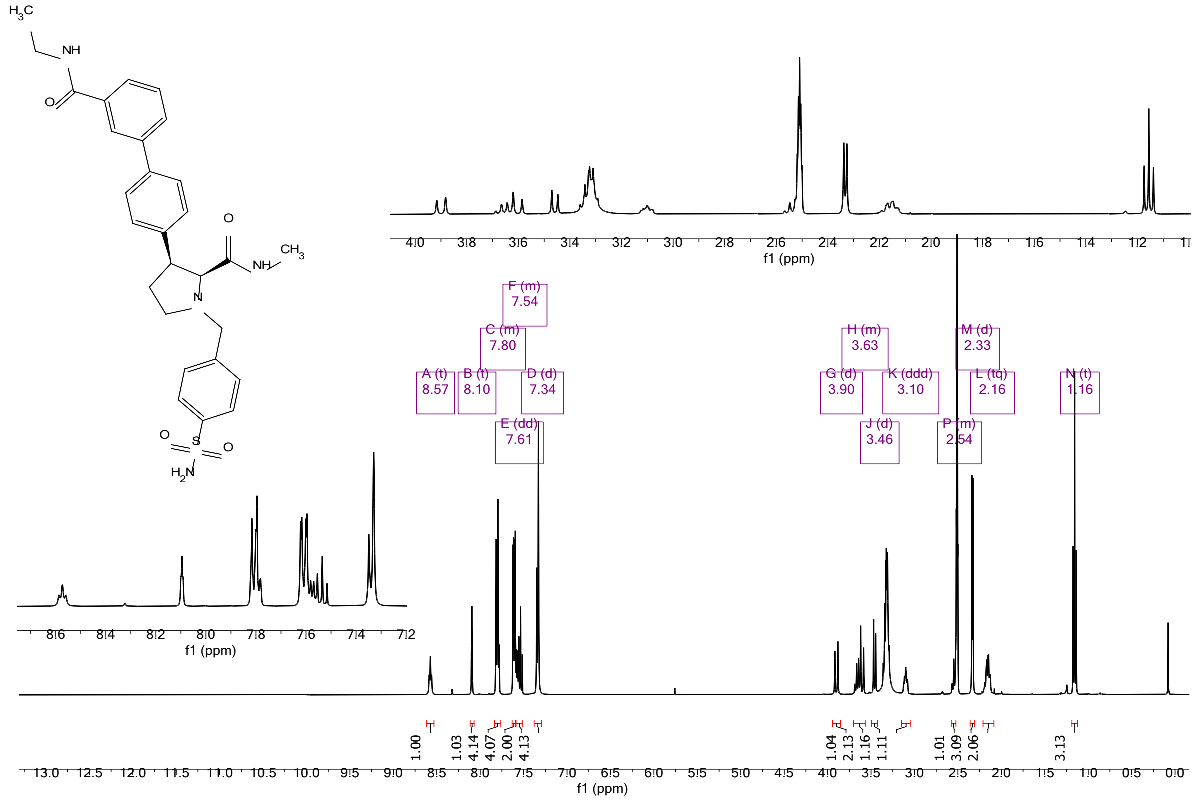

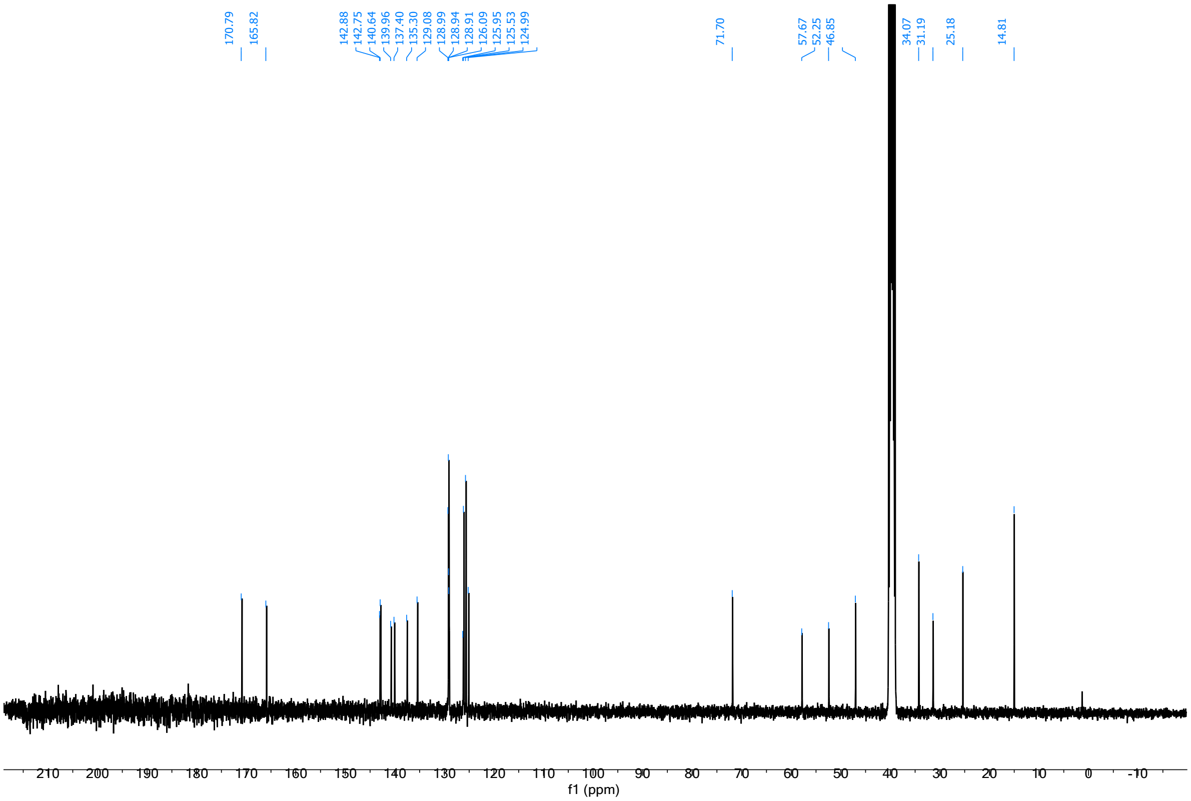

(2*R*,3*S*)-3-(3'-ethoxy-[1,1'-biphenyl]-3-yl)-*N*-methyl-1-((5-sulfamoylfuran-3-yl)methyl)pyrrolidine-2-carboxamide (**48**)

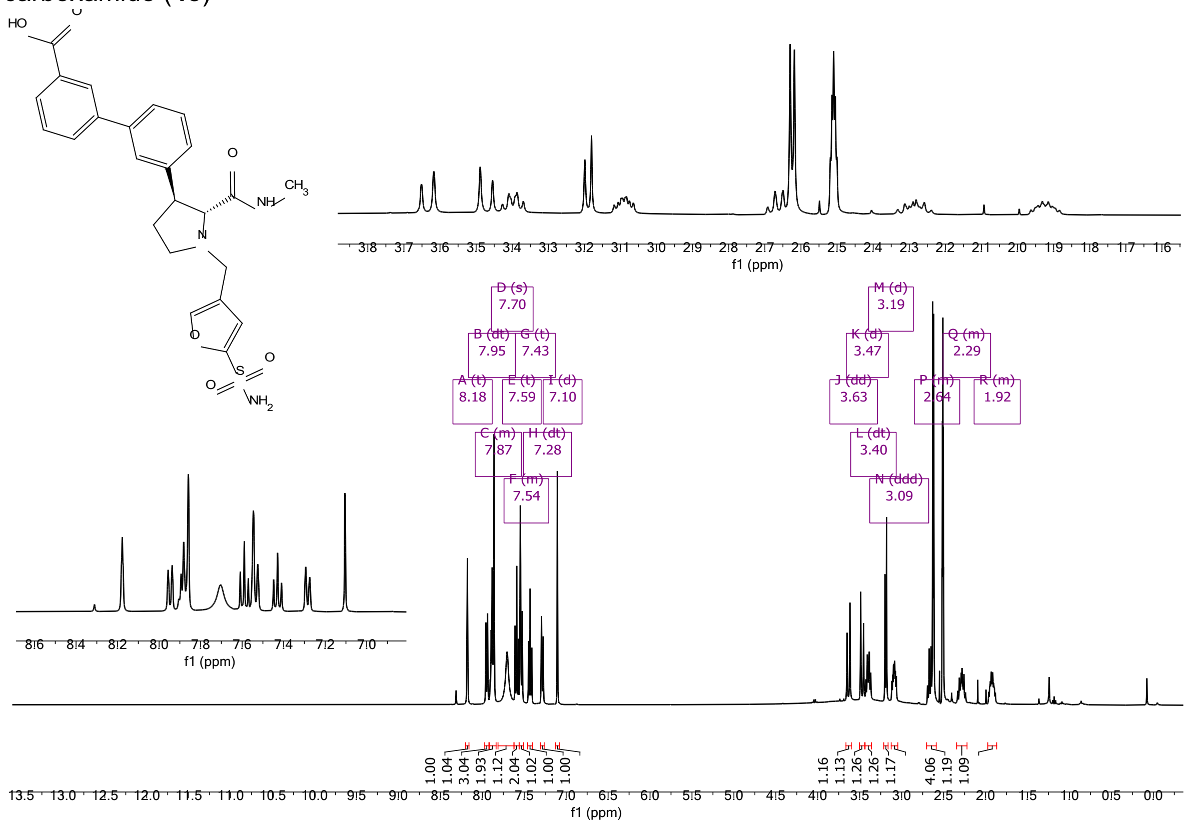

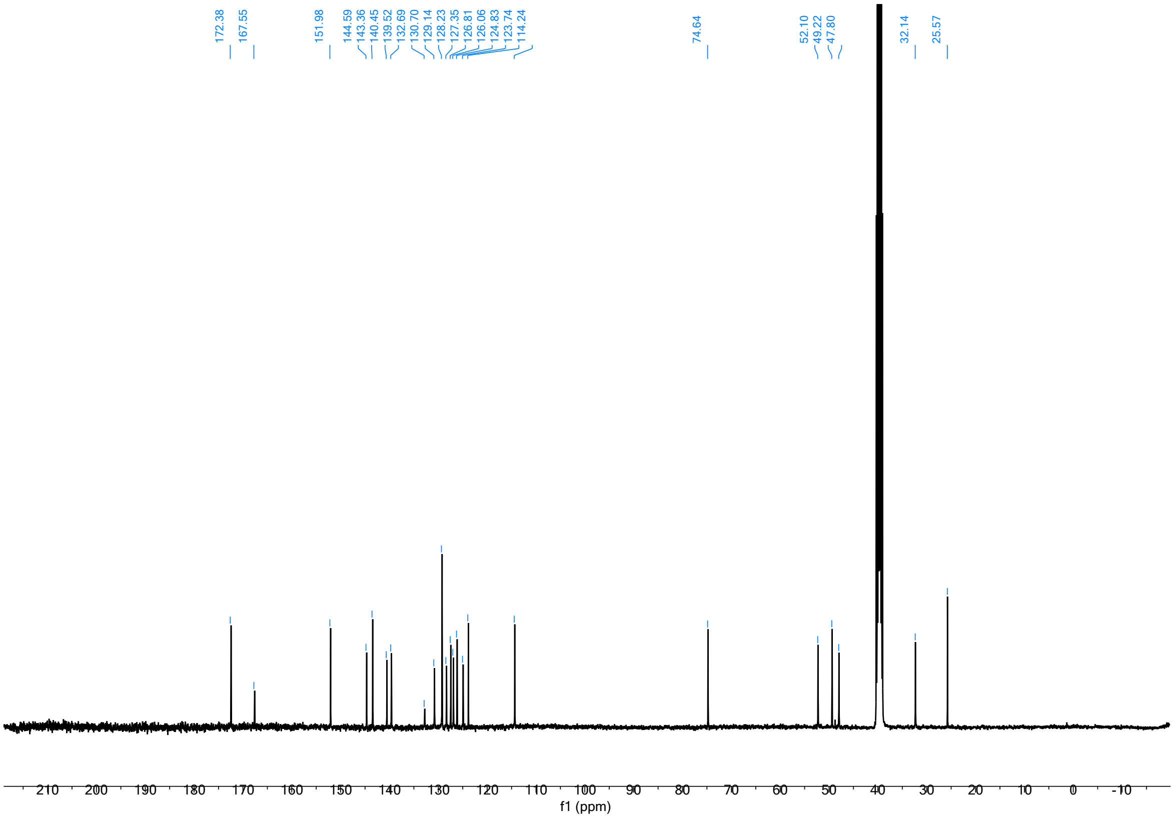

(2*R*,3*S*)-3-(3'-ethoxy-[1,1'-biphenyl]-4-yl)-*N*-methyl-1-((5-sulfamoylfuran-3-yl)methyl)pyrrolidine-2-carboxamide (**49**)

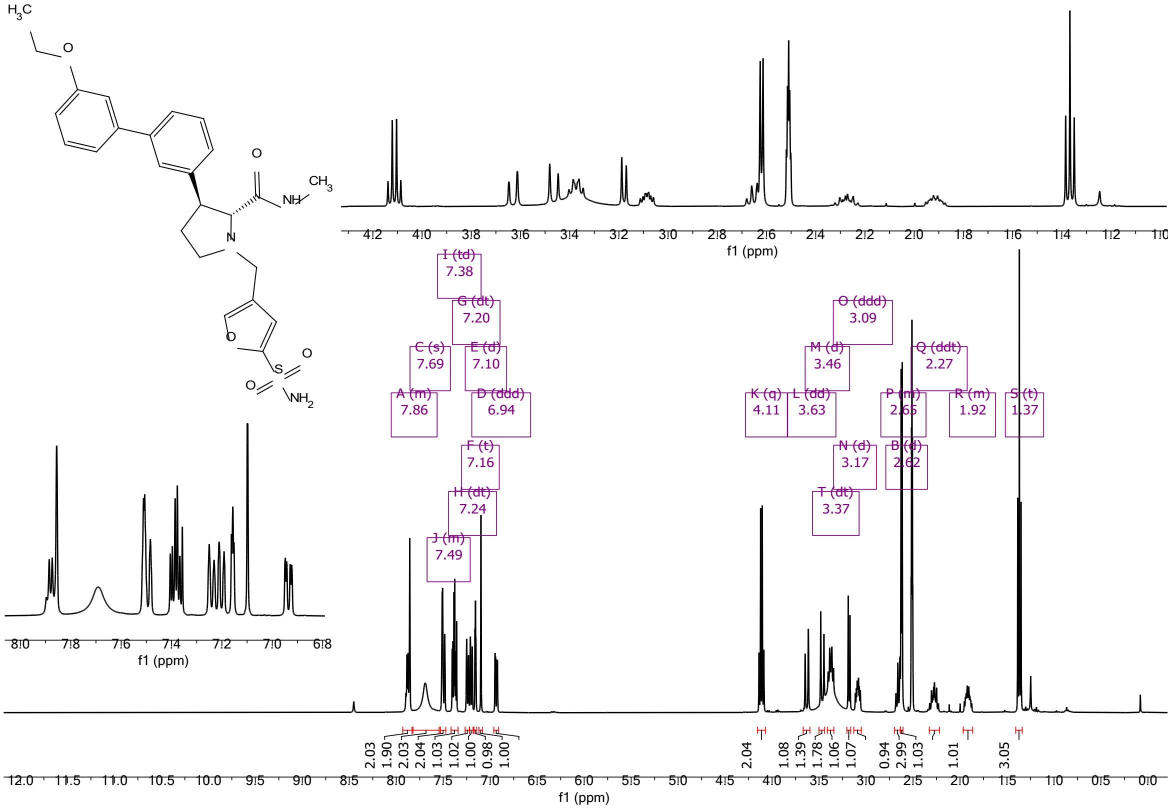

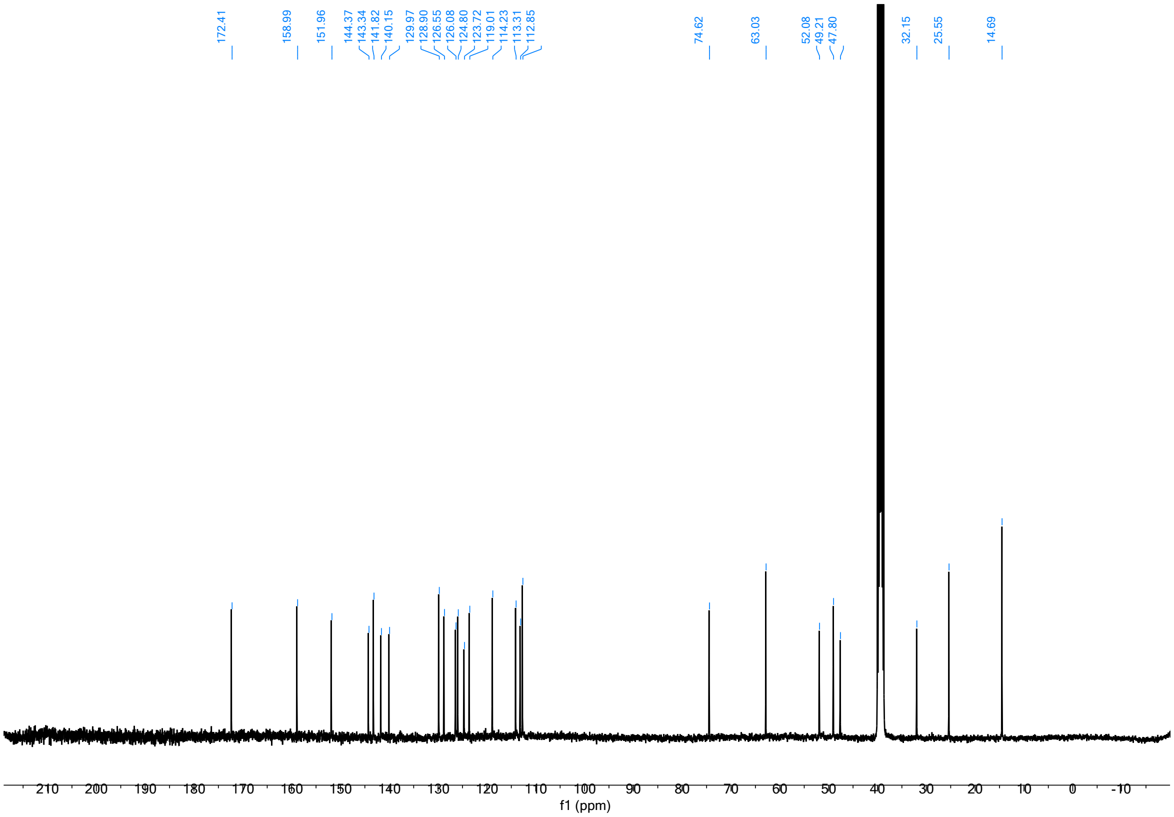

(2*S*,3*R*)-3-(3'-ethoxy-[1,1'-biphenyl]-4-yl)-*N*-methyl-1-((5-sulfamoylfuran-3-yl)methyl)azetidine-2-carboxamide (**50**)

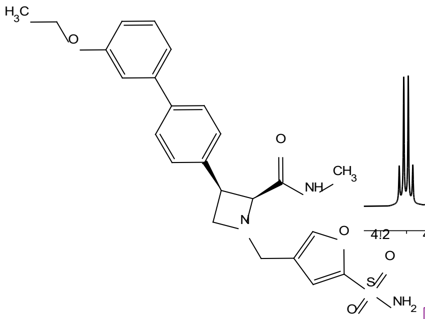

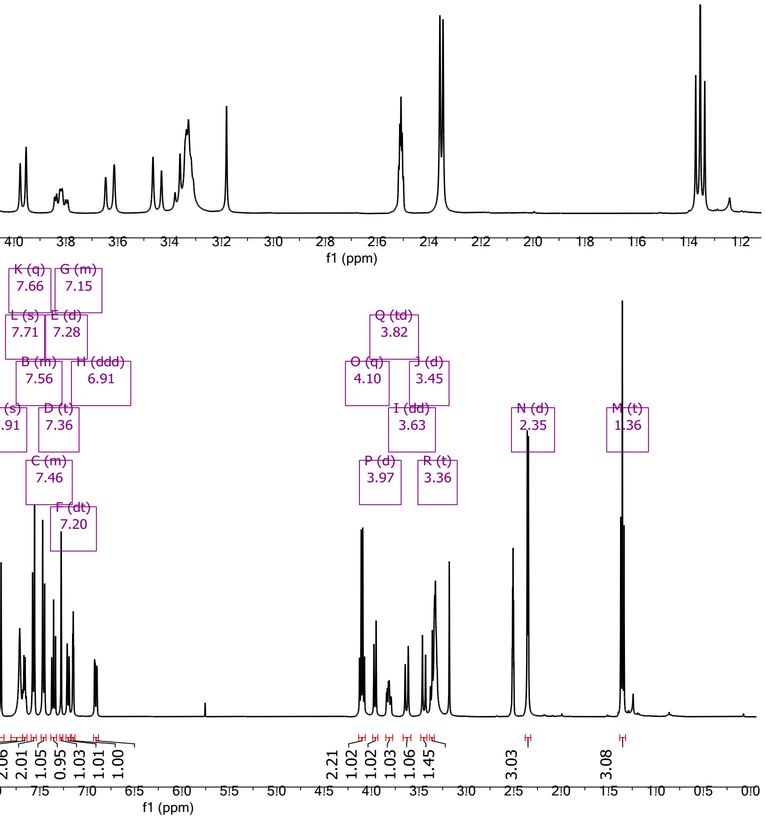

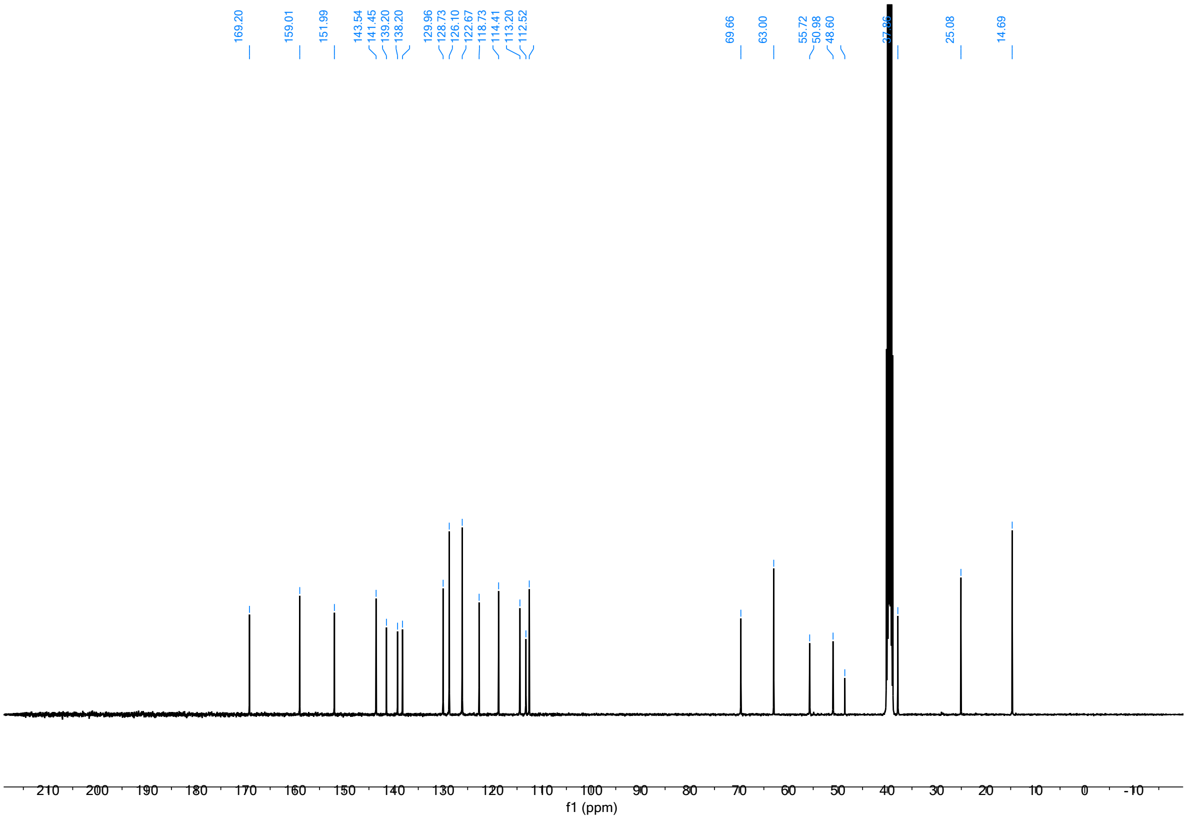

(2*S*,3*S*)-3-(5'-chloro-2'-methyl-[1,1'-biphenyl]-3-yl)-*N*-methyl-1-((5-sulfamoylfuran-3-yl)methyl)piperidine-2-carboxamide (**51**)

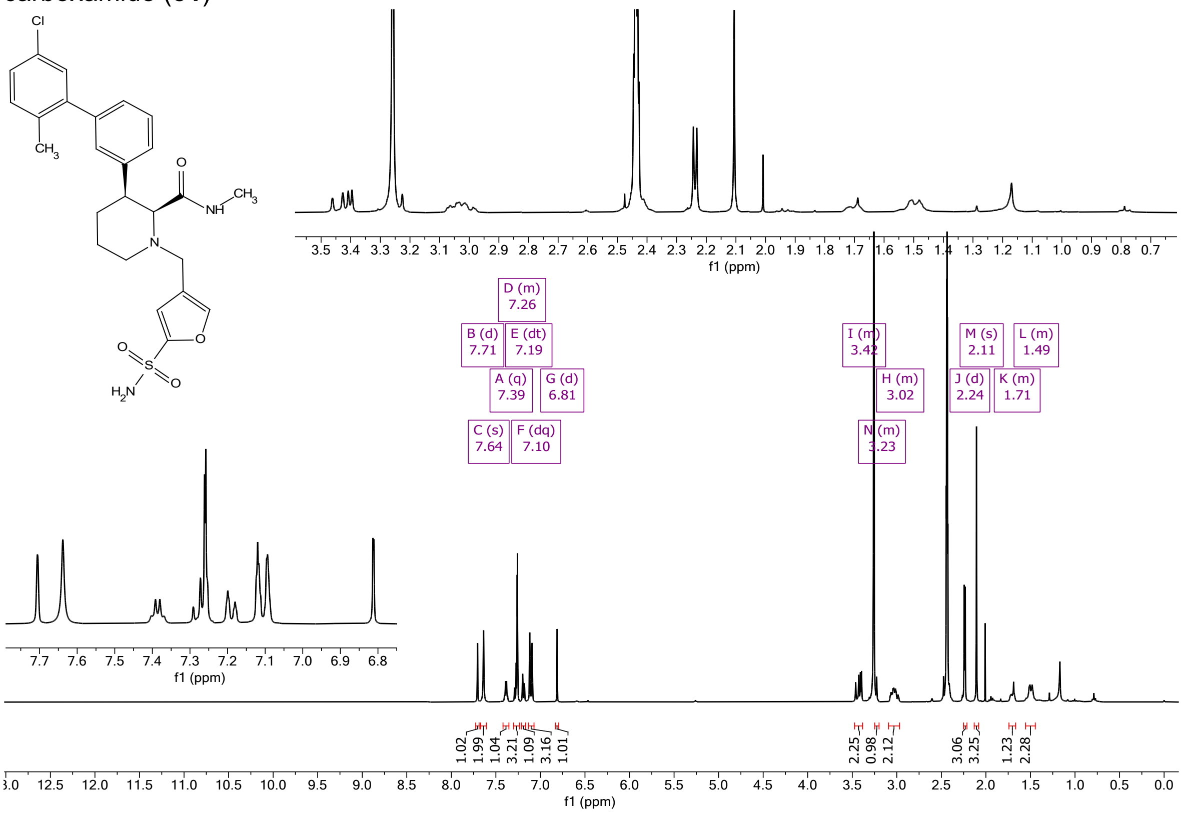

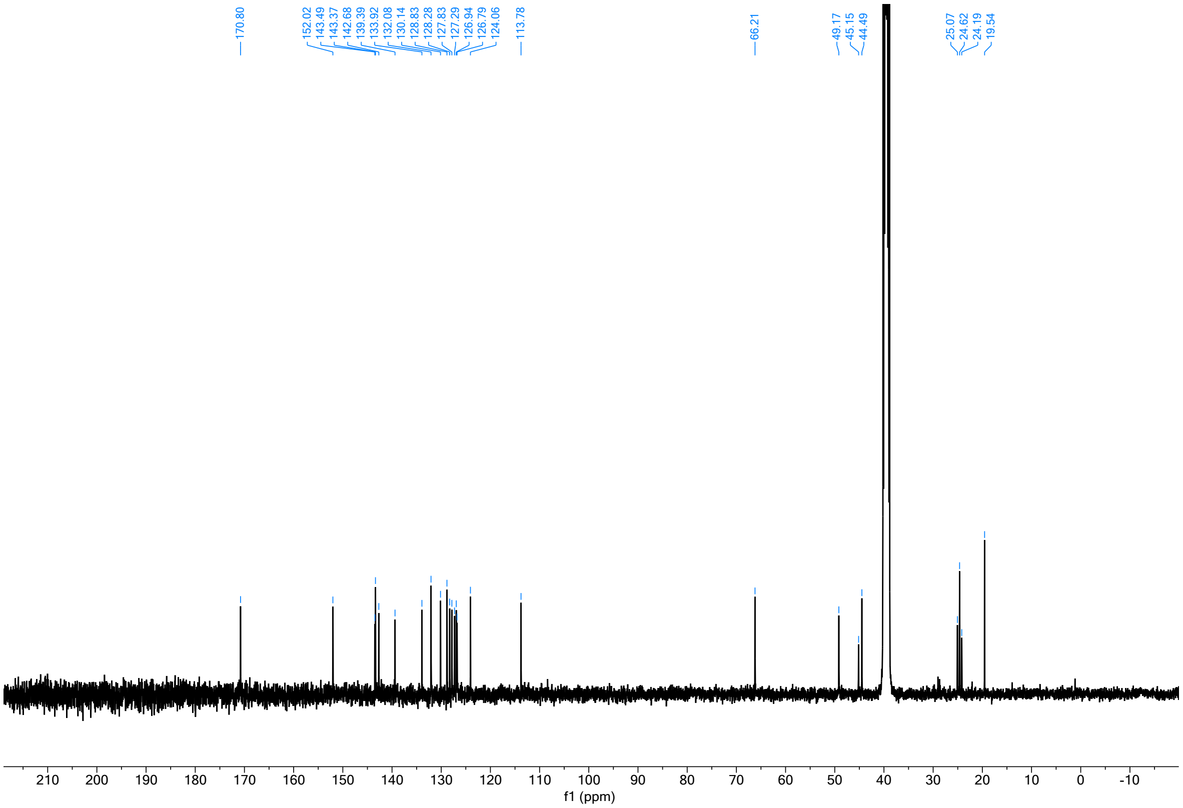

(2*R*,4*S*)-1-((6,7-dihydro-5H-pyrrolo[1,2-a]imidazol-2-yl)methyl)-*N*-methyl-4-((4'-sulfamoyl-[1,1'-biphenyl]-3-yl)oxy)pyrrolidine-2-carboxamide (**52**)

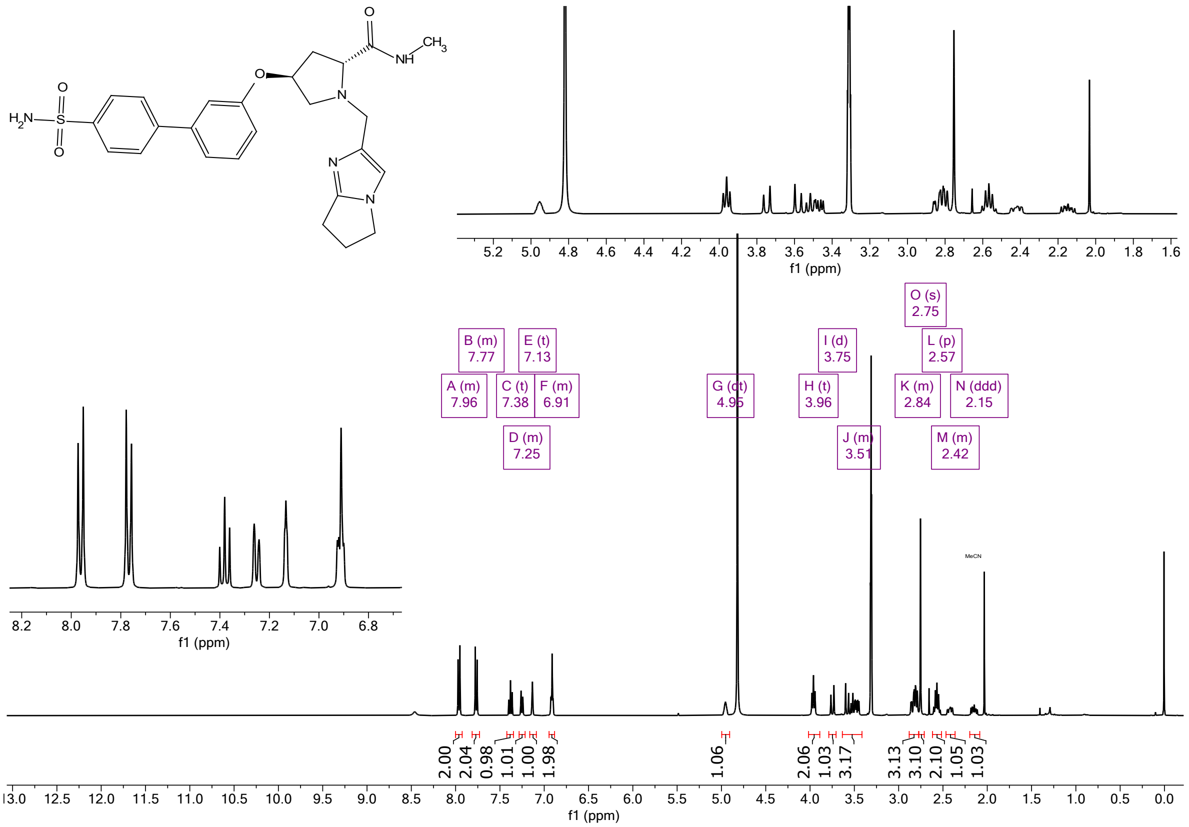

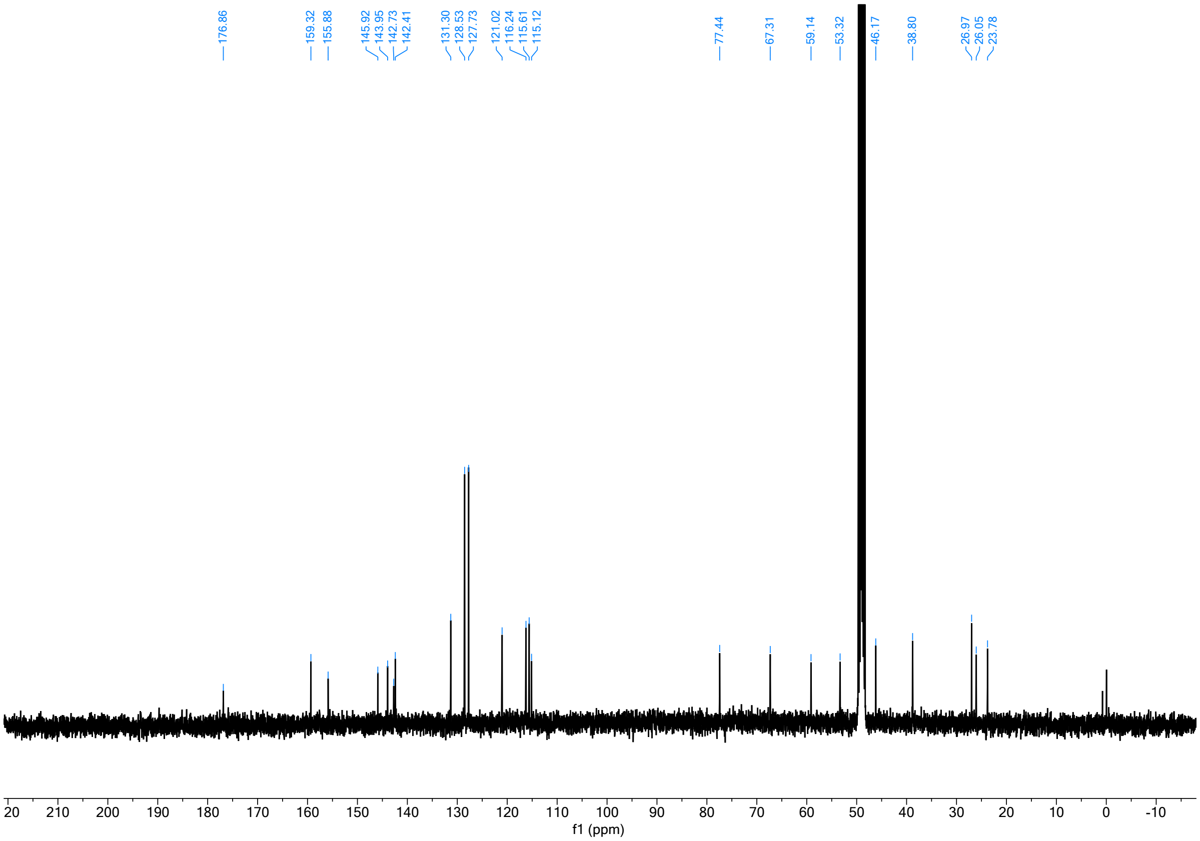

(*R*)-4'-(6-acetyl-2-(3-methyl-5,6,7,8-tetrahydroimidazo[1,5-*a*]pyridine-1-carbonyl)-2,6-diazaspiro[3.4]octan-8-yl)-[1,1'-biphenyl]-3-sulfonamide (**53**)

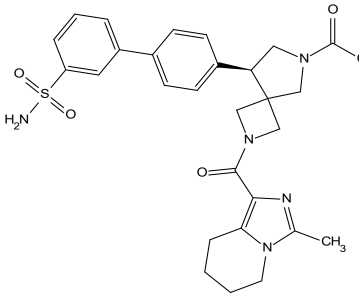

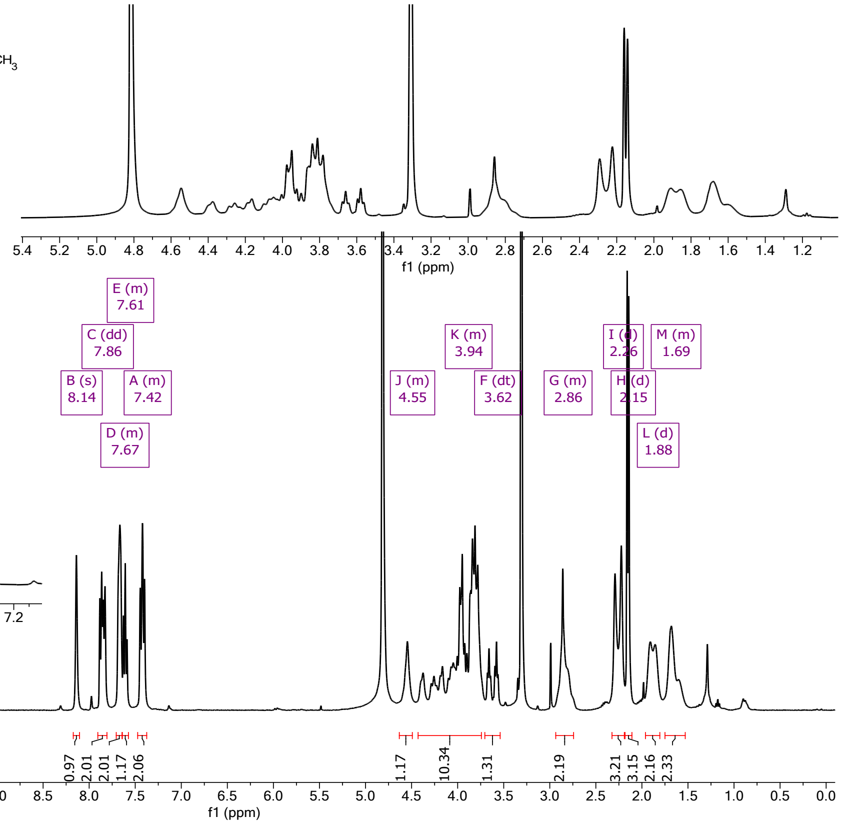

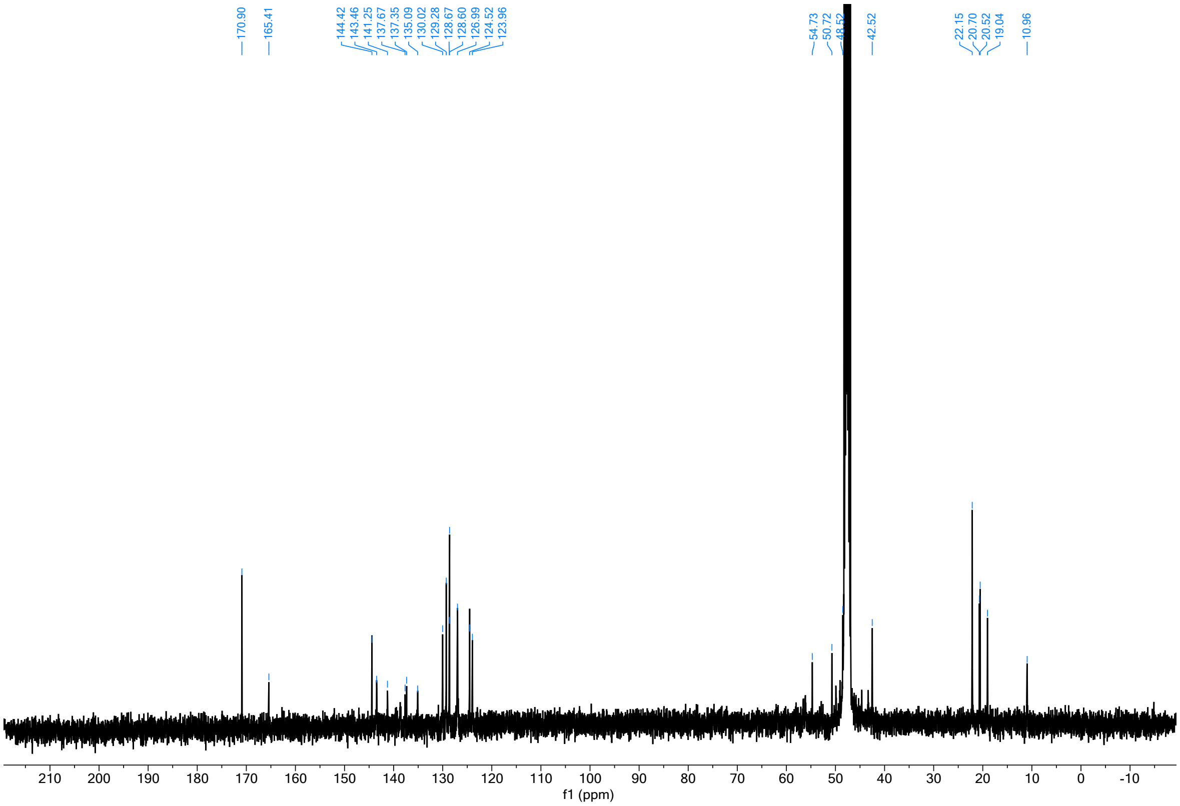

4'-((8*S*)-6-Acetyl-2-(2-(3-cyanophenoxy)propanoyl)-2,6-diazaspiro[3.4]octan-8-yl)-[1,1'-biphenyl]-4-sulfonamide (**54**)

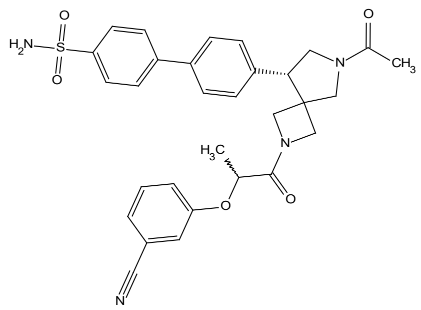

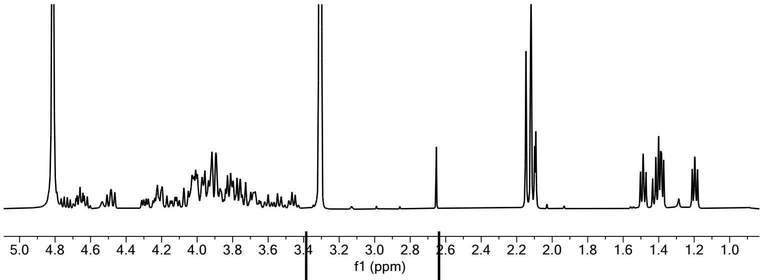

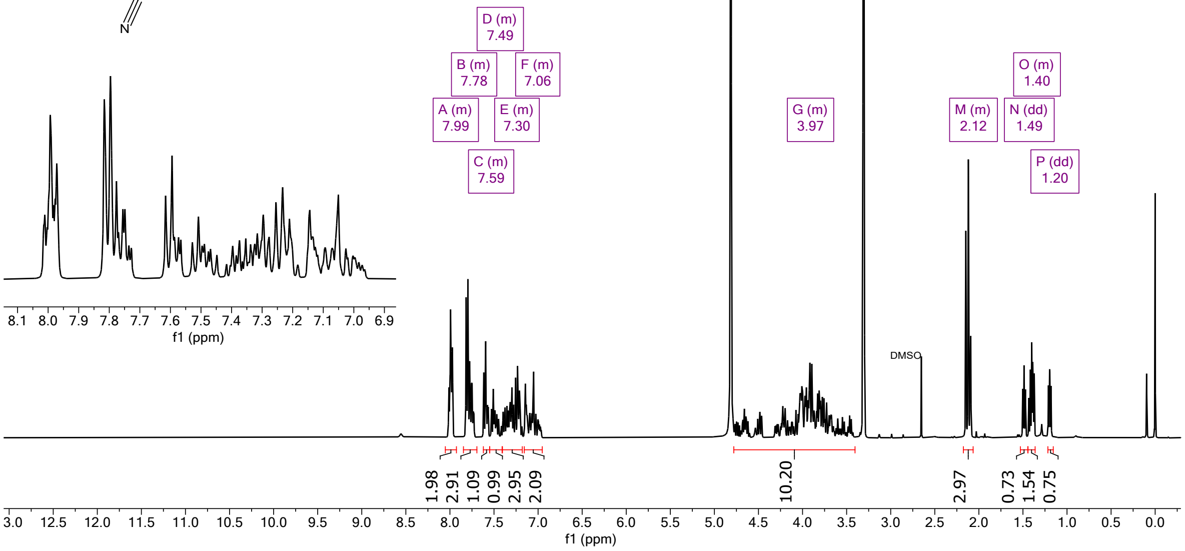

3-(3'-((2S,3R)-2-(Methylcarbamoyl)-1-(4-sulfamoylbenzoyl)pyrrolidin-3-yl)-[1,1'-biphenyl]-3-yl)propanoic acid (**55**)

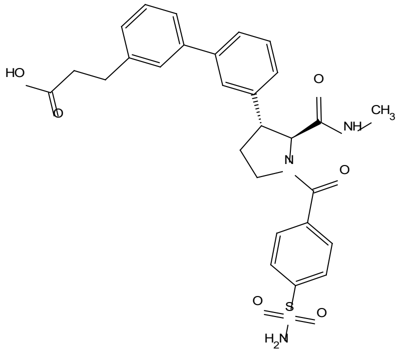

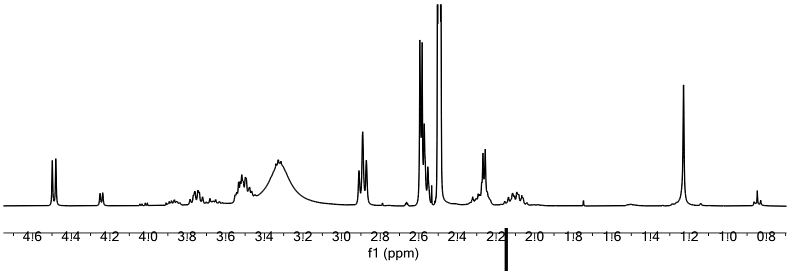

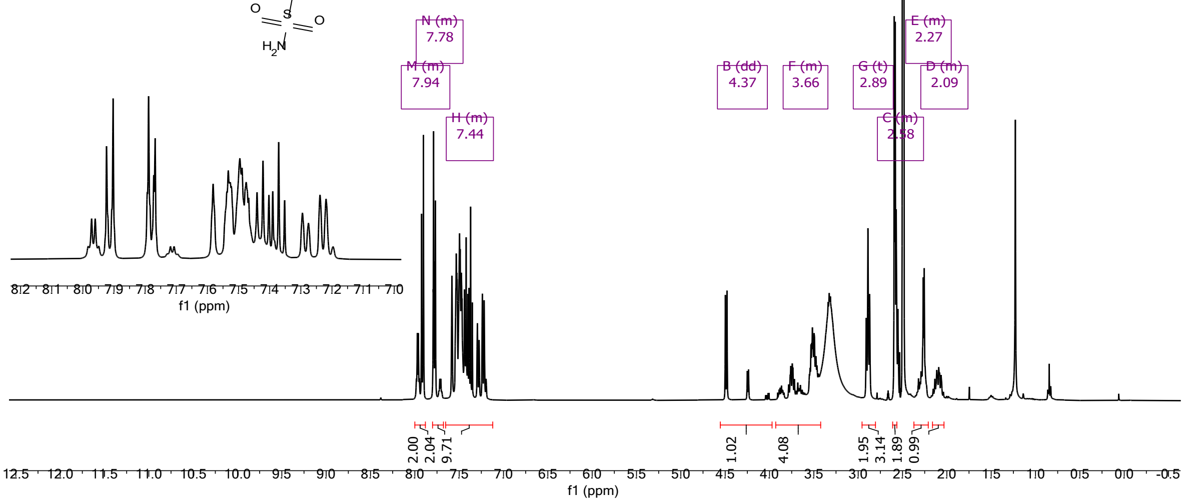

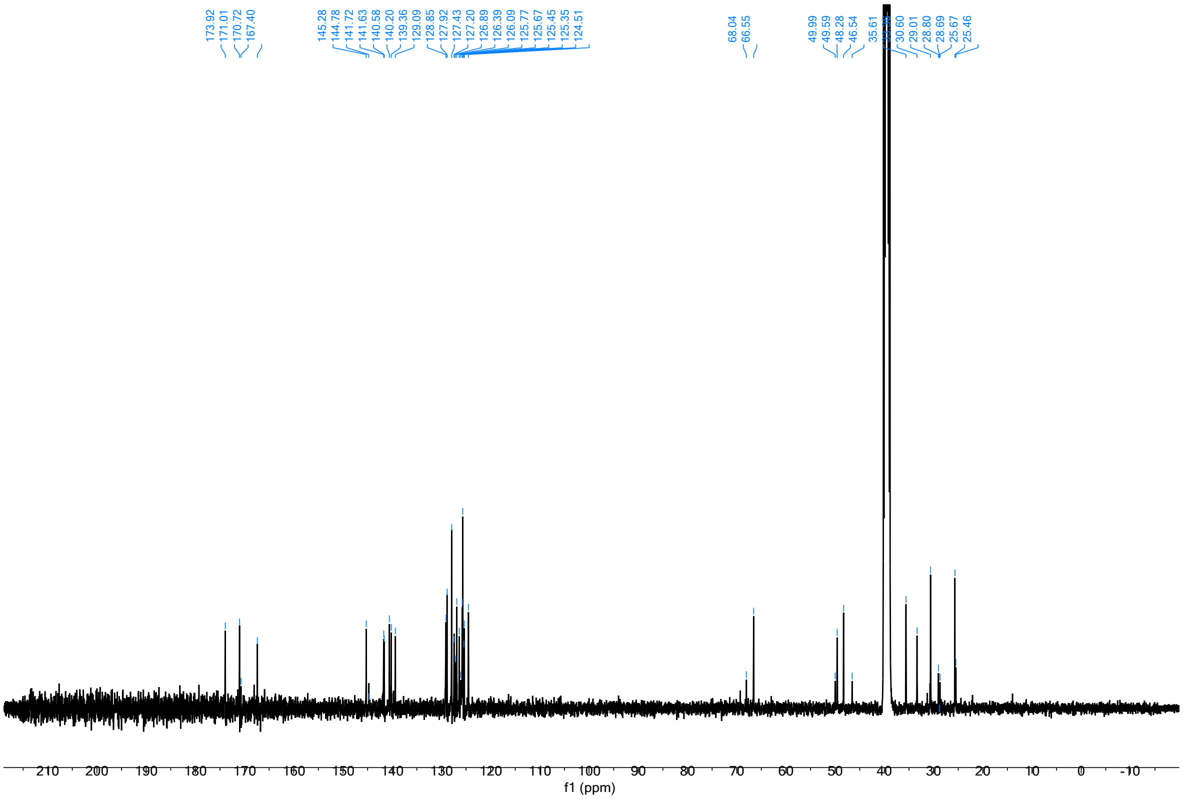

*N*-(((4*S*,5*R*)-2-((*R*)-1-hydroxypropan-2-yl)-4-methyl-1,1-dioxido-8-(3-sulfamoylphenyl)-2,3,4,5-tetrahydrobenzo[*b*][1,4,5]oxathiazocin-5-yl)methyl)-*N*,4-dimethylbenzenesulfonamide (**56**)

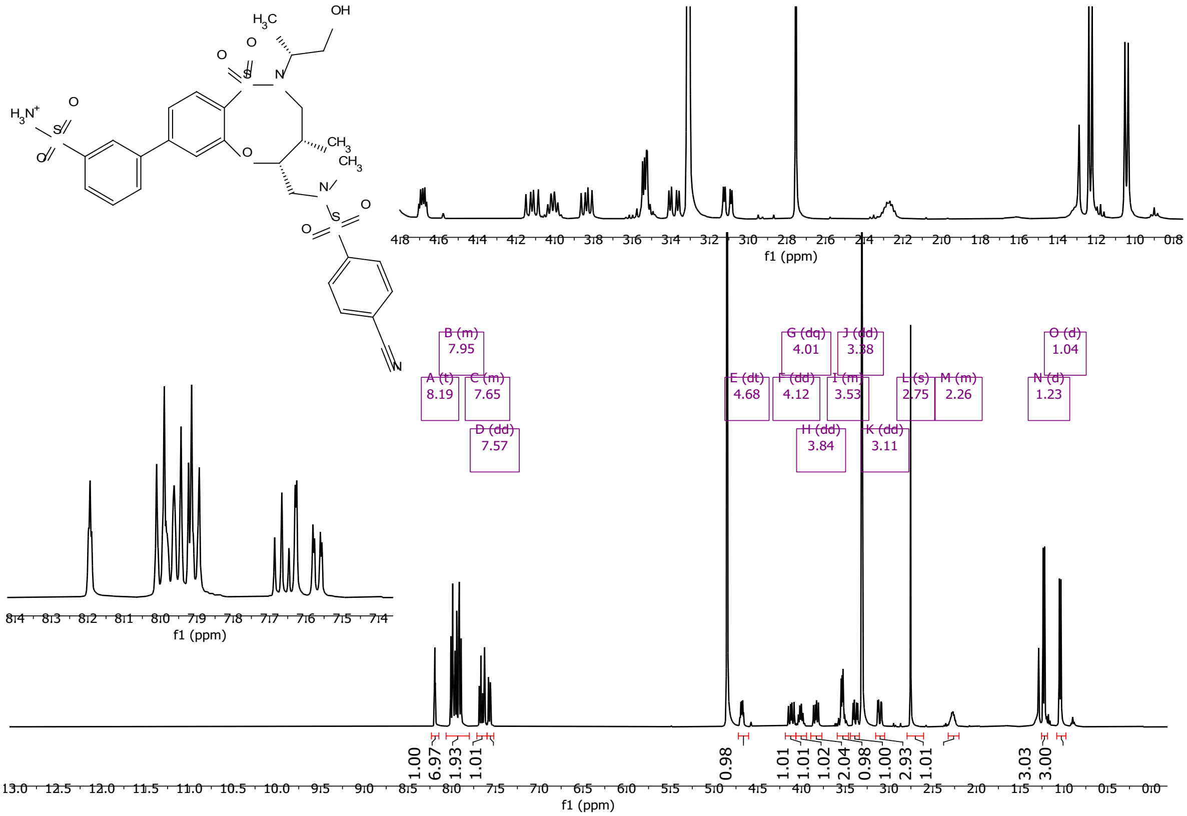

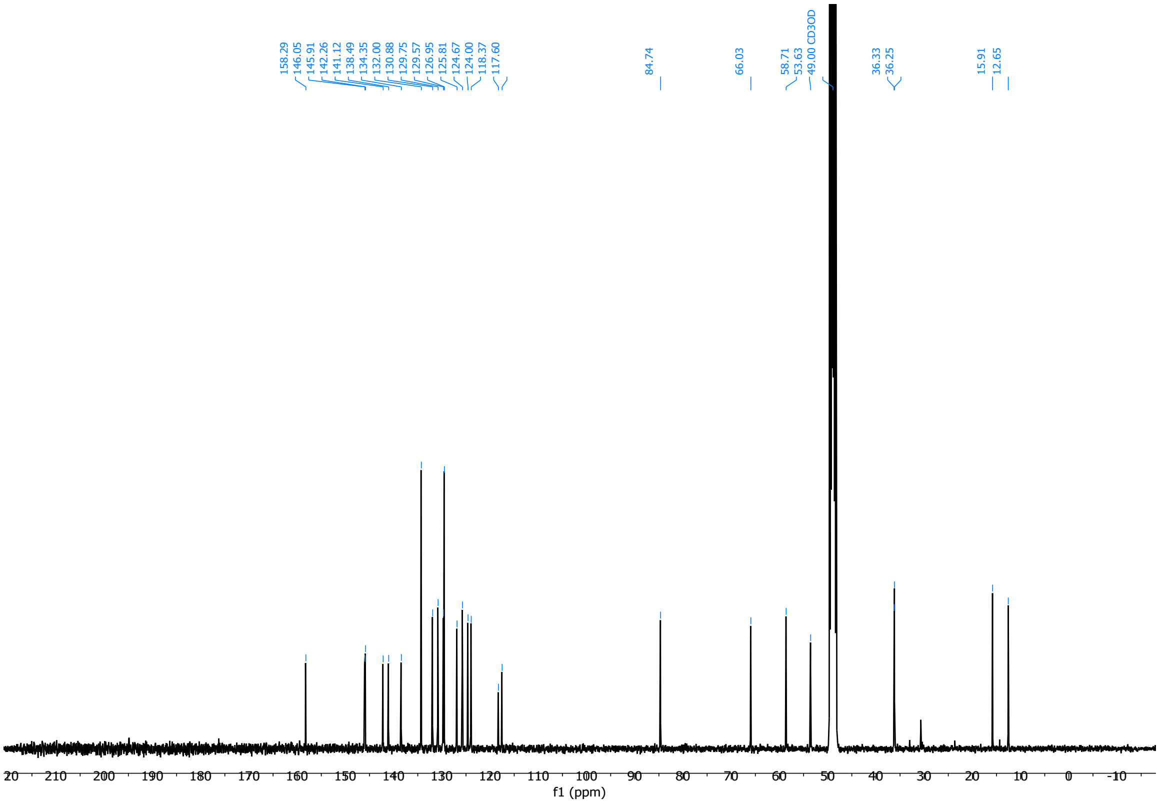

5-(Furan-2-yl)-*N*-(((4*S*,5*R*)-2-((*S*)-1-hydroxypropan-2-yl)-4-methyl-1,1-dioxido-8-(4-sulfamoylphenyl)-2,3,4,5-tetrahydrobenzo[*b*][1,4,5]oxathiazocin-5-yl)methyl)-*N*-methylisoxazole-4-carboxamide (**57**)

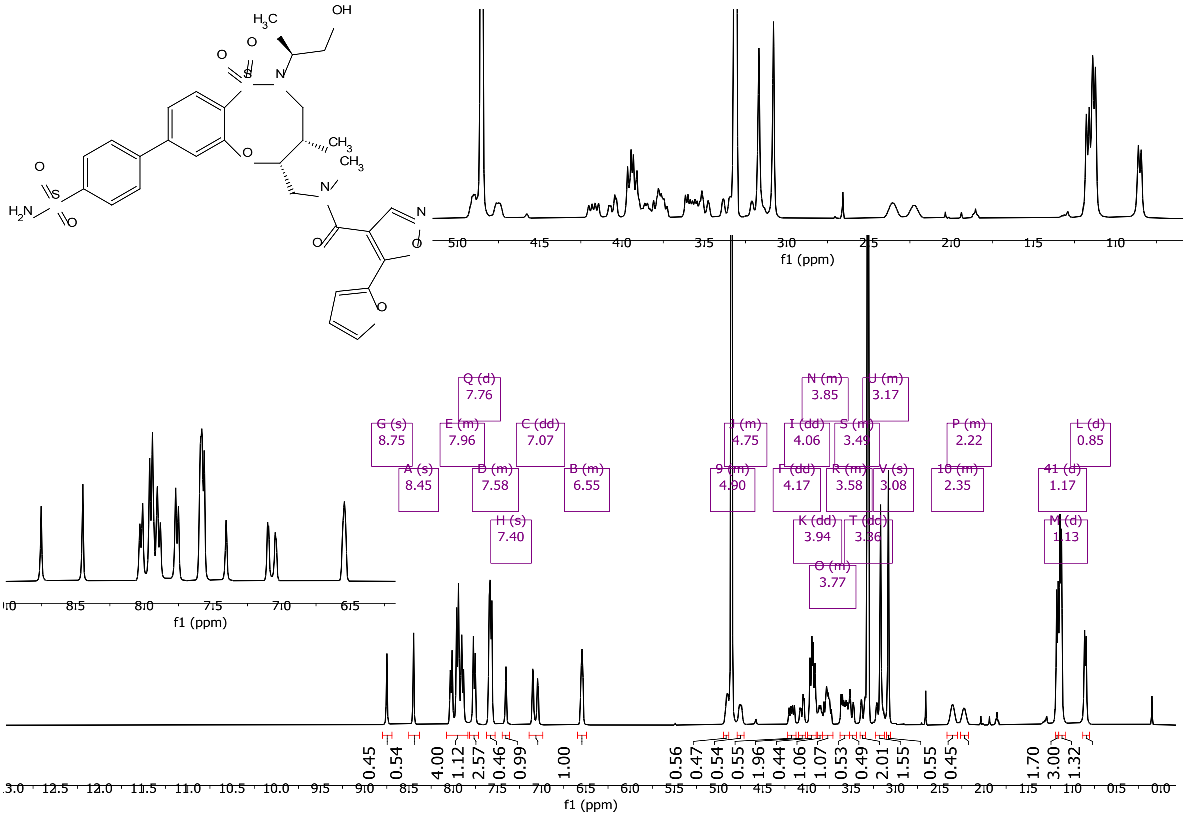

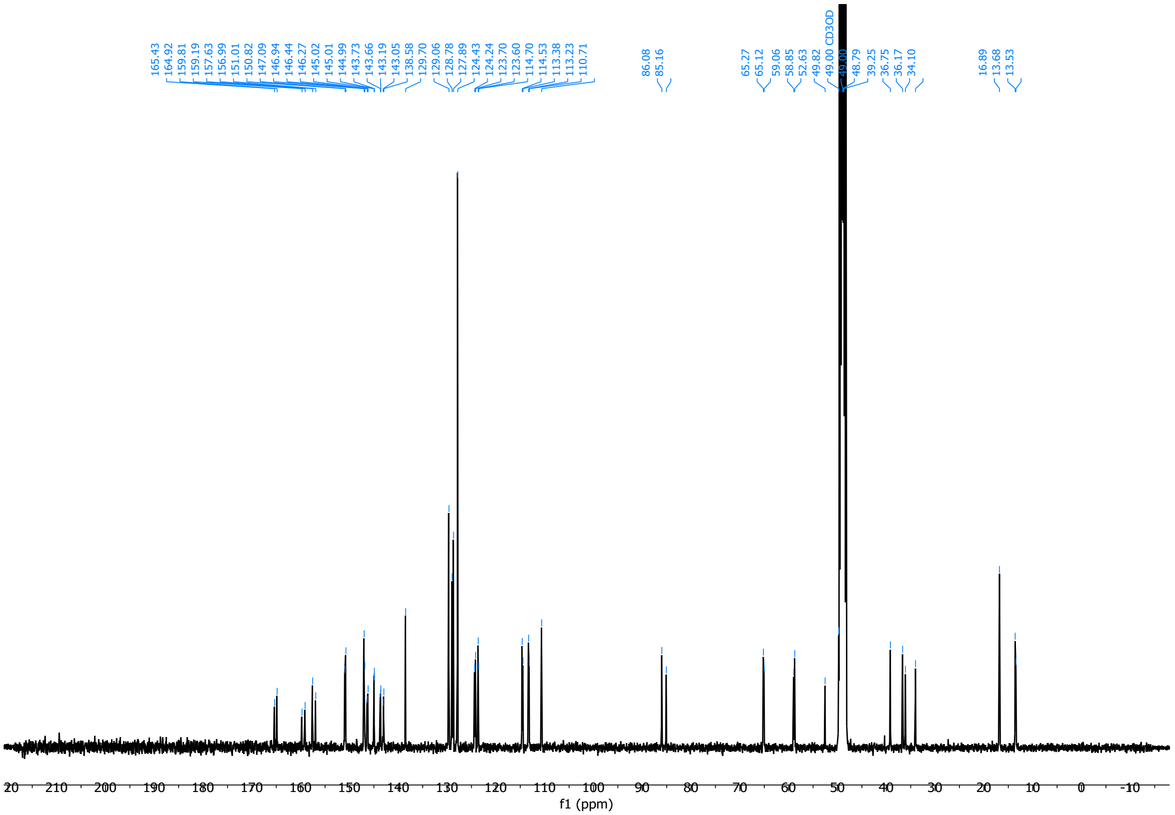

*N*-(((4*S*,5*S*)-2-((*R*)-1-hydroxypropan-2-yl)-4-methyl-1,1-dioxido-8-(4-sulfamoylphenyl)-2,3,4,5-tetrahydrobenzo[*b*][1,4,5]oxathiazocin-5-yl)methyl)-*N*-methyl-5-(methylsulfonyl)furan-3-carboxamide (**58**)

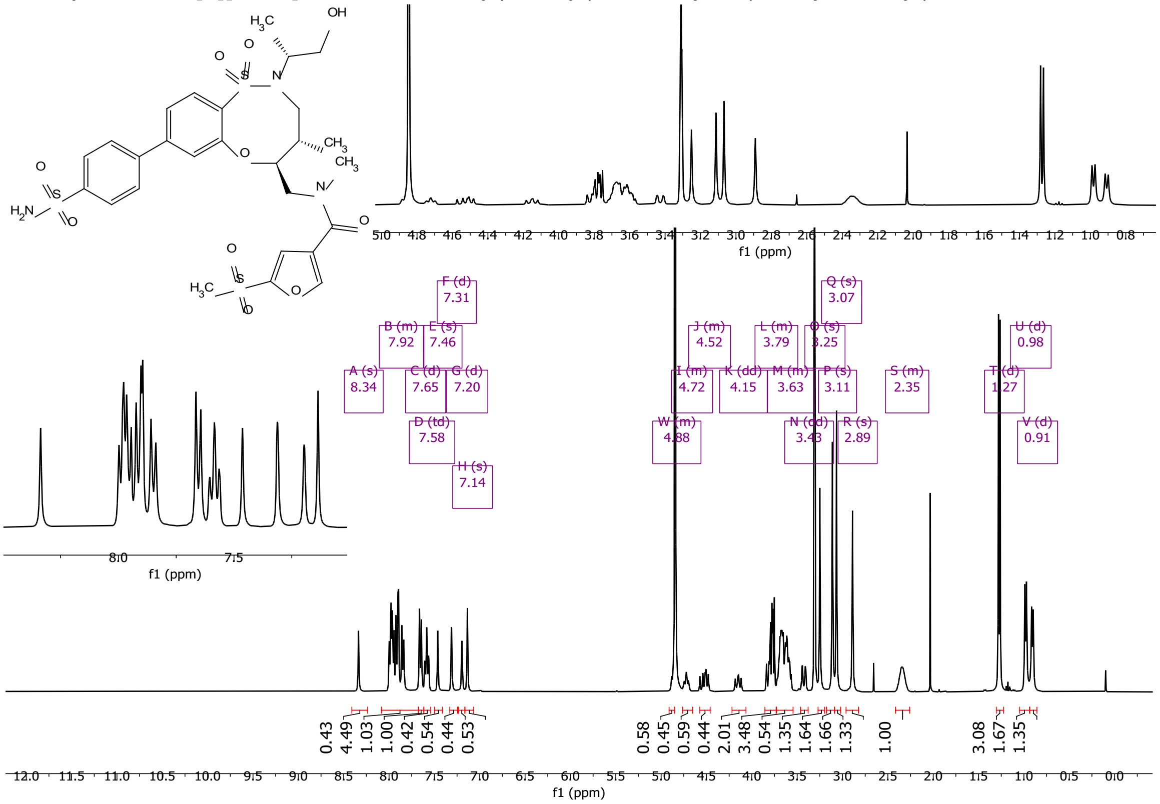

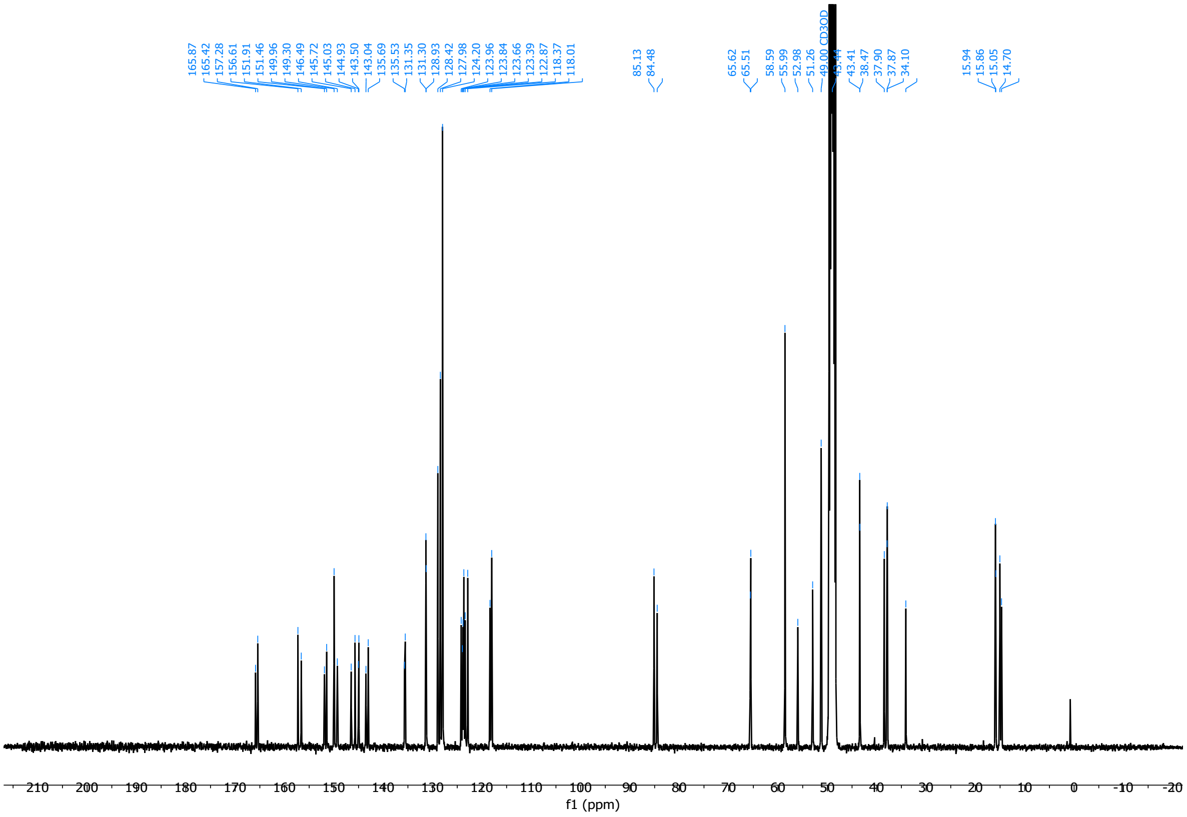

5-(Furan-2-yl)-*N*-(((4*R*,5*R*)-2-((*S*)-1-hydroxypropan-2-yl)-4-methyl-1,1-dioxido-8-(4-sulfamoylphenyl)-2,3,4,5-tetrahydrobenzo[*b*][1,4,5]oxathiazocin-5-yl)methyl)-*N*-methylisoxazole-4-carboxamide (**59**)

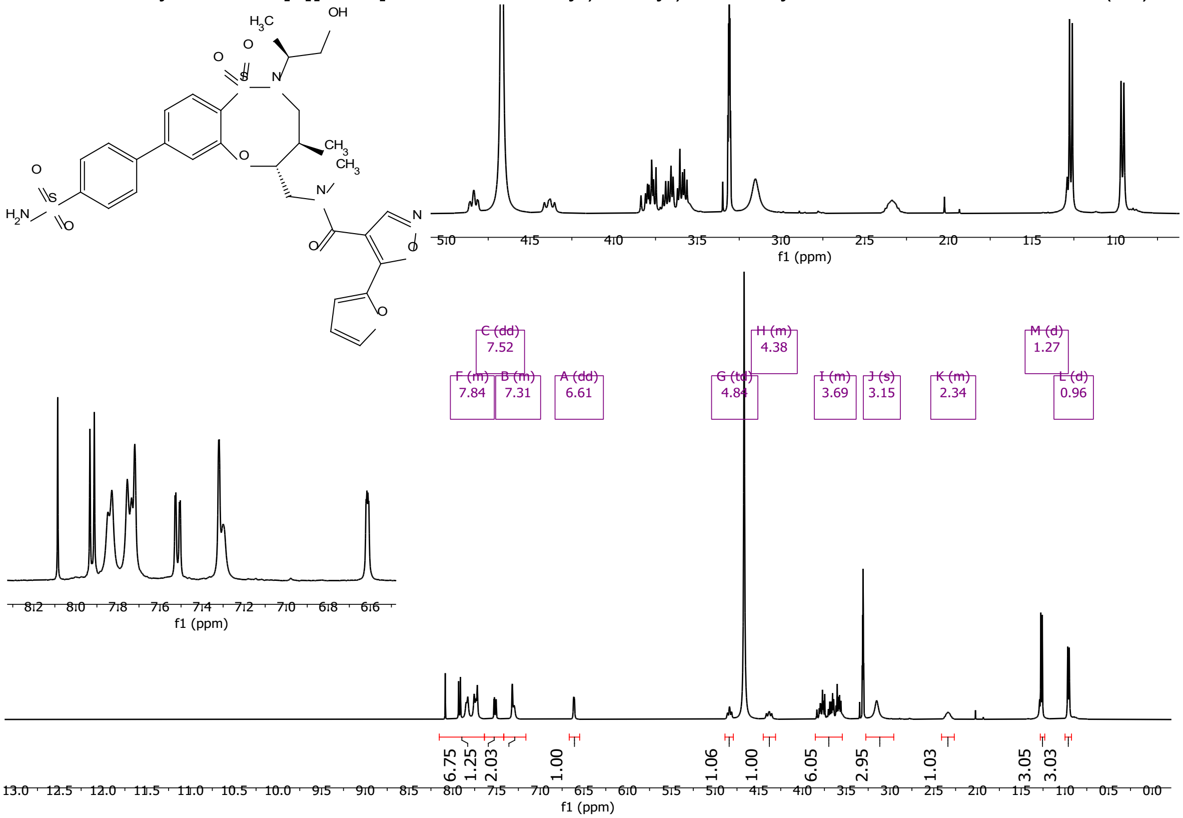

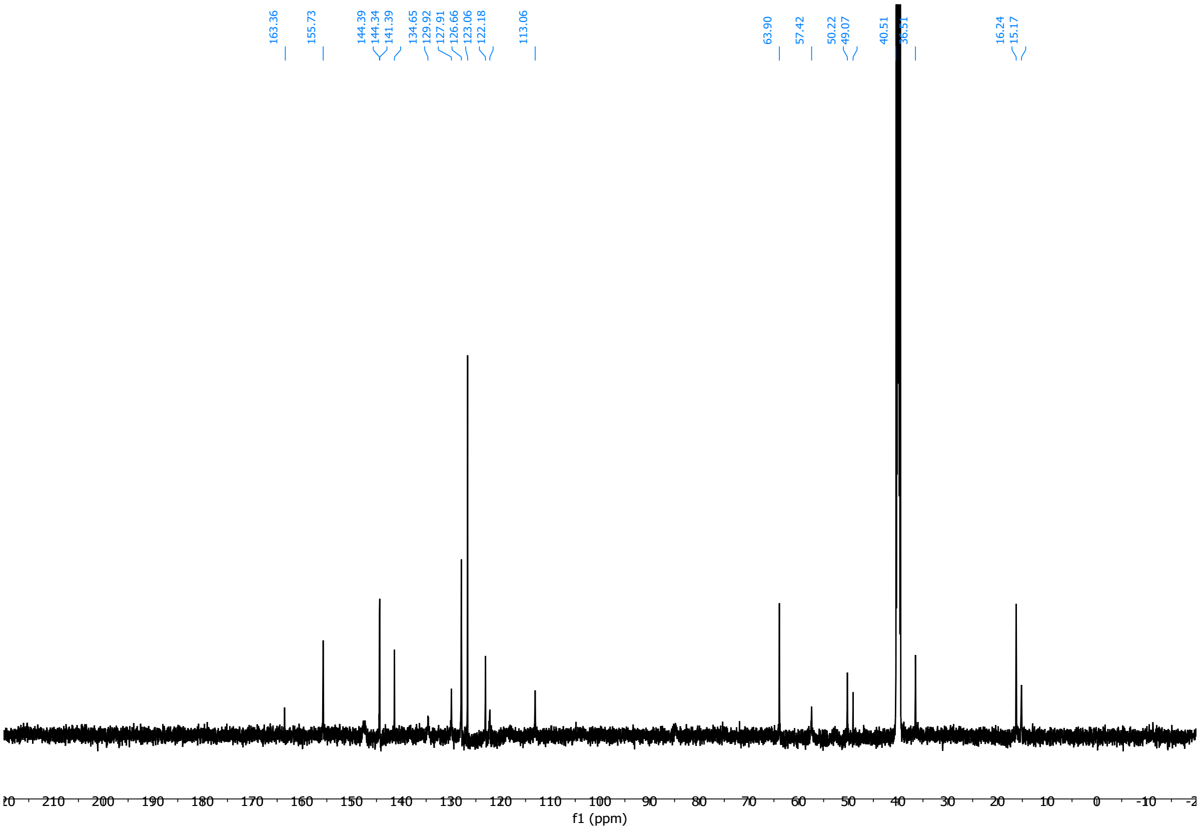

(*R*,*E*)-4-(4-(2-cyclohexylvinyl)phenyl)-2-((3,5-dichlorophenyl)sulfonyl)-3-(2-hydroxyethyl)-*N*-methyl-2,3-dihydro-1H-pyrrolo[3,4-*c*]pyridine-6-carboxamide (**60**)

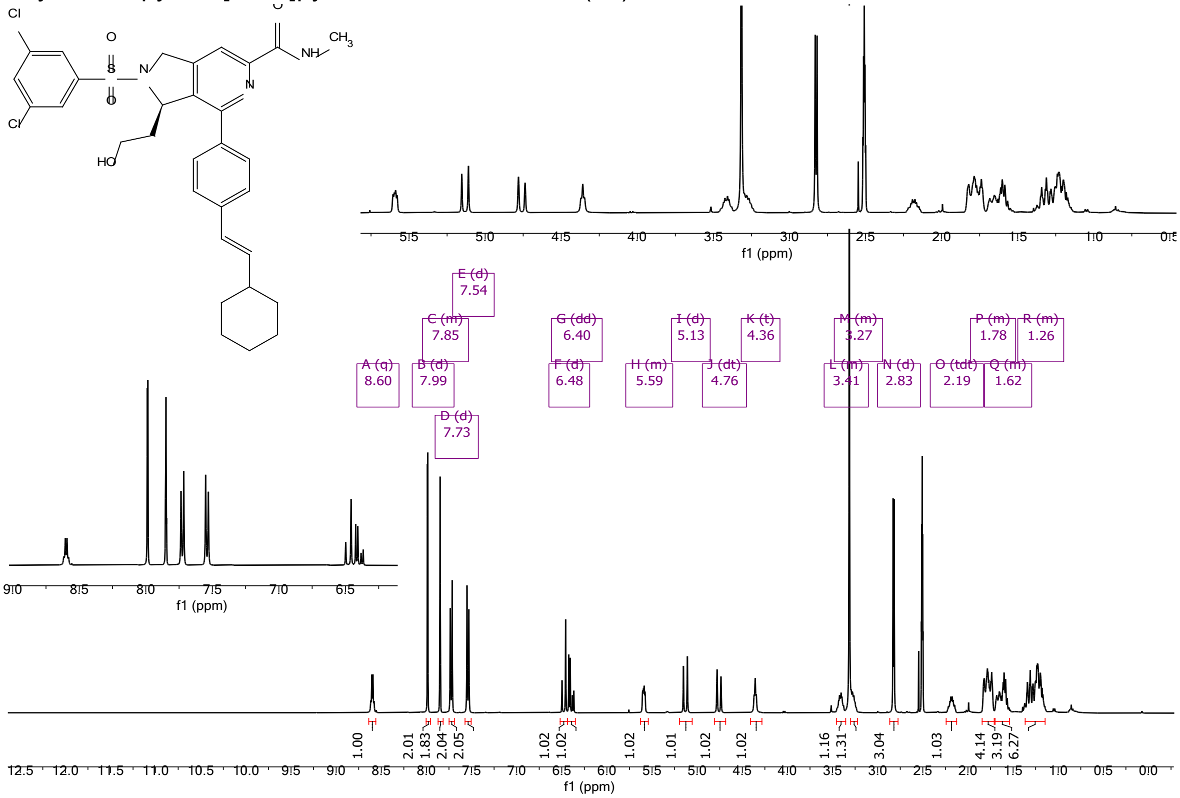

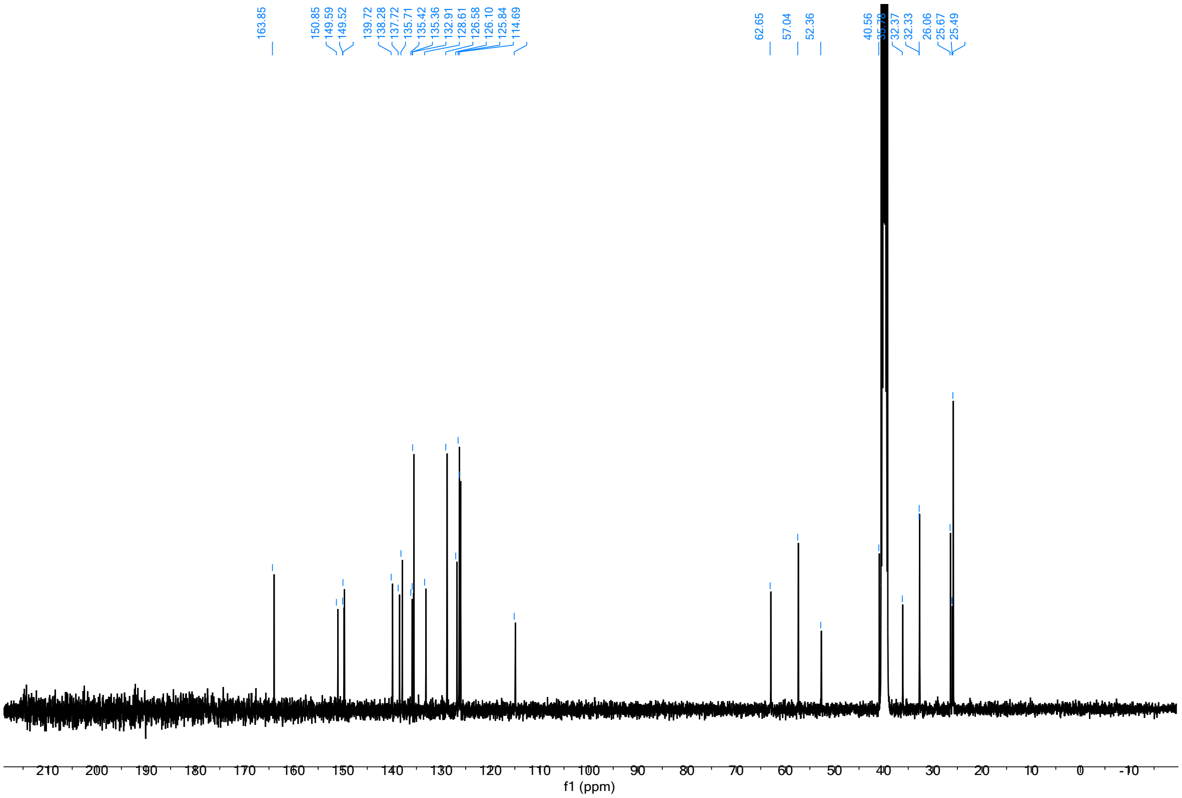

((8*R*,9*R*,10*S*)-9-(4-((*E*)-2-cyclohexylvinyl)phenyl)-6-((2-methoxyphenyl)sulfonyl)-1,6-diazabicyclo[6.2.0]decan-10-yl)methanol (**61**)

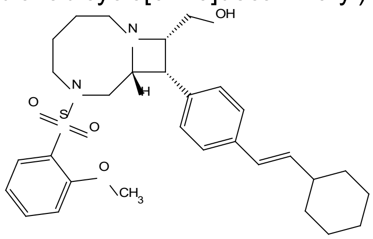

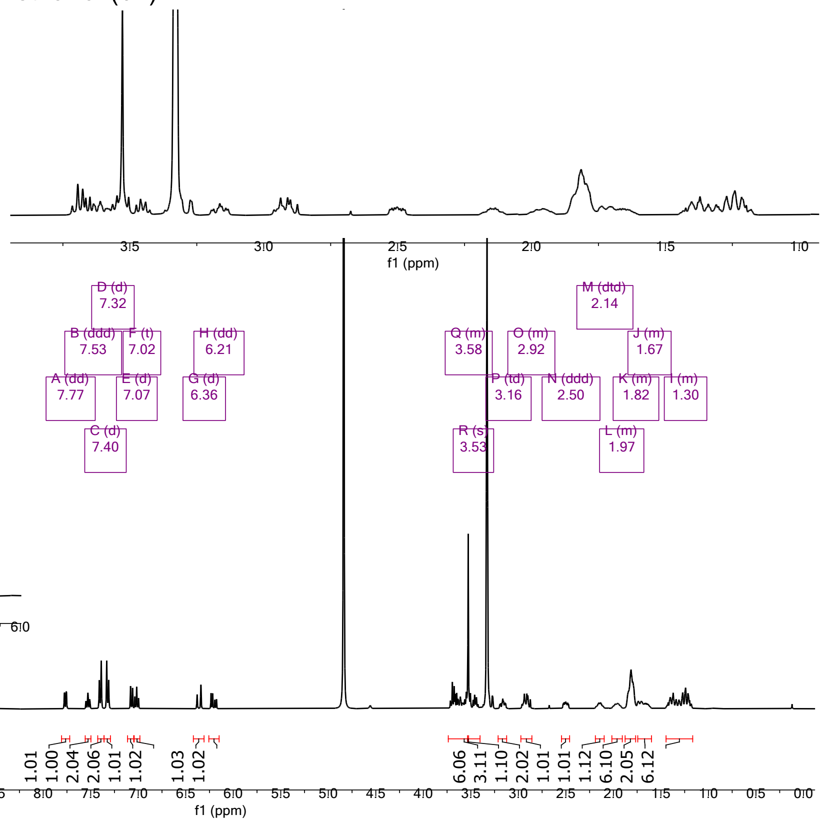

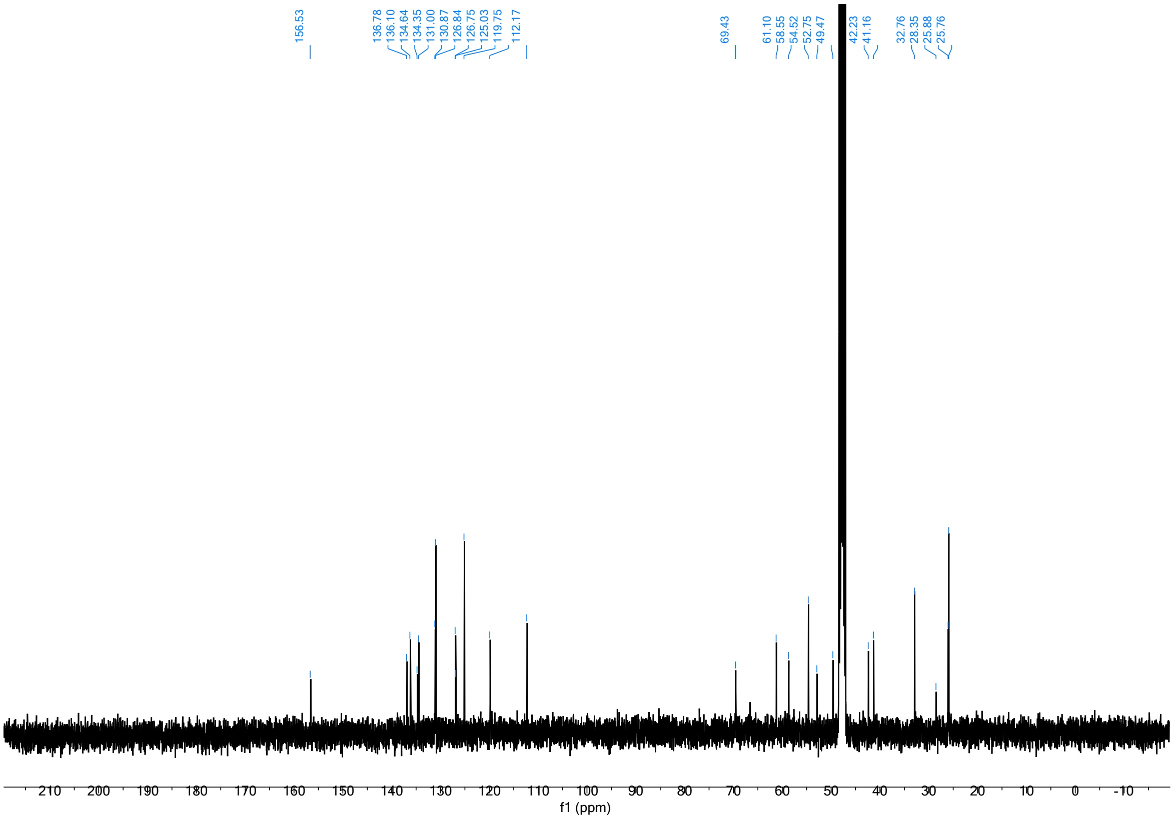

(2*R*,4*S*)-4-(3-((*E*)-2-cyclohexylvinyl)phenoxy)-1-((3,5-dichlorophenyl)sulfonyl)-*N*-methylpyrrolidine-2-carboxamide (**62**)

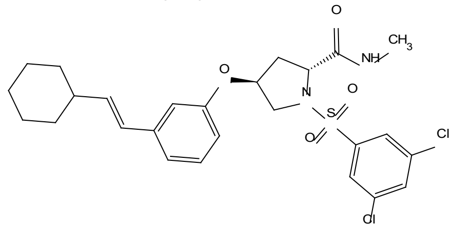

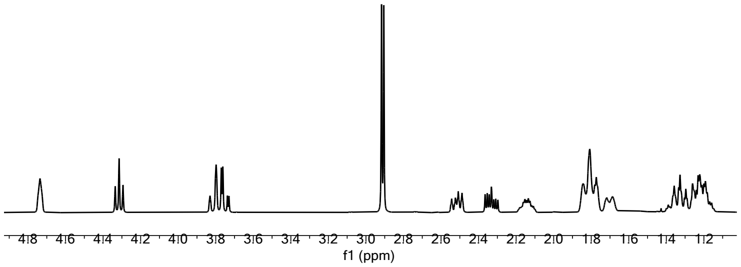

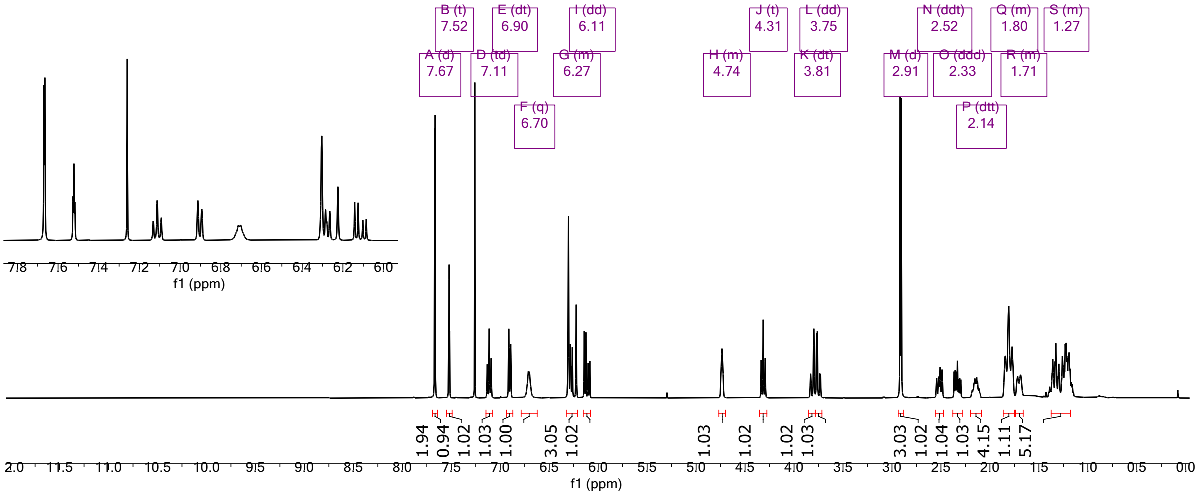

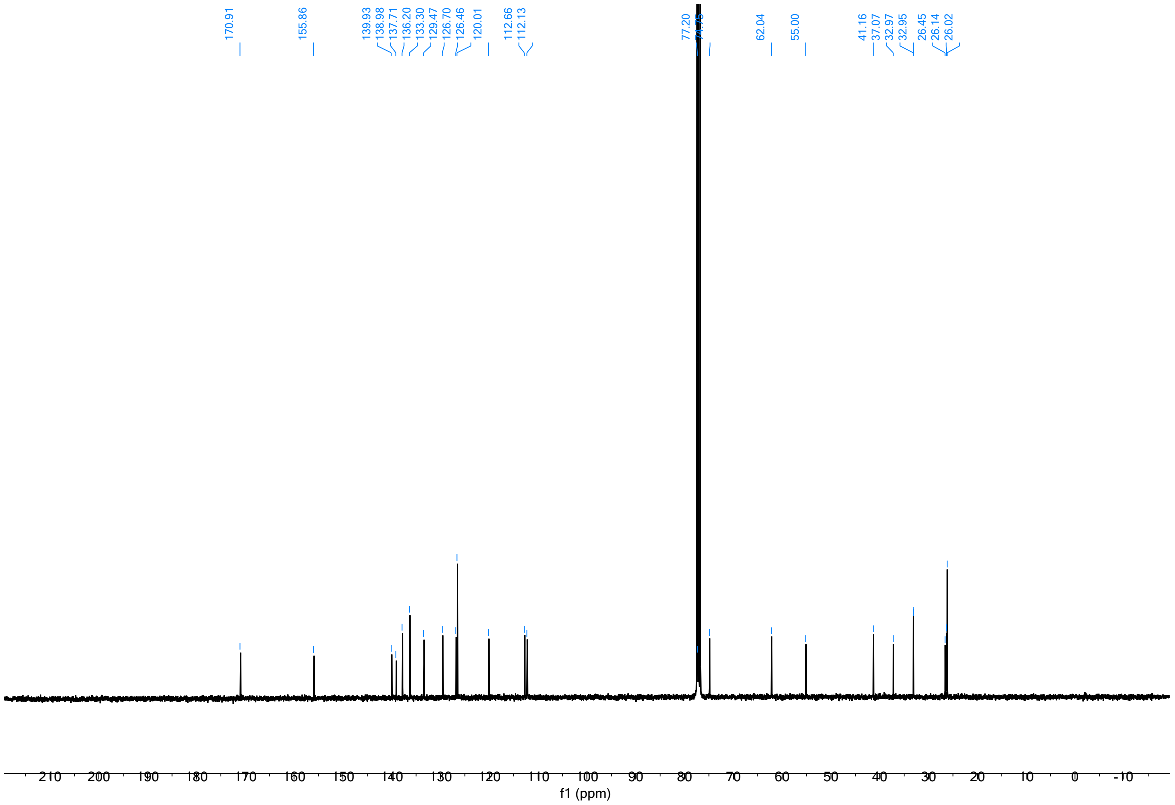

(2*S*,4*S*)-4-(4-((*E*)-2-cyclohexylvinyl)phenoxy)-1-(3-ethynylbenzyl)-*N*-methylpyrrolidine-2-carboxamide (**63**)

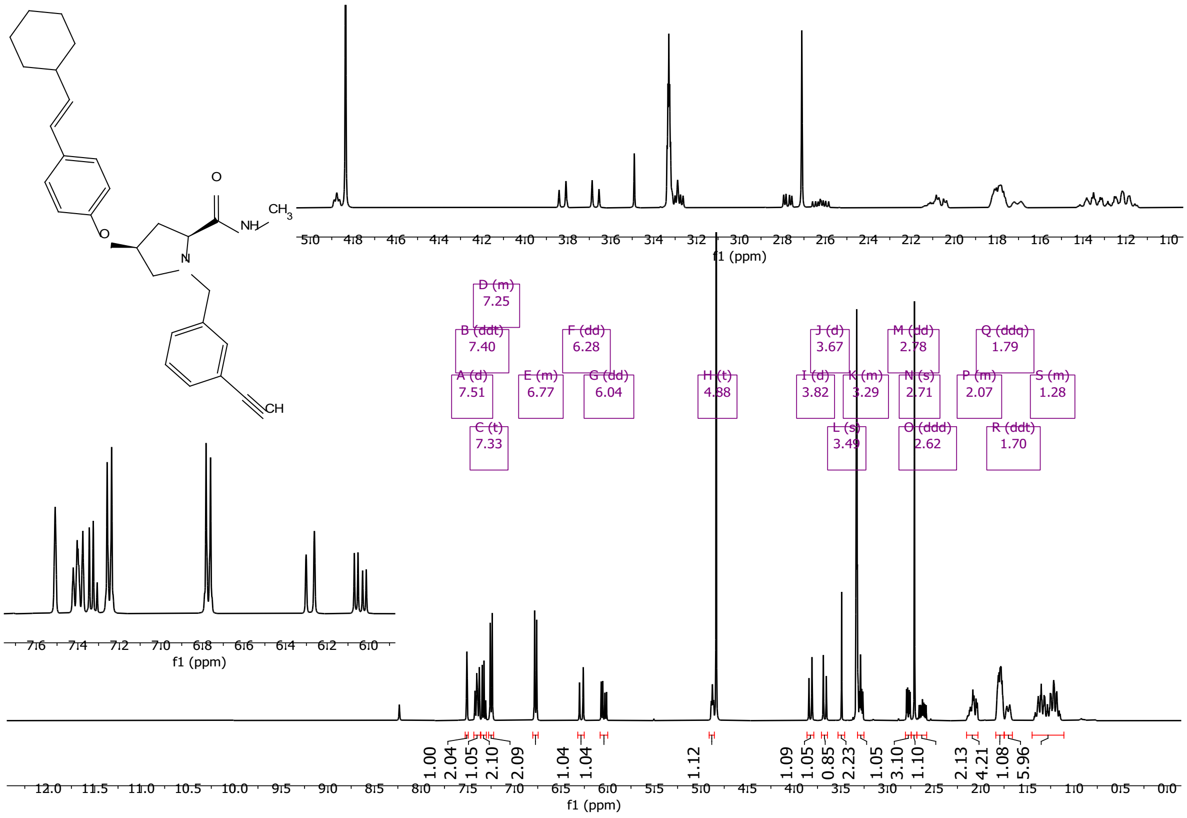

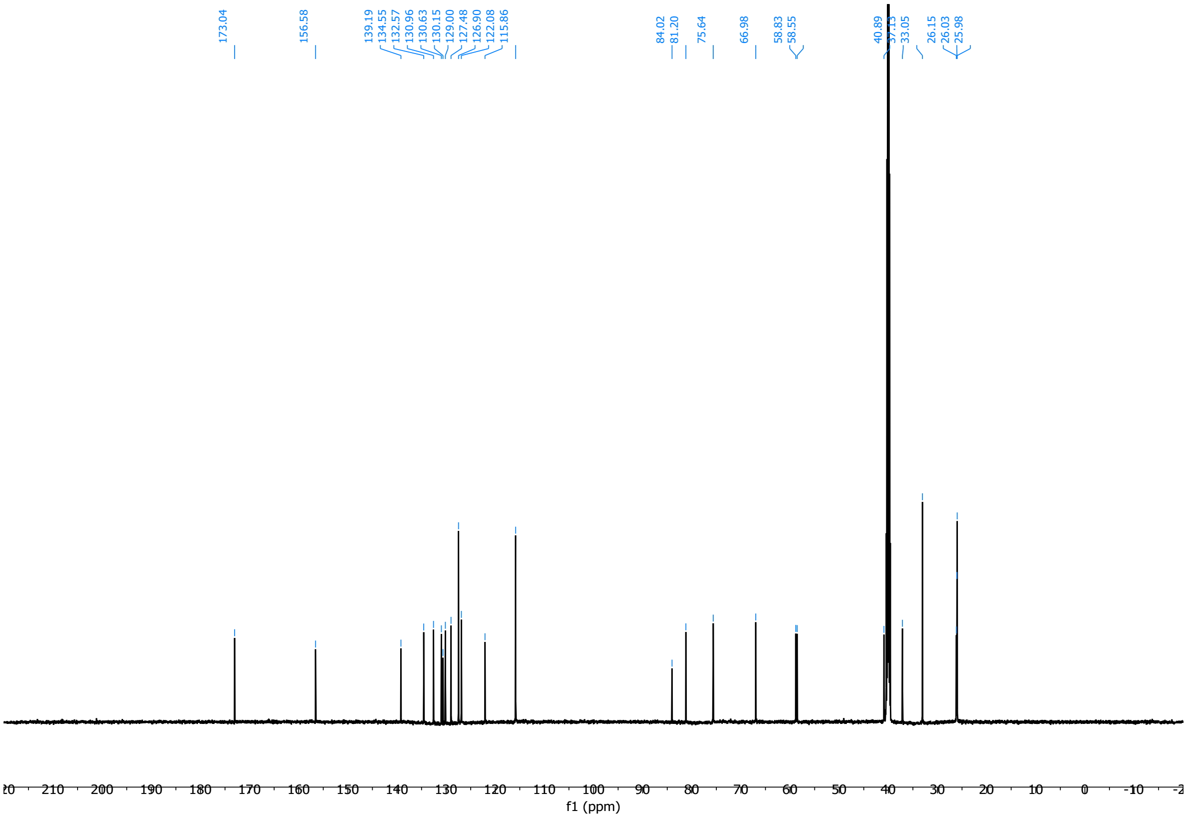

### 8. On-DNA analytical data

#### 8.1. DNA-skeleton conjugate purity

**Supplementary Table 11 | LCMS data for purified skeleton-AOP-headpiece conjugates.** [3H] denotes DNA attachment point

| i | Tag (5’-3’) | smiles | %AUC | rt_min |
|---|---|---|---|---|
| 1 | GCACCATCT | [3H]CNC(=O)[C@@H]1NC[C@@H]1c1ccc(I)cc1 | 100 | 2.31 |
| 2 | GCACTCTCT | [3H]CNC(=O)[C@H]1NC[C@H]1c1ccc(I)cc1 | 100 | 2.31 |
| 3 | TCTCGCTCT | [3H]CNC(=O)[C@H]1NC[C@@H]1c1ccc(I)cc1 | 96 | 2.4 |
| 4 | ATCTCGGCT | [3H]CNC(=O)[C@@H]1NC[C@H]1c1ccc(I)cc1 | 98 | 2.41 |
| 5 | ATGGAGGCT | [3H]CNC(=O)[C@@H]1NCC[C@@H]1c1ccc(I)cc1 | 100 | 2.34 |
| 6 | CGCACAACT | [3H]CNC(=O)[C@H]1NCC[C@H]1c1ccc(I)cc1 | 100 | 2.34 |
| 7 | CGCACTTCT | [3H]CNC(=O)[C@H]1NCC[C@@H]1c1ccc(I)cc1 | 99 | 2.4 |
| 8 | TGGAAGCCT | [3H]CNC(=O)[C@@H]1NCC[C@H]1c1ccc(I)cc1 | 97 | 2.4 |
| 9 | TGGAGACCT | [3H]CNC(=O)[C@@H]1NC[C@@H]1c1cccc(I)c1 | 99 | 2.29 |
| 10 | AGATGGCCT | [3H]CNC(=O)[C@H]1NC[C@H]1c1cccc(I)c1 | 99 | 2.29 |
| 11 | AGGACTGCT | [3H]CNC(=O)[C@H]1NC[C@@H]1c1cccc(I)c1 | 96 | 2.33 |
| 12 | AGTACCGCT | [3H]CNC(=O)[C@@H]1NC[C@H]1c1cccc(I)c1 | 100 | 2.33 |
| 13 | CTCTACGCT | [3H]CNC(=O)[C@@H]1NCC[C@@H]1c1cccc(I)c1 | 100 | 2.33 |
| 14 | GACAGACCT | [3H]CNC(=O)[C@H]1NCC[C@H]1c1cccc(I)c1 | 100 | 2.33 |
| 15 | GGCAACACT | [3H]CNC(=O)[C@H]1NCC[C@@H]1c1cccc(I)c1 | 100 | 2.35 |

| | | | | |
|---|---|---|---|---|
| 16 | GACCGAACT | [3H]CNC(=O)[C@@H]1NCC[C@H]1c1cccc(I)c1 | 100 | 2.35 |
| 17 | GAGCTGACT | [3H]CNC(=O)[C@@H]1NCCC[C@@H]1c1cccc(I)c1 | 100 | 2.39 |
| 18 | GCGAAGACT | [3H]CNC(=O)[C@H]1NCCC[C@H]1c1cccc(I)c1 | 100 | 2.39 |
| 19 | GGCGATTCT | [3H]CNC(=O)[C@H]1NCCC[C@@H]1c1cccc(I)c1 | 100 | 2.4 |
| 20 | TAGACCGCT | [3H]CNC(=O)[C@@H]1NCCC[C@H]1c1cccc(I)c1 | 100 | 2.4 |
| 21 | TCACGACCT | [3H]COC[C@@H]1[C@@H](c2ccc(Br)cc2)[C@@H]2CNCCCCN12 | 100 | 2.37 |
| 22 | TCCACCACT | [3H]COC[C@@H]1[C@@H](c2ccc(Br)cc2)[C@@H]2CNCC(=O)N12 | 100 | 2.33 |
| 23 | TCGCTGTCT | [3H]COC[C@H]1[C@@H](c2ccc(Br)cc2)[C@H]2CNCCCCN21 | 99 | 2.3 |
| 24 | TCTTGGCCT | [3H]COC[C@@H]1[C@H](c2ccc(Br)cc2)[C@@H]2CNCC(=O)N12 | 100 | 2.27 |
| 25 | TGACGGACT | [3H]COC[C@@H]1[C@@H](c2ccc(Br)cc2)[C@H]2CNCC(=O)N21 | 99 | 2.32 |
| 26 | GATGCTCCT | [3H]COC[C@H]1[C@H](c2ccc(Br)cc2)[C@@H]2CNCC(=O)N21 | 100 | 2.32 |
| 27 | GAGGTTCCT | [3H]COC[C@H]1[C@H](c2ccc(Br)cc2)[C@H]2CNCC(=O)N21 | 99 | 2.32 |
| 28 | TACGGTGCT | [3H]COC[C@@H]1[C@H](c2ccc(Br)cc2)[C@H]2CNCCCCN21 | 98 | 2.24 |
| 29 | GCTGCAACT | [3H]COC[C@@H]1[C@H](c2ccc(Br)cc2)[C@@H]2CNCCCCN12 | 100 | 2.3 |
| 30 | GCTTAGGCT | [3H]COC[C@@H]1[C@@H](c2ccc(Br)cc2)[C@H]2CNCCCCN21 | 99 | 2.25 |
| 31 | GGTCTGACT | [3H]COC[C@H](C)N1C[C@H](C)[C@H](CNC)Oc2cc(Br)ccc2S1(=O)=O | 100 | 2.26 |
| 32 | ACTTCGCCT | [3H]COC[C@@H](C)N1C[C@H](C)[C@H](CNC)Oc2cc(Br)ccc2S1(=O)=O | 99 | 2.31 |
| 33 | AGACGCACT | [3H]COC[C@H](C)N1C[C@@H](C)[C@@H](CNC)Oc2cc(Br)ccc2S1(=O)=O | 99 | 2.31 |
| 34 | AGCGAGACT | [3H]COC[C@@H](C)N1C[C@@H](C)[C@@H](CNC)Oc2cc(Br)ccc2S1(=O)=O | 100 | 2.26 |
| 35 | AGTCGTGCT | [3H]COC[C@@H](C)N1C[C@H](C)[C@@H](CNC)Oc2cc(Br)ccc2S1(=O)=O | 93 | 2.34 |
| 36 | ATCCAGCCT | [3H]COC[C@@H](C)N1C[C@@H](C)[C@H](CNC)Oc2cc(Br)ccc2S1(=O)=O | 100 | 2.31 |
| 37 | CGAGTCTCT | [3H]COC[C@H](C)N1C[C@H](C)[C@@H](CNC)Oc2cc(Br)ccc2S1(=O)=O | 100 | 2.31 |
| 38 | CAGCGATCT | [H]OC(=O)c1cc2c(c(-c3ccc(Br)cc3)n1)[C@@H](CCOC[3H])NC2 | 100 | 2.24 |
| 39 | CAGCTACCT | [H]OC(=O)c1cc2c(c(-c3cccc(Br)c3)n1)[C@H](CCOC[3H])NC2 | 100 | 2.25 |
| 40 | CATACCGCT | [3H]CNC(=O)c1cc2c(c(-c3ccc(Br)cc3)n1)[C@@H](CCO)NC2 | 91 | 2.34 |
| 41 | CGACAGTCT | [3H]CNC(=O)c1cc2c(c(-c3cccc(Br)c3)n1)[C@H](CCO)NC2 | 90 | 2.33 |
| 42 | CGACCAACT | [3H]CC(=O)N1C[C@H](c2ccc(I)cc2)C2(CNC2)C1 | 96 | 2.23 |
| 43 | GCTTCCACT | [3H]CC(=O)N1C[C@@H](c2ccc(I)cc2)C2(CNC2)C1 | 98 | 2.23 |
| 44 | CTGAGGACT | [3H]CC(=O)N1CC2(CNC[C@@H]2c2ccc(I)cc2)C1 | 99 | 2.21 |
| 45 | GACCGTTCT | [3H]CC(=O)N1CC2(CNC[C@H]2c2ccc(I)cc2)C1 | 99 | 2.21 |
| 46 | GAGTAGCCT | [3H]CNC(=O)[C@@H]1C[C@H](Oc2ccc(I)cc2)CN1 | 97 | 2.37 |
| 47 | GCTCGAACT | [3H]CNC(=O)[C@@H]1C[C@@H](Oc2ccc(I)cc2)CN1 | 99 | 2.39 |
| 48 | GCTGCTTCT | [3H]CNC(=O)[C@H]1C[C@@H](Oc2ccc(I)cc2)CN1 | 99 | 2.37 |
| 49 | GCTGTCTCT | [3H]CNC(=O)[C@H]1C[C@H](Oc2ccc(I)cc2)CN1 | 99 | 2.4 |

| | | | | |
|---|---|---|---|---|
| 50 | TACGAGCCT | [3H]CNC(=O)[C@@H]1C[C@H](Oc2cccc(I)c2)CN1 | 100 | 2.4 |
| 51 | TACGCGACT | [3H]CNC(=O)[C@@H]1C[C@@H](Oc2cccc(I)c2)CN1 | 100 | 2.4 |
| 52 | TCCTCTGCT | [3H]CNC(=O)[C@H]1C[C@@H](Oc2cccc(I)c2)CN1 | 99 | 2.38 |
| 53 | TCGGCATCT | [3H]CNC(=O)[C@H]1C[C@H](Oc2cccc(I)c2)CN1 | 100 | 2.4 |
| 54 | TCGGTCACT | [3H]CNC(=O)[C@@H]1C[C@H](OCc2ccc(I)cc2)CN1 | 100 | 2.41 |
| 55 | TCTGCCTCT | [3H]CNC(=O)[C@@H]1C[C@@H](OCc2ccc(I)cc2)CN1 | 100 | 2.41 |
| 56 | TTGGCGTCT | [3H]CNC(=O)[C@H]1C[C@@H](OCc2ccc(I)cc2)CN1 | 100 | 2.41 |
| 57 | TTGGTGCCT | [3H]CNC(=O)[C@H]1C[C@H](OCc2ccc(I)cc2)CN1 | 100 | 2.41 |
| 58 | AAGGCCACT | [3H]CNC(=O)[C@@H]1C[C@H](OCc2cccc(I)c2)CN1 | 100 | 2.41 |
| 59 | ACACCGACT | [3H]CNC(=O)[C@@H]1C[C@@H](OCc2cccc(I)c2)CN1 | 100 | 2.41 |
| 60 | AGAACGGCT | [3H]CNC(=O)[C@H]1C[C@@H](OCc2cccc(I)c2)CN1 | 99 | 2.41 |
| 61 | AGAGTGCCT | [3H]CNC(=O)[C@H]1C[C@H](OCc2cccc(I)c2)CN1 | 100 | 2.41 |

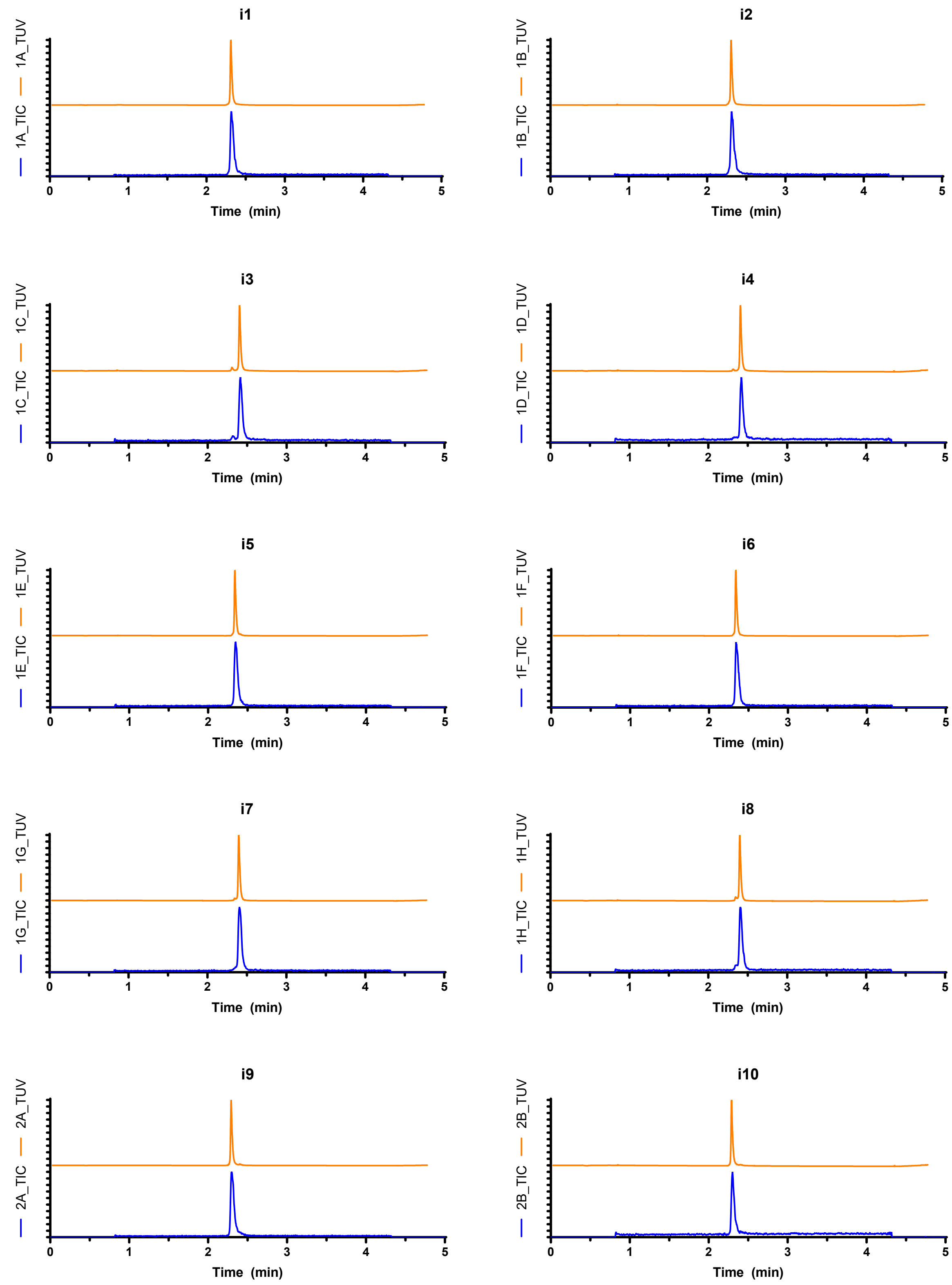
i1
1A_TIC
1A_TUV
Time (min)
i2
1B_TIC
1B_TUV
Time (min)
i3
1C_TIC
1C_TUV
Time (min)
i4
1D_TIC
1D_TUV
Time (min)
i5
1E_TIC
1E_TUV
Time (min)
i6
1F_TIC
1F_TUV
Time (min)
i7
1G_TIC
1G_TUV
Time (min)
i8
1H_TIC
1H_TUV
Time (min)
i9
2A_TIC
2A_TUV
Time (min)
i10
2B_TIC
2B_TUV
Time (min)

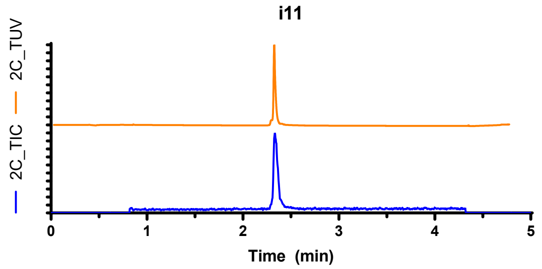
i11
2C_TUV
2C_TIC
0 1 2 3 4 5
Time (min)

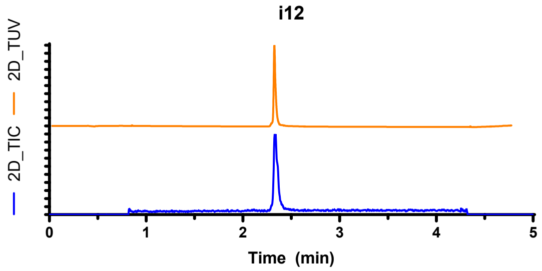
i12
2D_TUV
2D_TIC
0 1 2 3 4 5
Time (min)

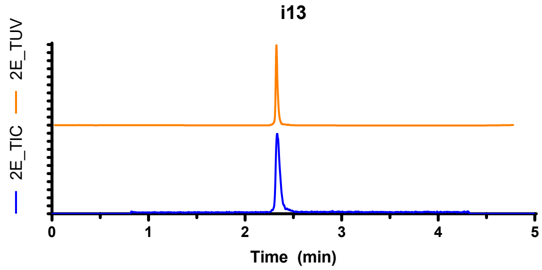
i13
2E_TUV
2E_TIC
0 1 2 3 4 5
Time (min)

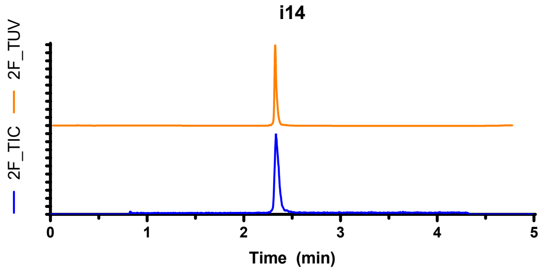
i14
2F_TUV
2F_TIC
0 1 2 3 4 5
Time (min)

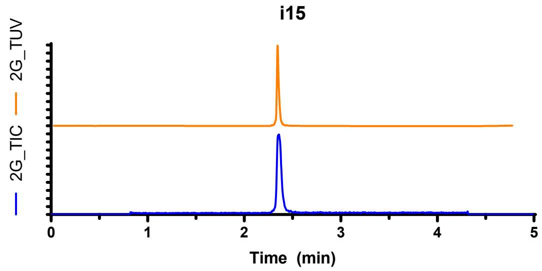
i15
2G_TUV
2G_TIC
0 1 2 3 4 5
Time (min)

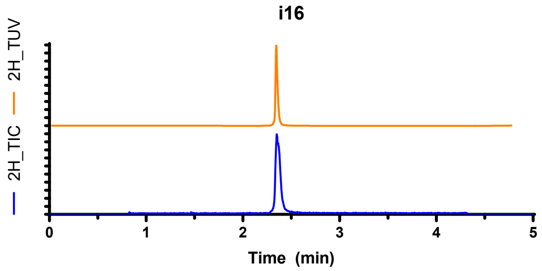
i16
2H_TUV
2H_TIC
0 1 2 3 4 5
Time (min)

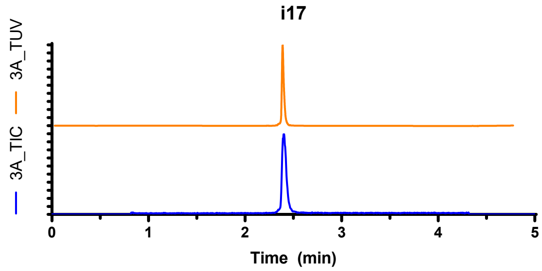
i17
3A_TUV
3A_TIC
0 1 2 3 4 5
Time (min)

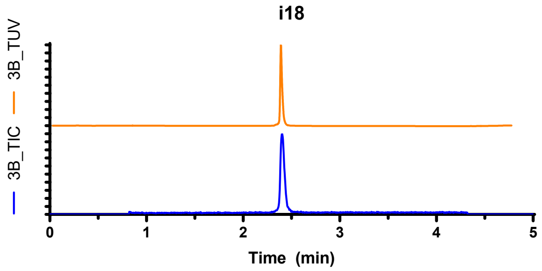
i18
3B_TUV
3B_TIC
0 1 2 3 4 5
Time (min)

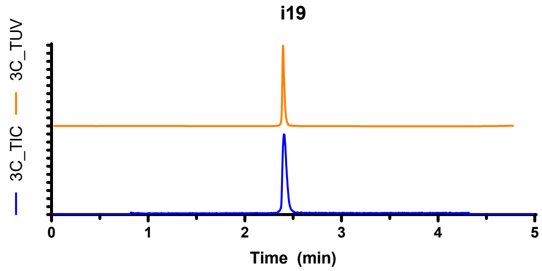
i19
3C_TUV
3C_TIC
0 1 2 3 4 5
Time (min)

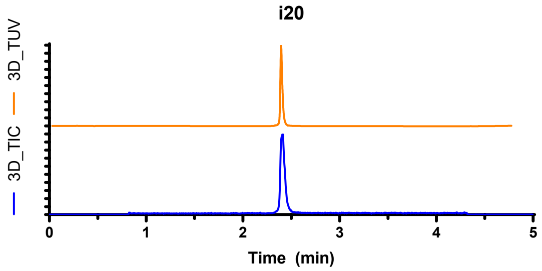
i20
3D_TUV
3D_TIC
0 1 2 3 4 5
Time (min)

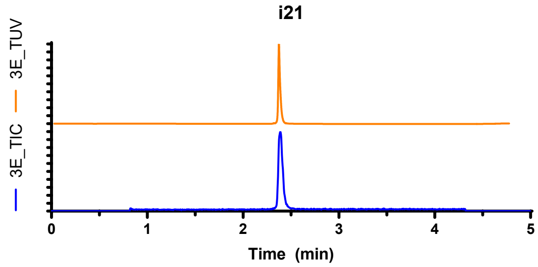
i21
3E_TIC
3E_TUV
0 1 2 3 4 5
Time (min)

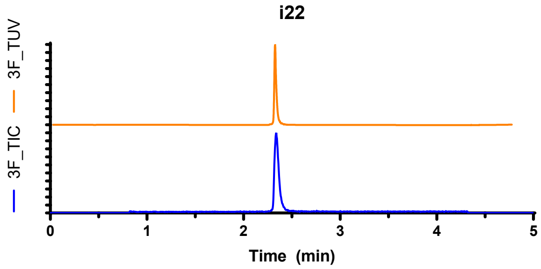
i22
3F_TIC
3F_TUV
0 1 2 3 4 5
Time (min)

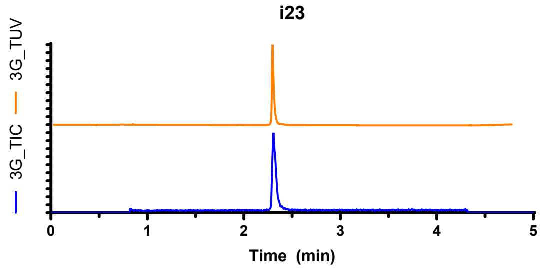
i23
3G_TIC
3G_TUV
0 1 2 3 4 5
Time (min)

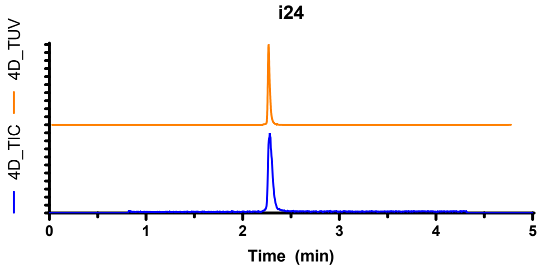
i24
4D_TIC
4D_TUV
0 1 2 3 4 5
Time (min)

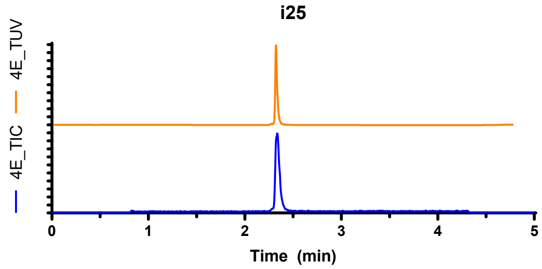
i25
4E_TIC
4E_TUV
0 1 2 3 4 5
Time (min)

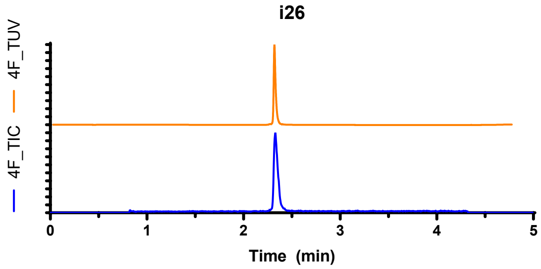
i26
4F_TIC
4F_TUV
0 1 2 3 4 5
Time (min)

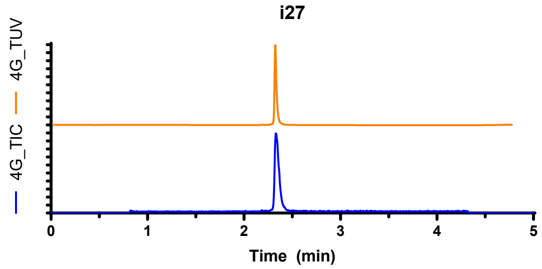
i27
4G_TIC
4G_TUV
0 1 2 3 4 5
Time (min)

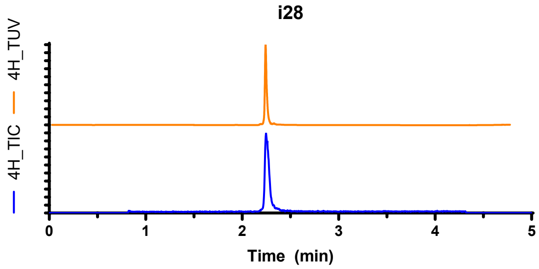
i28
4H_TIC
4H_TUV
0 1 2 3 4 5
Time (min)

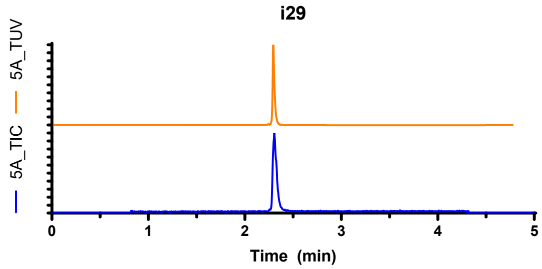
i29
5A_TIC
5A_TUV
0 1 2 3 4 5
Time (min)

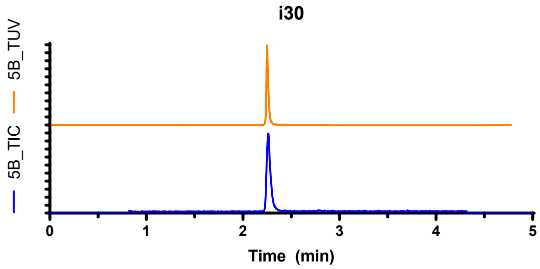
i30
5B_TIC
5B_TUV
0 1 2 3 4 5
Time (min)

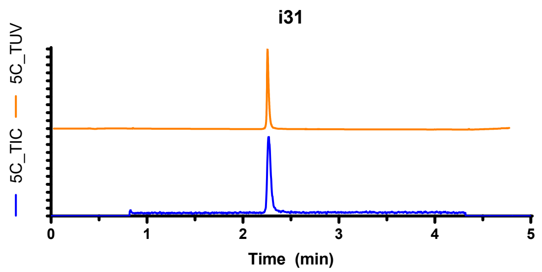
i31
5C_TUV
5C_TIC
0
1
2
3
4
5
Time (min)

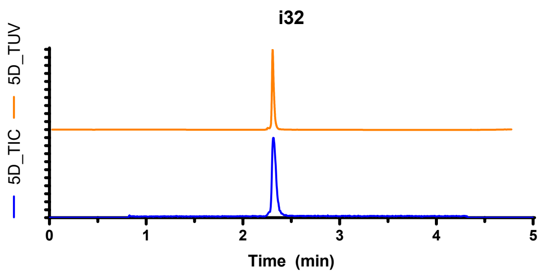
i32
5D_TUV
5D_TIC
0
1
2
3
4
5
Time (min)

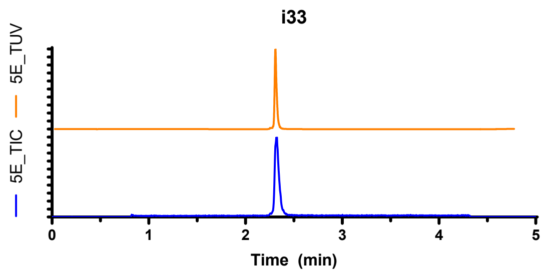
i33
5E_TUV
5E_TIC
0
1
2
3
4
5
Time (min)

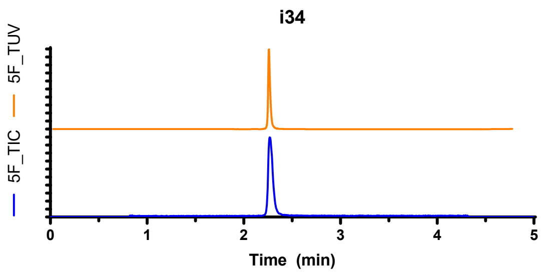
i34
5F_TUV
5F_TIC
0
1
2
3
4
5
Time (min)

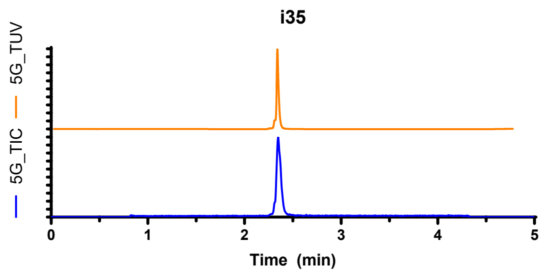
i35
5G_TUV
5G_TIC
0
1
2
3
4
5
Time (min)

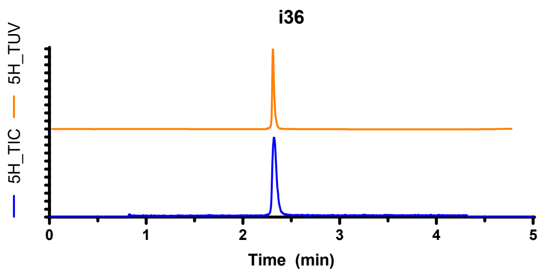
i36
5H_TUV
5H_TIC
0
1
2
3
4
5
Time (min)

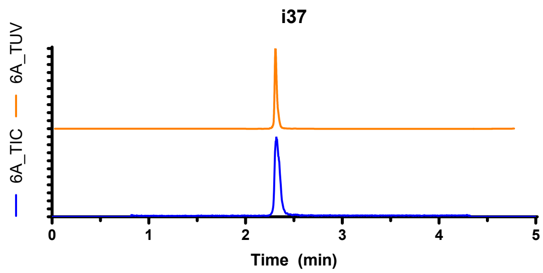
i37
6A_TUV
6A_TIC
0
1
2
3
4
5
Time (min)

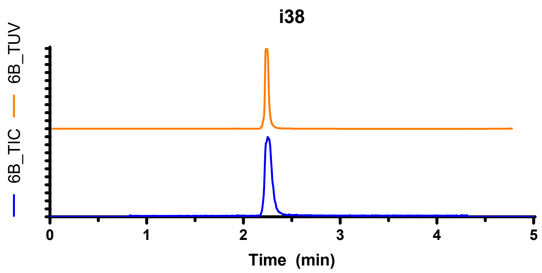
i38
6B_TUV
6B_TIC
0
1
2
3
4
5
Time (min)

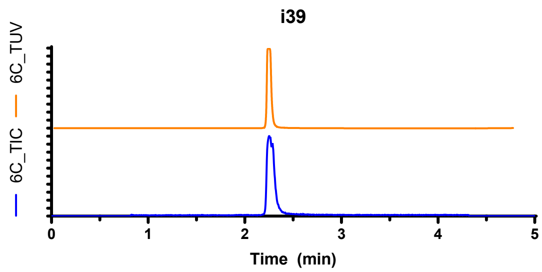
i39
6C_TUV
6C_TIC
0
1
2
3
4
5
Time (min)

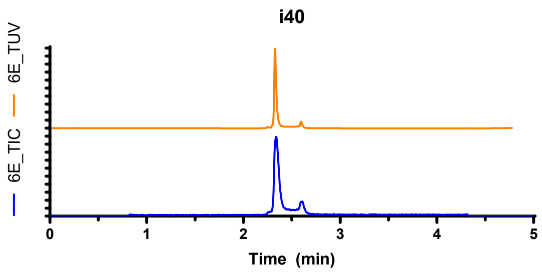
i40
6E_TUV
6E_TIC
0
1
2
3
4
5
Time (min)

i41

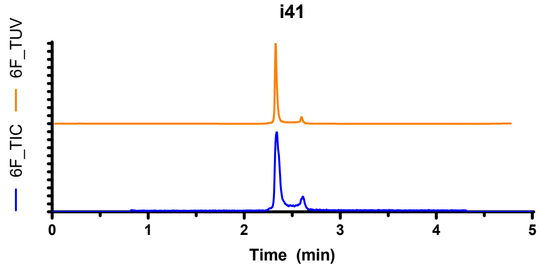

i42

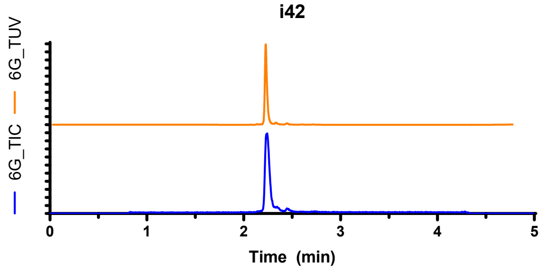

i43

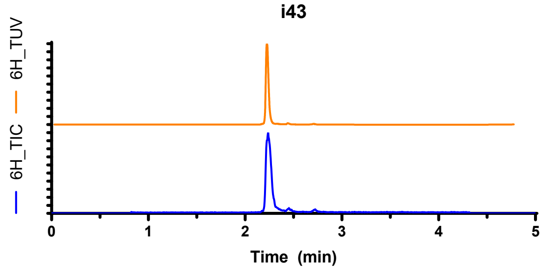

i44

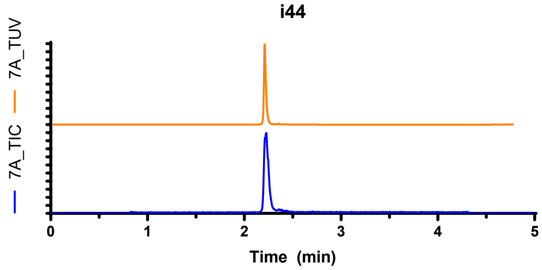

i45

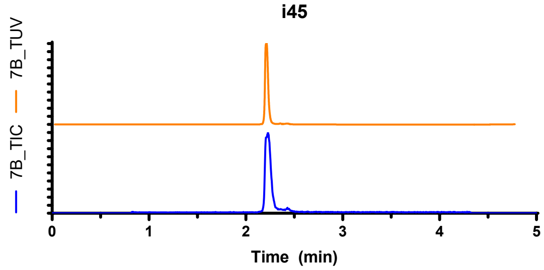

i46

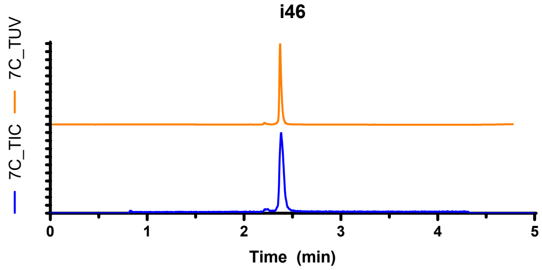

i47

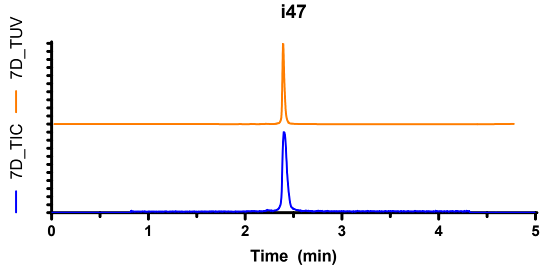

i48

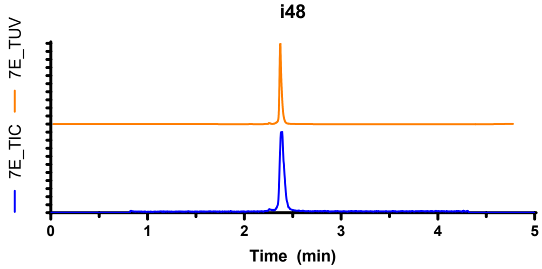

i49

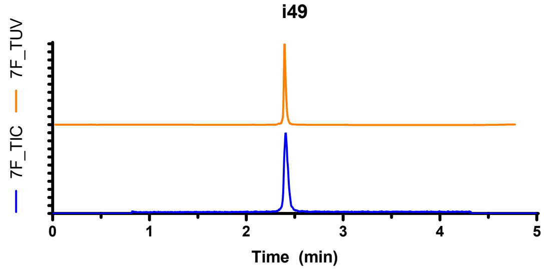

i50

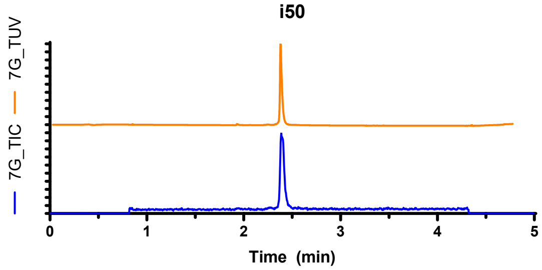

i51

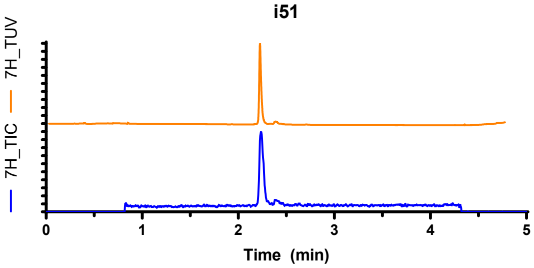

i52

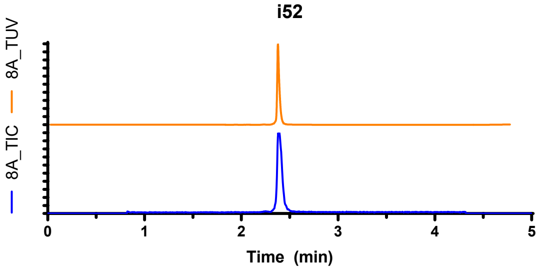

i53

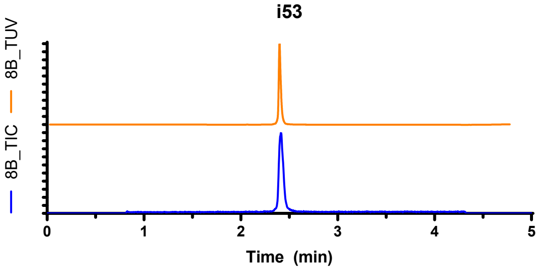

i54

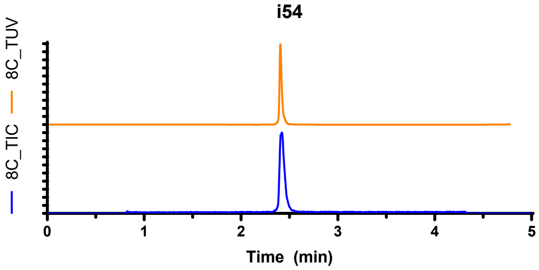

i55

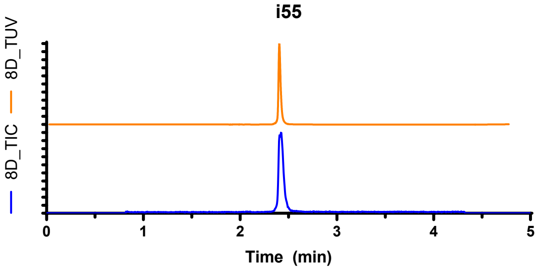

i56

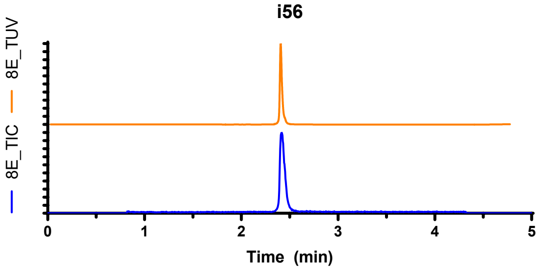

i57

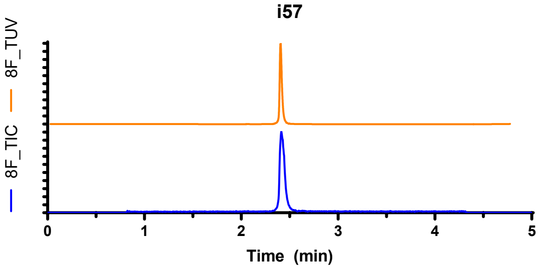

i58

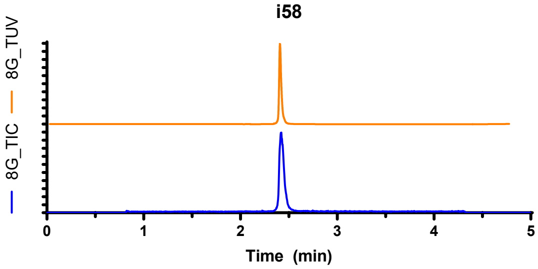

i59

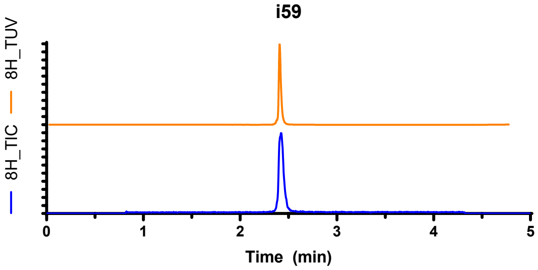

i60

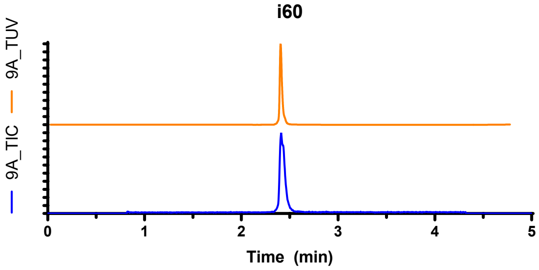

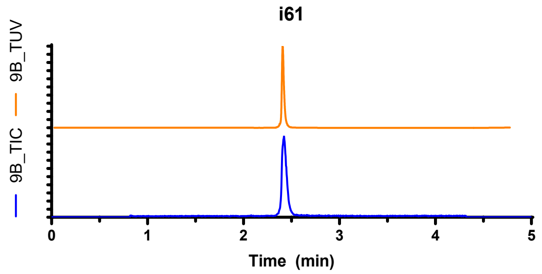

#### 8.2. Amine capping building block validation data

Building block validation data presented below only displays the two calculated factors “P_adj%” and “unknown_adj%”. “P_adj%” is the %AUC of product, or known product-derived species (e.g., carboxylic ester hydrolysis or TFA protecting group removal) when corrected for non-DNA species which may absorb at 260 nm and therefore be integrated as peaks. Non-DNA species were manually discounted based on having no analogous peak in the TIC trace, i.e., with a mass below the lower threshold of m/z monitoring (m/z = 500). “unknown_adj%” is the %AUC of any unknown DNA-based species (i.e., the mass data obtained was confounding, or simply could not be assigned to a known species) when corrected for peaks assigned as residual small molecule building block.

**Supplementary Table 12 | Amine capping building block validation data**

| Structure | P_adj% | unknown_adj% |
|---|---|---|
| **Cn1cnn(CCC(=O)O)c1=O** | 100 | 0 |
| **C#CC1(O)CC(C(=O)O)C1** | 100 | 0 |
| **O=C1Cc2c(C(=O)O)ccnc2N1** | 100 | 0 |
| **C/C(=C\c1ccccn1)C(=O)O** | 100 | 0 |
| **Cn1ccc(-c2cc(C(=O)O)n[nH]2)c1** | 100 | 0 |
| **Cc1nc2ccc(C(=O)O)cc2n1C** | 100 | 0 |
| **Cn1nnc2c1NCCC2C(=O)O** | 99 | 0 |
| **O=C(O)c1nc2n(n1)CCCN2** | 99 | 0 |
| **O=C(O)c1cn(C2CCC2O)nn1** | 99 | 1 |
| **Cn1nnc2c(C(=O)O)ccnc21** | 99 | 1 |
| **O=C(O)c1ncoc1C1CCCO1** | 99 | 0 |
| **CCC(O)Cn1cc(C(=O)O)nn1** | 99 | 1 |
| **COCc1nc(C(=O)O)co1** | 99 | 1 |
| **CCn1nc(C)c(C(=O)O)n1** | 99 | 1 |
| **O=C(O)c1cc(C2CC2)[nH]c(=O)c1** | 99 | 0 |
| **Cn1cc(C(=O)O)c(=O)n(C)c1=O** | 99 | 1 |
| **CC1Cc2cccc(C(=O)O)c2O1** | 99 | 1 |
| **O=C(O)c1cccc(-c2nnco2)c1** | 99 | 0 |
| **CS(=O)(=O)c1cc(C(=O)O)co1** | 99 | 1 |
| **CC1=C(C(=O)O)C(=O)OC1(C)C** | 99 | 0 |
| **Cn1ncc(CCC(=O)O)n1** | 99 | 1 |
| **O=C(O)CC1COc2ccccc21** | 99 | 1 |
| **O=C(O)c1ncc2c(F)cccn12** | 99 | 0 |
| **O=C(O)c1n[nH]c2c1COCC2** | 99 | 1 |
| **Cc1csc(/C=C/C(=O)O)n1** | 99 | 0 |
| **C/C(=C(/F)C(=O)O)C1CC1** | 99 | 0 |

| Structure | P_adj% | unknown_adj% |
|---|---|---|
| **O=C(O)c1coc2c1C(=O)CCC2** | 99 | 1 |
| **Cn1nc(CC(=O)O)c2ccccc21** | 99 | 1 |
| **Cc1cccc2nc(C(=O)O)cn12** | 99 | 1 |
| **CCOc1cccc(CC(=O)O)n1** | 99 | 0 |
| **O=C(O)c1cccc(-n2ccnn2)c1** | 99 | 1 |
| **O=C(O)c1nsc2ccccc12** | 99 | 1 |
| **O=C(O)CCc1cc2ccccc2o1** | 99 | 1 |
| **O=C(O)c1ccc2scnc2c1** | 99 | 1 |
| **O=C(O)c1coc(-c2ccccn2)n1** | 99 | 1 |
| **O=C(O)C1=C[C@@H](O)[C@@H](O)[C@H](O)C1** | 98 | 0 |
| **O=C(O)c1cc([N+](=O)[O-])cnc1O** | 98 | 0 |
| **Cc1onc(CO)c1C(=O)O** | 98 | 0 |
| **CC(=O)Nc1cccc(C(=O)O)n1** | 98 | 0 |
| **O=C1NC(=O)C2(CC(C(=O)O)C2)N1** | 98 | 1 |
| **O=C(O)c1csnc1Cl** | 98 | 2 |
| **O=C(O)CC1=CCOCC1** | 98 | 2 |
| **O=C(O)C1(n2cccn2)CC1** | 98 | 1 |
| **O=C(O)c1cnc2[nH]c(=O)[nH]c2c1** | 98 | 2 |
| **O=C(O)c1cc2n(n1)CCCC2O** | 98 | 1 |
| **Cc1cc(C2(C(=O)O)CC2)on1** | 98 | 3 |
| **COCc1ccc(C(=O)O)cc1F** | 98 | 1 |
| **N#Cc1cccc(C2CC2C(=O)O)c1** | 98 | 2 |
| **O=C(O)Cc1csc(C2CC2)n1** | 98 | 0 |
| **Cc1ccsc1C1CC1C(=O)O** | 98 | 0 |
| **O=C(O)C1=C(c2ccccc2)CCC1** | 98 | 2 |
| **O=C(O)c1ncoc1-c1ccon1** | 97 | 3 |
| **CS(=O)(=O)N1CC(C(=O)O)C1** | 97 | 0 |
| **O=C(O)c1[nH]nc([N+](=O)[O-])c1Cl** | 97 | 2 |
| **O=C(O)CCNC(=O)C1CCCO1** | 97 | 3 |
| **Cc1ncsc1COCC(=O)O** | 97 | 3 |
| **O=C(O)CC1CCC2(CCC2)CO1** | 97 | 3 |
| **O=C(O)C1=NN(CCO)C(=O)CC1** | 97 | 0 |
| **CCC(=O)NC1CCC(C(=O)O)C1** | 97 | 3 |
| **O=C(O)CCc1cc(Cl)cs1** | 97 | 3 |
| **COC(C)c1nc(C(=O)O)cs1** | 97 | 3 |
| **CCOc1oc(C)nc1C(=O)O** | 97 | 3 |
| **O=C(O)c1cnc(CC2CC2)s1** | 97 | 1 |
| **Cc1nc2c(C(=O)O)cccc2o1** | 97 | 0 |
| **CN(C)c1cc(C#N)cc(C(=O)O)c1** | 97 | 3 |
| **Cc1nc(CC(=O)O)c(C)s1** | 97 | 3 |
| **O=C(O)c1cc(-n2cccn2)ccn1** | 97 | 1 |
| **O=C(O)c1cc(-c2ccccc2)co1** | 97 | 1 |
| **CC(=O)[C@@H]1C[C@H](CC(=O)O)C1(C)C** | 96 | 2 |
| **O=C(O)COCc1cc[nH]c(=O)c1** | 96 | 4 |
| **O=C(O)c1ccnc(-c2ncc[nH]2)c1** | 96 | 4 |
| **Cc1cc(C)cc(CC(=O)O)c1** | 96 | 4 |
| **O=C(O)/C=C/c1ccc2c(c1)CCO2** | 96 | 0 |
| **Cc1sc(Cl)c(C(=O)O)c1C** | 96 | 0 |
| **O=C(O)Cn1ncc2cccnc21** | 96 | 2 |

| Structure | P_adj% | unknown_adj% |
|---|---|---|
| **O=C(O)C1CCc2nnc(O)n2C1** | 95 | 0 |
| **CC1(C)CC2(CCO1)CC2C(=O)O** | 95 | 0 |
| **O=C(O)c1cc2c(s1)CCOC2** | 95 | 1 |
| **O=C(O)Cc1n[nH]c2c1CCCC2** | 95 | 4 |
| **CC(Oc1cccc(C#N)c1)C(=O)O** | 95 | 0 |
| **O=C(O)CC1C(=O)Nc2ccccc21** | 95 | 0 |
| **O=C(O)CCn1ccc2ccccc21** | 95 | 3 |
| **Cc1nc(CC2CC2)oc1C(=O)O** | 95 | 5 |
| **Cc1c(C(=O)O)nn2c1OCCC2** | 95 | 5 |
| **O=C(O)CN1Cc2ccccc2C1=O** | 95 | 5 |
| **O=C(O)/C=C/c1ccc2[nH]ccc2c1** | 95 | 1 |
| **COc1ccc2cc(C(=O)O)[nH]c2c1** | 95 | 5 |
| **Cc1cc(C(=O)O)c(C)n1C1CC1** | 95 | 5 |
| **O=C(O)/C=C1\CN2CCC1CC2** | 94 | 1 |
| **O=C(O)c1ccc2oc(=O)[nH]c2n1** | 94 | 3 |
| **Cc1oc(C2CC2)nc1C(=O)O** | 94 | 6 |
| **C#CCOc1ccc(C(=O)O)cc1** | 94 | 6 |
| **Cn1nnc(C(=O)O)c1C(F)F** | 93 | 0 |
| **CCN1C(=O)CCC1CC(=O)O** | 93 | 1 |
| **Cc1nn(C)c(C)c1OCC(=O)O** | 93 | 1 |
| **COC1(C(=O)O)CS(=O)(=O)C1** | 93 | 2 |
| **O=C(O)c1conc1C1CCCC1** | 93 | 3 |
| **O=C(O)C1CCc2nnnn2CC1** | 93 | 7 |
| **O=C(O)c1cnoc1-c1ccco1** | 93 | 7 |
| **O=C(O)CCC1CCc2ccccc21** | 93 | 7 |
| **O=C(O)c1ccc2c(c1)B(O)OC2** | 93 | 5 |
| **Cc1cnc2nc(C(=O)O)nn2c1** | 93 | 7 |
| **CCn1nc(C2CC2)cc1C(=O)O** | 93 | 7 |
| **Cc1nn(CC(=O)O)c(=O)o1** | 92 | 8 |
| **COCCN1CC(C(=O)O)CC1=O** | 92 | 8 |
| **Cc1n[nH]c(C)c1C(C)C(=O)O** | 92 | 0 |
| **CC1CCc2nc(C(=O)O)cn2C1** | 92 | 3 |
| **CC(C)Cc1nc(C(=O)O)c[nH]1** | 92 | 1 |
| **O=C(O)c1nc2c(s1)CCC2** | 92 | 4 |
| **O=C(O)c1cc2occc2s1** | 92 | 3 |
| **Cc1nc2cccnn2c1C(=O)O** | 92 | 8 |
| **COc1cc(C)sc1C(=O)O** | 92 | 8 |
| **Cc1nc(C)n(CCCC(=O)O)n1** | 91 | 9 |
| **CCc1cc(C(=O)O)cc(Cl)n1** | 91 | 1 |
| **Cc1ncc(C(=O)O)c(C(C)C)n1** | 91 | 0 |
| **Cc1nc2c([nH]1)CCC(C(=O)O)C2** | 90 | 0 |
| **N#Cc1cc(C(=O)O)cs1** | 90 | 10 |
| **Cc1cc(C(=O)O)c2[nH]ncc2c1** | 90 | 2 |
| **CCNc1cccnc1C(=O)O** | 90 | 10 |
| **CC(C)Oc1cnc(C(=O)O)cn1** | 89 | 11 |
| **Cc1cc2ccc(C(=O)O)cn2c1** | 89 | 0 |
| **O=C(O)C1(Cl)CC2CCC1O2** | 88 | 12 |
| **O=C(O)c1c[nH]nc1C1CCOC1** | 88 | 6 |
| **C#CCNc1ncc(C(=O)O)c(C)n1** | 87 | 1 |

| Structure | P_adj% | unknown_adj% |
|---|---|---|
| **CC1(C)C(C#N)C1(C#N)C(=O)O** | 86 | 7 |
| **O=C(O)c1ccc(=O)n(CCF)n1** | 86 | 8 |
| **O=C(O)C1CNc2ccccc2O1** | 85 | 15 |
| **CS(=O)c1ccccc1C(=O)O** | 84 | 0 |
| **Cc1cn2c(n1)CC(C(=O)O)CC2** | 84 | 1 |
| **O=C(O)c1cccc[n+]1[O-]** | 84 | 11 |
| **O=C(O)C1=NOC(c2ccccc2)C1** | 84 | 1 |
| **CC1(c2cc(C(=O)O)no2)CC1** | 84 | 16 |
| **CC1=C(C(=O)O)CCS1(=O)=O** | 83 | 17 |
| **O=C(O)C1CN2CCCC2CO1** | 82 | 2 |
| **O=C(O)c1cccnc1-n1cncn1** | 82 | 18 |
| **O=C(O)c1cc2sccc2[nH]1** | 82 | 7 |
| **N#Cc1cc(C(=O)O)n2c1CCCC2** | 82 | 18 |
| **CC1=C(C(=O)O)Cc2ccccc2O1** | 81 | 6 |
| **Cc1ccn(C(C)CC(=O)O)n1** | 80 | 2 |
| **Cc1nc[nH]c(=O)c1CCC(=O)O** | 80 | 20 |
| **C=CCNc1ncc(C(=O)O)s1** | 79 | 6 |
| **Cc1nc(C(=O)O)c2n1CCCC2** | 79 | 0 |
| **COCCn1cc(C(=O)O)c(C)n1** | 78 | 1 |
| **Cc1cc(C(=O)O)c2c(C)n[nH]c2n1** | 78 | 0 |
| **N#Cc1nc(C(=O)O)ccc1Cl** | 78 | 7 |
| **O=C(O)C1CC1Cc1ccccc1** | 77 | 10 |
| **O=C(O)c1ccc2n[nH]c(=O)n2c1** | 77 | 23 |
| **O=C(O)c1ocnc1C(F)(F)F** | 75 | 12 |
| **O=C(O)CNC(=O)Cn1cncn1** | 75 | 1 |
| **Cc1nc2sccn2c1C(=O)O** | 75 | 10 |
| **Cn1cccc1/C=C/C(=O)O** | 75 | 6 |
| **O=C(O)c1cc(-c2ccoc2)n[nH]1** | 75 | 25 |
| **CO[C@H]1C[C@H](C(=O)O)N(C(C)=O)C1** | 74 | 4 |
| **O=C1CCC(C(=O)O)n2cccc21** | 73 | 4 |
| **O=C(O)C#Cc1ccc2c(c1)OCO2** | 73 | 3 |
| **Cc1nnc(CCCC(=O)O)n1C** | 72 | 4 |
| **O=C(O)c1c[nH]c2ncncc12** | 72 | 16 |
| **C=C(C)CC(C)C(=O)O** | 72 | 2 |
| **CC(C)c1ccc(C(=O)O)c(=O)[nH]1** | 72 | 0 |
| **O=C(O)c1ccc(-n2ccnc2)nn1** | 72 | 4 |
| **CC(C)(C(=O)O)n1cc(Cl)cn1** | 71 | 1 |
| **CCn1cc(CC(=O)O)nn1** | 71 | 28 |
| **O=C(O)Cn1cc(C2CCC2)nn1** | 71 | 0 |
| **O=C(O)c1coc2cccc(O)c12** | 71 | 2 |
| **O=C(O)c1ncnc2sccc12** | 68 | 1 |
| **CN(C)Cc1ccc(C(=O)O)cn1** | 67 | 0 |
| **Cn1cncc1C1CC1C(=O)O** | 66 | 5 |
| **C#CCCN(C)CC(=O)O** | 65 | 6 |
| **CC(C)c1nnsc1C(=O)O** | 63 | 3 |
| **CC1=NNC(=O)C1CCC(=O)O** | 62 | 0 |
| **O=C(O)CNC(=O)c1ccsc1** | 59 | 1 |
| **Cn1cnc2ccc(C(=O)O)nc21** | 59 | 6 |
| **O=C(O)Cn1cc(CCCO)cn1** | 57 | 0 |

| Structure | P_adj% | unknown_adj% |
|---|---|---|
| **Cc1cc(O)cc(=O)n1CC(=O)O** | 57 | 24 |
| **CC(Cc1cnc[nH]1)C(=O)O** | 57 | 0 |
| **CC(C)(C)n1nnc(C(=O)O)n1** | 56 | 8 |
| **O=C(O)CCCCc1c[nH]nn1** | 56 | 26 |
| **CC(=O)N1CCCCC1CC(=O)O** | 55 | 21 |
| **CC(C)N(C)C(=O)C1(C(=O)O)CC1** | 55 | 6 |
| **O=C1CCC(C(=O)O)O1** | 52 | 29 |
| **O=C(O)c1nc(=O)[nH][nH]1** | 52 | 5 |
| **O=C(O)C#Cc1cccs1** | 52 | 33 |
| **CN(C)c1nc(O)cc(C(=O)O)n1** | 51 | 49 |
| **CC(C)CC1(C(=O)O)CCNC1=O** | 50 | 2 |
| **O=C(O)c1ccc(-c2nnn[nH]2)cn1** | 50 | 0 |
| **CCC/C=C/C(O)C(=O)O** | 48 | 8 |
| **COc1cccc(C(O)C(=O)O)c1** | 47 | 9 |
| **Cn1nc(C2CC2)nc1CC(=O)O** | 46 | 0 |
| **O=C(O)C#CC1CC(=O)C1** | 45 | 10 |
| **CC(C(=O)O)N(C)CC(F)(F)F** | 45 | 1 |
| **CC(=O)c1c(C)[nH]c(C(=O)O)c1C** | 45 | 55 |
| **O=C1NC2(C(=O)O)CC3CC1C2C3** | 42 | 1 |
| **O=C(O)C1(Nc2ccccc2)CC1** | 42 | 36 |
| **CCc1cccc2c(C(=O)O)c[nH]c12** | 41 | 19 |
| **O=C(O)C1OCCn2ccnc21** | 40 | 7 |
| **CC(C)c1nnnn1CC(=O)O** | 39 | 3 |
| **CC1(C)[C@@H]2CC[C@@]1(C(=O)O)C(=O)C2** | 36 | 4 |
| **O=C1Nc2c(C(=O)O)cccc2C1=O** | 36 | 17 |
| **O=C(O)CCC(=O)c1ccc[nH]1** | 35 | 1 |
| **CC(C)c1nnc(/C=C/C(=O)O)o1** | 33 | 56 |
| **C=CC(C)(CCOC)C(=O)O** | 32 | 1 |
| **CC1C(C(=O)O)C(O)C(=O)N1C** | 30 | 2 |
| **O=C(O)Cn1[nH]c(=O)ccc1=O** | 27 | 0 |
| **COC1(C)CN(CC(=O)O)C1** | 22 | 0 |
| **O=C(O)Cc1csc2nccn12** | 20 | 2 |
| **CN1CCC(C(=O)O)c2ccccc21** | 19 | 2 |
| **O=C(O)C(=O)c1ccc(Cl)s1** | 19 | 14 |
| **CN1N=C(CC(=O)O)CC1=O** | 18 | 0 |
| **CCC(C(=O)O)n1cnc(C#N)n1** | 16 | 0 |
| **COCC(=O)N(C)CC(=O)O** | 15 | 8 |
| **CCn1c(C(=O)O)cnc1O** | 13 | 87 |
| **CC(C)(C(=O)O)N1CCOC1=O** | 12 | 0 |
| **O=C(O)CCc1nc2cccnc2[nH]1** | 12 | 36 |
| **O=C(O)c1nc2cccnc2s1** | 11 | 27 |
| **O=C(O)CC(=O)NCC1CC1** | 11 | 89 |
| **N=c1sccn1CC(=O)O** | 8 | 0 |
| **CC(C)(C)c1nc[nH]c1C(=O)O** | 8 | 21 |
| **CCCCN1CCOCC1C(=O)O** | 7 | 9 |
| **COc1cc(CC(=O)O)on1** | 3 | 29 |
| **Cn1ncnc1COCC(=O)O** | 0 | 1 |
| **CCCc1nc(C)c(C(=O)O)s1** | 0 | 3 |
| **N=C1CCCN1CC(=O)O** | 0 | 6 |

| Structure | P_adj% | unknown_adj% |
|---|---|---|
| **CCS(=O)(=O)NCC(=O)O** | 0 | 11 |
| **O=C(O)c1csc2cncn12** | 0 | 26 |
| **CCS(=O)CC(=O)O** | 0 | 49 |
| **Cn1cc(-c2c[nH]cc2C(=O)O)cn1** | 0 | 65 |
| **O=C(O)c1ccnnc1NC1CC1** | 0 | 71 |
| **CC(C)CCCC(=O)C(=O)O** | 0 | 79 |
| **O=C(O)c1cc(=O)n2ncnc2[nH]1** | 0 | 86 |
| **Cc1nn(C)c(C(=O)O)c1[N+](=O)[O-]** | 0 | 86 |
| **Cc1oc([N+](=O)[O-])cc1C(=O)O** | 0 | 100 |
| **CC1=NOC(=O)C1CC(=O)O** | 0 | 100 |
| **CC1=CC(=O)N(CC(=O)O)C1=O** | 0 | 100 |
| **Cc1nc(O)n(CCC(=O)O)c1C** | 0 | 100 |
| **CCCc1noc(CCC(=O)O)n1** | 0 | 100 |
| **O=C(O)c1c[nH]c(=O)s1** | 0 | 100 |
| **CC1(C)NC(=O)N(CC(=O)O)C1=O** | 0 | 0 |
| **CN1C(=O)Cc2ccc(C(=O)O)cc21** | 0 | 0 |
| **CC(O)Cc1c(C(=O)O)[nH][nH]c1=O** | 0 | 1 |
| **Cc1nnc2cc(C(=O)O)ccn12** | 0 | 1 |
| **CN1CCC(C(=O)O)S1(=O)=O** | 0 | 2 |
| **Cc1nc([N+](=O)[O-])cn1CC(=O)O** | 0 | 2 |
| **C[Si](C)(C)C#CCC(=O)O** | 0 | 2 |
| **CC1=NS(=O)(=O)NC=C1C(=O)O** | 0 | 5 |
| **O=C(O)c1cc(=O)c2ccncc2o1** | 0 | 8 |
| **O=C(O)c1cnon1** | 0 | 9 |
| **C=CCCC1(C(=O)O)CC1** | 0 | 9 |
| **CN(C1CCCC1)C(C)(C)C(=O)O** | 0 | 14 |
| **CNC(=O)c1cnn(C)c1C(=O)O** | 0 | 20 |
| **CC1CCN(C(=O)C(=O)O)C1** | 0 | 49 |
| **Cc1[nH]c(=O)[nH]c(=S)c1C(=O)O** | 0 | 91 |
| **CC(OCC(=O)O)c1nccn1C** | 0 | 0 |
| **O=C(O)c1c[nH]c(=S)n1C1CC1** | 0 | 4 |
| **O=C(O)c1ccc(O)c2ncccc12** | 0 | 4 |
| **Cc1cc(NC(=O)C(=O)O)no1** | 0 | 7 |
| **O=C(O)c1ncnc2nc[nH]c12** | 0 | 13 |
| **O=C(O)Cn1cnc2c1CCCC2** | 0 | 20 |
| **CN(C)C(C(=O)O)c1cncn1C** | 0 | 0 |
| **O=C(O)c1[nH]nc(O)c1CCO** | 0 | 0 |
| **CN(CC(=O)O)c1ccc(C#N)cn1** | 0 | 2 |
| **CC(C)(NCCC#N)C(=O)O** | 0 | 38 |
| **CC(NC1(CO)CCCC1)C(=O)O** | 0 | 0 |
| **CCC(C)c1cc(C(=O)O)on1** | 0 | 61 |
| **CN1CCN(CCC(=O)O)CC1** | 0 | 4 |
| **Cc1sc(O)nc1C(=O)O** | 0 | 55 |
| **C#Cc1cccc(NC(C)C(=O)O)c1** | 0 | 19 |
| **O=C(O)CNc1ccc(Cl)nn1** | 0 | 0 |
| **O=C(O)C1CCCc2cccnc21** | 0 | 20 |
| **O=C(O)c1ccc2[nH]nnc2c1O** | 0 | 9 |
| **Cc1ccc(NCCC(=O)O)nc1** | 0 | 12 |
| **CNc1nnc(CC(=O)O)s1** | 0 | 18 |

| Structure | P_adj% | unknown_adj% |
| --- | --- | --- |
| **CC(OCc1ccncc1)C(=O)O** | 0 | 19 |
| **CN(C)Cc1ccsc1C(=O)O** | 0 | 16 |
| **Cn1cc(NCC(=O)O)cn1** | 0 | 32 |
| **CC(C(=O)O)n1cnc2ccccc21** | 0 | 1 |
| **O=C(O)COc1ccc(Cl)cn1** | 0 | 100 |
| **Cn1c(C(=O)O)nc2ncccc21** | 0 | 1 |
| **CCn1cc(S(=O)(=O)Cl)cn1** | 100 | 0 |
| **COc1ccc(S(=O)(=O)Cl)cc1** | 100 | 0 |
| **COc1cccc(S(=O)(=O)Cl)c1** | 100 | 0 |
| **O=C(O)c1ccc(S(=O)(=O)Cl)c([N+](=O)[O-])c1** | 100 | 0 |
| **Nc1ncc(S(=O)(=O)Cl)cn1** | 100 | 0 |
| **O=S(=O)(Cl)c1nnc[nH]1** | 100 | 0 |
| **Cn1cnc(S(=O)(=O)Cl)c1** | 100 | 0 |
| **Cn1nncc1S(=O)(=O)Cl** | 100 | 0 |
| **Cc1nc(S(=O)(=O)Cl)cn1C** | 100 | 0 |
| **Cn1ccnc1S(=O)(=O)Cl** | 100 | 0 |
| **CCn1cnc(S(=O)(=O)Cl)c1** | 100 | 0 |
| **N#Cc1cccc(S(=O)(=O)Cl)c1** | 100 | 0 |
| **COc1cncc(S(=O)(=O)Cl)c1** | 100 | 0 |
| **CC(C)n1cc(S(=O)(=O)Cl)cn1** | 100 | 0 |
| **O=[N+]([O-])c1ccc(S(=O)(=O)Cl)c(Cl)c1** | 100 | 0 |
| **O=S(=O)(Cl)c1cc(F)cc(F)c1** | 100 | 0 |
| **Cc1cc(S(=O)(=O)Cl)c(C)o1** | 100 | 0 |
| **COc1cc(C)c(Cl)cc1S(=O)(=O)Cl** | 100 | 0 |
| **Cc1c(S(=O)(=O)Cl)cnn1C** | 100 | 0 |
| **COc1ccc(S(=O)(=O)Cl)cn1** | 100 | 0 |
| **Cc1cc(S(=O)(=O)Cl)cs1** | 100 | 0 |
| **CC(C)n1cnc(S(=O)(=O)Cl)c1** | 100 | 0 |
| **Cc1ccc(S(=O)(=O)Cl)c(C)c1** | 100 | 0 |
| **Cc1ccc(S(=O)(=O)Cl)cc1C** | 100 | 0 |
| **Cc1cc(C)cc(S(=O)(=O)Cl)c1** | 100 | 0 |
| **Cc1ccc(C)c(S(=O)(=O)Cl)c1** | 100 | 0 |
| **O=S(=O)(Cl)c1cc(Cl)cc(Cl)c1** | 100 | 0 |
| **Cc1ccc(S(=O)(=O)Cl)cc1** | 100 | 0 |
| **C=C(C)COc1cccc(S(=O)(=O)Cl)c1** | 100 | 0 |
| **Cc1cccc(S(=O)(=O)Cl)c1** | 100 | 0 |
| **O=S(=O)(Cl)c1ccc(Br)cc1C(F)(F)F** | 100 | 0 |
| **O=C1COc2cc(S(=O)(=O)Cl)ccc2N1** | 100 | 0 |
| **O=S(=O)(Cl)c1cc(F)cc(Cl)c1** | 100 | 0 |
| **C#CCOc1cccc(S(=O)(=O)Cl)c1** | 100 | 0 |
| **O=c1[nH]c2ccc(S(=O)(=O)Cl)cc2s1** | 100 | 0 |
| **C=Cc1ccc(S(=O)(=O)Cl)cc1** | 100 | 0 |
| **O=c1[nH]c(=O)c2cc(S(=O)(=O)Cl)cnc2[nH]1** | 100 | 0 |
| **O=c1ccc2ccc(S(=O)(=O)Cl)cc2[nH]1** | 100 | 0 |
| **Cc1noc(C)c1S(=O)(=O)Cl** | 99 | 0 |
| **O=S(=O)(Cl)c1cc(F)ccc1F** | 99 | 0 |
| **Cc1ccncc1S(=O)(=O)Cl** | 99 | 1 |
| **CCn1cc(S(=O)(=O)Cl)nc1C** | 99 | 1 |
| **O=S(=O)(Cl)C=Cc1ccccc1** | 99 | 1 |

| Structure | P_adj% | unknown_adj% |
|---|---|---|
| **CCn1ncc(S(=O)(=O)Cl)c1C** | 98 | 2 |
| **O=S(=O)(Cl)c1cnc2n1CCC2** | 98 | 2 |
| **Cc1cnc(N)c(S(=O)(=O)Cl)c1** | 97 | 3 |
| **O=Cc1ccc(S(=O)(=O)Cl)cc1** | 97 | 3 |
| **COc1ccccc1S(=O)(=O)Cl** | 96 | 4 |
| **N#Cc1ccc(S(=O)(=O)Cl)cc1** | 96 | 4 |
| **CCc1ccc(S(=O)(=O)Cl)cc1** | 96 | 4 |
| **O=C(N1CCc2ccc(S(=O)(=O)Cl)cc2C1)C(F)(F)F** | 96 | 4 |
| **C=C/C=C/S(=O)(=O)Cl** | 94 | 6 |
| **O=S(=O)(Cl)/C=C/c1ccccc1** | 94 | 6 |
| **N#Cc1ncccc1S(=O)(=O)Cl** | 92 | 8 |
| **O=S(=O)(Cl)c1ccc(F)c(Cl)c1** | 92 | 8 |
| **N#Cc1ccc(S(=O)(=O)Cl)cn1** | 91 | 9 |
| **N#Cc1cncc(S(=O)(=O)Cl)c1** | 80 | 20 |
| **Cc1ccc(S(=O)(=O)Cl)cc1C(=O)O** | 75 | 25 |
| **Cc1ccccc1S(=O)(=O)Cl** | 73 | 1 |
| **COc1ccncc1S(=O)(=O)Cl** | 72 | 28 |
| **CC(=O)OCCS(=O)(=O)Cl** | 56 | 0 |
| **CC(C)(C#N)CS(=O)(=O)Cl** | 51 | 8 |
| **CCCCS(=O)(=O)Cl** | 43 | 1 |
| **O=S(=O)(Cl)CCOCC1CC1** | 40 | 10 |
| **CC(C)(C)OCCS(=O)(=O)Cl** | 38 | 2 |
| **CCC1(S(=O)(=O)Cl)CC1** | 28 | 7 |
| **CS(=O)(=O)Cl** | 25 | 0 |
| **COCC1(S(=O)(=O)Cl)CC1** | 24 | 76 |
| **COC(=O)CCCS(=O)(=O)Cl** | 24 | 2 |
| **O=S(=O)(Cl)CC1CCOCC1** | 21 | 0 |
| **O=S(=O)(Cl)C1CCCCCC1** | 20 | 23 |
| **CN1CCN(S(=O)(=O)Cl)CC1** | 19 | 0 |
| **CC1(CS(=O)(=O)Cl)N=N1** | 18 | 7 |
| **CC(=O)OCCCS(=O)(=O)Cl** | 18 | 3 |
| **CC(C)C1(S(=O)(=O)Cl)CC1** | 18 | 30 |
| **O=S(=O)(Cl)c1cn[nH]c1** | 17 | 4 |
| **CC(C)(C)CS(=O)(=O)Cl** | 10 | 3 |
| **O=S(=O)(Cl)CC1CCCO1** | 5 | 4 |
| **O=S(=O)(Cl)N1CCCOCC1** | 4 | 3 |
| **CCN(CC1CC1)S(=O)(=O)Cl** | 0 | 6 |
| **CCN(C1CC1)S(=O)(=O)Cl** | 0 | 11 |
| **COC1(CS(=O)(=O)Cl)CCC1** | 0 | 20 |
| **CO[C@@H]1CCC[C@@H]1S(=O)(=O)Cl** | 0 | 22 |
| **COC(C)(C)CCS(=O)(=O)Cl** | 0 | 39 |
| **N#Cc1ccccc1S(=O)(=O)Cl** | 0 | 100 |
| **O=C1C=C(S(=O)(=O)Cl)C(=O)O1** | 0 | 100 |
| **COC(=O)/C=C/S(=O)(=O)Cl** | 0 | 100 |
| **O=[N+]([O-])c1ccccc1S(=O)(=O)Cl** | 0 | 100 |
| **Cn1c(=O)oc2ccc(S(=O)(=O)Cl)cc21** | 0 | 100 |
| **CCOC(=O)CS(=O)(=O)Cl** | 0 | 2 |
| **O=C(O)c1cc(S(=O)(=O)Cl)c[nH]1** | 0 | 2 |
| **COC(=O)NS(=O)(=O)Cl** | 0 | 2 |

| Structure | P_adj% | unknown_adj% |
|---|---|---|
| **O=C1OCCN1S(=O)(=O)Cl** | 0 | 3 |
| **CN(C)C1(CS(=O)(=O)Cl)CC1** | 0 | 3 |
| **CCN(CC)CCS(=O)(=O)Cl** | 0 | 3 |
| **COC(=O)CCS(=O)(=O)Cl** | 0 | 4 |
| **COC(=O)C(C)(C)S(=O)(=O)Cl** | 0 | 4 |
| **O=C1CCC(S(=O)(=O)Cl)CC1** | 0 | 4 |
| **O=S(=O)(Cl)N1CCCCCC1** | 0 | 5 |
| **CCOC(=O)CCS(=O)(=O)Cl** | 0 | 5 |
| **O=S(=O)(Cl)[C@@H]1CCC[C@H]1O** | 0 | 5 |
| **COCCOCCS(=O)(=O)Cl** | 0 | 14 |
| **CC(C)(C)NS(=O)(=O)Cl** | 0 | 3 |
| **N#CCC1(CS(=O)(=O)Cl)CC1** | 0 | 3 |
| **CC(C)OC(=O)CS(=O)(=O)Cl** | 0 | 3 |
| **O=S(=O)(Cl)c1ccccc1Cn1cncn1** | 0 | 4 |
| **C=CCS(=O)(=O)Cl** | 0 | 5 |
| **CN(C)S(=O)(=O)Cl** | 0 | 6 |
| **COCC(C)(C)CS(=O)(=O)Cl** | 0 | 7 |
| **CC(C)(C#N)S(=O)(=O)Cl** | 0 | 10 |
| **CC1(CS(=O)(=O)Cl)COC1** | 0 | 2 |
| **CC1CCN(S(=O)(=O)Cl)CC1** | 0 | 3 |
| **CC(C)(C)OC(=O)N1CCC(S(=O)(=O)Cl)CC1** | 0 | 4 |
| **CO[C@H]1COC[C@H]1S(=O)(=O)Cl** | 0 | 0 |
| **CO[C@@H]1CCC[C@H]1S(=O)(=O)Cl** | 0 | 3 |
| **Cc1ccc(O)c(S(=O)(=O)Cl)c1** | 0 | 8 |
| **CC(C)(C)S(=O)(=O)Cl** | 0 | 0 |
| **O=S(=O)(Cl)Cc1ccc(Cl)cc1Cl** | 0 | 3 |
| **O=S(=O)(Cl)c1nc2ccccc2s1** | 0 | 4 |
| **Cc1ccccc1CS(=O)(=O)Cl** | 0 | 3 |
| **Cc1ccc(CS(=O)(=O)Cl)cc1** | 0 | 3 |
| **Cc1cccc(CS(=O)(=O)Cl)c1** | 0 | 3 |
| **Cc1cccc(S(=O)(=O)Cl)c1C** | 0 | 3 |
| **COc1ccc(S(=O)(=O)Cl)c(C)c1C** | 0 | 4 |
| **O=[N+]([O-])c1cc(S(=O)(=O)Cl)ccc1Cl** | 0 | 100 |
| **O=Cc1c[nH]nn1** | 100 | 0 |
| **CCCC=O** | 100 | 0 |
| **O=CC1CCC1** | 100 | 0 |
| **CC(C)(C)OC(=O)N1CC(C=O)C1** | 100 | 0 |
| **O=CCC1CCOCC1** | 100 | 0 |
| **O=CCOCc1ccccc1** | 100 | 0 |
| **N#Cc1ccc(CCC=O)cc1** | 100 | 0 |
| **NS(=O)(=O)c1cc(C=O)co1** | 100 | 0 |
| **O=Cc1nccn1-c1ccccc1** | 100 | 0 |
| **CS(=O)(=O)c1ccc(C=O)o1** | 100 | 0 |
| **Cc1ccncc1CCC=O** | 100 | 0 |
| **Cc1cccc(C=O)n1** | 100 | 0 |
| **O=Cc1cn(-c2ccccc2)cn1** | 100 | 0 |
| **Cn1cc(-c2cccc(C=O)c2)cn1** | 100 | 0 |
| **O=Cc1ccc(-c2ccn[nH]2)o1** | 100 | 0 |
| **N#CCOc1ccccc1C=O** | 100 | 0 |

| Structure | P_adj% | unknown_adj% |
|---|---|---|
| **O=Cc1ccn(-c2cccnc2)c1** | 100 | 0 |
| **O=Cc1nc(-c2ccccc2)c[nH]1** | 100 | 0 |
| **CC(C)(CC=O)C1CC1** | 99 | 0 |
| **O=Cc1cn[nH]c1** | 99 | 0 |
| **O=Cc1csnn1** | 99 | 0 |
| **O=CC(=O)O** | 99 | 1 |
| **Cc1cc(C=O)cnc1-n1cccn1** | 99 | 0 |
| **O=Cc1ccnn1-c1ccccn1** | 99 | 0 |
| **CCCn1ccc(C=O)n1** | 99 | 0 |
| **O=Cc1ccccc1-n1cccn1** | 99 | 0 |
| **Cc1cccc(-c2[nH]ncc2C=O)c1** | 99 | 0 |
| **O=Cc1cc(C(=O)O)ccc1O** | 98 | 0 |
| **O=Cc1cccc(Cc2c[nH]cn2)c1** | 98 | 0 |
| **O=Cc1cnn(Cc2ccccc2)c1** | 98 | 0 |
| **CNC(=O)c1ccc(C=O)cc1** | 98 | 0 |
| **CS(=O)c1ccc(C=O)cc1** | 98 | 0 |
| **O=Cc1ccc(-n2ccnc2)cc1** | 98 | 0 |
| **O=Cc1ccc(-n2ccnc2)nc1** | 98 | 0 |
| **O=Cc1ccc(C(=O)NC2CC2)cc1** | 98 | 2 |
| **Cn1cc(C=O)cc1C#N** | 98 | 0 |
| **Cn1cc(-n2ccc(C=O)c2)cn1** | 98 | 2 |
| **O=Cc1ccc2cn[nH]c2c1** | 98 | 2 |
| **O=Cc1cnc[nH]1** | 97 | 1 |
| **CS(=O)(=O)c1cccc(C=O)c1** | 97 | 0 |
| **COc1ncccc1C=O** | 97 | 0 |
| **COC(=O)c1cc(C=O)cn1C** | 97 | 3 |
| **O=Cc1cn(-c2ccccc2)nn1** | 97 | 3 |
| **Cc1nn(-c2ccncc2)cc1C=O** | 97 | 3 |
| **CC(C)(O)c1ccc(C=O)cc1** | 97 | 0 |
| **C#Cc1cccc(C=O)c1** | 97 | 3 |
| **Cn1cc(C=O)cn1** | 96 | 0 |
| **Cc1nc2ccccc2nc1C=O** | 96 | 0 |
| **O=Cc1ccc2c(n1)NC(=O)CO2** | 96 | 0 |
| **O=Cc1n[nH]cc1-c1ccccc1** | 96 | 2 |
| **O=Cc1cc(CO)ccc1O** | 95 | 0 |
| **COC(=O)CCC=O** | 95 | 5 |
| **O=CC1CC2(CC2)C1** | 95 | 0 |
| **COc1cncc(CCC=O)c1** | 95 | 5 |
| **COC(C)(C)CC=O** | 95 | 5 |
| **NS(=O)(=O)c1ccc(C=O)o1** | 94 | 5 |
| **COc1cccnc1C=O** | 94 | 3 |
| **O=Cc1ccc(-c2cccnc2)o1** | 94 | 6 |
| **Cc1ccc2cc(C=O)ccc2n1** | 94 | 6 |
| **O=Cc1ccccc1-n1cccc1** | 94 | 0 |
| **CCc1ccc(C=O)cc1[N+](=O)[O-]** | 94 | 0 |
| **O=Cc1cc(C2CC2)n[nH]1** | 94 | 5 |
| **CC(=O)OCc1ccc(C=O)o1** | 93 | 7 |
| **CC(C)(C)c1ccc(CC=O)cc1** | 93 | 1 |
| **Cc1ncc(C=O)n1-c1ccccc1** | 93 | 0 |

| Structure | P_adj% | unknown_adj% |
|---|---|---|
| **CC(C)(C#N)c1ccc(C=O)cc1** | 93 | 5 |
| **COC(=O)c1[nH]cc(C=O)c1C** | 93 | 3 |
| **Cc1ccccc1-n1cc(C=O)cn1** | 93 | 6 |
| **Cn1cc(C=O)c(-c2cccnc2)n1** | 93 | 5 |
| **O=Cc1ccnc(-c2ccccc2)n1** | 93 | 7 |
| **O=CC1CCCCCCC1** | 92 | 0 |
| **COc1cccc2cc(C=O)oc12** | 92 | 8 |
| **COC(=O)c1ccc(C=O)o1** | 92 | 8 |
| **Cc1cc(C)c(C(=O)O)cc1C=O** | 92 | 4 |
| **COc1nc(C)ccc1C=O** | 92 | 6 |
| **O=Cc1c[nH]c(-c2ccccc2)n1** | 92 | 6 |
| **O=Cc1cc(-n2ccnc2)cs1** | 92 | 6 |
| **O=Cc1cnn(-c2ccccn2)c1** | 92 | 7 |
| **O=Cc1ccc2cccc(O)c2n1** | 92 | 8 |
| **O=CC1CCCCCC1** | 91 | 0 |
| **O=Cc1cccc(-c2ccccn2)c1** | 91 | 9 |
| **O=Cc1ccc(O)c(F)c1** | 90 | 0 |
| **COc1nccc(C=O)n1** | 90 | 8 |
| **O=Cc1cn2c(n1)CCC2** | 90 | 1 |
| **O=Cc1cccc2ccoc12** | 90 | 7 |
| **N#CCCCOc1cccc(C=O)c1** | 90 | 7 |
| **Cc1[nH]nc(Cl)c1C=O** | 89 | 0 |
| **CCOC(=O)c1cc(C=O)[nH]n1** | 89 | 3 |
| **O=Cc1cc(-c2ccccn2)n[nH]1** | 89 | 11 |
| **O=CC[C@@H](O)[C@@H](O)CO** | 88 | 0 |
| **O=CC[C@H](O)[C@H](O)CO** | 88 | 0 |
| **COC(=O)c1cccc(C=O)n1** | 88 | 12 |
| **COc1nc2ccccc2cc1C=O** | 88 | 5 |
| **O=Cc1cccc(-c2nn[nH]n2)c1** | 88 | 3 |
| **Cn1nc(-c2ccccc2)cc1C=O** | 88 | 1 |
| **O=Cc1ccc(-c2nn[nH]n2)cc1** | 87 | 0 |
| **O=Cc1occc1C(=O)O** | 86 | 0 |
| **Cn1cnc(C=O)c1** | 86 | 0 |
| **NS(=O)(=O)c1ccc(C=O)cc1** | 86 | 11 |
| **CC(C)(C)c1cc(O)cc(C=O)c1** | 86 | 11 |
| **O=Cc1c[nH]nc1-c1ccccc1** | 86 | 6 |
| **Cc1cc(C=O)cc2cccnc12** | 86 | 12 |
| **O=Cc1ccn(-c2ccccc2)n1** | 86 | 14 |
| **O=Cc1c[nH]nc1-c1cccs1** | 85 | 15 |
| **O=CC[C@@H](O)[C@H](O)[C@H](O)CO** | 84 | 0 |
| **Cc1ccc(CCC=O)cn1** | 84 | 0 |
| **CCOc1cc(C=O)ccc1OC** | 84 | 0 |
| **O=CC1CC1** | 83 | 0 |
| **O=Cc1cc(C(F)(F)F)[nH]n1** | 83 | 6 |
| **O=Cc1ccc2c(c1)CCC2** | 80 | 0 |
| **N#Cc1csc(C=O)c1** | 80 | 0 |
| **O=Cc1cc(O)c2ccsc2c1** | 80 | 8 |
| **O=Cc1ccnc(O)c1** | 79 | 0 |
| **Cc1c(C(F)(F)F)n[nH]c1C=O** | 79 | 4 |

| Structure | P_adj% | unknown_adj% |
|---|---|---|
| **O=Cc1cc(O)c2ccoc2c1** | 77 | 21 |
| **O=Cc1cc2cc[nH]c2cn1** | 76 | 9 |
| **CCCn1cc(C=O)c(C2CC2)n1** | 76 | 0 |
| **O=Cc1ccc2nccnc2c1** | 75 | 0 |
| **COc1ccc(C=O)c(OC)c1** | 73 | 16 |
| **O=C[C@H](O)CO** | 72 | 0 |
| **O=Cc1cc(Cl)c(O)cc1F** | 71 | 26 |
| **Cn1nccc1-c1ccccc1C=O** | 71 | 3 |
| **O=Cc1ccc([N+](=O)[O-])cn1** | 69 | 8 |
| **Cn1cc(-c2ncc(C=O)cn2)cn1** | 67 | 15 |
| **CC(C)c1cc(O)ccc1C=O** | 66 | 0 |
| **COc1cc(Cl)cc(C=O)c1O** | 62 | 2 |
| **CN(CCO)c1ccc(C=O)cn1** | 62 | 0 |
| **Cc1ccc(C=O)c2ccccc12** | 61 | 11 |
| **O=Cc1cc[n+]([O-])cc1** | 60 | 1 |
| **N#CC1(C=O)CC1** | 60 | 4 |
| **N#Cc1ccc(C=O)s1** | 60 | 0 |
| **COc1cc(C)c(C=O)cc1C** | 59 | 0 |
| **O=Cc1ccc(N2CCCC2)nc1** | 59 | 0 |
| **CCCCc1nc(C=O)c(Cl)[nH]1** | 59 | 0 |
| **O=Cc1nccn1Cc1ccccc1** | 57 | 1 |
| **O=Cc1ccc(O)nc1** | 56 | 0 |
| **CCOC(=O)c1[nH]c(C=O)cc1C** | 56 | 2 |
| **CC(C)C=O** | 54 | 1 |
| **Nc1ccc2cc(C=O)ccc2n1** | 54 | 3 |
| **O=Cc1cnc(N2CCCC2)nc1** | 53 | 0 |
| **COc1ccc(C=O)cc1C#N** | 51 | 2 |
| **O=Cc1cc2sccc2s1** | 50 | 0 |
| **O=Cc1ncc[nH]1** | 49 | 0 |
| **CS(=O)(=O)n1cccc1C=O** | 47 | 0 |
| **O=C[C@H](O)[C@H](O)CO** | 44 | 11 |
| **CC1(C=O)COC1** | 44 | 0 |
| **Cn1cc(-c2n[nH]cc2C=O)cn1** | 41 | 59 |
| **O=Cc1cc(F)c2[nH]ccc2c1** | 39 | 8 |
| **O=Cc1ccc2c(c1)CCCC2** | 36 | 1 |
| **O=Cc1cc2ccccc2[nH]1** | 36 | 0 |
| **Cc1occc1C=O** | 35 | 9 |
| **O=Cc1c[nH]c(C(F)(F)F)c1** | 35 | 65 |
| **C[C@]1(C=O)C[C@@H]1c1ccccc1** | 33 | 0 |
| **Nc1ncncc1C=O** | 32 | 1 |
| **COc1cc(C=O)oc(=O)c1** | 32 | 0 |
| **COCc1ccc(C=O)cc1** | 28 | 8 |
| **Nc1ncccc1C=O** | 28 | 0 |
| **O=Cc1cncc2ccccc12** | 27 | 0 |
| **Cc1cc(O)c(C=O)cc1C** | 27 | 3 |
| **O=Cc1ccn(Cc2ccccc2)c1** | 27 | 1 |
| **Cc1nn(C2CCCC2)cc1C=O** | 26 | 0 |
| **CN(C)c1ccncc1C=O** | 26 | 0 |
| **N#Cc1cccc(C=O)c1O** | 25 | 2 |

| Structure | P_adj% | unknown_adj% |
|---|---|---|
| **Cn1cc(CCC=O)cn1** | 25 | 4 |
| **O=Cc1c(O)cccc1[N+](=O)[O-]** | 25 | 0 |
| **O=Cc1cccc2ccc(O)cc12** | 25 | 11 |
| **O=Cc1c[nH]c2ncccc12** | 18 | 1 |
| **O=Cc1cccc2c1NCC2** | 17 | 0 |
| **COc1ccc(F)c(C=O)n1** | 16 | 3 |
| **Nc1ccc(C=O)cn1** | 16 | 39 |
| **Cc1nc(F)ccc1C=O** | 16 | 10 |
| **Cc1ccc2ccccc2c1C=O** | 16 | 0 |
| **Cc1ncc(C=O)c(N(C)C)n1** | 16 | 14 |
| **Cc1ccc(C=O)c(N)n1** | 16 | 3 |
| **O=Cc1cc[nH]c1** | 15 | 0 |
| **O=Cc1nccc2ccccc12** | 15 | 3 |
| **Cc1ncc(C=O)c(N)n1** | 14 | 0 |
| **O=Cc1cnc(O)nc1O** | 14 | 8 |
| **CC(C)(C)C(=O)CC=O** | 14 | 17 |
| **COc1ncncc1C=O** | 13 | 0 |
| **Cn1cc(C=O)c2ccccc21** | 13 | 0 |
| **O=Cc1cnon1** | 12 | 4 |
| **COc1ccnc(C=O)n1** | 12 | 0 |
| **O=Cc1scc2c1OCCO2** | 12 | 0 |
| **Cc1cc(C=O)ncn1** | 11 | 6 |
| **CN(C)c1nccc(C=O)n1** | 11 | 1 |
| **O=CCc1ccc(F)cc1Cl** | 11 | 3 |
| **Cc1cc(C=O)ccc1Cl** | 11 | 0 |
| **O=Cc1c(F)cccc1Cl** | 10 | 0 |
| **Cc1cc(C)c(C)c(C=O)c1C** | 10 | 0 |
| **Cc1cc(C)c(C=O)c(C)c1** | 10 | 0 |
| **Cc1ccc(C=O)nn1** | 8 | 16 |
| **Cc1ccc2[nH]cc(C=O)c2c1** | 8 | 2 |
| **CC(C)(C=O)c1ccccc1** | 6 | 0 |
| **Cc1ccccc1-n1nccc1C=O** | 6 | 0 |
| **Cn1cccc1C=O** | 6 | 1 |
| **CCn1nc(C=O)c2ccccc21** | 5 | 0 |
| **CC(=O)Nc1ccccc1C=O** | 4 | 1 |
| **CCN(CC)c1ncc(C=O)s1** | 4 | 1 |
| **CC(C)(C)c1ncc(C=O)cn1** | 3 | 0 |
| **N#Cc1ccc2[nH]cc(C=O)c2c1** | 3 | 0 |
| **COc1ccc2[nH]cc(C=O)c2c1** | 3 | 0 |
| **N#Cc1ccc(C=O)nc1** | 0 | 4 |
| **CCOCc1ccc(C=O)o1** | 0 | 9 |
| **CCc1ccc(C=O)cn1** | 0 | 17 |
| **O=CCCc1ccccc1CO** | 0 | 25 |
| **CN(C)c1ccc(C=O)o1** | 0 | 48 |
| **COc1ccc(C=O)nc1** | 0 | 100 |
| **N#Cc1ccc(C=O)o1** | 0 | 100 |
| **CC(C)c1nn(C)c(Cl)c1C=O** | 0 | 100 |
| **COc1cccc(C=O)c1Cl** | 0 | 100 |
| **Cc1ccc2occ(C=O)c(=O)c2c1** | 0 | 100 |

| Structure | P_adj% | unknown_adj% |
| --- | --- | --- |
| **Cc1c[nH]nc1C=O** | 0 | 100 |
| **O=Cc1cnc2[nH]ccc2c1** | 0 | 100 |
| **O=CC1CCOCC1** | 0 | 100 |
| **CN(C)C(C)(C)C=O** | 0 | 0 |
| **O=Cc1ccc(C(F)(F)F)[nH]1** | 0 | 0 |
| **Cn1c(C=O)cnc1Cl** | 0 | 0 |
| **Cc1cccc(C=O)c1OC(F)F** | 0 | 0 |
| **O=Cc1ccsc1Cl** | 0 | 0 |
| **CCn1cnc(Cl)c1C=O** | 0 | 0 |
| **COc1nc(Cl)c(C=O)n1C** | 0 | 0 |
| **Cn1ncc(C=O)c1Cl** | 0 | 1 |
| **CC(C)n1cncc1C=O** | 0 | 1 |
| **Cc1cc(C=O)ccc1C#N** | 0 | 1 |
| **O=Cc1cnc(C2CC2)nc1** | 0 | 1 |
| **N[C@@H](C=O)[C@@H](O)[C@H](O)[C@H](O)CO** | 0 | 2 |
| **O=Cc1ccc([N+](=O)[O-])o1** | 0 | 3 |
| **O=Cc1cscc1Cl** | 0 | 5 |
| **C#Cc1ccsc1C=O** | 0 | 10 |
| **Cn1ncc(C=O)c1C1CC1** | 0 | 11 |
| **CC(C)(C=O)N1CCOCC1** | 0 | 0 |
| **C#CCn1cccc1C=O** | 0 | 0 |
| **COc1ccc(C)cc1C=O** | 0 | 0 |
| **CCc1cccc(C=O)c1O** | 0 | 0 |
| **CC(C)(C=O)Oc1ccccc1** | 0 | 0 |
| **O=CCCC1CC1** | 0 | 0 |
| **Cc1nccc(C=O)n1** | 0 | 0 |
| **CCCn1ccc(C=O)c1** | 0 | 0 |
| **CCc1nn(C)c(OC)c1C=O** | 0 | 0 |
| **COC(=O)c1ccsc1C=O** | 0 | 0 |
| **O=Cc1cscc1C(F)(F)F** | 0 | 0 |
| **Cc1cc(Cl)cc(C=O)c1O** | 0 | 0 |
| **O=Cc1cccn1C1CC1** | 0 | 0 |
| **O=Cc1ccc(Cl)nn1** | 0 | 0 |
| **CCc1cccc(F)c1C=O** | 0 | 0 |
| **CCC(C)(C=O)OC** | 0 | 0 |
| **COCCCn1nccc1C=O** | 0 | 0 |
| **O=Cc1ccc(Cl)o1** | 0 | 0 |
| **O=Cc1ccnc(Cl)c1F** | 0 | 0 |
| **Cc1ccc(F)cc1C=O** | 0 | 0 |
| **CC(C)Cn1nccc1C=O** | 0 | 0 |
| **O=Cc1sccc1Cl** | 0 | 0 |
| **CCc1cc(OC)ccc1C=O** | 0 | 0 |
| **O=Cc1ccnn1C1CCCC1** | 0 | 0 |
| **CCOc1cccc(C=O)n1** | 0 | 0 |
| **Cc1ccoc1C=O** | 0 | 0 |
| **N#Cc1ccc(C=O)c(F)c1** | 0 | 0 |
| **N#Cc1c(F)ccc(C=O)c1Cl** | 0 | 0 |
| **CCCn1nc(C)c(Cl)c1C=O** | 0 | 0 |
| **O=Cc1ccccc1N1CCCCC1** | 0 | 0 |

| Structure | P_adj% | unknown_adj% |
|---|---|---|
| **Cc1ccc(C=O)nc1** | 0 | 0 |
| **CC(C)Oc1ccccc1C=O** | 0 | 0 |
| **O=Cc1nccnc1Cl** | 0 | 0 |
| **O=Cc1ccncc1[N+](=O)[O-]** | 0 | 0 |
| **CC(C)(C=O)CN1CCOCC1** | 0 | 0 |
| **COCCOc1ccccc1C=O** | 0 | 1 |
| **COC(=O)C1(C=O)CC1** | 0 | 1 |
| **O=C[C@H](O)[C@H]1OC(=O)[C@@H](O)[C@H]1O** | 0 | 1 |
| **COc1nccnc1C=O** | 0 | 2 |
| **CC(C)(O)C=O** | 0 | 2 |
| **O=CCC1CCCOC1** | 0 | 3 |
| **Cn1nc(C2CC2)cc1C=O** | 0 | 3 |
| **COc1ccc(F)c(C=O)c1** | 0 | 3 |
| **O=Cc1ncccc1C(F)(F)F** | 0 | 3 |
| **COC1(C=O)CCOC1** | 0 | 3 |
| **Cc1ccc(O)c(C=O)n1** | 0 | 4 |
| **CC(C)(C)C(=O)C=O** | 0 | 4 |
| **Cn1cc(C=O)c(Cl)n1** | 0 | 5 |
| **CCn1cc(C=O)c(C)n1** | 0 | 5 |
| **Cc1cc(C=O)cc(Cl)n1** | 0 | 6 |
| **COCC=O** | 0 | 19 |
| **Cc1ncccc1C=O** | 0 | 61 |
| **O=Cc1ccccc1C1CC1** | 0 | 0 |
| **O=Cc1ccc([N+](=O)[O-])s1** | 0 | 0 |
| **Cc1oc2ccccc2c1C=O** | 0 | 0 |
| **Cn1ncc(Cl)c1C=O** | 0 | 0 |
| **CCCc1ccc(C=O)s1** | 0 | 0 |
| **O=Cc1c(O)ccc(Cl)c1F** | 0 | 0 |
| **CC(C)CCn1cccc1C=O** | 0 | 0 |
| **Cc1c(C=O)cnn1C(C)C** | 0 | 0 |
| **CCn1ccnc1C=O** | 0 | 0 |
| **COCC1(C=O)CCC1** | 0 | 0 |
| **COC(C)(C=O)C1CC1** | 0 | 0 |
| **O=CC1(F)CC1** | 0 | 0 |
| **O=Cc1ccc(Cl)s1** | 0 | 0 |
| **O=CC1(C(F)F)CC1** | 0 | 0 |
| **C[C@H](O)[C@@H](O)[C@@H](O)[C@H](O)C=O** | 0 | 0 |
| **C[C@H](O)[C@H](O)[C@@H](O)[C@@H](O)C=O** | 0 | 0 |
| **N#Cc1cccc(C=O)n1** | 0 | 1 |
| **Cc1ccc2oc(C=O)cc2c1** | 0 | 1 |
| **COc1cc(C)c(C=O)c(C)c1C** | 0 | 1 |
| **CCOCCC=O** | 0 | 2 |
| **COc1cc(O)ccc1C=O** | 0 | 2 |
| **C#Cc1ccc(C=O)cc1** | 0 | 3 |
| **COc1nc(C=O)ccc1C** | 0 | 5 |
| **N#Cc1cnc(Cl)c(C=O)c1** | 0 | 5 |
| **C[C@@H](O)[C@@H](O)[C@H](O)[C@H](O)C=O** | 0 | 7 |
| **O=CC1C[C@@H]2CC[C@H]1O2** | 0 | 22 |
| **CCOc1ccc(Cl)cc1C=O** | 0 | 0 |

| Structure | P_adj% | unknown_adj% |
|---|---|---|
| **CN(C)c1ncc(C=O)n1C** | 0 | 0 |
| **O=Cc1ccc(OCC2CC2)cc1** | 0 | 0 |
| **C[C@@H](O)[C@H](O)[C@H](O)[C@@H](O)C=O** | 0 | 0 |
| **O=Cc1c(O)cc(O)cc1O** | 0 | 0 |
| **O=Cc1cnc(Cl)nc1** | 0 | 10 |
| **C#CCn1ccnc1C=O** | 0 | 0 |
| **COc1cc(C)nc(C=O)c1** | 0 | 0 |
| **COc1cc(C)c(C=O)c(O)c1** | 0 | 0 |
| **CN(C)c1ncccc1C=O** | 0 | 0 |
| **Cn1c(Cl)cnc1C=O** | 0 | 0 |
| **Nc1ncc(C=O)c(N)n1** | 0 | 0 |
| **CC(C)(C)C=O** | 0 | 1 |
| **O=CC1CCCOC1** | 0 | 2 |
| **CC1(C)[C@H]2C[C@@](C)(C=O)C[C@H]21** | 0 | 0 |
| **Cc1cc(C=O)cs1** | 0 | 2 |
| **Cc1c(O)cc(O)c(C=O)c1O** | 0 | 9 |
| **Cn1cc(C=O)c(C(C)(C)C)n1** | 0 | 3 |
| **Cc1ccncc1C=O** | 0 | 5 |
| **CC(C)(C=O)S(C)(=O)=O** | 0 | 3 |
| **O=Cc1cc(O)c(O)cc1O** | 0 | 7 |
| **O=CC1(c2ccccc2)CCOC1** | 0 | 9 |
| **CC(C)C(C#N)C=O** | 0 | 6 |
| **O=Cc1ccc(O)c(O)c1O** | 0 | 0 |
| **CCc1ccc(C=O)o1** | 0 | 7 |
| **CC1=C(C=O)C(C)(C)CC=C1** | 0 | 0 |
| **Cc1nc(C)c(C=O)c(N)n1** | 0 | 4 |
| **CC(C)c1ccc(CC=O)cc1** | 0 | 30 |
| **Cc1ccccc1CC=O** | 0 | 39 |
| **Cc1cc(O)cc(O)c1C=O** | 0 | 9 |
| **O=CC1=COCCC1** | 0 | 11 |
| **O=Cc1ccc(O)c2ncccc12** | 0 | 1 |
| **Cc1[nH]c2ccc(F)cc2c1C=O** | 0 | 1 |
| **Cc1c(C=O)c2ccccn2c1C#N** | 0 | 0 |
| **O=Cn1nnc2ccccc21** | 0 | 0 |
| **CCc1cccc2c(C=O)c[nH]c12** | 0 | 2 |
| **O=Cc1ccc(O)c2ccccc12** | 0 | 4 |
| **CN(C)c1ncnc(Cl)c1C=O** | 0 | 0 |

#### 8.3. Suzuki coupling building block validation data

In the same manner as outlined in **Supplementary Section 8.2,** the abridged Suzuki coupling validation data shown below only reports the key calculated factors of adjusted %AUC product (or derivatives) and adjusted %AUC unknown DNA species, in both cases correcting for any residual small molecule building block peaks integrated in the LCMS data.

Scores were assigned from 1 to 4, indicating a preference for use of a given building block across ArI and ArBr validation campaigns according to the below hierarchical rules:

((ArI_adj%P >= 70) AND (ArBr_adj%P >= 70)) AND ((ArI_unknown_adj% <= 10) AND (ArBr_unknown_adj% <= 10)) => 4

(((ArI_adj%P >= 70) AND (ArBr_adj%P >= 50)) XOR ((ArI_adj%P >= 50) AND (ArBr_adj%P >= 70))) AND ((ArI_unknown_adj% <= 10) AND (ArBr_unknown_adj% <= 10)) => 3

((ArI_adj%P >= 70) AND (ArBr_adj%P >= 70)) AND ((ArI_unknown_adj% <= 10) XOR (ArBr_unknown_adj% <= 10)) => 2

NOT MISSING Structure => 1

**Supplementary Table 13 | Suzuki coupling building block validation data**

| Structure | ArI_adj %P | ArI_unknown_ adj% | ArBr_adj %P | ArBr_unknown _adj% | Score |
|---|---|---|---|---|---|
| **O=C(O)COc1cc(F)cc(B(O)O)c1** | 100 | 0 | 96 | 0 | 4 |
| **CN(C)CCc1ccc(B(O)O)cc1** | 100 | 0 | 94 | 0 | 4 |
| **Cn1c(=O)[nH]c2ccc(B3OC(C)(C)C(C)(C)O3)cc21** | 100 | 0 | 92 | 0 | 4 |
| **OB(O)c1ccc2c(c1)OCO2** | 100 | 0 | 92 | 0 | 4 |
| **CC1(C)OB(c2ccc3c(c2)COC(=O)N3)OC1(C)C** | 100 | 0 | 87 | 0 | 4 |
| **Cn1nccc1B(O)OC(C)(C)C(C)(C)O** | 100 | 0 | 84 | 0 | 4 |
| **COC(=O)c1ccc(B(O)O)c(F)c1** | 100 | 0 | 81 | 0 | 4 |
| **CC(C)(C)C1CC=C(B(O)O)CC1** | 100 | 0 | 76 | 0 | 4 |
| **COc1c(C)cc(B(O)O)cc1C** | 99 | 0 | 99 | 0 | 4 |
| **CCOC(=O)c1cccc(B(O)O)c1** | 99 | 0 | 94 | 1 | 4 |
| **OB(O)c1cccc(-c2ccccc2)c1** | 99 | 0 | 93 | 0 | 4 |
| **Nc1cccc(B(O)O)c1** | 99 | 0 | 91 | 1 | 4 |
| **COC(=O)CCc1cccc(B(O)O)c1** | 99 | 0 | 90 | 0 | 4 |
| **CCCOc1ccc(B(O)O)cc1Cl** | 99 | 0 | 85 | 0 | 4 |
| **CC(C)c1cccc(B(O)OCCO)c1** | 99 | 0 | 84 | 0 | 4 |
| **OB(O)c1ccc2ccccc2c1** | 99 | 0 | 81 | 0 | 4 |
| **CC1(C)OB(c2cc3ccccc3o2)OC1(C)C** | 99 | 0 | 75 | 0 | 4 |
| **CC(=O)Nc1cccc(B(O)O)c1** | 98 | 0 | 100 | 0 | 4 |
| **COc1ccc2cc(B(O)O)ccc2c1** | 98 | 0 | 97 | 0 | 4 |
| **OCc1cccc(B(O)O)c1** | 98 | 0 | 97 | 0 | 4 |
| **Cc1cc(B(O)O)ccc1Cl** | 98 | 0 | 97 | 1 | 4 |
| **CN(C)Cc1cc(B(O)O)ccc1F** | 98 | 0 | 95 | 0 | 4 |
| **COc1cc(B(O)O)ccc1C** | 98 | 0 | 95 | 0 | 4 |
| **CNC(=O)c1ccc(B(O)O)cc1** | 98 | 0 | 94 | 0 | 4 |
| **O=C(O)c1ccc(B(O)O)cc1Cl** | 98 | 0 | 94 | 0 | 4 |
| **CC1(C)OB(c2cnc(N)nc2)OC1(C)C** | 98 | 0 | 93 | 0 | 4 |
| **Cc1ccc(B(O)O)cc1Cl** | 98 | 0 | 91 | 0 | 4 |
| **OB(O)c1cccc(F)c1** | 98 | 0 | 91 | 0 | 4 |
| **Cc1ccc(B(O)O)cc1** | 98 | 0 | 90 | 0 | 4 |
| **COc1ccc(B(O)O)cc1Cl** | 98 | 0 | 90 | 8 | 4 |
| **CC(C)Cc1ccc(B(O)O)cc1** | 98 | 0 | 88 | 0 | 4 |
| **CCc1cccc(B(O)O)c1** | 98 | 0 | 88 | 0 | 4 |
| **Cc1cc(B(O)O)ccc1F** | 98 | 0 | 88 | 0 | 4 |
| **Cc1ccc2cc(B(O)O)ccc2n1** | 98 | 0 | 88 | 0 | 4 |
| **CC(C)(C)OCc1ccc(B(O)O)cc1** | 98 | 0 | 87 | 0 | 4 |
| **OCc1ccc(B(O)O)cc1** | 98 | 0 | 87 | 1 | 4 |

| Structure | ArI_adj %P | ArI_unknown_adj% | ArBr_adj %P | ArBr_unknown_adj% | Score |
|---|---|---|---|---|---|
| **[2H]c1c([2H])c([2H])c(B(O)O)c([2H])c1[2H]** | 98 | 0 | 87 | 1 | 4 |
| **CC(C)Oc1ccc(B(O)O)cc1F** | 98 | 0 | 86 | 0 | 4 |
| **O=C(O)c1ccc(B(O)O)c(Cl)c1** | 98 | 0 | 85 | 0 | 4 |
| **CCOc1ccc(B(O)O)cc1C** | 98 | 0 | 83 | 0 | 4 |
| **CC(C)(C)c1cccc(B(O)O)c1** | 98 | 0 | 82 | 1 | 4 |
| **COC(=O)c1cc(B2OC(C)(C)C(C)(C)O2)ccc1C** | 98 | 0 | 81 | 0 | 4 |
| **OB(O)c1ccc(-c2ccccc2)cc1** | 98 | 0 | 79 | 3 | 4 |
| **OB(O)c1cccc2ccccc12** | 98 | 0 | 78 | 0 | 4 |
| **CCCCc1ccc(B(O)O)cc1** | 98 | 0 | 71 | 0 | 4 |
| **O=C(O)Cc1cccc(B(O)O)c1** | 97 | 0 | 98 | 0 | 4 |
| **OB(O)c1ccc(Cl)c(Cl)c1** | 97 | 0 | 98 | 0 | 4 |
| **Cc1cccc(B(O)O)c1** | 97 | 0 | 96 | 0 | 4 |
| **O=C(O)c1cc(B(O)O)ccc1Cl** | 97 | 0 | 96 | 0 | 4 |
| **CNC(=O)c1ccc(B2OC(C)(C)C(C)(C)O2)cc1C** | 97 | 1 | 96 | 1 | 4 |
| **OB(O)c1ccc2c(c1)CCC2** | 97 | 0 | 94 | 0 | 4 |
| **OB(O)c1cccc(Oc2ccccc2)c1** | 97 | 0 | 94 | 0 | 4 |
| **CCn1ccc(B2OC(C)(C)C(C)(C)O2)n1** | 97 | 0 | 92 | 0 | 4 |
| **COc1cc(B(O)O)ccc1Cl** | 97 | 0 | 91 | 3 | 4 |
| **COc1ccc(B(O)O)cc1F** | 97 | 0 | 90 | 0 | 4 |
| **COc1cc(B(O)O)cc(C(=O)O)c1** | 97 | 0 | 90 | 7 | 4 |
| **Cc1cc(C)cc(B(O)O)c1** | 97 | 0 | 89 | 0 | 4 |
| **OB(O)c1ccc(F)c(C(F)(F)F)c1** | 97 | 0 | 89 | 0 | 4 |
| **O=C(O)c1cccc(B(O)O)c1** | 97 | 0 | 89 | 3 | 4 |
| **CC1(C)OB(c2ccc(C3(CO)CC3)cc2)OC1(C)C** | 97 | 1 | 88 | 0 | 4 |
| **CCCc1ccc(B(O)O)cc1** | 97 | 0 | 86 | 0 | 4 |
| **CCc1ccc(B(O)O)cc1** | 97 | 0 | 86 | 0 | 4 |
| **CC1(C)OB(c2cccc(Cn3cccn3)c2)OC1(C)C** | 97 | 0 | 84 | 1 | 4 |
| **CC(=O)c1cccc(B(O)O)c1** | 97 | 0 | 79 | 0 | 4 |
| **COC(=O)c1cc(F)cc(B(O)O)c1** | 97 | 0 | 75 | 0 | 4 |
| **COc1ccc(C(C)C)cc1B(O)O** | 97 | 0 | 72 | 0 | 4 |
| **O=C(O)c1cc(Cl)cc(B(O)O)c1** | 96 | 0 | 100 | 0 | 4 |
| **CC1(C)OB(c2cnn(-c3ccncc3)c2)OC1(C)C** | 96 | 0 | 99 | 0 | 4 |
| **OB(O)c1cc(Cl)cc(Cl)c1** | 96 | 0 | 98 | 0 | 4 |
| **COc1cc(F)cc(B(O)O)c1** | 96 | 0 | 95 | 0 | 4 |
| **COc1cc(OC)cc(B(O)O)c1** | 96 | 0 | 95 | 0 | 4 |
| **CCNC(=O)c1cccc(B(O)O)c1** | 96 | 0 | 95 | 3 | 4 |
| **OB(O)c1cc(F)cc(Cl)c1** | 96 | 0 | 93 | 0 | 4 |
| **COc1ccc(B(O)O)cc1C** | 96 | 0 | 93 | 1 | 4 |
| **O=C(O)c1ccc(B(O)O)cc1F** | 96 | 0 | 93 | 1 | 4 |
| **CC1(C)OB(c2ccc3c(c2)CCC(=O)N3)OC1(C)C** | 96 | 1 | 93 | 0 | 4 |
| **CCOc1cccc(B(O)O)c1** | 96 | 0 | 92 | 2 | 4 |

| Structure | ArI_adj %P | ArI_unknown_adj% | ArBr_adj %P | ArBr_unknown_adj% | Score |
|---|---|---|---|---|---|
| **CC1(C)OB(c2cccc(S(N)(=O)=O)c2)OC1(C)C** | 96 | 4 | 90 | 10 | 4 |
| **Cc1cccc(B(O)O)c1C** | 96 | 0 | 88 | 0 | 4 |
| **OB(O)c1ccc(Oc2ccccc2)cc1** | 96 | 0 | 85 | 0 | 4 |
| **COc1ccc(F)c(B(O)O)c1** | 96 | 0 | 83 | 0 | 4 |
| **OB(O)c1cccc(OC(F)(F)F)c1** | 96 | 0 | 82 | 0 | 4 |
| **CC1(C)OB(C2=CCNCC2)OC1(C)C** | 96 | 3 | 76 | 6 | 4 |
| **CC1(C)OB(c2cnc3occc3c2)OC1(C)C** | 96 | 0 | 75 | 0 | 4 |
| **OB(O)c1cccc(Cl)c1** | 95 | 0 | 98 | 0 | 4 |
| **OCc1ccccc1B(O)O** | 95 | 0 | 98 | 0 | 4 |
| **OCCc1cccc(B(O)O)c1** | 95 | 0 | 94 | 1 | 4 |
| **OB(O)c1ccc(F)c(Cl)c1** | 95 | 2 | 92 | 5 | 4 |
| **COc1ncccc1B(O)O** | 95 | 0 | 91 | 0 | 4 |
| **O=C(O)CCc1cccc(B(O)O)c1** | 95 | 0 | 89 | 1 | 4 |
| **CC1(C)OB(c2ccc3c(c2)CCOC3)OC1(C)C** | 95 | 1 | 87 | 0 | 4 |
| **Cc1cc(F)ccc1B(O)O** | 95 | 0 | 86 | 0 | 4 |
| **NC(=O)c1ccc(B(O)O)cc1** | 95 | 5 | 84 | 2 | 4 |
| **COc1ccc(Cl)cc1B(O)O** | 95 | 0 | 76 | 0 | 4 |
| **O=C(O)c1cccc(B(O)O)c1F** | 95 | 5 | 71 | 0 | 4 |
| **Nc1cc(B(O)O)ccc1F** | 94 | 0 | 100 | 0 | 4 |
| **CCNC(=O)c1ccc(B(O)O)cc1** | 94 | 0 | 97 | 1 | 4 |
| **COc1ccc(C#N)cc1B(O)O** | 94 | 0 | 96 | 1 | 4 |
| **O=C(NC1CC1)c1cccc(B(O)O)c1** | 94 | 0 | 95 | 0 | 4 |
| **OB(O)c1ccc2c(c1)CCCO2** | 94 | 0 | 94 | 0 | 4 |
| **COc1cc(C(=O)O)ccc1B(O)O** | 94 | 0 | 92 | 3 | 4 |
| **CC(=O)c1ccc(B(O)O)cc1** | 94 | 0 | 92 | 4 | 4 |
| **OB(O)c1ccc(Cl)cc1** | 94 | 0 | 90 | 1 | 4 |
| **Cc1ccc(B(O)O)c(F)c1** | 94 | 0 | 89 | 0 | 4 |
| **OCc1ccc(Cl)c(B(O)O)c1** | 94 | 0 | 89 | 0 | 4 |
| **OB(O)c1ccc(Cl)c(F)c1** | 94 | 0 | 87 | 6 | 4 |
| **CCOc1ncccc1B(O)O** | 94 | 0 | 85 | 3 | 4 |
| **CC1(C)OB(c2cccc(O)c2F)OC1(C)C** | 94 | 0 | 81 | 0 | 4 |
| **Cc1ccc(F)cc1B(O)O** | 94 | 0 | 79 | 0 | 4 |
| **CC1(C)OB(c2cccc3c2CCO3)OC1(C)C** | 94 | 1 | 78 | 0 | 4 |
| **OB(O)c1ccc(OC(F)(F)F)cc1** | 94 | 1 | 78 | 0 | 4 |
| **COCCc1cccc(B2OC(C)(C)C(C)(C)O2)c1** | 94 | 0 | 75 | 0 | 4 |
| **Nc1cc(B(O)O)ccc1Cl** | 93 | 0 | 95 | 0 | 4 |
| **CC1(C)OB(c2ccc3ccc(=O)[nH]c3c2)OC1(C)C** | 93 | 3 | 95 | 0 | 4 |
| **CC1(C)OB(c2ccc(C3CCC(=O)N3)cc2)OC1(C)C** | 93 | 0 | 93 | 0 | 4 |
| **OB(O)c1cccc2c1OCCO2** | 93 | 0 | 92 | 0 | 4 |
| **CNC(=O)Nc1ccc(B2OC(C)(C)C(C)(C)O2)cc1** | 93 | 0 | 89 | 0 | 4 |

| Structure | ArI_adj %P | ArI_unknown_adj% | ArBr_adj %P | ArBr_unknown_adj% | Score |
|---|---|---|---|---|---|
| **CC(C)(O)c1ccc(B2OC(C)(C)C(C)(C)O2)c(F)c1** | 93 | 0 | 88 | 0 | 4 |
| **OB(O)c1cc(F)c(F)c(F)c1** | 93 | 0 | 88 | 2 | 4 |
| **CC(C)NC(=O)c1cccc(B(O)O)c1** | 93 | 0 | 83 | 2 | 4 |
| **CC(C)OCc1cccc(B(O)O)c1** | 93 | 1 | 83 | 1 | 4 |
| **COc1ncc(B2OC(C)(C)C(C)(C)O2)cc1C** | 93 | 0 | 80 | 0 | 4 |
| **Cc1c(Cl)cccc1B(O)O** | 93 | 0 | 80 | 0 | 4 |
| **COc1ncc(Cl)cc1B(O)O** | 93 | 0 | 71 | 0 | 4 |
| **CC1(C)OB(c2cnc3ccnn3c2)OC1(C)C** | 93 | 7 | 71 | 0 | 4 |
| **COC(=O)c1cc(O)cc(B2OC(C)(C)C(C)(C)O2)c1** | 92 | 6 | 94 | 0 | 4 |
| **OB(O)c1cccc2c1OCCCO2** | 92 | 0 | 87 | 0 | 4 |
| **COc1ccc(C)cc1B(O)O** | 92 | 0 | 87 | 4 | 4 |
| **CCCNC(=O)c1cccc(B(O)O)c1** | 92 | 0 | 85 | 0 | 4 |
| **CCOc1cc(F)ccc1B(O)O** | 92 | 0 | 83 | 0 | 4 |
| **CC1(C)OB(c2cccc(CN)c2)OC1(C)C** | 92 | 1 | 83 | 1 | 4 |
| **Cc1cc2cc(B3OC(C)(C)C(C)(C)O3)ccc2[nH]1** | 92 | 3 | 80 | 0 | 4 |
| **COC(=O)c1ccc(B2OC(C)(C)C(C)(C)O2)o1** | 92 | 4 | 79 | 0 | 4 |
| **CC1(C)OB(c2cccc3ccoc23)OC1(C)C** | 92 | 6 | 79 | 9 | 4 |
| **CC(C)Oc1ccc(F)cc1B(O)O** | 92 | 0 | 77 | 0 | 4 |
| **COc1ncc(B(O)O)cc1Cl** | 91 | 0 | 99 | 0 | 4 |
| **Cn1nnc2ccc(B3OC(C)(C)C(C)(C)O3)cc21** | 91 | 5 | 96 | 1 | 4 |
| **Cc1ccc(Cl)cc1B(O)O** | 91 | 0 | 91 | 4 | 4 |
| **CC1(C)OB(c2ccc3c(c2)CCCNC3=O)OC1(C)C** | 91 | 6 | 91 | 0 | 4 |
| **COC(=O)Nc1ccc(B(O)O)cc1** | 91 | 0 | 89 | 0 | 4 |
| **CCOC(=O)c1ccc(B(O)O)cc1F** | 91 | 0 | 88 | 0 | 4 |
| **CCN(CC)c1ncc(B(O)O)cn1** | 91 | 2 | 87 | 0 | 4 |
| **CCCOc1cccc(B(O)O)c1** | 91 | 5 | 86 | 2 | 4 |
| **OB(O)c1cccc(Cl)c1F** | 91 | 0 | 81 | 0 | 4 |
| **OB(O)c1cc(F)cc(C(F)(F)F)c1** | 90 | 0 | 97 | 0 | 4 |
| **COc1ccc(B(O)O)cc1C#N** | 90 | 6 | 92 | 0 | 4 |
| **OB(O)c1ccc(C(F)(F)F)c(F)c1** | 90 | 0 | 91 | 0 | 4 |
| **Cc1nc2ccc(B3OC(C)(C)C(C)(C)O3)cc2o1** | 90 | 7 | 91 | 3 | 4 |
| **OCc1ccc(B(O)O)cc1Cl** | 90 | 8 | 84 | 0 | 4 |
| **CC1(C)OB(c2ccc(C3CC(=O)C3)cc2)OC1(C)C** | 90 | 10 | 79 | 3 | 4 |
| **CS(=O)(=O)c1ccc(B(O)O)cc1** | 89 | 0 | 100 | 0 | 4 |
| **Cc1ccc(C(N)=O)cc1B1OC(C)(C)C(C)(C)O1** | 89 | 0 | 100 | 0 | 4 |
| **CC1(C)OB(c2cccc3c2CNC3=O)OC1(C)C** | 89 | 3 | 100 | 0 | 4 |
| **OB(O)c1cc(F)cc(F)c1** | 89 | 0 | 98 | 0 | 4 |

| Structure | ArI_adj %P | ArI_unknown_adj% | ArBr_adj %P | ArBr_unknown_adj% | Score |
|---|---|---|---|---|---|
| **O=C(O)CCc1ccc(B(O)O)cc1** | 89 | 0 | 91 | 0 | 4 |
| **COC(=O)c1cc(B(O)O)ccc1Cl** | 89 | 7 | 91 | 0 | 4 |
| **CC(C)(C)c1ccc(B(O)O)cc1** | 89 | 2 | 86 | 0 | 4 |
| **COc1cccc(B(O)O)c1OC** | 89 | 1 | 82 | 0 | 4 |
| **COc1cccc2cc(B3OC(C)(C)C(C)(C)O3)oc12** | 89 | 8 | 71 | 0 | 4 |
| **COc1ccc(B(O)O)cc1OC(C)C** | 88 | 0 | 94 | 0 | 4 |
| **OB(O)c1ccc(F)nc1** | 88 | 3 | 94 | 0 | 4 |
| **OB(O)c1ccc(C(F)(F)F)cc1** | 88 | 0 | 86 | 0 | 4 |
| **CCOc1ccc(B(O)O)cc1** | 87 | 0 | 88 | 0 | 4 |
| **N#Cc1cc(B(O)O)ccc1Cl** | 87 | 5 | 85 | 0 | 4 |
| **Cc1ccc(C(=O)O)cc1B1OC(C)(C)C(C)(C)O1** | 87 | 0 | 83 | 0 | 4 |
| **Cc1cc(Cl)ccc1B(O)O** | 87 | 0 | 80 | 0 | 4 |
| **OB(O)c1cccc2c1OCC2** | 87 | 0 | 79 | 0 | 4 |
| **CN(C)c1ncc(B2OC(C)(C)C(C)(C)O2)cc1Cl** | 87 | 7 | 77 | 3 | 4 |
| **COc1cc(CO)ccc1B1OC(C)(C)C(C)(C)O1** | 86 | 0 | 88 | 0 | 4 |
| **CCOc1ccccc1B(O)O** | 86 | 0 | 78 | 0 | 4 |
| **N#Cc1ccc(B(O)O)cc1F** | 84 | 0 | 85 | 0 | 4 |
| **COc1ccc(B2OC(C)(C)C(C)(C)O2)cn1** | 83 | 0 | 71 | 0 | 4 |
| **CC1Cc2cc(B(O)O)ccc2O1** | 81 | 0 | 96 | 0 | 4 |
| **OB(O)c1cncnc1** | 81 | 4 | 91 | 0 | 4 |
| **CC(C)(C)n1cc(B2OC(C)(C)C(C)(C)O2)cn1** | 81 | 0 | 71 | 0 | 4 |
| **COC(=O)c1ccc(B2OC(C)(C)C(C)(C)O2)cc1F** | 80 | 9 | 84 | 0 | 4 |
| **CCN(CC)Cc1ccc(B(O)O)cc1** | 79 | 0 | 100 | 0 | 4 |
| **OB(O)c1ccc2ccc(O)cc2c1** | 78 | 0 | 91 | 0 | 4 |
| **OB(O)c1ccc2cc[nH]c2c1** | 77 | 0 | 95 | 3 | 4 |
| **COC(=O)c1cc(B2OC(C)(C)C(C)(C)O2)ccc1O** | 72 | 0 | 73 | 4 | 4 |
| **COC(=O)c1cccc(B(O)O)c1F** | 99 | 0 | 69 | 0 | 3 |
| **OB(O)c1ccc2c(c1)OCCCO2** | 99 | 0 | 58 | 0 | 3 |
| **Cc1cc(B(O)O)cc(C(C)(C)C)c1** | 98 | 0 | 57 | 0 | 3 |
| **OB(O)c1ccsc1** | 98 | 1 | 55 | 0 | 3 |
| **OB(O)c1cccc(N2CCCCC2)c1** | 98 | 0 | 52 | 0 | 3 |
| **Cn1ncc2ncc(B(O)O)cc21** | 97 | 0 | 51 | 0 | 3 |
| **CC1(C)OB(c2ccc(S(C)(=O)=O)cc2)OC1(C)C** | 96 | 0 | 69 | 0 | 3 |
| **COC(=O)c1ccc(C)c(B2OC(C)(C)C(C)(C)O2)c1** | 96 | 0 | 54 | 0 | 3 |
| **CNc1cccc(B2OC(C)(C)C(C)(C)O2)c1** | 96 | 0 | 53 | 0 | 3 |
| **CN(C)Cc1ccc(F)c(B2OC(C)(C)C(C)(C)O2)c1** | 96 | 0 | 52 | 0 | 3 |
| **COc1ncc(B2OC(C)(C)C(C)(C)O2)cn1** | 95 | 2 | 66 | 0 | 3 |

| Structure | ArI_adj %P | ArI_unknown_adj% | ArBr_adj %P | ArBr_unknown_adj% | Score |
|---|---|---|---|---|---|
| **Cc1ccc(B(O)O)cc1[N+](=O)[O-]** | 95 | 0 | 62 | 0 | 3 |
| **CC1(c2ccc(B3OC(C)(C)C(C)(C)O3)cc2)CC1** | 95 | 2 | 58 | 1 | 3 |
| **OB(O)C=CC1CCCCC1** | 94 | 0 | 62 | 0 | 3 |
| **COc1cc(B(O)O)cc(OC)c1OC** | 94 | 0 | 60 | 0 | 3 |
| **CCc1ccccc1B(O)O** | 93 | 0 | 67 | 0 | 3 |
| **Cc1ccc(OC(C)C)c(B(O)O)c1** | 93 | 1 | 64 | 2 | 3 |
| **COc1cc(F)c(F)cc1B(O)O** | 93 | 0 | 61 | 0 | 3 |
| **CCOCc1ccc(B2OC(C)(C)C(C)(C)O2)cc1F** | 93 | 0 | 59 | 0 | 3 |
| **CC(C)Oc1cc(F)ccc1B(O)O** | 93 | 0 | 57 | 0 | 3 |
| **Cc1cc(B(O)O)ccc1OC(C)C** | 92 | 0 | 62 | 0 | 3 |
| **COC(=O)c1cc(Cl)cc(B(O)O)c1** | 92 | 0 | 60 | 0 | 3 |
| **Cc1cccc(B(O)O)c1Cl** | 92 | 0 | 54 | 0 | 3 |
| **COc1c(Cl)cccc1B(O)O** | 90 | 0 | 59 | 0 | 3 |
| **COc1c(F)cc(F)cc1B(O)O** | 90 | 0 | 56 | 0 | 3 |
| **Cc1ccc(B(O)O)c(Cl)c1** | 87 | 0 | 67 | 0 | 3 |
| **CC1(C)Cc2cc(B3OC(C)(C)C(C)(C)O3)ccc2O1** | 87 | 0 | 62 | 0 | 3 |
| **CC(C)COc1ccc(B(O)O)cc1** | 87 | 0 | 58 | 3 | 3 |
| **COc1cc(C)c(B(O)O)cc1C** | 85 | 0 | 53 | 2 | 3 |
| **COc1ccc(B(O)O)c(C)c1** | 81 | 0 | 50 | 0 | 3 |
| **CN(C)c1cc(B2OC(C)(C)C(C)(C)O2)ccn1** | 80 | 0 | 68 | 0 | 3 |
| **Cc1cc(B(O)O)c(C)cc1F** | 80 | 0 | 68 | 0 | 3 |
| **CC(C)c1nn(C)cc1B1OC(C)(C)C(C)(C)O1** | 76 | 8 | 68 | 1 | 3 |
| **CC1(C)OB(c2cccc(O)c2)OC1(C)C** | 69 | 0 | 84 | 0 | 3 |
| **COC(=O)Cc1ccc(B(O)O)cc1** | 64 | 2 | 93 | 0 | 3 |
| **CC1(C)OB(c2ccc3occc3c2)OC1(C)C** | 50 | 5 | 73 | 3 | 3 |
| **NC(=O)c1ccc(B(O)O)cc1Cl** | 96 | 0 | 74 | 22 | 2 |
| **COc1ccc(B(O)O)cc1C(=O)O** | 89 | 11 | 88 | 0 | 2 |
| **CONC(=O)c1ccc(B(O)O)cc1** | 88 | 12 | 96 | 0 | 2 |
| **OB(O)c1ccc2c(c1)OC(F)(F)O2** | 85 | 13 | 95 | 0 | 2 |
| **CC(O)c1ccc(B(O)O)cc1** | 85 | 13 | 89 | 5 | 2 |
| **COC(=O)c1cccc(B(O)O)c1** | 83 | 14 | 76 | 0 | 2 |
| **O=C(O)c1ccc(B(O)O)c(F)c1** | 82 | 14 | 84 | 0 | 2 |
| **CC1(C)OB(c2ccc(C3(N)CCCC3)cc2)OC1(C)C** | 82 | 13 | 80 | 4 | 2 |
| **CC(=O)Nc1ccc(B2OC(C)(C)C(C)(C)O2)cn1** | 79 | 21 | 89 | 5 | 2 |
| **N#Cc1cc(Cl)cc(B(O)O)c1** | 74 | 24 | 96 | 0 | 2 |
| **OB(O)c1ccc(N2CCCC2)nc1** | 74 | 25 | 87 | 0 | 2 |
| **CC1(C)OB(c2ccc(F)c(C#N)c2)OC1(C)C** | 73 | 22 | 79 | 0 | 2 |
| **OB(O)/C=C/c1ccc(Cl)cc1** | 100 | 0 | 51 | 20 | 1 |
| **OB(O)c1ccc2nncn2c1** | 100 | 0 | 40 | 0 | 1 |
| **CCOC(=O)c1cccc(B(O)O)c1F** | 100 | 0 | 33 | 6 | 1 |
| **Nc1ccc(B(O)O)cn1** | 100 | 0 | 27 | 44 | 1 |

| Structure | ArI_adj %P | ArI_unknown_adj% | ArBr_adj %P | ArBr_unknown_adj% | Score |
|---|---|---|---|---|---|
| **CC1(C)OB(c2cnn(-c3ccccn3)c2)OC1(C)C** | 100 | 0 | 26 | 0 | 1 |
| **CC1(C)OB(c2ccc(N)cc2F)OC1(C)C** | 100 | 0 | 17 | 0 | 1 |
| **COC(=O)/C=C/c1cccc(B(O)O)c1** | 100 | 0 | 13 | 0 | 1 |
| **CC1(C)OB(c2cnn3cccnc23)OC1(C)C** | 100 | 0 | 8 | 1 | 1 |
| **Nc1ccccc1B(O)O** | 100 | 0 | 0 | 0 | 1 |
| **Cc1cc(OC(C)C)ccc1B(O)O** | 100 | 0 | 0 | 3 | 1 |
| **COc1cccc(C#N)c1B1OC(C)(C)C(C)(C)O1** | 100 | 0 | 0 | 6 | 1 |
| **COc1ccc(/C=C/B(O)O)cc1** | 100 | 0 | 0 | 100 | 1 |
| **Cc1cc(B(O)O)cs1** | 99 | 0 | 48 | 0 | 1 |
| **CSc1ccc(B(O)O)cc1** | 99 | 0 | 29 | 0 | 1 |
| **CC1(C)OB(c2cccc3cc[nH]c23)OC1(C)C** | 99 | 0 | 7 | 0 | 1 |
| **OB(O)/C=C/c1cc(F)cc(F)c1** | 98 | 0 | 41 | 15 | 1 |
| **CC1(C)OB(c2cnc3[nH]ccc3c2)OC1(C)C** | 98 | 0 | 37 | 0 | 1 |
| **CSc1ccccc1B(O)O** | 98 | 0 | 33 | 0 | 1 |
| **Cc1cc(B(O)O)cnc1O** | 98 | 2 | 9 | 0 | 1 |
| **OB(O)c1cncc(F)c1** | 98 | 0 | 0 | 0 | 1 |
| **COC/C=C/B(O)OC(C)(C)C(C)(C)O** | 98 | 0 | 0 | 100 | 1 |
| **CC1(C)OB(c2cc[nH]c2)OC1(C)C** | 97 | 3 | 41 | 0 | 1 |
| **CC1(C)OB(c2ccc(C3CC3)c(C#N)c2)OC1(C)C** | 97 | 0 | 23 | 2 | 1 |
| **OB(O)c1cnc2ccccc2c1** | 97 | 0 | 12 | 0 | 1 |
| **CC1(C)OB(c2cccc(Cn3ccnc3)c2)OC1(C)C** | 97 | 0 | 0 | 0 | 1 |
| **Cc1n[nH]c2ccc(B3OC(C)(C)C(C)(C)O3)cc12** | 97 | 0 | 0 | 0 | 1 |
| **OB(O)c1cccc2cccnc12** | 97 | 0 | 0 | 2 | 1 |
| **Cc1ccc(/C=C/B(O)O)cc1** | 97 | 3 | 0 | 100 | 1 |
| **OB(O)/C=C/c1cccc(F)c1** | 96 | 0 | 40 | 27 | 1 |
| **Cc1ccc(Cl)c(B(O)O)c1** | 96 | 0 | 20 | 5 | 1 |
| **CCOc1ccc(B(O)O)cn1** | 96 | 0 | 8 | 0 | 1 |
| **CC1(C)OB(c2cncc3sccc23)OC1(C)C** | 96 | 0 | 0 | 1 | 1 |
| **O=[N+]([O-])c1ccc(B(O)O)cn1** | 96 | 0 | 0 | 3 | 1 |
| **O=C(O)/C=C/c1ccc(B(O)O)cc1** | 96 | 0 | 0 | 100 | 1 |
| **N#Cc1cc(B(O)O)cs1** | 96 | 2 | 0 | 0 | 1 |
| **C/C=C/B(O)O** | 95 | 0 | 49 | 2 | 1 |
| **Cc1nc(N)ccc1B1OC(C)(C)C(C)(C)O1** | 95 | 0 | 35 | 0 | 1 |
| **CC(C)Oc1cncc(B2OC(C)(C)C(C)(C)O2)c1** | 95 | 0 | 6 | 1 | 1 |
| **CC1(C)OB(/C=C/c2ccccn2)OC1(C)C** | 95 | 0 | 0 | 0 | 1 |
| **Cc1cncc(B2OC(C)(C)C(C)(C)O2)c1** | 95 | 0 | 0 | 0 | 1 |
| **OB(O)c1ccc2nonc2c1** | 95 | 0 | 0 | 1 | 1 |
| **CC(=O)n1ncc2ccc(B3OC(C)(C)C(C)(C)O3)cc21** | 95 | 0 | 0 | 3 | 1 |

| Structure | ArI_adj %P | ArI_unknown_adj% | ArBr_adj %P | ArBr_unknown_adj% | Score |
|---|---|---|---|---|---|
| **COc1ccc(C=CB2OC(C)(C)C(C)(C)O2)cc1** | 95 | 4 | 0 | 100 | 1 |
| **CC(C)Oc1ccc(Cl)cc1B(O)O** | 94 | 0 | 44 | 2 | 1 |
| **COCc1cc(B2OC(C)(C)C(C)(C)O2)cs1** | 94 | 4 | 40 | 0 | 1 |
| **CC(C)COc1cccc(B(O)O)c1F** | 94 | 0 | 36 | 0 | 1 |
| **CC(C)c1ccccc1B(O)O** | 94 | 0 | 21 | 19 | 1 |
| **CC1(C)OB(c2ccc(C(N)=O)s2)OC1(C)C** | 94 | 1 | 20 | 0 | 1 |
| **OB(O)c1ccc(Cl)cc1Cl** | 94 | 0 | 14 | 3 | 1 |
| **Cc1cccc(B2OC(C)(C)C(C)(C)O2)c1C#N** | 94 | 0 | 11 | 0 | 1 |
| **OB(O)c1cc(Cl)ccc1Cl** | 94 | 0 | 5 | 0 | 1 |
| **OB(O)c1cccc2cnccc12** | 94 | 1 | 0 | 0 | 1 |
| **Cc1nc2cc(B3OC(C)(C)C(C)(C)O3)ccc2n1C** | 94 | 3 | 0 | 8 | 1 |
| **N#Cc1cncc(B(O)O)c1** | 94 | 6 | 0 | 1 | 1 |
| **COc1c(B(O)O)ccc(F)c1F** | 93 | 0 | 38 | 7 | 1 |
| **CC1(C)OB(c2ccnc3[nH]ccc23)OC1(C)C** | 93 | 3 | 27 | 0 | 1 |
| **OB(O)c1ccc(F)cc1Cl** | 93 | 0 | 12 | 0 | 1 |
| **OB(O)c1csc(Cl)c1** | 93 | 1 | 11 | 0 | 1 |
| **Cc1cc(F)ncc1B(O)O** | 93 | 0 | 0 | 0 | 1 |
| **OB(O)c1ccc(O)nc1F** | 93 | 0 | 0 | 0 | 1 |
| **NCc1cc(F)ccc1B(O)O** | 93 | 0 | 0 | 49 | 1 |
| **COc1c(C)cccc1B(O)O** | 92 | 0 | 46 | 2 | 1 |
| **OB(O)c1ccccc1CC(F)(F)F** | 92 | 0 | 34 | 4 | 1 |
| **CC(=O)Nc1ccccc1B(O)O** | 92 | 8 | 26 | 0 | 1 |
| **OB(O)c1cscc1Cl** | 92 | 4 | 13 | 0 | 1 |
| **OB(O)c1cccc(Cl)c1Cl** | 92 | 0 | 0 | 4 | 1 |
| **O=[N+]([O-])c1cc(B(O)O)ccc1Cl** | 92 | 0 | 0 | 17 | 1 |
| **O=c1[nH]cc(B(O)O)cc1F** | 92 | 7 | 0 | 5 | 1 |
| **CC(N)c1ccc(B2OC(C)(C)C(C)(C)O2)cc1F** | 91 | 9 | 29 | 0 | 1 |
| **OB(O)c1cccc(C(F)(F)F)c1F** | 91 | 0 | 26 | 1 | 1 |
| **OB(O)c1cc(Cl)ccc1F** | 91 | 0 | 20 | 3 | 1 |
| **CC1(C)OB(c2ccc3ncoc3c2)OC1(C)C** | 91 | 6 | 17 | 0 | 1 |
| **OB(O)c1ccccc1OC(F)(F)F** | 91 | 0 | 15 | 7 | 1 |
| **Cn1c(B2OC(C)(C)C(C)(C)O2)cnc1Cl** | 91 | 0 | 0 | 0 | 1 |
| **CC1(C)OB(c2cnc(C3CC3)o2)OC1(C)C** | 91 | 0 | 0 | 4 | 1 |
| **CN(C)Cc1ccc(B2OC(C)(C)C(C)(C)O2)s1** | 91 | 7 | 0 | 0 | 1 |
| **NCc1cccc(B(O)O)c1F** | 90 | 0 | 41 | 0 | 1 |
| **Cc1c(F)ccc(B(O)O)c1C** | 90 | 0 | 39 | 0 | 1 |
| **COc1ccc(B(O)O)c(OC)n1** | 90 | 0 | 37 | 0 | 1 |
| **O=[N+]([O-])c1cccc(B(O)O)c1** | 90 | 0 | 25 | 68 | 1 |

| Structure | ArI_adj %P | ArI_unknown_adj% | ArBr_adj %P | ArBr_unknown_adj% | Score |
|---|---|---|---|---|---|
| **CN(C)C(=O)c1cncc(B2OC(C)(C)C(C)(C)O2)c1** | 90 | 1 | 23 | 0 | 1 |
| **CC(C)Oc1ccc(B(O)O)c(Cl)c1** | 90 | 0 | 10 | 2 | 1 |
| **CS(=O)(=O)c1cncc(B(O)O)c1** | 90 | 0 | 0 | 3 | 1 |
| **OB(O)c1csc2ccccc12** | 90 | 0 | 0 | 3 | 1 |
| **CN(C)c1ccc(/C=C/B2OC(C)(C)C(C)(C)O2)cc1** | 90 | 0 | 0 | 33 | 1 |
| **Cc1c(B2OC(C)(C)C(C)(C)O2)cnn1C** | 90 | 1 | 0 | 2 | 1 |
| **N#Cc1cc(F)c(B(O)O)cc1F** | 90 | 7 | 0 | 8 | 1 |
| **O=[N+]([O-])c1ccc(B(O)O)cc1** | 89 | 0 | 44 | 0 | 1 |
| **OB(O)c1ccc(C2(O)CCC2)cc1** | 89 | 0 | 44 | 39 | 1 |
| **CCOc1ccc(B(O)O)c(Cl)c1** | 89 | 0 | 26 | 0 | 1 |
| **CC1(C)OB(c2cncc(OC(F)F)c2)OC1(C)C** | 89 | 0 | 0 | 1 | 1 |
| **CC1(C)OB(c2ccco2)OC1(C)C** | 89 | 0 | 0 | 7 | 1 |
| **Cc1ccncc1B1OC(C)(C)C(C)(C)O1** | 89 | 1 | 0 | 0 | 1 |
| **CC1(C)OB(c2cncc(C(F)F)c2)OC1(C)C** | 88 | 0 | 0 | 0 | 1 |
| **CC1(C)OB(c2ccc3nccnc3c2)OC1(C)C** | 88 | 4 | 0 | 0 | 1 |
| **Cn1cc(B2OC(C)(C)C(C)(C)O2)cn1** | 87 | 2 | 30 | 0 | 1 |
| **[B]c1cc(C(F)(F)F)ccc1F** | 87 | 1 | 29 | 0 | 1 |
| **CCCOc1cncc(B2OC(C)(C)C(C)(C)O2)c1** | 87 | 0 | 2 | 0 | 1 |
| **O=[N+]([O-])c1cc(B(O)O)ccc1F** | 87 | 0 | 0 | 0 | 1 |
| **CC1(C)OB(c2cncc(-n3cccn3)c2)OC1(C)C** | 87 | 0 | 0 | 17 | 1 |
| **CC(=O)c1ccc(Cl)c(B(O)O)c1** | 87 | 1 | 0 | 9 | 1 |
| **COc1cc(C#N)ccc1B(O)O** | 86 | 5 | 41 | 0 | 1 |
| **Cn1cc(B2OC(C)(C)C(C)(C)O2)c2ccccc21** | 86 | 9 | 16 | 0 | 1 |
| **OB(O)c1ccc(F)cc1O** | 86 | 0 | 0 | 6 | 1 |
| **CC1(C)OB(c2ccc(OC3CC3)cc2)OC1(C)C** | 85 | 6 | 39 | 4 | 1 |
| **CC1(C)OB(c2ccc(C(F)(F)F)nc2)OC1(C)C** | 85 | 1 | 35 | 0 | 1 |
| **Cc1ccc2cccnc2c1B1OC(C)(C)C(C)(C)O1** | 85 | 0 | 0 | 3 | 1 |
| **Cc1occc1B(O)O** | 84 | 6 | 29 | 0 | 1 |
| **OB(O)c1cccc2c1OC(F)(F)O2** | 84 | 1 | 26 | 1 | 1 |
| **Cc1cc(OC(F)F)ccc1B1OC(C)(C)C(C)(C)O1** | 84 | 0 | 17 | 4 | 1 |
| **OB(O)c1cccnc1Cl** | 84 | 0 | 0 | 2 | 1 |
| **COCCn1cc(B2OC(C)(C)C(C)(C)O2)cn1** | 83 | 0 | 48 | 0 | 1 |
| **Cc1cc(B2OC(C)(C)C(C)(C)O2)cnc1Cl** | 83 | 10 | 39 | 0 | 1 |
| **Cc1c(B(O)O)ccnc1F** | 83 | 2 | 26 | 0 | 1 |
| **CN(Cc1cccc(B(O)O)c1)C1CC1** | 83 | 17 | 0 | 100 | 1 |
| **COc1cc(C)ccc1B(O)O** | 82 | 0 | 34 | 7 | 1 |

| Structure | ArI_adj %P | ArI_unknown_adj% | ArBr_adj %P | ArBr_unknown_adj% | Score |
|---|---|---|---|---|---|
| **OCc1ccc(B(O)O)s1** | 82 | 2 | 5 | 0 | 1 |
| **COCCn1nc(C)c(B2OC(C)(C)C(C)(C)O2)c1C** | 82 | 3 | 0 | 3 | 1 |
| **OB(O)c1cccnc1F** | 81 | 9 | 25 | 2 | 1 |
| **COC(=O)Cc1ccccc1B1OC(C)(C)C(C)(C)O1** | 81 | 6 | 0 | 4 | 1 |
| **Cn1cnc2ccc(B3OC(C)(C)C(C)(C)O3)cc21** | 81 | 15 | 0 | 0 | 1 |
| **Cc1cc(B(O)O)co1** | 80 | 5 | 41 | 0 | 1 |
| **[B]c1c(OC)ccc(OC)c1OC** | 80 | 0 | 0 | 14 | 1 |
| **CCOC(=O)c1cc(B2OC(C)(C)C(C)(C)O2)[nH]n1** | 80 | 11 | 0 | 1 | 1 |
| **Cc1cnc(F)cc1B(O)O** | 79 | 4 | 31 | 0 | 1 |
| **CC1(C)OB(c2c[nH]c3ncccc23)OC1(C)C** | 79 | 1 | 0 | 0 | 1 |
| **OB(O)c1ccc(Cl)nc1** | 79 | 16 | 0 | 48 | 1 |
| **OB(O)c1cccc(F)c1C(F)(F)F** | 78 | 2 | 6 | 0 | 1 |
| **CC1(C)OB(c2cccc(F)c2C#N)OC1(C)C** | 78 | 5 | 0 | 5 | 1 |
| **CC1(C)OB(c2cccc(S(=O)(=O)F)c2)OC1(C)C** | 77 | 23 | 53 | 47 | 1 |
| **N#Cc1ccc(F)c(B(O)O)c1** | 77 | 0 | 13 | 1 | 1 |
| **CC1(C)OB(c2cccc3scnc23)OC1(C)C** | 77 | 11 | 2 | 0 | 1 |
| **OB(O)c1cc(F)nc(F)c1** | 77 | 0 | 0 | 0 | 1 |
| **CSc1ncc(B(O)O)cn1** | 77 | 0 | 0 | 2 | 1 |
| **CC(C)Oc1cccc(F)c1B(O)O** | 77 | 0 | 0 | 3 | 1 |
| **CC(C)(C)Sc1cccc(B(O)O)c1** | 76 | 0 | 7 | 4 | 1 |
| **N#Cc1cccc(B(O)O)c1Cl** | 76 | 0 | 0 | 0 | 1 |
| **Nc1ccncc1B(O)O** | 76 | 0 | 0 | 0 | 1 |
| **OB(O)c1ccc(F)cc1C(F)(F)F** | 76 | 0 | 0 | 9 | 1 |
| **N#Cc1cc(B(O)O)ccc1O** | 75 | 0 | 39 | 25 | 1 |
| **COc1cccc(Cl)c1B(O)O** | 75 | 0 | 19 | 0 | 1 |
| **Cc1nn(C(C)C)cc1B1OC(C)(C)C(C)(C)O1** | 75 | 2 | 9 | 1 | 1 |
| **OB(O)c1cc(F)ccc1C(F)(F)F** | 75 | 0 | 0 | 8 | 1 |
| **OB(O)c1ccccc1OCC1CC1** | 75 | 5 | 0 | 18 | 1 |
| **CC1(C)OB(c2cc(C(F)(F)F)n[nH]2)OC1(C)C** | 75 | 13 | 0 | 1 | 1 |
| **Cc1noc(C)c1B1OC(C)(C)C(C)(C)O1** | 74 | 2 | 0 | 10 | 1 |
| **COc1ncncc1B1OC(C)(C)C(C)(C)O1** | 74 | 7 | 0 | 11 | 1 |
| **O=C(O)c1cncc(B(O)O)c1** | 73 | 16 | 71 | 14 | 1 |
| **Cc1c(B2OC(C)(C)C(C)(C)O2)ccc2c1CCO2** | 72 | 3 | 16 | 0 | 1 |
| **OB(O)c1ccoc1** | 71 | 11 | 49 | 0 | 1 |
| **CC1(C)OB(c2csc(C(F)(F)F)n2)OC1(C)C** | 71 | 15 | 9 | 0 | 1 |
| **CCCn1nc(C)c(B2OC(C)(C)C(C)(C)O2)c1C** | 71 | 0 | 6 | 9 | 1 |
| **CC1(C)OB(c2cc(Cl)ccc2N)OC1(C)C** | 71 | 4 | 0 | 12 | 1 |

| Structure | ArI_adj %P | ArI_unknown_adj% | ArBr_adj %P | ArBr_unknown_adj% | Score |
|---|---|---|---|---|---|
| **CC1(C)OB(c2c(Cl)cccc2C#N)OC1(C)C** | 69 | 4 | 0 | 0 | 1 |
| **CC1(C)OB(C2=CC(=O)CCC2)OC1(C)C** | 68 | 3 | 0 | 100 | 1 |
| **CC1(C)OB(c2cc(F)ccc2C#N)OC1(C)C** | 68 | 6 | 0 | 0 | 1 |
| **O=[N+]([O-])c1ccccc1B(O)O** | 67 | 0 | 0 | 0 | 1 |
| **C/C=C\B(O)O** | 67 | 6 | 0 | 9 | 1 |
| **Cc1cccc(C#N)c1B1OC(C)(C)C(C)(C)O1** | 65 | 7 | 0 | 4 | 1 |
| **COc1ccc(B2OC(C)(C)C(C)(C)O2)c(C#N)c1** | 65 | 9 | 0 | 0 | 1 |
| **COc1cccc(OC)c1B(O)O** | 64 | 0 | 33 | 0 | 1 |
| **CC1(C)OB(c2cccc(OC(F)F)c2)OC1(C)C** | 64 | 0 | 10 | 3 | 1 |
| **N#Cc1c(Cl)ccc(B(O)O)c1F** | 63 | 2 | 0 | 0 | 1 |
| **COc1ncc(B(O)O)c(OC)n1** | 62 | 38 | 58 | 4 | 1 |
| **CC1(C)OB(c2ccsc2C#N)OC1(C)C** | 62 | 24 | 27 | 1 | 1 |
| **CCOc1cccc(F)c1B(O)O** | 62 | 22 | 9 | 7 | 1 |
| **N#Cc1ccc(Cl)cc1B(O)O** | 62 | 0 | 0 | 3 | 1 |
| **CCc1cccc(CC)c1B(O)O** | 62 | 0 | 0 | 6 | 1 |
| **Cc1ccc2[nH]ncc2c1B(O)O** | 60 | 0 | 0 | 0 | 1 |
| **CC1(C)OB(c2ccncc2Cl)OC1(C)C** | 60 | 0 | 0 | 2 | 1 |
| **OB(O)c1ccnc(Cl)c1** | 60 | 14 | 0 | 0 | 1 |
| **CC1(C)OB(c2cc(C#N)ccc2O)OC1(C)C** | 59 | 2 | 12 | 3 | 1 |
| **Cn1nc(C(F)(F)F)cc1B1OC(C)(C)C(C)(C)O1** | 54 | 1 | 0 | 0 | 1 |
| **CN(C)c1ccccc1B(O)O** | 53 | 0 | 0 | 1 | 1 |
| **COc1cc(/C=C/B2OC(C)(C)C(C)(C)O2)ccc1O** | 53 | 48 | 0 | 100 | 1 |
| **CC1(C)OB(c2ccncc2F)OC1(C)C** | 52 | 43 | 0 | 9 | 1 |
| **CC1(C)OB(c2ccc3c(c2)CCC3=O)OC1(C)C** | 49 | 0 | 87 | 0 | 1 |
| **CC1(C)OB(c2ccc(N)c(CO)c2)OC1(C)C** | 49 | 0 | 31 | 0 | 1 |
| **CC1(C)OB(c2ccc(CN)s2)OC1(C)C** | 43 | 28 | 0 | 0 | 1 |
| **COc1ccc(F)c(F)c1B(O)O** | 39 | 0 | 0 | 5 | 1 |
| **CC#Cc1cccc(B(O)O)c1** | 37 | 9 | 0 | 19 | 1 |
| **COc1cnccc1B(O)O** | 36 | 12 | 0 | 0 | 1 |
| **COc1cccc(F)c1B(O)O** | 33 | 14 | 0 | 12 | 1 |
| **CCCOc1ccc(B(O)O)c(C)c1** | 32 | 34 | 9 | 4 | 1 |
| **Cc1cc(C)c(C)c(B(O)O)c1C** | 29 | 0 | 0 | 1 | 1 |
| **Cc1cc(C)c(B(O)O)c(C)c1** | 29 | 6 | 0 | 5 | 1 |
| **COc1cc(C)cc(C)c1B(O)O** | 28 | 0 | 0 | 0 | 1 |
| **OB(O)c1ccc(C(F)(F)F)s1** | 25 | 11 | 0 | 3 | 1 |
| **CC1(C)OB(c2cnc3c(c2)CC(=O)N3)OC1(C)C** | 24 | 63 | 0 | 0 | 1 |
| **OB(O)C1CCCc2ccccc21** | 21 | 32 | 6 | 1 | 1 |

| Structure | ArI_adj %P | ArI_unknown_adj% | ArBr_adj %P | ArBr_unknown_adj% | Score |
|---|---|---|---|---|---|
| **COc1ccc(B(O)O)c(OC)c1** | 20 | 0 | 0 | 5 | 1 |
| **CB(O)O** | 19 | 32 | 0 | 37 | 1 |
| **Cc1cocc1B1OC(C)(C)C(C)(C)O1** | 18 | 47 | 0 | 10 | 1 |
| **CCOc1ccc(/C=C/B(O)O)cc1** | 16 | 29 | 0 | 100 | 1 |
| **CS(=O)c1ccc(B(O)O)cc1** | 15 | 0 | 0 | 0 | 1 |
| **CCCOc1ccc([B-](O)(O)O)nc1** | 14 | 7 | 0 | 14 | 1 |
| **COc1ccc(B(O)O)nc1** | 13 | 12 | 0 | 8 | 1 |
| **OB(O)c1c(F)cccc1F** | 12 | 0 | 0 | 1 | 1 |
| **OB(O)c1ccccc1CN1CCCC1** | 11 | 0 | 0 | 0 | 1 |
| **CCn1nccc1B1OC(C)(C)C(C)(C)O1** | 8 | 10 | 0 | 8 | 1 |
| **Cn1ccc2cccc(B(O)O)c21** | 7 | 4 | 0 | 0 | 1 |
| **OB(O)c1ccc(-c2ccccc2)s1** | 6 | 0 | 0 | 1 | 1 |
| **CC1(C)OB(c2ccc(OC(F)F)nc2)OC1(C)C** | 6 | 10 | 0 | 0 | 1 |
| **N#Cc1ccccc1B(O)O** | 4 | 0 | 0 | 1 | 1 |
| **CC1(C)COc2ccc(B3OC(C)(C)C(C)(C)O3)cc21** | 3 | 5 | 57 | 2 | 1 |
| **CC1(C)OB(c2cncc(C(F)(F)F)c2)OC1(C)C** | 3 | 10 | 0 | 4 | 1 |
| **OB(O)c1c(F)ccc2c1OCO2** | 3 | 11 | 0 | 0 | 1 |
| **O=[N+]([O-])c1ccc(Cl)cc1B(O)O** | 1 | 0 | 0 | 4 | 1 |
| **OB(O)C1=Cc2ccccc2C1** | 0 | 100 | 97 | 2 | 1 |
| **CC(=O)c1ccc(B(O)O)cc1F** | 0 | 0 | 94 | 0 | 1 |
| **OB(O)c1ccc2occc2c1** | 0 | 0 | 91 | 9 | 1 |
| **OB(O)c1ccc(C2CCCCC2)cc1** | 0 | 97 | 91 | 2 | 1 |
| **CCCCOc1cccc(B(O)O)c1** | 0 | 0 | 90 | 0 | 1 |
| **COc1ccc(OC)c(B(O)O)c1** | 0 | 0 | 88 | 0 | 1 |
| **CCOc1ccc(F)cc1B(O)O** | 0 | 0 | 87 | 0 | 1 |
| **CN(C)c1ccc(B(O)O)cc1** | 0 | 11 | 87 | 2 | 1 |
| **OB(O)c1ccc(F)c(F)c1** | 0 | 0 | 85 | 0 | 1 |
| **CC1(C)OB(c2ccc3c(c2)NC(=O)C3)OC1(C)C** | 0 | 100 | 85 | 0 | 1 |
| **CC(C)COc1cccc(B(O)O)c1** | 0 | 100 | 84 | 10 | 1 |
| **COc1ccc(B(O)O)cc1CO** | 0 | 0 | 83 | 0 | 1 |
| **COc1cc(Cl)cc(B(O)O)c1** | 0 | 100 | 78 | 5 | 1 |
| **Nc1ccc(B(O)O)cc1** | 0 | 5 | 74 | 12 | 1 |
| **Cc1cccc(B(O)O)c1O** | 0 | 40 | 72 | 0 | 1 |
| **COC(=O)c1ccc(F)c(B(O)O)c1** | 0 | 0 | 68 | 0 | 1 |
| **COc1nscc1B1OC(C)(C)C(C)(C)O1** | 0 | 0 | 68 | 1 | 1 |
| **OB(O)c1cccc(F)c1O** | 0 | 0 | 61 | 0 | 1 |
| **Cc1cc(C(F)(F)F)ccc1B(O)O** | 0 | 100 | 58 | 0 | 1 |
| **OB(O)c1ccc(Cc2ccccc2)cc1** | 0 | 100 | 55 | 9 | 1 |
| **CC(=O)c1ccc(B(O)O)c(F)c1** | 0 | 0 | 48 | 0 | 1 |
| **OB(O)c1ccc(O)c(Cl)c1** | 0 | 0 | 46 | 0 | 1 |
| **Cc1cc(B2OC(C)(C)C(C)(C)O2)ccn1** | 0 | 2 | 45 | 5 | 1 |
| **OB(O)c1ccccc1-c1ccccc1** | 0 | 0 | 35 | 0 | 1 |
| **Cc1ccc(B(O)O)c2ccccc12** | 0 | 100 | 35 | 0 | 1 |
| **O=C(O)c1ccc(B(O)O)s1** | 0 | 100 | 29 | 0 | 1 |

| Structure | ArI_adj %P | ArI_unknown_adj% | ArBr_adj %P | ArBr_unknown_adj% | Score |
|---|---|---|---|---|---|
| **CC(C)B(O)O** | 0 | 0 | 21 | 6 | 1 |
| **Cc1cccc2cc(B(O)O)cnc12** | 0 | 2 | 18 | 0 | 1 |
| **COC(=O)c1ccc(Cl)c(B(O)O)c1** | 0 | 0 | 15 | 4 | 1 |
| **COc1ccc(B2OC(C)(C)C(C)(C)O2)c(CN)c1** | 0 | 0 | 14 | 0 | 1 |
| **CC(=O)c1ccccc1B(O)O** | 0 | 0 | 7 | 0 | 1 |
| **COc1ccc(C)c(F)c1B(O)O** | 0 | 68 | 5 | 4 | 1 |
| **C=C(C)B1OC(C)(C)C(C)(C)O1** | 0 | 0 | 0 | 0 | 1 |
| **CC1(C)OB(C#CC2CC2)OC1(C)C** | 0 | 0 | 0 | 0 | 1 |
| **CC1(C)OB(c2ccc(-c3cn[nH]c3)cc2)OC1(C)C** | 0 | 0 | 0 | 0 | 1 |
| **CC1(C)OB(c2cccc(-c3ncc[nH]3)c2)OC1(C)C** | 0 | 0 | 0 | 0 | 1 |
| **COC(=O)c1ccc(F)cc1B(O)O** | 0 | 0 | 0 | 0 | 1 |
| **COC(=O)c1ccccc1B(O)O** | 0 | 0 | 0 | 0 | 1 |
| **COc1cc(B(O)O)sn1** | 0 | 0 | 0 | 0 | 1 |
| **Cc1ccc(B(O)O)cc1N** | 0 | 0 | 0 | 0 | 1 |
| **Cc1ccc(CNCC2CCCO2)s1** | 0 | 0 | 0 | 0 | 1 |
| **O=C(O)c1ccc(F)cc1B(O)O** | 0 | 0 | 0 | 0 | 1 |
| **OB(O)c1c(F)cc(F)cc1F** | 0 | 0 | 0 | 0 | 1 |
| **OB(O)c1cc2ccccc2o1** | 0 | 0 | 0 | 0 | 1 |
| **OB(O)c1ccc2[nH]cnc2c1** | 0 | 0 | 0 | 0 | 1 |
| **O=C(O)c1ccccc1B(O)O** | 0 | 0 | 0 | 1 | 1 |
| **O=[N+]([O-])c1ccc(Cl)c(B(O)O)c1** | 0 | 0 | 0 | 1 | 1 |
| **OB(O)c1ccccc1C(F)(F)F** | 0 | 0 | 0 | 1 | 1 |
| **OB(O)c1c(F)cc(O)cc1F** | 0 | 0 | 0 | 2 | 1 |
| **OB(O)/C=C/c1ccccc1** | 0 | 0 | 0 | 4 | 1 |
| **OB(O)c1c(F)ccc(O)c1F** | 0 | 0 | 0 | 4 | 1 |
| **CC(C)=CB(O)O** | 0 | 0 | 0 | 5 | 1 |
| **CCn1nc(C)cc1B1OC(C)(C)C(C)(C)O1** | 0 | 0 | 0 | 5 | 1 |
| **Cc1ccc(B2OC(C)(C)C(C)(C)O2)o1** | 0 | 0 | 0 | 6 | 1 |
| **N#CC=Cc1ccc(B(O)O)cc1** | 0 | 0 | 0 | 6 | 1 |
| **OB(O)c1cnn(Cc2ccccc2)c1** | 0 | 0 | 0 | 6 | 1 |
| **OB(O)c1c(F)cccc1Cl** | 0 | 0 | 0 | 7 | 1 |
| **CC(C)CB(O)O** | 0 | 0 | 0 | 8 | 1 |
| **CC(C)(O)C(C)(C)OB(O)C1=CCCCO1** | 0 | 0 | 0 | 11 | 1 |
| **OB(O)c1cccc(C(F)(F)F)c1O** | 0 | 0 | 0 | 15 | 1 |
| **CC(C)(O)C(C)(C)OB(O)/C=C/C1CC1** | 0 | 0 | 0 | 60 | 1 |
| **COc1cccc(/C=C/B(O)O)c1** | 0 | 0 | 0 | 100 | 1 |
| **CC1(C)OB(c2[nH]ncc2C(F)(F)F)OC1(C)C** | 0 | 2 | 0 | 0 | 1 |
| **COC(=O)c1ccc(Cl)cc1B(O)O** | 0 | 2 | 0 | 0 | 1 |
| **OB(O)c1ccc(O)cc1F** | 0 | 3 | 0 | 1 | 1 |
| **Cc1nn(C)cc1B1OC(C)(C)C(C)(C)O1** | 0 | 4 | 0 | 0 | 1 |
| **CCCB(O)O** | 0 | 4 | 0 | 2 | 1 |
| **CC(C)=C(C)B(O)O** | 0 | 4 | 0 | 100 | 1 |
| **[B]c1c(F)cccc1C(F)(F)F** | 0 | 5 | 0 | 3 | 1 |

| Structure | ArI_adj %P | ArI_unknown_adj% | ArBr_adj %P | ArBr_unknown_adj% | Score |
|---|---|---|---|---|---|
| **Cn1nncc1B1OC(C)(C)C(C)(C)O1** | 0 | 5 | 0 | 9 | 1 |
| **OB(O)c1c(F)ccc(F)c1F** | 0 | 6 | 0 | 0 | 1 |
| **CC(C)(C)C=CB(O)O** | 0 | 7 | 0 | 0 | 1 |
| **CC1(C)OB(c2cnsc2)OC1(C)C** | 0 | 7 | 0 | 3 | 1 |
| **Cc1ccc(B2OC(C)(C)C(C)(C)O2)cc1Cl** | 0 | 8 | 0 | 3 | 1 |
| **CCB(O)O** | 0 | 9 | 0 | 3 | 1 |
| **NC(=O)c1cccc(B(O)O)c1F** | 0 | 9 | 0 | 5 | 1 |
| **CC1(C)OB(C2(C#C[Si](C)(C)C)CC2)OC1(C)C** | 0 | 9 | 0 | 8 | 1 |
| **Cn1ncc(Cl)c1B1OC(C)(C)C(C)(C)O1** | 0 | 10 | 0 | 0 | 1 |
| **Cc1ccc(Cl)c(B(O)O)c1F** | 0 | 10 | 0 | 2 | 1 |
| **CC1(C)OB(c2ccsc2C(F)(F)F)OC1(C)C** | 0 | 10 | 0 | 3 | 1 |
| **Cc1ccc(B2OC(C)(C)C(C)(C)O2)cn1** | 0 | 10 | 0 | 6 | 1 |
| **CC1(C)OB(c2ccc(F)s2)OC1(C)C** | 0 | 11 | 0 | 0 | 1 |
| **Cc1nc(C)c(B2OC(C)(C)C(C)(C)O2)s1** | 0 | 11 | 0 | 6 | 1 |
| **Cn1nccc1B1OC(C)(C)C(C)(C)O1** | 0 | 11 | 0 | 8 | 1 |
| **O[B-](O)(O)c1ccccn1** | 0 | 12 | 0 | 0 | 1 |
| **O[B-](O)(O)c1cnccn1** | 0 | 12 | 0 | 0 | 1 |
| **CC1(C)OB(c2ccc(Cl)s2)OC1(C)C** | 0 | 13 | 0 | 0 | 1 |
| **CCOC(=O)c1ccccc1B(O)O** | 0 | 13 | 0 | 0 | 1 |
| **CC1(C)OB(c2cncs2)OC1(C)C** | 0 | 13 | 0 | 2 | 1 |
| **C=C(B1OC(C)(C)C(C)(C)O1)C(F)(F)F** | 0 | 13 | 0 | 3 | 1 |
| **Cn1ncc(C(F)(F)F)c1B1OC(C)(C)C(C)(C)O1** | 0 | 13 | 0 | 3 | 1 |
| **Cc1cc(B2OC(C)(C)C(C)(C)O2)on1** | 0 | 13 | 0 | 4 | 1 |
| **CC1(C)OB(c2cc(Cl)cs2)OC1(C)C** | 0 | 14 | 0 | 0 | 1 |
| **CC1(C)OB(C2=CCOC2)OC1(C)C** | 0 | 14 | 0 | 50 | 1 |
| **Cc1cc(B2OC(C)(C)C(C)(C)O2)sn1** | 0 | 15 | 0 | 13 | 1 |
| **CC1(C)OB(c2ccc(F)cc2C2CC2)OC1(C)C** | 0 | 16 | 0 | 0 | 1 |
| **N#Cc1ccc(O)c(B(O)O)c1** | 0 | 16 | 0 | 3 | 1 |
| **CC1(C)OB(c2cnc(C(F)(F)F)s2)OC1(C)C** | 0 | 16 | 0 | 6 | 1 |
| **OB(O)/C=C/c1ccc(F)cc1** | 0 | 16 | 0 | 30 | 1 |
| **Cc1ccc(B2OC(C)(C)C(C)(C)O2)s1** | 0 | 17 | 0 | 9 | 1 |
| **COCc1noc(C)c1B(O)O** | 0 | 18 | 0 | 0 | 1 |
| **C#CB1OC(C)(C)C(C)(C)O1** | 0 | 18 | 0 | 4 | 1 |
| **OB(O)C=Cc1ccccc1** | 0 | 18 | 0 | 7 | 1 |
| **COc1ccc([B-](O)(O)O)nc1** | 0 | 19 | 0 | 0 | 1 |
| **CC1(C)OB(c2cnn(C(F)(F)F)c2)OC1(C)C** | 0 | 20 | 0 | 0 | 1 |
| **Cc1ccsc1B1OC(C)(C)C(C)(C)O1** | 0 | 20 | 0 | 0 | 1 |
| **CC1(C)OB(c2cccs2)OC1(C)C** | 0 | 20 | 0 | 4 | 1 |
| **Cc1ncc(B2OC(C)(C)C(C)(C)O2)s1** | 0 | 20 | 0 | 5 | 1 |
| **CC1(C)OB(c2cnc(Cl)s2)OC1(C)C** | 0 | 20 | 0 | 7 | 1 |
| **OB(O)c1ccc(O)c(F)c1** | 0 | 21 | 0 | 1 | 1 |

| Structure | ArI_adj %P | ArI_unknown_adj% | ArBr_adj %P | ArBr_unknown_adj% | Score |
|---|---|---|---|---|---|
| **CN1CC=C(B2OC(C)(C)C(C)(C)O2)CC1** | 0 | 22 | 0 | 0 | 1 |
| **Cc1coc(B2OC(C)(C)C(C)(C)O2)c1** | 0 | 22 | 0 | 4 | 1 |
| **OB(O)c1cccs1** | 0 | 24 | 0 | 1 | 1 |
| **CC1(C)OB(C=C2CCCCC2)OC1(C)C** | 0 | 25 | 0 | 0 | 1 |
| **Cc1ccc([N+](=O)[O-])cc1B(O)O** | 0 | 27 | 0 | 2 | 1 |
| **C=C(CO)B1OC(C)(C)C(C)(C)O1** | 0 | 32 | 0 | 100 | 1 |
| **OB(O)c1cscc1C(F)F** | 0 | 33 | 0 | 0 | 1 |
| **OB(O)c1cnc(Cl)nc1** | 0 | 37 | 0 | 0 | 1 |
| **Cn1c(B(O)O)cc2ccc(Cl)cc21** | 0 | 37 | 0 | 14 | 1 |
| **CC1(C)OB(c2ccc(C#N)s2)OC1(C)C** | 0 | 42 | 0 | 13 | 1 |
| **CC1(C)OB(C2=CCCCC2)OC1(C)C** | 0 | 42 | 0 | 100 | 1 |
| **CC1(C)OB(C2=CCOCC2)OC1(C)C** | 0 | 46 | 0 | 2 | 1 |
| **O=Cc1ccsc1B(O)O** | 0 | 79 | 0 | 0 | 1 |
| **CC(C)=CB1OC(C)(C)C(C)(C)O1** | 0 | 100 | 0 | 11 | 1 |
| **CC1(C)OB(c2ccoc2)OC1(C)C** | 0 | 100 | 0 | 17 | 1 |
| **CC1(C)OB(C2=CCCOC2)OC1(C)C** | 0 | 100 | 0 | 100 | 1 |
| **OB(O)c1ccc(B(O)O)cc1** | 0 | 100 | 0 | 100 | 1 |
| **OB(O)c1cccc(B(O)O)c1** | 0 | 100 | 0 | 100 | 1 |

#### 9. Building block to sequence association information

**Supplementary Table 14 | Building block to sequence association information**

| Cycle | Tag (5’-3’) | Smiles |
|---|---|---|
| 1 | GCACCATCT | O=C(O)CCCCCNC(=O)[C@@H]1NC[C@@H]1c1ccc(I)cc1 |
| 1 | GCACTCTCT | O=C(O)CCCCCNC(=O)[C@H]1NC[C@H]1c1ccc(I)cc1 |
| 1 | TCTCGCTCT | O=C(O)CCCCCNC(=O)[C@H]1NC[C@@H]1c1ccc(I)cc1 |
| 1 | ATCTCGGCT | O=C(O)CCCCCNC(=O)[C@@H]1NC[C@H]1c1ccc(I)cc1 |
| 1 | ATGGAGGCT | O=C(O)CCCCCNC(=O)[C@@H]1NCC[C@@H]1c1ccc(I)cc1 |
| 1 | CGCACAACT | O=C(O)CCCCCNC(=O)[C@H]1NCC[C@H]1c1ccc(I)cc1 |
| 1 | CGCACTTCT | O=C(O)CCCCCNC(=O)[C@H]1NCC[C@@H]1c1ccc(I)cc1 |
| 1 | TGGAAGCCT | O=C(O)CCCCCNC(=O)[C@@H]1NCC[C@H]1c1ccc(I)cc1 |
| 1 | TGGAGACCT | O=C(O)[C@@H]1NC[C@@H]1c1cccc(I)c1 |
| 1 | AGATGGCCT | O=C(O)[C@H]1NC[C@H]1c1cccc(I)c1 |
| 1 | AGGACTGCT | O=C(O)[C@H]1NC[C@@H]1c1cccc(I)c1 |
| 1 | AGTACCGCT | O=C(O)[C@@H]1NC[C@H]1c1cccc(I)c1 |
| 1 | CTCTACGCT | O=C(O)[C@@H]1NCC[C@@H]1c1cccc(I)c1 |
| 1 | GACAGACCT | O=C(O)[C@H]1NCC[C@H]1c1cccc(I)c1 |
| 1 | GGCAACACT | O=C(O)[C@H]1NCC[C@@H]1c1cccc(I)c1 |
| 1 | GACCGAACT | O=C(O)[C@@H]1NCC[C@H]1c1cccc(I)c1 |
| 1 | GAGCTGACT | O=C(O)[C@@H]1NCCC[C@@H]1c1cccc(I)c1 |
| 1 | GCGAAGACT | O=C(O)[C@H]1NCCC[C@H]1c1cccc(I)c1 |
| 1 | GGCGATTCT | O=C(O)[C@H]1NCCC[C@@H]1c1cccc(I)c1 |
| 1 | TAGACCGCT | O=C(O)[C@@H]1NCCC[C@H]1c1cccc(I)c1 |
| 1 | CGACCAACT | O=C(O)CCCC(=O)N1C[C@H](c2ccc(I)cc2)C2(CNC2)C1 |
| 1 | GCTTCCACT | O=C(O)CCCC(=O)N1C[C@@H](c2ccc(I)cc2)C2(CNC2)C1 |
| 1 | CTGAGGACT | O=C(O)CCCC(=O)N1CC2(CNC[C@@H]2c2ccc(I)cc2)C1 |
| 1 | GACCGTTCT | O=C(O)CCCC(=O)N1CC2(CNC[C@H]2c2ccc(I)cc2)C1 |
| 1 | GAGTAGCCT | O=C(O)[C@@H]1C[C@H](Oc2ccc(I)cc2)CN1 |
| 1 | GCTCGAACT | O=C(O)[C@@H]1C[C@@H](Oc2ccc(I)cc2)CN1 |
| 1 | GCTGCTTCT | O=C(O)[C@H]1C[C@@H](Oc2ccc(I)cc2)CN1 |
| 1 | GCTGTCTCT | O=C(O)[C@H]1C[C@H](Oc2ccc(I)cc2)CN1 |
| 1 | TACGAGCCT | O=C(O)[C@@H]1C[C@H](Oc2cccc(I)c2)CN1 |
| 1 | TACGCGACT | O=C(O)[C@@H]1C[C@@H](Oc2cccc(I)c2)CN1 |
| 1 | TCCTCTGCT | O=C(O)[C@H]1C[C@@H](Oc2cccc(I)c2)CN1 |
| 1 | TCGGCATCT | O=C(O)[C@H]1C[C@H](Oc2cccc(I)c2)CN1 |
| 1 | TCGGTCACT | O=C(O)[C@@H]1C[C@H](OCc2ccc(I)cc2)CN1 |
| 1 | TCTGCCTCT | O=C(O)[C@@H]1C[C@@H](OCc2ccc(I)cc2)CN1 |
| 1 | TTGGCGTCT | O=C(O)[C@H]1C[C@@H](OCc2ccc(I)cc2)CN1 |
| 1 | TTGGTGCCT | O=C(O)[C@H]1C[C@H](OCc2ccc(I)cc2)CN1 |
| 1 | AAGGCCACT | O=C(O)[C@@H]1C[C@H](OCc2cccc(I)c2)CN1 |
| 1 | ACACCGACT | O=C(O)[C@@H]1C[C@@H](OCc2cccc(I)c2)CN1 |
| 1 | AGAACGGCT | O=C(O)[C@H]1C[C@@H](OCc2cccc(I)c2)CN1 |
| 1 | AGAGTGCCT | O=C(O)[C@H]1C[C@H](OCc2cccc(I)c2)CN1 |
| 1 | TCACGACCT | OC[C@@H]1[C@@H](c2ccc(Br)cc2)[C@@H]2CNCCCCN12 |
| 1 | TCCACCACT | O=C1CNC[C@H]2[C@H](c3ccc(Br)cc3)[C@@H](CO)N12 |
| 1 | TCGCTGTCT | OC[C@H]1[C@@H](c2ccc(Br)cc2)[C@H]2CNCCCCN21 |
| 1 | TCTTGGCCT | O=C1CNC[C@H]2[C@@H](c3ccc(Br)cc3)[C@@H](CO)N12 |
| 1 | TGACGGACT | O=C1CNC[C@@H]2[C@H](c3ccc(Br)cc3)[C@@H](CO)N12 |

| Cycle | Tag (5’-3’) | Smiles |
|---|---|---|
| 1 | GATGCTCCT | O=C1CNC[C@H]2[C@@H](c3ccc(Br)cc3)[C@H](CO)N12 |
| 1 | GAGGTTCCT | O=C1CNC[C@@H]2[C@@H](c3ccc(Br)cc3)[C@H](CO)N12 |
| 1 | TACGGTGCT | OC[C@@H]1[C@H](c2ccc(Br)cc2)[C@H]2CNCCCCN12 |
| 1 | GCTGCAACT | OC[C@@H]1[C@H](c2ccc(Br)cc2)[C@@H]2CNCCCCN12 |
| 1 | GCTTAGGCT | OC[C@@H]1[C@@H](c2ccc(Br)cc2)[C@H]2CNCCCCN12 |
| 1 | GGTCTGACT | CNC[C@H]1Oc2cc(Br)ccc2S(=O)(=O)N([C@@H](C)CO)C[C@H]1C |
| 1 | ACTTCGCCT | CNC[C@H]1Oc2cc(Br)ccc2S(=O)(=O)N([C@H](C)CO)C[C@H]1C |
| 1 | AGACGCACT | CNC[C@@H]1Oc2cc(Br)ccc2S(=O)(=O)N([C@@H](C)CO)C[C@@H]1C |
| 1 | AGCGAGACT | CNC[C@@H]1Oc2cc(Br)ccc2S(=O)(=O)N([C@H](C)CO)C[C@@H]1C |
| 1 | AGTCGTGCT | CNC[C@@H]1Oc2cc(Br)ccc2S(=O)(=O)N([C@H](C)CO)C[C@H]1C |
| 1 | ATCCAGCCT | CNC[C@H]1Oc2cc(Br)ccc2S(=O)(=O)N([C@H](C)CO)C[C@@H]1C |
| 1 | CGAGTCTCT | CNC[C@@H]1Oc2cc(Br)ccc2S(=O)(=O)N([C@@H](C)CO)C[C@H]1C |
| 1 | CAGCGATCT | CCOC(=O)c1cc2c(c(-c3ccc(Br)cc3)n1)[C@@H](CCO)NC2 |
| 1 | CAGCTACCT | CCOC(=O)c1cc2c(c(-c3cccc(Br)c3)n1)[C@H](CCO)NC2 |
| 1 | CATACCGCT | O=C(O)c1cc2c(c(-c3ccc(Br)cc3)n1)[C@@H](CCO)NC2 |
| 1 | CGACAGTCT | O=C(O)c1cc2c(c(-c3cccc(Br)c3)n1)[C@H](CCO)NC2 |
| 2 | ACAGCAGAC | Cc1nc2cccnn2c1C(=O)O |
| 2 | CGCCTATAC | Cn1ncc(CCC(=O)O)n1 |
| 2 | GCCAATGAC | N#Cc1cccc([C@H]2C[C@H]2C(=O)O)c1 |
| 2 | GTTCGTCAC | Cc1nc(CC(=O)O)c(C)s1 |
| 2 | ACTTGCCAC | O=C(O)[C@@H]1CCc2nnnn2CC1 |
| 2 | CGTCATCAC | O=C(O)c1coc(-c2ccccn2)n1 |
| 2 | GGATTGCAC | O=C1Cc2c(C(=O)O)ccnc2N1 |
| 2 | TGATGCGAC | Cc1nc2ccc(C(=O)O)cc2n1C |
| 2 | ACGCTTGAC | Cc1ccsc1[C@@H]1C[C@@H]1C(=O)O |
| 2 | CGCTATGAC | COc1ccc2cc(C(=O)O)[nH]c2c1 |
| 2 | GCTGCATAC | CCn1nc(C)c(C(=O)O)n1 |
| 2 | TCCAGAGAC | O=C(O)CC[C@@H]1CCc2ccccc21 |
| 2 | AGAGACGAC | CCn1nc(C2CC2)cc1C(=O)O |
| 2 | CGTGGTTAC | C[C@@H]1CCc2nc(C(=O)O)cn2C1 |
| 2 | GGCTATCAC | O=C(O)c1cc([N+](=O)[O-])cnc1O |
| 2 | TGCCACTAC | O=C(O)c1ccc(-n2ccnc2)nn1 |
| 2 | AGTACGGAC | C/C(=C\c1ccccn1)C(=O)O |
| 2 | CGCTTAGAC | CCNc1cccnc1C(=O)O |
| 2 | GGACCATAC | O=C(O)C[C@@H]1CCC2(CCC2)CO1 |
| 2 | TCGTCACAC | CCOc1oc(C)nc1C(=O)O |
| 2 | AGCAAGGAC | O=C(O)c1ncoc1-c1ccon1 |
| 2 | CTACCAGAC | O=C(O)c1ccc2oc(=O)[nH]c2n1 |
| 2 | GGTGGATAC | O=C(O)C1=NN(CCO)C(=O)CC1 |
| 2 | TGCTCAGAC | Cc1cn2c(n1)C[C@H](C(=O)O)CC2 |
| 2 | ATCTGCCAC | O=C(O)C1=C(c2ccccc2)CCC1 |
| 2 | CGTCAACAC | Cc1cc(C(=O)O)c(C)n1C1CC1 |
| 2 | GGCAGTAAC | C=CCNc1ncc(C(=O)O)s1 |
| 2 | TGCCTCTAC | Cn1nc(CC(=O)O)c2ccccc21 |
| 2 | AGCGTAGAC | Cc1cnc2nc(C(=O)O)nn2c1 |
| 2 | CTACGCTAC | O=C(O)c1ccnc(-c2ncc[nH]2)c1 |
| 2 | GTGTACGAC | O=C(O)C[C@@H]1COc2ccccc21 |
| 2 | TGGAGTGAC | O=C(O)COCc1cc[nH]c(=O)c1 |
| 2 | CAACGTGAC | O=C(O)c1nsc2ccccc12 |

| Cycle | Tag (5'-3') | Smiles |
|---|---|---|
| 2 | CTATGGCAC | Cc1nc(CC2CC2)oc1C(=O)O |
| 2 | GGCATACAC | O=C(O)CN1Cc2ccccc2C1=O |
| 2 | TGGAACCAC | O=C(O)Cn1cc(C2CCC2)nn1 |
| 2 | ATCGACGAC | Cn1ccc(-c2cc(C(=O)O)[nH]n2)c1 |
| 2 | CTGGTAGAC | CS(=O)(=O)c1cc(C(=O)O)co1 |
| 2 | GTGTGAGAC | Cc1ccn([C@H](C)CC(=O)O)n1 |
| 2 | TGTGAGCAC | C#CCOc1ccc(C(=O)O)cc1 |
| 2 | CACACAGAC | O=C(O)c1coc2cccc(O)c12 |
| 2 | CTCACCAAC | O=C(O)c1ccc(=O)n(CCF)n1 |
| 2 | GGTAACGAC | Cn1cc(C(=O)O)c(=O)n(C)c1=O |
| 2 | TTCGGCAAC | CCC(=O)N[C@@H]1CC[C@@H](C(=O)O)C1 |
| 2 | ATGTCGCAC | O=C(O)c1cccc(-n2ccnn2)c1 |
| 2 | CTGTAGCAC | Cc1nc(C)n(CCCC(=O)O)n1 |
| 2 | GTTCACCAC | O=C(O)c1cn([C@H]2CC[C@H]2O)nn1 |
| 2 | TTAGGCGAC | CCc1cc(C(=O)O)cc(Cl)n1 |
| 2 | CACAGAGAC | CO[C@H]1C[C@H](C(=O)O)N(C(C)=O)C1 |
| 2 | CTCGTGAAC | CS(=O)(=O)N1CC(C(=O)O)C1 |
| 2 | GGTATGGAC | CC1(C)C[C@]2(CCO1)C[C@@H]2C(=O)O |
| 2 | AACCGACAC | Cc1cccc2nc(C(=O)O)cn12 |
| 2 | ATTCGGCAC | Cc1oc(C2CC2)nc1C(=O)O |
| 2 | CTTCGGAAC | Cc1cc(C)cc(CC(=O)O)c1 |
| 2 | TACTCGCAC | CC[C@H](O)Cn1cc(C(=O)O)nn1 |
| 2 | TTGGAGCAC | Cc1cc(C2(C(=O)O)CC2)on1 |
| 2 | CACGCATAC | N#Cc1nc(C(=O)O)ccc1Cl |
| 2 | CTGAACCAC | Cc1ncsc1COCC(=O)O |
| 2 | GGTGGTAAC | O=C(O)c1n[nH]c2c1COCC2 |
| 2 | AACCTCCAC | COCCN1C[C@@H](C(=O)O)CC1=O |
| 2 | CACCTCAAC | O=C(O)CCNC(=O)[C@@H]1CCCO1 |
| 2 | GCACCTAAC | O=C(O)CC1=CCOCC1 |
| 2 | TAGCCTGAC | O=C(O)/C=C/c1ccc2c(c1)CCO2 |
| 2 | AACAGCGAC | O=C(O)c1c[nH]nc1[C@@H]1CCOC1 |
| 2 | CACGCTAAC | O=C(O)c1conc1C1CCCC1 |
| 2 | CTTCGCTAC | COCCn1cc(C(=O)O)c(C)n1 |
| 2 | GTCACTCAC | CN(C)c1cc(C#N)cc(C(=O)O)c1 |
| 2 | AACGGACAC | O=C1N=c2cc(C(=O)O)cnc2=N1 |
| 2 | CAGTCCTAC | O=C1NC(=O)C2(CC(C(=O)O)C2)N1 |
| 2 | GCCACATAC | CC1=C(C(=O)O)C(=O)OC1(C)C |
| 2 | TAGTGCGAC | O=C(O)c1cnoc1-c1ccco1 |
| 2 | AACGCAGAC | Cc1ncc(C(=O)O)c(C(C)C)n1 |
| 2 | CAGAACGAC | Cc1cc(C(=O)O)c2n[nH]cc2c1 |
| 2 | CTTGCACAC | O=C(O)Cc1n[nH]c2c1CCCC2 |
| 2 | GTCGCTAAC | O=C(O)CCc1cc2ccccc2o1 |
| 2 | ACATCGCAC | O=C(O)C#Cc1ccc2c(c1)OCO2 |
| 2 | CGAACACAC | CC(C)Cc1nc(C(=O)O)c[nH]1 |
| 2 | GCGTGATAC | Cc1nn(C)c(C)c1OCC(=O)O |
| 2 | TCCTCACAC | O=C(O)[C@@H]1CN2CCC[C@@H]2CO1 |
| 2 | AAGTCGCAC | Cc1cc(C(=O)O)c2c(C)n[nH]c2n1 |
| 2 | CGAAGACAC | O=C(O)c1cc2occc2s1 |
| 2 | GCAGACTAC | Cn1nnc2c1NCC[C@@H]2C(=O)O |

| Cycle | Tag (5'-3') | Smiles |
|---|---|---|
| 2 | GTCTGCTAC | O=C(O)c1cc(C2CC2)[nH]c(=O)c1 |
| 2 | ACCTCTGAC | C[C@H](Oc1cccc(C#N)c1)C(=O)O |
| 2 | CGTAGTGAC | Cc1nn(CC(=O)O)c(=O)o1 |
| 2 | GCTACGAAC | C#CC1(O)CC(C(=O)O)C1 |
| 2 | TCGTCCTAC | CS(=O)c1ccccc1C(=O)O |
| 2 | ACACACCAC | O=C(O)CCn1ccc2ccccc21 |
| 2 | CGCCATAAC | C[C@@H]1Cc2cccc(C(=O)O)c2O1 |
| 2 | GCATGTCAC | O=C(O)c1n[nH]c([N+](=O)[O-])c1Cl |
| 2 | GTGTCCTAC | O=C(O)c1coc2c1C(=O)CCC2 |
| 2 | ACGACCAAC | CC1(C)[C@H](C#N)[C@@]1(C#N)C(=O)O |
| 2 | CGTATCGAC | C#CCNc1ncc(C(=O)O)c(C)n1 |
| 2 | GCTCATCAC | O=C(O)c1nc2n(n1)CCCN2 |
| 2 | TCTCGTGAC | COCc1ccc(C(=O)O)cc1F |
| 2 | ACTCCGAAC | O=C(O)c1cc2n(n1)CCC[C@H]2O |
| 2 | AGACTCGAC | CC(=O)Nc1cccc(C(=O)O)n1 |
| 2 | CACCTCTAC | O=C(O)c1ncoc1[C@@H]1CCCO1 |
| 2 | GCAGGTTAC | CC(C)(C(=O)O)n1cc(Cl)cn1 |
| 2 | GTTCACGAC | O=C(O)c1cccc(-c2nnco2)c1 |
| 2 | ACAGCGAAC | CCN1C(=O)CC[C@H]1CC(=O)O |
| 2 | ATGGTGGAC | O=C(O)c1ccc2scnc2c1 |
| 2 | GGAACCAAC | Cc1sc(Cl)c(C(=O)O)c1C |
| 2 | TCGCAAGAC | O=C(O)C1(n2cccn2)CC1 |
| 2 | AGAGAGGAC | O=C(O)Cn1ncc2cccnc21 |
| 2 | CATCCAGAC | COC1(C(=O)O)CS(=O)(=O)C1 |
| 2 | GCCTGTAAC | CC(C)c1ccc(C(=O)O)c(=O)[nH]1 |
| 2 | TAAGCCGAC | O=C(O)CNC(=O)Cn1cncn1 |
| 2 | ACAGGACAC | O=C(O)c1ncc2c(F)cccn12 |
| 2 | CAAGGACAC | Cc1n[nH]c(C)c1[C@H](C)C(=O)O |
| 2 | GGACGTTAC | Cc1nc2c(C(=O)O)cccc2o1 |
| 2 | TCTACCGAC | Cc1nc(C(=O)O)c2n1CCCC2 |
| 2 | AGAGGACAC | CC1=C(C(=O)O)Cc2ccccc2O1 |
| 2 | CGAACCTAC | O=C(O)c1cnc(CC2CC2)s1 |
| 2 | GCGTAACAC | O=C(O)c1cc(-n2cccn2)ccn1 |
| 2 | TATGCGGAC | O=C(O)Cc1csc(C2CC2)n1 |
| 2 | ACATGCCAC | Cn1cnn(CCC(=O)O)c1=O |
| 2 | CACTGACAC | Cc1nnc(CCCC(=O)O)n1C |
| 2 | GGCAATCAC | O=C(O)c1cc2sccc2[nH]1 |
| 2 | TGCGTGAAC | Cc1nc2c([nH]1)CC[C@H](C(=O)O)C2 |
| 2 | AGAGTGGAC | C/C(=C(/F)C(=O)O)C1CC1 |
| 2 | CGTATCCAC | O=C(O)C[C@@H]1C(=O)Nc2ccccc21 |
| 2 | GCGTATCAC | Cc1cc2ccc(C(=O)O)cn2c1 |
| 2 | TCACCTCAC | COCc1nc(C(=O)O)co1 |
| 2 | ACCGACAAC | O=C(O)/C=C1\CN2CCC1CC2 |
| 2 | CAGCCTAAC | CC(=O)[C@@H]1C[C@H](CC(=O)O)C1(C)C |
| 2 | GGTAAGCAC | O=C(O)C1=NO[C@@H](c2ccccc2)C1 |
| 2 | TGCTCACAC | Cc1onc(CO)c1C(=O)O |
| 2 | AGGACAGAC | O=C(O)c1ccc2c(c1)B(O)OC2 |
| 2 | CTACGAGAC | O=C(O)C1=C[C@@H](O)[C@@H](O)[C@H](O)C1 |
| 2 | GCTTGAGAC | O=C(O)[C@@H]1CCc2nnc(O)n2C1 |

| Cycle | Tag (5'-3') | Smiles |
|---|---|---|
| 2 | TCACGCTAC | CO[C@H](C)c1nc(C(=O)O)cs1 |
| 2 | ACTGCGAAC | CCOc1cccc(CC(=O)O)n1 |
| 2 | CTCGCTAAC | O=C(O)c1nc2c(s1)CCC2 |
| 2 | GGTACACAC | O=C(O)c1csnc1Cl |
| 2 | TGGAGAGAC | O=C(O)c1cc2c(s1)CCOC2 |
| 2 | AGTCCTGAC | C=C(C)C[C@@H](C)C(=O)O |
| 2 | CTCCTACAC | Cn1nnc(C(=O)O)c1C(F)F |
| 2 | GGAGATGAC | O=C1CC[C@@H](C(=O)O)n2cccc21 |
| 2 | TCGCGTTAC | Cc1c(C(=O)O)nn2c1OCCC2 |
| 2 | AGACCACAC | Cn1nnc2c(C(=O)O)ccnc21 |
| 2 | CTGACTGAC | NS(=O)(=O)c1ccc(C(=O)O)cc1 |
| 2 | GGTATGCAC | CC(C)(C)OC(=O)N1CC(C=O)C1 |
| 2 | TGTCAGCAC | COc1cncc(CCC=O)c1 |
| 2 | AGTGCTCAC | O=Cc1ccc2nccnc2c1 |
| 2 | CTCGTCTAC | Cc1ccc(CCC=O)cn1 |
| 2 | GGTAGTGAC | O=Cc1cc(-n2ccnc2)cs1 |
| 2 | TGAAGGCAC | O=Cc1cc(CO)ccc1O |
| 2 | AGCGAAGAC | CS(=O)c1ccc(C=O)cc1 |
| 2 | CTGTTCCAC | O=Cc1cc(C(F)(F)F)[nH]n1 |
| 2 | GGTCTCAAC | Cc1nn(-c2ccncc2)cc1C=O |
| 2 | TGTGGTCAC | Cn1cnc(C=O)c1 |
| 2 | AGTGGTCAC | Cc1n[nH]c(Cl)c1C=O |
| 2 | CTCTTGGAC | Cn1nccc1-c1ccccc1C=O |
| 2 | GGTCACTAC | O=Cc1occc1C(=O)O |
| 2 | TGACGAGAC | O=Cc1n[nH]cc1-c1ccccc1 |
| 2 | AGGAGAGAC | O=Cc1ccc(C(=O)NC2CC2)cc1 |
| 2 | CTTGGTGAC | O=Cc1cnc[nH]1 |
| 2 | GGTGGTTAC | N#Cc1ccc(CCC=O)cc1 |
| 2 | TTCCAGGAC | O=Cc1cccc(Cc2cnc[nH]2)c1 |
| 2 | ATGGACGAC | CS(=O)(=O)c1cccc(C=O)c1 |
| 2 | CTGCATCAC | NS(=O)(=O)c1ccc(C=O)o1 |
| 2 | GGTGCTAAC | O=Cc1cnc(-c2ccccc2)[nH]1 |
| 2 | TGCTGAGAC | Cn1nc(-c2ccccc2)cc1C=O |
| 2 | AGGATGCAC | Cc1ncc(C=O)n1-c1ccccc1 |
| 2 | GCATCCAAC | O=Cc1cnn(Cc2ccccc2)c1 |
| 2 | GTGGAAGAC | Cc1c(C(F)(F)F)n[nH]c1C=O |
| 2 | TTCTCGGAC | COC(=O)c1[nH]cc(C=O)c1C |
| 2 | CAACCTGAC | O=Cc1cn[nH]c1 |
| 2 | CTGCTTCAC | O=Cc1ccc2cn[nH]c2c1 |
| 2 | GTAACGCAC | O=Cc1ccnn1-c1ccccn1 |
| 2 | TGGCACAAC | N#Cc1csc(C=O)c1 |
| 2 | ATAGGCGAC | O=Cc1ccnc(-c2ccccc2)n1 |
| 2 | GCATTGGAC | O=Cc1cc(C2CC2)[nH]n1 |
| 2 | GTGTCTCAC | COc1cccc2cc(C=O)oc12 |
| 2 | TTGAGCGAC | N#CCOc1ccccc1C=O |
| 2 | CAAGGTGAC | O=Cc1cccc(-c2ccccn2)c1 |
| 2 | CTGGTTGAC | O=Cc1ccc(-n2ccnc2)cc1 |
| 2 | GTCTAGGAC | NS(=O)(=O)c1cc(C=O)co1 |
| 2 | TGGCTGAAC | O=Cc1cc(O)c2ccsc2c1 |

| Cycle | Tag (5'-3') | Smiles |
|---|---|---|
| 2 | ATCCTCGAC | Cn1cc(-c2cccc(C=O)c2)cn1 |
| 2 | GCCATTGAC | O=Cc1ccc(O)c(F)c1 |
| 2 | TAAGGCGAC | Cc1cc(C=O)cnc1-n1cccn1 |
| 2 | TTGGCACAC | Cn1cc(C=O)c(-c2cccnc2)n1 |
| 2 | CACAGACAC | Cc1ccccc1-n1cc(C=O)cn1 |
| 2 | GCACACAAC | Cc1nc2ccccc2nc1C=O |
| 2 | GTGACCTAC | COC(=O)c1cc(C=O)cn1C |
| 2 | AACCGTCAC | O=Cc1ccn(-c2cccnc2)c1 |
| 2 | ATGCGAGAC | CC(C)(C#N)c1ccc(C=O)cc1 |
| 2 | GCCTCATAC | O=Cc1cn[nH]c1-c1ccccc1 |
| 2 | TAGCCTCAC | Cc1ccc2cc(C=O)ccc2n1 |
| 2 | AAGGCTGAC | COc1nc2ccccc2cc1C=O |
| 2 | ACACCAGAC | O=Cc1nc(-c2ccccc2)c[nH]1 |
| 2 | CGCTACAAC | CCOC(=O)c1cc(C=O)[nH]n1 |
| 2 | GGACTTGAC | O=Cc1cn(-c2ccccc2)cn1 |
| 2 | TCTGAGGAC | O=Cc1nccn1-c1ccccc1 |
| 2 | AGACCTGAC | O=Cc1ccccc1-n1cccn1 |
| 2 | ATGCACGAC | Cn1cc(-n2ccc(C=O)c2)cn1 |
| 2 | CTTGCAGAC | Cc1cccc(-c2n[nH]cc2C=O)c1 |
| 2 | TGTGCCAAC | CCCn1cc(C=O)c(C2CC2)n1 |
| 2 | AGACGCTAC | CNC(=O)c1ccc(C=O)cc1 |
| 2 | CGCTCATAC | O=Cc1cnn(-c2ccccn2)c1 |
| 2 | GGAGCATAC | Cc1ccncc1CCC=O |
| 2 | TCTTGCGAC | O=Cc1ccc(-n2ccnc2)nc1 |
| 2 | AGATGCGAC | O=Cc1ccc(-c2cccnc2)o1 |
| 2 | ATGGCGAAC | N#CCCCOc1cccc(C=O)c1 |
| 2 | GCACTCAAC | CCOc1cc(C=O)ccc1OC |
| 2 | TGTGTCCAC | O=CC1CCCCCCC1 |
| 2 | AGCCGATAC | CCCn1ccc(C=O)n1 |
| 2 | CGTAGGAAC | COc1nccc(C=O)n1 |
| 2 | GGATCTGAC | O=CCC1CCOCC1 |
| 2 | TGAGGACAC | O=Cc1cn2c(n1)CCC2 |
| 2 | AGCATGGAC | O=Cc1cc(C(=O)O)ccc1O |
| 2 | CACGTTGAC | O=CC(=O)O |
| 2 | GCATGAGAC | COc1cccnc1C=O |
| 2 | TGTTGCCAC | C#Cc1cccc(C=O)c1 |
| 2 | AGTCGTCAC | O=Cc1ccnc(O)c1 |
| 2 | CGTCAAGAC | O=Cc1ccc2c(c1)CCC2 |
| 2 | GGCTATGAC | O=Cc1cn(-c2ccccc2)nn1 |
| 2 | TGCGGTAAC | O=Cc1cnn[nH]1 |
| 2 | AGCGTGAAC | O=CC[C@@H](O)[C@H](O)[C@H](O)CO |
| 2 | CAGTCCAAC | O=CC1CCC1 |
| 2 | GCTTACCAC | CC(=O)OCc1ccc(C=O)o1 |
| 2 | TTGGCTGAC | CCCC=O |
| 2 | ATCGAGGAC | O=C[C@H](O)CO |
| 2 | CGTCACAAC | O=CCOCc1ccccc1 |
| 2 | GTCAGGAAC | O=CC[C@@H](O)[C@@H](O)CO |
| 2 | TTCGTGGAC | COC(=O)CCC=O |
| 2 | AGGAACCAC | O=CC[C@H](O)[C@H](O)CO |

| Cycle | Tag (5'-3') | Smiles |
|---|---|---|
| 2 | CATCGCTAC | O=Cc1ccccc1-n1cccc1 |
| 2 | GGAGTCAAC | COc1ncccc1C=O |
| 2 | TTGGTGGAC | Cn1cc(C=O)cn1 |
| 2 | CACTTCGAC | Cc1cccc(C=O)n1 |
| 2 | CGTCCTTAC | O=Cc1cc2cc[nH]c2cn1 |
| 2 | GTCGTGAAC | O=Cc1cccc(-c2nn[nH]n2)c1 |
| 2 | AACTGGCAC | O=Cc1ccc(-c2nn[nH]n2)cc1 |
| 2 | AGGTAGCAC | COc1nc(C)ccc1C=O |
| 2 | CGAAGTCAC | O=CC1CC1 |
| 2 | GGTCCTAAC | CC(C)(CC=O)C1CC1 |
| 2 | AAGCACCAC | O=CC1CCCCC1 |
| 2 | CAGTTCCAC | CC(C)(C)c1ccc(CC=O)cc1 |
| 2 | CGTCGTAAC | CC(C)(O)c1ccc(C=O)cc1 |
| 2 | GTGGTAGAC | O=CC1CC2(CC2)C1 |
| 2 | AATCGCCAC | COC(C)(C)CC=O |
| 2 | AGGTGAGAC | O=Cc1ccc2c(n1)NC(=O)CO2 |
| 2 | CGAATGCAC | Cc1cc(C)c(C(=O)O)cc1C=O |
| 2 | GTTGGAGAC | O=Cc1cccc2ccoc12 |
| 2 | AAGGCACAC | O=Cc1csnn1 |
| 2 | CGAGAAGAC | NS(=O)(=O)c1ccc(C=O)cc1 |
| 2 | CTGCGATAC | O=S(=O)(Cl)c1cc(F)ccc1F |
| 2 | GTTGAGCAC | COc1cc(C)c(Cl)cc1S(=O)(=O)Cl |
| 2 | ACAGGCTAC | O=C1COc2cc(S(=O)(=O)Cl)ccc2N1 |
| 2 | AGGTGCTAC | Cc1noc(C)c1S(=O)(=O)Cl |
| 2 | CGAGTGAAC | Cc1ccc(S(=O)(=O)Cl)cc1 |
| 2 | TCAGGTCAC | O=S(=O)(Cl)c1cc(F)cc(F)c1 |
| 2 | ACCTCAGAC | O=S(=O)(Cl)c1cc(Cl)cc(Cl)c1 |
| 2 | CGAGACTAC | O=[N+]([O-])c1ccc(S(=O)(=O)Cl)c(Cl)c1 |
| 2 | CTTGGCAAC | O=S(=O)(Cl)c1ccc(F)c(Cl)c1 |
| 2 | TAGCTGCAC | O=c1ccc2ccc(S(=O)(=O)Cl)cc2[nH]1 |
| 2 | ACCATCCAC | O=c1[nH]c(=O)c2cc(S(=O)(=O)Cl)cnc2[nH]1 |
| 2 | AGTAGGCAC | O=c1[nH]c2ccc(S(=O)(=O)Cl)cc2s1 |
| 2 | CGTTCTGAC | O=C(N1CCc2ccc(S(=O)(=O)Cl)cc2C1)C(F)(F)F |
| 2 | TCGGTGAAC | C#CCOc1cccc(S(=O)(=O)Cl)c1 |
| 2 | ACGGTAGAC | Cc1ccncc1S(=O)(=O)Cl |
| 2 | CGAGGTTAC | C=C(C)COc1cccc(S(=O)(=O)Cl)c1 |
| 2 | GCACTGAAC | N#Cc1ccc(S(=O)(=O)Cl)cc1 |
| 2 | TAGGCACAC | O=S(=O)(Cl)c1cc(F)cc(Cl)c1 |
| 2 | ACGCAAGAC | O=C(O)c1ccc(S(=O)(=O)Cl)c([N+](=O)[O-])c1 |
| 2 | AGTTGCGAC | Cn1nncc1S(=O)(=O)Cl |
| 2 | CGTTGACAC | CCc1ccc(S(=O)(=O)Cl)cc1 |
| 2 | TCTCCGAAC | O=Cc1ccc(S(=O)(=O)Cl)cc1 |
| 2 | AGAAGGCAC | Cn1ccnc1S(=O)(=O)Cl |
| 2 | CGCAGATAC | Nc1ncc(S(=O)(=O)Cl)cn1 |
| 2 | GCTACAGAC | O=S(=O)(Cl)c1nnc[nH]1 |
| 2 | TCATGGCAC | COc1cccc(S(=O)(=O)Cl)c1 |
| 2 | ACGCCTTAC | CC(C)n1cnc(S(=O)(=O)Cl)c1 |
| 2 | ATCCGACAC | Cc1ccccc1S(=O)(=O)Cl |
| 2 | CTTAGCCAC | Cc1c(S(=O)(=O)Cl)cnn1C |

| Cycle | Tag (5'-3') | Smiles |
|---|---|---|
| 2 | TGAGCTGAC | COc1ccccc1S(=O)(=O)Cl |
| 2 | AGACTGCAC | CCn1cc(S(=O)(=O)Cl)cn1 |
| 2 | CGCATCTAC | O=S(=O)(Cl)c1cnc2n1CCC2 |
| 2 | GGAACGAAC | N#Cc1ncccc1S(=O)(=O)Cl |
| 2 | TCCGTAGAC | Cn1cnc(S(=O)(=O)Cl)c1 |
| 2 | ACTTCCGAC | N#Cc1ccc(S(=O)(=O)Cl)cn1 |
| 2 | ATCGCTGAC | CCn1cnc(S(=O)(=O)Cl)c1 |
| 2 | CTTCAGCAC | N#Cc1cccc(S(=O)(=O)Cl)c1 |
| 2 | TGGATCGAC | COc1cncc(S(=O)(=O)Cl)c1 |
| 2 | AGGACCAAC | CCn1cc(S(=O)(=O)Cl)nc1C |
| 2 | AGGACTCAC | NS(=O)(=O)c1ccc(S(=O)(=O)Cl)cc1 |
| 2 | AGGAGCAAC | Cc1nc(S(=O)(=O)Cl)cn1C |
| 2 | AGTCTCGAC | CC(C)n1cc(S(=O)(=O)Cl)cn1 |
| 2 | AGTGGTGAC | Cc1cnc(N)c(S(=O)(=O)Cl)c1 |
| 2 | ATCACGCAC | COc1ccc(S(=O)(=O)Cl)cn1 |
| 2 | ATCGGTGAC | COc1ccc(S(=O)(=O)Cl)cc1 |
| 2 | ATGCGACAC | CCn1ncc(S(=O)(=O)Cl)c1C |
| 3 | CAACGAGTT | COC(=O)c1cc(B2OC(C)(C)C(C)(C)O2)ccc1C |
| 3 | TACTCGGTT | CC1(C)OB(c2ccc(S(C)(=O)=O)cc2)OC1(C)C |
| 3 | TGGAGCATT | Cc1ccc(C(=O)O)cc1B1OC(C)(C)C(C)(C)O1 |
| 3 | GAGTCTGTT | OB(O)c1ccc(Cl)cc1 |
| 3 | CAATCCGTT | COc1cccc(B(O)O)c1OC |
| 3 | GACTCTCTT | CC1(C)OB(c2cccc(S(N)(=O)=O)c2)OC1(C)C |
| 3 | TATCGCCTT | CN(C)CCc1ccc(B(O)O)cc1 |
| 3 | TGGATGCTT | Cn1nnc2ccc(B3OC(C)(C)C(C)(C)O3)cc21 |
| 3 | GTCGTCATT | CC1(C)OB(c2cccc3ccoc23)OC1(C)C |
| 3 | CAGAAGGTT | COC(=O)c1cc(B2OC(C)(C)C(C)(C)O2)ccc1O |
| 3 | GGCATGTTT | CC1(C)Cc2cc(B3OC(C)(C)C(C)(C)O3)ccc2O1 |
| 3 | CGACTCTTT | Cc1cc2cc(B3OC(C)(C)C(C)(C)O3)ccc2[nH]1 |
| 3 | GCAGATGTT | CC1(C)OB(c2ccc(C3(CO)CC3)cc2)OC1(C)C |
| 3 | CAGGAAGTT | Cc1nc2ccc(B3OC(C)(C)C(C)(C)O3)cc2o1 |
| 3 | TCACACGTT | CC1(C)OB(c2cccc3c2CCO3)OC1(C)C |
| 3 | CAACAGCTT | CC(C)OCc1cccc(B(O)O)c1 |
| 3 | CGTCGAATT | COC(=O)c1ccc(B2OC(C)(C)C(C)(C)O2)o1 |
| 3 | GTGCCAATT | OCc1ccc(B(O)O)cc1Cl |
| 3 | CAGTCGTTT | CC1(C)OB(c2cnc3ccnn3c2)OC1(C)C |
| 3 | CACCGTATT | COc1ccc(B2OC(C)(C)C(C)(C)O2)cn1 |
| 3 | CGTCTCATT | CC1(C)OB(c2ccc3ccc(=O)[nH]c3c2)OC1(C)C |
| 3 | GCTAACCTT | CC1(C)OB(c2ccc3c(c2)CCCNC3=O)OC1(C)C |
| 3 | GTGCGTATT | OB(O)c1cccc2c1OCCCO2 |
| 3 | GTGAGGTTT | CC(C)c1nn(C)cc1B1OC(C)(C)C(C)(C)O1 |
| 3 | CAGTCTCTT | CC1(C)OB(c2cccc(Cn3cccn3)c2)OC1(C)C |
| 3 | GTGGCTATT | COc1cc(CO)ccc1B1OC(C)(C)C(C)(C)O1 |
| 3 | GCACAGTTT | CC1(C)OB(c2ccc(C3(N)CCCC3)cc2)OC1(C)C |
| 3 | TCTGGTGTT | OCc1cccc(B(O)O)c1 |
| 3 | CCACAACTT | CC1(C)OB(c2cccc(O)c2)OC1(C)C |
| 3 | GCTGTAGTT | CC1(C)OB(c2ccc(C3CC(=O)C3)cc2)OC1(C)C |
| 3 | GCAGTCATT | CC1(C)OB(c2cccc3c2CNC3=O)OC1(C)C |
| 3 | TGAGGTCTT | COc1cc(B(O)O)ccc1Cl |

| Cycle | Tag (5'-3') | Smiles |
|---|---|---|
| 3 | CCACTCATT | CC1(C)OB(c2cccc(CN)c2)OC1(C)C |
| 3 | CTCCGATTT | CC1(C)OB(c2ccc3c(c2)COC(=O)N3)OC1(C)C |
| 3 | CCTTGCTTT | OB(O)c1ccc(F)nc1 |
| 3 | GCCAGTATT | CC1(C)OB(c2cnc(N)nc2)OC1(C)C |
| 3 | GTGTCGTTT | Cn1c(=O)[nH]c2ccc(B3OC(C)(C)C(C)(C)O3)cc21 |
| 3 | TGATCGGTT | CC1(C)OB(c2ccc([C@@H]3CCC(=O)N3)cc2)OC1(C)C |
| 3 | CTCTCCTTT | Cc1cc(F)ccc1B(O)O |
| 3 | TACCGAGTT | CC(C)(C)n1cc(B2OC(C)(C)C(C)(C)O2)cn1 |
| 3 | CGCTGTATT | Cc1cccc(B(O)O)c1C |
| 3 | GTTCTCCTT | Cc1ccc2cc(B(O)O)ccc2n1 |
| 3 | CCATAGGTT | COc1ncc(B2OC(C)(C)C(C)(C)O2)cn1 |
| 3 | GCTAAGGTT | Cc1ccc(C(N)=O)cc1B1OC(C)(C)C(C)(C)O1 |
| 3 | TGCGACTTT | CN(C)Cc1cc(B(O)O)ccc1F |
| 3 | CTGTGAGTT | CNC(=O)c1ccc(B2OC(C)(C)C(C)(C)O2)cc1C |
| 3 | TCCAACGTT | CC(=O)c1cccc(B(O)O)c1 |
| 3 | GTTGGTCTT | CC1(C)OB(c2cnn(-c3ccncc3)c2)OC1(C)C |
| 3 | TGGACCTTT | CNC(=O)Nc1ccc(B2OC(C)(C)C(C)(C)O2)cc1 |
| 3 | CCGCAATTT | OB(O)c1ccc2c(c1)CCC2 |
| 3 | CTGTGCTTT | CN(C)c1cc(B2OC(C)(C)C(C)(C)O2)ccn1 |
| 3 | TCGGTAGTT | OB(O)c1ccc(F)c(Cl)c1 |
| 3 | CGAGAACTT | OB(O)c1cccc(Cl)c1F |
| 3 | GGAACCTTT | CC1(C)OB(c2cc3ccccc3o2)OC1(C)C |
| 3 | TAGCAGGTT | COCCc1cccc(B2OC(C)(C)C(C)(C)O2)c1 |
| 3 | CAACCACTT | Cc1ccc(Cl)cc1B(O)O |
| 3 | CGACCTATT | COc1ncc(Cl)cc1B(O)O |
| 3 | GAGTTCGTT | NC(=O)c1ccc(B(O)O)cc1 |
| 3 | GTCCACATT | O=C(O)c1cc(B(O)O)ccc1Cl |
| 3 | TCGTTGGTT | CN(C)c1ncc(B2OC(C)(C)C(C)(C)O2)cc1Cl |
| 3 | CGATTGCTT | C[C@@H]1Cc2cc(B(O)O)ccc2O1 |
| 3 | TAGCGCATT | Cc1c(Cl)cccc1B(O)O |
| 3 | GCAACAGTT | OB(O)c1cccc2c1OCC2 |
| 3 | TCTGCGTTT | COc1ccc(B(O)O)cc1C#N |
| 3 | GGTACAGTT | CC1(C)OB(c2ccc3c(c2)CCC(=O)N3)OC1(C)C |
| 3 | TCATCGCTT | CC1(C)OB(c2ccc3c(c2)CCOC3)OC1(C)C |
| 3 | CAGACCATT | CC1(C)OB(c2cccc(O)c2F)OC1(C)C |
| 3 | GCAAGACTT | OB(O)c1ccc2c(c1)CCCO2 |
| 3 | GTGGTGTTT | COc1ccc(B(O)O)cc1F |
| 3 | TGCCAAGTT | COC(=O)c1cc(O)cc(B2OC(C)(C)C(C)(C)O2)c1 |
| 3 | GGTGAAGTT | COc1ncc(B2OC(C)(C)C(C)(C)O2)cc1C |
| 3 | CAGGACATT | CC1(C)OB(C2=CCNCC2)OC1(C)C |
| 3 | TGCTTGCTT | OB(O)c1ccc2c(c1)OCO2 |
| 3 | GGTTAGGTT | Nc1cccc(B(O)O)c1 |
| 3 | TCGTGGTTT | COC(=O)c1ccc(B2OC(C)(C)C(C)(C)O2)cc1F |
| 3 | CTACGACTT | OB(O)c1cncnc1 |
| 3 | GCGGTATTT | CC(C)(O)c1ccc(B2OC(C)(C)C(C)(C)O2)c(F)c1 |
| 3 | GTTGCTGTT | C[C@H](O)c1ccc(B(O)O)cc1 |
| 3 | CAACTCGTT | CC1(C)OB(c2cnc3occc3c2)OC1(C)C |
| 3 | CGTAGAGTT | Nc1cc(B(O)O)ccc1Cl |
| 3 | GTGGAACTT | CCN(CC)c1ncc(B(O)O)cn1 |

| Cycle | Tag (5'-3') | Smiles |
|---|---|---|
| 3 | TCTCGACTT | OB(O)c1ccc2ccc(O)cc2c1 |
| 3 | CAGTCAGTT | COc1cccc2cc(B3OC(C)(C)C(C)(C)O3)oc12 |
| 3 | CTCATCCTT | CCn1ccc(B2OC(C)(C)C(C)(C)O2)n1 |
| 3 | GCGTTCTTT | OB(O)c1cccc2ccccc12 |
| 3 | TACGTCCTT | COC(=O)CCc1cccc(B(O)O)c1 |
| 3 | CAAGGCTTT | COc1ccc(C)cc1B(O)O |
| 3 | GTGGTTCTT | CCc1ccccc1B(O)O |
| 3 | TGAGCAGTT | OCc1ccc(B(O)O)cc1 |
| 3 | CAGTGTGTT | OB(O)c1cccc(Cl)c1 |
| 3 | CTCATGGTT | COc1ccc2cc(B(O)O)ccc2c1 |
| 3 | TAGCTCCTT | CC(C)Oc1ccc(B(O)O)cc1F |
| 3 | CTCGCAATT | COC(=O)c1cccc(B(O)O)c1 |
| 3 | GTTGCACTT | OB(O)c1ccc(Cl)c(F)c1 |
| 3 | TGCCTTCTT | CCCc1ccc(B(O)O)cc1 |
| 3 | CATCCTGTT | O=C(O)CCc1ccc(B(O)O)cc1 |
| 3 | CTGAGCATT | COc1ccc(B(O)O)cc1C(=O)O |
| 3 | TCACCAGTT | CC(C)(C)c1ccc(B(O)O)cc1 |
| 3 | CCACACTTT | OB(O)c1ccc(Oc2ccccc2)cc1 |
| 3 | CTGCCTATT | COc1ccc(F)c(B(O)O)c1 |
| 3 | TACAGCGTT | CCOc1ccc(B(O)O)cc1C |
| 3 | TGGAGGTTT | OB(O)c1cccc(OC(F)(F)F)c1 |
| 3 | GAATGCGTT | CC(C)Cc1ccc(B(O)O)cc1 |
| 3 | GGCATCATT | OB(O)c1cc(F)c(F)c(F)c1 |
| 3 | CCTAGTGTT | Cc1cc(C)cc(B(O)O)c1 |
| 3 | CCATACCTT | OB(O)c1ccc2ccccc2c1 |
| 3 | GACACAGTT | CC(C)(C)c1cccc(B(O)O)c1 |
| 3 | GTAGAGGTT | O=C(O)c1ccc(B(O)O)c(F)c1 |
| 3 | CCTTCTGTT | O=C(O)c1ccc(B(O)O)c(Cl)c1 |
| 3 | TGTCCAGTT | CCOc1ccc(B(O)O)cc1 |
| 3 | GACCTGTTT | CCCOc1cccc(B(O)O)c1 |
| 3 | GTAGGCTTT | CCOc1cc(F)ccc1B(O)O |
| 3 | TCCACTGTT | Cc1ccc(B(O)O)c(Cl)c1 |
| 3 | CGACGATTT | Cn1nccc1B(O)OC(C)(C)C(C)(C)O |
| 3 | GCGTAGTTT | OB(O)c1cccc(-c2ccccc2)c1 |
| 3 | CCGTCAATT | COc1ccc(C(C)C)cc1B(O)O |
| 3 | GACTCGTTT | OCc1ccccc1B(O)O |
| 3 | GTATCGCTT | OB(O)c1ccc2cc[nH]c2c1 |
| 3 | TCTGCTCTT | CCc1ccc(B(O)O)cc1 |
| 3 | TGCTCCTTT | OB(O)/C=C/C1CCCCC1 |
| 3 | GAACTCCTT | N#Cc1cc(B(O)O)ccc1Cl |
| 3 | GCTCTCATT | Cc1ccc(OC(C)C)c(B(O)O)c1 |
| 3 | TCCGATGTT | COc1c(C)cc(B(O)O)cc1C |
| 3 | CGCTATCTT | OB(O)c1cccc(F)c1 |
| 3 | GAGCTTCTT | CCc1cccc(B(O)O)c1 |
| 3 | GTTCCAGTT | O=C(NC1CC1)c1cccc(B(O)O)c1 |
| 3 | CCGTGATTT | CCOc1ncccc1B(O)O |
| 3 | GACGATGTT | O=C(O)c1cc(Cl)cc(B(O)O)c1 |
| 3 | GGAAGTCTT | CC(C)c1cccc(B(O)OCCO)c1 |
| 3 | TCGCTTCTT | CCOC(=O)c1cccc(B(O)O)c1 |

| Cycle | Tag (5'-3') | Smiles |
|---|---|---|
| 3 | CGCTGATTT | Cc1cccc(B(O)O)c1 |
| 3 | GAGTCCATT | COc1ccc(Cl)cc1B(O)O |
| 3 | TAACCGCTT | COc1cc(F)c(F)cc1B(O)O |
| 3 | TGGTGAGTT | COC(=O)Nc1ccc(B(O)O)cc1 |
| 3 | CCTGTCTTT | OB(O)c1cc(F)cc(Cl)c1 |
| 3 | GAGGTAGTT | CCOc1ccccc1B(O)O |
| 3 | GGAGTAGTT | O=C(O)c1cccc(B(O)O)c1 |
| 3 | TCGGACATT | CS(=O)(=O)c1ccc(B(O)O)cc1 |
| 3 | CGGAGAATT | OB(O)c1ccc(C(F)(F)F)cc1 |
| 3 | GATAGCGTT | Nc1cc(B(O)O)ccc1F |
| 3 | TATGCCGTT | COC(=O)c1cc(B(O)O)ccc1Cl |
| 3 | CACCAAGTT | O=C(O)Cc1cccc(B(O)O)c1 |
| 3 | CGAGGTATT | OB(O)c1ccc(F)c(C(F)(F)F)c1 |
| 3 | GGTGTTGTT | Cc1cc(Cl)ccc1B(O)O |
| 3 | TGCGTCATT | OB(O)c1ccc(Cl)c(Cl)c1 |
| 3 | CGGATACTT | OB(O)c1cc(Cl)cc(Cl)c1 |
| 3 | GCAGAACTT | O=C(O)c1cccc(B(O)O)c1F |
| 3 | TCACAGCTT | COc1cc(B(O)O)cc(C(=O)O)c1 |
| 3 | CACGACTTT | OB(O)c1cccc(Oc2ccccc2)c1 |
| 3 | CGCTCAATT | CC(C)(C)[C@H]1CC=C(B(O)O)CC1 |
| 3 | GATGTCCTT | OB(O)c1cccc2c1OCCO2 |
| 3 | GTCATGCTT | Cc1cc(B(O)O)ccc1OC(C)C |
| 3 | TGCTAGGTT | OCCc1cccc(B(O)O)c1 |
| 3 | CGGTAGTTT | N#Cc1ccc(B(O)O)cc1F |
| 3 | GCATGGTTT | OB(O)c1ccc(C(F)(F)F)c(F)c1 |
| 3 | TCAGAGGTT | COC(=O)c1ccc(B(O)O)c(F)c1 |
| 3 | CGGCTAATT | OB(O)c1ccc2c(c1)OC(F)(F)O2 |
| 3 | GATTGCCTT | Cc1ccc(F)cc1B(O)O |
| 3 | CTCTTGCTT | OB(O)c1cc(F)cc(C(F)(F)F)c1 |
| 3 | GGATGAGTT | Cc1cc(B(O)O)ccc1Cl |
| 3 | TCCTGTGTT | COC(=O)c1cc(Cl)cc(B(O)O)c1 |
| 3 | CGTACTCTT | COc1ncc(B(O)O)cc1Cl |
| 3 | GTGTGTCTT | OB(O)c1cc(F)cc(F)c1 |
| 3 | CAGGTTGTT | Cc1cc(B(O)O)c(C)cc1F |
| 3 | CTTCCTCTT | Cc1ccc(B(O)O)cc1 |
| 3 | GGTAGACTT | COc1ccc(B(O)O)cc1C |
| 3 | TCGTTCCTT | COc1cc(B(O)O)cc(OC)c1OC |
| 3 | CATAGGCTT | Cc1ccc(B(O)O)c(F)c1 |
| 3 | CGTGACTTT | CCOc1cccc(B(O)O)c1 |
| 3 | GCAATCGTT | COc1cc(F)cc(B(O)O)c1 |
| 3 | CATCGGTTT | CCN(CC)Cc1ccc(B(O)O)cc1 |
| 3 | CTTCTCGTT | CC(=O)Nc1cccc(B(O)O)c1 |
| 3 | GGTGACATT | O=C(O)COc1cc(F)cc(B(O)O)c1 |
| 3 | TGACGCATT | CCNC(=O)c1cccc(B(O)O)c1 |
| 3 | CATCGACTT | COc1ccc(C#N)cc1B(O)O |
| 3 | CTAAGCGTT | CNC(=O)c1ccc(B(O)O)cc1 |
| 3 | GCAGGATTT | COc1cc(OC)cc(B(O)O)c1 |
| 3 | TACCACCTT | CC(C)(C)OCc1ccc(B(O)O)cc1 |
| 3 | CCAACACTT | CCCCc1ccc(B(O)O)cc1 |

| Cycle | Tag (5'-3') | Smiles |
|---|---|---|
| 3 | GTAACGGTT | CCNC(=O)c1ccc(B(O)O)cc1 |
| 3 | TGAGTGCTT | O=C(O)c1ccc(B(O)O)cc1Cl |
| 3 | CCACCATTT | COc1cc(C(=O)O)ccc1B(O)O |
| 3 | GCCTAACTT | O=C(O)c1ccc(B(O)O)cc1F |
| 3 | TACGAGGTT | CC(C)Oc1ccc(F)cc1B(O)O |
| 3 | CCAACCTTT | Cc1cc(B(O)O)ccc1F |
| 3 | GAACGCTTT | COc1ccc(B(O)O)cc1Cl |
| 3 | GTCTACCTT | CC(C)NC(=O)c1cccc(B(O)O)c1 |
| 3 | TGCCAGTTT | CCCOc1ccc(B(O)O)cc1Cl |
| 3 | CCGAACATT | COC(=O)c1cccc(B(O)O)c1F |
| 3 | CTCACTGTT | OB(O)c1ccc(-c2ccccc2)cc1 |
| 3 | GCTCCTATT | OB(O)c1ccccc1 |
| 3 | TAGGACCTT | CC(=O)c1ccc(B(O)O)cc1 |
| 3 | CCACGTATT | COc1ncccc1B(O)O |
| 3 | GACACCATT | OCc1ccc(Cl)c(B(O)O)c1 |
| 3 | GTCTCAGTT | O=C(O)CCc1cccc(B(O)O)c1 |
| 3 | TGCTCTCTT | CONC(=O)c1ccc(B(O)O)cc1 |
| 3 | CCGAATGTT | CCCNC(=O)c1cccc(B(O)O)c1 |
| 3 | GCTCTACTT | COC(=O)c1cc(F)cc(B(O)O)c1 |
| 3 | CCATCAGTT | CCOC(=O)c1ccc(B(O)O)cc1F |
| 3 | GAGCAGTTT | Cc1ccc(B(O)O)cc1Cl |
| 3 | GTGTTGCTT | COc1cc(B(O)O)ccc1C |
| 3 | TGTAGCCTT | OB(O)c1ccc(OC(F)(F)F)cc1 |
| 3 | TGTGCTGTT | COC(=O)Cc1ccc(B(O)O)cc1 |
| 3 | TGTGGAGTT | COc1ccc(B(O)O)cc1OC(C)C |
| 3 | TGTGGCTTT | NS(=O)(=O)c1ccc(B(O)O)cc1 |
| 3 | TGTGTCGTT | Null-reaction; heated with Pd 'catalyst' |
| 3 | CACGGAATT | Null-reaction; not heated without Pd 'catalyst' |
